# A lipid compendium of a metabolically compromised bacterium provides insights into lipid acquisition, biosynthesis, and metabolism

**DOI:** 10.64898/2026.05.22.727245

**Authors:** Poulami Chatterjee, Hyejin Esther Shin, Miles I. Tuncel, Isaac A. Paddy, Alysha K. Lee, Joshua McCausland, Paula V. Welander, Christine Jacobs-Wagner, Laura M. K. Dassama

## Abstract

The Lyme disease agent *Borrelia burgdorferi* belongs to a class of metabolically compromised bacteria that cannot survive without host-derived lipids. Survival of the agent in tick and vertebrate hosts requires substantial nutrient acquisition and potential cell envelope remodeling. While prior studies identified cholesterol, cholesterol glycolipids, and phosphatidylcholines as membrane lipids in *B. burgdorferi*, the identity of many other membrane lipids, their origin, and their physiological relevance remain unknown. Here, we used a suite of untargeted and targeted high-resolution mass spectrometry methods to reveal a complex lipid profile of the pathogen and to identify the origin of its lipids. The analysis detected more than 500 lipids in *B. burgdorferi*, the majority of which are sourced from the environment. However, the bacterium selectively accumulates certain lipids while excluding others, suggesting discriminatory uptake. These include cholesteryl esters and triglycerides that are organized in foci within the pathogen. Intriguingly, the pathogen also synthesizes predominantly eukaryotic lipids such as the lysosomal bis(monoacylglycerol)phosphate and the plant glycolipid sulfoquinovosyl diacylglycerol (SQDG). The biosynthesis of the latter is carried out by enzymes that exhibit structural homology to plant oxidoreductases and galactosyltransferases, yet their closest orthologs are found in bacteria. This hints that the capability of SQDG synthesis is more widespread in spirochaetes and other bacteria. Together, the comprehensive lipid profiling we report here uncovers novel aspects of the physiology of the metabolically challenged *B. burgdorferi* and highlights lipid acquisition and synthesis pathways as potentially critical for pathogen survival.

## Introduction

Tick borne infections are on the rise, with Lyme disease being the most prevalent tick-borne illness in North America and Europe.^1^ The incidence of the disease has surged by 250-300% over roughly two decades in the north-central and northeastern regions of the United States alone.^2^ The economic burden on the U.S. healthcare system is estimated to be between $712 million and $1.3 billion annually, with the number of Lyme disease cases ranging from 240,000 to 440,000.^3^ When detected early, Lyme disease can be effectively treated with antibiotics, leading to the resolution of symptoms. However, delays in treatment can allow the pathogen to disseminate throughout the body, resulting in complications such as meningitis, arthritis, and carditis; Lyme arthritis sometimes persists for many years.^4^ Significant efforts have been made to develop protein and mRNA-based vaccines targeting the outer surface proteins (Osp) A and B of the causative agent, *Borrelia burgdorferi*.^5^ While higher OspA-specific immunoglobulin G titers compared to previous protein-based vaccine formulations have been observed, definitive therapeutic efficacy data in human patients are currently unavailable.

With a genome size of approximately ∼1.5 Mbp, *B. burgdorferi’s* survival is closely tied to its metabolic opportunities.^6^ This metabolic minimalism is also shared by other pathogenic spirochetes, including the syphilis agent *Treponema pallidum*. *B. burgdorferi* relies on its tick and vertebrate hosts for access to multiple essential metabolites.^7^ The bacterium can metabolize carbohydrates through glycolysis and can utilize alternative carbon sources such as mannose, *N*-acetylglucosamine, maltose, chitobiose, and glycerol, enhancing its adaptability in various environments.^8^ However, it lacks pathways for the de novo synthesis of amino acids, making its growth rate entirely dependent on and limited by the rate of peptide and amino acid acquisition from the environment.^9, 10^ Unusually, the outer membrane of *B. burgdorferi* lacks lipopolysaccharides (LPS), components that are typically crucial for maintaining cell integrity and are highly immunogenic. Instead, the pathogen encodes a large number of surface exposed lipoproteins.^11^ Surprisingly, *B. burgdorferi* requires cholesterol, a lipid that is sparsely used in the bacterial domain, for growth and viability.^12^ Cholesterol comprises rafts within the membrane of the pathogen and is modified to glycoforms. Beyond cholesterol glycolipids, *B. burgdorferi* performs limited biosynthesis of additional lipids using exogenous precursors (e.g., phosphatidyl choline using exogenous choline and fatty acids),^13^ and the predominantly plant-based glycolipid, monogalactosyl diacyl glycerol (MGDG).^14^ Other complex lipids and free fatty acids are thought to be acquired from the environment.^15^

While these findings highlight the critical role of specific lipids in *B. burgdorferi* biology, the broader landscape of its lipidome has remained largely unmapped. A recent study^16^ provided a metabolic rationale for this complexity, demonstrating via untargeted mass spectrometry that *B. burgdorferi* preferentially scavenges environmental lipids to avoid the high energetic cost of synthesizing glycerol phosphate precursors. Crucially, the authors established that lipid acquisition directly regulates the pathogen’s virulence profile, specifically the reciprocal expression of OspA and OspC. Despite these insights into the energetic “logic” of scavenging, the full identity of the lipid species and their origins, whether selectively acquired or synthesized via specialized internal pathways, were not fully elucidated. Given that lipids define the interface between the pathogen and the host, a comprehensive compendium of the *B. burgdorferi* lipidome is essential for unraveling precise metabolic strategies, novel biosynthetic capabilities, and functional insights.

Our use of untargeted and the more precise targeted tandem mass spectrometry together identified more than 500 lipid species in *B. burgdorferi.* A significant fraction is identical to those present in the growth medium and are composed of prevalent eukaryotic lipids such as sphingolipids and sterols. Several lipids are significantly depleted (600-1500-fold change) during the pathogen’s growth. These include phospholipids but also neutral lipids, which are known to organize in lipid droplets in other organisms and serve a variety of functions ranging from lipid storage to transcriptional regulation.^17^ Use of microscopy and a fluorescent probe revealed foci of neutral lipids in *B. burgdorferi.* Surprisingly, a fraction of the lipids detected in *B. burgdorferi* were not found in the medium, indicating that they are likely products of biosynthesis or modification using precursors. One is the anionic sulfoquinovosyl diacyl glycerol (SQDG), a lipid that is primarily found in photosynthetic organisms. Using structural homology methods with the plant biosynthetic enzymes as baits, we discovered two enzymes, BB0444 and BB0454, in *B. burgdorferi* that are sufficient for the synthesis of SQDG. Interestingly, orthologs of these enzymes are widespread in the bacterial and archaeal domains, hinting at a preponderance of SQDG synthesis that was not previously known. Together, this resource provides a complete overview of the lipids present within cultures of *B. burgdorferi* and maps their origins. It provides novel insights into previously unidentified biosynthetic pathways and metabolism in an obligate bacterial parasite, which may enable the discovery of protein targets for the selective disruption of lipid-related processes in *B. burgdorferi* and related opportunistic pathogens.

## Results and discussion

### Samples and analytical methods

A comprehensive investigation of the *B. burgdorferi* lipids and their origins required lipid extraction from cells, the BSK-II culture medium supplemented with 6% rabbit serum (complete medium), rabbit serum alone and the spent medium collected after bacterial cultivation. These samples were first evaluated qualitatively via thin-layer chromatography **(Fig. S1)** and subsequently characterized through mass spectrometry methods (**Scheme 1**; see Methods for more details). Given that the chemical structures of lipids impact their ionization, we employed multiple tandem mass spectrometry methods. Free fatty acids were detected via gas chromatography with a 5977 A Series mass selective detector (MSD) with electron ionization (EI). To broadly capture as many species as possible in an untargeted fashion, we first used a Thermo Orbitrap ID-X Tribrid mass spectrometer paired with a Vanquish UHPLC operating in both negative and positive ion modes of electrospray ionization (ESI, **Figs. S2 and S3)**. This setup enabled the annotation of more than 500 lipids using lipid databases. Following this untargeted profiling, we used multiple reaction monitoring (MRM) to validate the presence of each lipid species identified in *B. burgdorferi*; this targeted mass spectrometry method was performed using the highly sensitive Agilent mini Ultivo triple quadrupole (QQQ) mass spectrometer. To detect sterols that are not sufficiently ionized with ESI, we used Atmospheric Pressure Chemical Ionization (APCI) on an Agilent 6530 Q-TOF with an Infinity II UHPLC. Together, these analytical methods provided both a broad overview and a highly sensitive means to confidently assign lipids in the different samples used in the study.

**Scheme 1.**
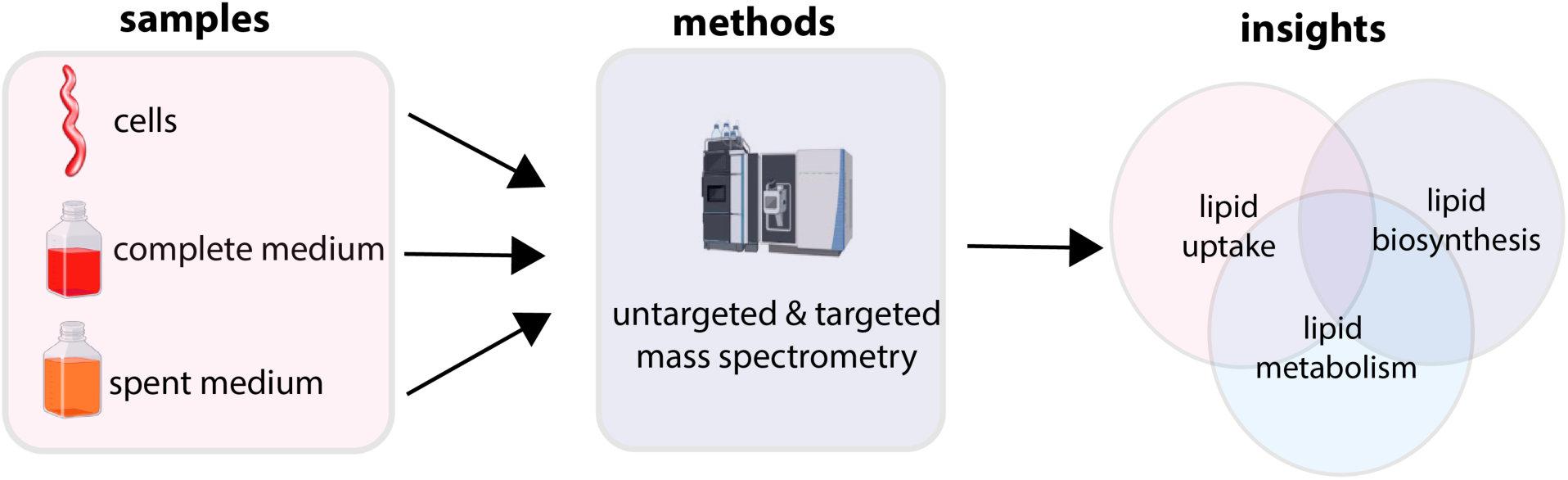
Schematic overview describing the project. Lipids extracted from cultured *B. burgdorferi*, BSK-II supplemented with 6% rabbit serum (complete medium), and the spent medium were subjected to untargeted and targeted tandem mass spectrometry. The resulting analysis provided insights into the identity of lipids taken up from the medium and synthesized by *B. burgdorferi* using precursors present in the medium. It also revealed previously unrecognized forms of lipid metabolism in the pathogen.

### The free fatty acid analysis of *B. burgdorferi* is consistent with previous findings

We employed gas chromatography (GC) to assess the free fatty acid (FA) composition of our samples (**Fig. S4**). Although our data shows that intact complex lipids are taken from the medium, the limited biosynthesis that *B. burgdorferi* performs requires FA. A previous report,^18^ showed the enrichment of myristic (C14:0), palmitic (C16:0), palmitoleic (C16:1), stearic (C18:0), oleic (C18:1), and linoleic (C18:2) acids in the pathogen. Our analysis revealed a similar profile, with stearic (C18:0), oleic (C18:1), palmitic (C16:0), and linoleic (C18:2) acids being predominant free FAs.^16^ Also detected in small amounts were heptadecanoic (C17:0), arachidic (C20:0), myristic (C14:0), and pentadecanoic (C15:0) acids. Collectively, six saturated and two unsaturated fatty acids (FAs) were identified. While FAs were abundant in the rabbit serum and complete medium, myristic acid (C14:0) was a notable exception. Its low level in the complete medium and absence in spent media (**Fig. S4C**) suggest that it is utilized by *B. burgdorferi* during growth.

### Untargeted profiling of the growth medium and its components reveals a diversity of lipids

Untargeted profiling of lipids extracted from rabbit serum, the complete medium (CM), and the spent medium (SPM) revealed a multitude of lipids (735 species) that ranged from free fatty acids to complex phospholipids (**Fig. S5**). Several members of the major classes of lipids, including glycerophospholipids, glycerolipids, sphingolipids, esters of different lipids, and free fatty acids were all enriched in the rabbit serum (**Figs. S5A, B**), the CM (**Figs. S5C, D**), and the SPM (**Figs. S5E, F**) when compared to a lipid extraction control. Nearly all lipids enriched in CM were also detected in the rabbit serum, indicating that the rabbit serum is the primary source of exogenous lipids in cultured *B. burgdorferi* (**Fig. S6A, B**).

Comparative analyses revealed that the lipid composition of the SPM closely mirrored that of the CM for lipids ionized in negative mode (**Fig. S7A**). In contrast, a significant difference was observed for many positively ionized lipids during *B. burgdorferi* growth (**Fig. S7B**). Differential abundance analysis using stringent criteria (adjusted *p* value 0.05, log2 fold change [CM/SPM] ≥ 2) identified marked depletion across multiple lipid classes. Neutral lipids, including cholesteryl esters (ChE), triacylglycerols (TG), and diacylglycerols (DG), exhibited large fold changes: ChE levels decreased by as much as 1160-fold, TG by up to 613-fold, and DG by up to 100-fold in the SPM compared to levels in the CM. Similarly, phosphatidylcholines (PC), lysophosphatidylcholines (LPC), and sphingomyelins (SM) were substantially reduced (4 to 1500-fold) during cultivation (**Fig. S7B**, **Supplementary file 1**). The significant reduction of these lipids in the spent medium provides strong evidence that they were incorporated by *B. burgdorferi* and are likely critical for pathogen growth.

### *B. burgdorferi* has a lipid composition that closely mirrors that of the growth medium

To determine what fraction of the lipids detected in the CM and depleted in the SPM were incorporated into *B. burgdorferi*, we performed a similar untargeted profiling of lipids extracted from the pathogen. Of the 594 lipids annotated in samples of *B. burgdorferi*, 543 were also present in the CM, suggesting that ∼ 95% of the lipids in the pathogen originate in CM. Four primary classes of lipids were shown to be enriched in *B. burgdorferi* when compared to the lipid extraction control (**Figs. 1A-D**). These include glycerophospholipids, for which PC, PE, and phosphatidyl glycerol (PG) are examples previously shown to exist in *B. burgdorferi*.^19^ Also present were glycerolipids such as TGs and DGs, the sphingolipids- SM, ceramides (Cer), and sphingosine (Sph), free fatty acids with varying degrees of saturation, and many others (**Figs. 1C, D**).

**Fig. 1.**
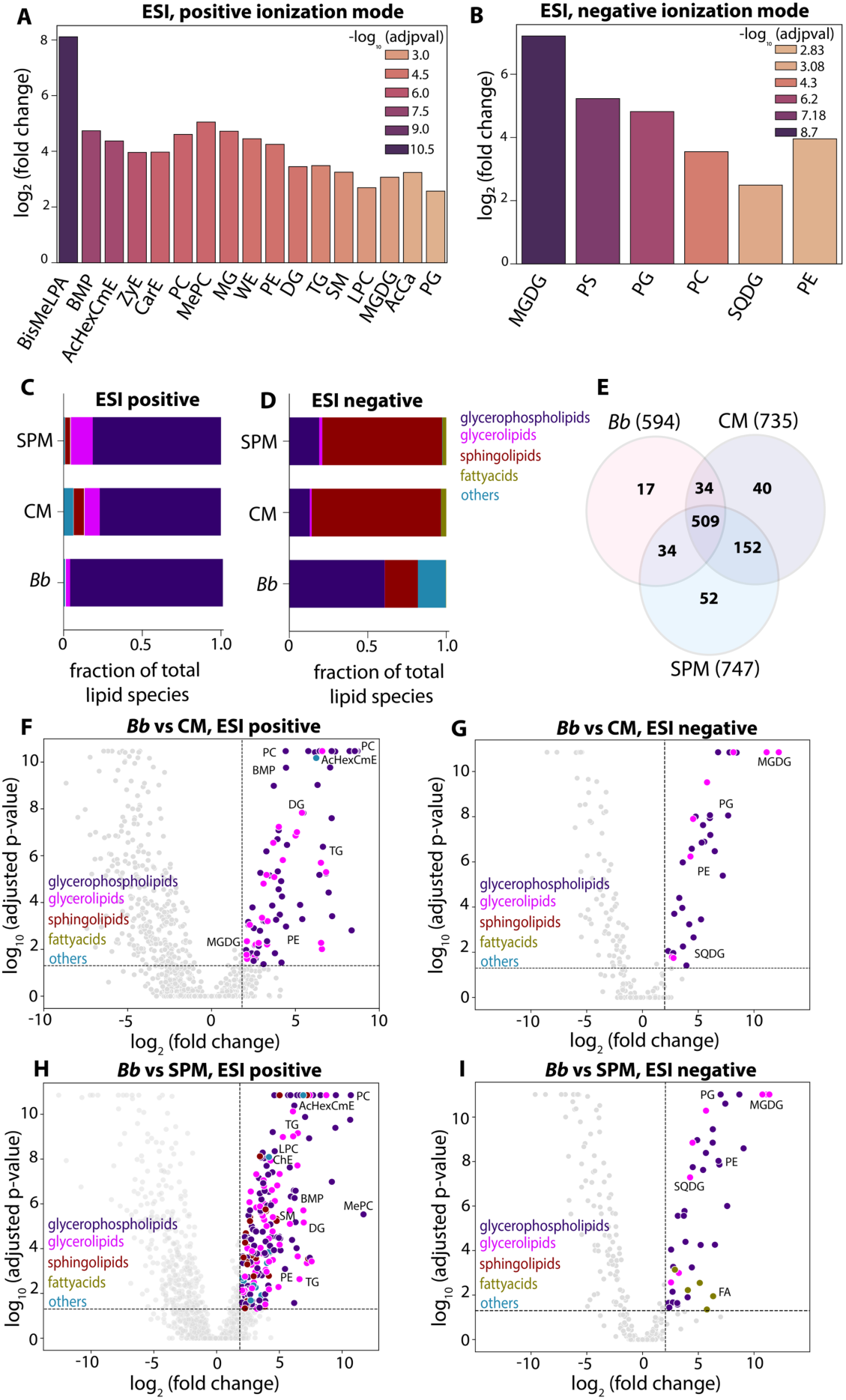
Untargeted lipidomic profiling of *B. burgdorferi* and BSK-II culture medium. Lipid classes identified in the exponential growth phase of *B. burgdorferi* were compared to a lipid extraction control, analyzed via positive **(A)** and negative **(B)** ionization modes; full lipid names are provided in **Table 1**. **(C, D)** Fractional distribution of lipid species across *B. burgdorferi* (*Bb*), complete medium (CM) and spent medium (SPM). **(E)** Venn diagram depicting the number of lipid species detected in *Bb*, CM, and SPM, with overlapping regions indicating shared species. Volcano plots identifying lipids enriched in *Bb* relative to CM in positive (**F**) and negative (**G**) ionization modes); *Bb* relative to the SPM is provided for positive (**H**) and negative (**I**) modes. Data were normalized using the median ratios normalization method and depict three biological replicates (each with three technical replicates) per sample.

**Table 1:** Lipid abbreviations, full names, and higher-order lipid class assignments.

|  |  |  |
| --- | --- | --- |
| lipid | full form | lipid class |
| AcHexChE | acylhexosylcampesterol ester | others |
| BisMeLPA | bis-methyl phosphatidic acid | phosphoglycerolipid |
| BMP | bis(monoacylglycero)phosphate | phosphoglycerolipid |
| CarE | carnitine ester | others |
| Cer | ceramide | sphingolipid |
| ChE | cholesterol ester | others |
| DG | diglyceride | glycerolipid |
| FA | fatty acid | fattyacid |
| Hex1Cer | hexosylceramide | sphingolipid |
| LdMePE | lysodimethylphosphatidylethanolamine | phosphoglycerolipid |
| LPC | lysophosphatidylcholine | phosphoglycerolipid |
| MePC | methylated Phosphatidylcholine | phosphoglycerolipid |
| MG | monoglyceride | glycerolipid |
| MGDG | monogalactosyldiacylglycerol | glycerolipid |
| PC | phosphatidylcholine | phosphoglycerolipid |
| PE | phosphatidylethanolamine | phosphoglycerolipid |
| PG | phosphatidylglycerol | phosphoglycerolipid |
| PS | phosphatidylserine | phosphoglycerolipid |
| SM | sphingomyelin | sphingolipid |
| SPH | sphingosine | sphingolipid |
| SQDG | sulfoquinovosyldiacylglycerol | glycerolipid |
| TG | triglyceride | glycerolipid |
| WE | wax ester | others |
| ZyE | zymosterol ester | others |

Of the lipids detected in *B. burgdorferi* via ESI positive ionization, glycerophospholipids dominate, followed by glycerolipids and sphingolipids. As expected, this distribution closely resembled that of the CM and the SPM (**Fig. 1C**). However, lipids that ionize in negative mode of ESI did not follow the same pattern (**Fig 1D**). Whereas glycerophospholipids are still abundant, the distribution of the classes of lipids detected in *B. burgdorferi* is distinct from what is observed in the medium. This indicates that while the pathogen is mostly reliant on the medium for lipids, it does discriminate against the uptake of some lipids. A Venn diagram of the lipid distributions (**Fig. 1E**) more clearly illustrates these findings: while there is overlap in some species, the lipid composition of *B. burgdorferi* does not perfectly mirror that of the CM, with the pathogen containing 17 lipid species found in neither CM nor SPM.

As before, comparative analyses more clearly highlighted lipids enriched in *B. burgdorferi* when compared with the CM. These included some lipids that were depleted from the CM (TG, ChE and DG) and lipids previously reported^14^ to be synthesized by the pathogen (e.g., the galactolipid, monogalactosyl diacylglycerol-MGDG). Other enriched lipids were unexpected, including the predominantly eukaryotic bis(monoacylglycerol)phosphate (LBPA/BMP), sulfoquinovosyl diacylglycerol (SQDG), and an acylatedhexosylcampesterol ester (AcHexCmE) (**Figs. 1F-H**, **Supplementary files 2 and 3**). These findings were surprising because they have not been previously reported in *B. burgdorferi* and SQDG has only been found in plants and photosynthetic bacteria^20–23^. While many of the lipids enriched in the pathogen when compared to the CM are also enriched when *B. burgdorferi* is compared to the SPM (**Figs. 1H, I**), the greater number of lipids in **Fig. 1H** (*vs.* **Fig. 1F**) lends additional credence to our observation that some lipids are depleted during growth (**Supplementary file 1**).

### *B. burgdorferi* accumulates neutral lipids from the growth medium

Given the reliance of the untargeted approach on accurate annotation using databases, we also used multiple reaction monitoring (MRM) to detect transitions from precursor to product ions for all lipids identified in the pathogen. This method enhances the accuracy and reliability of lipid species identification.^24^ MRM results revealed that, as indicated by the untargeted approach, TGs, ChEs and DG are indeed present in *B. burgdorferi*. Our targeted lipidomics profiling confirmed the presence of 15 species of ChEs (**Figs. 2A and S8**), 8 TGs (**Figs. 2B and S9A-H**) and one DG (**Fig. S9I**), all with diverse fatty acyl tails; these exact lipid species were all also present in the CM, hinting that they are acquired via uptake **(Figs. 2A, B, Figs. S8, S9, Supplementary file 1)**. The presence of neutral lipids in *B. burgdorferi* is unusual, as most bacteria do not store these lipids.^25^ In eukaryotes, neutral lipid domains function in lipid storage by primarily organizing into lipid droplets.^26^ The pathogen *Mycobacterium tuberculosis* has been observed to utilize host-derived triacylglycerols to synthesize polar lipids during dormancy.^27^ Beyond storage, lipid droplets have been implicated in maintaining cellular communication, stress adaptation, redox homeostasis and gene regulation.^28–30^ Given that *B. burgdorferi* resides in the midgut of tick between blood meals (which can be months or a year apart), it is plausible that the pathogen employs a similar mechanism for survival during periods of nutrient limitation.^31^

**Fig. 2.**
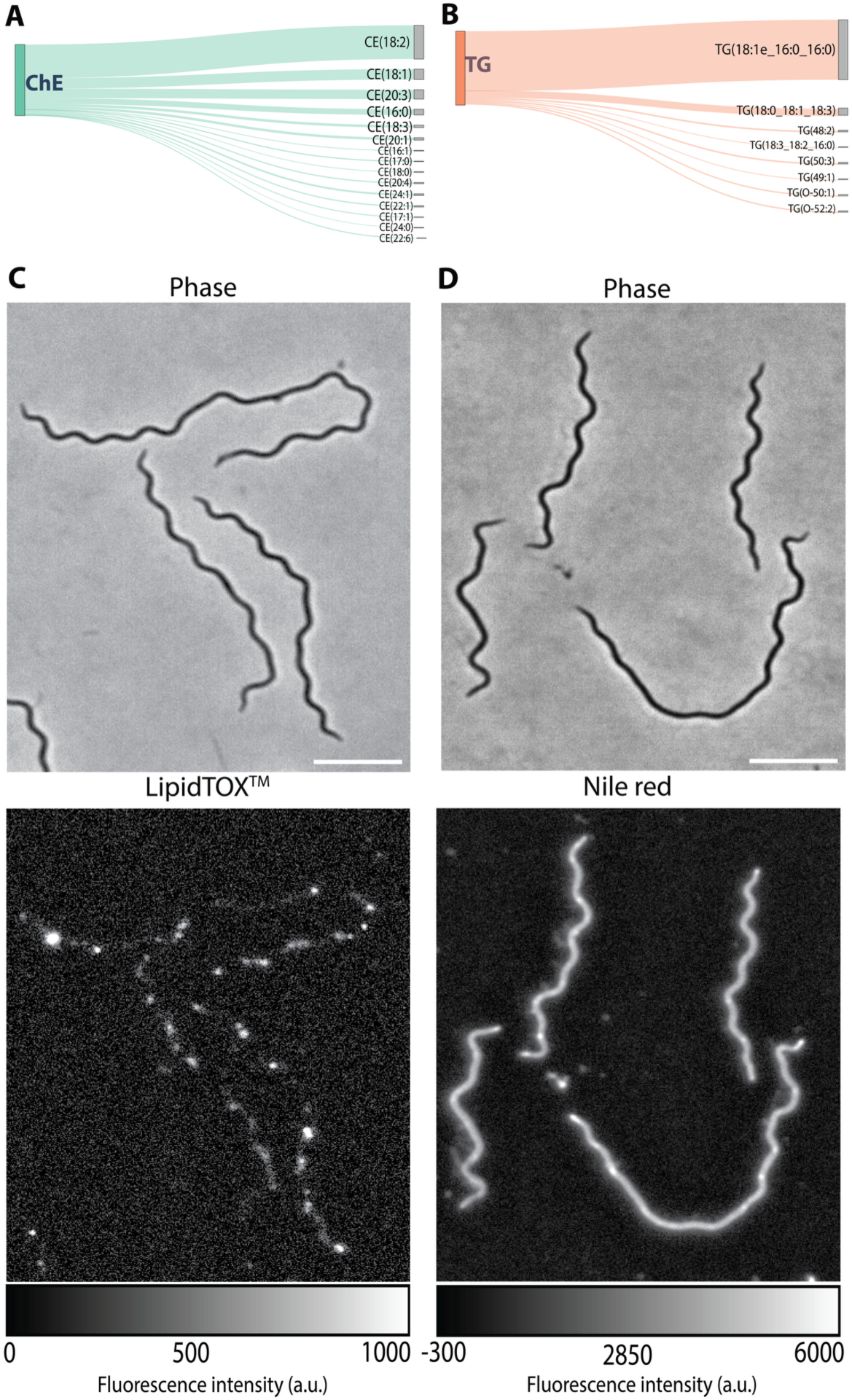
Neutral lipids in *B. burgdorferi.* **(A, B)** Sankey diagrams illustrating the relative abundance of neutral lipids identified via tandem mass spectrometry. The width of each flow is proportional to the integrated peak area under the curve of the daughter ion, with the distribution of cholesteryl esters (ChE, **A**) and triglycerides (TG, **B**). **(C, D)** Widefield epifluorescence microscopy images of fixed *B. burgdorferi* cells show lipid staining with either the neutral lipid-specific LipidTOX^TM^ (**C**) or the non-specific dye Nile Red (**D**). Scale bar = 5 µm.

To investigate whether the ChEs and TGs detected in *B. burgdorferi* might organize into defined domains, we fixed and stained exponentially growing *B. burgdorferi* cells with the neutral lipid-specific LipidTOX;^32^ the non-specific lipid dye Nile Red^33^ was used as a control. We then used wide-field epifluorescence microscopy to image *B. burgdorferi* cells. These data revealed distinct lipid foci in *B. burgdorferi* stained with LipidTOX (**Fig. 2C**) when compared to the DMSO control (**Fig. S10A**). By contrast, cells stained with Nile Red showed more homogeneously distributed fluorescence, as would be expected for a molecule that engages with most membrane lipids (**Figs. 2D and S10B**). While it is difficult to assign a particular location (membrane *vs.* cytoplasm) to the neutral lipid domains in *B. burgdorferi*, it is not unreasonable to assume that they at least function in lipid storage. It has been reported that *B. burgdorferi* can metabolize TGs, and that BB0646 is a lipase capable of catalyzing the release of fatty acids from TGs.^34^ A more recent study revealed BB0562 as another lipase whose knockout resulted in growth defects that were reversed by FA supplementation but not cholesterol.^35^ These findings align with the functional role of FAs (whether taken from the medium or liberated from DGs and TGs) as precursors of more complex lipids synthesized by the pathogen.

### Phosphatidylcholines are abundant in *B. burgdorferi*

The untargeted and targeted mass spectrometry approaches indicated that *B. burgdorferi* contains a significant amount of phospholipids. We identified 97 unique phosphatidylcholine (PC) species in the pathogen. The vast majority of these (94 species) were also present in the complete medium (CM), with only PC (16:1e/22:5), PC (26:1), and PC (17:0/18:2) found exclusively in the pathogen (**Figs. 3A, S11, and S12**). Phosphatidylcholine (PC) is predominantly a eukaryotic membrane lipid,^36^ occurring in a mere 15% of all bacteria. Consequently, the prominent presence of PC in pathogens like *Borrelia burgdorferi*^13^ and *Legionella pneumophila*,^37^ where it serves as a primary membrane phospholipid highlights a highly unusual and specialized evolutionary trait among these bacteria. Consistent with previous reports, our data confirm that cholesterol and its glycolipids constitute a significant portion (∼36%) of the *B. burgdorferi* membrane. However, our findings also reveal that phosphatidylcholines (PCs) represent another major, previously underappreciated lipid class in *B. burgdorferi.*^38^ Although, the pathogen possesses the biosynthetic enzymes necessary for PC synthesis from exogenous choline and fatty acids,^13^ our data imply that in vitro cultured *B. burgdorferi* primarily acquires these lipids from the medium.

**Fig. 3.**
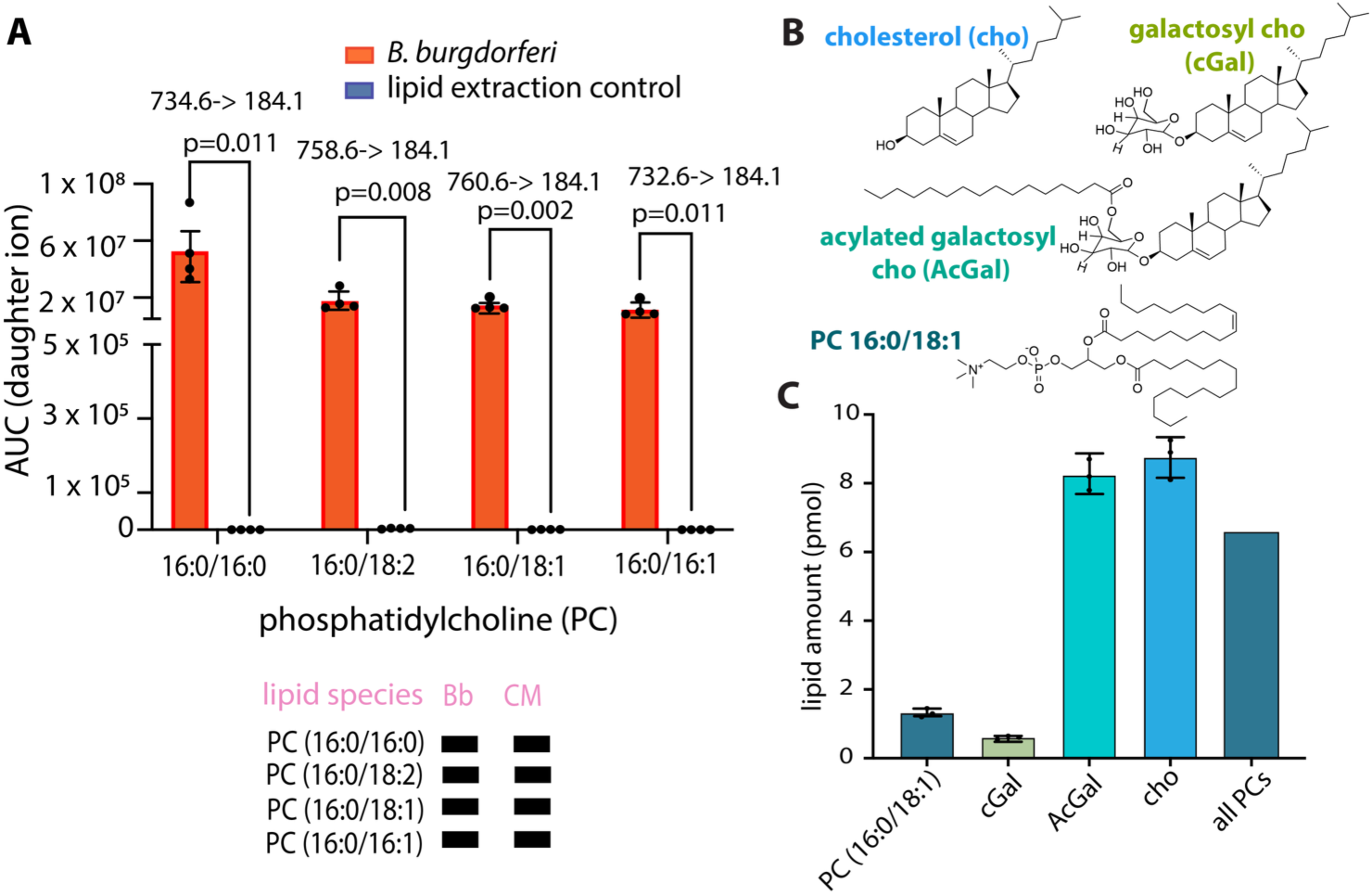
Targeted lipidomics analysis of key lipids in *B. burgdorferi*. (**A**) The upper left panel displays the abundance of the four most abundant PCs alongside the expected m/z values for fragmented ions. The lower left panel depicts the lipid species found in *B. burgdorferi*, indicating their presence (filled rectangles) or absence (empty rectangles) in the complete medium. Extracted lipids were analyzed using multiple reaction monitoring with identification based on retention time and distinct fragmentation patterns. (**B**) The top right panel shows the chemical structures of membrane lipids previously reported to be abundant. (**C**) The bottom right panel shows the quantification of POPC (PC 16:0/18:1), cholesterol, and cholesterol glycolipids performed using standards, alongside the estimation of all detected PCs using RF analysis.

To interrogate whether PCs represent a major lipid component in *B. burgdorferi*, we quantified one of the most abundant PC species (POPC) alongside free cholesterol, cholesteryl 6-O-acyl-β-D-galactopyranoside (AcGal or BbGL-I), and cholesteryl β-D-galactopyranoside (cGal) (**Fig. 3B**). Cholesterol emerged as the most abundant lipid (8.75 ± 0.95 pmoles/4 million cells), followed by AcGal (8.25 ± 0.63 pmoles/4 million cells), POPC (1.32 ± 0.09 pmoles/4 million cells), and cGal (0.60 ± 0.08 pmoles/4 million cells) (**Fig. 3C**). Our untargeted analysis indicates that the AcGal (C16:0) with m/z 804.6 species is consistently the most abundant, followed by three other species with stearic, oleic and linoleic acid^39, 40^ (**Fig. S11B**). Given its predominance and the availability of a high-quality commercial standard, we used the C16:0 species as a representative proxy for the total AcGal pool **(Fig. S11B)**.

Although absolute quantification of all 97 PC species was not possible due to the lack of commercial standards, we estimated their combined abundance using response factor calculations^41,42^ (see Methods). Validation of this approach using AcGal and POPC showed response factor values within 1–2-fold of the measured amounts (**Fig. S13**). The response factor-calculated total PC content (6.56 pmoles/4 million cells) remained lower than cholesterol levels (8.75 pmoles/4 million cells) but indicates that PCs, along with other phospholipids, might collectively constitute a significant proportion of *B. burgdorferi* lipidome.

### *B. burgdorferi* contains sphingolipids

Our data revealed the presence of other eukaryotic lipids in *B. burgdorferi* that appear to be acquired from the medium. Prominent amongst these are the sphingolipids sphingomyelin (SM) and ceramides (Cer) (**Figs. 4, S14 and S15**). While we do not currently understand the significance of these lipids in *B. burgdorferi*, sphingolipids play important roles in the physiology of eukaryotic membranes. SM is observed to be enriched on the outer leaflet of the plasma membrane of eukaryotic cells where it serves a structural role, but its breakdown products are also involved in signaling.^43^ Cer similarly function in providing membrane structure and signaling.^44^ *B. burgdorferi* was reported to bind ceramides as a means to adhere to host cells.^45^ More importantly*, B. burgdorferi* is known to contain lipid rafts,^12,40^ which are microdomains composed of cholesterol and sphingolipids that were first identified in eukaryotic cells.^46^ While it has long been established that *B. burgdorferi* incorporates an abundance of cholesterol in its membranes, our data now provide evidence for the presence of sphingolipids that could be stabilized by cholesterol in the observed lipid rafts.^47^

**Fig. 4.**
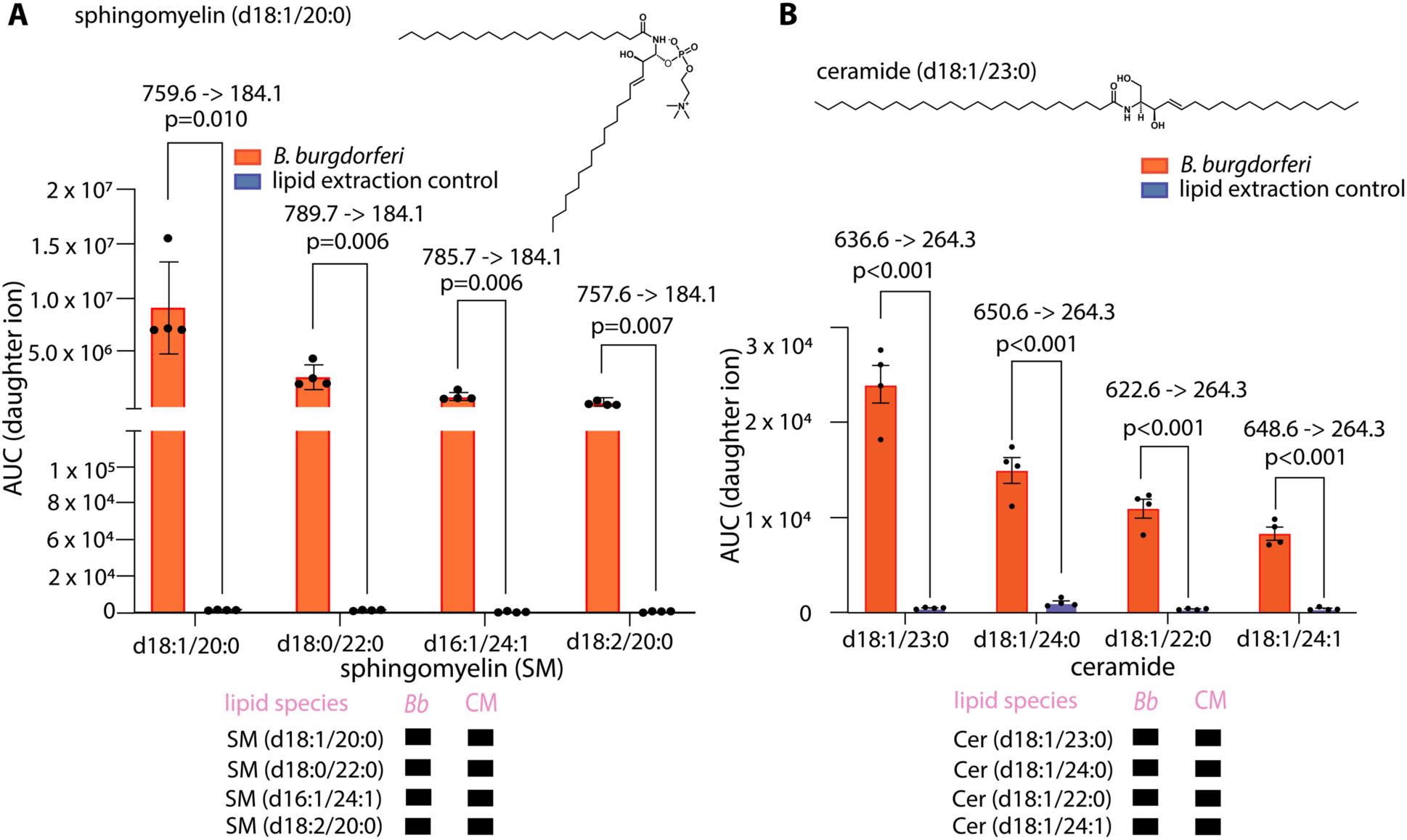
Targeted lipidomics analysis of predominantly eukaryotic lipids (**A**) sphingomyelin and (**B**) ceramides, identified in *B. burgdorferi*. Top panels show the chemical structures of sphingolipids in *B. burgdorferi*, while the middle panels show the abundance of each lipid species in the class and the expected m/z of fragmented ions identified in *B. burgdorferi*. The bottom panels show species detected in *B. burgdorferi* that were also present (filled rectangle) in the complete medium. Extracted lipids were analyzed using MRM, and species were identified on the basis of retention time and characteristic fragmentation patterns.

### *B. burgdorferi* performs lipid biosyntheses

The first indication that *B. burgdorferi* can synthesize lipids beyond what has been reported was our detection of 11 unique phosphatidyl glycerol (PG) species in the pathogen that are all absent in the CM. *B. burgdorferi* has the biosynthetic enzymes required for PG synthesis, and unlike the case with PCs, it might rely on those to synthesize this anionic lipid (**Figs. 5A and S16).** Also confirmed via MRM were three species of the zwitterionic phosphatidyl ethanolamine that were not detected in the CM (**Figs. 5B and S17**). The unusual bis(monoacylglycerol)phosphate (BMP) is another predominantly lysosomal lipid present in *B. burgdorferi*. We identified four BMPs in *B. burgdorferi* that were entirely absent from the CM. While distinguishing BMPs from their PG isomers can be challenging, their distinct retention time (RT) trendlines, driven by isomeric headgroup differences, strongly support these assignments.^48^ The fatty acyl tails of the BMPs detected all contained one or more unsaturation and carbon chain lengths 18-22 (**Figs. 5C and S18**). BMP use and synthesis is exceedingly rare in bacteria. One example of a BMP-synthesizing microbe is the plant pathogen *Agrobacterium tumefaciens* that uses lyso-PG as a substrate to make the diacylated BMP.^49^ Similarly, the plant symbiont *Sinorhizobium meliloti* can use exogenous lyso-PG to synthesize BMP.^49^ Because lyso-PG has not been detected in *B. burgdorferi* or in the medium, the synthesis pathway of BMPs in this organism remains unclear. A potential pathway may proceed via either the de-acylation, stereo-inversion, and re-acylation, or a direct stereo-inversion of PGs. While the role of BMPs in *B. burgdorferi* remains the focus of ongoing work, it has been speculated that the BMPs produced by bacteria are involved in membrane plasticity.^49^ They could also protect membranes from degradation by pH-sensitive lipases.

**Fig. 5.**
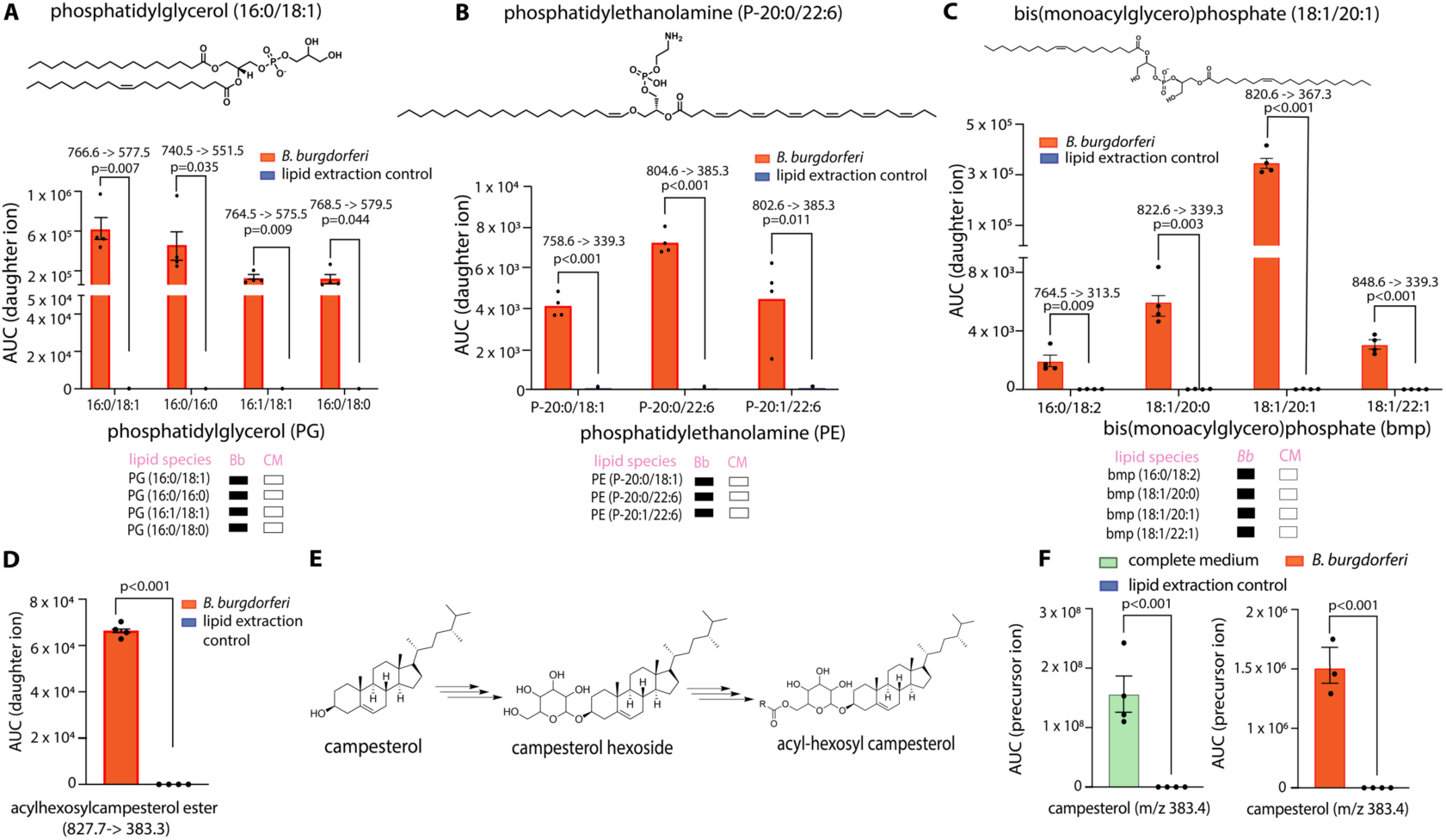
Targeted lipidomics analysis of glycerophospholipids that might result from biosynthesis. Top panels show the chemical structures of the lipids, while the middle panels show the abundance of each lipid species and the expected m/z of fragmented ions. The bottom panels show species detected in *B. burgdorferi* that were also present (filled rectangle) or absent (empty rectangle) in the complete medium. The four most abundant PGs (**A**), PEs (**B**), and BMPs (**C**) in *B. burgdorferi*. (**D**) Tandem mass spectrometry detection of acylhexosylcampesterol ester in *B. burgdorferi*. (**E**) Proposed biosynthetic route to the modified sterol, and (**F**) identification of campesterol in the growth medium.

### *B. burgdorferi* synthesizes glycolipids which are predominantly found in plants

MGDG is a glycolipid found in many plants and was previously detected in *B. burgdorferi*.^14^ Moreover, the *B. burgdorferi* protein BB0454 was identified as a galactosyltransferase sufficient for its biosynthesis.^14^ Consistent with this report, we detected multiple species of MGDG (fatty acid compositions 16:0/16:1, 16:0/18:1, 16:0/18:2, 18:1/18:1) (**Fig. S19 A-D**) in *B. burgdorferi*. Also, serum from individuals with late-stage Lyme disease shows an antibody response to MGDG and other glycolipids, suggesting a potential role in inflammation and immunity.^11,14^ However, our profiling revealed that MGDG is not the only plant glycolipid that is synthesized by *B. burgdorferi* (vide infra), and the importance of this finding or the relevance of these lipids on the pathogen’s physiology is ongoing work.

Acylhexosylcampesterol is an acylated and glycosylated steroidal lipid observed in the *B. burgdorferi*, which, to our knowledge, has never been detected in bacteria before. To validate this finding, we performed targeted lipidomics and confirmed the presence of the acylhexosylcampesterol exclusively in *B. burgdorferi* and not in the complete medium (CM) **(Fig. 5D)**. Given the absence of sterol biosynthesis in *B. burgdorferi*, we screened the culture medium for potential sterol precursor, campesterol. Using APCI-QTOF mass spectrometry, we confirmed the presence of campesterol, a phytosterol commonly found in plants^50^ in the CM and also detected in *B. burgdorferi* (**Fig. 5F**). As observed, the bacterium likely takes up the phytosterol, then glycosylates it to make a glycolipid and additionally acylates the sugar moiety in a manner similar to how it makes the cholesterol galactolipid AcGal (**Fig. 5E**).^11^

The untargeted lipidomics also revealed a 3-fold enrichment of the anionic sulfolipid sulfoquinovosyldiacylglycerol (SQDG) in *B. burgdorferi* when compared to the complete and spent media. Targeted lipidomics confirmed this through detection of both the precursor and product ions for three species (**Fig. 6A**). While SQDG is primarily associated with plants and photosynthetic bacteria,^22, 51–54^ its enrichment in *B. burgdorferi* and not in the CM suggests a biosynthetic origin. For plants and photosynthetic bacteria, SQDG is one of the least abundant lipid components in photosynthetic membranes and plays a crucial role in chloroplast development under phosphate-limiting conditions.^22, 51–54^ In *Arabidopsis thaliana,* the synthesis of SQDG requires at least two enzymes: UDP-sulfoquinovose synthase (SQD1), which utilizes sulfite, UDP-glucose, and NAD^+^ to produce UDP-sulfoquinovose, and a glycosyltransferase (SQD2) that transfers the modified sugar moiety to diacylglycerol (**Fig. 6B**). Structural bioinformatic analyses identified BB0444 and BB0454 as *B. burgdorferi* homologs of *A. thaliana* SQD1 and SQD2; BB0454 was previously reported to be a galactosyltransferase sufficient for MGDG synthesis.^14^ Structural alignment of AlphaFold2 models BB0444 with the crystal structure *A. thaliana* SQD1 (PDB 1QRR, **Fig. 6C**) and BB0454 with the AlphaFold model of *A. thaliana* SQD2 (**Fig. S20A**) shows strong homology, with a root mean square deviation (RMSD) of 1.071 Å over 233 pruned atom pairs for SQD1 and 1.088 Å over 235 pruned atom pairs for SQD2. For SQD1 (*A. thaliana*) and BB0444 (*B. burgdorferi*), the catalytic triad of Thr 145, Lys186, and Tyr 182 from *A. thaliana* align well with Thr 144, Lys 173 and Tyr 169 of BB0444 (**Fig. 6C**). In SQD1, the reactive 4′-hydroxyl of UDP-glucose clearly interacts directly with both Thr 145 and Tyr 182. Around the nicotinamide ribose moiety, the conserved Tyr 182 and Lys 186 side chains interact with the ribose hydroxyls, as expected in an enzyme from the short-chain dehydrogenase/reductase family.^55^

**Fig. 6.**
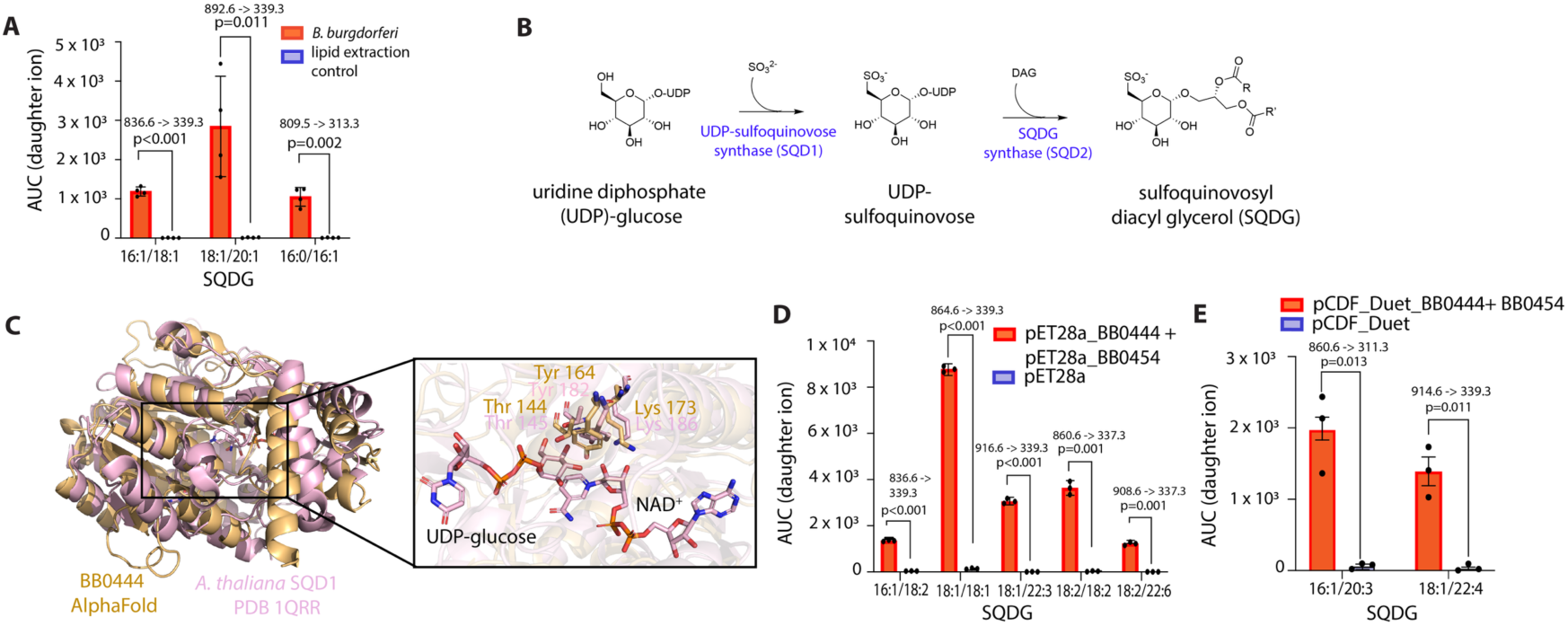
Lipid biosynthesis and modification in *B. burgdorferi*. (**A**) Tandem mass spectrometry detection of SQDG in *B. burgdorferi* and a proposed biosynthetic route to the lipid using *A. thaliana* as a model (**B**). (**C**) Structural alignment of *A. thaliana*’s SQD1 with the AlphaFold predicted structure of BB0444; essential active site residues shown in the inset. Reconstitution of SQDG biosynthesis via heterologously expressed BB0444 and BB0454 in *E. coli* (**D, E**).

To determine whether the *B. burgdorferi* enzymes BB0444 and BB0454 catalyze SQDG synthesis, we heterologously expressed these proteins in *Escherichia coli*, an organism for which SQDG production has not been reported (**Figs. S20B**). Co-incubation of lysates expressing BB0444 and BB0454 individually resulted in significant enrichment of five SQDG species compared to both empty vector controls and lysates containing either enzyme alone (**Fig. 6D**). This activity was additionally confirmed by co-expressing BB0444 and BB0454 in pETDuet-1 vector (**Fig. S20C**). Despite lower overall protein levels (**Figs. 6E, S20D and E**), lipid extracts from co-expressing cells showed a marked enrichment of two SQDG species (**Fig. 6E**), confirming that BB0444 and BB0454 are sufficient for SQDG biosynthesis in vitro. As before, SQDG biosynthesis was absent in lysates containing only BB0444 or BB0454 individually. This finding underscores the functional interdependence of BB0444 and BB0454 in synthesizing the sulfolipid.

To ascertain how widespread the synthesis of this lipid might be in the bacterial and archaeal domain, we created a protein sequence similarity network (SSN) with homologs of the *B. burgdorferi* proteins BB0444 and BB0454. This network included 9,441 sequences for BB0444 and 9,172 sequences for BB0454, all sharing an alignment score of 95 for BB0444 and 110 for BB0454, equating to a sequence identity cutoff ∼ 40%. With this cutoff, proteins are generally expected to retain a degree of functional homology, albeit with potential subtle differences.^56^ Both networks were visualized such that each node represents a unique protein sequence or sequences that are 100% identical (**Fig. S21** for BB0444 and **Fig. S22** for BB0454). To our knowledge, this SSN is the first to comprehensively display all annotated homologs of BB0444 and BB0454 in the bacterial and archaeal domains. The SSN of BB0454 highlighted several biochemically characterized homologs linked to SQDG biosynthesis, including *Sulfolobus acidocaldarius*,^21^ *Rhodobacter sphaeroides*,^22^ *Thermosynechococcus elongatus*^54^ (**Fig. S22A**). While the majority of proteins in the network are annotated as glycosyltransferases, others have annotated functions such as mannosylfructose-phosphate synthase and phosphatidylinositol α-mannosyltransferase activities. However, annotations are not always equivalent to biochemical or genetic validation. Regardless, it is encouraging that several homologs that cluster away from the *B. burgdorferi* enzyme have been reported to function in SQDG biosynthesis. It is therefore likely that the network contains true SQD2 homologs and some other glycosyltransferases (**Fig. S22**).

In contrast, no member of the BB0444 network has been characterized. It is notable that *A. thaliana* SQD1 and SQD2 were not included in the SSNs because they share low sequence identity to the *B. burgdorferi* proteins (12.67% for BB0444 and 16.71% for BB0454). The two SSNs additionally illustrate the conservation of these proteins within spirochetes, as homologs in spirochetes which are distinctly clustered in the network (highlighted in pink in **Figs. S21A, S22A**). Notably, these proteins have homologs in more than 30 species and subspecies within the *Borrelia* genus (**Figs. S21D, S22D**). It is therefore possible that the broad conservation of these proteins across pathogenic *Borrelia* species highlights their potential as targets for disrupting lipid synthesis or membrane biogenesis in *Borrelia*.

We infer that the presence of assorted plant glycolipids in *B. burgdorferi* highlights a sophisticated metabolic integration with its environment. These lipids are likely derived from precursors in the herbivorous rabbit serum used to supplement the BSK-II growth medium. If so, this ability to blend endogenous biosynthetic pathways with scavenged environmental resources allows *B. burgdorferi* to operate as a “lipid chimera”. The pathogen’s synthesis of SQDG, MGDG, and cholesterol galactolipids, paired with its modification of phytosterols like campesterol, reflects a dual strategy of metabolic economy and adaptation, ensuring survival across the diverse chemical landscapes of the tick vector and vertebrate host.

## Conclusion

This study delivers the most comprehensive and definitive lipidomic characterization of the metabolically restricted pathogen *Borrelia burgdorferi* to date. By integrating multi-platform untargeted and targeted liquid chromatography-mass spectrometry (Orbitrap, triple quadrupole, and quadrupole time-of-flight) with gas chromatography-mass spectrometry, we have constructed a high-resolution molecular database that fundamentally redefines our understanding of spirochetal metabolic adaptability. Our comparative analyses demonstrate that *B. burgdorferi* does not merely tolerate its stripped-down metabolic state; rather, it actively and precisely sculpts its cellular envelope by scavenging a sophisticated array of environmental and host-derived lipids. The discovery that this pathogen incorporates host-derived signaling molecules, specifically complex sphingolipids and ceramides, transcends the traditional view of these molecules as basic structural elements or simple adhesion factors.^45^ Instead, it opens entirely new horizons for investigating lipid-mediated host-pathogen communication during infection. Furthermore, our rigorous quantitative framework firmly establishes phosphatidylcholine as one of the principal membrane constituents alongside cholesterol and cholesterol galactolipids, rewriting the established model of the *B. burgdorferi* membrane envelope. Equally transformative are the structural and compartmental insights uncovered herein. We provide the first visual and molecular evidence of structured lipid storage within *B. burgdorferi*, demonstrating that neutral lipids are organized into distinct intracellular foci, a major paradigm shift for this minimal bacterium. Additionally, the identification of BMP, lipids typically restricted to eukaryotic endolysosomal systems, suggests an unexpected, sophisticated mechanism driving membrane plasticity and stress adaptation.^49^ Mechanistically, we successfully mapped out the biosynthetic architecture of the unique sulfolipid SQDG, pinpointing the enzymes BB0444 and BB0454 as the central catalysts of this pathway. Beyond *B. burgdorferi*, large-scale bioinformatic tracking revealed that these biosynthetic homologs are widely distributed across diverse bacterial phyla and tightly co-occur throughout the spirochete domain, highlighting a potentially universal evolutionary strategy for envelope integrity.

In summary, this compendium redefines the model of lipid metabolism in *B. burgdorferi*, showcasing a sophisticated dual strategy of intensive lipid scavenging and unique biosynthetic adaptations. By uncovering these pathways, our resource lays the groundwork needed to functionally interrogate lipid pathways during infection and proliferation. Key processes like sulfolipid synthesis and neutral lipid storage are now highlighted as compelling targets for future studies. Ultimately, the multi-platform lipidomics workflow developed here establishes a robust blueprint for discovering novel biomarkers and therapeutic targets in other metabolically dependent pathogens, opening powerful new avenues for innovative and targeted antimicrobial strategies.

## Resource availability

### Lead contact

Further information and requests for resources and reagents should be directed to Prof. Laura M. K. Dassama.

### Materials availability

Materials and reagents reported in this manuscript will be shared by the lead contact upon request.

### Data and code availability

All custom analysis workflows, including Python scripts, are available on the Dassama lab GitHub repository (https://github.com/dassamalab/Borrelia-lipidomics_Chatterjee_et_al_2026). Any additional information required to reanalyze the data reported in this paper is available from the lead contact upon request.

## Acknowledgments

We thank the Abu-Rameileh laboratory for helpful discussions and access to the LC-triple quadrupole mass spectrometer used for targeted lipidomics. The authors acknowledge the use of facilities and technical assistance of the Stanford Doerr School of Sustainability Geomicrobiology Shared Labs (GML) (RRID: SCR_025000). This work was supported in part by NIH grant R35GM150910 (to L.M.K.D.) and by the Howard Hughes Medical Institute Emerging Pathogens Initiative (to C.J.W. and L.M.K.D.). L.M.K.D. was additionally supported by a Terman Fellowship from Stanford University and MAC3 Impact Philanthropies Faculty Fellowship from the Sarafan ChEM-H Institute. I.A.P. is an NSF GRFP and HHMI Gilliam Fellow. M.I.T is supported by an NSF GRFP. C.J.-W. is an investigator of the Howard Hughes Medical Institute.

## Methods

### Bacterial strains and growth conditions

*B. burgdorferi* strain B31 MI clone A3 (B31-A3))^57^ cells were cultured in complete Barbour-Stoenner-Kelly (BSK)-II medium in humidified incubators at 34°C under a 5% CO_2_ atmosphere. The complete BSK-II medium consisted of 50 g/L bovine serum albumin (BSA, Millipore #81-003), 9.7 g/L CMRL-1066 (US Biological #C5900-01), 5 g/L neopeptone (Thermo Fisher #211681), 6 g/L HEPES acid (Millipore #391338), 5 g/L D-glucose (Sigma #G7021), 2 g/L yeastolate (Difco #255772), 0.7 g/L sodium citrate (Sigma #C7254), 0.8 g/L sodium pyruvate (Sigma #P5280), 2.2 g/L sodium bicarbonate (Sigma #S5761), and 0.4 g/L N-acetyl-glucosamine (Sigma #A3286). The pH of the medium was adjusted to 7.6 and then supplemented with 60 mL/L of heat-inactivated rabbit serum. The heat inactivation process involved incubating the serum at 50°C in a water bath for 30 minutes prior to its use in medium preparation. Unless otherwise specified, all *B. burgdorferi* cultures were maintained in the exponential phase at a density below 5 × 10^7^ cells/mL. Cell density was determined through direct counting using disposable Petroff-Hausser chambers and darkfield microscopy, as previously described by Takacs et al. (2018). For sample collection, 90 mL of culture was centrifuged at 8,000 × g for 10 minutes. The supernatant (spent medium) was collected, and the resulting cell pellets were washed three times with H-N buffer (50 mM NaCl, 10 mM HEPES pH 8.0) before being stored at -80°C.

### Total lipid extraction and subsequent analysis by thin layer chromatography (TLC)

Lipids from *B*. *burgdorferi* B31-A3 cells, complete medium (complete BSKII medium), and spent medium were extracted using the Bligh-Dyer method.^58^ In brief, 2 mL of complete media and spent media were used, along with cell pellets that were dissolved in water. To account for potential background noise introduced during the lipid extraction process, parallel extraction controls were prepared by subjecting 2 mL of water to the identical extraction protocol (termed lipid extraction control). To prepare the samples for extraction, a methanol and dichloromethane (DCM) mixture was prepared in a 2:1 ratio within a glass tube. The sample, supplemented with a deuterated lipid internal standard (Splashmix, Avanti #330709) at a 1:1000 ratio, was then introduced into the solvent mixture. The final volumetric ratio was adjusted to maintain a methanol: DCM: water proportion of 10:5:4. The samples were vortexed and sonicated in a water bath for at least one hour. Additional DCM and water were added in a 1:1 ratio, followed by another vortexing step, and the samples were stored at -20°C overnight. The glass tubes containing the solvents were then centrifuged at 2,800 RCF for 10 minutes at 4°C in a swinging bucket rotor to fully separate the layers. The organic phase was carefully transferred into a clean, baked glass tube using a baked Pasteur pipette.

For the qualitative analysis of lipids using thin layer chromatography, three primary solutions were prepared: mobile phase 1, consisting of ethanol, chloroform, triethylamine, and water in a ratio of 40:35:35:9 (v/v/v/v); mobile phase 2, a mixture of n-hexane and ethyl acetate at 5:1 (v/v); and a visualization reagent composed of 20% sulfuric acid (H_2_SO_4_) in ethanol. To ensure safety, the ethanol was measured first before the H_2_SO_4_ was slowly added. Silica glass plates were prepared by marking a horizontal origin line for sample application. Lipid extracts dissolved in DCM were applied to these points using a capillary tube, with multiple applications performed on the same spot, allowing each to dry completely before the next to ensure sufficient concentration without spreading.

Chromatographic separation was achieved through a sequential two-stage development process. First, the plates were placed in a 400 mL glass beaker containing mobile phase 1 and a filter paper liner for chamber saturation. The solvent was allowed to migrate via capillary action until the front reached one inch from the top of the plate, ensuring the initial solvent level did not submerge the origin line. After removal and room-temperature drying, the plates were developed a second time in a 400 mL beaker saturated with mobile phase 2. This second run was allowed to proceed further than the first to optimize the separation of neutral lipids. Following development, the plates were dried at room temperature, with the aid of N_2_ gas if necessary. For visualization, the plates were quickly immersed in a petri dish containing the H_2_SO_4_ solution and subsequently charred in an oven at 200°C for five minutes until the lipid species appeared as distinct spots.

### Untargeted fatty acid profiling using gas chromatography mass spectrometry (GC-MS)

Total lipid extracts were massed and resuspended at a concentration of 1 mg/mL in dichloromethane. Samples were then dried under a nitrogen stream and derivatized to trimethylsilyl ethers by adding 1:1 (vol: vol) Bis(trimethylsilyl)trifluoroacetamide: pyridine and heating at 70°C for 1 hour. 2 μg from each sample was injected into an Agilent 7890B Series GC in splitless mode at 250°C. Lipids were separated using a 60 m Agilent DB17HT column (60 m x 0.25 mm i.d. x 0.1 μm film thickness) with helium as the carrier gas at constant flow of 1.1 mL/min and programed as follows: 100°C for 2 minutes; then 8°C/minute to 250°C and held for 10 minutes; then 3°C/minutes to 330°C and held for 17 minutes. The GC was coupled to a 5977 A Series MSD with the ion source held at 230°C and operated at 70 eV in EI mode scanning from 50-850 Da in 0.5 s. Collected data was analyzed using Agilent MassHunter Qualitative Analysis (B.06.00). Fatty acids were identified on retention time and spectra by comparison to previously confirmed laboratory standards or spectra deposited in the National Institute of Standards and Technology (NIST) databases.

### Untargeted tandem mass spectrometry using ID-X Tribrid mass spectrometer

For mass spectrometry sample preparations, the lipids were concentrated at 35°C under nitrogen gas. The dried lipids were resuspended in a lipid reconstitution buffer composed of acetonitrile, 2-propanol, and water in a 13:6:1 (v/v/v) ratio. The mixture was vortexed for 10 minutes and centrifuged at 13,000 RCF at 4°C for 15 minutes. The upper layer was carefully transferred into glass insert vials for mass spectrometry. Three biological replicates and three technical replicates were prepared for this experiment.

Lipid analysis was performed using an ID-X Tribrid mass spectrometer (Thermo Fisher Scientific) equipped with a heated electrospray ionization (HESI) probe. Lipid separation was achieved using an Agilent C18 column (150 x 2.1 mm). The mobile phases consisted of water and acetonitrile for Phase A, and 2-propanol and acetonitrile for Phase B, both supplemented with 10 mM ammonium formate and 0.1 % formic acid for positive ion mode, and ammonium acetate (and 0.1 % formic acid) for negative mode. A specific gradient was employed to separate the lipids, starting with isocratic elution and gradually increasing Phase B to 97% over 21 minutes. The flow rate was set to 0.26 mL/min, with an injection volume of 4 µL, and the column temperature maintained at 55°C.

For mass spectrometry, the ion transfer tube and vaporizer temperatures were set to 300°C and 375°C, respectively. The resolutions for MS^1^ and MS^2^ were 120,000 and 30,000, covering a mass-to-charge ratio (m/z) range of 150-1500. Ionization was performed in both positive and negative modes, with fragmentation achieved using stepped HCD at 15%, 25%, and 35%. Data-dependent tandem MS^2^ (ddMS^2^) was utilized with a cycle time of 1.5 seconds. The results were analyzed using LipidSearch (https://www.thermofisher.com/us/en/home/industrial/mass-spectrometry/liquid-chromatography-mass-spectrometry-lc-ms/lc-ms-software/multi-omics-data-analysis/lipid-search-software.html) and Compound Discoverer for differential lipid analysis, with annotations based on precursor and product tolerances of 5 ppm. The mass list was exported for further quantification, and data processing included specific settings for peak intensity, retention time, and background filtering.^59^

### Identification of sterols and quantification of POPC, cholesterol and cholesterol glycolipids

The following lipid standards were purchased from Avanti: cholesteryl 6-O-palmitoyl-β-D-galactopyranoside (AcGal, #700193), Galactosyl Cholesterol (cGal, #700187), and 16:0/18:1 PC (POPC, #850457). Free cholesterol was obtained from Thermo Scientific Chemicals (#A1147018). The quantification of cholesterol from four million *B. burgdorferi* B31 cells, was estimated using atmospheric pressure chemical ionization (APCI) on an Agilent QTOF 6530 mass spectrometer, coupled with an Agilent 1290 Infinity II ultra-high-pressure liquid chromatography (UHPLC) system. Atmospheric pressure chemical ionization (APCI) was selected for the analysis of native sterols. Sterols, such as cholesterol, are more apolar and lack easily ionizable functional groups (e.g., primary amines or carboxylic acids). Consequently, they are poorly ionized by electrospray ionization (ESI), a technique that relies on the formation of pre-charged ions in solution, which leads to significantly lower sensitivity for these compounds. In contrast, APCI is a gas-phase ionization method that is highly effective for analyzing less polar and thermally stable molecules. It ionizes neutral analytes via proton transfer reactions with charged solvent molecules in the gas phase. For sterols, this process typically yields a highly abundant characteristic protonated and water-loss fragment [M+H-H₂O]⁺, providing both sensitive detection and structural confirmation. While ESI can be employed for sterol analysis following chemical derivatization to introduce a charged group, APCI offers the distinct advantage of allowing for the direct, high-sensitivity analysis of these lipids in their native form, thus simplifying the analytical workflow.^60^ Campesterol was also detected in complete media using the same method as cholesterol. In contrast, POPC, cGal, and AcGal were quantified using electrospray ionization (ESI).

Samples were separated on a C18 column with a pore size of 95 Å, a particle size of 1.8 μm, and dimensions of 2.1 x 50 mm (Agilent #959757-902). External mass calibration was performed before every set of sample run using a QTOF standard tuning mix. The buffers used were (A) LCMS-grade water (#W6500, Fisher Chemical) and (B) methanol (#A4564, Fisher Chemical), with a gradient from 80% to 100% B over 7 minutes for APCI. The column temperature was maintained at 40°C throughout the experiment (for both APCI and ESI). A sample volume of 5 microliters was injected per run, with a flow rate of 0.6 mL/min. For ESI, the mobile phases consisted of (A) water (#W6500, Fisher Chemical) and acetonitrile (#A955-4, Fisher Chemical), and (B) 2-propanol (#102781, Millipore Sigma) and acetonitrile, each containing 10 mM ammonium formate (#A666, Fisher Chemical) and 0.1 % formic acid (#A117-50, Thermo Fisher Scientific). A linear gradient from 15% to 99% B was applied over 7 minutes at a flow rate of 0.5 mL/min. MS/MS data were acquired in both positive and negative ion modes over a mass-to-charge (m/z) range of 100–1700. Source parameters included gas temperatures of 325°C for both ESI positive and APCI modes, a vaporizer temperature of 350 °C for ESI positive and APCI positive modes, and drying gas flows of 12 L/min for ESI and 4 L/min for APCI. Capillary voltage settings were ±3500 V for ESI and ±2000 V for APCI. The MS scan rate was set to 4 spectra/s, while the MS/MS spectra were collected at 2 spectra/s. The fragmentor voltage was set to 150 V for MS, with collision energies of 10, 20, and 40 V (positive mode) for MS/MS data collection. The autosampler was kept at 4°C throughout. The concentration (picomoles) of each lipid in *B. burgdorferi* B31 was calculated by running a standard curve for each sample.^61^

### Targeted tandem mass spectrometry using triple quadrupole

Targeted analyses were performed with complete medium, spent medium, and *B. burgdorferi* B31 samples using methods previously described.^62,63^ Lipids were extracted and prepared as mentioned above. The samples were separated on an Agilent RRHD Eclipse Plus C18, 2.1 × 100 mm, 1.8 µm column with an Agilent guard holder (UHPLC Grd; Ecl. Plus C18; 2.1 mm; 1.8 µm), The liquid chromatography system was linked to an Ultivo triple quadrupole (QQQ) mass spectrometer with a liquid chromatography–electrospray ionization probe. External mass calibration was performed weekly using a QQQ standard tuning mix. The column compressor and autosampler were held at 45 °C and 4 °C, respectively. For lipidomics, the mass spectrometer parameters included a capillary voltage of 4.4 kV in positive mode and 5.5 kV in negative mode, and the gas temperature and sheath gas flow were held at 200 °C and 275 °C, respectively. The gas flow and sheath gas flow were 10 and 11 l min^−1^, respectively, whereas the nebulizer was maintained at 45 psi. The nozzle voltages were maintained at 500 in the positive mode and 1,000 in the negative mode. These conditions were held constant for both ionization mode acquisition. For lipid measurements, 5 µL injection volumes (equivalent to lipid extract from four million Borrelial cells) were used for each sample for polarity switching for each method with a total ten methods being created. A mobile phase with two distinct components (A and B) was used in the chromatographic process. Mobile phase A was a mixture of acetonitrile:water (2:3 (v/v)), whereas mobile phase B was composed of isopropanol:acetonitrile (9:1 (v/v)), both containing 0.1% formic acid and 10 mM ammonium formate. The elution gradient was carried out over a total of 16 min, with an isocratic elution of 15% B for the first minute, followed by a gradual increase to 70% of B over 3 min and then to 100% of B from 3 to 14 min. Subsequently, this was maintained from 14 to 15 min, after which solvent B was reduced to 15% and maintained for 1 min, followed by an extra 2 min for column re-equilibration. The flow rate was set to 0.400 ml min^−1^. The QQQ was set to operate in multiple reaction monitoring (MRM) to analyze compounds of interest. Standard lipids including glycerophosphodiesters were optimized using the MassHunter Optimizer MRM, a software used for automated method development. For most species, the one or two most abundant transitions were selected to detect it. The precursor–product ion pairs (*m*/*z*) of the compounds used for MRM are listed in **Supplementary file 4**. Lipids were annotated, and their abundances were measured using the Qualitative MassHunter acquisition software and QQQ quantitative analysis software (Quant My-Way), respectively. The peak areas of all metabolites were integrated using the retention time and MRM method, and the resulting raw abundances were exported to Microsoft Excel for further analysis. The raw abundances of internal controls and highly abundant lipids were examined to ensure the accuracy of the analysis. Control samples were consistently measured to ensure optimal instrument performance and linearity.

### Semi-quantitative profiling of phosphatidylcholines using calculated response factors

To quantify the lipid composition of *B. burgdorferi*, we employed a semi-quantitative approach based on the principle of equimolar response within the different PCs.^41,42^ High-sensitivity Multiple Reaction Monitoring (MRM) transitions were utilized on a triple quadrupole mass spectrometer (QQQ) to identify and profile 97 individual PC species with diverse fatty acid compositions.

To convert raw peak areas into molar amounts, a class-specific response factor (RF) was determined. Calibration was performed on a QTOF instrument to leverage its high mass accuracy and wide linear dynamic range. A representative standard, 1-palmitoyl-2-oleoyl-glycero-3-phosphocholine (POPC) (Avanti, POPC, #850457), was used to generate a multi-point calibration curve. Following established methodologies,^41,42^ the RF was calculated. This RF (expressed in picomoles per area unit) was subsequently applied to the peak areas of all PC species identified in the QQQ analysis to estimate their respective molar abundances. This calculation assumes that species within the same headgroup class exhibit near-uniform ionization efficiencies (daughter ion for PC was m/z =184.1).^41,42^ To ensure analytical consistency, the same sample pool, derived from four million *B. burgdorferi* cells, was utilized for both the QTOF calibration and the QQQ profiling. The total PC pool was defined as the sum of all estimated molar amounts. These values were integrated with the absolute molar concentration of cholesterol, which was determined independently via APCI mass spectrometry using QTOF. This combined dataset provided a semiquantitative estimation of the total PC pool relative to cholesterol, establishing the experimental baseline for the molar proportions of these lipids in *B. burgdorferi*.

### Widefield epifluorescence microscopy to observe lipids in *B. burgdorferi*

For the preparation of BSK-II agarose pads, concentrated BSK-II medium without phenol red was made. This medium contains three times the concentration of all BSK-II components, except for BSA and the rabbit serum, which were respectively in 2x and 1.5x concentration compared to the regular BSK-II medium recipe due to solubility issues. To avoid boiling of the BSK-II medium, low-melting-temperature SeaPlaque® GTG® agarose (Lonza, 50111) was resuspended in MilliQ water to create a 4.5% agarose solution. This agarose solution was initially polymerized by incubation at 80°C for at least 1 hour. Subsequently, both the agarose solution and 3x BSK-II medium were incubated at 45°C in separate tubes before mixing. Once heated to 45°C, an appropriate volume of the 4.5% low temperature agarose solution was mixed with the concentrated BSK-II medium to achieve a final 3% agarose concentration. The mixture was vortexed and then centrifuged briefly to remove bubbles. The mixture was returned to the heat block and intermittently mixed with gentle flicking until it appeared homogeneous. The BSK-II agarose mixture was not left on the heat block for more than 5 minutes to avoid overheating the components of the BSK-II medium. Finally, 100 μL of the BSK-II agarose mixture was quickly pipetted onto a glass slide and sandwiched with another glass slide to create a 1.5 mm thick pad. The pad was dried for approximately 30 minutes before spotting the cell sample.

To prepare the sample for lipid staining, *B. burgdorferi* B31-A3 cells were cultured in complete BSK-II medium without phenol red at 34°C under 5% CO₂. For each sample, B31-A3 cultures were grown to mid-exponential phase (cell density ≤ 5 x 10^7^ cells/mL). Cell density was determined using a Petroff-Hausser chamber (INCYTO C-Chip disposable hemocytometer) and darkfield microscopy. A total of 3 x 10^7^ cells were collected by centrifugation at 5,000 × g for 5 minutes, and the medium was aspirated. Cells were washed twice with 1 mL of 1× PBS supplemented with 1 mM MgCl_2_. Cells were then fixed in 1 mL of 4% formaldehyde solution prepared in 1× PBS supplemented with 1 mM MgCl_2_. The 4% formaldehyde fixing solution was diluted from a 16% stock, which was made by diluting methanol-free 20% formaldehyde (Ladd Research, CAT#20300) in 120 mM sodium phosphate buffer, pH 7.4 (sodium phosphate buffer: 1 M NaH_2_PO_4_, 1 M Na_2_HPO_4_, pH 7.4). The cells were incubated in the fixing solution for 5 minutes at room temperature and then for 10 minutes on ice. Fixed cells were centrifuged, the fixing solution was aspirated, and the cells were washed twice with 1 mL of 1× PBS supplemented with 1 mM MgCl_2_. Cells were resuspended in 100 μL of either 1:200 HCS LipidTOX™ Deep Red Neutral Lipid Stain (Invitrogen, #H34477) or 1:200 Nile Red from a 1 mg/mL stock (Sigma-Aldrich, #19123-10MG). An equivalent volume of DMSO (0.5% final concentration) was added to the control samples to match the dye preparation. After a 30-minute incubation with the stains, cells were washed twice with 1× PBS supplemented with 1 mM MgCl_2_. Finally, the cell pellet was resuspended in 40 μL of 1× PBS supplemented with 1 mM MgCl_2_. A 1 μL aliquot of each sample was then immobilized on BSK-II agarose pads prepared as mentioned above. To reduce background fluorescence, coverslips (no. 1.5, 22 x 50 mm, VWR, 48393-195) and glass slides (Fisherbrand, 125444) were cleaned by immersing them in 2% Hellmanex III, 1M KOH, and 200-proof ethanol, with each immersion followed by sonication for 30 minutes. Cover slips and glass slides were rinsed in deionized water between each immersion. Finally, they were plasma-cleaned (Tergeo Pro plasma system, PIE Scientific).

Darkfield microscopy was performed using a Nikon Eclipse E600 microscope equipped with a 40×/0.55 NA Ph2 air objective (Nikon) to determine cell density. For widefield epifluorescence microscopy experiments, snapshots were taken using a Nikon Ti2-E inverted microscope equipped with a Plan Apo 100×/1.45 NA Ph3 oil objective (Nikon), a Photometrics Prime BSI back-illuminated sCMOS camera (2048×2048 pixels sensor with a pixel size of 6.5 μm), and a Lumencor Spectra III Light Engine. The microscope featured a Perfect Focus system, a motorized stage, and a microscope enclosure with dark panels (Okolab). Images were acquired using NIS-Elements AR (version 5.21.03) Microscope Imaging software. For Cy3 (Nile Red) and Cy5 (LipidTOX deep red) channels, a polychroic mirror (Semrock, FF-409/493/573/652/759-Di01-25×36) combined with a triple-pass emitter (Semrock, FF01-432/515/595/681/809-25) was used and for Cy3 (Nile red) imaging, an additional emission filter (Semrock, FF01-595/31-25) was applied in the optical path. Cy3 (Nile red) and Cy5 (LipidTOX deep red) were excited using 555/28nm and 637/12nm, respectively with both light sources set to 40% transmission; exposure times for Cy3 and Cy5 were 200 ms and 1 s, respectively.

For image analysis, the background fluorescence across the entire field of view was reconstructed using the background correction pipeline as previously described.^64^ The analysis pipeline for background subtraction is available at the Jacobs-Wagner lab GitHub at https://github.com/JacobsWagnerLab/published/blob/master/Papagiannakis_2025/snap <u>shots_analysis_UNET_ghv.py</u>. Images were further analyzed with Fiji.^65^ For visualization and accurate comparison, the Look-Up Tables (LUTs) were scaled relative to the control images. Images were cropped for the figure preparation.

### Biochemical characterization of SQD1 and SQD2 enzyme activity

The DNA sequences encoding BB0444 and BB0454 (**Table S1**) were synthesized without stop codons and codon-optimized for *E. coli*. Each gene was individually cloned into the pET-28a vector using the NheI and XhoI restriction sites, incorporating an N-terminal His-tag, for heterologous expression in *E. coli* BL21(DE3) cells. Additionally, both genes were co-cloned into the pETDuet-1 vector: BB0444 was inserted between the BamHI and EcoRI restriction sites with N-terminal histidine tag, while BB0454 was cloned into the KpnI and XhoI sites with C-terminal strep-tactin tag. Transformed *E. coli* cells were cultured at 37°C in Terrific Broth,^66^ without phosphate until reaching an OD 600_nm_ of 0.6. Protein expression was induced by adding 0.5 mM isopropyl β-D-1-thiogalactopyranoside (IPTG), followed by incubation at 18°C while shaking for 12 hours. Cells were harvested via centrifugation (8,000 × g, 15 min) and resuspended in lysis buffer containing 25 mM Tris-HCl (pH 8.0), 200 mM NaCl, 2-mercaptoethanol (pH=8.2), and 1 mM phenylmethylsulfonyl fluoride (PMSF). Cell lysis was performed using a microfluidizer at 17,000 psi for four passages. The lysate was clarified by centrifugation at 15,000 RPM for 30 minutes. Western blotting with anti-polyHistidine-Peroxidase antibody, Mouse monoclonal (Sigma-Aldrich (#A7058), 1:10000, and HRP visualization) was performed to confirm expression of recombinant BB0444 and BB0454.

For enzyme activity assays, lysates (∼5g of cells) from individual clones of BB0444 and BB0454 (cloned into the pET-28a vector) were incubated on ice for 2 hours, and reactions were quenched with methanol and dichloromethane (DCM) to extract lipids. The pETDuet-1 vector containing both DNA constructs was processed using the same protocol, with the exception that lipid extraction was performed immediately after cell lysis. Western blot analysis was also conducted to confirm protein expression, followed by targeted mass spectrometry using a triple quadrupole mass spectrometer to analyze lipid profiles.

### Sequence similarity network analysis of BB0444 and BB0454

Protein sequence similarity network (SSN) of BB0444 and BB0454 depicting size and functional clustering. The SSN was generated via the Enzyme Function Initiatives-Enzyme Similarity tool (EFI-EST, https://efi.igb.illinois.edu/efi-est/) and visualized in Cytoscape45 with an “organic” layout. The SSN was generated by employing a single sequence blast of *B. burgdorferi* BB0444 and BB0454 and tailored so that the nodes represent sequences with 100% identity, an e-value of 10^10^, and an alignment score of 150 and 95 for BB0444 and BB0454 respectively. This analysis included 9,441 sequences for BB0444 and 9,172 sequences for BB0454. BB0444 had 474,291 edges (9,433 nodes) while BB0454 had 806,370 edges (8,721 nodes).

### Computational analyses

Bar plot: Differential feature tables comparing *B. burgdorferi* to the lipid-extraction control (water blank) in positive- and negative-ion electrospray (ESI) modes were exported from Compound Discoverer to Excel and analyzed in Python (v3.12.4) using pandas,^67^ NumPy,^68^ seaborn,^69^ and matplotlib^70^ For each feature, log2 fold change and Benjamini–Hochberg adjusted P values were extracted, and adjusted P values were transformed to −log10(adjusted P) for visualization. Statistical significance was defined as adjusted *p* < 0.05 with |log2 fold change| ≥ 2; entries lacking compound annotation were excluded. Lipid-class enrichment was summarized by assigning lipid class from the compound name prefix and calculating, within each class, the mean log2 fold change and mean −log10(adjusted P) across retained significant features. Class-level enrichment was visualized as bar plots of mean log2 fold change, with bar color mapped to mean −log10(adjusted P) using a continuous “flare” color scale. (**Figs 1 A and B**)

Volcano plots: Differential feature tables were exported to Excel from Compound Discoverer and analyzed in Python (v3.12.4; pandas, NumPy, seaborn, matplotlib). Log2 fold change values and Benjamini–Hochberg adjusted P values were extracted; adjusted P values were transformed to −log10(adjusted P) for plotting. Features were considered significant at adjusted *p* < 0.05 and |log2 fold change| ≥ 2. Entries lacking compound annotation were excluded. For significant features with log2 fold change ≥ 2, lipid class was assigned from the compound name prefix preceding the first parenthesis and points were colored by lipid class using a predefined color palette. Volcano plots were generated as scatter plots of log2 fold change versus −log10(adjusted P), with non-significant features shown in gray and significant features overlaid in class-specific colors. A dashed horizontal line indicating adjusted P = 0.05 was included (**Figs 1F-I, S5, S6 and S7**)

EIC plots: For representative species, extracted-ion chromatograms (EICs) were generated in Agilent Mass Hunter Qualitative Analysis and exported as TXT files. EICs were processed in Python (v3.12.4) using a custom script using pandas, NumPy, seaborn, and matplotlib to compare four biological replicates per condition (*B. burgdorferi*, complete medium, and lipid-extraction control). Chromatograms were restricted to predefined retention-time windows (rt_min–rt_max), interpolated onto a common retention-time grid, and summarized as mean extracted-ion counts with variability shown as mean ± SEM (SEM = SD/√n) using shaded bands (**Figs S8-S9, S11-S12, and S14-S19**).

## Supporting Information for

### Supporting figures

**Fig. S1.**
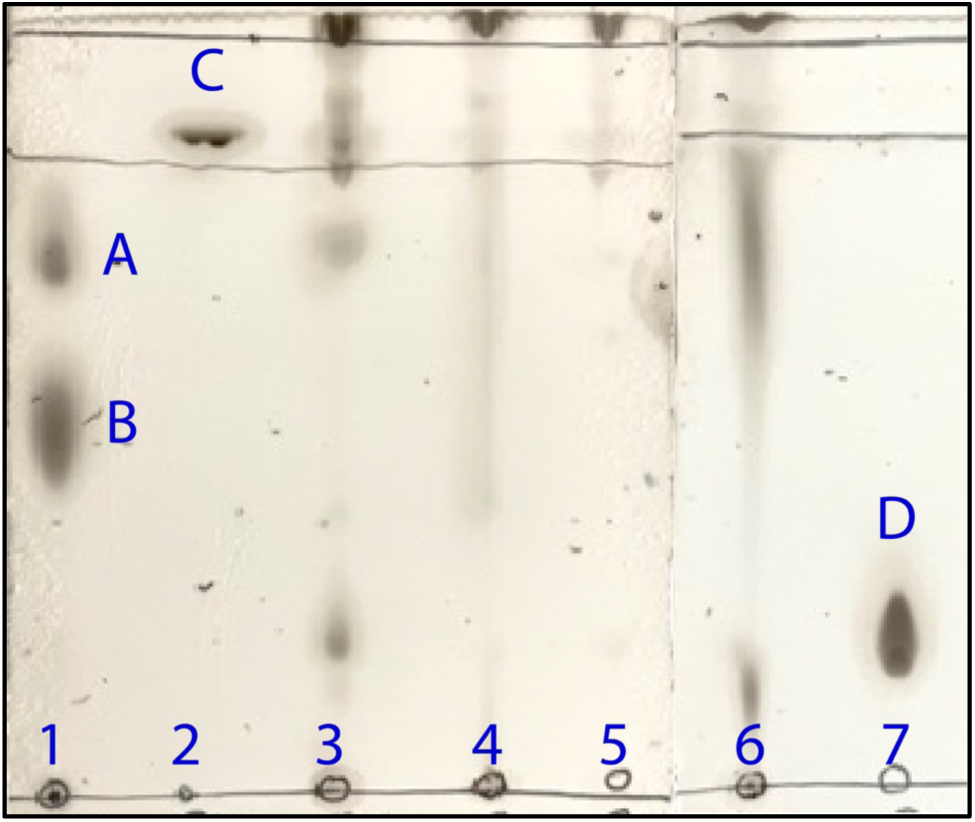
Qualitative analysis of lipids was performed in rabbit serum, complete medium, spent medium, *Borrelia burgdorferi*, and lipid standards using Thin Layer Chromatography (TLC). The lanes are designated as follows: total polar lipid extract of *Escherichia coli* (standard lipids are annotated as observed in the #Avanti research (1), cholesterol standard (2), total lipid from rabbit serum (3), total lipid from complete medium (4), total lipid from spent medium (5), total lipid from *Borrelia burgdorferi* (6), and phosphatidylcholine standard (7). On the TLC plate, phosphatidylglycerol is labeled A, phosphatidylethanolamine B, cholesterol C, and phosphatidylcholine D.

**Fig. S2.**
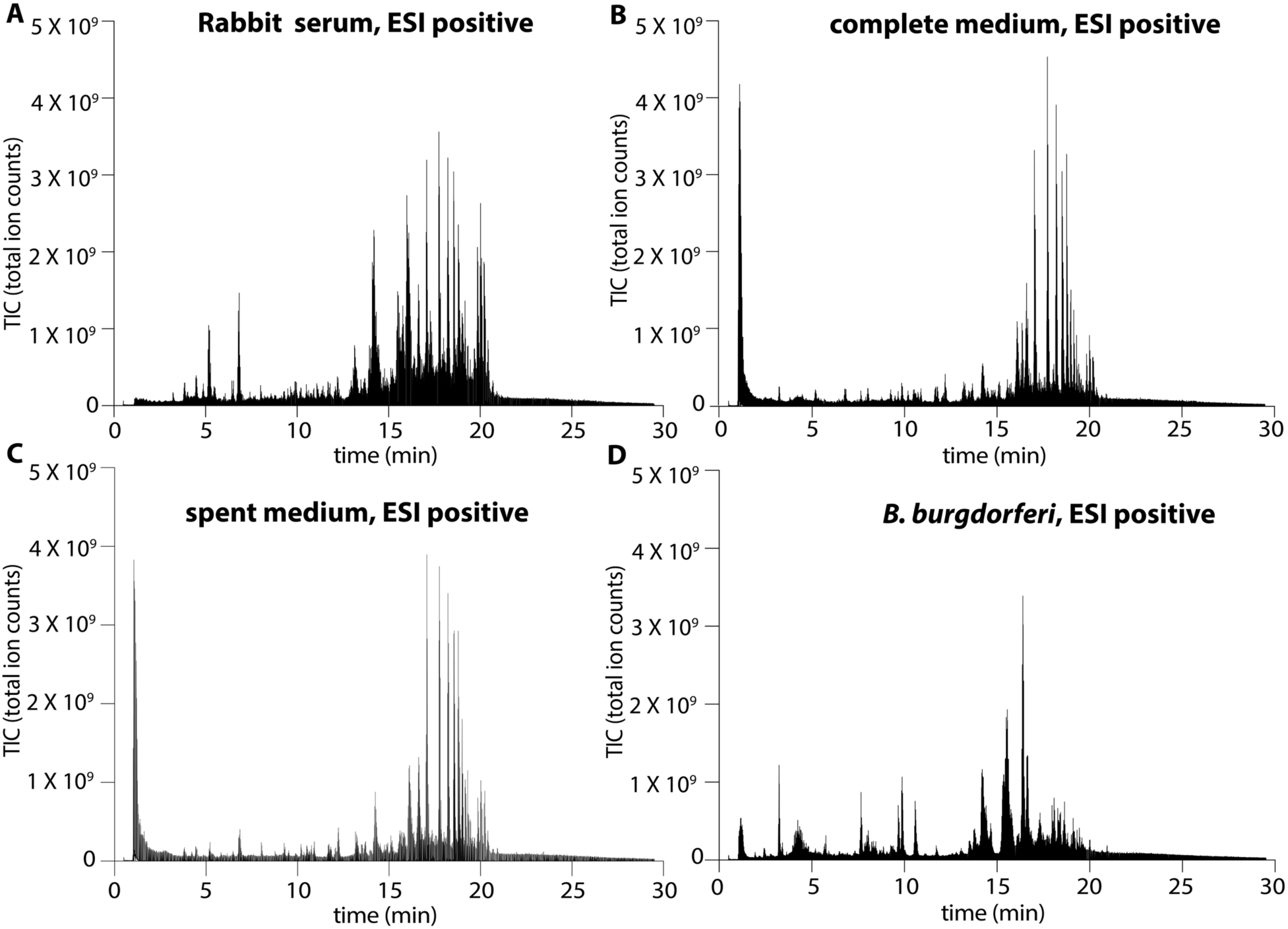
Untargeted lipidomic profiling of rabbit serum, growth media, and *B. burgdorferi*. Representative total ion chromatograms acquired in positive electrospray ionization (ESI positive) mode are shown for **(A)** rabbit serum, **(B)** complete medium, **(C)** spent medium, and **(D)** *B. burgdorferi*.

**Fig. S3.**
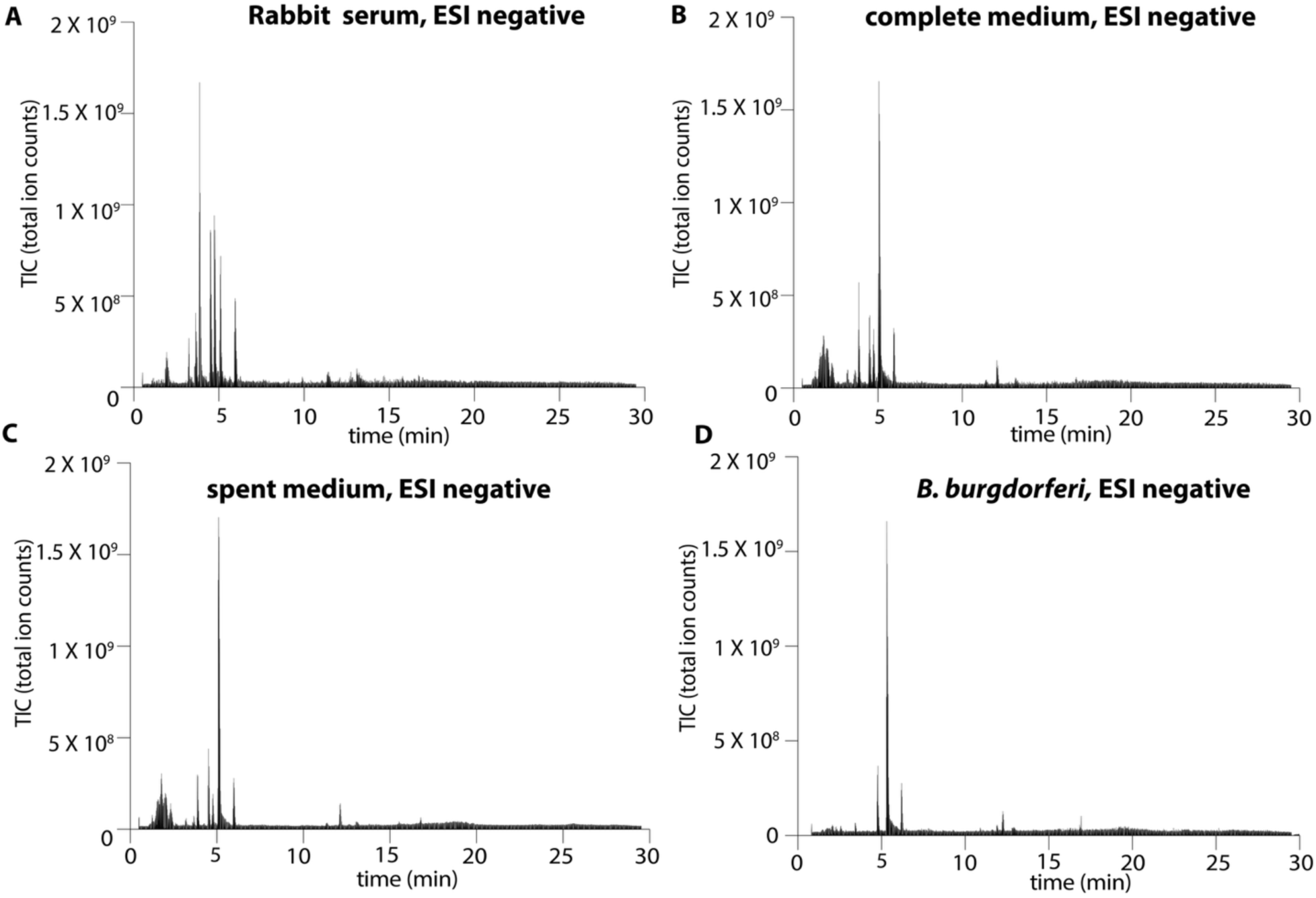
Untargeted lipidomic profiling of rabbit serum, growth media, and *B. burgdorferi*. Representative total ion chromatograms acquired in negative electrospray ionization (ESI negative) mode are shown for **(A)** rabbit serum, **(B)** complete medium, **(C)** spent medium, and **(D)** *B. burgdorferi*.

**Fig. S4.**
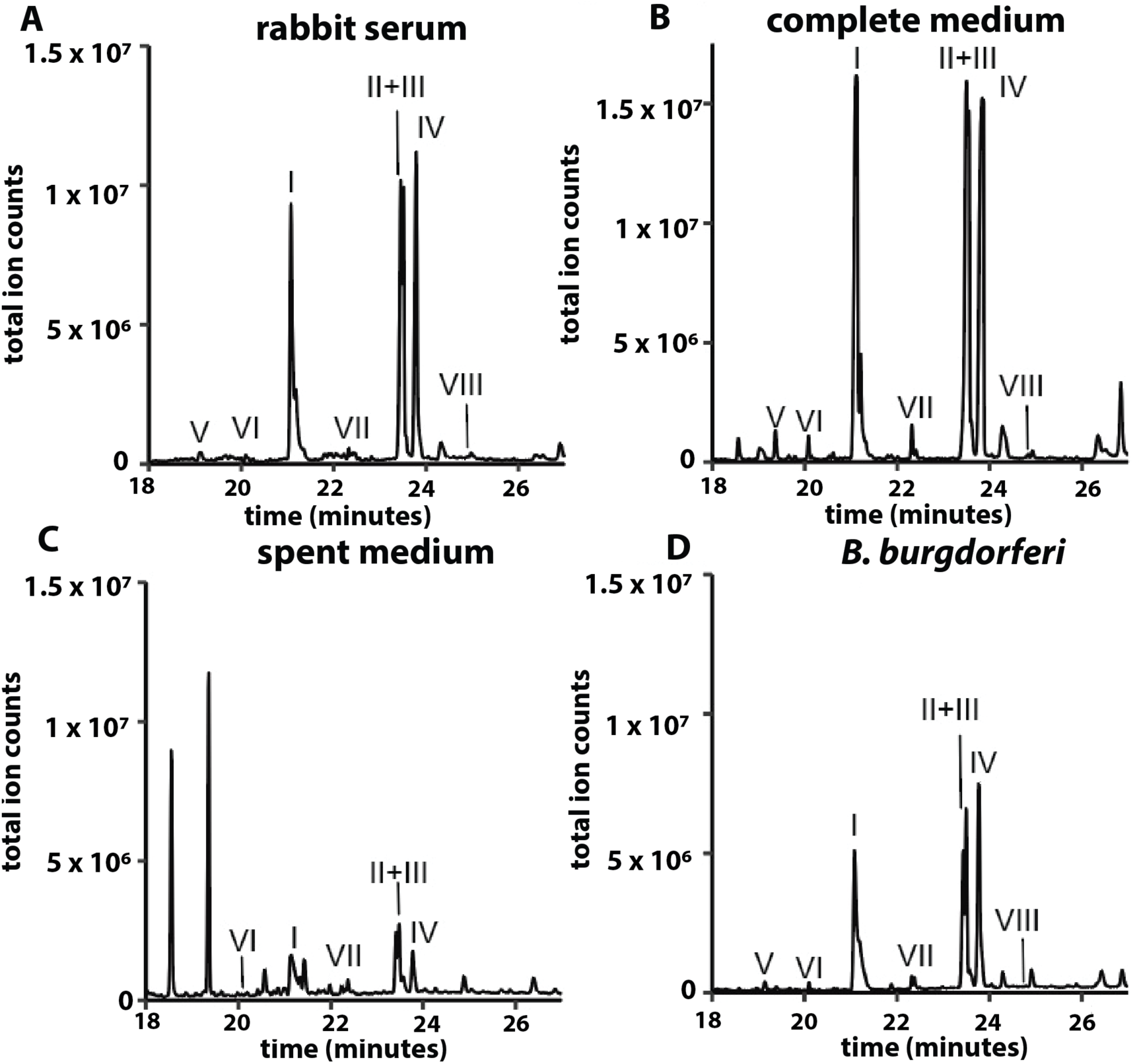
Fatty acid profiling of rabbit serum, complete medium, spent medium, and *Borrelia burgdorferi*. Samples were analyzed via GC. Peaks were annotated on the basis of retention time and corresponding lipid standards. The identified fatty acids were: I: Palmitic acid, II: Stearic acid, III: Oleic acid, IV: Linoleic acid, V: Myristic acid, VI: Pentadecanoic acid, VII: Heptadecanoic acid, VIII: Arachidonic acid.

**Fig. S5.**
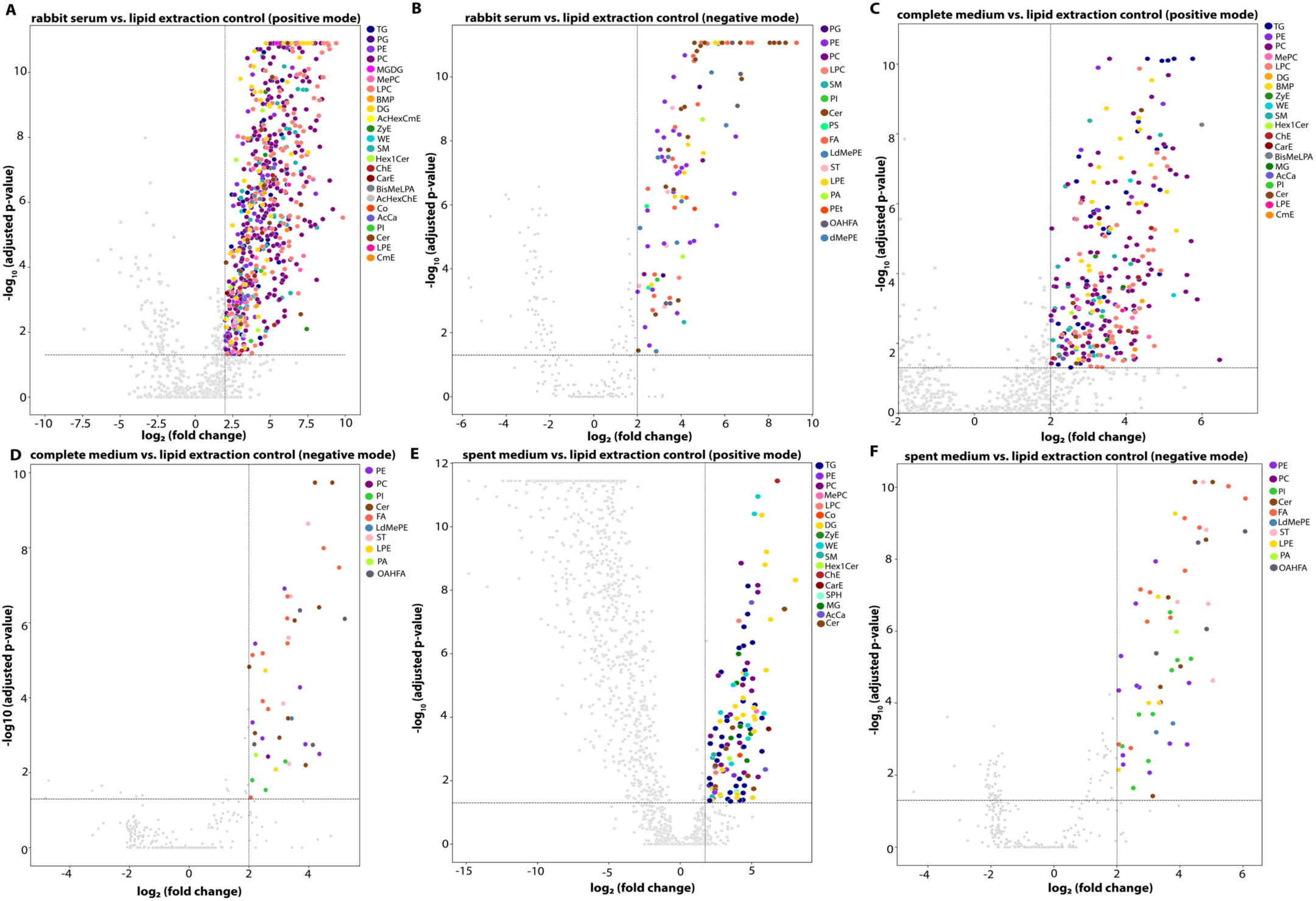
Untargeted lipidomics of rabbit serum and media to assess lipid abundance and distribution. Volcano plots show lipids significantly enriched in rabbit serum **(A, B)**, complete medium **(C, D)** and spent medium **(E, F)** compared to a lipid extraction control.

**Fig. S6.**
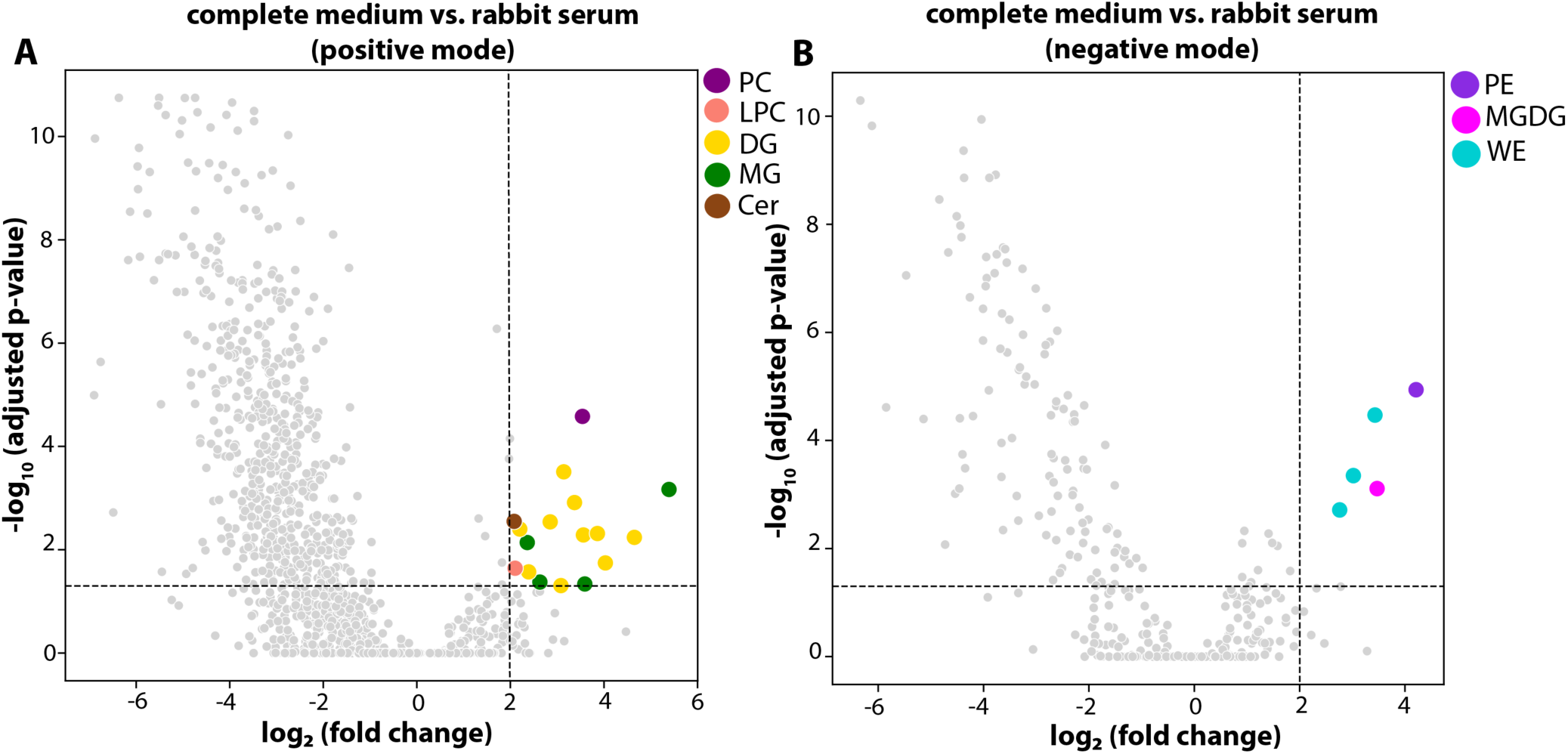
Untargeted comparative lipidomics of rabbit serum and the complete medium. Volcano plots compare lipid profiles between rabbit serum and complete medium in **(A)** positive and **(B)** negative electrospray ionization modes. Most lipids in complete medium originate from rabbit serum, with limited medium-specific lipid enrichment (highlighted in colors). The plots demonstrate significant overlap in lipid composition between rabbit serum and complete medium.

**Fig. S7.**
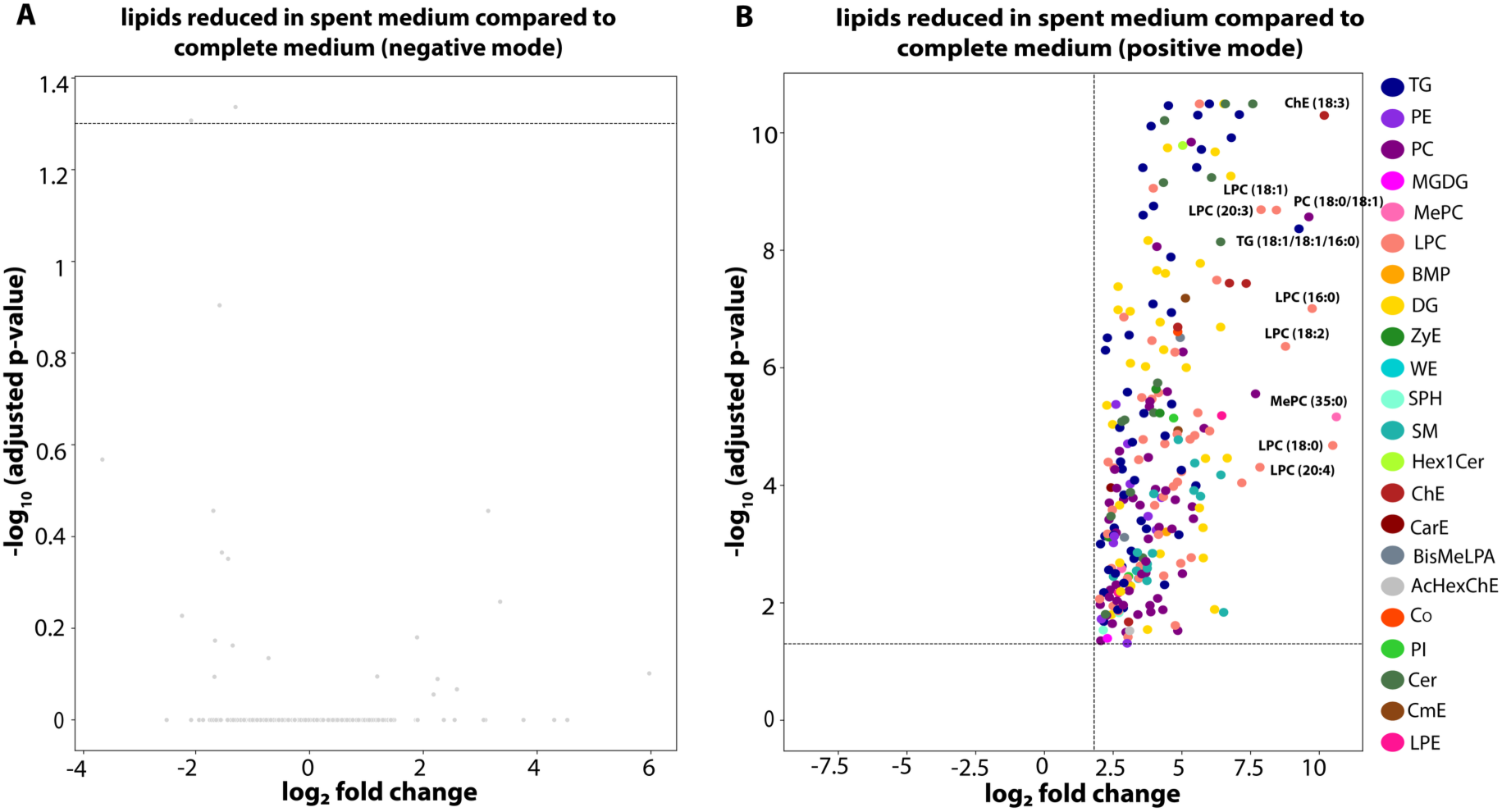
Lipid depletion in the spent medium. Volcano plots showing lipids enriched in the complete medium that are absent in the spent medium. No substantial differential enrichment was observed for lipids that ionize in negative ionization mode (**A**) but a substantial number of lipids that ionize in positive ionization mode was detected (**B**); the ten most significantly reduced lipids are labeled.

**Fig. S8.**
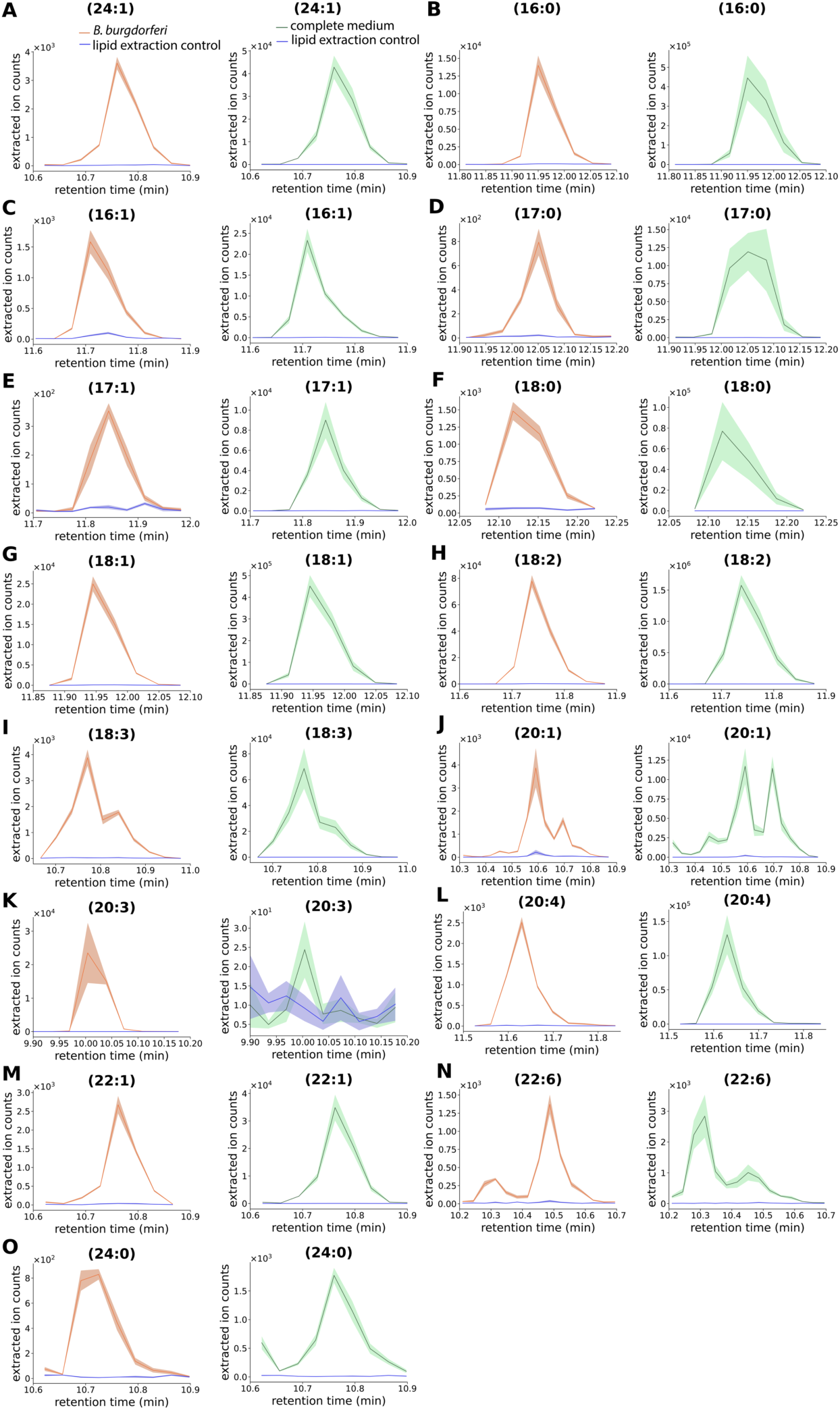
Cholesteryl esters in *B. burgdorferi* and the complete medium. Extracted ion chromatograms (EICs) of diagnostic product ions from targeted MS/MS analysis of selected CE species are shown. Orange traces represent *B. burgdorferi* samples, green traces represent BSKII complete medium, and blue traces represent lipid extraction controls. The solid line indicates the mean of four biological replicates, and the shaded area denotes the standard error of the mean (SEM).

**Fig. S9.**
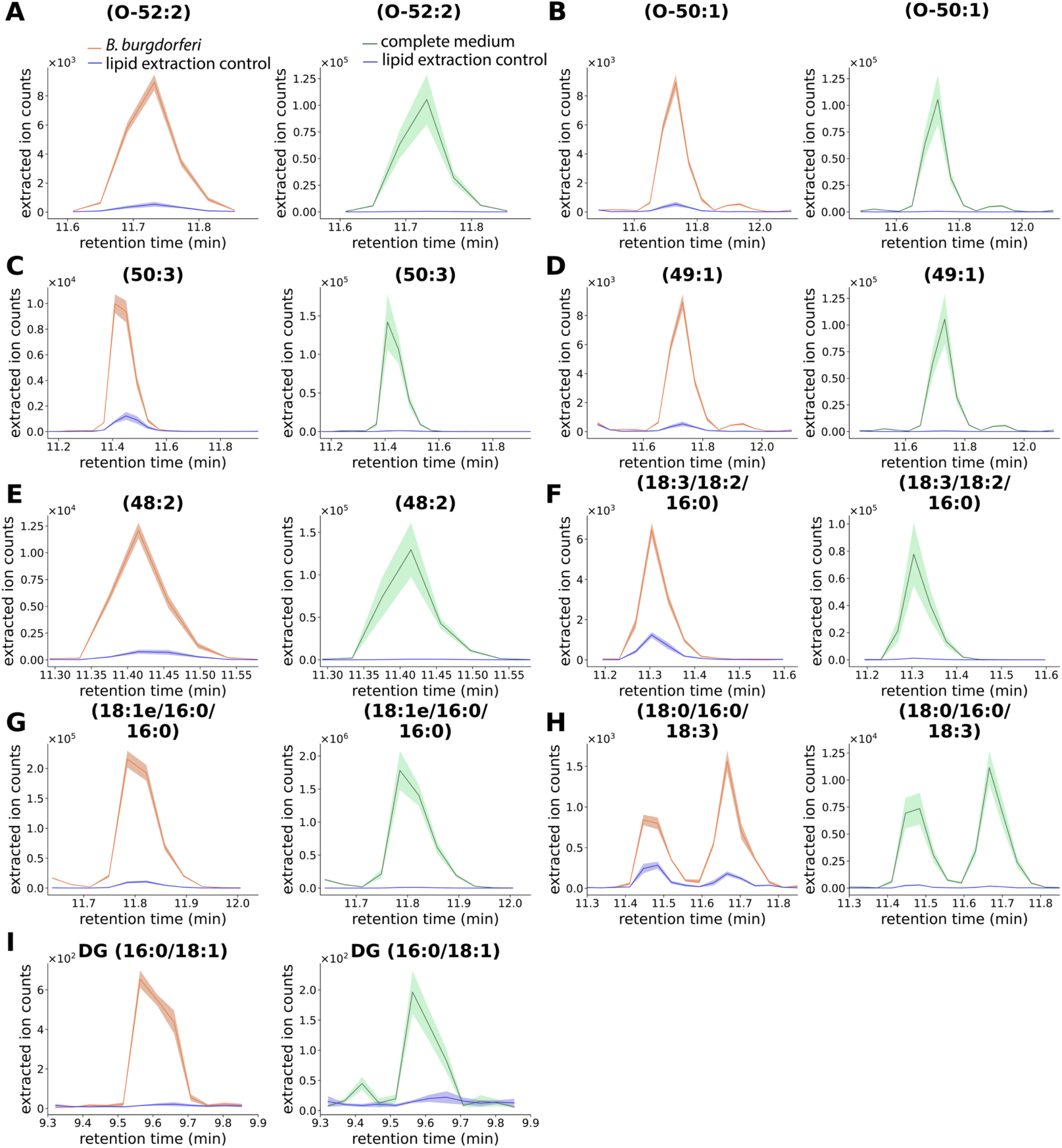
Neutral lipids in *B. burgdorferi* and the complete medium. EICs of diagnostic product ions from targeted MS/MS analysis of the indicated neutral lipid species are shown. Orange traces represent *B. burgdorferi* samples, green traces represent BSKII complete medium, and blue traces represent lipid extraction controls. The solid line indicates the mean of four biological replicates, and the shaded area denotes the SEM.

**Fig. S10.**
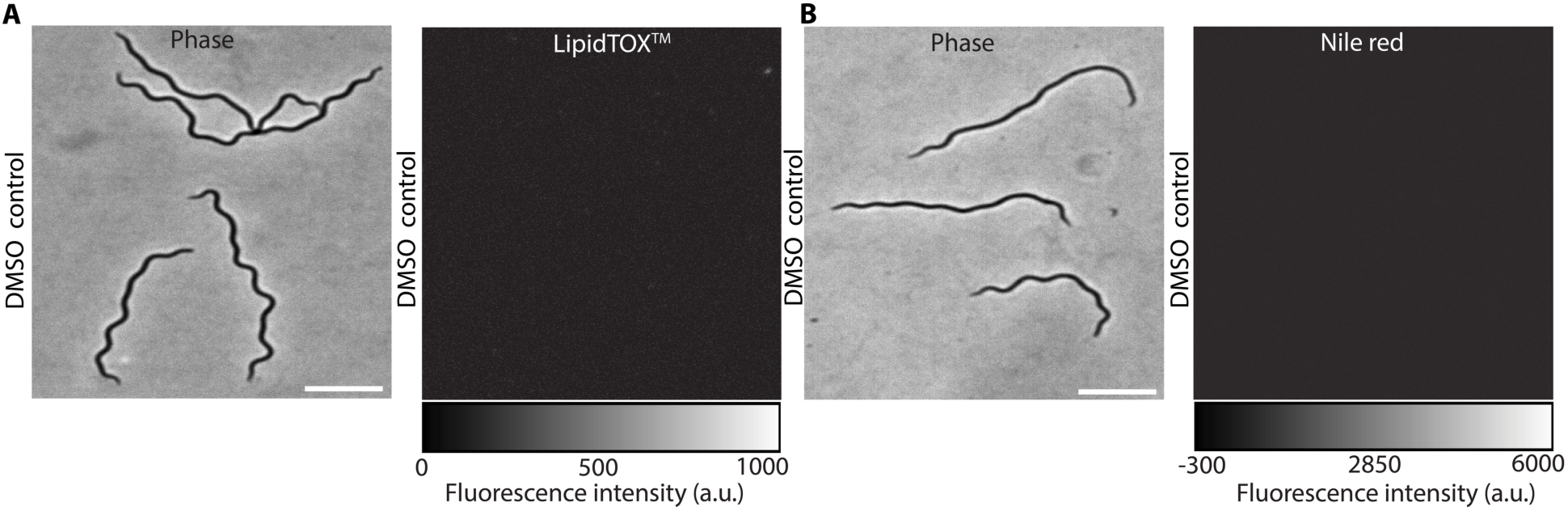
**Microscopy of *B. burgdorferi***. Widefield epifluorescence microscopy images of fixed *B. burgdorferi* cells treated with DMSO control at a 1:200 ratio in phosphate-buffered saline with MgCl_2_, observed under the phase contrast and the LipidTOX™ fluorescence channel (**A**) and the phase contrast and Nile Red fluorescence channel. Scale bar = 5 µm.

**Fig. S11.**
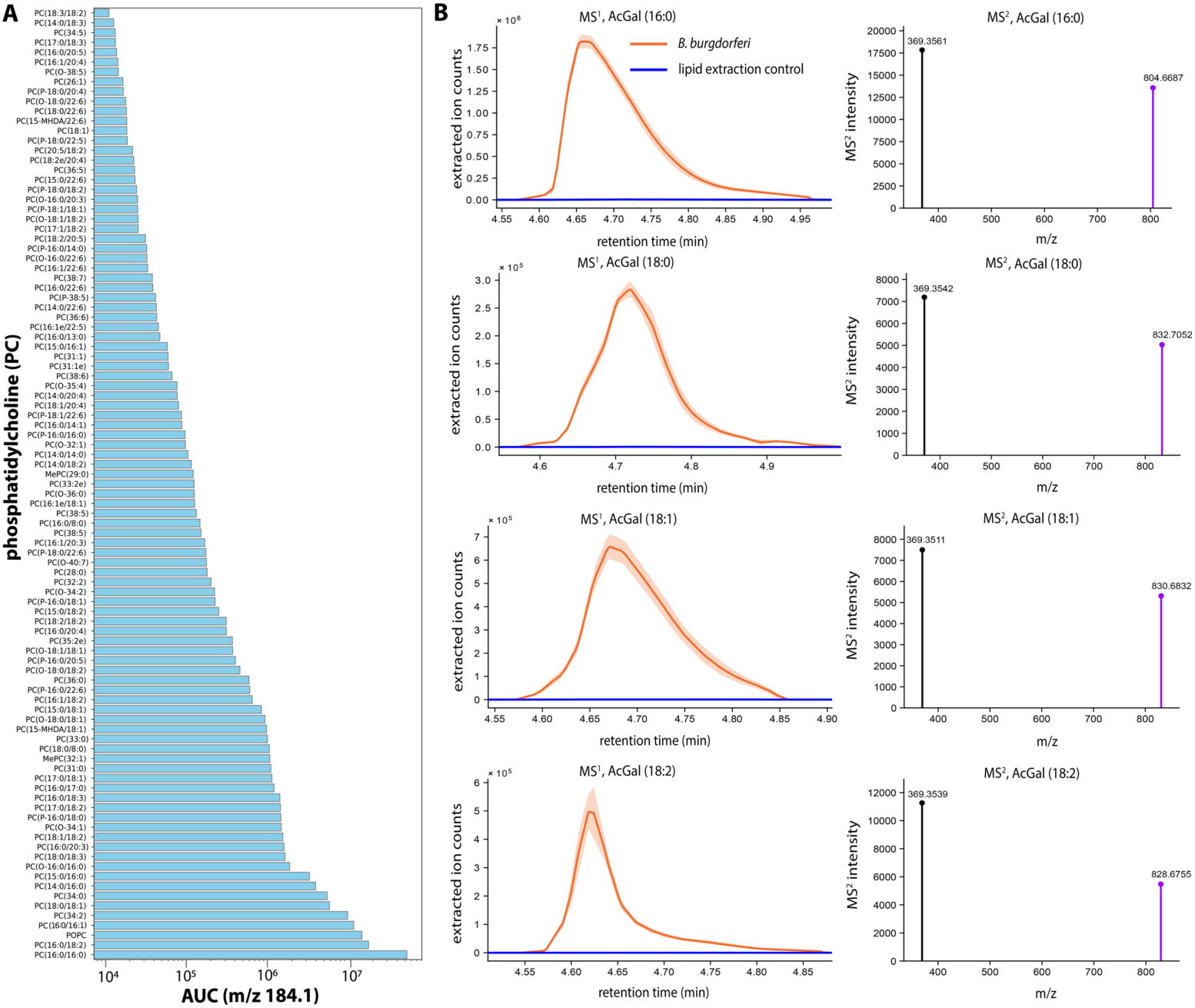
Targeted profiling of phosphatidylcholines (PCs) and acylated cholesterol galactolipids (AcGals) in *B. burgdorferi*. (**A**) Ranked abundances of the area under the curve (AUC) of 97 PC species identified in *B. burgdorferi*. Abundances were calculated from the extracted ion chromatograms of the diagnostic PC product ion at m/z 184.1. (**B**) Left panel: EICs show the signal for each AcGal precursor ion in *B. burgdorferi* (orange) compared to a lipid extraction control (blue). Right panel: Corresponding mass spectra showing the intact ammonium-adducted precursor ion ([M+NH₄]⁺, purple) and characteristic fragment ions.

**Fig. S12.**
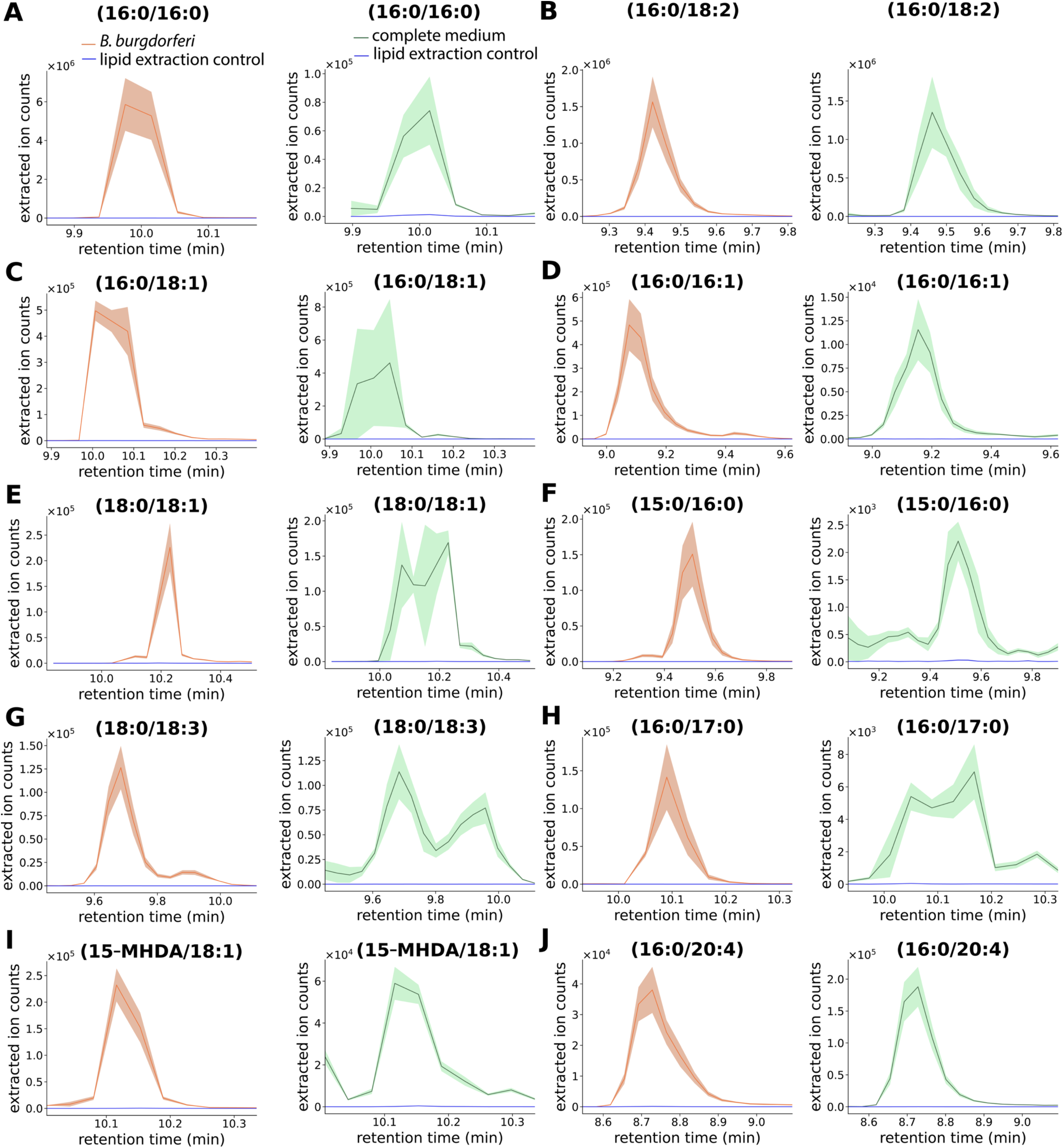
Phosphatidylcholines in *B. burgdorferi* and the complete medium. EICs of diagnostic product ions from targeted MS/MS analysis of the ten selected PC species in *B. burgdorferi*. Orange traces represent *B. burgdorferi* samples, green traces represent BSKII complete medium, and blue traces represent lipid extraction controls. The solid line indicates the mean of four biological replicates, and the shaded area denotes the SEM.

**Fig. S13.**
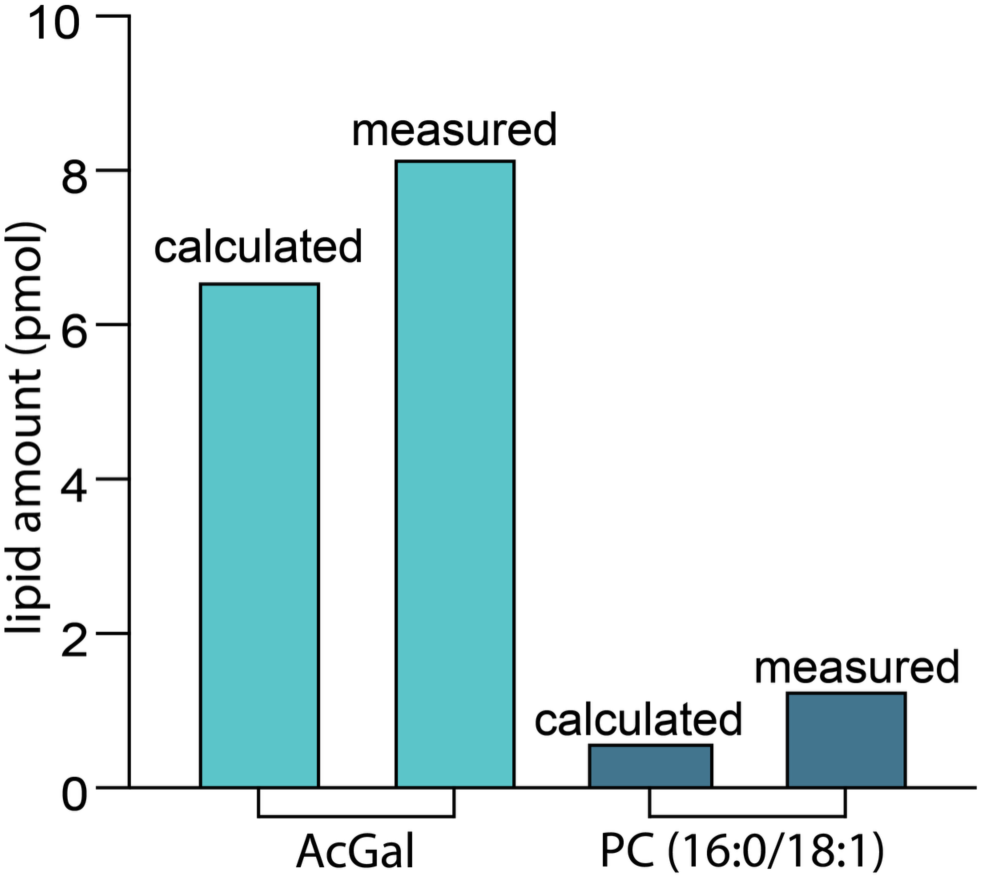
Validation of the response factor calculation. The amounts of AcGal (16:0) and PC (16:0/18:1) calculated for data acquired via the QQQ (calculated) are similar to the amounts measured using a multipoint calibration curve-based performed on a Q-TOF instrument (measured).

**Fig. S14.**
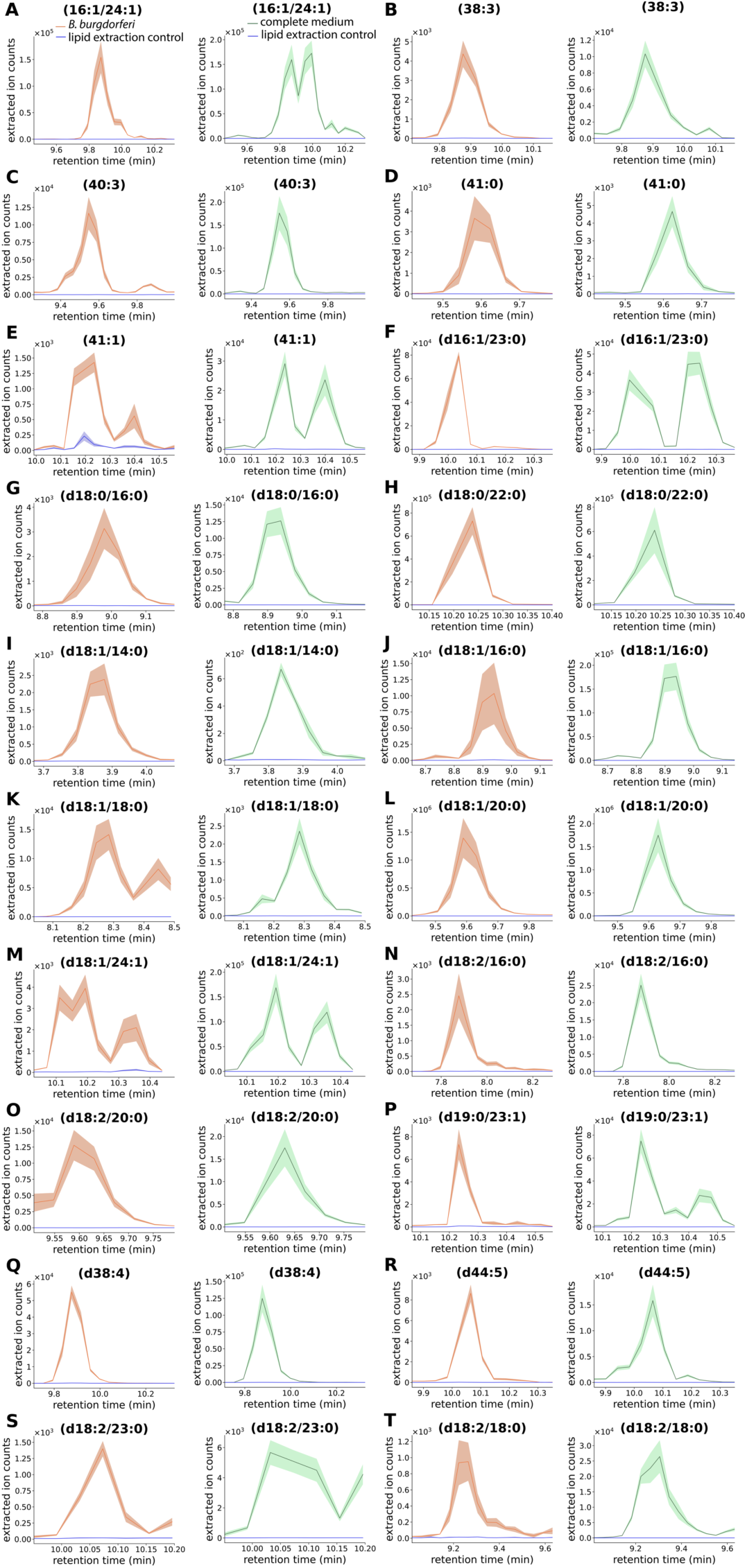
Sphingomyelins in *B. burgdorferi* and the complete medium. EICs of diagnostic product ions from targeted MS/MS analysis of sphingomyelin species are shown. Orange traces represent *B. burgdorferi* samples, green traces represent BSKII complete medium, and blue traces represent lipid extraction controls. The solid line indicates the mean of four biological replicates, and the shaded area denotes the SEM.

**Fig. S15.**
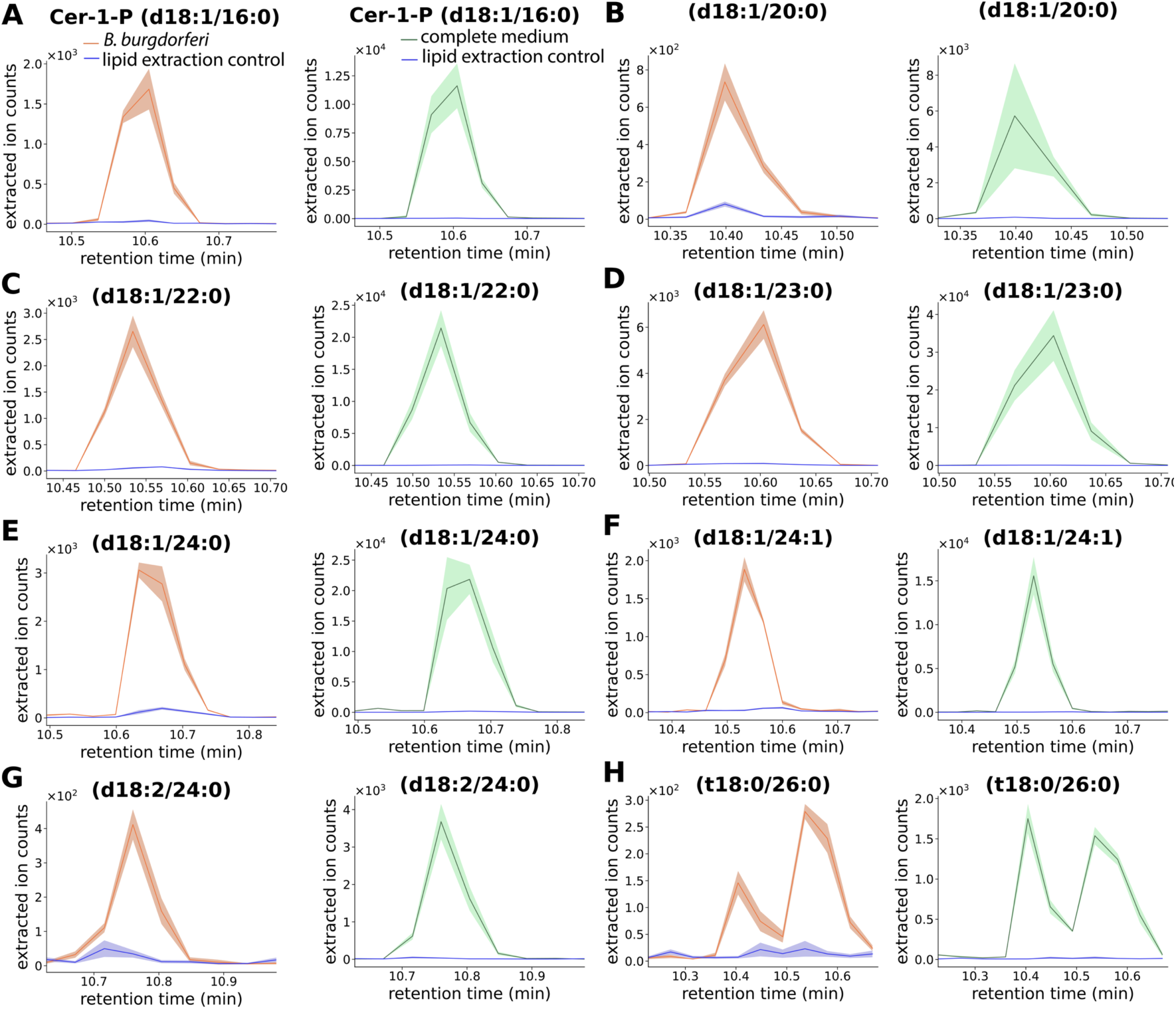
Analysis of ceramides in *B. burgdorferi* and the complete medium. EICs of diagnostic product ions from targeted MS/MS analysis of ceramide species are shown. Panels correspond to individual ceramide species. Orange traces represent *B. burgdorferi* samples, green traces represent BSKII complete medium, and blue traces represent lipid extraction controls. The solid line indicates the mean of four biological replicates, and the shaded area denotes the SEM.

**Fig. S16.**
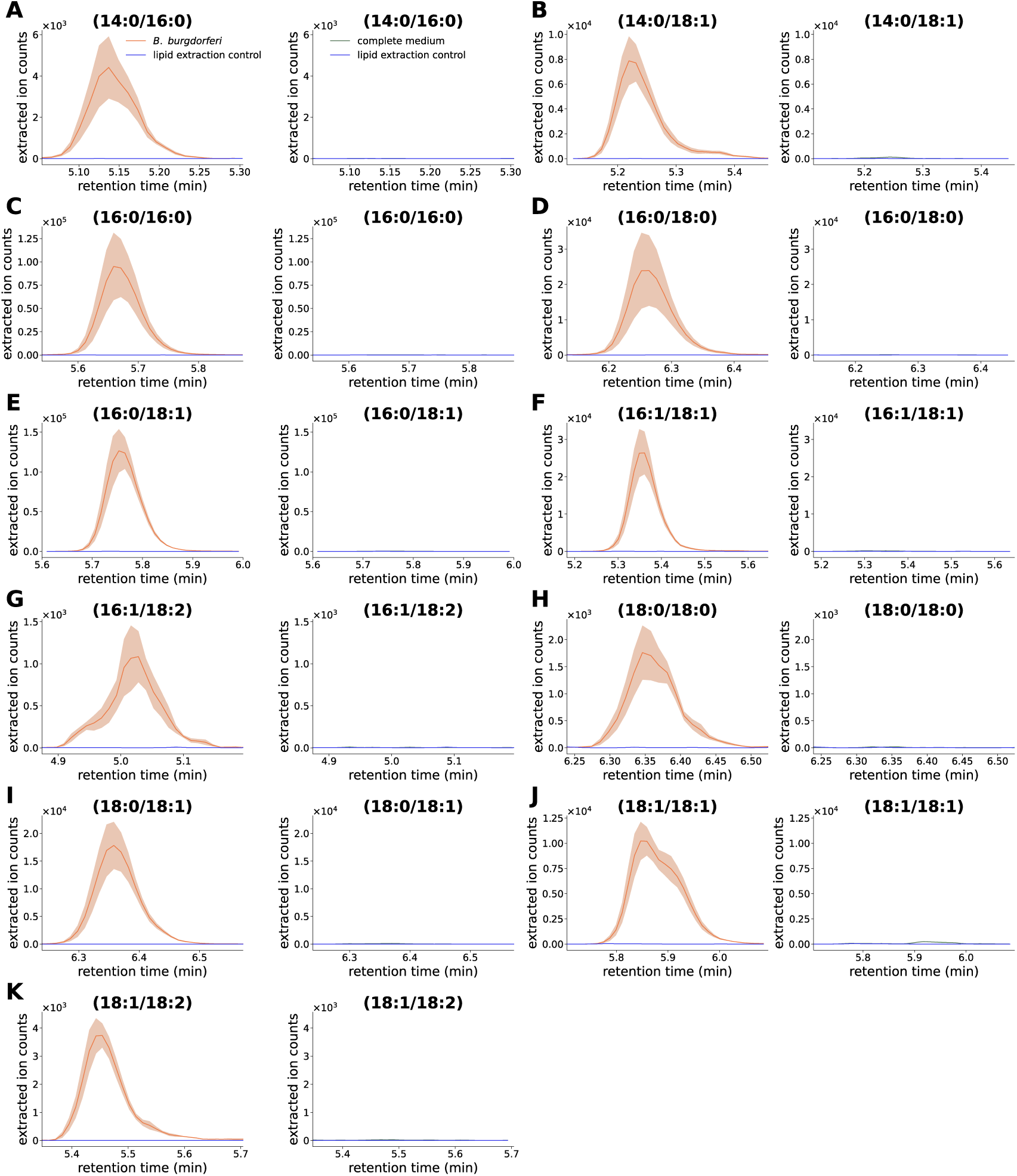
Analysis of phosphatidylglycerols in *B. burgdorferi* and the complete medium. EICs of diagnostic product ions from targeted MS/MS analysis of phosphatidylglycerol species are shown. Orange traces represent *B. burgdorferi* samples, green traces represent BSKII complete medium, and blue traces represent lipid extraction controls. The solid line indicates the mean of four biological replicates, and the shaded area denotes the SEM.

**Fig. S17.**
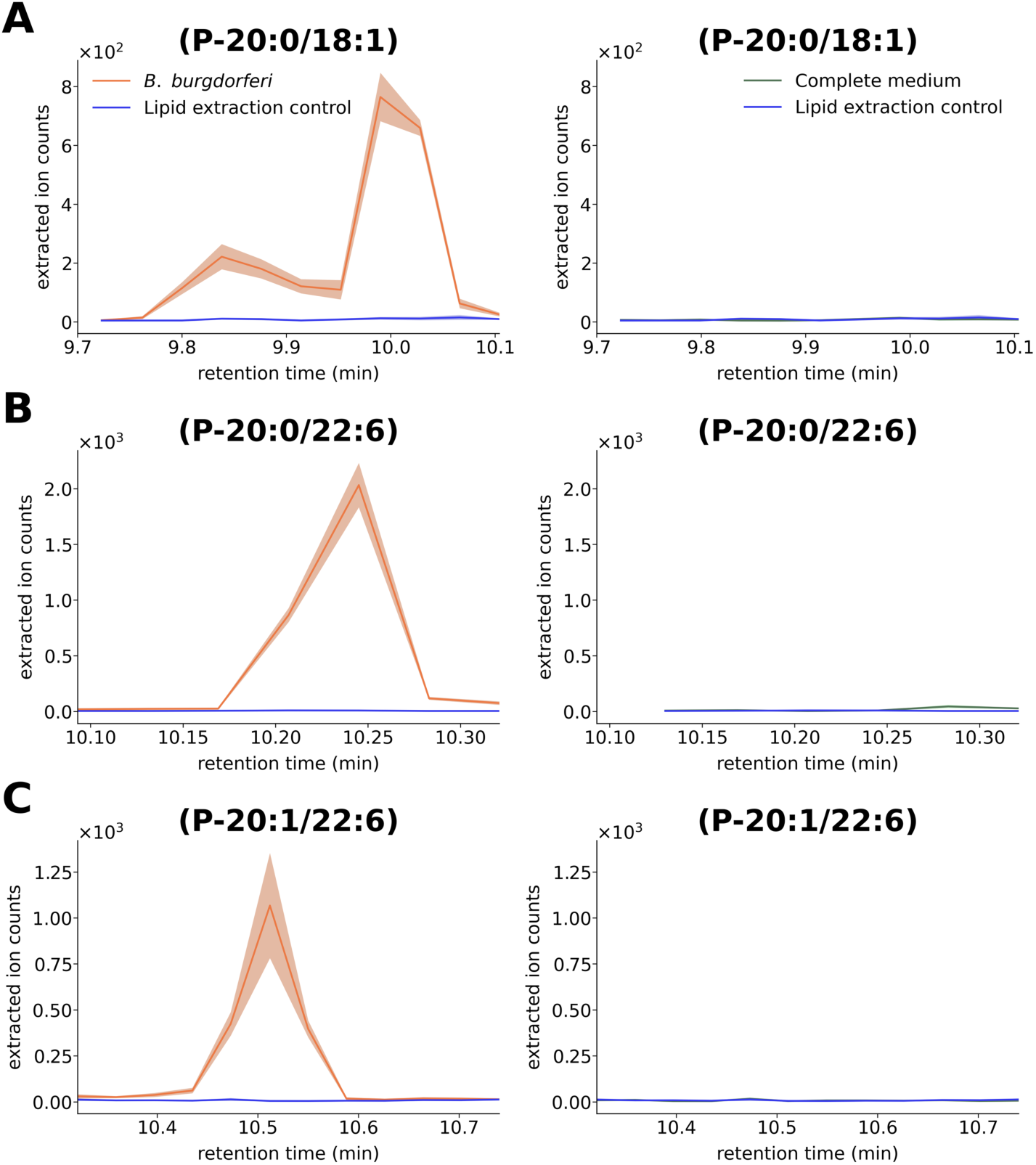
Analysis of phosphatidylethanolamines in *B. burgdorferi* and the complete medium. EICs of diagnostic product ions from targeted MS/MS analysis of phosphatidylethanolamine species are shown. Orange traces represent *B. burgdorferi* samples, green traces represent BSKII complete medium, and blue traces represent lipid extraction controls. The solid line indicates the mean of four biological replicates, and the shaded area denotes the SEM.

**Fig. S18.**
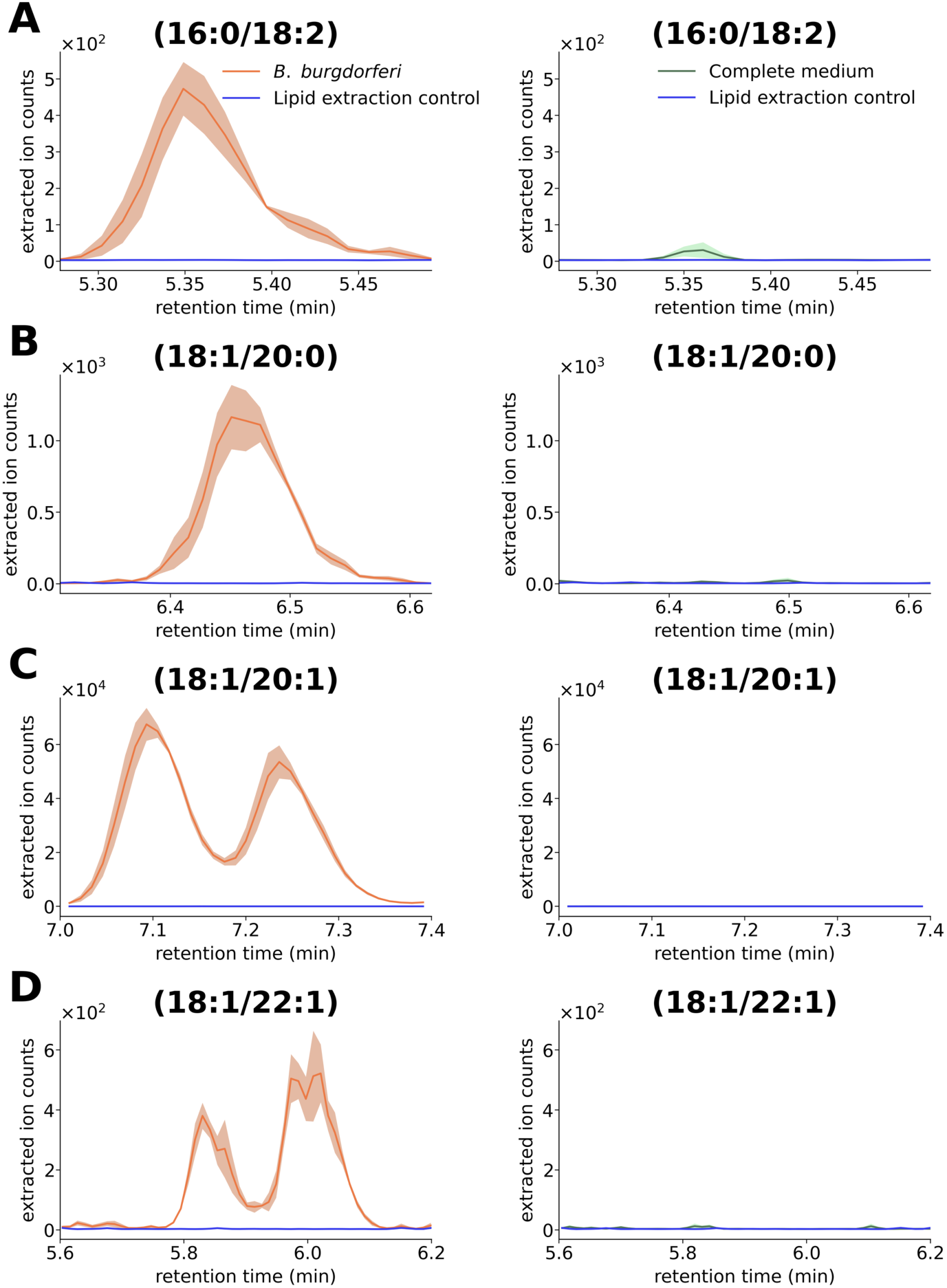
Analysis of bis(monoacylglycero)phosphates (BMPs) in *B. burgdorferi* and the complete medium. EICs of diagnostic product ions from targeted MS/MS analysis of BMPs species are shown. Orange traces represent *B. burgdorferi* samples, green traces represent BSKII complete medium, and blue traces represent lipid extraction controls. The solid line indicates the mean of four biological replicates, and the shaded area denotes the SEM.

**Fig. S19.**
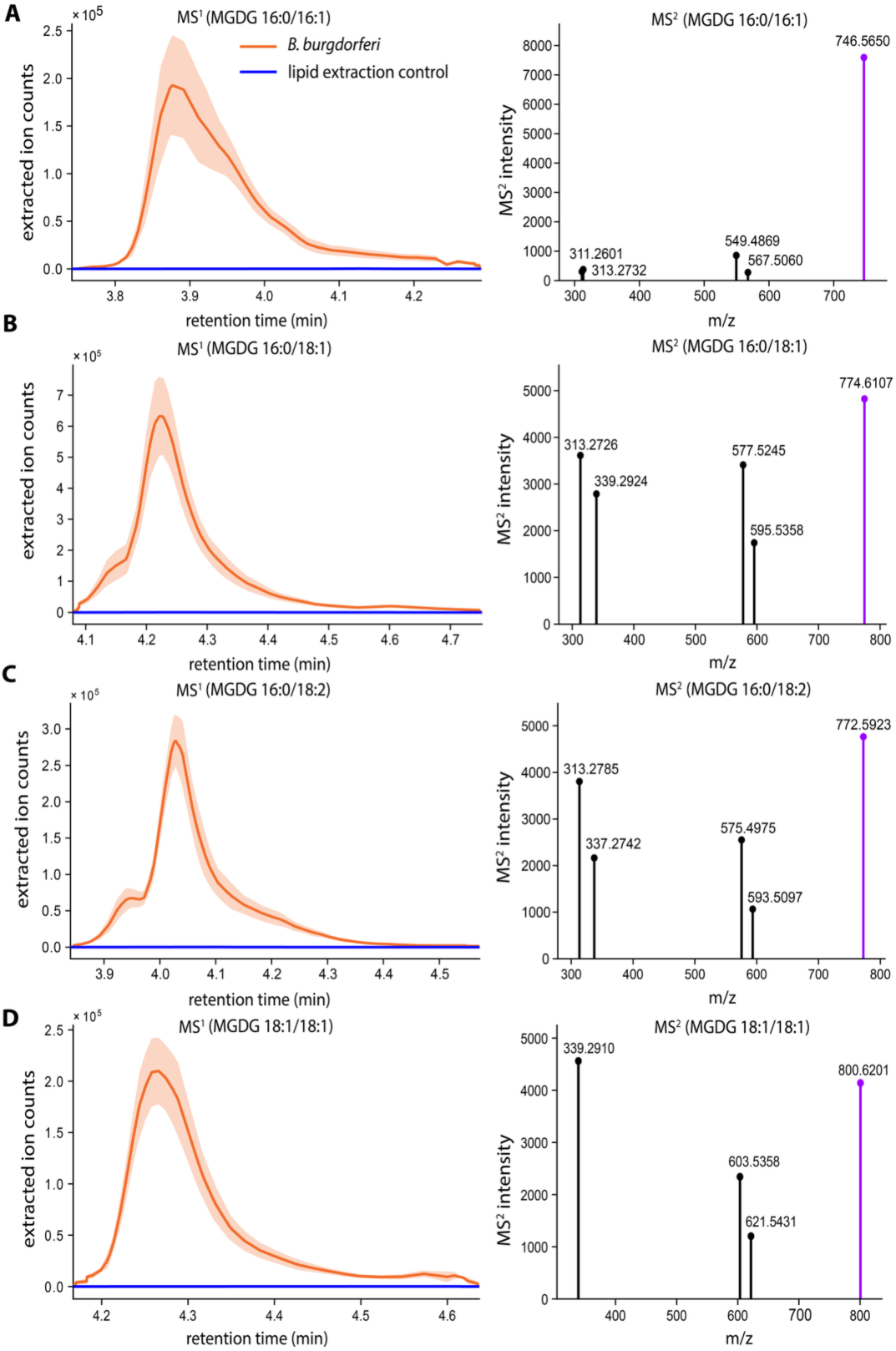
Representative mass spectrometry data for four distinct monogalactosyldiacylglycerol (MGDG) species in *B. burgdorferi*. MGDGs were characterized by their different fatty acid compositions (A: 16:0/16:1, B: 16:0/18:1, C: 16:0/18:2, and D: 18:1/18:1). Left panel: EICs show the signal for each MGDG precursor ion in *B. burgdorferi* (orange) compared to a lipid extraction control (blue). Right panel: Corresponding mass spectra showing the intact ammonium-adducted precursor ion ([M+NH₄]⁺, purple) and characteristic fragment ions.

**Fig. S20.**
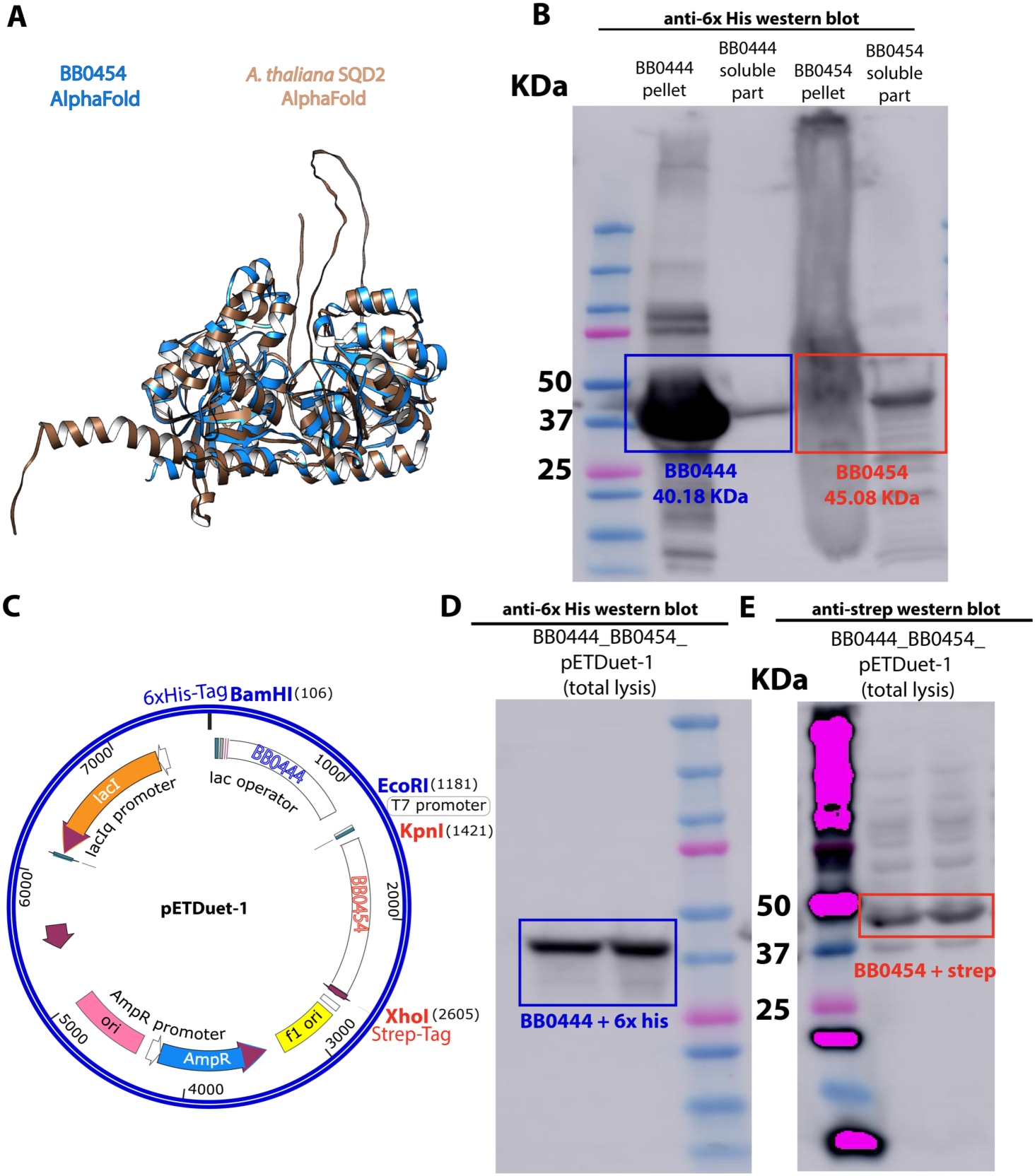
SQDG synthesis in *B. burgdorferi*. (**A**) An alignment of AlphaFold2 models of BB0454 and the SQD2 protein from *A. thaliana*. (**B**) Heterologously expressed BB0444 and BB0454 cloned separately in pET28a vectors and bearing hexa-histidine affinity tags. (**C**) Plasmid map of pETDuet vector containing BB0444 and BB0454. (**D, E**) Heterologously expressed of BB0444 and BB0454 in verified through immunoblotting using affinity tags appended to each protein in pETDuet vector.

**Fig. S21.**
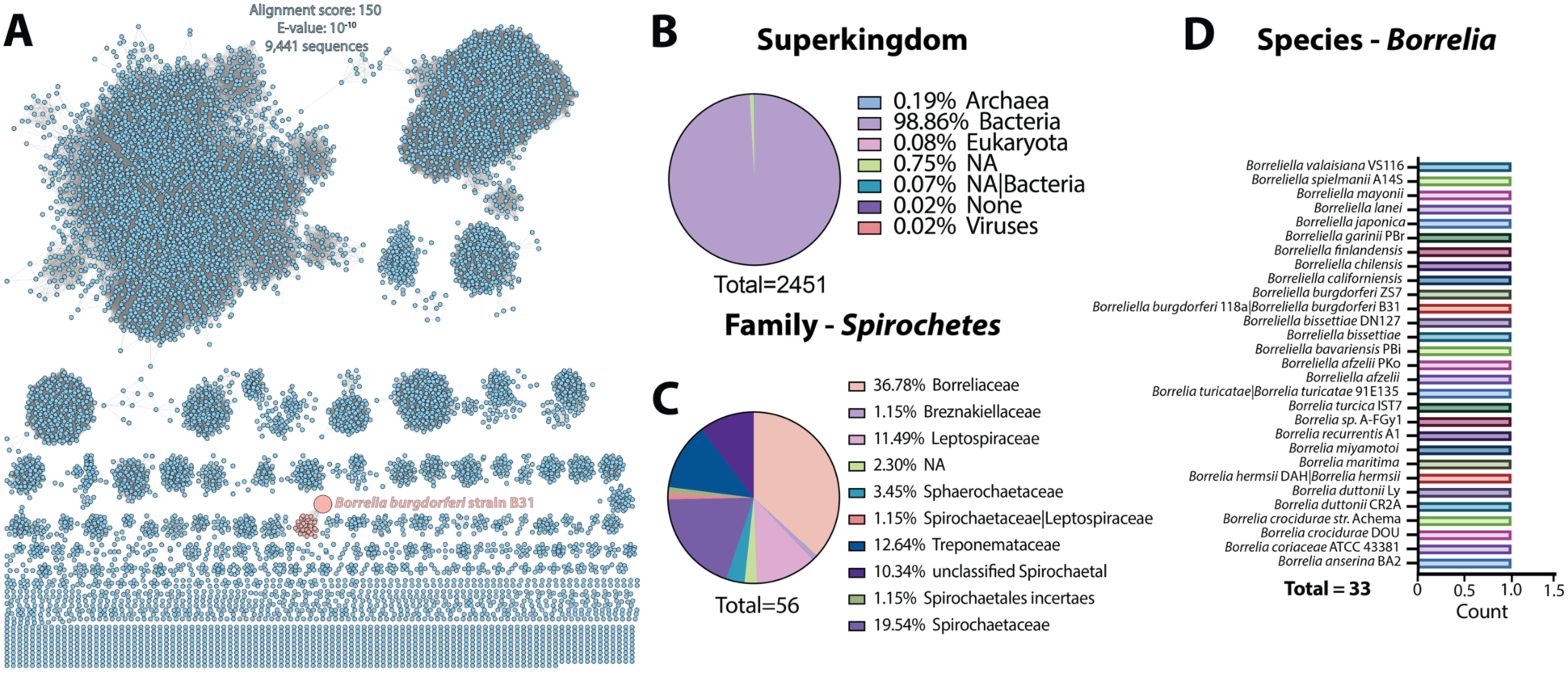
Bioinformatics analyses of BB0444. (**A**) Protein sequence similarity network (SSN) the BB0444 protein from *B. burgdorferi*, showing separate clustering from homologs with a minimum of 90% identity. (**B**) Distribution of BB0444 homologs suggest they are widespread in bacteria, including in spirochetes (**C**) where a total of 33 Borrelia species encode a BB0444 homolog (**D**).

**Fig. S22.**
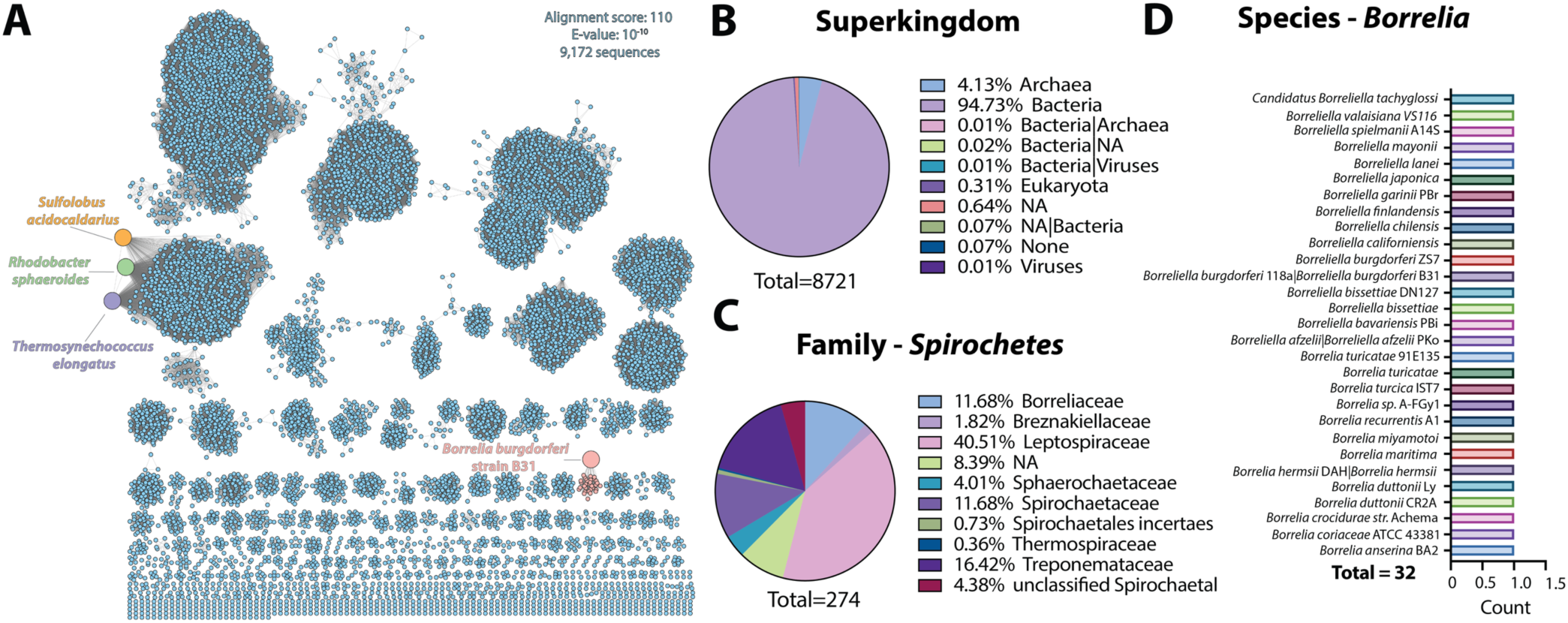
Bioinformatics analyses of BB0454. (**A**) Protein sequence similarity network (SSN) the BB0454 protein from *B. burgdorferi*, showing separate clustering from homologs with a minimum of 90% identity. (**B**) Distribution of BB0454 homologs suggest they are widespread in bacteria, including in spirochetes (**C**). A total of 32 Borrelia species encode a BB0454 homolog (**D**).

### Supporting table

**Table S1:**
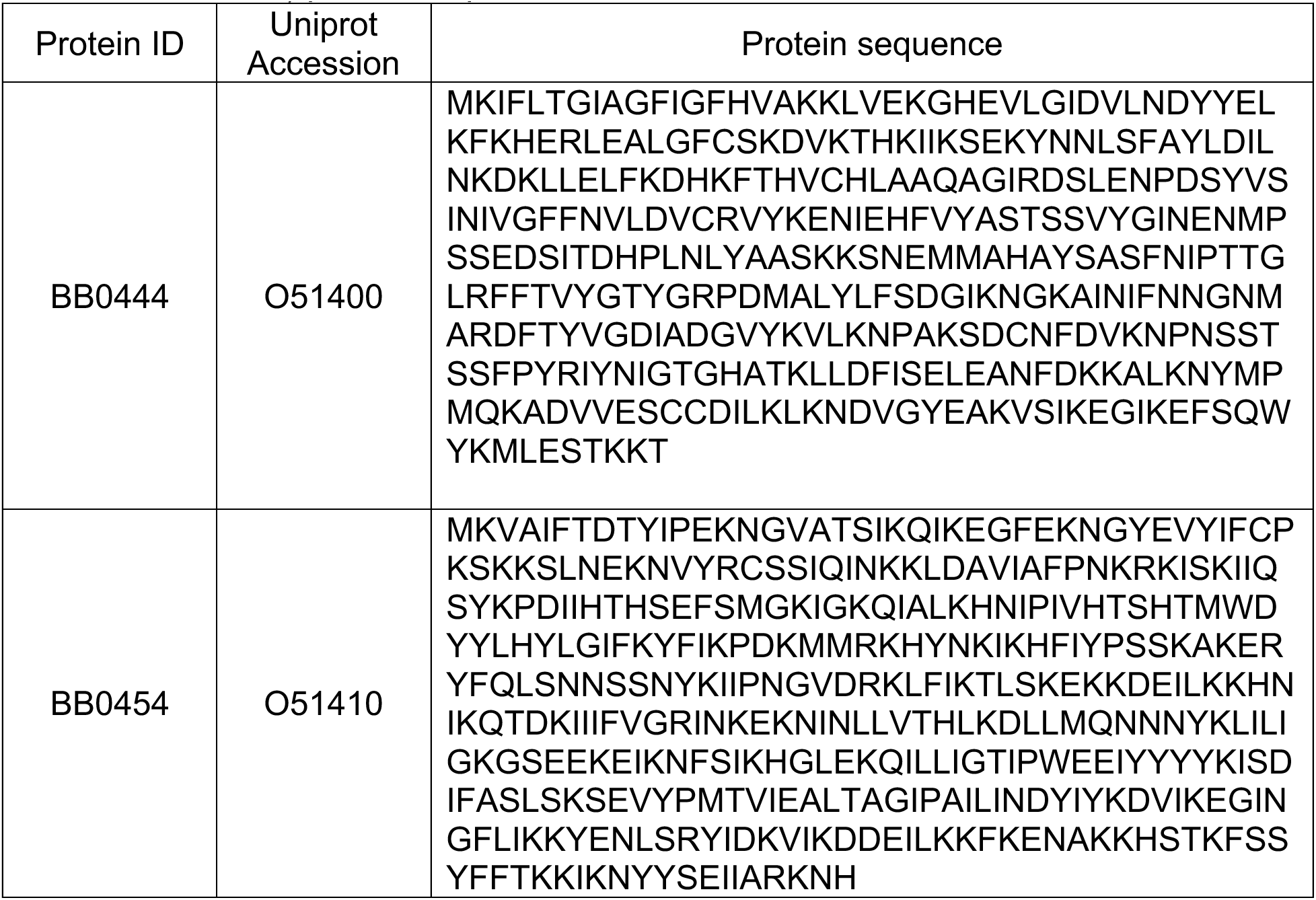
Amino acid sequences of *Borrelia burgdorferi* (strain ATCC 35210 / DSM 4680 / CIP 102532 / B31) protein sequences.

