## Supplementary file 1 for "A lipid compendium of a metabolically compromised bacterium provides insights into lipid acquisition, biosynthesis, and metabolism"

### Supplementary Spreadsheet 1: Comparative analysis of lipid species significantly re[illegible]

| Name | lipid_class | Group Area: CM_pos_12022024_Newanalysis |
|---|---|---|
| AcHexChE(15:0) | AcHexChE | 4754849.891 |
| BisMeLPA(12:0e) | BisMeLPA | 16693909.26 |
| BisMeLPA(10:0e) | BisMeLPA | 2145335.997 |
| BMP(18:0_18:1) | BMP | 10292743.78 |
| CarE(16:1) | CarE | 23250441.29 |
| Cer(d18:1_23:0) | Cer | 66567694.82 |
| Cer(t17:0_26:0+O) | Cer | 28393969.27 |
| Cer(d18:1_24:1) | Cer | 45209902.01 |
| Cer(d18:1_22:0) | Cer | 43985993.19 |
| Cer(t20:0_25:0+O) | Cer | 6251081.394 |
| Cer(t42:0) | Cer | 4456662.961 |
| Cer(d18:1_24:2) | Cer | 8002096.473 |
| Cer(d18:1_20:0) | Cer | 10760299.53 |
| Cer(d18:1_16:0) | Cer | 8903255.662 |
| Cer(m36:0) | Cer | 4235536.527 |
| Cer(d18:1_24:0) | Cer | 2313140.643 |
| Cer(d19:2_31:0+O) | Cer | 1670027.682 |
| Cer(m18:1_22:0) | Cer | 1711355.254 |
| ChE(18:3) | ChE | 592096919.7 |
| ChE(20:5) | ChE | 88554223.05 |
| ChE(18:2) | ChE | 66016579.57 |
| ChE(20:4) | ChE | 15822942.41 |
| CmE(18:3) | CmE | 16788578.1 |
| CmE(18:2) | CmE | 10284411.74 |
| Co(Q10) | Co | 8799182.022 |
| DG(18:1_16:0) | DG | 144194289.9 |
| DG(32:2e) | DG | 94099416.24 |
| DG(34:3e) | DG | 72666614.08 |
| DG(34:4e) | DG | 68938875.66 |

| | | |
|---|---|---|
| DG(18:2_16:0) | DG | 83035866.2 |
| DG(36:2e) | DG | 923308186 |
| DG(18:0_16:0) | DG | 24530659.76 |
| DG(34:2e) | DG | 484143510.9 |
| DG(36:4e) | DG | 35940976.92 |
| DG(36:5) | DG | 83035866.2 |
| DG(36:5e) | DG | 33078450.18 |
| DG(38:2e) | DG | 18342256.25 |
| DG(16:0_18:3) | DG | 12357206.54 |
| DG(35:2e) | DG | 9081236.961 |
| DG(36:6) | DG | 2395343.124 |
| DG(18:1_18:1) | DG | 10779000.61 |
| DG(36:3e) | DG | 10928828.82 |
| DG(36:6e) | DG | 9435531.407 |
| DG(18:4_18:1) | DG | 5990051.713 |
| DG(18:1_18:2) | DG | 154204355.3 |
| DG(18:1_18:3) | DG | 8323241.929 |
| DG(16:0_16:1) | DG | 4775491.649 |
| DG(18:0_18:1) | DG | 38454971.96 |
| DG(28:5) | DG | 3792649.136 |
| Hex1Cer(d18:1_16:0) | Hex1Cer | 4709144.279 |
| LPC(18:0) | LPC | 179369466.1 |
| LPC(16:0) | LPC | 205449825.3 |
| LPC(18:2) | LPC | 72490370.65 |
| LPC(18:1) | LPC | 53022168.68 |
| LPC(20:3) | LPC | 20114721.3 |
| LPC(20:4) | LPC | 50971693.25 |
| LPC(18:1e) | LPC | 20498145.2 |
| LPC(18:0e) | LPC | 9324409.353 |
| LPC(18:3) | LPC | 8065158.096 |

| | | |
|---|---|---|
| LPC(16:0e) | LPC | 25751206.26 |
| LPC(16:1) | LPC | 19733330.37 |
| LPC(15:0) | LPC | 15969773.3 |
| LPC(17:1) | LPC | 6550003.59 |
| LPC(19:0) | LPC | 30343263.09 |
| LPC(17:0) | LPC | 18094312.51 |
| LPC(18:2e) | LPC | 1955041.82 |
| LPC(14:0) | LPC | 3118045.517 |
| LPC(19:1) | LPC | 3899804.56 |
| LPC(22:5) | LPC | 4494355.48 |
| LPC(20:2) | LPC | 12715397.37 |
| LPC(20:0) | LPC | 3202887.295 |
| LPC(20:1) | LPC | 1186185.708 |
| LPC(22:4) | LPC | 7657502.836 |
| LPC(16:1e) | LPC | 15891664.54 |
| LPC(18:4) | LPC | 1827795.88 |
| LPC(18:1D7) | LPC | 1037178.289 |
| LPC(20:5) | LPC | 1656916.303 |
| LPC(31:0) | LPC | 2568692.659 |
| LPC(16:2e) | LPC | 672779.2774 |
| LPC(20:1e) | LPC | 1869035.57 |
| LPC(20:2e) | LPC | 348352.4492 |
| LPE(18:0) | LPE | 10736796.6 |
| MePC(35:0) | MePC | 154673422.3 |
| MePC(14:1e) | MePC | 1474300.644 |
| MGDG(19:1_19:1) | MGDG | 1903868.151 |
| PC(18:0_18:1) | PC | 1337465516 |
| PC(18:1_18:2) | PC | 395367558.9 |
| PC(19:1_19:1) | PC | 154021498.4 |
| PC(18:1e_20:4) | PC | 31778660.35 |

| | | |
|---|---|---|
| PC(18:0_16:0) | PC | 98592128.24 |
| PC(16:0_18:3) | PC | 7444233.464 |
| PC(18:0_22:4) | PC | 195387267.2 |
| PC(20:2_18:2) | PC | 60043432.42 |
| PC(18:1_18:1) | PC | 11741736033 |
| PC(35:3e) | PC | 7571577.017 |
| PC(16:0e_18:1) | PC | 130349353.3 |
| PC(38:7D5) | PC | 182239479.2 |
| PC(21:4) | PC | 2570705.572 |
| PC(40:9) | PC | 2647353.422 |
| PC(16:0_17:0) | PC | 4662525.431 |
| PC(17:0_18:2) | PC | 23404118.53 |
| PC(18:0e_18:2) | PC | 60992470.75 |
| PC(18:0_18:2) | PC | 2468337.394 |
| PC(40:7) | PC | 67187826.31 |
| PC(32:3e) | PC | 2193902.088 |
| PC(18:3_18:2) | PC | 19733188.68 |
| PC(36:2) | PC | 11484193.38 |
| PC(16:1e_16:0) | PC | 18423515.03 |
| PC(20:0_18:2) | PC | 11626906.23 |
| PC(22:5_18:2) | PC | 1469881.203 |
| PC(19:0_18:1) | PC | 40026385.11 |
| PC(18:0_20:4) | PC | 1548569796 |
| PC(17:0_18:1) | PC | 137347847 |
| PC(18:0_22:5) | PC | 112296724.3 |
| PC(34:2) | PC | 1418438.406 |
| PC(34:1) | PC | 5343852.992 |
| PC(19:2) | PC | 5979774.758 |
| PC(16:0_18:2) | PC | 2266312.668 |
| PC(19:1_18:1) | PC | 189309592.4 |

| | | |
|---|---|---|
| PC(22:3) | PC | 2571435.011 |
| PC(18:1_18:3) | PC | 1735463.393 |
| PC(16:1_22:6) | PC | 1266657.902 |
| PC(20:5_18:2) | PC | 1406292.356 |
| PC(38:7) | PC | 1255944.116 |
| PC(26:2) | PC | 531005.6265 |
| PC(15:0_22:5) | PC | 2188536.966 |
| PE(18:0p_18:2) | PE | 29482139.05 |
| PE(62:1) | PE | 3806842.427 |
| PE(16:0p_22:5) | PE | 14133148.23 |
| PE(18:0_20:4) | PE | 31389961.83 |
| PE(16:0_18:2) | PE | 13581587.76 |
| PE(18:0_18:1) | PE | 13272329.91 |
| PE(16:0_20:4) | PE | 4380392.943 |
| PE(18:2e) | PE | 1024116.13 |
| PE(16:0e) | PE | 1391980.662 |
| PE(21:4) | PE | 394434.6412 |
| PI(18:0_20:3) | PI | 3415821.896 |
| PI(18:0_20:4) | PI | 4880165.634 |
| SM(t40:4) | SM | 13538933.85 |
| SM(d18:1_24:2) | SM | 85495183.96 |
| SM(d42:2) | SM | 74399840.63 |
| SM(d34:0) | SM | 11982240.08 |
| SM(d42:4) | SM | 5342996.376 |
| SM(d18:1_23:1) | SM | 39527740.72 |
| SM(d20:1_20:1) | SM | 32898716.84 |
| SM(d41:3) | SM | 11923037.91 |
| SM(d39:2) | SM | 11866471.63 |
| SM(d35:1) | SM | 2752230.736 |
| SM(d17:1_21:1) | SM | 25625273.25 |

| | | |
|---|---|---|
| SM(d44:5) | SM | 9092973.819 |
| SPH(d18:1) | SPH | 7374550.555 |
| SPH(t18:0) | SPH | 2021184.298 |
| TG(18:1_18:1_16:0) | TG | 2552811830 |
| TG(16:0_18:1_18:3) | TG | 134212209.6 |
| TG(12:1e_6:0_18:3) | TG | 82660984.49 |
| TG(16:0_16:0_18:3) | TG | 46877977.02 |
| TG(18:1_18:2_20:3) | TG | 27957296.59 |
| TG(12:0e_6:0_20:5) | TG | 36769913.92 |
| TG(16:0_16:0_18:2) | TG | 74107661.35 |
| TG(18:1_18:1_18:3) | TG | 48644297.61 |
| TG(20:0e_16:0_18:0) | TG | 5165636.531 |
| TG(20:2e_18:3_20:4) | TG | 77104076.4 |
| TG(18:0e_18:2_16:0) | TG | 5149099.001 |
| TG(18:1e_16:0_16:0) | TG | 5658015.756 |
| TG(36:4) | TG | 26351842.44 |
| TG(18:0_18:1_22:6) | TG | 22153313.8 |
| TG(20:0e_18:0_18:1) | TG | 3449872.455 |
| TG(18:2_10:2_10:2) | TG | 12439798.59 |
| TG(18:4_16:0_18:2) | TG | 19928923.94 |
| TG(8:0_10:2_18:2) | TG | 11923544.25 |
| TG(18:3_18:2_18:2) | TG | 18454018.82 |
| TG(18:0_18:1_18:3) | TG | 7993082.734 |
| TG(6:0_12:2_18:1) | TG | 12439798.59 |
| TG(6:0_14:4_18:1) | TG | 1973435.269 |
| TG(20:0e_18:0_18:2) | TG | 4783997.818 |
| TG(12:0_12:0_14:0) | TG | 3076023.251 |
| TG(18:0_18:3_18:3) | TG | 6174031.977 |
| TG(14:0_18:2_18:3) | TG | 7112587.295 |
| TG(18:3e_18:4_22:3) | TG | 8563018.831 |

| | | |
|---|---|---|
| TG(18:1_18:2_18:3) | TG | 18191741.19 |
| TG(6:0_11:2_18:3) | TG | 3048748.858 |
| TG(6:0_14:4_18:2) | TG | 5073929.784 |
| TG(16:0_6:0_12:2) | TG | 8461888.435 |
| TG(8:0_10:2_18:1) | TG | 6285059.372 |
| TG(20:0e_16:0_16:0) | TG | 7911119.205 |
| TG(6:0_11:1_14:2) | TG | 2297189.461 |
| TG(20:3e_18:1_18:3) | TG | 15813622.1 |
| TG(6:0_12:2_18:2) | TG | 3809935.047 |
| TG(8:0_10:1_18:1) | TG | 3069349.426 |
| TG(20:0e_16:0_17:0) | TG | 3045627.992 |
| TG(18:2_18:2_16:0) | TG | 5779864.361 |
| TG(20:0e_18:2_16:0) | TG | 9356858.575 |
| WE(5:0_15:2) | WE | 47727155.77 |
| ZyE(18:3) | ZyE | 12483440.36 |
| ZyE() | ZyE | 149416174.3 |
| ZyE(0:0) | ZyE | 3997474.597 |

duced in spent media relative to complete media, including peak area quantification and statistical si

| Group Area: SM_pos_12022024_Newanalyis | FoldChange_CM_SM | log2FC |
|---|---|---|
| 556264.8658 | 8.5741877 | 3.1 |
| 543135.4869 | 30.69645182 | 4.94 |
| 286357.5009 | 7.516181994 | 2.91 |
| 473478.5476 | 21.70566924 | 4.44 |
| 4343146.588 | 5.351710219 | 2.42 |
| 347634.9058 | 191.3407038 | 7.58 |
| 296106.2091 | 95.67035191 | 6.58 |
| 532719.6069 | 85.0358921 | 6.41 |
| 649714.7414 | 67.6491546 | 6.08 |
| 302742.6198 | 20.67764529 | 4.37 |
| 221217.7613 | 20.11221399 | 4.33 |
| 461679.3612 | 17.3877578 | 4.12 |
| 683779.6215 | 15.77972327 | 3.98 |
| 748940.5333 | 11.87618857 | 3.57 |
| 482874.1689 | 8.75434961 | 3.13 |
| 305666.1131 | 7.568461174 | 2.92 |
| 236045.5276 | 7.06162397 | 2.82 |
| 366326.3953 | 4.658934346 | 2.22 |
| 511606.1759 | 1160.073099 | 10.18 |
| 547455.5313 | 162.016844 | 7.34 |
| 622560.9412 | 106.1529019 | 6.73 |
| 550212.9304 | 28.8400148 | 4.85 |
| 479095.4584 | 35.01739844 | 5.13 |
| 354856.7114 | 29.04061297 | 4.86 |
| 304643.8749 | 28.8400148 | 4.85 |
| 1258492.012 | 114.5632091 | 6.84 |
| 937316.5875 | 100.4267645 | 6.65 |
| 785824.7705 | 92.41146851 | 6.53 |
| 812990.3708 | 85.0358921 | 6.41 |

| | | |
|---|---|---|
| 1125362.044 | 74.02804377 | 6.21 |
| 12622007.3 | 73.00887782 | 6.19 |
| 421578.8123 | 58.08122594 | 5.86 |
| 8826864.755 | 54.94818793 | 5.78 |
| 658900.0384 | 54.56863307 | 5.77 |
| 1630449.005 | 50.91433496 | 5.67 |
| 664963.6559 | 49.8665331 | 5.64 |
| 513058.5965 | 35.75318842 | 5.16 |
| 553809.7134 | 22.31589866 | 4.48 |
| 430366.2119 | 21.11212657 | 4.4 |
| 118200.1874 | 20.2521055 | 4.34 |
| 582145.7671 | 18.50701094 | 4.21 |
| 589321.0758 | 18.50701094 | 4.21 |
| 553391.7157 | 17.02992292 | 4.09 |
| 440095.7847 | 13.64215827 | 3.77 |
| 11436611.13 | 13.45434264 | 3.75 |
| 649102.4726 | 12.81711804 | 3.68 |
| 698059.8871 | 6.821079134 | 2.77 |
| 5737373.248 | 6.680703355 | 2.74 |
| 570481.4159 | 6.634556367 | 2.73 |
| 144333.0486 | 32.67238802 | 5.03 |
| 124962.8102 | 1438.151553 | 10.49 |
| 240452.7873 | 855.1300295 | 9.74 |
| 165488.5066 | 436.5490646 | 8.77 |
| 152456.8809 | 347.2907078 | 8.44 |
| 85628.46588 | 235.5680386 | 7.88 |
| 221860.2372 | 229.1264182 | 7.84 |
| 266425.4614 | 77.17170097 | 6.27 |
| 186688.9579 | 49.8665331 | 5.64 |
| 168741.1407 | 47.83517596 | 5.58 |

| | | |
|---|---|---|
| 579999.952 | 44.32350298 | 5.47 |
| 634413.4754 | 31.12495832 | 4.96 |
| 556788.9407 | 28.64080227 | 4.84 |
| 230464.3112 | 28.4429658 | 4.83 |
| 1128302.74 | 26.90868529 | 4.75 |
| 701452.2912 | 25.8125363 | 4.69 |
| 93832.29659 | 20.82146969 | 4.38 |
| 155004.9056 | 20.11221399 | 4.33 |
| 219364.847 | 17.75311155 | 4.15 |
| 278436.4802 | 16.1112888 | 4.01 |
| 846065.8357 | 15.03236399 | 3.91 |
| 213618.541 | 15.03236399 | 3.91 |
| 98679.4065 | 12.04197398 | 3.59 |
| 709216.242 | 10.77786861 | 3.43 |
| 1639544.212 | 9.713559075 | 3.28 |
| 212260.1849 | 8.633825892 | 3.11 |
| 125964.5756 | 8.224910613 | 3.04 |
| 204777.6927 | 8.111675838 | 3.02 |
| 377725.3466 | 6.821079134 | 2.77 |
| 116460.314 | 5.775716782 | 2.53 |
| 330016.6515 | 5.656854249 | 2.5 |
| 86232.94787 | 4.0278222 | 2.01 |
| 122677.5884 | 87.42657643 | 6.45 |
| 98216.73907 | 1573.760186 | 10.62 |
| 210830.5495 | 7.012845771 | 2.81 |
| 389371.0307 | 4.890561111 | 2.29 |
| 1702239.462 | 786.8800928 | 9.62 |
| 1927297.384 | 205.0738887 | 7.68 |
| 2748468.373 | 56.10276616 | 5.81 |
| 744250.083 | 42.81368175 | 5.42 |

| | | |
|---|---|---|
| 2359871.892 | 41.64293937 | 5.38 |
| 183651.9958 | 40.50421101 | 5.34 |
| 5948923.284 | 32.89964245 | 5.04 |
| 1845717.698 | 32.44670335 | 5.02 |
| 407872321.7 | 28.8400148 | 4.85 |
| 278883.9911 | 27.09585 | 4.76 |
| 5219257.796 | 24.93326655 | 4.64 |
| 8227254.232 | 22.16175149 | 4.47 |
| 120933.064 | 21.25897303 | 4.41 |
| 133574.982 | 19.8353232 | 4.31 |
| 258740.2403 | 18.00093576 | 4.17 |
| 1350353.591 | 17.3877578 | 4.12 |
| 3696420.806 | 16.44982123 | 4.04 |
| 169220.2821 | 14.6213032 | 3.87 |
| 4707351.307 | 14.32040113 | 3.84 |
| 153514.5384 | 14.32040113 | 3.84 |
| 1399349.406 | 14.12324794 | 3.82 |
| 834972.6153 | 13.73704698 | 3.78 |
| 1342182.362 | 13.73704698 | 3.78 |
| 901040.975 | 12.90626815 | 3.69 |
| 113537.2042 | 12.90626815 | 3.69 |
| 3447693.754 | 11.63178014 | 3.54 |
| 147118556.5 | 10.55606329 | 3.4 |
| 14782350.8 | 9.317868692 | 3.22 |
| 13078540.76 | 8.5741877 | 3.1 |
| 181299.9514 | 7.835362381 | 2.97 |
| 727928.4849 | 7.361501205 | 2.88 |
| 819093.0431 | 7.310651602 | 2.87 |
| 342755.4723 | 6.634556367 | 2.73 |
| 30952063.56 | 6.105036836 | 2.61 |

| | | |
|---|---|---|
| 421309.0099 | 6.105036836 | 2.61 |
| 294720.7519 | 5.897076869 | 2.56 |
| 228285.1951 | 5.540437872 | 2.47 |
| 274613.6296 | 5.13370359 | 2.36 |
| 245529.3688 | 5.098242509 | 2.35 |
| 111068.9737 | 4.789914818 | 2.26 |
| 528556.2983 | 4.141059695 | 2.05 |
| 839330.5501 | 35.01739844 | 5.13 |
| 198723.6099 | 19.15965927 | 4.26 |
| 839950.4487 | 16.79546694 | 4.07 |
| 3622163.058 | 8.6938789 | 3.12 |
| 1658593.525 | 8.168097006 | 3.03 |
| 1651479.8 | 8.0556444 | 3.01 |
| 721228.3503 | 6.062866266 | 2.6 |
| 176577.9386 | 5.815890069 | 2.54 |
| 243689.7782 | 5.696200782 | 2.51 |
| 93932.86983 | 4.198866734 | 2.07 |
| 131871.2749 | 25.99207668 | 4.7 |
| 583516.9341 | 8.339726087 | 3.06 |
| 147691.691 | 91.77313587 | 6.52 |
| 998586.6192 | 85.62736351 | 6.42 |
| 1449428.785 | 51.26847217 | 5.68 |
| 270660.432 | 44.32350298 | 5.47 |
| 123153.4225 | 43.41133848 | 5.44 |
| 1353986.341 | 29.24260641 | 4.87 |
| 2091001.154 | 15.77972327 | 3.98 |
| 891895.1532 | 13.36140671 | 3.74 |
| 887663.7489 | 13.36140671 | 3.74 |
| 207695.5744 | 13.26911273 | 3.73 |
| 2516236.994 | 10.19648502 | 3.35 |

| | | |
|---|---|---|
| 1601516.113 | 5.696200782 | 2.51 |
| 1127459.323 | 6.543216468 | 2.71 |
| 458216.9637 | 4.407620464 | 2.14 |
| 4177550.338 | 613.1090968 | 9.26 |
| 986004.3944 | 136.2393834 | 7.09 |
| 740855.0225 | 111.4304721 | 6.8 |
| 730946.0085 | 64 | 6 |
| 534943.3553 | 52.34573175 | 5.71 |
| 771281.0831 | 47.83517596 | 5.58 |
| 1630907.48 | 45.56960626 | 5.51 |
| 1543080.557 | 31.55944654 | 4.98 |
| 174056.9685 | 29.65081798 | 4.89 |
| 3103167.851 | 24.7610399 | 4.63 |
| 208745.4208 | 24.5900029 | 4.62 |
| 232871.5306 | 24.25146506 | 4.6 |
| 1153358.938 | 22.78480313 | 4.51 |
| 1059797.092 | 20.96629446 | 4.39 |
| 166462.1922 | 20.67764529 | 4.37 |
| 793544.8337 | 15.67072476 | 3.97 |
| 1287667.636 | 15.45498126 | 3.95 |
| 808773.0261 | 14.72300241 | 3.88 |
| 1412411.188 | 13.08643294 | 3.71 |
| 650210.5517 | 12.29500145 | 3.62 |
| 1033137.153 | 12.04197398 | 3.59 |
| 164991.6246 | 11.95879399 | 3.58 |
| 416051.6702 | 11.47164198 | 3.52 |
| 317504.5701 | 9.713559075 | 3.28 |
| 688789.0554 | 8.938297105 | 3.16 |
| 841039.1836 | 8.456144324 | 3.08 |
| 1158141.452 | 7.412704495 | 2.89 |

| | | |
|---|---|---|
| 2488852.751 | 7.310651602 | 2.87 |
| 432015.41 | 7.06162397 | 2.82 |
| 759895.5514 | 6.680703355 | 2.74 |
| 1318744.301 | 6.408559021 | 2.68 |
| 990569.0879 | 6.36429187 | 2.67 |
| 1364564.068 | 5.815890069 | 2.54 |
| 399905.3708 | 5.735820992 | 2.52 |
| 3113069.061 | 5.063026376 | 2.34 |
| 777678.6775 | 4.890561111 | 2.29 |
| 670110.5412 | 4.59479342 | 2.2 |
| 681484.2473 | 4.469148552 | 2.16 |
| 1289028.558 | 4.469148552 | 2.16 |
| 2277039.583 | 4.112455307 | 2.04 |
| 4453806.951 | 10.70342044 | 3.42 |
| 679030.3142 | 18.37917368 | 4.2 |
| 8947578.13 | 16.67945217 | 4.06 |
| 789545.098 | 5.063026376 | 2.34 |

gnificance metrics.

| Adj. P-value: (CM_pos_12022024_Newanalysis) / (SM_pos_12022024_Newanalyis) | Significant |
|---|---|
| 0.029971518 | TRUE |
| 3.07581E-07 | TRUE |
| 0.000769149 | TRUE |
| 0.000620209 | TRUE |
| 0.000109321 | TRUE |
| 3.21958E-11 | TRUE |
| 3.21958E-11 | TRUE |
| 7.19757E-09 | TRUE |
| 5.77401E-10 | TRUE |
| 6.1671E-11 | TRUE |
| 7.0499E-10 | TRUE |
| 1.80399E-06 | TRUE |
| 5.81838E-06 | TRUE |
| 0.001704921 | TRUE |
| 0.000131244 | TRUE |
| 7.70933E-06 | TRUE |
| 8.21587E-06 | TRUE |
| 0.015706223 | TRUE |
| 5.05684E-11 | TRUE |
| 3.66868E-08 | TRUE |
| 3.62528E-08 | TRUE |
| 2.03285E-07 | TRUE |
| 6.54609E-08 | TRUE |
| 1.16917E-05 | TRUE |
| 2.44673E-07 | TRUE |
| 5.78869E-10 | TRUE |
| 3.44703E-05 | TRUE |
| 3.21958E-11 | TRUE |
| 2.03418E-07 | TRUE |

| | |
|---|---|
| 2.11062E-10 | TRUE |
| 0.012971296 | TRUE |
| 3.48733E-05 | TRUE |
| 0.001723598 | TRUE |
| 0.000527338 | TRUE |
| 1.67081E-08 | TRUE |
| 0.000242621 | TRUE |
| 9.92014E-07 | TRUE |
| 1.80571E-10 | TRUE |
| 2.47474E-08 | TRUE |
| 4.94793E-07 | TRUE |
| 1.67811E-07 | TRUE |
| 0.00146678 | TRUE |
| 2.20289E-08 | TRUE |
| 6.83902E-09 | TRUE |
| 0.028574293 | TRUE |
| 9.49564E-07 | TRUE |
| 0.006428085 | TRUE |
| 0.002083661 | TRUE |
| 0.000218227 | TRUE |
| 1.64636E-10 | TRUE |
| 2.10258E-05 | TRUE |
| 9.82558E-08 | TRUE |
| 4.33783E-07 | TRUE |
| 2.07031E-09 | TRUE |
| 2.03921E-09 | TRUE |
| 4.91585E-05 | TRUE |
| 3.21957E-08 | TRUE |
| 3.21958E-11 | TRUE |
| 5.83738E-06 | TRUE |

| | |
|---|---|
| 1.41511E-05 | TRUE |
| 0.002126778 | TRUE |
| 8.74477E-05 | TRUE |
| 1.35139E-05 | TRUE |
| 5.42967E-07 | TRUE |
| 0.000104489 | TRUE |
| 1.96788E-05 | TRUE |
| 0.000156689 | TRUE |
| 2.6644E-06 | TRUE |
| 0.000218949 | TRUE |
| 3.39723E-06 | TRUE |
| 3.45541E-07 | TRUE |
| 1.66026E-05 | TRUE |
| 3.67159E-05 | TRUE |
| 0.001717704 | TRUE |
| 0.005579733 | TRUE |
| 0.038255801 | TRUE |
| 0.003843454 | TRUE |
| 0.012240684 | TRUE |
| 5.04328E-05 | TRUE |
| 0.011341514 | TRUE |
| 0.008615779 | TRUE |
| 6.51812E-06 | TRUE |
| 6.85544E-06 | TRUE |
| 0.002615311 | TRUE |
| 0.040305904 | TRUE |
| 2.70781E-09 | TRUE |
| 2.77373E-06 | TRUE |
| 1.07081E-05 | TRUE |
| 0.00037033 | TRUE |

| | |
|---|---|
| 0.000230207 | TRUE |
| 1.43773E-10 | TRUE |
| 5.37349E-07 | TRUE |
| 0.003190271 | TRUE |
| 0.029771282 | TRUE |
| 0.000176405 | TRUE |
| 0.000551205 | TRUE |
| 2.54891E-06 | TRUE |
| 0.000122973 | TRUE |
| 0.013156242 | TRUE |
| 0.000515997 | TRUE |
| 0.008398202 | TRUE |
| 0.000116159 | TRUE |
| 0.014322846 | TRUE |
| 3.73298E-06 | TRUE |
| 0.011046097 | TRUE |
| 4.57006E-06 | TRUE |
| 3.35141E-05 | TRUE |
| 0.000816137 | TRUE |
| 0.00307318 | TRUE |
| 0.001973351 | TRUE |
| 0.003229801 | TRUE |
| 0.015858725 | TRUE |
| 0.000164031 | TRUE |
| 0.005025985 | TRUE |
| 0.031684156 | TRUE |
| 0.000174417 | TRUE |
| 0.010967885 | TRUE |
| 2.62731E-05 | TRUE |
| 0.004911986 | TRUE |

| | |
|---|---|
| 0.000641955 | TRUE |
| 5.37213E-05 | TRUE |
| 0.022653896 | TRUE |
| 0.000198227 | TRUE |
| 0.000380772 | TRUE |
| 0.016417447 | TRUE |
| 0.04417354 | TRUE |
| 2.09823E-09 | TRUE |
| 0.000162034 | TRUE |
| 0.000577809 | TRUE |
| 9.48465E-05 | TRUE |
| 1.96712E-05 | TRUE |
| 0.048894017 | TRUE |
| 4.2037E-06 | TRUE |
| 0.00073715 | TRUE |
| 0.000955815 | TRUE |
| 0.018968559 | TRUE |
| 7.1973E-06 | TRUE |
| 0.003544484 | TRUE |
| 0.014531446 | TRUE |
| 6.63775E-05 | TRUE |
| 0.000153454 | TRUE |
| 4.17827E-05 | TRUE |
| 0.00012238 | TRUE |
| 1.67732E-05 | TRUE |
| 0.000139131 | TRUE |
| 0.002180119 | TRUE |
| 0.002578532 | TRUE |
| 0.004200484 | TRUE |
| 0.002851188 | TRUE |

| | |
|---|---|
| 0.003566022 | TRUE |
| 0.014800638 | TRUE |
| 0.029117655 | TRUE |
| 4.31502E-09 | TRUE |
| 4.88888E-11 | TRUE |
| 1.21359E-10 | TRUE |
| 3.21958E-11 | TRUE |
| 1.92324E-10 | TRUE |
| 4.9916E-11 | TRUE |
| 0.000101123 | TRUE |
| 5.49754E-05 | TRUE |
| 0.000693021 | TRUE |
| 4.14816E-06 | TRUE |
| 1.15002E-07 | TRUE |
| 1.3025E-08 | TRUE |
| 3.43884E-11 | TRUE |
| 1.43992E-05 | TRUE |
| 0.004934026 | TRUE |
| 1.76643E-09 | TRUE |
| 8.22174E-08 | TRUE |
| 7.6992E-11 | TRUE |
| 0.000549912 | TRUE |
| 5.95667E-06 | TRUE |
| 2.52171E-09 | TRUE |
| 3.93178E-10 | TRUE |
| 0.000401457 | TRUE |
| 8.17059E-05 | TRUE |
| 0.001313559 | TRUE |
| 2.78331E-07 | TRUE |
| 0.004609884 | TRUE |

| | |
|---|---|
| 0.012218122 | TRUE |
| 0.002422859 | TRUE |
| 1.04491E-05 | TRUE |
| 0.009925619 | TRUE |
| 0.013126995 | TRUE |
| 0.000529542 | TRUE |
| 0.000708552 | TRUE |
| 0.002750229 | TRUE |
| 3.09895E-07 | TRUE |
| 0.000732679 | TRUE |
| 0.020787141 | TRUE |
| 0.006682818 | TRUE |
| 0.001000021 | TRUE |
| 0.003873774 | TRUE |
| 5.91208E-06 | TRUE |
| 2.30305E-06 | TRUE |
| 0.000763856 | TRUE |

| Reduced_in_SM |
| --- |
| TRUE |
| TRUE |
| TRUE |
| TRUE |
| TRUE |
| TRUE |
| TRUE |
| TRUE |
| TRUE |
| TRUE |
| TRUE |
| TRUE |
| TRUE |
| TRUE |
| TRUE |
| TRUE |
| TRUE |
| TRUE |
| TRUE |
| TRUE |
| TRUE |
| TRUE |
| TRUE |
| TRUE |
| TRUE |
| TRUE |
| TRUE |
| TRUE |
| TRUE |

| |
|---|
| TRUE |
| TRUE |
| TRUE |
| TRUE |
| TRUE |
| TRUE |
| TRUE |
| TRUE |
| TRUE |
| TRUE |
| TRUE |
| TRUE |
| TRUE |
| TRUE |
| TRUE |
| TRUE |
| TRUE |
| TRUE |
| TRUE |
| TRUE |
| TRUE |
| TRUE |
| TRUE |
| TRUE |
| TRUE |
| TRUE |
| TRUE |
| TRUE |
| TRUE |
| TRUE |

TRUE
TRUE
TRUE
TRUE
TRUE
TRUE
TRUE
TRUE
TRUE
TRUE
TRUE
TRUE
TRUE
TRUE
TRUE
TRUE
TRUE
TRUE
TRUE
TRUE
TRUE
TRUE
TRUE
TRUE
TRUE
TRUE
TRUE
TRUE
TRUE
TRUE

TRUE
TRUE
TRUE
TRUE
TRUE
TRUE
TRUE
TRUE
TRUE
TRUE
TRUE
TRUE
TRUE
TRUE
TRUE
TRUE
TRUE
TRUE
TRUE
TRUE
TRUE
TRUE
TRUE
TRUE
TRUE
TRUE
TRUE
TRUE
TRUE
TRUE

| |
|---|
| TRUE |
| TRUE |
| TRUE |
| TRUE |
| TRUE |
| TRUE |
| TRUE |
| TRUE |
| TRUE |
| TRUE |
| TRUE |
| TRUE |
| TRUE |
| TRUE |
| TRUE |
| TRUE |
| TRUE |
| TRUE |
| TRUE |
| TRUE |
| TRUE |
| TRUE |
| TRUE |
| TRUE |
| TRUE |
| TRUE |
| TRUE |
| TRUE |
| TRUE |
| TRUE |

TRUE
TRUE
TRUE
TRUE
TRUE
TRUE
TRUE
TRUE
TRUE
TRUE
TRUE
TRUE
TRUE
TRUE
TRUE
TRUE
TRUE
TRUE
TRUE
TRUE
TRUE
TRUE
TRUE
TRUE
TRUE
TRUE
TRUE
TRUE
TRUE
TRUE

TRUE
TRUE
TRUE
TRUE
TRUE
TRUE
TRUE
TRUE
TRUE
TRUE
TRUE
TRUE
TRUE
TRUE
TRUE
TRUE
TRUE
