## Supplementary file 2 for "A lipid compendium of a metabolically compromised bacterium provides insights into lipid acquisition, biosynthesis, and metabolism"

### Supplementary spreadsheet 2: Untargeted lipidomics analysis of ESI-positive mode data obtaine

| Name | Formula | Annot. DeltaM | Calc. MW | m/z | RT [min] | MS Depth |
|---|---|---|---|---|---|---|
| ZyE(20:5) | C47 H72 O2 | 1.53 | 668.55425 | 669.5615 | 17.299 | 2 |
| ZyE(18:3) | C45 H72 O2 | 1.36 | 644.55411 | 645.56139 | 18.665 | 2 |
| ZyE(0:0) | C27 H44 O | 1.12 | 384.33965 | 385.34692 | 12.666 | 2 |
| ZyE() | C27 H44 O | 2.88 | 384.34032 | 367.33704 | 24.547 | 2 |
| WE(5:0_19:2) | C24 H44 O2 | 1.89 | 364.33482 | 365.3421 | 14.246 | 2 |
| WE(5:0_16:1) | C21 H40 O2 | 0.5 | 324.30299 | 325.31027 | 9.722 | 2 |
| WE(5:0_15:2) | C20 H36 O2 | 0.87 | 308.2718 | 309.27908 | 7.342 | 2 |
| WE(4:0_22:2) | C26 H48 O2 | 2.41 | 392.36638 | 393.37366 | 16.099 | 2 |
| WE(4:0_19:1) | C23 H44 O2 | 1.37 | 352.33461 | 353.34189 | 14.535 | 1 |
| WE(4:0_16:3) | C20 H34 O2 | 1.01 | 306.25619 | 307.26346 | 8.581 | 1 |
| WE(4:0_14:2) | C18 H32 O2 | 0.59 | 280.24039 | 281.24768 | 4.925 | 1 |
| WE(3:0_20:2) | C23 H42 O2 | 2.14 | 350.31923 | 383.35275 | 8.378 | 2 |
| WE(3:0_17:2) | C20 H36 O2 | 0.2 | 308.27159 | 309.27887 | 6.847 | 2 |
| WE(3:0_16:3) | C19 H32 O2 | 0.87 | 292.24048 | 293.24775 | 5.284 | 2 |
| WE(2:0_14:2) | C16 H28 O2 | 0.2 | 252.20898 | 253.21626 | 4.086 | 1 |
| TG(9:0_9:0_9:0) | C30 H56 O6 | -5.98 | 512.40462 | 513.41382 | 11.52 | 2 |
| TG(8:0_8:0_10:3) | C29 H48 O6 | -4.42 | 492.34291 | 493.35019 | 10.358 | 1 |
| TG(8:0_10:2_18:2) | C39 H66 O6 | -0.48 | 630.48564 | 631.49253 | 11.854 | 2 |
| TG(8:0_10:2_18:1) | C39 H68 O6 | 1.13 | 632.5023 | 633.50956 | 12.965 | 2 |
| TG(8:0_10:1_18:1) | C39 H70 O6 | 0.94 | 634.51784 | 652.55249 | 12.526 | 1 |
| TG(6:0_6:0_13:0) | C28 H52 O6 | -2.54 | 484.37516 | 485.38251 | 8.246 | 1 |
| TG(6:0_14:4_18:2) | C41 H66 O6 | -2.79 | 654.48411 | 655.49139 | 11.083 | 2 |
| TG(6:0_14:4_18:1) | C41 H68 O6 | -2.15 | 656.50018 | 657.50745 | 12.639 | 2 |
| TG(6:0_12:2_18:2) | C39 H66 O6 | 0.82 | 630.48646 | 631.49352 | 11.541 | 2 |
| TG(6:0_12:2_18:1) | C39 H68 O6 | 0.38 | 632.50183 | 655.49119 | 11.293 | 2 |
| TG(6:0_11:2_18:3) | C38 H62 O6 | -0.9 | 614.45409 | 615.46051 | 7.519 | 1 |
| TG(6:0_11:2_18:2) | C38 H64 O6 | -2.18 | 616.46894 | 617.47632 | 8.353 | 2 |
| TG(6:0_11:1_18:2) | C38 H66 O6 | -2.55 | 618.48436 | 619.49164 | 9.13 | 1 |
| TG(6:0_11:1_14:2) | C34 H58 O6 | -2.02 | 562.4222 | 563.42914 | 8.217 | 1 |
| TG(6:0_10:3_10:4) | C29 H40 O6 | 0.08 | 484.28253 | 485.28995 | 19.407 | 2 |
| TG(4:0_6:0_17:1) | C30 H54 O6 | 1.11 | 510.3926 | 511.40003 | 8.697 | 2 |
| TG(4:0_6:0_11:1) | C24 H42 O6 | 0.78 | 426.29847 | 427.30575 | 18.413 | 2 |
| TG(4:0_16:0_18:2) | C41 H74 O6 | 8.29 | 662.55403 | 663.56131 | 19.407 | 2 |

| | | | | | | |
|---|---|---|---|---|---|---|
| TG(4:0_16:0_18:1) | C41 H76 O6 | 9.03 | 664.57019 | 665.57747 | 19.646 | 2 |
| TG(36:4) | C39 H66 O6 | -2.4 | 630.48442 | 631.4917 | 12.386 | 2 |
| TG(22:1_10:3_10:3) | C45 H72 O6 | 1.14 | 708.5337 | 709.54104 | 13.824 | 2 |
| TG(20:3e_18:1_18:3) | C59 H102 O5 | 5.17 | 890.77733 | 891.78461 | 19.27 | 1 |
| TG(20:3e_11:3_20:3) | C54 H88 O5 | -3.71 | 816.66014 | 817.66742 | 18.478 | 3 |
| TG(20:3e_11:2_18:3) | C52 H86 O5 | -3.44 | 790.64481 | 791.6517 | 18.366 | 3 |
| TG(20:2e_18:3_20:4) | C61 H102 O5 | 4.23 | 914.7766 | 915.78387 | 18.773 | 2 |
| TG(20:2e_11:2_18:3) | C52 H88 O5 | -3.07 | 792.66074 | 793.6675 | 18.66 | 3 |
| TG(20:2e_10:2_12:3) | C45 H74 O5 | 0.85 | 694.55422 | 695.56039 | 17.33 | 2 |
| TG(20:1e_18:3_20:4) | C61 H104 O5 | 2.21 | 916.7904 | 917.79768 | 18.729 | 1 |
| TG(20:1e_11:3_11:3) | C45 H74 O5 | 0.19 | 694.55376 | 695.56104 | 14.644 | 1 |
| TG(20:1_18:1_18:1) | C59 H108 O6 | -1.27 | 912.81343 | 913.82071 | 21.967 | 1 |
| TG(20:0e_18:2_16:0) | C57 H108 O5 | 0.81 | 872.82038 | 890.8542 | 20.931 | 3 |
| TG(20:0e_18:1_16:0) | C57 H110 O5 | 0.63 | 874.83588 | 892.8697 | 21.183 | 3 |
| TG(20:0e_18:0_18:2) | C59 H112 O5 | 0.53 | 900.85145 | 918.88526 | 21.226 | 3 |
| TG(20:0e_18:0_18:1) | C59 H114 O5 | 0.74 | 902.86729 | 920.90112 | 21.508 | 3 |
| TG(20:0e_17:0_18:1) | C58 H112 O5 | 0.33 | 888.85127 | 906.88503 | 21.344 | 3 |
| TG(20:0e_16:0_18:0) | C57 H112 O5 | 0.86 | 876.85173 | 894.88557 | 21.52 | 3 |
| TG(20:0e_16:0_17:0) | C56 H110 O5 | 1.13 | 862.8363 | 880.87012 | 21.342 | 3 |
| TG(20:0e_16:0_16:0) | C55 H108 O5 | 0.69 | 848.82026 | 866.85409 | 21.193 | 3 |
| TG(20:0e_15:0_16:0) | C54 H106 O5 | 3.55 | 834.80699 | 852.83847 | 21.045 | 3 |
| TG(20:0_16:0_16:0) | C55 H106 O6 | 1.01 | 862.79981 | 880.83337 | 19.566 | 2 |
| TG(19:1_18:1_18:4) | C58 H100 O6 | 3.09 | 892.75475 | 893.76202 | 19.18 | 1 |
| TG(19:1_12:4_12:4) | C46 H70 O6 | -3.54 | 718.5147 | 719.52197 | 14.627 | 1 |
| TG(19:0) | C22 H40 O6 | 0.99 | 400.28288 | 401.29016 | 18.622 | 1 |
| TG(18:4_6:0_18:4) | C45 H70 O6 | 1.15 | 706.51805 | 707.52533 | 12.228 | 1 |
| TG(18:4_18:2_18:2) | C57 H94 O6 | 0.02 | 874.70506 | 875.71234 | 16.305 | 2 |
| TG(18:4_17:1_18:1) | C56 H96 O6 | 4.38 | 864.72447 | 865.73175 | 18.747 | 1 |
| TG(18:4_16:1_17:1) | C54 H92 O6 | 4.07 | 836.6928 | 837.70007 | 19.126 | 1 |
| TG(18:4_16:0_18:2) | C55 H94 O6 | 0.86 | 850.70577 | 851.71305 | 16.385 | 1 |
| TG(18:4_16:0_18:1) | C55 H96 O6 | 0.75 | 852.72133 | 853.7286 | 18.696 | 2 |
| TG(18:3e_18:4_22:3) | C61 H100 O5 | 6.48 | 912.76299 | 913.77026 | 18.602 | 1 |
| TG(18:3_18:3_16:0) | C55 H94 O6 | -2.2 | 850.70317 | 851.71045 | 20.706 | 1 |
| TG(18:3_18:2_18:2) | C57 H96 O6 | -2.62 | 876.71839 | 877.72567 | 19.873 | 1 |
| TG(18:3_18:2_16:0) | C55 H96 O6 | 0.58 | 852.72118 | 870.75502 | 20.584 | 3 |

| | | | | | | |
|---|---|---|---|---|---|---|
| TG(18:2_18:2_20:4) | C59 H98 O6 | 5.26 | 902.74109 | 903.74837 | 19.515 | 1 |
| TG(18:2_18:2_20:1) | C59 H104 O6 | -1.19 | 908.78221 | 909.78949 | 20.491 | 1 |
| TG(18:2_18:2_18:2) | C57 H98 O6 | 0.7 | 878.73695 | 879.74363 | 19.728 | 2 |
| TG(18:2_18:2_18:1) | C57 H100 O6 | -1.94 | 880.75028 | 881.75728 | 21.165 | 1 |
| TG(18:2_18:2_17:0) | C56 H100 O6 | 2.61 | 868.75426 | 869.76153 | 19.865 | 1 |
| TG(18:2_18:2_16:0) | C55 H98 O6 | 0.6 | 854.73685 | 872.77068 | 20.764 | 3 |
| TG(18:2_18:2_15:0) | C54 H96 O6 | -4.62 | 840.71681 | 919.73597 | 19.632 | 1 |
| TG(18:2_10:2_11:4) | C42 H64 O6 | 2.27 | 664.47179 | 665.47907 | 9.294 | 1 |
| TG(18:2_10:2_10:2) | C41 H66 O6 | -2.57 | 654.48426 | 655.49153 | 11.55 | 2 |
| TG(18:2_10:1_10:4) | C41 H64 O6 | -1.66 | 652.46921 | 653.47611 | 10.592 | 2 |
| TG(18:1e_16:0_16:0) | C53 H102 O5 | 0.86 | 818.77343 | 836.80725 | 20.628 | 3 |
| TG(18:1_18:2_20:3) | C59 H102 O6 | 5.44 | 906.77257 | 907.77985 | 18.41 | 1 |
| TG(18:1_18:2_18:3) | C57 H98 O6 | 1.78 | 878.7379 | 879.74522 | 18.807 | 2 |
| TG(18:1_18:2_18:2) | C57 H100 O6 | -0.08 | 880.75192 | 881.75922 | 19.929 | 2 |
| TG(18:1_18:2_18:1) | C57 H102 O6 | -2.51 | 882.76543 | 883.77271 | 21.412 | 1 |
| TG(18:1_18:2_16:0) | C55 H100 O6 | 0.7 | 856.75259 | 857.75996 | 20.045 | 3 |
| TG(18:1_18:1_22:6) | C61 H102 O6 | 4.57 | 930.7719 | 931.77917 | 18.646 | 1 |
| TG(18:1_18:1_22:4) | C61 H106 O6 | 3.71 | 934.80241 | 917.79919 | 18.728 | 1 |
| TG(18:1_18:1_20:1) | C59 H108 O6 | -2.23 | 912.81256 | 913.81984 | 20.949 | 1 |
| TG(18:1_18:1_18:3) | C57 H100 O6 | 0.72 | 880.75263 | 881.75988 | 18.989 | 2 |
| TG(18:1_18:1_18:2) | C57 H102 O6 | 0.41 | 882.768 | 883.77501 | 20.399 | 2 |
| TG(18:1_18:1_18:1) | C57 H104 O6 | 1.06 | 884.78422 | 885.78947 | 20.256 | 3 |
| TG(18:1_18:1_16:0) | C55 H102 O6 | -0.23 | 858.76744 | 876.80127 | 20.221 | 3 |
| TG(18:0e_18:2_16:0) | C55 H104 O5 | 0.43 | 844.78874 | 862.82265 | 20.636 | 3 |
| TG(18:0e_18:1_16:0) | C55 H106 O5 | 1.07 | 846.80493 | 864.83874 | 20.911 | 3 |
| TG(18:0e_16:0_16:0) | C53 H104 O5 | 0.49 | 820.78878 | 838.82256 | 20.891 | 3 |
| TG(18:0_18:3_18:3) | C57 H98 O6 | 0.45 | 878.73673 | 879.74401 | 17.481 | 1 |
| TG(18:0_18:1_22:6) | C61 H104 O6 | 4.67 | 932.78765 | 933.79492 | 19.133 | 2 |
| TG(18:0_18:1_18:3) | C57 H102 O6 | 0.65 | 882.76822 | 883.77548 | 19.238 | 2 |
| TG(18:0_16:0_18:3) | C55 H100 O6 | 0.31 | 856.75225 | 857.75954 | 19.2 | 2 |
| TG(18:0_16:0_16:0) | C53 H102 O6 | 0.46 | 834.76803 | 835.7756 | 19.362 | 3 |
| TG(17:0_18:1D7_17:1) | C55 H95 D7 O | 0.96 | 865.81241 | 866.81763 | 20.193 | 3 |
| TG(16:2e_6:0_12:2) | C37 H64 O5 | 0.96 | 588.47594 | 589.48322 | 12.206 | 2 |
| TG(16:1_18:2_18:3) | C55 H94 O6 | 0.53 | 850.70549 | 851.71249 | 19.355 | 2 |
| TG(16:1_18:2_16:0) | C53 H96 O6 | -0.17 | 828.72055 | 829.72832 | 19.852 | 2 |

| | | | | | | |
|---|---|---|---|---|---|---|
| TG(16:1_18:2_15:0) | C52 H94 O6 | -6.03 | 814.70013 | 815.71345 | 19.597 | 2 |
| TG(16:1_18:1_20:5) | C57 H96 O6 | 1.13 | 876.72168 | 877.72891 | 16.846 | 2 |
| TG(16:1_17:1_18:2) | C54 H96 O6 | 2.32 | 840.72264 | 841.72992 | 20.966 | 1 |
| TG(16:1_14:1_15:0) | C48 H88 O6 | 2.78 | 760.6602 | 778.69409 | 20.264 | 1 |
| TG(16:1_12:0_14:0) | C45 H84 O6 | 1.39 | 720.62779 | 721.63507 | 20.342 | 1 |
| TG(16:0_8:0_10:2) | C37 H66 O6 | 0.98 | 606.48653 | 607.49372 | 12.937 | 2 |
| TG(16:0_8:0_10:0) | C37 H70 O6 | 0.25 | 610.51739 | 628.55121 | 20.855 | 2 |
| TG(16:0_6:0_17:0) | C42 H80 O6 | -2.76 | 680.59361 | 681.60089 | 20.12 | 2 |
| TG(16:0_6:0_12:2) | C37 H66 O6 | 1.85 | 606.48706 | 607.49426 | 12.186 | 2 |
| TG(16:0_18:2_22:4) | C59 H102 O6 | 4.22 | 906.77147 | 907.77875 | 20.474 | 1 |
| TG(16:0_18:2_17:1) | C54 H98 O6 | -4.57 | 842.73249 | 921.75214 | 19.879 | 1 |
| TG(16:0_18:2_16:0) | C53 H98 O6 | -0.25 | 830.73613 | 831.74341 | 21.142 | 1 |
| TG(16:0_18:1_22:6) | C59 H100 O6 | 3.38 | 904.75504 | 905.76093 | 19.13 | 1 |
| TG(16:0_18:1_18:3) | C55 H98 O6 | 1.84 | 854.73791 | 855.74493 | 19.38 | 1 |
| TG(16:0_16:0_18:3) | C53 H96 O6 | 1.03 | 828.72154 | 829.72885 | 18.914 | 2 |
| TG(16:0_16:0_18:2) | C53 H98 O6 | -3.17 | 830.7337 | 831.73721 | 19.777 | 1 |
| TG(16:0_16:0_16:0) | C51 H98 O6 | -2.23 | 806.73454 | 807.74182 | 19.593 | 1 |
| TG(16:0_14:0_22:5) | C55 H96 O6 | -2.54 | 852.71852 | 853.7258 | 20.92 | 1 |
| TG(16:0_10:3_11:4) | C40 H62 O6 | -2.67 | 638.45293 | 671.48633 | 11.053 | 2 |
| TG(16:0_10:2_11:4) | C40 H64 O6 | 0.69 | 640.47073 | 658.50485 | 10.191 | 2 |
| TG(15:0_6:0_12:1) | C36 H66 O6 | 1.21 | 594.48666 | 612.52032 | 9.251 | 2 |
| TG(15:0_18:2_18:3) | C54 H94 O6 | -8.21 | 838.69815 | 839.70543 | 19.592 | 1 |
| TG(15:0_16:1_14:0) | C48 H90 O6 | 6.79 | 762.67892 | 763.6862 | 20.002 | 2 |
| TG(15:0_14:0_14:0) | C46 H88 O6 | -1.77 | 736.65679 | 737.66406 | 20.075 | 1 |
| TG(14:0_18:2_18:3) | C53 H92 O6 | 1.01 | 824.69022 | 825.69751 | 18.37 | 2 |
| TG(14:0_14:1_13:0) | C44 H82 O6 | -1.72 | 706.60992 | 707.6172 | 19.549 | 1 |
| TG(14:0_14:0_14:3) | C45 H80 O6 | -2.87 | 716.59343 | 717.60071 | 23.841 | 2 |
| TG(14:0_14:0_14:0) | C45 H86 O6 | -2.28 | 722.64079 | 723.64807 | 25.703 | 1 |
| TG(12:1e_6:0_18:4) | C39 H66 O5 | 1.33 | 614.49184 | 615.4993 | 11.75 | 2 |
| TG(12:1e_6:0_18:3) | C39 H68 O5 | 1.4 | 616.50754 | 634.54137 | 16.253 | 2 |
| TG(12:1e_12:2_12:3) | C39 H64 O5 | 1.5 | 612.47629 | 613.48357 | 11.808 | 1 |
| TG(12:0e_6:0_20:5) | C41 H70 O5 | -2.37 | 642.5208 | 643.52808 | 17.682 | 2 |
| TG(12:0e_6:0_10:3) | C31 H54 O5 | -3.6 | 506.3953 | 507.40251 | 13.294 | 2 |
| TG(12:0_14:0_16:0) | C45 H86 O6 | -2.36 | 722.64073 | 723.64801 | 19.209 | 1 |
| TG(12:0_12:0_14:0) | C41 H78 O6 | 9.97 | 666.58648 | 667.59376 | 20.209 | 1 |

| | | | | | | |
|---|---|---|---|---|---|---|
| TG(10:0_18:1_18:3) | C49 H86 O6 | 5.94 | 770.64702 | 771.6543 | 18.719 | 1 |
| SPH(t18:1) | C18 H37 N O3 | 2.52 | 315.27814 | 316.28522 | 4.534 | 2 |
| SPH(t18:0) | C18 H39 N O3 | -5.68 | 317.29119 | 318.30057 | 3.257 | 2 |
| SPH(d24:0) | C24 H51 N O2 | 3.04 | 385.39315 | 386.40042 | 12.19 | 2 |
| SPH(d22:0) | C22 H47 N O2 | 2.58 | 357.3616 | 358.36888 | 8.472 | 2 |
| SPH(d21:1) | C21 H43 N O2 | 0.61 | 341.32959 | 342.33686 | 9.043 | 2 |
| SPH(d20:1) | C20 H41 N O2 | 0.87 | 327.31402 | 328.32128 | 7.866 | 2 |
| SPH(d19:1) | C19 H39 N O2 | 0.16 | 313.29813 | 314.3054 | 7.843 | 2 |
| SPH(d18:1) | C18 H37 N O2 | 0.94 | 299.28271 | 341.3166 | 3.653 | 2 |
| SPH(d18:0) | C18 H39 N O2 | 1.23 | 301.29845 | 302.30573 | 4.329 | 2 |
| SPH(d16:1) | C16 H33 N O2 | 1.21 | 271.25146 | 272.25873 | 4.064 | 1 |
| SM(t42:4) | C47 H89 N2 O | 1.21 | 824.64173 | 825.64903 | 11.764 | 2 |
| SM(t42:3) | C47 H91 N2 O | 0.79 | 826.65704 | 827.66431 | 11.412 | 2 |
| SM(t40:4) | C45 H85 N2 O | -0.43 | 796.6091 | 797.61797 | 9.159 | 2 |
| SM(t34:2) | C39 H77 N2 O | 0.91 | 716.54749 | 717.55477 | 11.114 | 2 |
| SM(t34:1) | C39 H79 N2 O | 1.18 | 718.56334 | 719.57062 | 12.458 | 2 |
| SM(t34:0) | C39 H81 N2 O | -3.4 | 720.57569 | 762.60913 | 12.567 | 2 |
| SM(t18:0_16:1) | C39 H79 N2 O | 1.38 | 718.56348 | 719.57076 | 10.845 | 2 |
| SM(d44:5) | C49 H91 N2 O | 1.59 | 834.6628 | 835.67007 | 16.352 | 2 |
| SM(d43:3) | C48 H93 N2 O | 0.76 | 824.67775 | 825.68503 | 18.082 | 2 |
| SM(d43:2) | C48 H95 N2 O | 0.98 | 826.69359 | 827.70086 | 19.171 | 2 |
| SM(d43:1) | C48 H97 N2 O | 0.84 | 828.70912 | 829.7164 | 24.007 | 2 |
| SM(d42:5) | C47 H87 N2 O | 0.97 | 806.63096 | 807.63825 | 14.802 | 2 |
| SM(d42:4) | C47 H89 N2 O | 0.91 | 808.64656 | 809.65383 | 16.192 | 2 |
| SM(d42:3) | C47 H91 N2 O | -1.84 | 810.65998 | 811.66852 | 17.957 | 2 |
| SM(d42:2) | C47 H93 N2 O | 0.61 | 812.67762 | 813.68489 | 18.455 | 2 |
| SM(d41:3) | C46 H89 N2 O | 0.33 | 796.64609 | 797.6534 | 16.445 | 2 |
| SM(d41:2) | C46 H91 N2 O | -0.23 | 798.66129 | 799.66857 | 18.806 | 2 |
| SM(d41:1) | C46 H93 N2 O | 0.12 | 800.67722 | 801.6845 | 19.553 | 2 |
| SM(d40:4) | C45 H85 N2 O | 1.36 | 780.61559 | 781.62286 | 14.345 | 2 |
| SM(d40:3) | C45 H87 N2 O | 1.15 | 782.63108 | 783.63835 | 15.781 | 2 |
| SM(d40:1) | C45 H91 N2 O | 1.17 | 786.6624 | 787.66967 | 18.564 | 2 |
| SM(d39:2) | C44 H87 N2 O | 1.01 | 770.63096 | 771.63823 | 16.44 | 2 |
| SM(d39:1) | C44 H89 N2 O | 0.97 | 772.64658 | 773.65385 | 17.929 | 2 |
| SM(d38:5) | C43 H79 N2 O | -0.27 | 750.56737 | 751.57465 | 13.63 | 2 |

| | | | | | | |
|---|---|---|---|---|---|---|
| SM(d38:4) | C43 H81 N2 O | -2.9 | 752.58104 | 753.58832 | 15.519 | 1 |
| SM(d38:3) | C43 H83 N2 O | 1.01 | 754.59963 | 755.60692 | 13.739 | 2 |
| SM(d38:1) | C43 H87 N2 O | 0.87 | 758.63083 | 759.63811 | 17.002 | 2 |
| SM(d37:2) | C42 H83 N2 O | 1.07 | 742.59967 | 743.60703 | 14.583 | 2 |
| SM(d37:1) | C42 H85 N2 O | 0.25 | 744.61471 | 745.62199 | 15.986 | 2 |
| SM(d36:5) | C41 H75 N2 O | -1.79 | 722.53498 | 723.54226 | 11.855 | 2 |
| SM(d36:4) | C41 H77 N2 O | -2.38 | 724.5502 | 725.55709 | 13.327 | 2 |
| SM(d36:3) | C41 H79 N2 O | 1.4 | 726.56859 | 727.57587 | 12.653 | 2 |
| SM(d36:2D9) | C41 H72 D9 N | 1 | 737.64046 | 738.64774 | 13.523 | 2 |
| SM(d36:2) | C41 H81 N2 O | 0.91 | 728.58389 | 729.59121 | 13.992 | 2 |
| SM(d36:0) | C41 H85 N2 O | 1.9 | 732.61592 | 733.62319 | 15.653 | 2 |
| SM(d35:4) | C40 H75 N2 O | -3.05 | 710.53411 | 711.54138 | 12.693 | 1 |
| SM(d35:2) | C40 H79 N2 O | 1.45 | 714.56861 | 715.57589 | 12.715 | 2 |
| SM(d35:1) | C40 H81 N2 O | 0.85 | 716.58383 | 717.59111 | 14.225 | 2 |
| SM(d34:3) | C39 H75 N2 O | 0.95 | 698.53693 | 699.54421 | 10.662 | 2 |
| SM(d34:0) | C39 H81 N2 O | -0.15 | 704.58312 | 705.59039 | 14.165 | 2 |
| SM(d33:2) | C38 H75 N2 O | 0.93 | 686.53691 | 687.54419 | 11.045 | 2 |
| SM(d33:1) | C38 H77 N2 O | 0.89 | 688.55254 | 689.55972 | 12.73 | 2 |
| SM(d33:0) | C38 H79 N2 O | 0.99 | 690.56826 | 691.57554 | 13.038 | 2 |
| SM(d32:2) | C37 H73 N2 O | 1.37 | 672.52154 | 673.52883 | 10.192 | 2 |
| SM(d32:1) | C37 H75 N2 O | 1.05 | 674.53698 | 675.54426 | 11.472 | 2 |
| SM(d32:0) | C37 H77 N2 O | 1.38 | 676.55286 | 677.56015 | 12.103 | 2 |
| SM(d31:1) | C36 H73 N2 O | 1.56 | 660.52166 | 661.52893 | 10.652 | 2 |
| SM(d30:2) | C35 H69 N2 O | 1.8 | 644.49048 | 645.49772 | 8.754 | 2 |
| SM(d30:1) | C35 H71 N2 O | 1.4 | 646.50588 | 647.51316 | 9.904 | 2 |
| SM(d28:1) | C33 H67 N2 O | 1.56 | 618.47464 | 619.48193 | 8.56 | 2 |
| SM(d20:1_20:1) | C45 H89 N2 O | 1.21 | 784.64678 | 785.65404 | 16.972 | 2 |
| SM(d19:1_17:0) | C41 H83 N2 O | 0.31 | 730.5991 | 731.60685 | 15.459 | 2 |
| SM(d19:0_23:1) | C47 H95 N2 O | 1.86 | 814.69429 | 815.70156 | 23.147 | 2 |
| SM(d18:1_24:2) | C47 H91 N2 O | 1 | 810.66228 | 811.66957 | 17.155 | 2 |
| SM(d18:1_24:1) | C47 H93 N2 O | 0.51 | 812.67754 | 813.68482 | 19.534 | 2 |
| SM(d18:1_23:1) | C46 H91 N2 O | 1.37 | 798.66257 | 799.66985 | 17.757 | 2 |
| SM(d18:1_16:1) | C39 H77 N2 O | 1.87 | 700.55324 | 701.56051 | 11.789 | 2 |
| SM(d17:1_21:1) | C43 H85 N2 O | 1.46 | 756.61563 | 757.62291 | 15.443 | 2 |
| SM(d16:1_18:0) | C39 H79 N2 O | 1.41 | 702.56857 | 703.57585 | 13.56 | 2 |

| | | | | | | |
|---|---|---|---|---|---|---|
| SiE(18:3) | C47 H78 O2 | 2.98 | 674.60219 | 675.60794 | 19.178 | 2 |
| PI(18:1_20:4) | C47 H81 O13 | 0.4 | 884.54183 | 902.57568 | 11.183 | 2 |
| PI(18:1_18:2) | C45 H81 O13 | 0.23 | 860.54168 | 878.57559 | 11.526 | 2 |
| PI(18:0_22:6) | C49 H83 O13 | 0.99 | 910.55803 | 928.59183 | 12.144 | 2 |
| PI(18:0_20:5) | C47 H81 O13 | 0.87 | 884.54225 | 885.54946 | 9.896 | 2 |
| PI(18:0_20:4) | C47 H83 O13 | 0.33 | 886.55743 | 904.59116 | 12.728 | 2 |
| PI(18:0_20:3) | C47 H85 O13 | 0.23 | 888.57298 | 906.60675 | 13.317 | 2 |
| PI(18:0_20:2) | C47 H87 O13 | 0.7 | 890.58906 | 908.62294 | 14.06 | 2 |
| PI(18:0_18:2) | C45 H83 O13 | 0.08 | 862.5572 | 863.56444 | 12.873 | 2 |
| PI(18:0_18:1) | C45 H85 O13 | 0.58 | 864.57328 | 882.60713 | 13.771 | 2 |
| PI(16:0_22:6) | C47 H79 O13 | 1.23 | 882.52692 | 900.56104 | 10.721 | 2 |
| PI(16:0_20:4) | C45 H79 O13 | 1.09 | 858.52676 | 876.56067 | 10.985 | 2 |
| PI(16:0_18:2) | C43 H79 O13 | 1.33 | 834.52694 | 852.56077 | 11.11 | 2 |
| PG(36:2) | C42 H79 O10 | 1 | 774.54186 | 792.57593 | 13.374 | 1 |
| PG(20:4D8_18:1) | C44 H69 D8 O | -1.95 | 804.57408 | 805.58136 | 16.285 | 1 |
| PG(16:0_17:0) | C39 H77 O10 | 3.78 | 736.52822 | 737.5339 | 12.926 | 2 |
| PE(62:1) | C67 H132 N O | 3.02 | 1109.97241 | 1110.97969 | 21.79 | 1 |
| PE(46:1D4) | C51 H96 D4 N | -6.78 | 889.73773 | 890.74504 | 18.38 | 2 |
| PE(40:5p) | C45 H80 N O7 | 1.03 | 777.56804 | 778.57532 | 16.119 | 2 |
| PE(37:5e) | C42 H76 N O7 | 1.15 | 737.53679 | 738.54407 | 15.222 | 2 |
| PE(36:2e) | C41 H80 N O7 | 2.96 | 729.5694 | 730.57513 | 15.994 | 1 |
| PE(36:1D3) | C41 H77 D3 N | -1.63 | 748.57977 | 749.58789 | 16.697 | 2 |
| PE(35:1) | C40 H78 N O8 | 1.57 | 731.54766 | 732.55493 | 18.218 | 2 |
| PE(35:0e) | C40 H82 N O7 | 0.97 | 719.58359 | 720.59083 | 17.534 | 2 |
| PE(34:2e) | C39 H76 N O7 | 0.11 | 701.53602 | 702.54357 | 14.865 | 2 |
| PE(33:0e) | C38 H78 N O7 | 4.77 | 691.55489 | 692.56219 | 17.053 | 2 |
| PE(31:1) | C36 H70 N O8 | 4.56 | 675.48699 | 676.49426 | 20.72 | 3 |
| PE(31:0D7) | C36 H65 D7 N | -3.45 | 684.54113 | 748.55774 | 12.833 | 2 |
| PE(22:5e) | C27 H46 N O7 | 1.52 | 527.30199 | 528.30927 | 4.707 | 1 |
| PE(21:4) | C26 H44 N O8 | 1.27 | 529.28113 | 530.2884 | 4.348 | 1 |
| PE(20:4e) | C25 H44 N O7 | 0.49 | 501.28578 | 502.29306 | 4.467 | 2 |
| PE(20:4e_18:1) | C43 H78 N O7 | -1.77 | 751.55026 | 784.5838 | 18.064 | 1 |
| PE(20:3e) | C25 H46 N O7 | 0.77 | 503.30158 | 504.30885 | 4.838 | 2 |
| PE(20:0p_20:4) | C45 H82 N O7 | 1.12 | 779.58376 | 780.59116 | 16.87 | 2 |
| PE(20:0p_18:2) | C43 H82 N O7 | 1.48 | 755.58401 | 756.59083 | 17.32 | 2 |

| | | | | | | |
|---|---|---|---|---|---|---|
| PE(19:1_19:1) | C43 H82 N O8 | 1.77 | 771.57917 | 772.58641 | 16.318 | 1 |
| PE(19:1_18:1) | C42 H80 N O8 | 0.86 | 757.5628 | 758.57008 | 16.065 | 2 |
| PE(18:3e) | C23 H42 N O7 | 1.47 | 475.27059 | 476.27786 | 3.497 | 1 |
| PE(18:2p_18:2) | C41 H74 N O7 | 1.58 | 723.52143 | 724.52871 | 13.721 | 2 |
| PE(18:2e) | C23 H44 N O7 | 1.26 | 477.28614 | 478.29342 | 4.168 | 2 |
| PE(18:2_20:4) | C43 H74 N O8 | 1.58 | 763.51641 | 764.52369 | 12.57 | 2 |
| PE(18:2_18:2) | C41 H74 N O8 | 1.53 | 739.51634 | 740.52356 | 12.78 | 2 |
| PE(18:1p_22:6) | C45 H76 N O7 | -0.56 | 773.53551 | 774.54379 | 14.483 | 2 |
| PE(18:1p_20:5) | C43 H74 N O7 | 1.1 | 747.52112 | 748.52839 | 10.938 | 2 |
| PE(18:1p_18:2) | C41 H76 N O7 | 1.24 | 725.53684 | 726.5442 | 14.818 | 2 |
| PE(18:1e_20:4) | C43 H78 N O7 | 0.55 | 751.552 | 752.55923 | 14.025 | 2 |
| PE(18:1D7_15:0) | C38 H67 D7 N | 0.8 | 710.55971 | 711.56699 | 14.044 | 1 |
| PE(18:1_22:6) | C45 H76 N O8 | 1.46 | 789.532 | 790.53928 | 13.454 | 2 |
| PE(18:1_20:4) | C43 H76 N O8 | 1.21 | 765.53178 | 766.53906 | 13.719 | 2 |
| PE(18:1_18:2) | C41 H76 N O8 | 0.17 | 741.53098 | 742.53675 | 14.43 | 2 |
| PE(18:1_18:1) | C41 H78 N O8 | 1.21 | 743.54741 | 744.55471 | 14.036 | 2 |
| PE(18:0p_22:6) | C45 H78 N O7 | 0.89 | 775.55228 | 776.55955 | 15.67 | 2 |
| PE(18:0p_20:5) | C43 H76 N O7 | 1.16 | 749.53681 | 750.54408 | 12.576 | 2 |
| PE(18:0p_20:3) | C43 H80 N O7 | -0.04 | 753.56721 | 754.57449 | 16.794 | 1 |
| PE(18:0p_18:2) | C41 H78 N O7 | 1.15 | 727.55243 | 728.55973 | 16.201 | 2 |
| PE(18:0_20:5) | C43 H76 N O8 | 1.14 | 765.53173 | 766.53899 | 11.325 | 2 |
| PE(18:0_20:4) | C43 H78 N O8 | 0.86 | 767.54717 | 768.55446 | 15.506 | 2 |
| PE(18:0_18:3) | C41 H76 N O8 | -0.31 | 741.53063 | 742.5379 | 11.048 | 1 |
| PE(18:0_18:2) | C41 H78 N O8 | 1.28 | 743.54746 | 744.55475 | 15.694 | 2 |
| PE(18:0_18:1) | C41 H80 N O8 | 1.77 | 745.56347 | 746.57038 | 16.401 | 2 |
| PE(17:1_19:1) | C41 H78 N O8 | 1.14 | 743.54735 | 744.55463 | 18.039 | 2 |
| PE(17:1_18:1) | C40 H76 N O8 | 0.92 | 729.53153 | 730.5388 | 14.611 | 2 |
| PE(17:1_16:0) | C38 H74 N O8 | 4.28 | 703.51822 | 704.52549 | 13.868 | 2 |
| PE(17:0_20:4) | C42 H76 N O8 | -1.4 | 753.5298 | 754.53708 | 14.577 | 2 |
| PE(16:0p_22:6) | C43 H74 N O7 | 0.68 | 747.5208 | 748.52807 | 14.049 | 2 |
| PE(16:0p_22:5) | C43 H76 N O7 | 0.89 | 749.53661 | 750.54388 | 14.957 | 2 |
| PE(16:0p_22:4) | C43 H78 N O7 | 0.85 | 751.55223 | 752.55952 | 16.018 | 2 |
| PE(16:0p_20:5) | C41 H72 N O7 | 0.99 | 721.50536 | 722.51263 | 10.569 | 2 |
| PE(16:0p_20:4) | C41 H74 N O7 | 0.57 | 723.5207 | 724.52799 | 14.627 | 2 |
| PE(16:0p_18:2) | C39 H74 N O7 | 1.24 | 699.52116 | 700.52844 | 14.6 | 2 |

| | | | | | | |
|---|---|---|---|---|---|---|
| PE(16:0p_18:1) | C39 H76 N O7 | 0.79 | 701.53649 | 702.54381 | 15.769 | 2 |
| PE(16:0e) | C21 H44 N O7 | 1.56 | 453.28624 | 454.29352 | 5.047 | 2 |
| PE(16:0_22:6) | C43 H74 N O8 | 1.15 | 763.51608 | 764.52336 | 13.229 | 2 |
| PE(16:0_20:4) | C41 H74 N O8 | 1.31 | 739.51617 | 740.52348 | 13.844 | 2 |
| PE(16:0_18:2) | C39 H74 N O8 | -0.03 | 715.51518 | 716.52293 | 14.001 | 2 |
| PE(16:0_18:1) | C39 H76 N O8 | 1.33 | 717.53181 | 718.53907 | 14.9 | 2 |
| PC(8:0e_16:0) | C32 H66 N O7 | 1.68 | 607.45871 | 608.46598 | 12.19 | 2 |
| PC(8:0_20:3) | C36 H66 N O8 | 1.32 | 671.45349 | 672.46125 | 7.427 | 1 |
| PC(44:6e) | C52 H94 N O7 | 0.53 | 875.67725 | 876.68453 | 19.211 | 2 |
| PC(44:5e) | C52 H96 N O7 | 1.1 | 877.6934 | 878.70068 | 23.572 | 2 |
| PC(44:5) | C52 H94 N O8 | -0.07 | 891.67165 | 892.67892 | 12.948 | 2 |
| PC(44:4e) | C52 H98 N O7 | 0.57 | 879.70859 | 880.71587 | 25.829 | 2 |
| PC(44:2) | C52 H100 N O | -0.59 | 897.71813 | 898.7254 | 25.856 | 2 |
| PC(42:6e) | C50 H90 N O7 | 0.61 | 847.64601 | 848.65318 | 17.337 | 2 |
| PC(42:5e) | C50 H92 N O7 | 0.39 | 849.66147 | 850.66875 | 18.686 | 2 |
| PC(42:5) | C50 H90 N O8 | -8.2 | 863.63333 | 864.6406 | 11.221 | 2 |
| PC(42:4e) | C50 H94 N O7 | 1.1 | 851.67772 | 852.685 | 23.851 | 2 |
| PC(42:2e) | C50 H98 N O7 | 1.02 | 855.70897 | 856.71624 | 26.552 | 2 |
| PC(42:10) | C50 H80 N O8 | 0.61 | 853.56267 | 854.56995 | 12.223 | 2 |
| PC(40:9) | C48 H78 N O8 | -2.48 | 827.54445 | 828.55173 | 13.633 | 1 |
| PC(40:7) | C48 H82 N O8 | -2.47 | 831.57575 | 832.58303 | 15.795 | 2 |
| PC(40:4) | C48 H88 N O8 | 3.19 | 837.62743 | 838.63448 | 13.268 | 2 |
| PC(40:3e) | C48 H92 N O7 | 2.19 | 825.66295 | 826.67023 | 19.329 | 2 |
| PC(40:2e) | C48 H94 N O7 | 1.14 | 827.67773 | 828.685 | 24.741 | 2 |
| PC(40:1e) | C48 H96 N O7 | 0.92 | 829.69321 | 830.70048 | 25.945 | 2 |
| PC(40:1) | C48 H94 N O8 | -0.32 | 843.67143 | 844.67871 | 23.595 | 2 |
| PC(4:0_14:1) | C26 H50 N O8 | 1.58 | 535.32825 | 536.33552 | 2.863 | 2 |
| PC(38:8e) | C46 H78 N O7 | -2.36 | 787.54973 | 788.55701 | 14.831 | 2 |
| PC(38:7D5) | C46 H73 D5 N | 1.79 | 808.57933 | 809.58661 | 16.103 | 1 |
| PC(38:7) | C46 H78 N O8 | -2.75 | 803.5443 | 804.55157 | 13.148 | 2 |
| PC(38:6D8) | C46 H72 D8 N | 4.33 | 813.61589 | 814.62317 | 16.665 | 1 |
| PC(38:6D3) | C46 H77 D3 N | 0.05 | 808.58103 | 809.5883 | 13.637 | 1 |
| PC(38:5) | C46 H82 N O8 | 0.2 | 807.57796 | 808.58524 | 15.368 | 1 |
| PC(38:4) | C46 H84 N O8 | 0.78 | 809.59409 | 810.60137 | 16.281 | 2 |
| PC(37:6) | C45 H78 N O8 | 1.5 | 791.54769 | 792.55469 | 14.864 | 2 |

| | | | | | | |
|---|---|---|---|---|---|---|
| PC(37:5e) | C45 H82 N O7 | 0.4 | 779.5832 | 780.59048 | 15.847 | 2 |
| PC(37:5) | C45 H80 N O8 | 1.11 | 793.56304 | 794.57031 | 14.993 | 1 |
| PC(37:4e) | C45 H84 N O7 | 0.8 | 781.59917 | 782.60647 | 16.105 | 2 |
| PC(36:5D8) | C44 H70 D8 N | 4.58 | 787.60032 | 788.6076 | 15.968 | 1 |
| PC(36:5) | C44 H78 N O8 | -2.11 | 779.54486 | 780.55213 | 14.139 | 2 |
| PC(36:4D8) | C44 H72 D8 N | 5.16 | 789.61644 | 790.62372 | 17.643 | 3 |
| PC(36:4D3) | C44 H77 D3 N | 1.15 | 784.58189 | 785.58916 | 15.062 | 2 |
| PC(36:4) | C44 H80 N O8 | 1.66 | 781.56345 | 782.57073 | 11.258 | 2 |
| PC(36:2) | C44 H84 N O8 | 0.05 | 785.59349 | 786.60077 | 17.892 | 2 |
| PC(36:1) | C44 H86 N O8 | 1.32 | 787.61015 | 788.61742 | 12.548 | 2 |
| PC(35:4e) | C43 H80 N O7 | 0.48 | 753.5676 | 754.57488 | 14.667 | 2 |
| PC(35:4D7) | C43 H71 D7 N | -2.5 | 774.5885 | 775.59578 | 14.706 | 2 |
| PC(35:3e) | C43 H82 N O7 | -0.35 | 755.58263 | 756.5899 | 16.182 | 1 |
| PC(35:2e) | C43 H84 N O7 | 0.22 | 757.59871 | 758.60598 | 17.857 | 2 |
| PC(35:0) | C43 H86 N O8 | 0.65 | 775.60961 | 776.61688 | 18.955 | 1 |
| PC(34:5D8) | C42 H66 D8 N | 4.23 | 759.56863 | 760.57591 | 14.176 | 2 |
| PC(34:3e) | C42 H80 N O7 | -1.82 | 741.56589 | 742.57532 | 14.446 | 2 |
| PC(34:2) | C42 H80 N O8 | 1.9 | 757.5636 | 758.57003 | 10.232 | 2 |
| PC(34:1) | C42 H82 N O8 | 3.76 | 759.58066 | 760.58597 | 10.819 | 2 |
| PC(34:0) | C42 H84 N O8 | 3.55 | 761.59616 | 762.60136 | 14.529 | 2 |
| PC(33:3e) | C41 H78 N O7 | -1.21 | 727.55071 | 728.55798 | 14.418 | 2 |
| PC(33:3) | C41 H76 N O8 | 0.99 | 741.53158 | 742.53886 | 14.199 | 2 |
| PC(33:2e) | C41 H80 N O7 | -2.6 | 729.56534 | 730.57518 | 15.055 | 2 |
| PC(33:1e) | C41 H82 N O7 | 1.18 | 731.58376 | 732.59103 | 15.975 | 2 |
| PC(33:1D9) | C41 H71 D9 N | -4.24 | 754.61545 | 755.62272 | 15.005 | 1 |
| PC(33:1D7) | C41 H73 D7 N | -2.23 | 752.60441 | 753.61336 | 15.313 | 2 |
| PC(33:0) | C41 H82 N O8 | -0.05 | 747.57777 | 748.58504 | 16.831 | 2 |
| PC(32:4D8) | C40 H64 D8 N | 5.01 | 733.55345 | 734.56072 | 14.245 | 1 |
| PC(32:3e) | C40 H76 N O7 | 1.01 | 713.53666 | 714.54394 | 13.343 | 2 |
| PC(32:1D3) | C40 H75 D3 N | 1.95 | 734.56677 | 735.57406 | 15.5 | 2 |
| PC(32:1) | C40 H78 N O8 | 1.84 | 731.54785 | 732.55517 | 15.5 | 2 |
| PC(32:1_6:0) | C46 H90 N O8 | 0.95 | 815.64118 | 816.64845 | 19.172 | 2 |
| PC(31:1e) | C39 H78 N O7 | -4.33 | 703.54855 | 704.55974 | 15.449 | 2 |
| PC(31:1) | C39 H76 N O8 | 3.01 | 717.53302 | 718.5394 | 13.598 | 2 |
| PC(31:0) | C39 H78 N O8 | 4.12 | 719.54947 | 720.56075 | 15.111 | 1 |

| | | | | | | |
|---|---|---|---|---|---|---|
| PC(30:1) | C38 H74 N O8 | 1.44 | 703.51622 | 704.5236 | 12.879 | 2 |
| PC(26:3) | C34 H62 N O8 | 1.93 | 643.42255 | 644.42984 | 6.542 | 2 |
| PC(26:2) | C34 H64 N O8 | 0.53 | 645.4373 | 646.44457 | 8.126 | 2 |
| PC(26:1) | C34 H66 N O8 | 0.95 | 647.45322 | 648.4605 | 9.327 | 2 |
| PC(26:1_16:0) | C50 H98 N O8 | -0.79 | 871.70232 | 872.70959 | 26.109 | 3 |
| PC(24:5e) | C32 H56 N O7 | 0.62 | 597.37981 | 598.38708 | 6.254 | 1 |
| PC(24:0_20:4) | C52 H96 N O8 | 0.41 | 893.68772 | 894.695 | 24.277 | 2 |
| PC(24:0_18:2) | C50 H96 N O8 | 0.76 | 869.68802 | 870.69528 | 24.948 | 2 |
| PC(23:1) | C31 H60 N O8 | 5.43 | 605.40894 | 606.41622 | 6.286 | 1 |
| PC(22:5_18:2) | C48 H82 N O8 | 1.42 | 831.57898 | 832.58626 | 13.711 | 2 |
| PC(22:5_14:1) | C44 H76 N O8 | -1.95 | 777.52934 | 778.53644 | 13.128 | 2 |
| PC(22:4) | C30 H52 N O8 | 1.5 | 585.34393 | 586.35121 | 2.991 | 2 |
| PC(22:4_20:4) | C50 H84 N O8 | 0.7 | 857.59406 | 858.60134 | 13.838 | 2 |
| PC(22:3) | C30 H54 N O8 | 1.45 | 587.35956 | 588.36684 | 3.164 | 2 |
| PC(22:1_20:4) | C50 H90 N O8 | 0.59 | 863.64091 | 864.64819 | 17.316 | 2 |
| PC(21:4) | C29 H50 N O8 | 1.16 | 571.32806 | 572.33534 | 4.384 | 2 |
| PC(21:1) | C29 H56 N O8 | 1.58 | 577.37526 | 578.38254 | 5.85 | 1 |
| PC(21:0) | C29 H58 N O8 | 1.2 | 579.3907 | 580.3979 | 6.159 | 2 |
| PC(20:5_18:2) | C46 H78 N O8 | -2.81 | 803.54424 | 804.55152 | 14.041 | 2 |
| PC(20:4) | C28 H48 N O8 | -0.45 | 557.3115 | 558.31878 | 2.389 | 2 |
| PC(20:4_20:4) | C48 H80 N O8 | 1.7 | 829.56356 | 830.57084 | 12.545 | 2 |
| PC(20:3e_18:1) | C46 H86 N O7 | -0.77 | 795.61357 | 796.62085 | 17.339 | 2 |
| PC(20:3) | C28 H50 N O8 | 1.15 | 559.32805 | 560.33565 | 2.257 | 2 |
| PC(20:3_22:6) | C50 H82 N O8 | 0.74 | 855.57843 | 856.58571 | 13.292 | 2 |
| PC(20:3_18:2) | C46 H82 N O8 | 0.75 | 807.57841 | 808.5857 | 13.707 | 2 |
| PC(20:2) | C28 H52 N O8 | 1.92 | 561.34413 | 562.35107 | 2.514 | 1 |
| PC(20:2_18:2) | C46 H84 N O8 | 1.03 | 809.59429 | 810.60156 | 15.058 | 2 |
| PC(20:1) | C28 H54 N O8 | 1.93 | 563.35979 | 564.36678 | 3.098 | 2 |
| PC(20:0e_20:4) | C48 H90 N O7 | 0.8 | 823.64615 | 824.65341 | 18.812 | 2 |
| PC(20:0e_18:2) | C46 H90 N O7 | 0.42 | 799.64583 | 800.6531 | 19.702 | 2 |
| PC(20:0e_18:1) | C46 H92 N O7 | 0.05 | 801.66118 | 802.66846 | 23.326 | 2 |
| PC(20:0_18:2) | C46 H88 N O8 | 1.74 | 813.62617 | 814.63339 | 16.271 | 2 |
| PC(19:3e) | C27 H50 N O7 | -0.09 | 531.33244 | 532.33936 | 5.713 | 2 |
| PC(19:2) | C27 H50 N O8 | 1.22 | 547.32807 | 548.33535 | 4.51 | 2 |
| PC(19:1) | C27 H52 N O8 | 0.65 | 549.34341 | 550.35069 | 6.001 | 2 |

| | | | | | | |
|---|---|---|---|---|---|---|
| PC(19:1_19:1) | C46 H88 N O8 | 0.57 | 813.62522 | 814.6325 | 17.331 | 2 |
| PC(19:1_18:1) | C45 H86 N O8 | 0.93 | 799.60985 | 800.61713 | 19.688 | 2 |
| PC(19:0_20:4) | C47 H86 N O8 | 0.53 | 823.60954 | 824.61681 | 16.27 | 2 |
| PC(19:0_20:3) | C47 H88 N O8 | 0.16 | 825.62488 | 826.63217 | 17.387 | 2 |
| PC(19:0_18:1) | C45 H88 N O8 | 0.84 | 801.62543 | 802.6327 | 18.147 | 2 |
| PC(18:3e_24:0) | C50 H96 N O7 | 0.5 | 853.69287 | 854.70014 | 24.578 | 2 |
| PC(18:3e_20:4) | C46 H80 N O7 | 1.21 | 789.56819 | 790.57547 | 13.667 | 2 |
| PC(18:3e_18:2) | C44 H80 N O7 | 0.65 | 765.56774 | 766.57501 | 13.903 | 2 |
| PC(18:3_18:2) | C44 H78 N O8 | 1.55 | 779.54771 | 780.55499 | 12.055 | 2 |
| PC(18:2e_20:4) | C46 H82 N O7 | 1.57 | 791.58413 | 792.59141 | 14.03 | 2 |
| PC(18:2e_18:2) | C44 H82 N O7 | 0.58 | 767.58334 | 768.59061 | 14.382 | 2 |
| PC(18:2) | C26 H48 N O8 | 1.3 | 533.31245 | 534.31971 | 2.399 | 2 |
| PC(18:2_23:0) | C49 H94 N O8 | 0.47 | 855.67211 | 856.67938 | 23.683 | 1 |
| PC(18:2_20:4) | C46 H80 N O8 | 1.21 | 805.56313 | 806.57041 | 12.887 | 2 |
| PC(18:2_18:2) | C44 H80 N O8 | 1.03 | 781.56296 | 782.57024 | 14.292 | 2 |
| PC(18:1e_22:6) | C48 H84 N O7 | 0.97 | 817.59933 | 818.60661 | 16.23 | 2 |
| PC(18:1e_22:4) | C48 H88 N O7 | 0.82 | 821.63052 | 822.63779 | 16.996 | 2 |
| PC(18:1e_20:4) | C46 H84 N O7 | 0.4 | 793.59886 | 794.60613 | 16.142 | 2 |
| PC(18:1e_18:2) | C44 H84 N O7 | 1.66 | 769.59982 | 770.6071 | 16.937 | 2 |
| PC(18:1e_18:1) | C44 H86 N O7 | 0.7 | 771.61473 | 772.62199 | 18.177 | 2 |
| PC(18:1D7_15:0) | C41 H73 D7 N | 0.68 | 752.6066 | 753.61388 | 14.664 | 2 |
| PC(18:1) | C26 H50 N O8 | 1.9 | 535.32842 | 518.32513 | 2.065 | 1 |
| PC(18:1_24:0) | C50 H98 N O8 | -1.49 | 871.70171 | 872.70898 | 25.405 | 2 |
| PC(18:1_22:0) | C48 H94 N O8 | 0.34 | 843.67199 | 844.67927 | 24.331 | 2 |
| PC(18:1_20:4) | C46 H82 N O8 | -2.2 | 807.57603 | 808.5833 | 16.127 | 2 |
| PC(18:1_18:3) | C44 H80 N O8 | -9.41 | 781.5548 | 782.56217 | 9.551 | 1 |
| PC(18:1_18:2) | C44 H82 N O8 | 0.9 | 783.57851 | 784.58579 | 14.371 | 2 |
| PC(18:1_18:1) | C44 H84 N O8 | -0.19 | 785.5933 | 786.60058 | 16.494 | 2 |
| PC(18:0e_22:6) | C48 H86 N O7 | 1.68 | 819.61556 | 820.62284 | 16.606 | 2 |
| PC(18:0e_20:3) | C46 H88 N O7 | 1.24 | 797.63083 | 798.6381 | 17.774 | 2 |
| PC(18:0e_18:2) | C44 H86 N O7 | 0.85 | 771.61485 | 772.62213 | 17.256 | 2 |
| PC(18:0_8:0) | C34 H68 N O8 | 0.69 | 649.4687 | 650.47598 | 10.548 | 2 |
| PC(18:0_22:6) | C48 H84 N O8 | 0.95 | 833.59424 | 834.60153 | 14.849 | 2 |
| PC(18:0_22:5) | C48 H86 N O8 | 0.61 | 835.60961 | 836.6169 | 16.558 | 2 |
| PC(18:0_22:4) | C48 H88 N O8 | 0.57 | 837.62523 | 838.63251 | 17.449 | 2 |

| | | | | | | |
|---|---|---|---|---|---|---|
| PC(18:0_20:4) | C46 H84 N O8 | 1.6 | 809.59475 | 810.60203 | 15.207 | 2 |
| PC(18:0_20:3) | C46 H86 N O8 | 0.51 | 811.60952 | 812.61679 | 17.321 | 2 |
| PC(18:0_18:3) | C44 H82 N O8 | 0.51 | 783.5782 | 784.58547 | 15.358 | 2 |
| PC(18:0_18:2) | C44 H84 N O8 | 0.8 | 785.59408 | 786.60136 | 16.715 | 2 |
| PC(18:0_18:1) | C44 H86 N O8 | -0.14 | 787.609 | 788.61627 | 18.006 | 2 |
| PC(18:0_18:0) | C44 H88 N O8 | 0.63 | 789.62525 | 790.63253 | 19.181 | 3 |
| PC(18:0_16:0) | C42 H84 N O8 | 0.38 | 761.59374 | 762.60101 | 18.053 | 2 |
| PC(17:1_22:6) | C47 H80 N O8 | 1.68 | 817.56353 | 818.57104 | 12.942 | 2 |
| PC(17:1_20:4) | C45 H80 N O8 | 0.73 | 793.56273 | 794.57011 | 13.984 | 2 |
| PC(17:1_18:2) | C43 H80 N O8 | 1.12 | 769.56302 | 770.57032 | 13.724 | 2 |
| PC(17:0_22:6) | C47 H82 N O8 | 0.89 | 819.57853 | 820.58581 | 14.418 | 2 |
| PC(17:0_22:5) | C47 H84 N O8 | 1.05 | 821.59432 | 822.60161 | 15.006 | 2 |
| PC(17:0_20:4) | C45 H82 N O8 | 1.05 | 795.57864 | 796.58591 | 14.855 | 2 |
| PC(17:0_20:3) | C45 H84 N O8 | 0.68 | 797.59399 | 798.60127 | 16.194 | 2 |
| PC(17:0_18:3) | C43 H80 N O8 | 0.94 | 769.56288 | 770.57018 | 14.089 | 2 |
| PC(17:0_18:2) | C43 H82 N O8 | 1.25 | 771.57877 | 772.58571 | 15.781 | 1 |
| PC(17:0_18:1) | C43 H84 N O8 | 1.1 | 773.5943 | 774.60158 | 16.46 | 2 |
| PC(16:1e_22:6) | C46 H80 N O7 | 0.89 | 789.56794 | 790.57522 | 14.299 | 2 |
| PC(16:1e_22:5) | C46 H82 N O7 | 0.41 | 791.58322 | 792.59049 | 16.025 | 1 |
| PC(16:1e_22:4) | C46 H84 N O7 | 0.41 | 793.59886 | 794.60614 | 15.933 | 2 |
| PC(16:1e_20:5) | C44 H78 N O7 | 1.39 | 763.55265 | 764.55993 | 13.636 | 2 |
| PC(16:1e_20:4) | C44 H80 N O7 | 1.13 | 765.5681 | 766.57538 | 14.762 | 2 |
| PC(16:1e_20:3) | C44 H82 N O7 | 0.97 | 767.58364 | 768.59091 | 16.114 | 2 |
| PC(16:1e_18:3) | C42 H78 N O7 | 0.21 | 739.55174 | 740.55902 | 14.087 | 2 |
| PC(16:1e_18:2) | C42 H80 N O7 | 1.24 | 741.56816 | 742.57545 | 15.54 | 2 |
| PC(16:1e_18:1) | C42 H82 N O7 | -0.52 | 743.58251 | 744.58978 | 16.096 | 1 |
| PC(16:1e_16:1) | C40 H78 N O7 | 0.83 | 715.55218 | 716.55947 | 14.624 | 2 |
| PC(16:1e_16:0) | C40 H80 N O7 | 0.7 | 717.56775 | 718.5751 | 16.226 | 2 |
| PC(16:1_22:6) | C46 H78 N O8 | 1.24 | 803.5475 | 804.55478 | 12.2 | 2 |
| PC(16:1_20:3) | C44 H80 N O8 | -2.43 | 781.56025 | 782.56753 | 15.639 | 2 |
| PC(16:1_18:2) | C42 H78 N O8 | 0.41 | 755.54682 | 756.55409 | 13.909 | 1 |
| PC(16:1_18:1) | C42 H80 N O8 | 0.64 | 757.56264 | 758.56992 | 14.551 | 2 |
| PC(16:0e_20:4) | C44 H82 N O7 | 0.92 | 767.58359 | 768.59087 | 15.535 | 2 |
| PC(16:0e_20:3) | C44 H84 N O7 | 1.06 | 769.59936 | 770.60663 | 15.797 | 2 |
| PC(16:0e_18:2) | C42 H82 N O7 | 1.5 | 743.584 | 744.59129 | 15.495 | 2 |

| | | | | | | |
|---|---|---|---|---|---|---|
| PC(16:0e_18:1) | C42 H84 N O7 | 1.85 | 745.59992 | 746.60719 | 16.792 | 2 |
| PC(16:0e_16:0) | C40 H82 N O7 | 1.33 | 719.58385 | 720.59111 | 16.605 | 2 |
| PC(16:0e_14:0) | C38 H78 N O7 | 0.78 | 691.55213 | 692.5594 | 14.71 | 2 |
| PC(16:0D9_22:6) | C46 H71 D9 N | 0.97 | 814.61943 | 815.62671 | 16.95 | 1 |
| PC(16:0_8:0) | C32 H64 N O8 | 2.96 | 621.43879 | 644.42884 | 8.672 | 1 |
| PC(16:0_22:6) | C46 H80 N O8 | -2.29 | 805.56031 | 806.56759 | 14.675 | 2 |
| PC(16:0_22:5) | C46 H82 N O8 | 1.25 | 807.57882 | 808.58609 | 14.572 | 2 |
| PC(16:0_20:5) | C44 H78 N O8 | 1.06 | 779.54733 | 780.55461 | 12.462 | 2 |
| PC(16:0_19:0) | C43 H86 N O8 | 2.76 | 775.61125 | 776.61867 | 18.468 | 1 |
| PC(16:0_18:3) | C42 H78 N O8 | 1.15 | 755.54737 | 756.55465 | 13.265 | 2 |
| PC(16:0_18:2) | C42 H80 N O8 | 0.49 | 757.56253 | 758.5698 | 15.148 | 2 |
| PC(16:0_18:1) | C42 H82 N O8 | 0.78 | 759.57839 | 760.58567 | 16.113 | 2 |
| PC(16:0_17:0) | C41 H82 N O8 | 0.28 | 747.57802 | 748.5853 | 16.272 | 2 |
| PC(16:0_16:1) | C40 H78 N O8 | 1.13 | 731.54733 | 732.55461 | 14.337 | 2 |
| PC(16:0_16:0) | C40 H80 N O8 | 0.93 | 733.56284 | 734.57012 | 15.338 | 2 |
| PC(16:0_14:1) | C38 H74 N O8 | 1.59 | 703.51632 | 704.5236 | 12.023 | 2 |
| PC(16:0_14:0) | C38 H76 N O8 | 1.29 | 705.53177 | 706.53905 | 13.489 | 2 |
| PC(16:0_13:0) | C37 H74 N O8 | 1.32 | 691.51612 | 692.52339 | 12.597 | 2 |
| PC(15:0_22:6) | C45 H78 N O8 | 0.85 | 791.54718 | 792.5545 | 12.484 | 2 |
| PC(15:0_22:5) | C45 H80 N O8 | 1.36 | 793.56323 | 794.57052 | 13.708 | 2 |
| PC(15:0_20:5) | C43 H76 N O8 | -4.6 | 765.52733 | 766.53461 | 13.219 | 1 |
| PC(15:0_20:4) | C43 H78 N O8 | 1.58 | 767.54771 | 768.55521 | 12.561 | 2 |
| PC(15:0_18:3) | C41 H76 N O8 | 1.47 | 741.53195 | 742.53922 | 12.234 | 2 |
| PC(15:0_18:2) | C41 H78 N O8 | 1.49 | 743.54761 | 744.55491 | 13.349 | 2 |
| PC(15:0_18:1) | C41 H80 N O8 | 1.3 | 745.56312 | 746.5704 | 14.64 | 2 |
| PC(15:0_16:1) | C39 H76 N O8 | 1.06 | 717.53162 | 718.53892 | 12.889 | 2 |
| PC(15:0_16:0) | C39 H78 N O8 | 1.25 | 719.5474 | 720.55468 | 14.731 | 2 |
| PC(14:0e_18:2) | C40 H78 N O7 | 0.63 | 715.55204 | 716.55967 | 13.726 | 2 |
| PC(14:0_22:6) | C44 H76 N O8 | 1.21 | 777.5318 | 778.53906 | 11.865 | 2 |
| PC(14:0_20:5) | C42 H74 N O8 | 0.86 | 751.51585 | 752.52313 | 11.166 | 2 |
| PC(14:0_20:4) | C42 H76 N O8 | 1.51 | 753.53199 | 754.53926 | 12.212 | 2 |
| PC(14:0_18:3) | C40 H74 N O8 | 1.32 | 727.51616 | 728.52344 | 11.376 | 2 |
| PC(14:0_18:2) | C40 H76 N O8 | 1.32 | 729.53182 | 730.5391 | 12.446 | 2 |
| PC(14:0_14:0) | C36 H72 N O8 | 1.38 | 677.50049 | 678.50777 | 11.757 | 2 |
| PC(12:0_18:2) | C38 H72 N O8 | 1.42 | 701.50055 | 702.50782 | 10.81 | 2 |

| | | | | | | |
|---|---|---|---|---|---|---|
| PC(11:0_23:1) | C42 H82 N O8 | -5.17 | 759.57388 | 760.58567 | 19.081 | 1 |
| MGDG(19:1_19:1) | C47 H86 O10 | 5.05 | 810.62619 | 811.63347 | 18.329 | 1 |
| MGDG(18:2_18:2) | C45 H78 O10 | -9.41 | 778.55217 | 779.55945 | 13.183 | 1 |
| MGDG(16:1_15:0) | C40 H74 O10 | -9.76 | 714.52123 | 715.5285 | 13.537 | 2 |
| MGDG(16:0_16:1) | C41 H76 O10 | 2.5 | 728.54567 | 1480.0791 | 13.998 | 2 |
| MGDG(16:0_15:0) | C40 H76 O10 | -7.51 | 716.53847 | 717.54574 | 14.588 | 2 |
| MG(20:4e) | C23 H40 O3 | 0.97 | 364.2981 | 365.3054 | 8.251 | 2 |
| MG(18:3) | C21 H36 O4 | 1.62 | 352.26193 | 353.26931 | 3.23 | 2 |
| MG(18:1) | C21 H40 O4 | 1.67 | 356.29325 | 357.30027 | 6.393 | 2 |
| MG(18:0) | C21 H42 O4 | 0.98 | 358.30866 | 359.31586 | 1.004 | 2 |
| MG(16:2e) | C19 H36 O3 | 0.76 | 312.26668 | 313.27395 | 5.525 | 2 |
| MePC(36:4) | C45 H82 N O8 | 0.66 | 795.57833 | 796.5856 | 15.246 | 2 |
| MePC(35:0) | C44 H88 N O8 | 1.53 | 789.62596 | 790.63324 | 16.88 | 1 |
| MePC(34:4) | C43 H78 N O8 | -1.49 | 767.54536 | 768.55438 | 13.412 | 2 |
| MePC(34:3) | C43 H80 N O8 | 6.89 | 769.56746 | 770.5706 | 14.858 | 1 |
| MePC(33:3) | C42 H78 N O8 | 1.52 | 755.54766 | 756.55493 | 10.785 | 1 |
| MePC(32:1) | C41 H80 N O8 | 1.58 | 745.56334 | 746.57062 | 15.527 | 2 |
| MePC(30:2) | C39 H74 N O8 | 1.76 | 715.51647 | 716.52374 | 14.489 | 1 |
| MePC(29:0) | C38 H76 N O8 | 1.48 | 705.5319 | 706.53918 | 14.357 | 2 |
| MePC(21:5e) | C30 H52 N O7 | 0.65 | 569.34851 | 570.3558 | 5.5 | 2 |
| MePC(16:1e) | C25 H50 N O7 | 1.26 | 507.33313 | 508.34035 | 5.312 | 2 |
| MePC(14:1e) | C23 H46 N O7 | 1.22 | 479.30177 | 480.30905 | 5.082 | 2 |
| MePC(14:0e) | C23 H48 N O7 | 0.81 | 481.31723 | 482.32452 | 7.256 | 2 |
| LPE(22:6) | C27 H44 N O7 | 1.5 | 525.28633 | 526.29358 | 3.981 | 2 |
| LPE(18:0) | C23 H48 N O7 | 2.79 | 481.31818 | 482.3254 | 6.908 | 2 |
| LPC(38:1) | C46 H92 N O7 | 1.21 | 801.66211 | 802.66919 | 23.951 | 2 |
| LPC(32:1) | C40 H80 N O7 | 1.37 | 717.56822 | 718.5755 | 15.626 | 1 |
| LPC(31:0) | C39 H80 N O7 | 0.99 | 705.56794 | 706.57535 | 15.775 | 2 |
| LPC(24:1) | C32 H64 N O7 | 1.4 | 605.44288 | 606.45016 | 10.256 | 2 |
| LPC(22:6) | C30 H50 N O7 | 1.64 | 567.33342 | 568.3407 | 4 | 2 |
| LPC(22:5) | C30 H52 N O7 | 0.59 | 569.34848 | 570.35577 | 5.175 | 2 |
| LPC(22:4) | C30 H54 N O7 | 0.48 | 571.36406 | 572.37134 | 5.211 | 1 |
| LPC(22:2) | C30 H58 N O7 | 1.6 | 575.39601 | 576.40329 | 7.453 | 2 |
| LPC(22:1) | C30 H60 N O7 | 1.99 | 577.41189 | 578.41917 | 8.173 | 1 |
| LPC(22:0) | C30 H62 N O7 | 1.34 | 579.42717 | 580.43444 | 10.221 | 2 |

| | | | | | | |
|---|---|---|---|---|---|---|
| LPC(20:5) | C28 H48 N O7 | -2.42 | 541.31553 | 542.3224 | 4.995 | 2 |
| LPC(20:4e) | C28 H52 N O6 | -3.63 | 529.3513 | 530.35858 | 7.337 | 1 |
| LPC(20:4) | C28 H50 N O7 | 1.11 | 543.33309 | 544.34038 | 4.778 | 2 |
| LPC(20:3e) | C28 H54 N O6 | -2.63 | 531.36748 | 532.37476 | 8.273 | 1 |
| LPC(20:3) | C28 H52 N O7 | -2.7 | 545.34666 | 546.35394 | 7.438 | 2 |
| LPC(20:2e) | C28 H56 N O6 | 1.31 | 533.38523 | 534.3925 | 6.996 | 1 |
| LPC(20:2) | C28 H54 N O7 | 1.26 | 547.36448 | 548.37172 | 5.575 | 2 |
| LPC(20:1e) | C28 H58 N O6 | 1.63 | 535.40105 | 536.40833 | 9.575 | 1 |
| LPC(20:1) | C28 H56 N O7 | 1.65 | 549.38035 | 550.38764 | 7.017 | 2 |
| LPC(20:0e) | C28 H60 N O6 | 1.31 | 537.41653 | 538.4238 | 9.531 | 2 |
| LPC(20:0) | C28 H58 N O7 | 2.28 | 551.39635 | 552.40363 | 7.444 | 1 |
| LPC(19:1) | C27 H54 N O7 | 1.42 | 535.36455 | 536.37183 | 6.615 | 2 |
| LPC(19:0) | C27 H56 N O7 | 1.02 | 537.37999 | 538.38726 | 7.997 | 2 |
| LPC(18:4) | C26 H46 N O7 | 1.97 | 515.30221 | 516.30948 | 3.257 | 2 |
| LPC(18:3e) | C26 H50 N O6 | 1.63 | 503.3384 | 504.34567 | 5.049 | 2 |
| LPC(18:3) | C26 H48 N O7 | 1.26 | 517.31749 | 518.32477 | 2.083 | 1 |
| LPC(18:2e) | C26 H52 N O6 | 0.07 | 505.35326 | 506.36053 | 5.298 | 2 |
| LPC(18:2) | C26 H50 N O7 | 1.17 | 519.3331 | 520.34037 | 5.066 | 2 |
| LPC(18:1e) | C26 H54 N O6 | 1.8 | 507.36979 | 508.37706 | 6.573 | 2 |
| LPC(18:1D7) | C26 H45 D7 N | 0.76 | 528.39248 | 529.39976 | 5.639 | 2 |
| LPC(18:1) | C26 H52 N O7 | 1.6 | 521.34897 | 522.35625 | 5.734 | 2 |
| LPC(18:0e) | C26 H56 N O6 | 1.47 | 509.38527 | 510.39257 | 7.734 | 2 |
| LPC(18:0) | C26 H54 N O7 | 0.95 | 523.36429 | 524.37157 | 7.529 | 2 |
| LPC(17:1) | C25 H50 N O7 | 1.23 | 507.33311 | 508.34039 | 4.452 | 2 |
| LPC(17:0) | C25 H52 N O7 | 1.27 | 509.34878 | 510.35612 | 6.552 | 2 |
| LPC(16:2e) | C24 H48 N O6 | 6.67 | 477.32511 | 478.33239 | 5.152 | 1 |
| LPC(16:1e) | C24 H50 N O6 | 1.92 | 479.33849 | 480.34577 | 5.832 | 2 |
| LPC(16:1) | C24 H48 N O7 | 1.1 | 493.31738 | 494.32466 | 4.76 | 2 |
| LPC(16:0e) | C24 H52 N O6 | 1.33 | 481.35387 | 482.36114 | 6.608 | 2 |
| LPC(16:0) | C24 H50 N O7 | 1.43 | 495.3332 | 496.34048 | 5.643 | 2 |
| LPC(15:0) | C23 H48 N O7 | 2.01 | 481.31781 | 482.32509 | 4.816 | 2 |
| LPC(14:0) | C22 H46 N O7 | 2.07 | 467.30216 | 468.30944 | 3.6 | 2 |
| LPC(12:0) | C20 H42 N O7 | 2.13 | 439.27083 | 440.278 | 2.806 | 2 |
| LBPA(18:1_18:2) | C42 H77 O10 | 0.43 | 772.52576 | 790.55936 | 11.766 | 2 |
| LBPA(18:0_18:1) | C42 H81 O10 | 0.42 | 776.55706 | 777.56402 | 14.141 | 2 |

| | | | | | | |
|---|---|---|---|---|---|---|
| LBPA(16:0_18:2) | C40 H75 O10 | 0.69 | 746.5103 | 747.51778 | 11.536 | 2 |
| LBPA(16:0_18:1) | C40 H77 O10 | 1.57 | 748.52661 | 766.56055 | 16.303 | 2 |
| LBPA(16:0_18:0) | C40 H79 O10 | 0.52 | 750.54147 | 768.57519 | 13.853 | 2 |
| LBPA(16:0_16:1) | C38 H73 O10 | 0.88 | 720.49477 | 721.50228 | 11.181 | 2 |
| LBPA(16:0_16:0) | C38 H75 O10 | 1.76 | 722.51105 | 740.54519 | 12.454 | 2 |
| LBPA(14:0_16:0) | C36 H71 O10 | 1.5 | 694.47953 | 712.51343 | 10.923 | 2 |
| Hex2Cer(d18:1_16:0) | C46 H87 N O1 | 1.85 | 861.61933 | 862.62661 | 12.2 | 2 |
| Hex1Cer(m17:1_24:6) | C47 H79 N O7 | 6.14 | 769.59038 | 770.59766 | 15.736 | 1 |
| Hex1Cer(d18:1_24:1) | C48 H91 N O8 | 0.44 | 809.67482 | 810.68262 | 17.456 | 2 |
| Hex1Cer(d18:1_16:0) | C40 H77 N O8 | 0.87 | 699.56553 | 700.57277 | 13.38 | 2 |
| DG(8:0_12:2) | C23 H40 O5 | 0.84 | 396.28791 | 397.29517 | 19.317 | 2 |
| DG(38:5e) | C41 H72 O4 | 1.56 | 628.54404 | 629.55132 | 13.029 | 2 |
| DG(38:2e) | C41 H78 O4 | 2.41 | 634.59154 | 635.59795 | 19.03 | 2 |
| DG(36:6e) | C39 H66 O4 | 1.1 | 598.49677 | 599.50402 | 12.379 | 2 |
| DG(36:6) | C39 H64 O5 | 1.66 | 612.47639 | 613.48367 | 7.547 | 1 |
| DG(36:5e) | C39 H68 O4 | 1.88 | 600.51289 | 601.52016 | 20.063 | 2 |
| DG(36:5) | C39 H66 O5 | -2.64 | 614.4894 | 615.49668 | 16.886 | 2 |
| DG(36:4e) | C39 H70 O4 | 1.14 | 602.5281 | 603.53538 | 12.842 | 2 |
| DG(36:3e) | C39 H72 O4 | 2.22 | 604.5444 | 605.55168 | 13.876 | 1 |
| DG(36:2e) | C39 H74 O4 | 0.99 | 606.55931 | 607.56659 | 22.2 | 2 |
| DG(35:2e) | C38 H72 O4 | 0.85 | 592.54356 | 593.55084 | 18.404 | 2 |
| DG(34:4e) | C37 H66 O4 | 1.75 | 574.49712 | 575.50439 | 20.046 | 2 |
| DG(34:3e) | C37 H68 O4 | 1.06 | 576.51237 | 577.5195 | 17.977 | 1 |
| DG(34:2e) | C37 H70 O4 | 0.58 | 578.52775 | 579.53502 | 19.319 | 2 |
| DG(33:2e) | C36 H68 O4 | 0.72 | 564.51216 | 565.51944 | 13.079 | 2 |
| DG(32:3e) | C35 H64 O4 | 0.52 | 548.48074 | 549.48802 | 11.182 | 2 |
| DG(32:2e) | C35 H66 O4 | 0.98 | 550.49665 | 551.50393 | 18.343 | 2 |
| DG(31:2e) | C34 H64 O4 | 0.12 | 536.48053 | 537.48835 | 11.557 | 2 |
| DG(30:2e) | C33 H62 O4 | 1.05 | 522.46536 | 523.47264 | 10.974 | 2 |
| DG(28:7e) | C31 H48 O4 | -0.73 | 484.35491 | 485.36218 | 10.94 | 1 |
| DG(28:5) | C31 H50 O5 | -3.34 | 502.36415 | 503.37115 | 8.219 | 2 |
| DG(26:4e) | C29 H50 O4 | 0.38 | 462.37108 | 463.37836 | 8.003 | 2 |
| DG(25:2) | C28 H50 O5 | 1.39 | 466.36647 | 467.37375 | 8.508 | 2 |
| DG(23:3e) | C26 H46 O4 | -3.4 | 422.33817 | 423.34524 | 8.325 | 2 |
| DG(23:2) | C26 H46 O5 | -4.14 | 438.33271 | 439.33998 | 9.174 | 2 |

| | | | | | | |
|---|---|---|---|---|---|---|
| DG(21:2e) | C24 H44 O4 | 1.93 | 396.32472 | 397.332 | 12.539 | 2 |
| DG(21:2) | C24 H42 O5 | -4.26 | 410.30148 | 411.30872 | 7.407 | 2 |
| DG(20:3e) | C23 H40 O4 | -5.41 | 380.2906 | 381.29791 | 8.509 | 2 |
| DG(20:1_18:2) | C41 H74 O5 | -2.92 | 646.55174 | 647.55893 | 17.821 | 2 |
| DG(19:3e) | C22 H38 O4 | -5 | 366.27518 | 367.28268 | 7.523 | 2 |
| DG(18:4_18:2) | C39 H64 O5 | 1.11 | 612.47605 | 613.48333 | 10.537 | 2 |
| DG(18:4_18:1) | C39 H66 O5 | 1.46 | 614.49192 | 615.49921 | 11.515 | 2 |
| DG(18:4_16:0) | C37 H64 O5 | 1.1 | 588.47602 | 589.4833 | 11.544 | 2 |
| DG(18:3_18:2) | C39 H66 O5 | 1.88 | 614.49218 | 632.52601 | 15.006 | 3 |
| DG(18:2_18:2) | C39 H68 O5 | -0.91 | 616.50611 | 617.51223 | 18.354 | 2 |
| DG(18:2_16:1) | C37 H66 O5 | 0.95 | 590.49158 | 591.49898 | 16.17 | 3 |
| DG(18:2_16:0) | C37 H68 O5 | 0.75 | 592.50712 | 593.51431 | 14.433 | 2 |
| DG(18:1_20:4) | C41 H70 O5 | 1.99 | 642.52361 | 660.55749 | 16.912 | 3 |
| DG(18:1_20:3) | C41 H72 O5 | 1.71 | 644.53908 | 645.54692 | 17.339 | 3 |
| DG(18:1_18:3) | C39 H68 O5 | 1.68 | 616.50771 | 617.51505 | 12.6 | 2 |
| DG(18:1_18:2) | C39 H70 O5 | 0.79 | 618.52281 | 619.52994 | 14.682 | 2 |
| DG(18:1_18:1) | C39 H72 O5 | 0.91 | 620.53854 | 621.54579 | 15.815 | 2 |
| DG(18:1_18:0) | C39 H74 O5 | 1.16 | 622.55435 | 640.5882 | 23.172 | 3 |
| DG(18:1_16:0) | C37 H70 O5 | 1.38 | 594.52314 | 595.53017 | 18.403 | 2 |
| DG(18:1_14:0) | C35 H66 O5 | 1.01 | 566.4916 | 549.48834 | 14.001 | 2 |
| DG(18:0_22:0) | C43 H84 O5 | 0.51 | 680.63222 | 698.66595 | 24.313 | 3 |
| DG(18:0_20:0) | C41 H80 O5 | 0.7 | 652.60103 | 670.6348 | 19.012 | 3 |
| DG(18:0_18:3) | C39 H70 O5 | 0.84 | 618.52285 | 619.53008 | 14.028 | 2 |
| DG(18:0_18:1) | C39 H74 O5 | 1.11 | 622.55432 | 640.58813 | 19.092 | 2 |
| DG(18:0_18:0) | C39 H76 O5 | 1.22 | 624.57004 | 642.60391 | 19.538 | 2 |
| DG(18:0_17:0) | C38 H74 O5 | 0.87 | 610.55416 | 628.58807 | 22.582 | 3 |
| DG(18:0_16:0) | C37 H72 O5 | 1.7 | 596.53899 | 597.54626 | 18.715 | 1 |
| DG(17:0_18:2) | C38 H70 O5 | 1.1 | 606.52299 | 624.55682 | 17.476 | 3 |
| DG(17:0_18:1) | C38 H72 O5 | 1.36 | 608.5388 | 626.57271 | 17.923 | 3 |
| DG(17:0_16:0) | C36 H70 O5 | 1.35 | 582.52311 | 600.55693 | 17.854 | 3 |
| DG(16:1_18:3) | C37 H64 O5 | 1.24 | 588.4761 | 606.51 | 14.911 | 3 |
| DG(16:0_18:3) | C37 H66 O5 | 1.04 | 590.49164 | 591.49901 | 12.6 | 2 |
| DG(16:0_16:1) | C35 H66 O5 | 0.54 | 566.49133 | 584.5252 | 16.765 | 3 |
| DG(16:0_16:0) | C35 H68 O5 | 1.14 | 568.50732 | 586.54115 | 18.315 | 3 |
| DG(14:0_18:2) | C35 H64 O5 | 1.65 | 564.47631 | 582.51013 | 15.564 | 3 |

| | | | | | | |
|---|---|---|---|---|---|---|
| Co(Q10) | C59 H90 O4 | 1.19 | 862.68494 | 880.71881 | 20.422 | 2 |
| CmE(18:3) | C46 H76 O2 | 1.46 | 660.58549 | 661.59285 | 19.039 | 2 |
| CmE(18:2) | C46 H78 O2 | 0.36 | 662.60042 | 680.63424 | 20.542 | 2 |
| ChE(22:6) | C49 H76 O2 | 8.04 | 696.59014 | 697.59741 | 19.923 | 1 |
| ChE(20:5) | C47 H74 O2 | 1.52 | 670.5699 | 671.57719 | 18.851 | 2 |
| ChE(20:4) | C47 H76 O2 | 1.06 | 672.58524 | 673.59203 | 20.287 | 2 |
| ChE(20:3) | C47 H78 O2 | 0.75 | 674.60069 | 675.60797 | 20.037 | 2 |
| ChE(18:3) | C45 H74 O2 | 1.24 | 646.56969 | 647.57697 | 19.753 | 2 |
| ChE(18:2) | C45 H76 O2 | 1.68 | 648.58562 | 649.59261 | 20.415 | 2 |
| ChE(18:1) | C45 H78 O2 | 1.69 | 650.60128 | 651.60864 | 20.709 | 2 |
| Cer(t42:0) | C42 H85 N O4 | 1.3 | 667.64873 | 668.65601 | 18.438 | 1 |
| Cer(t20:0_26:0+O) | C46 H93 N O5 | 0.83 | 739.70599 | 740.71326 | 19.361 | 3 |
| Cer(t20:0_26:0) | C46 H93 N O4 | 0.74 | 723.711 | 724.71827 | 20.114 | 1 |
| Cer(t20:0_25:0+O) | C45 H91 N O5 | 0.02 | 725.68974 | 726.69702 | 19.346 | 1 |
| Cer(t20:0_24:0+O) | C44 H89 N O5 | -0.32 | 711.67385 | 712.682 | 19.477 | 3 |
| Cer(t18:0_26:0) | C44 H89 N O4 | 1.33 | 695.68009 | 696.6864 | 19.762 | 1 |
| Cer(t17:0_26:0+O) | C43 H87 N O5 | 0.05 | 697.65846 | 698.66554 | 19.042 | 1 |
| Cer(m39:0+O) | C39 H79 N O3 | 0.88 | 609.60653 | 610.61381 | 20.589 | 2 |
| Cer(m36:0) | C36 H73 N O2 | 1 | 551.56468 | 552.57196 | 18.117 | 1 |
| Cer(m34:0) | C34 H69 N O2 | 0.64 | 523.53316 | 524.54056 | 17.444 | 2 |
| Cer(m18:1_22:0) | C40 H79 N O2 | 1.22 | 605.61182 | 606.6191 | 18.312 | 2 |
| Cer(m18:0_18:0) | C36 H73 N O2 | 1.02 | 551.56469 | 552.57202 | 22.455 | 2 |
| Cer(m18:0_16:0) | C34 H69 N O2 | 0.79 | 523.53324 | 524.54053 | 22.003 | 2 |
| Cer(d35:4) | C35 H63 N O3 | 2.04 | 545.48191 | 546.48919 | 13.999 | 2 |
| Cer(d35:1) | C35 H69 N O3 | 0.62 | 551.52808 | 552.53534 | 22.138 | 2 |
| Cer(d34:0) | C34 H69 N O3 | 2.29 | 539.52898 | 540.53625 | 16.635 | 2 |
| Cer(d33:1) | C33 H65 N O3 | 0.35 | 523.49663 | 524.50391 | 21.723 | 2 |
| Cer(d32:0) | C32 H65 N O3 | 0.96 | 511.49693 | 512.50421 | 20.902 | 2 |
| Cer(d30:0) | C30 H61 N O3 | 1.62 | 483.46593 | 484.47321 | 12.933 | 2 |
| Cer(d28:0) | C28 H57 N O3 | 2.77 | 455.43511 | 456.44238 | 11.263 | 2 |
| Cer(d22:0_18:0) | C40 H81 N O3 | 1.72 | 623.62272 | 624.62998 | 19.341 | 2 |
| Cer(d20:2_26:0+O) | C46 H89 N O4 | 1.14 | 719.67998 | 720.68726 | 19.537 | 2 |
| Cer(d20:0_24:0) | C44 H89 N O3 | 1.29 | 679.68512 | 680.6924 | 20.044 | 2 |
| Cer(d19:2_31:0+O) | C50 H97 N O4 | 1.09 | 775.74261 | 776.74991 | 19.342 | 2 |
| Cer(d19:0_24:0) | C43 H87 N O3 | 1.7 | 665.66973 | 666.677 | 19.908 | 2 |

| | | | | | | |
|---|---|---|---|---|---|---|
| Cer(d18:2_28:0) | C46 H89 N O3 | 1.48 | 703.68529 | 704.69257 | 19.734 | 2 |
| Cer(d18:2_26:0+O) | C44 H85 N O4 | 1.79 | 691.6491 | 692.65637 | 19.241 | 2 |
| Cer(d18:2_24:0) | C42 H81 N O3 | 0.86 | 647.6222 | 648.62945 | 19.096 | 2 |
| Cer(d18:2_23:0) | C41 H79 N O3 | 1.3 | 633.60682 | 634.61411 | 17.983 | 2 |
| Cer(d18:2_22:0) | C40 H77 N O3 | -0.68 | 619.58992 | 620.59835 | 17.684 | 2 |
| Cer(d18:2_16:0) | C34 H65 N O3 | 1.37 | 535.49718 | 536.50442 | 13.747 | 2 |
| Cer(d18:1_26:0) | C44 H87 N O3 | 1.95 | 677.66992 | 678.67716 | 19.946 | 2 |
| Cer(d18:1_24:2) | C42 H79 N O3 | 0.84 | 645.60654 | 646.61407 | 17.727 | 2 |
| Cer(d18:1_24:1) | C42 H81 N O3 | -4.07 | 647.61901 | 648.63012 | 17.985 | 3 |
| Cer(d18:1_24:0) | C42 H83 N O3 | 1.34 | 649.63816 | 650.64543 | 18.679 | 2 |
| Cer(d18:1_23:0) | C41 H81 N O3 | 1.54 | 635.62263 | 636.62992 | 18.458 | 2 |
| Cer(d18:1_22:0) | C40 H79 N O3 | 1.46 | 621.60691 | 622.61418 | 18.201 | 2 |
| Cer(d18:1_20:0) | C38 H75 N O3 | 1.48 | 593.57557 | 594.58288 | 17.557 | 2 |
| Cer(d18:1_19:0) | C37 H73 N O3 | 1.46 | 579.55989 | 580.56727 | 17.2 | 1 |
| Cer(d18:1_18:0) | C36 H71 N O3 | 1.1 | 565.54402 | 566.55131 | 16.611 | 2 |
| Cer(d18:1_16:0) | C34 H67 N O3 | 1.39 | 537.51284 | 538.52016 | 15.222 | 2 |
| Cer(d18:0_16:0) | C34 H69 N O3 | 1.32 | 539.52846 | 540.53573 | 15.744 | 2 |
| CarE(20:0) | C27 H53 N O4 | 0.44 | 455.39766 | 456.40494 | 8.16 | 1 |
| CarE(16:1) | C23 H43 N O4 | 1.33 | 397.31974 | 398.32701 | 5.317 | 2 |
| CarE(14:1) | C21 H39 N O4 | 2.08 | 369.28868 | 370.29595 | 3.231 | 3 |
| CarE(14:0+O) | C21 H41 N O5 | 2.17 | 387.29931 | 388.30644 | 3.099 | 2 |
| BisMePA(26:2e) | C31 H59 O7 P | 5.53 | 574.40301 | 597.39185 | 11.136 | 1 |
| BisMeLPA(12:0e) | C17 H37 O6 P | 7.88 | 368.23568 | 369.24295 | 5.237 | 2 |
| BisMeLPA(10:0e) | C15 H33 O6 P | 1.67 | 340.20204 | 341.20932 | 3.358 | 2 |
| AcHexSiE(16:0) | C51 H90 O7 | -0.08 | 814.66859 | 815.67587 | 18.151 | 1 |
| AcHexCmE(18:3) | C52 H86 O7 | -3.02 | 822.63487 | 823.64215 | 17.61 | 1 |
| AcHexCmE(18:1) | C52 H90 O7 | -0.27 | 826.66843 | 844.7026 | 18.378 | 1 |
| AcHexChE(18:2) | C51 H86 O7 | -2.76 | 810.63512 | 811.6424 | 18.491 | 1 |
| AcHexChE(17:1) | C50 H86 O7 | -0.73 | 798.63677 | 799.64304 | 17.936 | 2 |
| AcHexChE(16:1) | C49 H84 O7 | 0.58 | 784.62216 | 785.62872 | 17.845 | 2 |
| AcHexChE(15:0) | C48 H84 O7 | -1.61 | 772.62046 | 790.65631 | 17.676 | 2 |
| AcHexChE(14:1) | C47 H80 O7 | 9.08 | 756.59728 | 757.60455 | 16.737 | 1 |
| AcHexChE(0:0) | C33 H56 O6 | 0.87 | 548.40817 | 549.41544 | 9.38 | 1 |
| AcCa(8:0) | C15 H29 N O4 | 6.26 | 287.21146 | 288.21731 | 1.43 | 2 |
| AcCa(24:0) | C31 H61 N O4 | 1.56 | 511.46086 | 512.46813 | 11.275 | 2 |

| | | | | | | |
|---|---|---|---|---|---|---|
| AcCa(20:4) | C27 H45 N O4 | 1.4 | 447.33548 | 448.34276 | 4.054 | 2 |
| AcCa(20:1) | C27 H51 N O4 | 0.75 | 453.38215 | 454.38941 | 6.089 | 2 |
| AcCa(18:3) | C25 H43 N O4 | 1.27 | 421.31974 | 422.32702 | 3.615 | 2 |
| AcCa(18:2) | C25 H45 N O4 | 2.08 | 423.33574 | 424.34302 | 4.134 | 2 |
| AcCa(18:1) | C25 H47 N O4 | 1.53 | 425.35116 | 426.35844 | 5.023 | 2 |
| AcCa(18:0) | C25 H49 N O4 | 1.6 | 427.36684 | 428.37412 | 6.286 | 3 |
| AcCa(16:0) | C23 H45 N O4 | 1.78 | 399.33557 | 400.34285 | 4.73 | 3 |
| AcCa(14:3) | C21 H35 N O4 | 1.37 | 365.25711 | 366.26443 | 2.46 | 2 |
| AcCa(12:3) | C19 H31 N O4 | 0.19 | 337.22537 | 355.25931 | 1.528 | 1 |
| AcCa(12:1) | C19 H35 N O4 | 1.03 | 341.25696 | 342.26438 | 2.203 | 2 |
| AcCa(12:0) | C19 H37 N O4 | 2.28 | 343.27304 | 344.2803 | 2.583 | 2 |
| AcCa(10:2) | C17 H29 N O4 | 1.25 | 311.21005 | 312.2173 | 1.522 | 2 |
| AcCa(10:1) | C17 H31 N O4 | 1.55 | 313.22579 | 314.23307 | 1.623 | 2 |
| AcCa(10:0) | C17 H33 N O4 | 0.8 | 315.24121 | 316.24845 | 1.691 | 2 |
| | | | 522.35423 | 523.36151 | 7.768 | 1 |

d via Orbitrap-IDX

| Reference Ion | Log2 Fold Change: (Bb_pos_12022024_Newanalysis) / (CM_pos_12022024_Newanalysis) |
|---|---|
| [M+H]+1 | -0.59 |
| [M+H]+1 | 1.04 |
| [M+H]+1 | 0.34 |
| [M+H-H2O]+1 | -5.26 |
| [M+H]+1 | 0.91 |
| [M+H]+1 | -0.12 |
| [M+H]+1 | 0.78 |
| [M+H]+1 | -2.12 |
| [M+H]+1 | 0.36 |
| [M+H]+1 | -0.82 |
| [M+H]+1 | -1.02 |
| [M+H+MeOH]+1 | 0.32 |
| [M+H]+1 | -1.95 |
| [M+H]+1 | -1.38 |
| [M+H]+1 | 0.13 |
| [M+H]+1 | 1.11 |
| [M+H]+1 | 0.45 |
| [M+H]+1 | -4.71 |
| [M+H]+1 | -1.46 |
| [M+NH4]+1 | -1.72 |
| [M+H]+1 | 0.12 |
| [M+H]+1 | -4.28 |
| [M+H]+1 | -4.08 |
| [M+H]+1 | -3.47 |
| [M+Na]+1 | -5.9 |
| [M+H]+1 | -1.42 |
| [M+H]+1 | 0.17 |
| [M+H]+1 | 0.84 |
| [M+H]+1 | 0.64 |
| [M+H]+1 | -0.79 |
| [M+H]+1 | 0.44 |
| [M+H]+1 | 6.52 |
| [M+H]+1 | 1.52 |

| | |
|---|---|
| [M+H]+1 | 0.73 |
| [M+H]+1 | -5.16 |
| [M+H]+1 | 1.97 |
| [M+H]+1 | -2.79 |
| [M+H]+1 | 6.58 |
| [M+H]+1 | 6.53 |
| [M+H]+1 | -6.08 |
| [M+H]+1 | 0.21 |
| [M+H]+1 | 0.42 |
| [M+H]+1 | -3.45 |
| [M+H]+1 | 1.03 |
| [M+H]+1 | 0.46 |
| [M+NH4]+1 | -4.22 |
| [M+NH4]+1 | -4.67 |
| [M+NH4]+1 | -4.54 |
| [M+NH4]+1 | -4.11 |
| [M+NH4]+1 | -2.78 |
| [M+NH4]+1 | -4.74 |
| [M+NH4]+1 | -3.77 |
| [M+NH4]+1 | -5.29 |
| [M+NH4]+1 | -2.74 |
| [M+NH4]+1 | 1.18 |
| [M+H]+1 | -1.13 |
| [M+H]+1 | 1.28 |
| [M+H]+1 | 1.7 |
| [M+H]+1 | -0.61 |
| [M+H]+1 | 1.39 |
| [M+H]+1 | -0.33 |
| [M+H]+1 | 0.7 |
| [M+H]+1 | 1.22 |
| [M+H]+1 | -5.99 |
| [M+H]+1 | -3.41 |
| [M+H]+1 | 0.07 |
| [M+H]+1 | -5.06 |
| [M+NH4]+1 | -2.28 |

| | |
|---|---|
| [M+H]+1 | -0.42 |
| [M+H]+1 | -1.05 |
| [M+H]+1 | -4.8 |
| [M+H]+1 | 0.55 |
| [M+H]+1 | -2.72 |
| [M+NH4]+1 | -3.8 |
| [M+DMSO+H]+1 | -0.36 |
| [M+H]+1 | 0.81 |
| [M+H]+1 | -5.49 |
| [M+H]+1 | 0.28 |
| [M+NH4]+1 | -4.7 |
| [M+H]+1 | -3.45 |
| [M+H]+1 | -4.27 |
| [M+H]+1 | -4.58 |
| [M+H]+1 | 0.26 |
| [M+H]+1 | -2.96 |
| [M+H]+1 | -1.39 |
| [M+H-H2O]+1 | -2.48 |
| [M+H]+1 | 0.23 |
| [M+H]+1 | -6.3 |
| [M+H]+1 | -2.36 |
| [M+H]+1 | -1.18 |
| [M+NH4]+1 | -5.34 |
| [M+NH4]+1 | -4.31 |
| [M+NH4]+1 | -2.39 |
| [M+NH4]+1 | -2.64 |
| [M+H]+1 | -2.49 |
| [M+H]+1 | -2.06 |
| [M+H]+1 | -4.98 |
| [M+H]+1 | -3.74 |
| [M+H]+1 | -0.38 |
| [M+H]+1 | -1.95 |
| [M+H]+1 | -2.05 |
| [M+H]+1 | -1.17 |
| [M+H]+1 | -2.49 |

| | |
|---|---|
| [M+H]+1 | -0.26 |
| [M+H]+1 | -2.29 |
| [M+H]+1 | 0.76 |
| [M+NH4]+1 | -2.56 |
| [M+H]+1 | -0.45 |
| [M+H]+1 | 0.87 |
| [M+NH4]+1 | -0.51 |
| [M+H]+1 | -2 |
| [M+H]+1 | -0.68 |
| [M+H]+1 | -3.86 |
| [M+DMSO+H]+1 | -2.73 |
| [M+H]+1 | 0.52 |
| [M+H]+1 | -2.05 |
| [M+H]+1 | -0.91 |
| [M+H]+1 | -6.43 |
| [M+H]+1 | -4.07 |
| [M+H]+1 | -0.02 |
| [M+H]+1 | 0.76 |
| [M+H+MeOH]+1 | -0.09 |
| [M+NH4]+1 | 0.18 |
| [M+NH4]+1 | 0.8 |
| [M+H]+1 | -0.45 |
| [M+H]+1 | -0.61 |
| [M+H]+1 | -2.65 |
| [M+H]+1 | -0.46 |
| [M+H]+1 | 0.04 |
| [M+H]+1 | 0.55 |
| [M+H]+1 | 0.67 |
| [M+H]+1 | -1.84 |
| [M+NH4]+1 | -4.05 |
| [M+H]+1 | -1.42 |
| [M+H]+1 | -1.27 |
| [M+H]+1 | 2.29 |
| [M+H]+1 | -2.4 |
| [M+H]+1 | -0.14 |

| | |
|---|---|
| [M+H]+1 | 0 |
| [M+H]+1 | 0.62 |
| [M+H]+1 | 0.99 |
| [M+H]+1 | -0.4 |
| [M+H]+1 | 0.74 |
| [M+H]+1 | 0.84 |
| [M+H]+1 | 0.5 |
| [M+H]+1 | 0.11 |
| [M+ACN+H]+1 | 1.87 |
| [M+H]+1 | 0.14 |
| [M+H]+1 | 0.99 |
| [M+H]+1 | 0.74 |
| [M+H]+1 | -2.11 |
| [M+H]+1 | 1.16 |
| [M+H]+1 | -0.4 |
| [M+H]+1 | -3.12 |
| [M+ACN+H]+1 | 2.17 |
| [M+H]+1 | -0.23 |
| [M+H]+1 | 1.43 |
| [M+H]+1 | 0.43 |
| [M+H]+1 | -0.79 |
| [M+H]+1 | -0.96 |
| [M+H]+1 | -0.67 |
| [M+H]+1 | -2.76 |
| [M+H]+1 | 0.18 |
| [M+H]+1 | -4.55 |
| [M+H]+1 | 1.04 |
| [M+H]+1 | -1.93 |
| [M+H]+1 | -0.36 |
| [M+H]+1 | 0.55 |
| [M+H]+1 | -1.57 |
| [M+H]+1 | -3.47 |
| [M+H]+1 | 1.04 |
| [M+H]+1 | -1.45 |
| [M+H]+1 | -1.88 |

| | |
|---|---|
| [M+H]+1 | 2.24 |
| [M+H]+1 | 0.79 |
| [M+H]+1 | -4.43 |
| [M+H]+1 | -1.74 |
| [M+H]+1 | -3.76 |
| [M+H]+1 | -1.45 |
| [M+H]+1 | -3.94 |
| [M+H]+1 | -0.42 |
| [M+H]+1 | 1.38 |
| [M+H]+1 | -0.3 |
| [M+H]+1 | 1.93 |
| [M+H]+1 | 0.44 |
| [M+H]+1 | -3.1 |
| [M+H]+1 | -2.37 |
| [M+H]+1 | 1.1 |
| [M+H]+1 | -0.6 |
| [M+H]+1 | 0.05 |
| [M+H]+1 | -2.15 |
| [M+H]+1 | -3.7 |
| [M+H]+1 | -1.29 |
| [M+H]+1 | -1.14 |
| [M+H]+1 | -1.52 |
| [M+H]+1 | 0.87 |
| [M+H]+1 | 0.25 |
| [M+H]+1 | 0.77 |
| [M+H]+1 | 0.01 |
| [M+H]+1 | -1.79 |
| [M+H]+1 | -2.33 |
| [M+H]+1 | -4.37 |
| [M+H]+1 | -4.14 |
| [M+H]+1 | -4.21 |
| [M+H]+1 | -3.9 |
| [M+H]+1 | -1.95 |
| [M+H]+1 | -1.21 |
| [M+H]+1 | -2.96 |

| | |
|---|---|
| [M+H]+1 | -4.11 |
| [M+NH4]+1 | -0.18 |
| [M+NH4]+1 | -0.56 |
| [M+NH4]+1 | -0.42 |
| [M+H]+1 | 0.77 |
| [M+NH4]+1 | -0.05 |
| [M+NH4]+1 | -0.75 |
| [M+NH4]+1 | -0.06 |
| [M+H]+1 | 0.29 |
| [M+NH4]+1 | 0.8 |
| [M+NH4]+1 | 0.66 |
| [M+NH4]+1 | -0.9 |
| [M+NH4]+1 | -1.79 |
| [M+NH4]+1 | -1.48 |
| [M+H]+1 | 2.37 |
| [M+H]+1 | 2.58 |
| [M+H]+1 | -3.71 |
| [M+H]+1 | 1.46 |
| [M+H]+1 | -2.12 |
| [M+H]+1 | 0.25 |
| [M+H]+1 | -2.39 |
| [M+H]+1 | 0.75 |
| [M+H]+1 | -1.32 |
| [M+H]+1 | -0.77 |
| [M+H]+1 | -1.63 |
| [M+H]+1 | -1.9 |
| [M+H]+1 | 0.07 |
| [M+ACN+Na]+1 | -0.09 |
| [M+H]+1 | 1.01 |
| [M+H]+1 | -0.19 |
| [M+H]+1 | 1.2 |
| [M+H+MeOH]+1 | 0.27 |
| [M+H]+1 | 0.73 |
| [M+H]+1 | 0.06 |
| [M+H]+1 | -2.43 |

| | |
|---|---|
| [M+H]+1 | 1.27 |
| [M+H]+1 | 0.32 |
| [M+H]+1 | 0.18 |
| [M+H]+1 | -1.04 |
| [M+H]+1 | -0.61 |
| [M+H]+1 | -0.75 |
| [M+H]+1 | -0.69 |
| [M+H]+1 | 0.15 |
| [M+H]+1 | -0.48 |
| [M+H]+1 | -2.44 |
| [M+H]+1 | 1.85 |
| [M+H]+1 | 0.84 |
| [M+H]+1 | 1.07 |
| [M+H]+1 | -1.03 |
| [M+H]+1 | 4.44 |
| [M+H]+1 | 2.88 |
| [M+H]+1 | 2.06 |
| [M+H]+1 | 3.1 |
| [M+H]+1 | -0.2 |
| [M+H]+1 | -5.58 |
| [M+H]+1 | -0.69 |
| [M+H]+1 | 3.38 |
| [M+H]+1 | 1.2 |
| [M+H]+1 | 2.04 |
| [M+H]+1 | 0.56 |
| [M+H]+1 | 0.13 |
| [M+H]+1 | -1.16 |
| [M+H]+1 | 1.07 |
| [M+H]+1 | 1.13 |
| [M+H]+1 | 0.39 |
| [M+H]+1 | -1.32 |
| [M+H]+1 | -3.48 |
| [M+H]+1 | 1.25 |
| [M+H]+1 | -1.6 |
| [M+H]+1 | -2.8 |

| | |
|---|---|
| [M+H]+1 | -0.04 |
| [M+H]+1 | -3.17 |
| [M+H]+1 | -1.01 |
| [M+H]+1 | 1.38 |
| [M+H]+1 | 0.07 |
| [M+H]+1 | -1.46 |
| [M+H]+1 | -2.64 |
| [M+H]+1 | 0.09 |
| [M+H]+1 | -0.67 |
| [M+H]+1 | -0.55 |
| [M+H]+1 | 1.45 |
| [M+H]+1 | 0.53 |
| [M+H]+1 | 0.53 |
| [M+H]+1 | 0.3 |
| [M+H]+1 | -1.06 |
| [M+H]+1 | 1.01 |
| [M+H]+1 | -2.43 |
| [M+H]+1 | 0.65 |
| [M+H]+1 | -2.41 |
| [M+H]+1 | -0.03 |
| [M+H]+1 | -5.37 |
| [M+H]+1 | 2.52 |
| [M+H]+1 | -0.38 |
| [M+H]+1 | -1.23 |
| [M+H]+1 | 0.29 |
| [M+H]+1 | 0.15 |
| [M+H]+1 | -0.38 |
| [M+H]+1 | -1.52 |
| [M+H]+1 | -2.65 |
| [M+H]+1 | -1.84 |
| [M+H]+1 | -1.79 |
| [M+H]+1 | -2.97 |
| [M+H]+1 | 3.13 |
| [M+H]+1 | -2.09 |
| [M+H]+1 | -0.91 |

| | |
|---|---|
| [M+H]+1 | -1.08 |
| [M+H]+1 | 1.73 |
| [M+H]+1 | -2.27 |
| [M+H]+1 | -1.16 |
| [M+H]+1 | 0.52 |
| [M+H]+1 | 4.06 |
| [M+H]+1 | -2.83 |
| [M+H]+1 | -2.8 |
| [M+H]+1 | 0.44 |
| [M+H]+1 | -1.97 |
| [M+H]+1 | -0.75 |
| [M+H]+1 | 1.16 |
| [M+H]+1 | -2.05 |
| [M+H]+1 | -2.31 |
| [M+H]+1 | 1.85 |
| [M+H]+1 | 1.02 |
| [M+H]+1 | 0.09 |
| [M+H]+1 | -3.26 |
| [M+H]+1 | -3.76 |
| [M+H]+1 | 2.33 |
| [M+H]+1 | -1.84 |
| [M+H]+1 | 2.94 |
| [M+H]+1 | -2.2 |
| [M+H]+1 | -2.67 |
| [M+H]+1 | -0.95 |
| [M+H]+1 | 2.02 |
| [M+H]+1 | 2.19 |
| [M+H]+1 | 3.95 |
| [M+H]+1 | -2.87 |
| [M+H]+1 | 7.19 |
| [M+H]+1 | 8.35 |
| [M+H]+1 | -1.66 |
| [M+H]+1 | 3.11 |
| [M+H]+1 | 6.32 |
| [M+H]+1 | 3.31 |

| | |
|---|---|
| [M+H]+1 | 2.91 |
| [M+H]+1 | -0.18 |
| [M+H]+1 | 3.93 |
| [M+H]+1 | 4.2 |
| [M+H]+1 | -0.03 |
| [M+H]+1 | 0.27 |
| [M+H]+1 | 0.64 |
| [M+H]+1 | -0.95 |
| [M+H]+1 | -0.21 |
| [M+H]+1 | -1.37 |
| [M+H]+1 | 2.39 |
| [M+H]+1 | -0.53 |
| [M+H]+1 | 0.69 |
| [M+H]+1 | 1.5 |
| [M+H]+1 | 0.16 |
| [M+H]+1 | -2.31 |
| [M+H]+1 | 1.34 |
| [M+H]+1 | 0.39 |
| [M+H]+1 | -2.59 |
| [M+H]+1 | 0.78 |
| [M+H]+1 | -3.14 |
| [M+H]+1 | -2.08 |
| [M+H]+1 | -1.08 |
| [M+H]+1 | -1.62 |
| [M+H]+1 | -2.84 |
| [M+H]+1 | 0.85 |
| [M+H]+1 | -1.7 |
| [M+H]+1 | -1.72 |
| [M+H]+1 | -1.59 |
| [M+H]+1 | -1.85 |
| [M+H]+1 | -0.42 |
| [M+H]+1 | -1.14 |
| [M+H]+1 | -0.7 |
| [M+H]+1 | -4.07 |
| [M+H]+1 | 0.46 |

| | |
|---|---|
| [M+H]+1 | 0.67 |
| [M+H]+1 | -0.39 |
| [M+H]+1 | 1 |
| [M+H]+1 | 0.39 |
| [M+H]+1 | -0.88 |
| [M+H]+1 | -0.07 |
| [M+H]+1 | -0.99 |
| [M+H]+1 | -0.69 |
| [M+H]+1 | -1.31 |
| [M+H]+1 | 1.62 |
| [M+H]+1 | -0.07 |
| [M+H]+1 | -0.12 |
| [M+H]+1 | 0.29 |
| [M+H]+1 | -3.71 |
| [M+H]+1 | -4.2 |
| [M+H]+1 | -1.87 |
| [M+H]+1 | -2.57 |
| [M+H]+1 | -3.78 |
| [M+H]+1 | -4.54 |
| [M+H]+1 | -2.79 |
| [M+H]+1 | 1.27 |
| [M+H-H2O]+1 | -1.32 |
| [M+H]+1 | -0.55 |
| [M+H]+1 | 0.27 |
| [M+H]+1 | -1.77 |
| [M+H]+1 | 2.56 |
| [M+H]+1 | 1.5 |
| [M+H]+1 | -3.14 |
| [M+H]+1 | 0.37 |
| [M+H]+1 | -2.65 |
| [M+H]+1 | -4.5 |
| [M+H]+1 | 4 |
| [M+H]+1 | -2.45 |
| [M+H]+1 | -3.83 |
| [M+H]+1 | -2.41 |

| | |
|---|---|
| [M+H]+1 | -2.23 |
| [M+H]+1 | -2.42 |
| [M+H]+1 | -1.1 |
| [M+H]+1 | -6.14 |
| [M+H]+1 | -0.09 |
| [M+H]+1 | 1.8 |
| [M+H]+1 | 2.89 |
| [M+H]+1 | 0.32 |
| [M+H]+1 | 0.05 |
| [M+H]+1 | 0.26 |
| [M+H]+1 | 0.41 |
| [M+H]+1 | -2.77 |
| [M+H]+1 | -4.4 |
| [M+H]+1 | -2.54 |
| [M+H]+1 | 1.02 |
| [M+H]+1 | -0.32 |
| [M+H]+1 | 2.62 |
| [M+H]+1 | 0.71 |
| [M+H]+1 | -2.81 |
| [M+H]+1 | -4.9 |
| [M+H]+1 | -1.2 |
| [M+H]+1 | -5.71 |
| [M+H]+1 | -2.39 |
| [M+H]+1 | -3.89 |
| [M+H]+1 | 0.09 |
| [M+H]+1 | -2.51 |
| [M+H]+1 | 0.38 |
| [M+H]+1 | -3.6 |
| [M+H]+1 | -2.09 |
| [M+H]+1 | 1.87 |
| [M+H]+1 | 1.13 |
| [M+H]+1 | 0.63 |
| [M+H]+1 | -1.64 |
| [M+H]+1 | -4.71 |
| [M+H]+1 | -3.87 |

| | |
|---|---|
| [M+H]+1 | -4.1 |
| [M+H]+1 | -2.01 |
| [M+H]+1 | 0.41 |
| [M+H]+1 | -2.08 |
| [M+Na]+1 | 0.64 |
| [M+H]+1 | 0.71 |
| [M+H]+1 | -4.21 |
| [M+H]+1 | -1.77 |
| [M+H]+1 | 1.11 |
| [M+H]+1 | 1.75 |
| [M+H]+1 | -0.98 |
| [M+H]+1 | 3.13 |
| [M+H]+1 | 5.77 |
| [M+H]+1 | 6.43 |
| [M+H]+1 | 6.98 |
| [M+H]+1 | 6.62 |
| [M+H]+1 | 8.23 |
| [M+H]+1 | 7.37 |
| [M+H]+1 | -1.5 |
| [M+H]+1 | -0.36 |
| [M+H]+1 | 0.97 |
| [M+H]+1 | 0.19 |
| [M+H]+1 | 1.98 |
| [M+H]+1 | 1.19 |
| [M+H]+1 | 4 |
| [M+H]+1 | 6.17 |
| [M+H]+1 | 7.16 |
| [M+H]+1 | -0.21 |
| [M+H]+1 | -0.68 |
| [M+H]+1 | -0.01 |
| [M+H]+1 | 0.1 |
| [M+H]+1 | 4.42 |
| [M+H]+1 | 4.5 |
| [M+H]+1 | 7.16 |
| [M+H]+1 | 2.81 |

| | |
|---|---|
| [M+H]+1 | 0.2 |
| [M+H]+1 | 2.11 |
| [M+H]+1 | -3.28 |
| [M+H]+1 | -2.11 |
| [2M+Na]+1 | 3.29 |
| [M+H]+1 | 0.57 |
| [M+H]+1 | 0.46 |
| [M+H]+1 | 0.59 |
| [M+H]+1 | 0.57 |
| [M+H]+1 | -1.45 |
| [M+H]+1 | 0.24 |
| [M+H]+1 | -1.38 |
| [M+H]+1 | 0.05 |
| [M+H]+1 | 0.25 |
| [M+H]+1 | -0.4 |
| [M+H]+1 | -2.68 |
| [M+H]+1 | 3.27 |
| [M+H]+1 | 0.89 |
| [M+H]+1 | 3.85 |
| [M+H]+1 | -0.29 |
| [M+H]+1 | -0.95 |
| [M+H]+1 | 0.82 |
| [M+H]+1 | 0.45 |
| [M+H]+1 | 0.42 |
| [M+H]+1 | -1.4 |
| [M+H]+1 | -0.59 |
| [M+H]+1 | 2.48 |
| [M+H]+1 | 0.47 |
| [M+H]+1 | -0.12 |
| [M+H]+1 | 0.46 |
| [M+H]+1 | -2.27 |
| [M+H]+1 | -4.19 |
| [M+H]+1 | -0.14 |
| [M+H]+1 | 1.37 |
| [M+H]+1 | -1.31 |

| | |
|---|---|
| [M+H]+1 | 0.17 |
| [M+H]+1 | 0.46 |
| [M+H]+1 | -0.7 |
| [M+H]+1 | 0.64 |
| [M+H]+1 | -1.3 |
| [M+H]+1 | -0.14 |
| [M+H]+1 | -2.98 |
| [M+H]+1 | 0.58 |
| [M+H]+1 | -5.72 |
| [M+H]+1 | -1.55 |
| [M+H]+1 | -0.13 |
| [M+H]+1 | -0.53 |
| [M+H]+1 | 0.89 |
| [M+H]+1 | 0.72 |
| [M+H]+1 | -1.01 |
| [M+H]+1 | -1.3 |
| [M+H]+1 | -2.1 |
| [M+H]+1 | -3.14 |
| [M+H]+1 | -1.11 |
| [M+H]+1 | 1.85 |
| [M+H]+1 | -5.92 |
| [M+H]+1 | -2.13 |
| [M+H]+1 | -5.33 |
| [M+H]+1 | 1.72 |
| [M+H]+1 | -0.15 |
| [M+H]+1 | 0.09 |
| [M+H]+1 | -5.31 |
| [M+H]+1 | -0.13 |
| [M+H]+1 | -0.02 |
| [M+H]+1 | -5.45 |
| [M+H]+1 | 0.91 |
| [M+H]+1 | -1.04 |
| [M+H]+1 | -0.62 |
| [M+NH4]+1 | 4.44 |
| [M+H]+1 | 0.52 |

| | |
|---|---|
| [M+H]+1 | 6.45 |
| [M+NH4]+1 | 0.09 |
| [M+NH4]+1 | 5.47 |
| [M+H]+1 | 1.91 |
| [M+NH4]+1 | 1.29 |
| [M+NH4]+1 | 0.75 |
| [M+H]+1 | -0.46 |
| [M+H]+1 | -1.78 |
| [M+H]+1 | -0.58 |
| [M+H]+1 | 1.3 |
| [M+H]+1 | 0.04 |
| [M+H]+1 | -0.95 |
| [M+H]+1 | 1.28 |
| [M+H]+1 | -1.78 |
| [M+H]+1 | -1.25 |
| [M+H]+1 | -3.92 |
| [M+H]+1 | -0.65 |
| [M+H]+1 | 2.58 |
| [M+H]+1 | 1.37 |
| [M+H]+1 | 0.57 |
| [M+H]+1 | 1.68 |
| [M+H]+1 | -4.88 |
| [M+H]+1 | 0.99 |
| [M+H]+1 | 0.01 |
| [M+H]+1 | 3.28 |
| [M+H]+1 | 3.74 |
| [M+H]+1 | 2.01 |
| [M+H]+1 | 0.51 |
| [M+H]+1 | 2.28 |
| [M+H]+1 | -0.25 |
| [M+H]+1 | -0.58 |
| [M+H]+1 | -0.58 |
| [M+H]+1 | 0.67 |
| [M+H]+1 | -0.13 |
| [M+H]+1 | 0.59 |

| | |
|---|---|
| [M+H]+1 | 0.93 |
| [M+H]+1 | 0.34 |
| [M+H]+1 | 0.86 |
| [M+H]+1 | -0.75 |
| [M+H]+1 | -0.46 |
| [M+H]+1 | 1.03 |
| [M+H]+1 | -4.39 |
| [M+H]+1 | -1.95 |
| [M+NH4]+1 | -2.29 |
| [M+H]+1 | 1.41 |
| [M+H]+1 | -0.21 |
| [M+H]+1 | 5.39 |
| [M+NH4]+1 | -1.36 |
| [M+H]+1 | 0.42 |
| [M+H]+1 | -4 |
| [M+H]+1 | 1.9 |
| [M+H]+1 | 4.01 |
| [M+NH4]+1 | 0.39 |
| [M+H]+1 | 2.12 |
| [M+H-H2O]+1 | 3.68 |
| [M+NH4]+1 | 0.71 |
| [M+NH4]+1 | 1.28 |
| [M+H]+1 | 0.14 |
| [M+NH4]+1 | -1.11 |
| [M+NH4]+1 | 0.72 |
| [M+NH4]+1 | 0.78 |
| [M+H]+1 | 1.42 |
| [M+NH4]+1 | -1.67 |
| [M+NH4]+1 | 1.07 |
| [M+NH4]+1 | 0.03 |
| [M+NH4]+1 | -2.2 |
| [M+H]+1 | -0.64 |
| [M+NH4]+1 | -0.38 |
| [M+NH4]+1 | -0.06 |
| [M+NH4]+1 | 0.51 |

| | |
|---|---|
| [M+NH4]+1 | -0.97 |
| [M+H]+1 | -4.95 |
| [M+NH4]+1 | -5.63 |
| [M+H]+1 | -2.37 |
| [M+H]+1 | -5.24 |
| [M+H]+1 | -2.98 |
| [M+H]+1 | -2.85 |
| [M+H]+1 | -2.71 |
| [M+H]+1 | -5.15 |
| [M+H]+1 | -2.68 |
| [M+H]+1 | -0.88 |
| [M+H]+1 | 0.02 |
| [M+H]+1 | -2.91 |
| [M+H]+1 | -0.15 |
| [M+H]+1 | 0.88 |
| [M+H]+1 | -0.05 |
| [M+H]+1 | 0 |
| [M+H]+1 | 4.11 |
| [M+H]+1 | 0.77 |
| [M+H]+1 | 1.21 |
| [M+H]+1 | 0.16 |
| [M+H]+1 | 0.66 |
| [M+H]+1 | 0.36 |
| [M+H]+1 | 0.23 |
| [M+H]+1 | 0.37 |
| [M+H]+1 | 0.37 |
| [M+H]+1 | 0.62 |
| [M+H]+1 | 0.9 |
| [M+H]+1 | -0.37 |
| [M+H]+1 | -0.37 |
| [M+H]+1 | -0.12 |
| [M+H]+1 | -0.17 |
| [M+H]+1 | -2.65 |
| [M+H]+1 | -1.24 |
| [M+H]+1 | -2.73 |

| | |
|---|---|
| [M+H]+1 | -0.39 |
| [M+H]+1 | 0.26 |
| [M+H]+1 | 0.76 |
| [M+H]+1 | 0.2 |
| [M+H]+1 | -0.93 |
| [M+H]+1 | 0.48 |
| [M+H]+1 | -0.91 |
| [M+H]+1 | -3.8 |
| [M+H]+1 | 0.52 |
| [M+H]+1 | -5.29 |
| [M+H]+1 | -5.15 |
| [M+H]+1 | -5.18 |
| [M+H]+1 | -4.57 |
| [M+H]+1 | -1.2 |
| [M+H]+1 | -0.32 |
| [M+H]+1 | -1.24 |
| [M+H]+1 | 0.12 |
| [M+H]+1 | -1.01 |
| [M+H]+1 | 0.63 |
| [M+H]+1 | 0.48 |
| [M+H]+1 | 2.17 |
| [M+Na]+1 | 0.83 |
| [M+H]+1 | 2.11 |
| [M+H]+1 | -0.82 |
| [M+H]+1 | 0.47 |
| [M+H]+1 | -0.66 |
| [M+NH4]+1 | 6.24 |
| [M+H]+1 | 2.68 |
| [M+H]+1 | 0.05 |
| [M+H]+1 | -2.45 |
| [M+NH4]+1 | -1.17 |
| [M+H]+1 | -1.84 |
| [M+H]+1 | 1.03 |
| [M+H]+1 | 0.58 |
| [M+H]+1 | -2.05 |

| | |
|---|---|
| [M+H]+1 | 0.67 |
| [M+H]+1 | 0.89 |
| [M+H]+1 | -2.19 |
| [M+H]+1 | -1.19 |
| [M+H]+1 | -0.91 |
| [M+H]+1 | -3.63 |
| [M+H]+1 | -3.01 |
| [M+H]+1 | -0.03 |
| [M+NH4]+1 | -2.22 |
| [M+H]+1 | -1.3 |
| [M+H]+1 | 0.11 |
| [M+H]+1 | 1.32 |
| [M+H]+1 | 1.9 |
| [M+H]+1 | 1.95 |
| [M+H]+1 | -4.54 |

| |
|---|
| Log2 Fold Change: (Bb_pos_12022024_Newanalysis) / (Rabbitserum_12022024_Newanalysis) |
| -2.8 |
| 0.41 |
| -4.78 |
| -8.31 |
| -0.98 |
| 0.26 |
| 2.67 |
| -3.26 |
| -1.6 |
| 0.89 |
| -0.25 |
| 1.34 |
| -4.59 |
| -1.04 |
| 1.26 |
| 3.28 |
| 0.09 |
| -6.81 |
| -4.43 |
| -5.36 |
| 0.12 |
| -6.53 |
| -5.44 |
| -6.96 |
| -7.64 |
| -0.21 |
| 0.11 |
| 1.18 |
| 1.54 |
| 0.4 |
| 1.55 |
| 5.81 |
| 2.9 |

| | |
|---|---|
| | 1.85 |
| | -6.15 |
| | -3.96 |
| | -6.24 |
| | 6.51 |
| | 5.6 |
| | -7.48 |
| | 0.34 |
| | -2.09 |
| | -3.49 |
| | -2.22 |
| | 1 |
| | -5.98 |
| | -6.69 |
| | -6.27 |
| | -5.38 |
| | -3.58 |
| | -5.81 |
| | -4.45 |
| | -6.91 |
| | -4.46 |
| | -0.05 |
| | -1.86 |
| | -1.7 |
| | 1.93 |
| | -2.73 |
| | -0.55 |
| | -3.35 |
| | 0 |
| | 1.28 |
| | -7.23 |
| | -4.22 |
| | -0.31 |
| | -3.94 |
| | -5.04 |

| | |
|---|---|
| | -4.6 |
| | -1.82 |
| | -2.84 |
| | 0.17 |
| | -4.01 |
| | -6.51 |
| | -3.6 |
| | 0.67 |
| | -7.2 |
| | -2.84 |
| | -6.84 |
| | -7.17 |
| | -6.17 |
| | -3.12 |
| | 0.86 |
| | -3.96 |
| | -7.56 |
| | -2.37 |
| | 1.14 |
| | -6.46 |
| | -2.84 |
| | -2.14 |
| | -6.4 |
| | -6.27 |
| | -4.43 |
| | -4.48 |
| | -4.52 |
| | -4.62 |
| | -6 |
| | -5.44 |
| | -0.76 |
| | -1.92 |
| | -5.91 |
| | 1.01 |
| | -3.44 |

| | |
|---|---|
| | 0.4 |
| | -4.3 |
| | 1.51 |
| | -3.52 |
| | -1.01 |
| | -0.63 |
| | 1.13 |
| | -2.69 |
| | -3.7 |
| | -2.5 |
| | -4.2 |
| | 0.57 |
| | -3.18 |
| | -0.99 |
| | -7.4 |
| | -5.55 |
| | -1.64 |
| | 0.15 |
| | -6.46 |
| | -1.77 |
| | 1.69 |
| | -1.8 |
| | -1.77 |
| | -3.54 |
| | -1.07 |
| | -2.77 |
| | 0.96 |
| | -0.56 |
| | -3.51 |
| | -7.09 |
| | -3.32 |
| | -0.54 |
| | 1.98 |
| | -3.67 |
| | -1.62 |

| | |
|---|---|
| | -0.34 |
| | -1.27 |
| | 1.53 |
| | -0.57 |
| | 1.47 |
| | 1.41 |
| | 1.04 |
| | -0.19 |
| | 2.9 |
| | -1.01 |
| | 0.97 |
| | -4.09 |
| | -7.13 |
| | -1.53 |
| | -4.55 |
| | -5.71 |
| | 2.67 |
| | -1.95 |
| | 3.04 |
| | -1.64 |
| | -3.62 |
| | -4.25 |
| | -5.08 |
| | -5.24 |
| | -1.53 |
| | -7.37 |
| | -0.4 |
| | -2.62 |
| | -3.89 |
| | -1.02 |
| | -5.11 |
| | -4.16 |
| | -1.21 |
| | -2.96 |
| | -5.72 |

| | |
|---|---|
| | 0.56 |
| | -4.11 |
| | -5.93 |
| | -6.48 |
| | -5.51 |
| | -4.97 |
| | -5.95 |
| | -3.63 |
| | 2.1 |
| | -7.2 |
| | -1.21 |
| | 0.15 |
| | -7.17 |
| | -4.03 |
| | 2.4 |
| | -4.06 |
| | -3.53 |
| | -3.63 |
| | -6.39 |
| | -4.25 |
| | -3.99 |
| | -3.75 |
| | 0.08 |
| | 0.14 |
| | 1.39 |
| | -0.93 |
| | -3.99 |
| | -0.58 |
| | -5.75 |
| | -7.85 |
| | -5.78 |
| | -5.9 |
| | -5.55 |
| | -5.09 |
| | -6.23 |

| | |
|---|---|
| | -4.98 |
| | -2.08 |
| | -1.65 |
| | -0.29 |
| | 1.21 |
| | -0.47 |
| | -2.39 |
| | -2.82 |
| | -1.23 |
| | -0.71 |
| | 1.33 |
| | -0.63 |
| | -2.77 |
| | -0.94 |
| | 1.94 |
| | 2.4 |
| | -3.09 |
| | -3.91 |
| | -3.04 |
| | -0.19 |
| | -4.54 |
| | -2.32 |
| | 1.08 |
| | -4.58 |
| | -5 |
| | -6.09 |
| | -0.31 |
| | -0.52 |
| | 1.59 |
| | -0.85 |
| | -1.21 |
| | 0.06 |
| | -1.4 |
| | -1.17 |
| | -4.99 |

| |
|---|
| 0.1 |
| -1.9 |
| 0.52 |
| -1.06 |
| -1.98 |
| -3.26 |
| -3.28 |
| -0.75 |
| -1.86 |
| -6.66 |
| 0.16 |
| -1.06 |
| -2.1 |
| -3.33 |
| 1.84 |
| -0.58 |
| 0.41 |
| 0.61 |
| 0.6 |
| -7.65 |
| -3.37 |
| 0.07 |
| -0.85 |
| 0.14 |
| -0.68 |
| -0.13 |
| -2.45 |
| -0.22 |
| -1.85 |
| -2.6 |
| -2.79 |
| -5.59 |
| 2.08 |
| -3.2 |
| -5.88 |

| |
|---|
| -3.51 |
| -6.42 |
| -2.82 |
| -3.86 |
| -3.18 |
| -3.03 |
| -6.04 |
| -0.37 |
| -1.43 |
| -4.59 |
| 0.27 |
| -2.74 |
| 0.87 |
| -1.15 |
| -0.87 |
| 0.11 |
| -7.02 |
| -2.13 |
| -2.21 |
| -2.36 |
| -6.19 |
| -2 |
| -1.07 |
| -6.29 |
| -3.97 |
| -3.74 |
| -3.34 |
| -4.11 |
| -1.15 |
| -4.39 |
| -4.26 |
| -5.53 |
| 1.23 |
| -3.65 |
| -1.71 |

| | |
|---|---|
| | -4.82 |
| | 0.86 |
| | -3.85 |
| | -4.61 |
| | 2.66 |
| | 2.32 |
| | -4.89 |
| | -6.98 |
| | -0.48 |
| | -5.18 |
| | -2.05 |
| | 2.74 |
| | -2.74 |
| | -2.84 |
| | 1.5 |
| | -1.25 |
| | -2.46 |
| | -5.62 |
| | -6.27 |
| | -2.49 |
| | -1.69 |
| | -0.22 |
| | -2.53 |
| | -6.5 |
| | -4.18 |
| | 2.04 |
| | 3.14 |
| | 0.96 |
| | -1.56 |
| | 3.6 |
| | 3.56 |
| | -4.8 |
| | -0.49 |
| | 4.3 |
| | 1.87 |

| |
|---|
| 1.28 |
| -1.09 |
| 5.85 |
| 4.55 |
| -4.26 |
| -0.58 |
| -2.75 |
| -5.24 |
| -2.93 |
| -3.64 |
| -0.91 |
| -3.76 |
| -3.44 |
| 1.52 |
| -0.6 |
| -3.45 |
| -0.34 |
| -2.55 |
| -4.89 |
| -2.64 |
| -5.92 |
| -6.08 |
| 0.6 |
| -4.47 |
| -6.04 |
| 0.71 |
| -4.12 |
| -4.16 |
| -4.89 |
| -5.38 |
| -3.21 |
| -4.27 |
| -3.52 |
| -4.69 |
| -0.78 |

| | |
|---|---|
| | -2.16 |
| | -1.48 |
| | -1.5 |
| | -2.34 |
| | -2.49 |
| | -3.8 |
| | -3.39 |
| | -4.46 |
| | -4.32 |
| | 0.46 |
| | -3.62 |
| | -5.83 |
| | -3.68 |
| | -7.2 |
| | -7.34 |
| | -2.6 |
| | -4.17 |
| | -5.46 |
| | -7.12 |
| | -4.2 |
| | 1.86 |
| | -2.34 |
| | -2.24 |
| | -4.16 |
| | -1.44 |
| | 1.76 |
| | -1.47 |
| | -4.66 |
| | 0.9 |
| | -4.59 |
| | -7.87 |
| | 5.11 |
| | -5.32 |
| | -5.69 |
| | -4.82 |

| | |
|---|---|
| | -5.32 |
| | -4.56 |
| | -3.91 |
| | -7.77 |
| | -2.47 |
| | 0.15 |
| | 2.06 |
| | -1.26 |
| | -4.1 |
| | -2.82 |
| | -1.41 |
| | -5.64 |
| | -7.89 |
| | -2.96 |
| | -2.56 |
| | -2.99 |
| | -0.22 |
| | -0.71 |
| | -4.42 |
| | -7.69 |
| | -2.84 |
| | -9.35 |
| | -4.82 |
| | -5.21 |
| | -2.28 |
| | -4.04 |
| | -3.99 |
| | -6.98 |
| | -2.52 |
| | 1.29 |
| | -2.11 |
| | -1.26 |
| | -5.34 |
| | -8.2 |
| | -7.38 |

| | |
|---|---|
| | -7.17 |
| | -4.9 |
| | -3.99 |
| | -1.75 |
| | -0.18 |
| | -0.67 |
| | -6.79 |
| | -4.56 |
| | -1.5 |
| | -2.85 |
| | -4.12 |
| | 0.99 |
| | 2.38 |
| | 5.16 |
| | 3.62 |
| | 5.84 |
| | 5.39 |
| | 6.72 |
| | -2.35 |
| | -2.97 |
| | -1.69 |
| | -3.3 |
| | -1.59 |
| | -2.58 |
| | 0.41 |
| | 3.33 |
| | 4.76 |
| | -3.69 |
| | -0.63 |
| | -1.41 |
| | -2.45 |
| | 2.27 |
| | 1.36 |
| | 6.79 |
| | 3.11 |

| | |
|---|---|
| | 0.82 |
| | 2.52 |
| | -2.54 |
| | -4.96 |
| | 0.28 |
| | -4.95 |
| | 3.41 |
| | 0.08 |
| | 2.6 |
| | 2.14 |
| | 2.12 |
| | -5.4 |
| | -5.4 |
| | -2.68 |
| | -3.16 |
| | -4.52 |
| | 0.32 |
| | -2.05 |
| | 2.6 |
| | -2.3 |
| | -4.33 |
| | -1.04 |
| | -0.81 |
| | 0.11 |
| | -3.71 |
| | -5.32 |
| | 0.25 |
| | -3.73 |
| | -3.71 |
| | -2.14 |
| | -4.6 |
| | -6.16 |
| | -5.65 |
| | 1.71 |
| | -5.32 |

| | |
|---|---|
| | -4.13 |
| | -2.62 |
| | -4.85 |
| | 2.74 |
| | -7.27 |
| | -2.31 |
| | -6.39 |
| | -0.58 |
| | -9.79 |
| | -6.92 |
| | -3.05 |
| | -4.45 |
| | -2.3 |
| | 2.71 |
| | -5.11 |
| | -2.08 |
| | -5.15 |
| | -7.88 |
| | -5.94 |
| | 2.78 |
| | -9.28 |
| | -6.57 |
| | -9.37 |
| | -2.86 |
| | -4.64 |
| | -2.65 |
| | -9.61 |
| | -2.32 |
| | -3.3 |
| | -8.97 |
| | -2.37 |
| | -2.75 |
| | 1.09 |
| | 4.36 |
| | 0.83 |

| | |
|---|---|
| | 1.49 |
| | -0.63 |
| | 0.5 |
| | 2.76 |
| | 4.11 |
| | 1.51 |
| | -0.57 |
| | -3.26 |
| | -3.46 |
| | 0.9 |
| | 1.38 |
| | -1.11 |
| | 4.42 |
| | -4.89 |
| | -3.05 |
| | -5.69 |
| | -0.74 |
| | 2.81 |
| | 0.41 |
| | 1.14 |
| | 2.99 |
| | -6.2 |
| | -0.97 |
| | 3.57 |
| | 2.06 |
| | 3.4 |
| | 3.56 |
| | 2.67 |
| | 3.24 |
| | -1.28 |
| | -0.21 |
| | -2.91 |
| | -2.15 |
| | 0.5 |
| | -0.2 |

| | |
|---|---|
| | 3.32 |
| | 0.66 |
| | 1.92 |
| | 2.4 |
| | 0.67 |
| | -1.18 |
| | -7.42 |
| | -4.71 |
| | -5.58 |
| | 0.85 |
| | -1.87 |
| | 2.67 |
| | -4.97 |
| | -1.1 |
| | -7.1 |
| | 4.98 |
| | 2.15 |
| | 0.09 |
| | 1.98 |
| | 0.67 |
| | 1.13 |
| | 4.66 |
| | -1.37 |
| | 0.41 |
| | 2.64 |
| | 1.26 |
| | 1.98 |
| | -4.91 |
| | -1.34 |
| | -0.74 |
| | -5.07 |
| | -2.23 |
| | -3.27 |
| | 1.86 |
| | -2.59 |

| | |
|---|---|
| | -2.06 |
| | -4.92 |
| | -7.51 |
| | -3.84 |
| | -6.14 |
| | -3.66 |
| | -3.74 |
| | -6.07 |
| | -6.41 |
| | -3.83 |
| | -2.15 |
| | 1.65 |
| | -3.46 |
| | -0.31 |
| | 2.82 |
| | -0.92 |
| | 0.69 |
| | 5.2 |
| | 2.85 |
| | 3.03 |
| | 0.46 |
| | 0.96 |
| | 0.93 |
| | -2.41 |
| | 0.82 |
| | 1.13 |
| | 0.91 |
| | 1.53 |
| | -1.28 |
| | 0.24 |
| | 1.44 |
| | 0.45 |
| | -3.54 |
| | -4.17 |
| | -3.84 |

| |
|---|
| -1.5 |
| 0.35 |
| 0.16 |
| -3.22 |
| -4.12 |
| -3.34 |
| -2.49 |
| -6.48 |
| -1.53 |
| -7 |
| -6.92 |
| -6.99 |
| -7.09 |
| -3.65 |
| -3.06 |
| -5.04 |
| -3.15 |
| -0.12 |
| 0.75 |
| -0.08 |
| 2.79 |
| -0.74 |
| 4.33 |
| -0.97 |
| -1.26 |
| -3.28 |
| 2.62 |
| 3.18 |
| -2.66 |
| -2.46 |
| -5.11 |
| -3.56 |
| 1.44 |
| 0.5 |
| -4.49 |

| |
|---|
| -0.78 |
| 1.59 |
| -0.91 |
| -0.94 |
| -1.62 |
| -6.52 |
| -5 |
| 0.46 |
| -1.47 |
| -2.42 |
| 0.59 |
| 1.65 |
| 1.02 |
| 1.77 |
| -11.28 |

| |
|---|
| Log2 Fold Change: (CM_pos_12022024_Newanalysis) / (Rabbitserum_12022024_Newanalysis) |
| -2.21 |
| -0.62 |
| -5.13 |
| -3.06 |
| -1.9 |
| 0.38 |
| 1.88 |
| -1.14 |
| -1.97 |
| 1.71 |
| 0.77 |
| 1.01 |
| -2.64 |
| 0.34 |
| 1.13 |
| 2.17 |
| -0.36 |
| -2.11 |
| -2.97 |
| -3.64 |
| 0 |
| -2.25 |
| -1.36 |
| -3.48 |
| -1.75 |
| 1.21 |
| -0.06 |
| 0.35 |
| 0.9 |
| 1.2 |
| 1.1 |
| -0.72 |
| 1.38 |

| | |
|---|---|
| | 1.12 |
| | -0.99 |
| | -5.92 |
| | -3.45 |
| | -0.06 |
| | -0.93 |
| | -1.39 |
| | 0.13 |
| | -2.51 |
| | -0.04 |
| | -3.25 |
| | 0.53 |
| | -1.76 |
| | -2.02 |
| | -1.73 |
| | -1.27 |
| | -0.81 |
| | -1.07 |
| | -0.68 |
| | -1.62 |
| | -1.72 |
| | -1.22 |
| | -0.74 |
| | -2.98 |
| | 0.23 |
| | -2.12 |
| | -1.94 |
| | -3.02 |
| | -0.7 |
| | 0.06 |
| | -1.24 |
| | -0.81 |
| | -0.38 |
| | 1.12 |
| | -2.77 |

| |
|---|
| -4.18 |
| -0.78 |
| 1.96 |
| -0.38 |
| -1.29 |
| -2.7 |
| -3.24 |
| -0.14 |
| -1.71 |
| -3.12 |
| -2.14 |
| -3.72 |
| -1.9 |
| 1.46 |
| 0.61 |
| -1 |
| -6.17 |
| 0.12 |
| 0.91 |
| -0.16 |
| -0.48 |
| -0.96 |
| -1.05 |
| -1.95 |
| -2.04 |
| -1.84 |
| -2.03 |
| -2.56 |
| -1.02 |
| -1.69 |
| -0.38 |
| 0.03 |
| -3.86 |
| 2.19 |
| -0.95 |

| |
|---|
| 0.66 |
| -2.01 |
| 0.75 |
| -0.96 |
| -0.56 |
| -1.5 |
| 1.64 |
| -0.69 |
| -3.01 |
| 1.37 |
| -1.47 |
| 0.05 |
| -1.13 |
| -0.08 |
| -0.97 |
| -1.48 |
| -1.62 |
| -0.61 |
| -6.37 |
| -1.95 |
| 0.89 |
| -1.36 |
| -1.16 |
| -0.89 |
| -0.61 |
| -2.81 |
| 0.4 |
| -1.23 |
| -1.67 |
| -3.04 |
| -1.9 |
| 0.73 |
| -0.31 |
| -1.27 |
| -1.48 |

| | |
|---|---|
| | -0.34 |
| | -1.89 |
| | 0.54 |
| | -0.17 |
| | 0.73 |
| | 0.57 |
| | 0.54 |
| | -0.31 |
| | 1.03 |
| | -1.16 |
| | -0.02 |
| | -4.83 |
| | -5.02 |
| | -2.68 |
| | -4.15 |
| | -2.6 |
| | 0.51 |
| | -1.72 |
| | 1.61 |
| | -2.07 |
| | -2.82 |
| | -3.29 |
| | -4.41 |
| | -2.48 |
| | -1.72 |
| | -2.82 |
| | -1.43 |
| | -0.69 |
| | -3.53 |
| | -1.57 |
| | -3.54 |
| | -0.69 |
| | -2.25 |
| | -1.51 |
| | -3.84 |

| |
|---|
| -1.68 |
| -4.9 |
| -1.51 |
| -4.74 |
| -1.76 |
| -3.51 |
| -2.01 |
| -3.21 |
| 0.72 |
| -6.9 |
| -3.14 |
| -0.29 |
| -4.07 |
| -1.67 |
| 1.29 |
| -3.46 |
| -3.58 |
| -1.48 |
| -2.7 |
| -2.96 |
| -2.85 |
| -2.23 |
| -0.8 |
| -0.11 |
| 0.62 |
| -0.95 |
| -2.2 |
| 1.75 |
| -1.38 |
| -3.71 |
| -1.57 |
| -2 |
| -3.6 |
| -3.88 |
| -3.26 |

| |
|---|
| -0.87 |
| -1.9 |
| -1.09 |
| 0.13 |
| 0.45 |
| -0.43 |
| -1.64 |
| -2.76 |
| -1.52 |
| -1.5 |
| 0.68 |
| 0.27 |
| -0.98 |
| 0.54 |
| -0.43 |
| -0.18 |
| 0.62 |
| -5.37 |
| -0.92 |
| -0.44 |
| -2.15 |
| -3.07 |
| 2.4 |
| -3.81 |
| -3.38 |
| -4.18 |
| -0.38 |
| -0.43 |
| 0.58 |
| -0.67 |
| -2.4 |
| -0.2 |
| -2.13 |
| -1.23 |
| -2.55 |

| |
|---|
| -1.17 |
| -2.22 |
| 0.34 |
| -0.02 |
| -1.37 |
| -2.51 |
| -2.59 |
| -0.91 |
| -1.38 |
| -4.22 |
| -1.69 |
| -1.9 |
| -3.17 |
| -2.3 |
| -2.6 |
| -3.46 |
| -1.65 |
| -2.48 |
| 0.79 |
| -2.07 |
| -2.69 |
| -3.31 |
| -2.05 |
| -1.89 |
| -1.24 |
| -0.26 |
| -1.3 |
| -1.3 |
| -2.98 |
| -2.99 |
| -1.48 |
| -2.12 |
| 0.83 |
| -1.59 |
| -3.08 |

| | |
|---|---|
| | -3.47 |
| | -3.24 |
| | -1.81 |
| | -5.24 |
| | -3.25 |
| | -1.57 |
| | -3.39 |
| | -0.45 |
| | -0.76 |
| | -4.04 |
| | -1.18 |
| | -3.27 |
| | 0.34 |
| | -1.45 |
| | 0.2 |
| | -0.9 |
| | -4.6 |
| | -2.78 |
| | 0.21 |
| | -2.33 |
| | -0.82 |
| | -4.52 |
| | -0.69 |
| | -5.07 |
| | -4.26 |
| | -3.89 |
| | -2.97 |
| | -2.59 |
| | 1.5 |
| | -2.55 |
| | -2.47 |
| | -2.57 |
| | -1.9 |
| | -1.56 |
| | -0.79 |

| | |
|---|---|
| | -3.73 |
| | -0.88 |
| | -1.58 |
| | -3.45 |
| | 2.15 |
| | -1.74 |
| | -2.06 |
| | -4.19 |
| | -0.92 |
| | -3.21 |
| | -1.3 |
| | 1.57 |
| | -0.69 |
| | -0.53 |
| | -0.35 |
| | -2.27 |
| | -2.55 |
| | -2.35 |
| | -2.52 |
| | -4.82 |
| | 0.15 |
| | -3.16 |
| | -0.32 |
| | -3.83 |
| | -3.23 |
| | 0.03 |
| | 0.95 |
| | -3 |
| | 1.32 |
| | -3.59 |
| | -4.79 |
| | -3.13 |
| | -3.6 |
| | -2.02 |
| | -1.44 |

| |
|---:|
| -1.63 |
| -0.92 |
| 1.92 |
| 0.35 |
| -4.23 |
| -0.85 |
| -3.39 |
| -4.29 |
| -2.72 |
| -2.27 |
| -3.3 |
| -3.22 |
| -4.14 |
| 0.02 |
| -0.76 |
| -1.14 |
| -1.68 |
| -2.95 |
| -2.3 |
| -3.41 |
| -2.78 |
| -4 |
| 1.68 |
| -2.85 |
| -3.2 |
| -0.15 |
| -2.42 |
| -2.43 |
| -3.3 |
| -3.53 |
| -2.8 |
| -3.14 |
| -2.82 |
| -0.62 |
| -1.24 |

| | |
|---|---|
| | -2.83 |
| | -1.08 |
| | -2.5 |
| | -2.73 |
| | -1.61 |
| | -3.73 |
| | -2.4 |
| | -3.77 |
| | -3.01 |
| | -1.16 |
| | -3.55 |
| | -5.71 |
| | -3.97 |
| | -3.48 |
| | -3.15 |
| | -0.73 |
| | -1.6 |
| | -1.68 |
| | -2.58 |
| | -1.41 |
| | 0.58 |
| | -1.02 |
| | -1.7 |
| | -4.43 |
| | 0.33 |
| | -0.79 |
| | -2.97 |
| | -1.52 |
| | 0.52 |
| | -1.94 |
| | -3.37 |
| | 1.11 |
| | -2.88 |
| | -1.86 |
| | -2.41 |

| | |
|---|---|
| | -3.09 |
| | -2.14 |
| | -2.81 |
| | -1.64 |
| | -2.38 |
| | -1.65 |
| | -0.83 |
| | -1.58 |
| | -4.15 |
| | -3.08 |
| | -1.82 |
| | -2.87 |
| | -3.49 |
| | -0.42 |
| | -3.58 |
| | -2.67 |
| | -2.84 |
| | -1.42 |
| | -1.62 |
| | -2.79 |
| | -1.64 |
| | -3.64 |
| | -2.43 |
| | -1.32 |
| | -2.37 |
| | -1.52 |
| | -4.37 |
| | -3.39 |
| | -0.43 |
| | -0.58 |
| | -3.23 |
| | -1.9 |
| | -3.7 |
| | -3.49 |
| | -3.51 |

| | |
|---|---|
| | -3.07 |
| | -2.89 |
| | -4.4 |
| | 0.32 |
| | -0.82 |
| | -1.38 |
| | -2.58 |
| | -2.79 |
| | -2.61 |
| | -4.6 |
| | -3.15 |
| | -2.14 |
| | -3.39 |
| | -1.28 |
| | -3.36 |
| | -0.78 |
| | -2.83 |
| | -0.65 |
| | -0.85 |
| | -2.61 |
| | -2.66 |
| | -3.5 |
| | -3.57 |
| | -3.77 |
| | -3.59 |
| | -2.84 |
| | -2.4 |
| | -3.48 |
| | 0.06 |
| | -1.4 |
| | -2.55 |
| | -2.15 |
| | -3.14 |
| | -0.37 |
| | 0.3 |

| |
|---:|
| 0.62 |
| 0.41 |
| 0.75 |
| -2.85 |
| -3.01 |
| -5.51 |
| 2.95 |
| -0.51 |
| 2.03 |
| 3.59 |
| 1.88 |
| -4.01 |
| -5.45 |
| -2.94 |
| -2.76 |
| -1.84 |
| -2.95 |
| -2.94 |
| -1.25 |
| -2.01 |
| -3.38 |
| -1.87 |
| -1.27 |
| -0.31 |
| -2.31 |
| -4.74 |
| -2.23 |
| -4.2 |
| -3.59 |
| -2.6 |
| -2.33 |
| -1.97 |
| -5.51 |
| 0.34 |
| -4.01 |

| | |
|---|---|
| | -4.31 |
| | -3.08 |
| | -4.15 |
| | 2.1 |
| | -5.97 |
| | -2.17 |
| | -3.41 |
| | -1.16 |
| | -4.07 |
| | -5.37 |
| | -2.92 |
| | -3.92 |
| | -3.2 |
| | 1.99 |
| | -4.09 |
| | -0.77 |
| | -3.05 |
| | -4.75 |
| | -4.83 |
| | 0.93 |
| | -3.36 |
| | -4.44 |
| | -4.04 |
| | -4.58 |
| | -4.49 |
| | -2.74 |
| | -4.3 |
| | -2.19 |
| | -3.28 |
| | -3.53 |
| | -3.28 |
| | -1.71 |
| | 1.71 |
| | -0.09 |
| | 0.32 |

| | |
|---|---|
| | -4.96 |
| | -0.72 |
| | -4.97 |
| | 0.84 |
| | 2.81 |
| | 0.77 |
| | -0.12 |
| | -1.48 |
| | -2.88 |
| | -0.4 |
| | 1.33 |
| | -0.16 |
| | 3.14 |
| | -3.11 |
| | -1.8 |
| | -1.77 |
| | -0.09 |
| | 0.23 |
| | -0.95 |
| | 0.57 |
| | 1.31 |
| | -1.32 |
| | -1.96 |
| | 3.56 |
| | -1.21 |
| | -0.34 |
| | 1.55 |
| | 2.17 |
| | 0.96 |
| | -1.03 |
| | 0.36 |
| | -2.33 |
| | -2.82 |
| | 0.63 |
| | -0.79 |

| | |
|---|---|
| | 2.39 |
| | 0.32 |
| | 1.06 |
| | 3.15 |
| | 1.13 |
| | -2.21 |
| | -3.03 |
| | -2.76 |
| | -3.29 |
| | -0.57 |
| | -1.67 |
| | -2.73 |
| | -3.61 |
| | -1.51 |
| | -3.1 |
| | 3.08 |
| | -1.86 |
| | -0.29 |
| | -0.15 |
| | -3.01 |
| | 0.43 |
| | 3.37 |
| | -1.51 |
| | 1.52 |
| | 1.92 |
| | 0.48 |
| | 0.56 |
| | -3.24 |
| | -2.42 |
| | -0.77 |
| | -2.87 |
| | -1.59 |
| | -2.89 |
| | 1.92 |
| | -3.11 |

| | |
|---|---|
| | -1.09 |
| | 0.04 |
| | -1.88 |
| | -1.47 |
| | -0.9 |
| | -0.68 |
| | -0.89 |
| | -3.36 |
| | -1.25 |
| | -1.15 |
| | -1.27 |
| | 1.64 |
| | -0.55 |
| | -0.16 |
| | 1.94 |
| | -0.87 |
| | 0.7 |
| | 1.1 |
| | 2.08 |
| | 1.82 |
| | 0.3 |
| | 0.31 |
| | 0.57 |
| | -2.64 |
| | 0.44 |
| | 0.76 |
| | 0.29 |
| | 0.63 |
| | -0.9 |
| | 0.61 |
| | 1.56 |
| | 0.61 |
| | -0.89 |
| | -2.93 |
| | -1.11 |

| | |
|---|---|
| | -1.11 |
| | 0.09 |
| | -0.6 |
| | -3.41 |
| | -3.19 |
| | -3.82 |
| | -1.58 |
| | -2.68 |
| | -2.05 |
| | -1.71 |
| | -1.77 |
| | -1.81 |
| | -2.51 |
| | -2.44 |
| | -2.74 |
| | -3.81 |
| | -3.27 |
| | 0.89 |
| | 0.13 |
| | -0.55 |
| | 0.61 |
| | -1.56 |
| | 2.22 |
| | -0.15 |
| | -1.73 |
| | -2.63 |
| | -3.63 |
| | 0.5 |
| | -2.71 |
| | -0.02 |
| | -3.94 |
| | -1.72 |
| | 0.41 |
| | -0.08 |
| | -2.43 |

| |
|---|
| -1.45 |
| 0.7 |
| 1.28 |
| 0.25 |
| -0.71 |
| -2.89 |
| -1.99 |
| 0.49 |
| 0.76 |
| -1.12 |
| 0.48 |
| 0.33 |
| -0.88 |
| -0.18 |
| -6.73 |

| |
|---|
| Log2 Fold Change: (Hs_pos_12022024_Newanalysis) / (Rabbitserum_12022024_Newanalysis) |
| 2.44 |
| -0.59 |
| -1.4 |
| -0.75 |
| -1.29 |
| 0 |
| -7.41 |
| -1.19 |
| 1.27 |
| -1.08 |
| 0.39 |
| -4.93 |
| -0.93 |
| 0.61 |
| 0.2 |
| 0.5 |
| 1.13 |
| -3.92 |
| -2.84 |
| -0.85 |
| -2.62 |
| -4.27 |
| -1.76 |
| -3.85 |
| -2.68 |
| 5.27 |
| 5.67 |
| 7.69 |
| 4.17 |
| -0.48 |
| -1.21 |
| 0.84 |
| -0.3 |

| | |
|---|---|
| | -0.6 |
| | -4.14 |
| | -2.08 |
| | -1.64 |
| | 1.35 |
| | -0.52 |
| | 0.53 |
| | -0.91 |
| | -1.7 |
| | 0.01 |
| | 0.45 |
| | -0.79 |
| | -5.5 |
| | -6.52 |
| | -6.01 |
| | -6.74 |
| | -5.25 |
| | -7.32 |
| | -6.12 |
| | -8.28 |
| | -5.9 |
| | -2.87 |
| | -1.01 |
| | 1.31 |
| | -1.47 |
| | 0.24 |
| | 0.56 |
| | -3.55 |
| | -0.88 |
| | 1.63 |
| | -0.25 |
| | 1.87 |
| | -1.64 |
| | -0.18 |
| | 1.35 |

| | |
|---|---|
| | 2.18 |
| | 0.02 |
| | 0.83 |
| | 0.8 |
| | -0.18 |
| | 2.14 |
| | -1.94 |
| | -2.44 |
| | -2.86 |
| | -2.49 |
| | -1.83 |
| | -1.44 |
| | 0.84 |
| | 1.9 |
| | -0.55 |
| | 0.85 |
| | 1.86 |
| | -0.08 |
| | -0.87 |
| | 0.99 |
| | -1.02 |
| | 0.19 |
| | 0.15 |
| | -1.48 |
| | -2.78 |
| | -4.24 |
| | -2.42 |
| | 2.27 |
| | -0.44 |
| | -2.01 |
| | -2.3 |
| | -1.5 |
| | -2.4 |
| | 0.03 |
| | 0.38 |

| | |
|---|---|
| | 0.98 |
| | -0.1 |
| | -1.06 |
| | 0.19 |
| | 0.28 |
| | -1.43 |
| | -1.53 |
| | -0.57 |
| | -3.12 |
| | -0.87 |
| | -0.36 |
| | -1.02 |
| | 0.2 |
| | -1.34 |
| | -2.51 |
| | -0.79 |
| | -1.31 |
| | -0.56 |
| | -4.34 |
| | -1.1 |
| | 3.74 |
| | -0.33 |
| | -0.6 |
| | -0.6 |
| | 1.65 |
| | 0.27 |
| | 0.71 |
| | -2.16 |
| | 0.61 |
| | -0.4 |
| | 1 |
| | -0.89 |
| | 0.12 |
| | -4.42 |
| | -0.5 |

| | |
|---|---|
| | -0.54 |
| | 1.67 |
| | -3.26 |
| | 2.71 |
| | 0.33 |
| | -1.56 |
| | 0.25 |
| | -2.88 |
| | 0.34 |
| | -2.33 |
| | 0.98 |
| | -2.02 |
| | -0.35 |
| | -1.61 |
| | 2.63 |
| | 1.6 |
| | 0.59 |
| | 2.83 |
| | 2.44 |
| | 2.24 |
| | 3.33 |
| | 2.87 |
| | 2.37 |
| | 1.85 |
| | 3.37 |
| | 2.73 |
| | 1.99 |
| | -0.29 |
| | 3.16 |
| | -0.16 |
| | -0.51 |
| | 2.48 |
| | 1 |
| | 3.15 |
| | -2.83 |

| | |
|---|---|
| | -0.17 |
| | 1.01 |
| | 2.96 |
| | -1.11 |
| | -0.18 |
| | 2.07 |
| | -1.03 |
| | 3.27 |
| | 1.39 |
| | -0.45 |
| | 0.57 |
| | 2.81 |
| | 2.09 |
| | -3.85 |
| | 5.51 |
| | -3.92 |
| | 3.98 |
| | 4.06 |
| | -1.59 |
| | 6.33 |
| | 5.48 |
| | 4.49 |
| | 4.85 |
| | 5.74 |
| | 7.46 |
| | 6.39 |
| | 4.51 |
| | 5.62 |
| | 3.74 |
| | 1.85 |
| | 2.49 |
| | 2.49 |
| | 3.29 |
| | 0.22 |
| | -1.77 |

| | |
|---|---|
| | -3.71 |
| | 4.49 |
| | 2.5 |
| | 5.87 |
| | 5.25 |
| | 5.14 |
| | 1.67 |
| | 1 |
| | 1.95 |
| | 1.16 |
| | 5.64 |
| | 7.22 |
| | 4.38 |
| | 2.35 |
| | -0.04 |
| | 1.32 |
| | -4.73 |
| | -1.56 |
| | 3.17 |
| | 1.43 |
| | -0.7 |
| | -0.54 |
| | -1.33 |
| | 0.44 |
| | 0.04 |
| | -7.06 |
| | -1.64 |
| | 2.61 |
| | 2.52 |
| | 1.73 |
| | 1.18 |
| | 2.79 |
| | 4.15 |
| | 2.07 |
| | -1.13 |

| |
|---|
| -0.45 |
| 2.32 |
| 0.1 |
| 0.63 |
| 3.82 |
| 3.72 |
| 3.42 |
| 1.87 |
| 4.05 |
| 2.51 |
| -0.21 |
| 1.32 |
| 4.33 |
| 2.1 |
| 1.62 |
| -0.27 |
| 4.86 |
| 3.77 |
| 1.06 |
| 1.18 |
| 3.27 |
| 0.6 |
| 2.62 |
| 0.84 |
| 1.39 |
| -2.01 |
| -0.52 |
| 1.7 |
| 0.49 |
| 4 |
| 4.06 |
| 3.77 |
| 6.12 |
| 1.15 |
| 1.85 |

| |
|---|
| 0.91 |
| 2.49 |
| 6.92 |
| -1.75 |
| 1.02 |
| 1.98 |
| 1.48 |
| 3.21 |
| 5.36 |
| 5.84 |
| 3.87 |
| 4.16 |
| 1.76 |
| 5.74 |
| 4.81 |
| -2.13 |
| 2.41 |
| 3.63 |
| 7.39 |
| 1.42 |
| -2.47 |
| 0.85 |
| 4.46 |
| 0.36 |
| 1.54 |
| 1.07 |
| 0.13 |
| -1.24 |
| -2.54 |
| -1.27 |
| 3.01 |
| 1.91 |
| -0.23 |
| 1.33 |
| 6.58 |

| | |
|---|---|
| | -1.77 |
| | -0.41 |
| | 1.96 |
| | 0.01 |
| | -0.66 |
| | 5.07 |
| | -0.33 |
| | 2.41 |
| | 0.53 |
| | 0.4 |
| | 0.62 |
| | 3.36 |
| | 0.04 |
| | -0.55 |
| | -0.43 |
| | -1.46 |
| | -1.23 |
| | 2.24 |
| | 0.96 |
| | -2.53 |
| | 0.7 |
| | 0.98 |
| | -0.6 |
| | -2.84 |
| | -2.11 |
| | 2.29 |
| | -0.53 |
| | -2.34 |
| | 0.05 |
| | -5.16 |
| | -0.98 |
| | 1.31 |
| | -0.98 |
| | 1.9 |
| | 0.77 |

| | |
|---|---|
| | 1.77 |
| | 3.91 |
| | -0.88 |
| | -0.99 |
| | 0.82 |
| | 2.31 |
| | 2.82 |
| | 1.17 |
| | 2.77 |
| | 4 |
| | -0.34 |
| | -1.06 |
| | 0.52 |
| | 0.08 |
| | 3.49 |
| | 1.12 |
| | 2.48 |
| | 2.08 |
| | -4.11 |
| | 0.74 |
| | 5.04 |
| | 0.13 |
| | 4.01 |
| | 1.21 |
| | 2.24 |
| | -0.06 |
| | 2.55 |
| | -2.21 |
| | 1.18 |
| | 0.05 |
| | 2.74 |
| | -5.97 |
| | 1.54 |
| | -0.97 |
| | 0.89 |

| |
|---|
| 0.98 |
| 0.55 |
| -0.14 |
| 0.22 |
| -4.46 |
| 3.76 |
| 2.48 |
| 1.81 |
| 2.01 |
| -1.45 |
| 1.94 |
| -2.12 |
| 0.61 |
| 1.96 |
| -1.21 |
| 4.4 |
| 3.55 |
| 1.86 |
| 1.64 |
| 1.16 |
| -0.06 |
| 0.25 |
| 1.36 |
| 1.26 |
| -1.11 |
| 2.47 |
| 2.21 |
| -2.01 |
| 6.13 |
| 2.74 |
| -0.47 |
| 3.02 |
| 2.74 |
| 1.28 |
| 1.51 |

| |
|---|
| 0.99 |
| 0.85 |
| 0.17 |
| -4.19 |
| -2.46 |
| 0.11 |
| -0.23 |
| 4.02 |
| -0.56 |
| 0 |
| 5.62 |
| 1.77 |
| 0.58 |
| 0.84 |
| -2.53 |
| -1.17 |
| 0.08 |
| 4.7 |
| -1.09 |
| 0.84 |
| 4.34 |
| 2.2 |
| -0.96 |
| -0.66 |
| -2.04 |
| -1.57 |
| 1.59 |
| 1.42 |
| 7.85 |
| -1.61 |
| 0.09 |
| -1.04 |
| -0.57 |
| -1.8 |
| -1.4 |

| | |
|---|---|
| | 0.54 |
| | 1.19 |
| | 0.96 |
| | 1.98 |
| | 2.31 |
| | 1.65 |
| | 3.29 |
| | 4.79 |
| | -0.91 |
| | -0.41 |
| | 0.17 |
| | 1.04 |
| | -0.98 |
| | 1.53 |
| | 0.22 |
| | 6.74 |
| | 4.09 |
| | 5.96 |
| | 3.74 |
| | 2.3 |
| | 1.38 |
| | 2.69 |
| | 2.1 |
| | 0.49 |
| | 0.11 |
| | 2.87 |
| | -0.94 |
| | -1.69 |
| | 9.03 |
| | 6.04 |
| | 6.74 |
| | 5.44 |
| | 4.49 |
| | 7.59 |
| | 6.85 |

| | |
|---|---|
| | -1.22 |
| | -1.92 |
| | 4.09 |
| | 2.32 |
| | 1.26 |
| | -1.93 |
| | -1.72 |
| | -0.73 |
| | 0.64 |
| | -1.74 |
| | 0.46 |
| | -0.42 |
| | -5.93 |
| | 2.21 |
| | 1.92 |
| | 0.15 |
| | 2.16 |
| | -0.45 |
| | 0.16 |
| | 1.15 |
| | 1.85 |
| | 1.79 |
| | -0.25 |
| | 5.87 |
| | -0.93 |
| | 1.38 |
| | 0.26 |
| | -2.44 |
| | 1.35 |
| | 6.04 |
| | 1.16 |
| | -2.33 |
| | -0.59 |
| | 0.95 |
| | 1.67 |

| | |
|---|---|
| | 0.51 |
| | -1.29 |
| | 3.16 |
| | 0.38 |
| | 0.61 |
| | -0.06 |
| | 0.72 |
| | -1.05 |
| | 0.76 |
| | -0.66 |
| | -1.98 |
| | -0.75 |
| | -2.54 |
| | 1.73 |
| | -0.35 |
| | 0.47 |
| | -1.31 |
| | 1.51 |
| | 1.6 |
| | 0.01 |
| | 2.35 |
| | 0.51 |
| | 0.71 |
| | 1.63 |
| | 0.68 |
| | 1.66 |
| | 1.44 |
| | 3.02 |
| | 0.43 |
| | 2.79 |
| | -0.26 |
| | 6.13 |
| | 5.84 |
| | 2.76 |
| | -0.6 |

| | |
|---|---|
| | -2.76 |
| | -0.52 |
| | -0.83 |
| | 0.06 |
| | -0.7 |
| | -1.27 |
| | 6.54 |
| | -2.5 |
| | 0.58 |
| | -0.79 |
| | -1.64 |
| | 2.26 |
| | -1.03 |
| | 1.07 |
| | -1.37 |
| | 0.98 |
| | -3.66 |
| | 2.48 |
| | 1.74 |
| | -0.98 |
| | -2.29 |
| | -0.66 |
| | -0.04 |
| | -1.33 |
| | 1.72 |
| | -0.1 |
| | -0.84 |
| | 0.5 |
| | 0.25 |
| | -1.33 |
| | -1.24 |
| | 0.91 |
| | -2.97 |
| | -5.51 |
| | -3.45 |

| | |
|---|---|
| | 1.13 |
| | -3.62 |
| | -2.74 |
| | -2.3 |
| | -4.48 |
| | 1.99 |
| | 0.16 |
| | -1.07 |
| | -0.9 |
| | -0.61 |
| | -0.65 |
| | -0.39 |
| | 0.95 |
| | -0.41 |
| | 1.18 |
| | 1.86 |
| | 0.57 |
| | -1.54 |
| | 1.56 |
| | 1.26 |
| | 0.42 |
| | -2.18 |
| | 0.02 |
| | -1.31 |
| | -4.21 |
| | -2.03 |
| | -0.54 |
| | -2.31 |
| | -1.76 |
| | -4.28 |
| | -1.32 |
| | -1.47 |
| | -0.51 |
| | -1.72 |
| | -1.02 |

| | |
|---|---|
| | -0.96 |
| | -5.62 |
| | -5.35 |
| | -0.37 |
| | 1.44 |
| | 0.97 |
| | -0.6 |
| | -3 |
| | -1.11 |
| | -2.36 |
| | -3.57 |
| | -0.52 |
| | -0.2 |
| | -2.52 |
| | -1.31 |
| | 0.71 |
| | -1.47 |
| | -0.69 |
| | -1.62 |
| | -1.87 |
| | 3.37 |
| | -0.5 |
| | -0.88 |
| | 0.77 |
| | -0.68 |
| | -0.67 |
| | -0.54 |
| | -1.06 |
| | 0.98 |
| | 1.35 |
| | -0.35 |
| | 0.81 |
| | -0.6 |
| | -2.72 |
| | -0.39 |

| | |
|---|---|
| | 0.85 |
| | -0.4 |
| | -0.89 |
| | 0.72 |
| | 1.78 |
| | -1.39 |
| | -0.45 |
| | -0.55 |
| | -1.34 |
| | -2.25 |
| | -3.46 |
| | -2.72 |
| | -3.02 |
| | -2.55 |
| | -4.21 |
| | -3.68 |
| | -2.16 |
| | -3.66 |
| | 0.77 |
| | 0.46 |
| | -3.14 |
| | 0.57 |
| | -0.67 |
| | 2.63 |
| | -0.56 |
| | 3.63 |
| | 0.87 |
| | 3.01 |
| | 1.98 |
| | -0.66 |
| | -0.77 |
| | -2.5 |
| | -0.72 |
| | 1.5 |
| | 1.76 |

| |
|---:|
| 3.35 |
| -2 |
| 3.53 |
| 6.47 |
| 3.77 |
| 0.23 |
| 2.21 |
| 1.99 |
| -3.1 |
| 4.22 |
| 5.47 |
| -4.1 |
| -3.79 |
| 4.48 |
| -2 |

| |
|---|
| Log2 Fold Change: (SM_pos_12022024_Newanalyis) / (Rabbitserum_12022024_Newanalysis) |
| -1.13 |
| -3.17 |
| -4.59 |
| -2.41 |
| -4.1 |
| -1.1 |
| -1.54 |
| -2.57 |
| 0.73 |
| 2.64 |
| 2.67 |
| 0.27 |
| -1.14 |
| 1.53 |
| 1.82 |
| 0.77 |
| 1.75 |
| -5.99 |
| -5.64 |
| -5.84 |
| -1.06 |
| -4.99 |
| -3.1 |
| -5.78 |
| -5.34 |
| 2.07 |
| 0.16 |
| 1.2 |
| -1.62 |
| 5.26 |
| 0.86 |
| -0.03 |
| 5.44 |

| | |
|---|---|
| | 3.63 |
| | -2.52 |
| | -4.44 |
| | -5.79 |
| | -0.08 |
| | -1.58 |
| | -6.03 |
| | 4.33 |
| | -0.18 |
| | 0.53 |
| | -0.84 |
| | 4.27 |
| | -3.8 |
| | -3.62 |
| | -5.26 |
| | -5.65 |
| | -1.97 |
| | -5.96 |
| | -2.84 |
| | -4.15 |
| | -2.99 |
| | -2.83 |
| | 2.5 |
| | -0.45 |
| | 2.45 |
| | -1.07 |
| | -3.67 |
| | 0.86 |
| | 4.03 |
| | -3.2 |
| | -3.13 |
| | -3.7 |
| | 2.82 |
| | -0.45 |
| | 5.76 |

| | |
|---|---|
| | 0.61 |
| | 0.15 |
| | 2.67 |
| | 5.23 |
| | -0.19 |
| | 5.48 |
| | 0.19 |
| | -1.79 |
| | -3.51 |
| | -3.86 |
| | -6.74 |
| | -9.43 |
| | -4.77 |
| | 3.07 |
| | 3.89 |
| | 0.38 |
| | -3.72 |
| | 1.68 |
| | 1.18 |
| | -5.14 |
| | 3.15 |
| | -0.67 |
| | -10.31 |
| | -6.58 |
| | -2.25 |
| | -2.27 |
| | -3.34 |
| | -1.29 |
| | -3.91 |
| | -2.01 |
| | -1.87 |
| | -1.1 |
| | -4.3 |
| | 4.58 |
| | 2.85 |

| |
|---|
| 1.19 |
| -3.45 |
| 3.69 |
| 0.72 |
| 1.36 |
| -1.03 |
| 2.63 |
| 1.47 |
| -3.98 |
| -1.18 |
| 1.16 |
| 3.71 |
| 0.51 |
| 3.16 |
| -6.97 |
| -6.99 |
| -2.35 |
| 3.58 |
| -5.6 |
| -3.42 |
| -0.13 |
| 3.02 |
| -2.05 |
| -0.21 |
| -3.69 |
| 0.39 |
| 0.47 |
| -1.15 |
| -2.5 |
| -9.84 |
| -3.51 |
| -4.85 |
| -1.17 |
| -1.84 |
| 0.09 |

| | |
|---|---|
| | 3.88 |
| | -0.16 |
| | -1.61 |
| | 1.21 |
| | 0.47 |
| | 0.29 |
| | -0.11 |
| | 1.06 |
| | 0.13 |
| | -0.61 |
| | 0.69 |
| | -2.12 |
| | -4.16 |
| | -2.69 |
| | -3.67 |
| | -2.31 |
| | 3.35 |
| | 0.13 |
| | -0.89 |
| | -1.32 |
| | 0.08 |
| | -1.87 |
| | -3.95 |
| | -2.68 |
| | 3.03 |
| | -8.5 |
| | -5.18 |
| | 0.07 |
| | -0.71 |
| | -3.26 |
| | -2.77 |
| | -0.54 |
| | -5.99 |
| | 0.28 |
| | -2.57 |

| | |
|---|---|
| | 0.82 |
| | -6.21 |
| | -5.77 |
| | -3.98 |
| | -2.83 |
| | -4.31 |
| | -0.72 |
| | -1.51 |
| | 1.44 |
| | -2.04 |
| | -5.38 |
| | 3.03 |
| | -3.91 |
| | -1.6 |
| | 0.64 |
| | -1.65 |
| | -2.2 |
| | 2.8 |
| | -2.08 |
| | -2.88 |
| | -1.99 |
| | -2.01 |
| | -2.25 |
| | 1.81 |
| | -1.27 |
| | -1.02 |
| | -4.09 |
| | 3.11 |
| | -1.7 |
| | -10.13 |
| | 0.99 |
| | -5.92 |
| | -2.8 |
| | -7.22 |
| | 1.46 |

| |
|---|
| -1.55 |
| -0.22 |
| 0.67 |
| 1.49 |
| -0.39 |
| 5.46 |
| 3.11 |
| -0.31 |
| 2.99 |
| 0.68 |
| 0.41 |
| 0.61 |
| 0.21 |
| 0.97 |
| -0.4 |
| -1.85 |
| -3.64 |
| -6.35 |
| -2.57 |
| -2.11 |
| -1.54 |
| -3.05 |
| -0.38 |
| 0.35 |
| -2.81 |
| -6.47 |
| 2.13 |
| -1.29 |
| -0.52 |
| -2.74 |
| -0.99 |
| 1.11 |
| -1.71 |
| -1.51 |
| -3.72 |

| | |
|---|---|
| | -0.42 |
| | -1.43 |
| | 0.76 |
| | -0.24 |
| | -3.93 |
| | -0.41 |
| | -2.04 |
| | 0.28 |
| | 0.25 |
| | -2.79 |
| | -1.31 |
| | -0.77 |
| | -2.98 |
| | -2.35 |
| | -0.44 |
| | -1.12 |
| | -3.63 |
| | 0.26 |
| | 1.47 |
| | -7.2 |
| | -1.25 |
| | 3.05 |
| | -2.87 |
| | 4.09 |
| | -4.25 |
| | 0.41 |
| | -1.52 |
| | -0.66 |
| | -0.32 |
| | -1.51 |
| | -0.4 |
| | -3.12 |
| | 0.34 |
| | -1.06 |
| | -2.33 |

| | |
|---|---|
| | -2.07 |
| | -5.76 |
| | -0.79 |
| | -0.99 |
| | 2.95 |
| | -1.2 |
| | -3.25 |
| | -0.1 |
| | 2.27 |
| | -3.29 |
| | 1.51 |
| | -3.18 |
| | 0.34 |
| | 0.76 |
| | 2.96 |
| | 1.65 |
| | -6.77 |
| | -2.43 |
| | 0.87 |
| | -2.19 |
| | -4.66 |
| | -5.57 |
| | 1.37 |
| | -3.81 |
| | -3.75 |
| | -3.99 |
| | -2.86 |
| | -2.98 |
| | -2.97 |
| | -2.23 |
| | -4.65 |
| | -2.44 |
| | -0.18 |
| | 2.92 |
| | -0.21 |

| | |
|---|---|
| | -2.65 |
| | -2.56 |
| | -2.76 |
| | -6.91 |
| | 2.21 |
| | -2 |
| | -1.43 |
| | -4.68 |
| | 5.54 |
| | -2.07 |
| | -0.44 |
| | 2.11 |
| | 1.04 |
| | 0.67 |
| | 4.28 |
| | -2.03 |
| | -2.19 |
| | -4.38 |
| | -4.18 |
| | -3.06 |
| | 2.83 |
| | -1.36 |
| | 0.84 |
| | -4.73 |
| | -2.97 |
| | 3.72 |
| | 2.98 |
| | -3.37 |
| | 0.98 |
| | -2.77 |
| | -2.62 |
| | -1.02 |
| | -2.13 |
| | -0.14 |
| | -0.8 |

| | |
|---|---|
| | -2.09 |
| | -2.01 |
| | -0.34 |
| | 3.18 |
| | -4.04 |
| | -0.35 |
| | -3.36 |
| | -3.16 |
| | -2.4 |
| | -1.91 |
| | -3.6 |
| | -2.84 |
| | -3.13 |
| | -2.59 |
| | 0.81 |
| | -5.55 |
| | -2.02 |
| | -3.01 |
| | -1.99 |
| | -2.47 |
| | -2.66 |
| | -2.4 |
| | 2.66 |
| | -3.93 |
| | -2.89 |
| | 0.37 |
| | -2.47 |
| | -3.5 |
| | -4.84 |
| | -0.63 |
| | -2.46 |
| | -6.83 |
| | -4.27 |
| | -3.49 |
| | 3.56 |

| | |
|---|---|
| | -0.06 |
| | 3.57 |
| | -4 |
| | -2.35 |
| | -5.15 |
| | -3.64 |
| | -3.29 |
| | -3.46 |
| | -6.83 |
| | -3.14 |
| | -2.76 |
| | -5.65 |
| | -4.14 |
| | -6.45 |
| | -0.78 |
| | -2.36 |
| | -2.92 |
| | -1.55 |
| | -3.16 |
| | 1.02 |
| | 1.22 |
| | 0.21 |
| | -2.7 |
| | -3.72 |
| | 0.95 |
| | 0.62 |
| | -10.65 |
| | 2.48 |
| | 0.27 |
| | -3.15 |
| | -7.41 |
| | 0.64 |
| | -1.41 |
| | -0.94 |
| | 2.02 |

| |
|---|
| -3.13 |
| 3.63 |
| 0.38 |
| 0.86 |
| 3 |
| 0.42 |
| 5.21 |
| -2.1 |
| -3.29 |
| -3.76 |
| -2.29 |
| -2.58 |
| -3.54 |
| 1.85 |
| -2.27 |
| 0.27 |
| -6.06 |
| -3.18 |
| 0.14 |
| -0.52 |
| -2.44 |
| -3.02 |
| -1.4 |
| 0.47 |
| 1.46 |
| 2.7 |
| -2.16 |
| -7.16 |
| -0.24 |
| -6.13 |
| -1.08 |
| -1.56 |
| -2.27 |
| -4.15 |
| -2.77 |

| | |
|---|---|
| | -7.72 |
| | -5.47 |
| | -3.48 |
| | 2.32 |
| | 1.54 |
| | -0.58 |
| | -0.82 |
| | -0.16 |
| | -3.06 |
| | -3.55 |
| | -1.32 |
| | 3.04 |
| | -4.03 |
| | 1.81 |
| | -2.66 |
| | 0.15 |
| | -2.29 |
| | 2.74 |
| | -0.71 |
| | -2.66 |
| | -4.55 |
| | -0.49 |
| | -4.12 |
| | -3.53 |
| | -3.26 |
| | -2.55 |
| | 0.85 |
| | -3.68 |
| | -1.19 |
| | -1.04 |
| | -2.99 |
| | -2.57 |
| | -2.81 |
| | -0.03 |
| | 2.06 |

| | |
|---|---|
| | 1.89 |
| | -1.88 |
| | 1.75 |
| | -2.56 |
| | -1.02 |
| | -2.78 |
| | 2.48 |
| | -2.42 |
| | 4.2 |
| | 3.21 |
| | 3.27 |
| | -2.09 |
| | -4.82 |
| | 1.25 |
| | -1.7 |
| | -2.49 |
| | -1.98 |
| | -0.27 |
| | -3.31 |
| | 1.86 |
| | -0.88 |
| | 2.18 |
| | 5.09 |
| | -0.21 |
| | -0.79 |
| | -3.89 |
| | -0.28 |
| | -6.97 |
| | -3.31 |
| | -2.69 |
| | -0.85 |
| | -2.94 |
| | -4.12 |
| | -0.95 |
| | -2.96 |

| | |
|---|---|
| | 1.02 |
| | 0.1 |
| | -0.45 |
| | 1.58 |
| | 2.19 |
| | -2.86 |
| | -4.28 |
| | 0.02 |
| | -7.61 |
| | -3.36 |
| | -1.26 |
| | -1.53 |
| | -1.12 |
| | -1.12 |
| | -4.23 |
| | 0.46 |
| | -2.13 |
| | 2.19 |
| | -2.64 |
| | 1.07 |
| | -1.59 |
| | -10.08 |
| | 3.21 |
| | -3.75 |
| | 2.05 |
| | -3.44 |
| | -7.58 |
| | 0.56 |
| | 4.54 |
| | 1.75 |
| | -1.07 |
| | -2.55 |
| | 1.82 |
| | -1.33 |
| | -4.12 |

| | |
|---|---|
| | -5.18 |
| | -1.38 |
| | -3.33 |
| | -0.1 |
| | 1.56 |
| | -0.19 |
| | 1.21 |
| | -0.86 |
| | -1.92 |
| | 2.66 |
| | -0.83 |
| | -1 |
| | -2.02 |
| | -4 |
| | -1.16 |
| | -2.29 |
| | -5.76 |
| | 2.06 |
| | 0.78 |
| | 0.75 |
| | -3.09 |
| | -3.35 |
| | 0.8 |
| | 3.85 |
| | -1.7 |
| | -0.39 |
| | 5.11 |
| | 0.55 |
| | 1.31 |
| | 3.75 |
| | -0.67 |
| | -0.84 |
| | -2.36 |
| | 0.53 |
| | 1.79 |

| | |
|---|---|
| | 3.33 |
| | -0.85 |
| | 1.26 |
| | 1.32 |
| | -1.85 |
| | -2.93 |
| | -5.29 |
| | -3.61 |
| | -3.2 |
| | 5.11 |
| | -3.42 |
| | -2.16 |
| | -4.66 |
| | 0.66 |
| | -3.52 |
| | 2.44 |
| | -1.59 |
| | -1.21 |
| | 3.11 |
| | -1.02 |
| | 0.49 |
| | 2.23 |
| | -0.3 |
| | 3.86 |
| | 3.87 |
| | -0.04 |
| | 3.88 |
| | -4.39 |
| | -0.56 |
| | 0.13 |
| | -3.8 |
| | 0.13 |
| | -3.12 |
| | 4.63 |
| | -3.79 |

| | |
|---|---|
| | 3.22 |
| | -5.09 |
| | -6.74 |
| | -0.73 |
| | -8.23 |
| | 2.14 |
| | 0.65 |
| | 3.81 |
| | 0.92 |
| | -2.26 |
| | -5.61 |
| | -0.09 |
| | -0.97 |
| | -2.58 |
| | 3.84 |
| | 2.33 |
| | -0.94 |
| | 1.37 |
| | -1.05 |
| | 0.6 |
| | -1.93 |
| | 0.36 |
| | 0.25 |
| | -1.34 |
| | 0.51 |
| | 0.76 |
| | 0.7 |
| | 3.69 |
| | 0.42 |
| | 3.14 |
| | 3.32 |
| | 2.05 |
| | -0.88 |
| | -5.75 |
| | -0.46 |

| | |
|---|---|
| | 2.42 |
| | 0.53 |
| | 6.42 |
| | -0.65 |
| | -3.81 |
| | -4.52 |
| | -0.44 |
| | -6.8 |
| | -0.44 |
| | -2.96 |
| | -9.35 |
| | -7.9 |
| | -6.49 |
| | -4.33 |
| | -3.69 |
| | -2.54 |
| | -4.28 |
| | 2.54 |
| | 1.97 |
| | -1.29 |
| | -1.23 |
| | -0.67 |
| | -2.72 |
| | -2.88 |
| | -2.56 |
| | -1.68 |
| | -4.28 |
| | 0.46 |
| | 0.58 |
| | 1.71 |
| | -7.04 |
| | -1.32 |
| | 0.08 |
| | 1.53 |
| | -1.46 |

| | |
|---|---|
| | -1.48 |
| | 0.82 |
| | -0.6 |
| | 1.26 |
| | -2.5 |
| | -3.28 |
| | -3.51 |
| | 0.76 |
| | 1.9 |
| | 0.37 |
| | 0.53 |
| | 1.87 |
| | -0.6 |
| | 0.6 |
| | 2.56 |

| |
|---|
| Log2 Fold Change: (Bb_pos_12022024_Newanalysis) / (SM_pos_12022024_Newanalyis) |
| -1.68 |
| 3.58 |
| -0.19 |
| -5.91 |
| 3.11 |
| 1.37 |
| 4.21 |
| -0.69 |
| -2.34 |
| -1.75 |
| -2.91 |
| 1.07 |
| -3.45 |
| -2.57 |
| -0.55 |
| 2.5 |
| -1.66 |
| -0.82 |
| 1.21 |
| 0.47 |
| 1.18 |
| -1.54 |
| -2.35 |
| -1.18 |
| -2.31 |
| -2.28 |
| -0.06 |
| -0.01 |
| 3.16 |
| -4.85 |
| 0.69 |
| 5.83 |
| -2.54 |

| | |
|---|---|
| | -1.77 |
| | -3.62 |
| | 0.48 |
| | -0.44 |
| | 6.6 |
| | 7.19 |
| | -1.45 |
| | -3.99 |
| | -1.91 |
| | -4.02 |
| | -1.38 |
| | -3.28 |
| | -2.18 |
| | -3.07 |
| | -1.01 |
| | 0.26 |
| | -1.61 |
| | 0.16 |
| | -1.61 |
| | -2.75 |
| | -1.47 |
| | 2.78 |
| | -4.36 |
| | -1.26 |
| | -0.52 |
| | -1.66 |
| | 3.13 |
| | -4.21 |
| | -4.03 |
| | 4.48 |
| | -4.1 |
| | -0.52 |
| | -3.13 |
| | -3.49 |
| | -10.8 |

| | |
|---|---|
| | -5.21 |
| | -1.97 |
| | -5.5 |
| | -5.05 |
| | -3.82 |
| | -11.98 |
| | -3.79 |
| | 2.45 |
| | -3.69 |
| | 1.02 |
| | -0.1 |
| | 2.26 |
| | -1.4 |
| | -6.2 |
| | -3.02 |
| | -4.34 |
| | -3.84 |
| | -4.05 |
| | -0.03 |
| | -1.32 |
| | -5.99 |
| | -1.47 |
| | 3.91 |
| | 0.31 |
| | -2.18 |
| | -2.2 |
| | -1.18 |
| | -3.32 |
| | -2.09 |
| | -3.43 |
| | 1.1 |
| | -0.82 |
| | -1.61 |
| | -3.56 |
| | -6.3 |

| | |
|---|---|
| | -0.79 |
| | -0.85 |
| | -2.18 |
| | -4.24 |
| | -2.37 |
| | 0.39 |
| | -1.5 |
| | -4.16 |
| | 0.28 |
| | -1.32 |
| | -5.35 |
| | -3.14 |
| | -3.7 |
| | -4.16 |
| | -0.42 |
| | 1.44 |
| | 0.71 |
| | -3.43 |
| | -0.87 |
| | 1.65 |
| | 1.82 |
| | -4.82 |
| | 0.29 |
| | -3.34 |
| | 2.62 |
| | -3.16 |
| | 0.49 |
| | 0.59 |
| | -1.02 |
| | 2.75 |
| | 0.2 |
| | 4.3 |
| | 3.16 |
| | -1.83 |
| | -1.71 |

| | |
|---|---|
| | -4.22 |
| | -1.11 |
| | 3.13 |
| | -1.78 |
| | 1 |
| | 1.12 |
| | 1.15 |
| | -1.26 |
| | 2.77 |
| | -0.41 |
| | 0.28 |
| | -1.97 |
| | -2.97 |
| | 1.16 |
| | -0.89 |
| | -3.41 |
| | -0.68 |
| | -2.08 |
| | 3.94 |
| | -0.32 |
| | -3.7 |
| | -2.37 |
| | -1.13 |
| | -2.56 |
| | -4.56 |
| | 1.13 |
| | 4.78 |
| | -2.69 |
| | -3.18 |
| | 2.23 |
| | -2.34 |
| | -3.62 |
| | 4.78 |
| | -3.24 |
| | -3.15 |

| |
|---|
| -0.26 |
| 2.1 |
| -0.17 |
| -2.5 |
| -2.68 |
| -0.66 |
| -5.23 |
| -2.12 |
| 0.66 |
| -5.16 |
| 4.16 |
| -2.88 |
| -3.25 |
| -2.43 |
| 1.75 |
| -2.41 |
| -1.32 |
| -6.44 |
| -4.32 |
| -1.37 |
| -2 |
| -1.74 |
| 2.33 |
| -1.67 |
| 2.66 |
| 0.08 |
| 0.11 |
| -3.69 |
| -4.05 |
| 2.28 |
| -6.77 |
| 0.03 |
| -2.75 |
| 2.13 |
| -7.68 |

| | |
|---|---|
| | -3.43 |
| | -1.86 |
| | -2.32 |
| | -1.78 |
| | 1.6 |
| | -5.93 |
| | -5.5 |
| | -2.51 |
| | -4.22 |
| | -1.39 |
| | 0.92 |
| | -1.24 |
| | -2.98 |
| | -1.91 |
| | 2.34 |
| | 4.24 |
| | 0.55 |
| | 2.43 |
| | -0.46 |
| | 1.92 |
| | -3 |
| | 0.73 |
| | 1.46 |
| | -4.93 |
| | -2.2 |
| | 0.39 |
| | -2.44 |
| | 0.77 |
| | 2.1 |
| | 1.88 |
| | -0.21 |
| | -1.05 |
| | 0.31 |
| | 0.35 |
| | -1.27 |

| | |
|---|---|
| | 0.52 |
| | -0.47 |
| | -0.23 |
| | -0.83 |
| | 1.95 |
| | -2.85 |
| | -1.24 |
| | -1.03 |
| | -2.11 |
| | -3.87 |
| | 1.47 |
| | -0.3 |
| | 0.88 |
| | -0.98 |
| | 2.28 |
| | 0.54 |
| | 4.04 |
| | 0.36 |
| | -0.87 |
| | -0.45 |
| | -2.12 |
| | -2.98 |
| | 2.02 |
| | -3.95 |
| | 3.57 |
| | -0.54 |
| | -0.93 |
| | 0.43 |
| | -1.53 |
| | -1.1 |
| | -2.39 |
| | -2.47 |
| | 1.74 |
| | -2.14 |
| | -3.55 |

| | |
|---|---|
| | -1.44 |
| | -0.66 |
| | -2.03 |
| | -2.88 |
| | -6.13 |
| | -1.83 |
| | -2.79 |
| | -0.27 |
| | -3.7 |
| | -1.3 |
| | -1.25 |
| | 0.44 |
| | 0.52 |
| | -1.91 |
| | -3.83 |
| | -1.54 |
| | -0.25 |
| | 0.3 |
| | -3.08 |
| | -0.16 |
| | -1.53 |
| | 3.57 |
| | -2.44 |
| | -2.48 |
| | -0.22 |
| | 0.25 |
| | -0.48 |
| | -1.13 |
| | 1.82 |
| | -2.17 |
| | 0.39 |
| | -3.1 |
| | 1.41 |
| | -6.57 |
| | -1.5 |

| | |
|---|---|
| | -2.17 |
| | 3.42 |
| | -1.1 |
| | 2.3 |
| | 0.46 |
| | 4.32 |
| | -3.46 |
| | -2.3 |
| | -6.02 |
| | -3.1 |
| | -1.61 |
| | 0.63 |
| | -3.78 |
| | -3.51 |
| | -2.78 |
| | 0.78 |
| | -0.27 |
| | -1.23 |
| | -2.1 |
| | 0.56 |
| | -4.52 |
| | 1.14 |
| | -3.36 |
| | -1.77 |
| | -1.21 |
| | -1.67 |
| | 0.16 |
| | 4.33 |
| | -2.53 |
| | 6.37 |
| | 6.18 |
| | -3.77 |
| | 1.63 |
| | 4.44 |
| | 2.67 |

| | |
|---|---|
| | 3.37 |
| | 0.91 |
| | 6.19 |
| | 1.37 |
| | -0.22 |
| | -0.22 |
| | 0.61 |
| | -2.08 |
| | -0.54 |
| | -1.73 |
| | 2.69 |
| | -0.91 |
| | -0.31 |
| | 4.11 |
| | -1.41 |
| | 2.1 |
| | 1.68 |
| | 0.45 |
| | -2.9 |
| | -0.16 |
| | -3.26 |
| | -3.68 |
| | -2.07 |
| | -0.54 |
| | -3.15 |
| | 0.34 |
| | -1.65 |
| | -0.66 |
| | -0.04 |
| | -4.75 |
| | -0.75 |
| | 2.55 |
| | 0.75 |
| | -1.2 |
| | -4.33 |

| | |
|---|---|
| | -2.09 |
| | -5.05 |
| | 2.5 |
| | 0.01 |
| | 2.66 |
| | -0.16 |
| | -0.1 |
| | -1 |
| | 2.51 |
| | 3.6 |
| | -0.86 |
| | -0.19 |
| | 0.46 |
| | -0.75 |
| | -6.56 |
| | -0.25 |
| | -1.25 |
| | -3.91 |
| | -3.96 |
| | -5.22 |
| | 0.64 |
| | -2.55 |
| | 0.45 |
| | -0.44 |
| | -2.39 |
| | 1.14 |
| | 9.18 |
| | -7.14 |
| | 0.63 |
| | -1.44 |
| | -0.46 |
| | 4.47 |
| | -3.91 |
| | -4.75 |
| | -6.84 |

| | |
|---|---|
| | -2.19 |
| | -8.19 |
| | -4.29 |
| | -8.63 |
| | -5.47 |
| | -0.27 |
| | -3.15 |
| | 0.85 |
| | -0.8 |
| | 0.94 |
| | 0.88 |
| | -3.06 |
| | -4.35 |
| | -4.81 |
| | -0.29 |
| | -3.26 |
| | 5.83 |
| | 2.47 |
| | -4.57 |
| | -7.17 |
| | -0.39 |
| | -6.33 |
| | -3.42 |
| | -5.68 |
| | -3.74 |
| | -6.73 |
| | -1.83 |
| | 0.18 |
| | -2.28 |
| | 7.42 |
| | -1.03 |
| | 0.3 |
| | -3.06 |
| | -4.05 |
| | -4.6 |

| | |
|---|---|
| | 0.55 |
| | 0.57 |
| | -0.51 |
| | -4.07 |
| | -1.73 |
| | -0.09 |
| | -5.97 |
| | -4.4 |
| | 1.56 |
| | 0.7 |
| | -2.81 |
| | -2.05 |
| | 6.41 |
| | 3.35 |
| | 6.29 |
| | 5.69 |
| | 7.68 |
| | 3.98 |
| | -1.64 |
| | -0.31 |
| | 2.86 |
| | -2.81 |
| | 2.53 |
| | 0.95 |
| | 3.67 |
| | 5.88 |
| | 3.91 |
| | -0.01 |
| | 0.56 |
| | -0.37 |
| | 0.54 |
| | 4.84 |
| | 4.17 |
| | 6.82 |
| | 1.05 |

| | |
|---|---|
| | -1.07 |
| | 4.4 |
| | -4.29 |
| | -2.39 |
| | 1.3 |
| | -2.16 |
| | 0.92 |
| | 2.5 |
| | -1.6 |
| | -1.07 |
| | -1.15 |
| | -3.31 |
| | -0.58 |
| | -3.93 |
| | -1.46 |
| | -2.03 |
| | 2.31 |
| | -1.79 |
| | 5.91 |
| | -4.16 |
| | -3.45 |
| | -3.22 |
| | -5.9 |
| | 0.32 |
| | -2.92 |
| | -1.44 |
| | 0.53 |
| | 3.24 |
| | -0.4 |
| | 0.56 |
| | -3.75 |
| | -3.22 |
| | -1.53 |
| | 2.66 |
| | -2.36 |

| | |
|---|---|
| | -5.15 |
| | -2.72 |
| | -4.4 |
| | 1.17 |
| | -9.46 |
| | 0.55 |
| | -2.11 |
| | -0.59 |
| | -2.17 |
| | -3.56 |
| | -1.79 |
| | -2.92 |
| | -1.19 |
| | 3.83 |
| | -0.87 |
| | -2.53 |
| | -3.02 |
| | -10.07 |
| | -3.3 |
| | 1.71 |
| | -7.69 |
| | 3.52 |
| | -12.58 |
| | 0.9 |
| | -6.7 |
| | 0.79 |
| | -2.03 |
| | -2.88 |
| | -7.83 |
| | -10.73 |
| | -1.3 |
| | -0.2 |
| | -0.73 |
| | 5.68 |
| | 4.96 |

| | |
|---|---|
| | 6.67 |
| | 0.75 |
| | 3.83 |
| | 2.86 |
| | 2.55 |
| | 1.7 |
| | -1.79 |
| | -2.4 |
| | -1.54 |
| | -1.76 |
| | 2.21 |
| | -0.11 |
| | 6.44 |
| | -0.89 |
| | -1.88 |
| | -3.4 |
| | 5.02 |
| | 0.76 |
| | -0.36 |
| | 0.39 |
| | 6.08 |
| | -2.86 |
| | -1.78 |
| | -0.28 |
| | 3.77 |
| | 3.79 |
| | -1.54 |
| | 2.12 |
| | 1.92 |
| | -5.02 |
| | 0.45 |
| | -2.06 |
| | 0.2 |
| | -0.03 |
| | -1.99 |

| | |
|---|---|
| | -0.01 |
| | 1.51 |
| | 0.66 |
| | 1.08 |
| | 2.52 |
| | 1.75 |
| | -2.12 |
| | -1.1 |
| | -2.38 |
| | -4.26 |
| | 1.55 |
| | 4.83 |
| | -0.31 |
| | -1.75 |
| | -3.58 |
| | 2.54 |
| | 3.74 |
| | 1.3 |
| | -1.13 |
| | 1.69 |
| | 0.64 |
| | 2.43 |
| | -1.07 |
| | -3.45 |
| | -1.23 |
| | 1.3 |
| | -1.9 |
| | -0.52 |
| | -0.78 |
| | -0.87 |
| | -1.27 |
| | -2.37 |
| | -0.15 |
| | -2.76 |
| | 1.19 |

| | |
|---|---|
| | -5.28 |
| | 0.18 |
| | -0.77 |
| | -3.12 |
| | 2.09 |
| | -5.8 |
| | -4.39 |
| | -9.88 |
| | -7.33 |
| | -1.57 |
| | 3.45 |
| | 1.74 |
| | -2.49 |
| | 2.27 |
| | -1.03 |
| | -3.25 |
| | 1.63 |
| | 3.84 |
| | 3.9 |
| | 2.43 |
| | 2.39 |
| | 0.6 |
| | 0.69 |
| | -1.08 |
| | 0.3 |
| | 0.37 |
| | 0.2 |
| | -2.16 |
| | -1.69 |
| | -2.9 |
| | -1.88 |
| | -1.61 |
| | -2.66 |
| | 1.59 |
| | -3.37 |

| | |
|---|---|
| | -3.93 |
| | -0.18 |
| | -6.26 |
| | -2.56 |
| | -0.31 |
| | 1.18 |
| | -2.05 |
| | 0.32 |
| | -1.09 |
| | -4.04 |
| | 2.43 |
| | 0.9 |
| | -0.6 |
| | 0.69 |
| | 0.63 |
| | -2.5 |
| | 1.13 |
| | -2.66 |
| | -1.22 |
| | 1.22 |
| | 4.02 |
| | -0.07 |
| | 7.05 |
| | 1.9 |
| | 1.3 |
| | -1.6 |
| | 6.9 |
| | 2.72 |
| | -3.24 |
| | -4.17 |
| | 1.93 |
| | -2.23 |
| | 1.36 |
| | -1.02 |
| | -3.03 |

| | |
|---|---|
| | 0.7 |
| | 0.77 |
| | -0.31 |
| | -2.19 |
| | 0.87 |
| | -3.24 |
| | -1.5 |
| | -0.3 |
| | -3.37 |
| | -2.79 |
| | 0.07 |
| | -0.22 |
| | 1.63 |
| | 1.17 |
| | -13.84 |

| |
|---|
| Log2 Fold Change: (CM_pos_12022024_Newanalysis) / (SM_pos_12022024_Newanalyis) |
| -1.09 |
| 2.55 |
| -0.54 |
| -0.65 |
| 2.2 |
| 1.49 |
| 3.42 |
| 1.43 |
| -2.7 |
| -0.93 |
| -1.9 |
| 0.75 |
| -1.5 |
| -1.19 |
| -0.69 |
| 1.4 |
| -2.11 |
| 3.88 |
| 2.67 |
| 2.2 |
| 1.06 |
| 2.74 |
| 1.74 |
| 2.29 |
| 3.59 |
| -0.86 |
| -0.22 |
| -0.85 |
| 2.52 |
| -4.06 |
| 0.24 |
| -0.69 |
| -4.06 |

| | |
|---|---|
| | -2.5 |
| | 1.53 |
| | -1.49 |
| | 2.34 |
| | 0.02 |
| | 0.65 |
| | 4.63 |
| | -4.19 |
| | -2.33 |
| | -0.57 |
| | -2.42 |
| | -3.74 |
| | 2.04 |
| | 1.6 |
| | 3.52 |
| | 4.37 |
| | 1.17 |
| | 4.89 |
| | 2.16 |
| | 2.54 |
| | 1.27 |
| | 1.6 |
| | -3.23 |
| | -2.54 |
| | -2.22 |
| | -1.05 |
| | 1.73 |
| | -3.88 |
| | -4.73 |
| | 3.26 |
| | 1.89 |
| | 2.89 |
| | -3.2 |
| | 1.57 |
| | -8.52 |

| |
|---|
| -4.79 |
| -0.92 |
| -0.71 |
| -5.6 |
| -1.1 |
| -8.18 |
| -3.43 |
| 1.64 |
| 1.8 |
| 0.74 |
| 4.6 |
| 5.71 |
| 2.87 |
| -1.62 |
| -3.28 |
| -1.37 |
| -2.45 |
| -1.57 |
| -0.27 |
| 4.98 |
| -3.63 |
| -0.29 |
| 9.26 |
| 4.62 |
| 0.21 |
| 0.44 |
| 1.31 |
| -1.27 |
| 2.89 |
| 0.32 |
| 1.48 |
| 1.13 |
| 0.44 |
| -2.39 |
| -3.8 |

| |
|---|
| -0.53 |
| 1.44 |
| -2.94 |
| -1.68 |
| -1.91 |
| -0.48 |
| -0.99 |
| -2.16 |
| 0.97 |
| 2.55 |
| -2.62 |
| -3.65 |
| -1.64 |
| -3.25 |
| 6 |
| 5.51 |
| 0.73 |
| -4.19 |
| -0.77 |
| 1.47 |
| 1.02 |
| -4.38 |
| 0.9 |
| -0.69 |
| 3.08 |
| -3.2 |
| -0.06 |
| -0.08 |
| 0.82 |
| 6.8 |
| 1.62 |
| 5.58 |
| 0.87 |
| 0.57 |
| -1.57 |

| | |
|---|---|
| | -4.21 |
| | -1.73 |
| | 2.14 |
| | -1.38 |
| | 0.26 |
| | 0.28 |
| | 0.65 |
| | -1.37 |
| | 0.9 |
| | -0.55 |
| | -0.71 |
| | -2.71 |
| | -0.86 |
| | 0.01 |
| | -0.48 |
| | -0.29 |
| | -2.85 |
| | -1.84 |
| | 2.51 |
| | -0.74 |
| | -2.9 |
| | -1.42 |
| | -0.46 |
| | 0.2 |
| | -4.74 |
| | 5.68 |
| | 3.74 |
| | -0.76 |
| | -2.82 |
| | 1.69 |
| | -0.77 |
| | -0.15 |
| | 3.74 |
| | -1.79 |
| | -1.27 |

| | |
|---|---|
| | -2.5 |
| | 1.31 |
| | 4.26 |
| | -0.76 |
| | 1.07 |
| | 0.8 |
| | -1.29 |
| | -1.7 |
| | -0.72 |
| | -4.86 |
| | 2.24 |
| | -3.32 |
| | -0.16 |
| | -0.06 |
| | 0.65 |
| | -1.81 |
| | -1.37 |
| | -4.28 |
| | -0.62 |
| | -0.08 |
| | -0.86 |
| | -0.22 |
| | 1.46 |
| | -1.92 |
| | 1.89 |
| | 0.07 |
| | 1.9 |
| | -1.36 |
| | 0.32 |
| | 6.42 |
| | -2.56 |
| | 3.93 |
| | -0.8 |
| | 3.35 |
| | -4.72 |

| | |
|---|---|
| | 0.68 |
| | -1.68 |
| | -1.76 |
| | -1.36 |
| | 0.83 |
| | -5.88 |
| | -4.76 |
| | -2.45 |
| | -4.5 |
| | -2.19 |
| | 0.26 |
| | -0.34 |
| | -1.19 |
| | -0.43 |
| | -0.03 |
| | 1.66 |
| | 4.26 |
| | 0.97 |
| | 1.65 |
| | 1.67 |
| | -0.61 |
| | -0.02 |
| | 2.78 |
| | -4.16 |
| | -0.57 |
| | 2.29 |
| | -2.51 |
| | 0.86 |
| | 1.09 |
| | 2.07 |
| | -1.41 |
| | -1.32 |
| | -0.43 |
| | 0.29 |
| | 1.17 |

| | |
|---|---|
| | -0.75 |
| | -0.79 |
| | -0.41 |
| | 0.22 |
| | 2.56 |
| | -2.1 |
| | -0.54 |
| | -1.18 |
| | -1.62 |
| | -1.43 |
| | -0.38 |
| | -1.14 |
| | -0.19 |
| | 0.05 |
| | -2.17 |
| | -2.34 |
| | 1.98 |
| | -2.74 |
| | -0.68 |
| | 5.13 |
| | -1.44 |
| | -6.37 |
| | 0.81 |
| | -5.99 |
| | 3.01 |
| | -0.68 |
| | 0.23 |
| | -0.64 |
| | -2.67 |
| | -1.48 |
| | -1.07 |
| | 1.01 |
| | 0.49 |
| | -0.53 |
| | -0.75 |

| | |
|---|---|
| | -1.4 |
| | 2.51 |
| | -1.02 |
| | -4.25 |
| | -6.21 |
| | -0.37 |
| | -0.14 |
| | -0.35 |
| | -3.02 |
| | -0.75 |
| | -2.7 |
| | -0.09 |
| | 0 |
| | -2.22 |
| | -2.76 |
| | -2.54 |
| | 2.18 |
| | -0.34 |
| | -0.67 |
| | -0.13 |
| | 3.84 |
| | 1.05 |
| | -2.06 |
| | -1.26 |
| | -0.51 |
| | 0.1 |
| | -0.11 |
| | 0.39 |
| | 4.47 |
| | -0.33 |
| | 2.18 |
| | -0.13 |
| | -1.72 |
| | -4.48 |
| | -0.59 |

| | |
|---|---|
| | -1.08 |
| | 1.68 |
| | 1.17 |
| | 3.46 |
| | -0.06 |
| | 0.26 |
| | -0.62 |
| | 0.49 |
| | -6.46 |
| | -1.14 |
| | -0.86 |
| | -0.53 |
| | -1.73 |
| | -1.2 |
| | -4.63 |
| | -0.23 |
| | -0.36 |
| | 2.03 |
| | 1.66 |
| | -1.76 |
| | -2.68 |
| | -1.8 |
| | -1.16 |
| | 0.9 |
| | -0.26 |
| | -3.69 |
| | -2.04 |
| | 0.37 |
| | 0.34 |
| | -0.81 |
| | -2.17 |
| | -2.11 |
| | -1.47 |
| | -1.88 |
| | -0.64 |

| |
|---|
| 0.45 |
| 1.09 |
| 2.26 |
| -2.83 |
| -0.18 |
| -0.49 |
| -0.03 |
| -1.12 |
| -0.33 |
| -0.35 |
| 0.3 |
| -0.38 |
| -1 |
| 2.61 |
| -1.57 |
| 4.41 |
| 0.35 |
| 0.06 |
| -0.31 |
| -0.94 |
| -0.12 |
| -1.6 |
| -0.99 |
| 1.08 |
| -0.31 |
| -0.52 |
| 0.05 |
| 1.07 |
| 1.54 |
| -2.9 |
| -0.33 |
| 3.69 |
| 1.45 |
| 2.87 |
| -4.79 |

| |
|---|
| -2.76 |
| -4.65 |
| 1.5 |
| -0.38 |
| 3.54 |
| -0.09 |
| 0.89 |
| -0.31 |
| 3.82 |
| 1.98 |
| -0.79 |
| -0.06 |
| 0.17 |
| 2.97 |
| -2.36 |
| 1.62 |
| 1.32 |
| -0.13 |
| 0.58 |
| -2.43 |
| -0.63 |
| -1.23 |
| 1 |
| -0.71 |
| -0.62 |
| -1.41 |
| 7.68 |
| -4 |
| 0.25 |
| 1.21 |
| 4.04 |
| 0.47 |
| -1.46 |
| -0.92 |
| -4.42 |

| | |
|---|---|
| | 0.04 |
| | -5.77 |
| | -3.19 |
| | -2.49 |
| | -5.38 |
| | -2.07 |
| | -6.04 |
| | 0.53 |
| | -0.86 |
| | 0.68 |
| | 0.47 |
| | -0.29 |
| | 0.05 |
| | -2.28 |
| | -1.31 |
| | -2.94 |
| | 3.22 |
| | 1.76 |
| | -1.76 |
| | -2.27 |
| | 0.8 |
| | -0.62 |
| | -1.02 |
| | -1.78 |
| | -3.83 |
| | -4.22 |
| | -2.21 |
| | 3.78 |
| | -0.2 |
| | 5.55 |
| | -2.15 |
| | -0.33 |
| | -1.43 |
| | 0.66 |
| | -0.73 |

| | |
|---|---|
| | 4.64 |
| | 2.58 |
| | -0.92 |
| | -1.99 |
| | -2.36 |
| | -0.81 |
| | -1.76 |
| | -2.63 |
| | 0.45 |
| | -1.05 |
| | -1.83 |
| | -5.17 |
| | 0.64 |
| | -3.08 |
| | -0.7 |
| | -0.93 |
| | -0.54 |
| | -3.39 |
| | -0.14 |
| | 0.04 |
| | 1.88 |
| | -3 |
| | 0.55 |
| | -0.24 |
| | -0.33 |
| | -0.28 |
| | -3.25 |
| | 0.2 |
| | 1.25 |
| | -0.36 |
| | 0.44 |
| | 0.42 |
| | -0.33 |
| | -0.34 |
| | -1.76 |

| | |
|---|---|
| | -1.27 |
| | 2.29 |
| | -1 |
| | -0.29 |
| | -1.99 |
| | -2.73 |
| | 0.47 |
| | 1.91 |
| | -2.17 |
| | 0.38 |
| | -1.39 |
| | -1.93 |
| | -0.63 |
| | -4.18 |
| | -1.06 |
| | 0.65 |
| | -0.96 |
| | -2.67 |
| | 2.06 |
| | -3.87 |
| | -2.5 |
| | -4.05 |
| | -6.35 |
| | -0.1 |
| | -1.52 |
| | -0.85 |
| | -1.95 |
| | 2.77 |
| | -0.28 |
| | 0.1 |
| | -1.48 |
| | 0.98 |
| | -1.39 |
| | 1.29 |
| | -1.05 |

| | |
|---|---|
| | -5.33 |
| | -3.18 |
| | -3.7 |
| | 0.53 |
| | -8.16 |
| | 0.69 |
| | 0.87 |
| | -1.18 |
| | 3.54 |
| | -2.01 |
| | -1.67 |
| | -2.39 |
| | -2.08 |
| | 3.11 |
| | 0.14 |
| | -1.23 |
| | -0.92 |
| | -6.93 |
| | -2.19 |
| | -0.14 |
| | -1.77 |
| | 5.64 |
| | -7.25 |
| | -0.82 |
| | -6.54 |
| | 0.7 |
| | 3.28 |
| | -2.75 |
| | -7.82 |
| | -5.28 |
| | -2.21 |
| | 0.84 |
| | -0.11 |
| | 1.24 |
| | 4.44 |

| |
|---|
| 0.21 |
| 0.66 |
| -1.63 |
| 0.94 |
| 1.26 |
| 0.95 |
| -1.33 |
| -0.62 |
| -0.96 |
| -3.06 |
| 2.17 |
| 0.84 |
| 5.16 |
| 0.89 |
| -0.64 |
| 0.52 |
| 5.67 |
| -1.83 |
| -1.73 |
| -0.18 |
| 4.4 |
| 2.02 |
| -2.76 |
| -0.29 |
| 0.49 |
| 0.05 |
| -3.56 |
| 1.61 |
| -0.35 |
| -4.77 |
| 1.03 |
| -1.49 |
| -0.46 |
| 0.1 |
| -2.58 |

| | |
|---|---|
| | -0.94 |
| | 1.17 |
| | -0.2 |
| | 1.83 |
| | 2.98 |
| | 0.72 |
| | 2.27 |
| | 0.85 |
| | -0.09 |
| | -5.68 |
| | 1.76 |
| | -0.56 |
| | 1.06 |
| | -2.17 |
| | 0.42 |
| | 0.64 |
| | -0.27 |
| | 0.92 |
| | -3.26 |
| | -1.99 |
| | -0.07 |
| | 1.14 |
| | -1.21 |
| | -2.34 |
| | -1.95 |
| | 0.52 |
| | -3.32 |
| | 1.15 |
| | -1.85 |
| | -0.91 |
| | 0.93 |
| | -1.72 |
| | 0.23 |
| | -2.7 |
| | 0.68 |

| |
|---:|
| -4.31 |
| 5.13 |
| 4.86 |
| -0.75 |
| 7.34 |
| -2.83 |
| -1.54 |
| -7.17 |
| -2.18 |
| 1.11 |
| 4.33 |
| 1.72 |
| 0.42 |
| 2.42 |
| -1.9 |
| -3.19 |
| 1.64 |
| -0.27 |
| 3.13 |
| 1.22 |
| 2.22 |
| -0.06 |
| 0.33 |
| -1.3 |
| -0.07 |
| -0.01 |
| -0.41 |
| -3.06 |
| -1.32 |
| -2.53 |
| -1.75 |
| -1.44 |
| -0.01 |
| 2.82 |
| -0.64 |

| | |
|---|---|
| | -3.54 |
| | -0.44 |
| | -7.02 |
| | -2.76 |
| | 0.62 |
| | 0.7 |
| | -1.14 |
| | 4.12 |
| | -1.6 |
| | 1.25 |
| | 7.58 |
| | 6.08 |
| | 3.98 |
| | 1.89 |
| | 0.95 |
| | -1.26 |
| | 1.01 |
| | -1.65 |
| | -1.84 |
| | 0.74 |
| | 1.85 |
| | -0.89 |
| | 4.94 |
| | 2.72 |
| | 0.83 |
| | -0.94 |
| | 0.65 |
| | 0.05 |
| | -3.28 |
| | -1.72 |
| | 3.1 |
| | -0.4 |
| | 0.33 |
| | -1.6 |
| | -0.97 |

| |
|---:|
| 0.03 |
| -0.12 |
| 1.88 |
| -1.01 |
| 1.78 |
| 0.39 |
| 1.51 |
| -0.27 |
| -1.15 |
| -1.48 |
| -0.05 |
| -1.53 |
| -0.27 |
| -0.78 |
| -9.3 |

| Log2 Fold Change: (Bb_pos_12022024_Newanalysis) / (Water_pos_12022024_Newanalysis) |
|---|
| -1.32 |
| 5.44 |
| 0.22 |
| -0.89 |
| 2.24 |
| 2.16 |
| 6.04 |
| -0.42 |
| -0.28 |
| 0.26 |
| -0.3 |
| 4.43 |
| -1.69 |
| 1.81 |
| -0.35 |
| 2.65 |
| -1.03 |
| -0.15 |
| -1.3 |
| 1.68 |
| 0.31 |
| -0.74 |
| 0.13 |
| -0.72 |
| -0.96 |
| -0.91 |
| -0.53 |
| -0.67 |
| 1.92 |
| -2.61 |
| -0.72 |
| 3.51 |
| -0.3 |

| |
|---|
| 2.61 |
| -0.83 |
| 1.79 |
| -1.83 |
| 5.31 |
| 4.66 |
| -3.35 |
| -0.54 |
| -1.45 |
| -1.31 |
| 0.75 |
| -1.27 |
| -1.26 |
| -1.88 |
| -1.43 |
| -1.5 |
| -1.35 |
| -1.5 |
| -1.35 |
| -1.88 |
| -1.36 |
| -0.19 |
| -3.11 |
| 0.73 |
| 0.6 |
| -0.73 |
| 2.75 |
| -3.92 |
| -0.13 |
| 4.09 |
| -1.11 |
| -2.08 |
| -1.12 |
| -3.96 |
| -1.36 |

| | |
|---|---|
| | -1.95 |
| | -1.55 |
| | -4.66 |
| | -1.88 |
| | -1.31 |
| | -2.74 |
| | -2 |
| | 0.62 |
| | -1.19 |
| | 1.87 |
| | -2.34 |
| | -2.06 |
| | -3.62 |
| | -4.63 |
| | -1.46 |
| | -1.71 |
| | -2.18 |
| | -2.74 |
| | -2.06 |
| | -4 |
| | -2.09 |
| | -2.94 |
| | -3.24 |
| | -1.89 |
| | -1.26 |
| | -0.77 |
| | -1.37 |
| | -2.41 |
| | -3.06 |
| | -3.14 |
| | -2.59 |
| | -1.73 |
| | -0.65 |
| | -2.79 |
| | -1.31 |

| |
|---|
| -2.59 |
| -1.48 |
| -1.26 |
| -2.18 |
| -1.23 |
| -0.23 |
| -1.76 |
| -1.3 |
| 0.31 |
| -3.46 |
| -1.29 |
| -1.84 |
| -2.41 |
| -2.52 |
| -1.17 |
| -1.23 |
| -2.35 |
| -1.26 |
| -0.76 |
| -1.29 |
| 0.48 |
| -2.52 |
| 0.72 |
| -1.51 |
| -0.64 |
| -2.23 |
| -1.25 |
| -1.14 |
| 0.51 |
| 0.76 |
| 0.17 |
| 1.87 |
| 1.76 |
| -3.39 |
| 0.58 |

| | |
|---|---|
| | -1.4 |
| | -0.56 |
| | 1.37 |
| | -0.82 |
| | -0.87 |
| | -0.85 |
| | -0.72 |
| | -2.31 |
| | 1.69 |
| | -1.25 |
| | 0.12 |
| | -0.57 |
| | -0.99 |
| | -0.59 |
| | -0.8 |
| | -1 |
| | 1.68 |
| | -0.73 |
| | 3.95 |
| | -0.47 |
| | -2.41 |
| | -1.22 |
| | -0.6 |
| | -1.21 |
| | -0.12 |
| | -4.22 |
| | 3.27 |
| | -3.62 |
| | -2.23 |
| | 1.96 |
| | 1.57 |
| | -3.31 |
| | 3.27 |
| | -0.19 |
| | 0.34 |

| | |
|---|---|
| | 3.57 |
| | 1.75 |
| | -2.48 |
| | 0.66 |
| | -0.49 |
| | -0.24 |
| | -1.24 |
| | -1.12 |
| | 6.27 |
| | -1.09 |
| | 2.86 |
| | -1.02 |
| | -0.89 |
| | 1.32 |
| | 2.5 |
| | 1.87 |
| | -0.76 |
| | -1.47 |
| | -0.66 |
| | -1.2 |
| | -0.23 |
| | -0.63 |
| | 2.61 |
| | -1.08 |
| | 1.87 |
| | -0.19 |
| | -1.41 |
| | -0.12 |
| | -3.91 |
| | -2.06 |
| | -3.18 |
| | -3.48 |
| | 0.53 |
| | 1.96 |
| | 1.89 |

| | |
|---|---|
| | -3.14 |
| | -1 |
| | -1.3 |
| | -0.86 |
| | 1.95 |
| | -1.08 |
| | -0.72 |
| | 0.05 |
| | -0.99 |
| | 1.96 |
| | 0.46 |
| | -0.91 |
| | -0.8 |
| | 0.62 |
| | 1.89 |
| | 2.92 |
| | -0.99 |
| | -0.19 |
| | -0.26 |
| | -1.06 |
| | -0.6 |
| | -0.87 |
| | -2.39 |
| | -1.61 |
| | -1.19 |
| | -2.22 |
| | -1.32 |
| | -1.04 |
| | 1.11 |
| | -1.29 |
| | 0.65 |
| | -0.4 |
| | 0.04 |
| | -0.89 |
| | -1.46 |

| | |
|---|---|
| | 2.75 |
| | 1.49 |
| | 0.03 |
| | -0.36 |
| | 0.61 |
| | -0.72 |
| | 1.21 |
| | -1.39 |
| | -0.83 |
| | -0.6 |
| | 4.18 |
| | 1.49 |
| | 0.47 |
| | 1.78 |
| | 5.4 |
| | 3.56 |
| | 1.92 |
| | 2.48 |
| | -1.07 |
| | -0.61 |
| | -1.06 |
| | 2.66 |
| | 1.83 |
| | 0.63 |
| | 1.09 |
| | -0.53 |
| | 0.95 |
| | 0.83 |
| | 1.14 |
| | -0.05 |
| | 0.05 |
| | -1.85 |
| | 2.59 |
| | -2.28 |
| | 0.47 |

| | |
|---|---|
| | 1.69 |
| | -1.14 |
| | -1.44 |
| | 1.33 |
| | -0.23 |
| | 0.17 |
| | -0.82 |
| | -0.86 |
| | -2.7 |
| | -1.38 |
| | 1.27 |
| | -1.27 |
| | -1.27 |
| | -1.54 |
| | -3.04 |
| | -0.31 |
| | -1.27 |
| | -1.38 |
| | -0.78 |
| | -0.52 |
| | -1.3 |
| | 2.3 |
| | -2.79 |
| | -1.27 |
| | -1.34 |
| | -1.14 |
| | -0.1 |
| | -0.63 |
| | 2.66 |
| | 2.83 |
| | -1.38 |
| | 0.86 |
| | 4.69 |
| | 3.62 |
| | -0.91 |

| | |
|---|---|
| | 1.07 |
| | 3.92 |
| | -0.4 |
| | 3.1 |
| | 4.72 |
| | 5.98 |
| | 1.77 |
| | -1.08 |
| | 2.84 |
| | -0.92 |
| | 0.79 |
| | 4.01 |
| | 0.35 |
| | -1.46 |
| | 1.3 |
| | 5.48 |
| | 1.59 |
| | -0.32 |
| | -0.69 |
| | 3.25 |
| | 1.28 |
| | 4.78 |
| | -0.31 |
| | -0.36 |
| | -0.68 |
| | 3.01 |
| | 1.17 |
| | 5.6 |
| | -0.03 |
| | 9.18 |
| | 9.04 |
| | -2.49 |
| | 3.67 |
| | 5.81 |
| | 2.76 |

| | |
|---|---|
| | 1.59 |
| | -0.65 |
| | 5.01 |
| | 3.08 |
| | -1.35 |
| | -1.05 |
| | -1.15 |
| | -1.42 |
| | -1.04 |
| | 1.38 |
| | 5.02 |
| | -0.51 |
| | 1.6 |
| | 4.19 |
| | -1.06 |
| | -3.46 |
| | 0.88 |
| | -0.8 |
| | 0.42 |
| | 1.11 |
| | -0.68 |
| | -1.43 |
| | -0.36 |
| | -0.7 |
| | 1.25 |
| | 1.34 |
| | -0.32 |
| | -0.5 |
| | -3.25 |
| | -2.44 |
| | -1.1 |
| | 1.15 |
| | 1.16 |
| | -3.85 |
| | -0.56 |

| | |
|---|---|
| | 1.21 |
| | -2.27 |
| | 2.83 |
| | 0.48 |
| | 1.75 |
| | -1.11 |
| | 0.54 |
| | 0.76 |
| | 3.69 |
| | 4.43 |
| | 1.53 |
| | 1.11 |
| | -1.14 |
| | 0.43 |
| | 1.15 |
| | -0.68 |
| | -1.85 |
| | -0.4 |
| | -1.55 |
| | -2.62 |
| | 6.38 |
| | -0.5 |
| | -1.21 |
| | -1.03 |
| | 3.83 |
| | 2.11 |
| | 5.48 |
| | 2.43 |
| | -1.38 |
| | -3.48 |
| | -1.04 |
| | 5.37 |
| | -0.61 |
| | -0.87 |
| | -0.42 |

| | |
|---|---|
| | 0.95 |
| | -1.56 |
| | 3.39 |
| | 0.32 |
| | 3.5 |
| | 0.73 |
| | 1.99 |
| | -0.66 |
| | 0.34 |
| | 4.98 |
| | 1.28 |
| | 0.23 |
| | 0.08 |
| | -0.74 |
| | 4.18 |
| | 2.91 |
| | 6.94 |
| | 2.09 |
| | -0.52 |
| | -0.71 |
| | 0.64 |
| | -1.74 |
| | -0.36 |
| | 0.27 |
| | 2.54 |
| | -0.36 |
| | 0.71 |
| | -0.25 |
| | -0.82 |
| | 5.89 |
| | 1.61 |
| | 4.02 |
| | 2.97 |
| | -0.12 |
| | 0.46 |

| | |
|---|---|
| | -0.55 |
| | -1.38 |
| | 1.09 |
| | -1.08 |
| | -1.3 |
| | 4.62 |
| | -1.11 |
| | 0.72 |
| | 0.42 |
| | 5.61 |
| | 3.86 |
| | 7.3 |
| | 8.7 |
| | 6.92 |
| | 9.08 |
| | 6.58 |
| | 9.24 |
| | 5.91 |
| | -1.26 |
| | 1.56 |
| | 1.32 |
| | -0.62 |
| | 3.15 |
| | 5.8 |
| | 7.54 |
| | 6.52 |
| | 8.41 |
| | 1.83 |
| | -0.78 |
| | -0.95 |
| | 1.37 |
| | 3.55 |
| | 6.32 |
| | 6.9 |
| | 2.37 |

| | |
|---|---|
| | -1.01 |
| | 1.01 |
| | -3.6 |
| | -0.16 |
| | 3.07 |
| | 0.47 |
| | 4.41 |
| | 0.82 |
| | -0.78 |
| | 0.36 |
| | -0.41 |
| | 1.82 |
| | -1.42 |
| | 0.57 |
| | -0.59 |
| | 0.06 |
| | 4.2 |
| | 0.79 |
| | 5.27 |
| | -0.77 |
| | -0.94 |
| | 1.47 |
| | -0.59 |
| | -0.64 |
| | -0.76 |
| | -1.25 |
| | 3.23 |
| | 1.91 |
| | -0.14 |
| | -0.75 |
| | -1.14 |
| | -0.52 |
| | -0.6 |
| | 1.61 |
| | -0.76 |

| | |
|---|---|
| | -0.35 |
| | -0.77 |
| | 0.19 |
| | 2.27 |
| | -0.81 |
| | -0.97 |
| | -0.42 |
| | -0.79 |
| | -0.95 |
| | -0.2 |
| | -0.86 |
| | -1.02 |
| | 2.18 |
| | 3.63 |
| | -0.33 |
| | -0.46 |
| | 0.59 |
| | -0.62 |
| | -0.68 |
| | 4.24 |
| | -2.07 |
| | 2.22 |
| | -0.42 |
| | 1.18 |
| | -0.07 |
| | 2.85 |
| | -0.73 |
| | 1.37 |
| | -1.02 |
| | -1.53 |
| | -0.31 |
| | -0.79 |
| | -0.32 |
| | 4.27 |
| | 4.94 |

| |
|---|
| 5.01 |
| 0.3 |
| 5.11 |
| -0.12 |
| 0.32 |
| -0.35 |
| -0.83 |
| 0.92 |
| -1.1 |
| 0.78 |
| -1.53 |
| 0.06 |
| -0.52 |
| -0.67 |
| -0.34 |
| -2 |
| 3.63 |
| 2.26 |
| 2.02 |
| -1.22 |
| 0.28 |
| -1.85 |
| -1.01 |
| -0.38 |
| 3.13 |
| 2.9 |
| 0.89 |
| 1.94 |
| 2.09 |
| -0.77 |
| 2.18 |
| 1.58 |
| -1.74 |
| 4.95 |
| -1.17 |

| | |
|---|---|
| | 2.14 |
| | -1.04 |
| | -1.03 |
| | -1.42 |
| | -1.61 |
| | 2.49 |
| | -1.25 |
| | -1 |
| | 0.82 |
| | -0.24 |
| | 0.18 |
| | 3.87 |
| | -1.72 |
| | -1.45 |
| | -0.72 |
| | 0.32 |
| | 3.8 |
| | -0.85 |
| | -0.27 |
| | 3.46 |
| | -1.03 |
| | -0.36 |
| | -1.02 |
| | -2.31 |
| | -0.62 |
| | -1.37 |
| | -0.26 |
| | -0.54 |
| | 2.81 |
| | 0.39 |
| | -0.7 |
| | -1.03 |
| | -0.69 |
| | -0.48 |
| | 3.54 |

| | |
|---|---|
| | -1.52 |
| | -3.09 |
| | -1.4 |
| | -1.12 |
| | -3.16 |
| | -2.3 |
| | -1.5 |
| | -2.48 |
| | -2.45 |
| | -1.3 |
| | -0.09 |
| | -2.15 |
| | -1.82 |
| | -2.11 |
| | -0.77 |
| | -2.48 |
| | -1.63 |
| | 3.43 |
| | 0.78 |
| | 1.96 |
| | -0.81 |
| | -1.42 |
| | -1.3 |
| | 0.64 |
| | -1.36 |
| | -1.55 |
| | -1.25 |
| | -0.97 |
| | -0.42 |
| | -0.12 |
| | -2.53 |
| | -2.56 |
| | -1.51 |
| | -2.79 |
| | -1.44 |

| | |
|---|---|
| | -2.49 |
| | -2.09 |
| | -0.17 |
| | -0.19 |
| | -1.62 |
| | 1.79 |
| | -1.12 |
| | -3 |
| | 0.02 |
| | -2.06 |
| | -0.75 |
| | -2.58 |
| | -1.89 |
| | -0.9 |
| | -1.42 |
| | 1.99 |
| | 1.43 |
| | -1.12 |
| | 0.71 |
| | 0.71 |
| | 2.9 |
| | 0 |
| | 8.11 |
| | 2.05 |
| | -0.75 |
| | -1.25 |
| | 4.37 |
| | 1.35 |
| | 0.01 |
| | -2.98 |
| | -1.86 |
| | -0.87 |
| | 0.21 |
| | -1.52 |
| | -1.19 |

| |
|---|
| 0.44 |
| 3.24 |
| -0.3 |
| 0.62 |
| 1.74 |
| -1.29 |
| -1.49 |
| 0.21 |
| -0.16 |
| -2.65 |
| -1.23 |
| -0.58 |
| -0.67 |
| -0.62 |
| -1.04 |

| |
|---|
| Log2 Fold Change: (CM_pos_12022024_Newanalysis) / (Water_pos_12022024_Newanalysis) |
| -0.74 |
| 4.4 |
| -0.13 |
| 4.37 |
| 1.33 |
| 2.28 |
| 5.25 |
| 1.69 |
| -0.65 |
| 1.08 |
| 0.72 |
| 4.11 |
| 0.27 |
| 3.19 |
| -0.49 |
| 1.54 |
| -1.48 |
| 4.56 |
| 0.16 |
| 3.4 |
| 0.18 |
| 3.54 |
| 4.21 |
| 2.75 |
| 4.94 |
| 0.51 |
| -0.69 |
| -1.51 |
| 1.28 |
| -1.82 |
| -1.16 |
| -3.01 |
| -1.82 |

| |
|---|
| 1.88 |
| 4.32 |
| -0.17 |
| 0.96 |
| -1.27 |
| -1.87 |
| 2.73 |
| -0.75 |
| -1.87 |
| 2.14 |
| -0.28 |
| -1.73 |
| 2.96 |
| 2.79 |
| 3.11 |
| 2.61 |
| 1.42 |
| 3.23 |
| 2.42 |
| 3.41 |
| 1.38 |
| -1.37 |
| -1.98 |
| -0.55 |
| -1.1 |
| -0.11 |
| 1.35 |
| -3.59 |
| -0.83 |
| 2.87 |
| 4.88 |
| 1.33 |
| -1.19 |
| 1.1 |
| 0.92 |

| |
|---|
| -1.53 |
| -0.5 |
| 0.14 |
| -2.43 |
| 1.41 |
| 1.07 |
| -1.64 |
| -0.19 |
| 4.3 |
| 1.59 |
| 2.36 |
| 1.39 |
| 0.65 |
| -0.05 |
| -1.71 |
| 1.26 |
| -0.79 |
| -0.26 |
| -2.29 |
| 2.3 |
| 0.27 |
| -1.75 |
| 2.11 |
| 2.43 |
| 1.13 |
| 1.87 |
| 1.12 |
| -0.35 |
| 1.92 |
| 0.6 |
| -2.2 |
| 0.22 |
| 1.4 |
| -1.62 |
| 1.19 |

| |
|---|
| -2.32 |
| 0.81 |
| -2.02 |
| 0.38 |
| -0.78 |
| -1.1 |
| -1.25 |
| 0.7 |
| 0.99 |
| 0.41 |
| 1.44 |
| -2.35 |
| -0.36 |
| -1.61 |
| 5.26 |
| 2.84 |
| -2.32 |
| -2.02 |
| -0.66 |
| -1.46 |
| -0.32 |
| -2.07 |
| 1.33 |
| 1.14 |
| -0.17 |
| -2.27 |
| -1.81 |
| -1.81 |
| 2.35 |
| 4.81 |
| 1.59 |
| 3.14 |
| -0.53 |
| -0.99 |
| 0.72 |

| | |
|---|---|
| | -1.39 |
| | -1.18 |
| | 0.37 |
| | -0.42 |
| | -1.61 |
| | -1.69 |
| | -1.21 |
| | -2.42 |
| | -0.18 |
| | -1.39 |
| | -0.87 |
| | -1.31 |
| | 1.12 |
| | -1.74 |
| | -0.39 |
| | 2.12 |
| | -0.49 |
| | -0.5 |
| | 2.52 |
| | -0.9 |
| | -1.62 |
| | -0.26 |
| | 0.07 |
| | 1.55 |
| | -0.31 |
| | 0.33 |
| | 2.23 |
| | -1.69 |
| | -1.87 |
| | 1.41 |
| | 3.14 |
| | 0.16 |
| | 2.23 |
| | 1.25 |
| | 2.23 |

| | |
|---|---|
| | 1.32 |
| | 0.96 |
| | 1.95 |
| | 2.4 |
| | 3.26 |
| | 1.21 |
| | 2.7 |
| | -0.7 |
| | 4.9 |
| | -0.8 |
| | 0.93 |
| | -1.46 |
| | 2.2 |
| | 3.69 |
| | 1.4 |
| | 2.47 |
| | -0.8 |
| | 0.68 |
| | 3.04 |
| | 0.09 |
| | 0.91 |
| | 0.89 |
| | 1.73 |
| | -1.33 |
| | 1.11 |
| | -0.2 |
| | 0.38 |
| | 2.21 |
| | 0.46 |
| | 2.07 |
| | 1.03 |
| | 0.43 |
| | 2.48 |
| | 3.17 |
| | 4.85 |

| | |
|---|---|
| | 0.96 |
| | -0.82 |
| | -0.74 |
| | -0.44 |
| | 1.19 |
| | -1.04 |
| | 0.03 |
| | 0.11 |
| | -1.27 |
| | 1.16 |
| | -0.2 |
| | -0.01 |
| | 1 |
| | 2.1 |
| | -0.48 |
| | 0.34 |
| | 2.72 |
| | -1.65 |
| | 1.85 |
| | -1.31 |
| | 1.79 |
| | -1.63 |
| | -1.07 |
| | -0.84 |
| | 0.44 |
| | -0.31 |
| | -1.39 |
| | -0.95 |
| | 0.1 |
| | -1.1 |
| | -0.54 |
| | -0.67 |
| | -0.69 |
| | -0.95 |
| | 0.97 |

| | |
|---|---|
| | 1.48 |
| | 1.18 |
| | -0.15 |
| | 0.69 |
| | 1.23 |
| | 0.02 |
| | 1.9 |
| | -1.54 |
| | -0.34 |
| | 1.84 |
| | 2.33 |
| | 0.65 |
| | -0.61 |
| | 2.81 |
| | 0.96 |
| | 0.68 |
| | -0.14 |
| | -0.61 |
| | -0.88 |
| | 4.97 |
| | -0.37 |
| | -0.72 |
| | 0.62 |
| | -1.41 |
| | 0.53 |
| | -0.67 |
| | 2.1 |
| | -0.24 |
| | 0.01 |
| | -0.44 |
| | 1.36 |
| | 1.63 |
| | 1.34 |
| | -0.67 |
| | 3.26 |

| | |
|---|---|
| | 1.73 |
| | 2.03 |
| | -0.43 |
| | -0.05 |
| | -0.3 |
| | 1.63 |
| | 1.82 |
| | -0.95 |
| | -2.02 |
| | -0.84 |
| | -0.18 |
| | -1.8 |
| | -1.8 |
| | -1.84 |
| | -1.97 |
| | -1.32 |
| | 1.16 |
| | -2.03 |
| | 1.63 |
| | -0.49 |
| | 4.07 |
| | -0.22 |
| | -2.41 |
| | -0.04 |
| | -1.63 |
| | -1.29 |
| | 0.28 |
| | 0.89 |
| | 5.31 |
| | 4.67 |
| | 0.4 |
| | 3.83 |
| | 1.56 |
| | 5.71 |
| | 0.01 |

| | |
|---|---|
| | 2.16 |
| | 2.19 |
| | 1.87 |
| | 4.26 |
| | 4.2 |
| | 1.92 |
| | 4.6 |
| | 1.72 |
| | 2.4 |
| | 1.05 |
| | 1.53 |
| | 2.85 |
| | 2.4 |
| | 0.85 |
| | -0.54 |
| | 4.47 |
| | 1.5 |
| | 2.94 |
| | 3.06 |
| | 0.92 |
| | 3.12 |
| | 1.84 |
| | 1.9 |
| | 2.31 |
| | 0.27 |
| | 0.99 |
| | -1.02 |
| | 1.65 |
| | 2.84 |
| | 2 |
| | 0.69 |
| | -0.83 |
| | 0.56 |
| | -0.51 |
| | -0.55 |

| | |
|---|---|
| | -1.32 |
| | -0.47 |
| | 1.08 |
| | -1.12 |
| | -1.32 |
| | -1.32 |
| | -1.79 |
| | -0.47 |
| | -0.83 |
| | 2.75 |
| | 2.62 |
| | 0.03 |
| | 0.91 |
| | 2.69 |
| | -1.22 |
| | -1.15 |
| | -0.46 |
| | -1.2 |
| | 3.01 |
| | 0.33 |
| | 2.46 |
| | 0.65 |
| | 0.72 |
| | 0.92 |
| | 4.09 |
| | 0.49 |
| | 1.37 |
| | 1.23 |
| | -1.66 |
| | -0.59 |
| | -0.68 |
| | 2.29 |
| | 1.86 |
| | 0.22 |
| | -1.02 |

| | |
|---|---|
| | 0.54 |
| | -1.88 |
| | 1.83 |
| | 0.08 |
| | 2.63 |
| | -1.04 |
| | 1.54 |
| | 1.46 |
| | 4.99 |
| | 2.81 |
| | 1.6 |
| | 1.23 |
| | -1.43 |
| | 4.14 |
| | 5.35 |
| | 1.19 |
| | 0.72 |
| | 3.38 |
| | 2.99 |
| | 0.17 |
| | 5.11 |
| | 0.82 |
| | -0.67 |
| | -1.31 |
| | 5.6 |
| | -0.45 |
| | 3.98 |
| | 5.57 |
| | -1.76 |
| | -0.83 |
| | 3.46 |
| | 1.38 |
| | 1.84 |
| | 2.96 |
| | 1.99 |

| | |
|---|---|
| | 3.18 |
| | 0.86 |
| | 4.48 |
| | 6.46 |
| | 3.58 |
| | -1.07 |
| | -0.9 |
| | -0.98 |
| | 0.28 |
| | 4.72 |
| | 0.87 |
| | 2.99 |
| | 4.48 |
| | 1.79 |
| | 3.15 |
| | 3.23 |
| | 4.33 |
| | 1.37 |
| | 2.29 |
| | 4.19 |
| | 1.84 |
| | 3.97 |
| | 2.03 |
| | 4.16 |
| | 2.44 |
| | 2.15 |
| | 0.33 |
| | 3.35 |
| | 1.27 |
| | 4.02 |
| | 0.49 |
| | 3.39 |
| | 4.6 |
| | 4.58 |
| | 4.33 |

| | |
|---|---|
| | 3.55 |
| | 0.63 |
| | 0.67 |
| | 1 |
| | -1.94 |
| | 3.91 |
| | 3.1 |
| | 2.5 |
| | -0.69 |
| | 3.85 |
| | 4.84 |
| | 4.18 |
| | 2.94 |
| | 0.49 |
| | 2.1 |
| | -0.03 |
| | 1.02 |
| | -1.46 |
| | 0.24 |
| | 1.92 |
| | 0.34 |
| | -0.82 |
| | 1.18 |
| | 4.61 |
| | 3.55 |
| | 0.35 |
| | 1.25 |
| | 2.04 |
| | -0.09 |
| | -0.94 |
| | 1.27 |
| | -0.86 |
| | 1.82 |
| | -0.26 |
| | -0.44 |

| | |
|---|---|
| | -1.2 |
| | -1.1 |
| | -0.32 |
| | 1.95 |
| | -0.22 |
| | -0.1 |
| | 3.95 |
| | 0.23 |
| | -1.35 |
| | 1.8 |
| | -0.65 |
| | 3.21 |
| | -1.48 |
| | 0.32 |
| | -0.19 |
| | 2.75 |
| | 0.93 |
| | -0.1 |
| | 1.41 |
| | -0.48 |
| | 0 |
| | 0.65 |
| | -1.04 |
| | -1.06 |
| | 0.64 |
| | -0.66 |
| | 0.75 |
| | 1.44 |
| | -0.02 |
| | -1.21 |
| | 1.12 |
| | 3.67 |
| | -0.47 |
| | 0.25 |
| | 0.55 |

| |
|---|
| -0.52 |
| -1.23 |
| 0.89 |
| 1.63 |
| 0.49 |
| -0.82 |
| 2.56 |
| -1.38 |
| 4.76 |
| 1.36 |
| -0.74 |
| -0.49 |
| 1.28 |
| 2.91 |
| 0.68 |
| 0.85 |
| 2.69 |
| 2.51 |
| 0.43 |
| 2.39 |
| 3.85 |
| 4.35 |
| 4.91 |
| -0.54 |
| 0.08 |
| 2.76 |
| 4.58 |
| 1.5 |
| -1 |
| 3.92 |
| -1.22 |
| 0.25 |
| 0.3 |
| -0.17 |
| 4.42 |

| | |
|---|---|
| | -1.45 |
| | 0.21 |
| | -0.35 |
| | -2.03 |
| | -0.97 |
| | -1.09 |
| | -0.37 |
| | 2.71 |
| | -0.52 |
| | -0.52 |
| | -1.57 |
| | 1.01 |
| | -1.8 |
| | 1.11 |
| | 0.91 |
| | 1.92 |
| | 4.28 |
| | -0.32 |
| | 0.65 |
| | -1.79 |
| | -1.4 |
| | 3.03 |
| | -2 |
| | -0.39 |
| | -0.14 |
| | -0.84 |
| | -1.12 |
| | 1.43 |
| | -0.18 |
| | -0.52 |
| | 2.75 |
| | 2.15 |
| | -2.41 |
| | 5.07 |
| | -1.76 |

| | |
|---|---|
| | 1.21 |
| | -1.38 |
| | -1.89 |
| | -0.67 |
| | -1.15 |
| | 1.46 |
| | 3.14 |
| | 0.94 |
| | 3.11 |
| | -1.65 |
| | 0.38 |
| | -1.52 |
| | -0.35 |
| | -1.87 |
| | 3.28 |
| | -1.58 |
| | -0.22 |
| | -1.24 |
| | -2.39 |
| | -0.22 |
| | -1.74 |
| | -1.64 |
| | -1.16 |
| | -1.2 |
| | -1.34 |
| | -2.14 |
| | -1.68 |
| | 1.13 |
| | 1.74 |
| | 0.35 |
| | 1.5 |
| | -0.38 |
| | -0.31 |
| | -0.42 |
| | 3.03 |

| | |
|---|---|
| | -0.55 |
| | 1.87 |
| | 4.23 |
| | 1.25 |
| | 2.08 |
| | 0.68 |
| | 1.34 |
| | 0.23 |
| | 2.71 |
| | 1.37 |
| | 0.79 |
| | -2.17 |
| | 1.09 |
| | -1.96 |
| | -1.65 |
| | -2.43 |
| | -1.63 |
| | -0.68 |
| | 0.01 |
| | 0.74 |
| | -0.97 |
| | -2.07 |
| | -1.66 |
| | 0.41 |
| | -1.73 |
| | -1.93 |
| | -1.86 |
| | -1.87 |
| | -0.04 |
| | 0.25 |
| | -2.41 |
| | -2.39 |
| | 1.14 |
| | -1.55 |
| | 1.29 |

| | |
|---|---|
| | -2.1 |
| | -2.34 |
| | -0.93 |
| | -0.38 |
| | -0.69 |
| | 1.31 |
| | -0.2 |
| | 0.8 |
| | -0.5 |
| | 3.23 |
| | 4.4 |
| | 2.6 |
| | 2.69 |
| | 0.31 |
| | -1.09 |
| | 3.22 |
| | 1.31 |
| | -0.11 |
| | 0.09 |
| | 0.23 |
| | 0.73 |
| | -0.83 |
| | 5.99 |
| | 2.87 |
| | -1.23 |
| | -0.59 |
| | -1.87 |
| | -1.33 |
| | -0.03 |
| | -0.54 |
| | -0.7 |
| | 0.97 |
| | -0.82 |
| | -2.1 |
| | 0.86 |

| |
|---|
| -0.23 |
| 2.35 |
| 1.89 |
| 1.8 |
| 2.65 |
| 2.34 |
| 1.51 |
| 0.24 |
| 2.06 |
| -1.35 |
| -1.34 |
| -1.89 |
| -2.57 |
| -2.57 |
| 3.51 |

| |
|---|
| Log2 Fold Change: (Hs_pos_12022024_Newanalysis) / (Water_pos_12022024_Newanalysis) |
| 3.91 |
| 4.44 |
| 3.6 |
| 6.67 |
| 1.93 |
| 1.9 |
| -4.04 |
| 1.65 |
| 2.59 |
| -1.71 |
| 0.33 |
| -1.84 |
| 1.97 |
| 3.47 |
| -1.42 |
| -0.13 |
| 0.01 |
| 2.74 |
| 0.29 |
| 6.2 |
| -2.44 |
| 1.53 |
| 3.82 |
| 2.39 |
| 4 |
| 4.57 |
| 5.04 |
| 5.83 |
| 4.54 |
| -3.49 |
| -3.47 |
| -1.46 |
| -3.49 |

| |
|---|
| 0.16 |
| 1.18 |
| 3.67 |
| 2.77 |
| 0.14 |
| -1.46 |
| 4.66 |
| -1.79 |
| -1.06 |
| 2.19 |
| 3.42 |
| -3.06 |
| -0.78 |
| -1.71 |
| -1.18 |
| -2.86 |
| -3.02 |
| -3.02 |
| -3.02 |
| -3.26 |
| -2.8 |
| -3.01 |
| -2.25 |
| 3.75 |
| -2.81 |
| 2.24 |
| 3.86 |
| -4.13 |
| -1.01 |
| 4.44 |
| 5.87 |
| 4.01 |
| -2.45 |
| -0.21 |
| 5.04 |

| | |
|---|---|
| | 4.83 |
| | 0.3 |
| | -0.99 |
| | -1.26 |
| | 2.52 |
| | 5.91 |
| | -0.34 |
| | -2.49 |
| | 3.14 |
| | 2.21 |
| | 2.67 |
| | 3.67 |
| | 3.39 |
| | 0.39 |
| | -2.87 |
| | 3.1 |
| | 7.24 |
| | -0.45 |
| | -4.07 |
| | 3.46 |
| | -0.27 |
| | -0.61 |
| | 3.31 |
| | 2.9 |
| | 0.39 |
| | -0.53 |
| | 0.73 |
| | 4.48 |
| | 2.5 |
| | 0.29 |
| | -4.12 |
| | -1.31 |
| | 2.86 |
| | -3.78 |
| | 2.52 |

| | |
|---|---|
| | -2.01 |
| | 2.72 |
| | -3.83 |
| | 1.53 |
| | 0.05 |
| | -1.02 |
| | -4.42 |
| | 0.81 |
| | 0.89 |
| | -1.83 |
| | 2.54 |
| | -3.43 |
| | 0.97 |
| | -2.87 |
| | 3.72 |
| | 3.53 |
| | -2.01 |
| | -1.96 |
| | 1.37 |
| | -0.61 |
| | 2.53 |
| | -1.04 |
| | 1.89 |
| | 1.43 |
| | 2.08 |
| | 0.81 |
| | -1.51 |
| | -2.74 |
| | 4.64 |
| | 7.46 |
| | 4.48 |
| | 1.53 |
| | -0.1 |
| | -4.14 |
| | 1.69 |

| | |
|---|---|
| | -1.6 |
| | 2.39 |
| | -3.43 |
| | 2.45 |
| | -2.01 |
| | -3.82 |
| | -1.5 |
| | -5 |
| | -0.86 |
| | -2.56 |
| | 0.13 |
| | 1.5 |
| | 5.8 |
| | -0.67 |
| | 6.39 |
| | 6.32 |
| | -0.4 |
| | 4.05 |
| | 3.35 |
| | 3.41 |
| | 4.53 |
| | 5.89 |
| | 6.85 |
| | 5.88 |
| | 4.78 |
| | 5.88 |
| | 5.66 |
| | -1.29 |
| | 4.82 |
| | 2.83 |
| | 6.17 |
| | 3.33 |
| | 5.47 |
| | 5.92 |
| | 3.23 |

| | |
|---|---|
| | 2.84 |
| | 6.87 |
| | 6.41 |
| | 6.03 |
| | 4.84 |
| | 6.79 |
| | 3.68 |
| | 5.78 |
| | 5.57 |
| | 5.66 |
| | 4.64 |
| | 1.64 |
| | 8.36 |
| | 1.51 |
| | 5.62 |
| | 2.02 |
| | 6.76 |
| | 6.22 |
| | 4.15 |
| | 9.38 |
| | 9.24 |
| | 7.61 |
| | 7.38 |
| | 4.52 |
| | 7.95 |
| | 7.13 |
| | 7.09 |
| | 6.07 |
| | 5.58 |
| | 7.64 |
| | 5.09 |
| | 4.91 |
| | 9.38 |
| | 7.27 |
| | 6.34 |

| | |
|---|---|
| | -1.87 |
| | 5.58 |
| | 2.86 |
| | 5.3 |
| | 5.99 |
| | 4.53 |
| | 3.34 |
| | 3.87 |
| | 2.19 |
| | 3.83 |
| | 4.76 |
| | 6.94 |
| | 6.36 |
| | 3.91 |
| | -0.08 |
| | 1.84 |
| | -2.63 |
| | 2.16 |
| | 5.95 |
| | 0.56 |
| | 3.24 |
| | 0.9 |
| | -4.81 |
| | 3.41 |
| | 3.86 |
| | -3.19 |
| | -2.65 |
| | 2.09 |
| | 2.05 |
| | 1.3 |
| | 3.04 |
| | 2.33 |
| | 5.6 |
| | 2.34 |
| | 2.39 |

| | |
|---|---|
| | 2.19 |
| | 5.71 |
| | -0.4 |
| | 1.34 |
| | 6.42 |
| | 6.25 |
| | 7.91 |
| | 1.23 |
| | 5.08 |
| | 8.56 |
| | 3.82 |
| | 3.87 |
| | 6.9 |
| | 7.21 |
| | 5.18 |
| | 3.87 |
| | 6.38 |
| | 5.64 |
| | -0.61 |
| | 8.22 |
| | 5.58 |
| | 3.19 |
| | 5.3 |
| | 1.33 |
| | 3.16 |
| | -2.41 |
| | 2.88 |
| | 2.75 |
| | 3.48 |
| | 6.56 |
| | 6.9 |
| | 7.52 |
| | 6.63 |
| | 2.07 |
| | 8.19 |

| | |
|---|---|
| | 6.11 |
| | 7.76 |
| | 8.3 |
| | 3.44 |
| | 3.97 |
| | 5.18 |
| | 6.7 |
| | 2.71 |
| | 4.09 |
| | 9.04 |
| | 4.87 |
| | 5.64 |
| | -0.37 |
| | 5.36 |
| | 2.64 |
| | -2.55 |
| | 8.16 |
| | 4.37 |
| | 8.81 |
| | 3.26 |
| | 2.42 |
| | 5.14 |
| | 2.74 |
| | 5.39 |
| | 4.17 |
| | 3.68 |
| | 3.37 |
| | 2.24 |
| | 1.27 |
| | 5.96 |
| | 5.88 |
| | 8.31 |
| | 3.22 |
| | 8.6 |
| | 7.38 |

| | |
|---|---|
| | 4.12 |
| | 2.66 |
| | 5.41 |
| | 7.72 |
| | 1.39 |
| | 8.73 |
| | 6.32 |
| | 8.32 |
| | 3.85 |
| | 4.66 |
| | 3.46 |
| | 4.63 |
| | 3.13 |
| | 0.83 |
| | -0.62 |
| | 5.27 |
| | 2.83 |
| | 7.53 |
| | 6.54 |
| | 3.21 |
| | 3.67 |
| | 5.98 |
| | 1.62 |
| | 3.3 |
| | 1.39 |
| | 3.25 |
| | -2.5 |
| | 2.3 |
| | 1.57 |
| | 0.42 |
| | 4.49 |
| | 3.62 |
| | 3.18 |
| | 3.42 |
| | 1.66 |

| | |
|---|---|
| | 2.09 |
| | 4.36 |
| | -1.72 |
| | -2.46 |
| | 3.73 |
| | 1.84 |
| | 4.42 |
| | 4.99 |
| | 4.66 |
| | 9.02 |
| | 5.59 |
| | 2.19 |
| | 5.56 |
| | 2.76 |
| | 3.04 |
| | 1.11 |
| | 3.7 |
| | 3.83 |
| | 1.2 |
| | 4.48 |
| | 10.27 |
| | 4.79 |
| | 3.05 |
| | 4.97 |
| | 9.53 |
| | 0.58 |
| | 6.35 |
| | 1.45 |
| | 2.81 |
| | 2.99 |
| | 4.86 |
| | -0.54 |
| | 6.23 |
| | -0.13 |
| | 1.11 |

| | |
|---|---|
| | 4.35 |
| | -0.24 |
| | 4.2 |
| | 3.03 |
| | -0.21 |
| | 6.45 |
| | 6.41 |
| | 7.03 |
| | 10.01 |
| | 2.52 |
| | 7.09 |
| | 4.82 |
| | 3.14 |
| | 9.59 |
| | 7.28 |
| | 6.32 |
| | 5.86 |
| | 6.92 |
| | 7.21 |
| | 2.74 |
| | 4.46 |
| | 2.09 |
| | 2.39 |
| | 4.38 |
| | 4.16 |
| | 2.82 |
| | 9.16 |
| | 5.09 |
| | 3.85 |
| | 3.85 |
| | 6.36 |
| | 3.29 |
| | 7.45 |
| | 6.1 |
| | 5.91 |

| | |
|---|---|
| | 7.27 |
| | 3.85 |
| | 7.46 |
| | 3.91 |
| | 3.5 |
| | 0.69 |
| | -0.3 |
| | 4.62 |
| | 3.87 |
| | 7.79 |
| | 8.31 |
| | 7.63 |
| | 8.55 |
| | 3.05 |
| | 4.2 |
| | 4.73 |
| | 7.24 |
| | 7.49 |
| | 2.81 |
| | 7.82 |
| | 7.81 |
| | 9.8 |
| | 3.5 |
| | 4.81 |
| | 2.78 |
| | 2.11 |
| | 6.29 |
| | 8.16 |
| | 9.55 |
| | 2.99 |
| | 3.81 |
| | 4.24 |
| | 7.74 |
| | 6.27 |
| | 6.44 |

| | |
|---|---|
| | 7.16 |
| | 4.71 |
| | 6.03 |
| | 2.66 |
| | 1.19 |
| | 6.94 |
| | 8.97 |
| | 10.08 |
| | 1.01 |
| | 8.04 |
| | 8.15 |
| | 7.35 |
| | 5.34 |
| | 3.3 |
| | 5.68 |
| | 7.48 |
| | 7.94 |
| | 5.15 |
| | 4.83 |
| | 6.83 |
| | 4.38 |
| | 5.37 |
| | 6.84 |
| | 8.86 |
| | 7.25 |
| | 6.06 |
| | 2.71 |
| | 3.83 |
| | 8.89 |
| | 6.49 |
| | 10.55 |
| | 6.73 |
| | 9.45 |
| | 7.7 |
| | 6.11 |

| |
|---|
| -3.05 |
| -3.43 |
| 3.03 |
| 7.12 |
| 4.05 |
| 3.49 |
| -0.72 |
| 0.02 |
| -2.74 |
| -3.52 |
| -2.07 |
| 6.8 |
| -1.95 |
| 5.47 |
| 4.5 |
| 4.74 |
| 6.04 |
| 2.39 |
| 2.83 |
| 2.68 |
| 5.23 |
| 4.31 |
| -0.03 |
| 5.12 |
| 2.02 |
| 5.45 |
| 3.23 |
| 3.21 |
| 4.92 |
| 7.42 |
| 4.61 |
| 3.31 |
| 4.46 |
| 0.86 |
| 6.23 |

| | |
|---|---|
| | 4.3 |
| | 0.56 |
| | 8.2 |
| | -0.1 |
| | 7.08 |
| | 1.28 |
| | 6.7 |
| | -1.27 |
| | 9.59 |
| | 6.06 |
| | 0.21 |
| | 2.68 |
| | 1.95 |
| | 2.65 |
| | 4.43 |
| | 2.09 |
| | 4.43 |
| | 8.77 |
| | 6.86 |
| | 1.47 |
| | 9.56 |
| | 9.3 |
| | 9.67 |
| | 5.67 |
| | 5.25 |
| | 7.15 |
| | 10.33 |
| | 6.71 |
| | 2.71 |
| | 10.24 |
| | 1.8 |
| | 8.09 |
| | 4.43 |
| | 2.68 |
| | 3.51 |

| |
|---|
| 0.75 |
| 0.42 |
| 3.79 |
| -2.82 |
| -4.49 |
| -3.13 |
| 6.29 |
| 1.68 |
| 2.94 |
| -0.91 |
| -4.55 |
| 3.42 |
| -5.97 |
| 5.29 |
| 1.34 |
| 4.66 |
| 0.71 |
| 1.93 |
| 3.34 |
| -3.34 |
| -4.99 |
| 3.7 |
| -0.09 |
| -5.27 |
| 2.78 |
| -0.6 |
| -3.51 |
| -0.23 |
| -0.9 |
| -0.83 |
| 1.15 |
| 5.4 |
| -2.56 |
| -1.06 |
| -4.43 |

| | |
|---|---|
| | -0.06 |
| | -5.33 |
| | -5.69 |
| | -6.12 |
| | -6.76 |
| | 5.66 |
| | 6.33 |
| | 2.64 |
| | 5.5 |
| | -1.69 |
| | 1.41 |
| | 0.82 |
| | 4.2 |
| | -0.76 |
| | 7.57 |
| | -2.8 |
| | 2.21 |
| | -2.49 |
| | -0.69 |
| | 4.05 |
| | -1.74 |
| | -7.19 |
| | 0.38 |
| | -4.02 |
| | -7.47 |
| | -4.66 |
| | -2.78 |
| | 2.06 |
| | 2.39 |
| | -3.15 |
| | 3.06 |
| | -0.27 |
| | 2.07 |
| | -4.05 |
| | 5.11 |

| | |
|---|---|
| | -0.42 |
| | -3.79 |
| | 0.76 |
| | 2.36 |
| | 4.42 |
| | 2.33 |
| | 1.63 |
| | 0.59 |
| | 2.85 |
| | 0.16 |
| | -1.51 |
| | -4.32 |
| | 1.44 |
| | -4.32 |
| | -4.9 |
| | -0.85 |
| | -3.79 |
| | -2.47 |
| | -3.7 |
| | -2.94 |
| | 2.1 |
| | -2.88 |
| | -3.12 |
| | 3.82 |
| | -2.85 |
| | -3.36 |
| | -2.7 |
| | -3.55 |
| | 1.84 |
| | 0.99 |
| | -4.32 |
| | -2.2 |
| | 1.43 |
| | -1.34 |
| | 2 |

| | |
|---|---|
| | -0.14 |
| | -2.84 |
| | -1.22 |
| | 3.75 |
| | 4.29 |
| | 3.74 |
| | 0.93 |
| | 2.93 |
| | 0.21 |
| | 2.7 |
| | 2.7 |
| | 1.69 |
| | 2.18 |
| | 0.2 |
| | -2.56 |
| | 3.35 |
| | 2.41 |
| | -4.65 |
| | 0.73 |
| | 1.24 |
| | -3.03 |
| | 1.3 |
| | 3.1 |
| | 5.65 |
| | -0.05 |
| | 5.67 |
| | 2.63 |
| | 1.18 |
| | 4.65 |
| | -1.18 |
| | 2.47 |
| | 0.19 |
| | -1.94 |
| | -0.53 |
| | 5.06 |

| |
|---|
| 4.57 |
| -0.35 |
| 4.14 |
| 8.03 |
| 7.13 |
| 5.46 |
| 5.71 |
| 1.74 |
| -1.79 |
| 3.99 |
| 3.65 |
| -6.32 |
| -5.49 |
| 2.09 |
| 8.24 |

| |
|---|
| Log2 Fold Change: (Rabbitserum_12022024_Newanalysis) / (Water_pos_12022024_Newanalysis) |
| 1.48 |
| 5.02 |
| 5 |
| 7.43 |
| 3.22 |
| 1.89 |
| 3.37 |
| 2.84 |
| 1.32 |
| -0.63 |
| -0.05 |
| 3.09 |
| 2.91 |
| 2.86 |
| -1.62 |
| -0.63 |
| -1.12 |
| 6.67 |
| 3.13 |
| 7.05 |
| 0.18 |
| 5.8 |
| 5.57 |
| 6.23 |
| 6.69 |
| -0.71 |
| -0.63 |
| -1.86 |
| 0.38 |
| -3.01 |
| -2.27 |
| -2.3 |
| -3.19 |

| | |
|---|---|
| | 0.76 |
| | 5.31 |
| | 5.75 |
| | 4.41 |
| | -1.21 |
| | -0.95 |
| | 4.13 |
| | -0.88 |
| | 0.64 |
| | 2.18 |
| | 2.97 |
| | -2.26 |
| | 4.72 |
| | 4.81 |
| | 4.84 |
| | 3.88 |
| | 2.23 |
| | 4.31 |
| | 3.1 |
| | 5.03 |
| | 3.1 |
| | -0.14 |
| | -1.25 |
| | 2.44 |
| | -1.34 |
| | 2.01 |
| | 3.29 |
| | -0.58 |
| | -0.13 |
| | 2.81 |
| | 6.11 |
| | 2.14 |
| | -0.81 |
| | -0.02 |
| | 3.68 |

| | |
|---|---|
| | 2.65 |
| | 0.28 |
| | -1.82 |
| | -2.05 |
| | 2.7 |
| | 3.77 |
| | 1.6 |
| | -0.05 |
| | 6.01 |
| | 4.71 |
| | 4.5 |
| | 5.11 |
| | 2.55 |
| | -1.5 |
| | -2.32 |
| | 2.25 |
| | 5.38 |
| | -0.37 |
| | -3.2 |
| | 2.47 |
| | 0.75 |
| | -0.79 |
| | 3.16 |
| | 4.38 |
| | 3.17 |
| | 3.7 |
| | 3.15 |
| | 2.2 |
| | 2.94 |
| | 2.29 |
| | -1.82 |
| | 0.19 |
| | 5.26 |
| | -3.8 |
| | 2.14 |

| | |
|---|---|
| | -2.99 |
| | 2.82 |
| | -2.77 |
| | 1.34 |
| | -0.22 |
| | 0.41 |
| | -2.89 |
| | 1.39 |
| | 4.01 |
| | -0.96 |
| | 2.9 |
| | -2.41 |
| | 0.77 |
| | -1.53 |
| | 6.23 |
| | 4.32 |
| | -0.71 |
| | -1.41 |
| | 5.71 |
| | 0.48 |
| | -1.21 |
| | -0.71 |
| | 2.49 |
| | 2.03 |
| | 0.43 |
| | 0.54 |
| | -2.21 |
| | -0.58 |
| | 4.02 |
| | 7.86 |
| | 3.48 |
| | 2.41 |
| | -0.22 |
| | 0.28 |
| | 2.2 |

| | |
|---|---|
| | -1.06 |
| | 0.72 |
| | -0.16 |
| | -0.25 |
| | -2.34 |
| | -2.26 |
| | -1.76 |
| | -2.12 |
| | -1.21 |
| | -0.23 |
| | -0.85 |
| | 3.52 |
| | 6.14 |
| | 0.94 |
| | 3.76 |
| | 4.72 |
| | -0.99 |
| | 1.22 |
| | 0.91 |
| | 1.17 |
| | 1.2 |
| | 3.02 |
| | 4.48 |
| | 4.03 |
| | 1.41 |
| | 3.15 |
| | 3.67 |
| | -1 |
| | 1.66 |
| | 2.98 |
| | 6.68 |
| | 0.85 |
| | 4.48 |
| | 2.77 |
| | 6.06 |

| |
|---|
| 3.01 |
| 5.86 |
| 3.45 |
| 7.14 |
| 5.02 |
| 4.72 |
| 4.71 |
| 2.51 |
| 4.18 |
| 6.1 |
| 4.07 |
| -1.17 |
| 6.28 |
| 5.36 |
| 0.11 |
| 5.93 |
| 2.77 |
| 2.16 |
| 5.73 |
| 3.05 |
| 3.77 |
| 3.12 |
| 2.53 |
| -1.22 |
| 0.49 |
| 0.74 |
| 2.58 |
| 0.46 |
| 1.83 |
| 5.79 |
| 2.6 |
| 2.42 |
| 6.08 |
| 7.05 |
| 8.11 |

| | |
|---|---|
| | 1.83 |
| | 1.08 |
| | 0.36 |
| | -0.57 |
| | 0.74 |
| | -0.61 |
| | 1.67 |
| | 2.87 |
| | 0.24 |
| | 2.66 |
| | -0.88 |
| | -0.28 |
| | 1.97 |
| | 1.56 |
| | -0.05 |
| | 0.52 |
| | 2.1 |
| | 3.72 |
| | 2.78 |
| | -0.87 |
| | 3.94 |
| | 1.44 |
| | -3.47 |
| | 2.97 |
| | 3.82 |
| | 3.87 |
| | -1.01 |
| | -0.52 |
| | -0.47 |
| | -0.43 |
| | 1.86 |
| | -0.47 |
| | 1.45 |
| | 0.28 |
| | 3.52 |

| | |
|---|---|
| | 2.65 |
| | 3.4 |
| | -0.5 |
| | 0.71 |
| | 2.59 |
| | 2.54 |
| | 4.49 |
| | -0.64 |
| | 1.03 |
| | 6.05 |
| | 4.03 |
| | 2.55 |
| | 2.57 |
| | 5.11 |
| | 3.56 |
| | 4.14 |
| | 1.52 |
| | 1.87 |
| | -1.67 |
| | 7.04 |
| | 2.31 |
| | 2.59 |
| | 2.68 |
| | 0.49 |
| | 1.77 |
| | -0.4 |
| | 3.4 |
| | 1.05 |
| | 2.99 |
| | 2.55 |
| | 2.84 |
| | 3.75 |
| | 0.51 |
| | 0.92 |
| | 6.34 |

| | |
|---|---|
| | 5.2 |
| | 5.27 |
| | 1.38 |
| | 5.19 |
| | 2.96 |
| | 3.2 |
| | 5.22 |
| | -0.5 |
| | -1.27 |
| | 3.2 |
| | 1 |
| | 1.47 |
| | -2.13 |
| | -0.39 |
| | -2.17 |
| | -0.42 |
| | 5.76 |
| | 0.75 |
| | 1.42 |
| | 1.84 |
| | 4.9 |
| | 4.3 |
| | -1.72 |
| | 5.02 |
| | 2.63 |
| | 2.6 |
| | 3.24 |
| | 3.48 |
| | 3.81 |
| | 7.22 |
| | 2.87 |
| | 6.39 |
| | 3.46 |
| | 7.27 |
| | 0.8 |

| | |
|---|---|
| | 5.89 |
| | 3.06 |
| | 3.45 |
| | 7.71 |
| | 2.05 |
| | 3.66 |
| | 6.66 |
| | 5.9 |
| | 3.32 |
| | 4.26 |
| | 2.84 |
| | 1.27 |
| | 3.08 |
| | 1.38 |
| | -0.19 |
| | 6.73 |
| | 4.05 |
| | 5.29 |
| | 5.58 |
| | 5.74 |
| | 2.97 |
| | 5 |
| | 2.22 |
| | 6.14 |
| | 3.5 |
| | 0.96 |
| | -1.97 |
| | 4.64 |
| | 1.53 |
| | 5.58 |
| | 5.47 |
| | 2.31 |
| | 4.16 |
| | 1.52 |
| | 0.89 |

| | |
|---|---|
| | 0.31 |
| | 0.45 |
| | -0.84 |
| | -1.47 |
| | 2.91 |
| | -0.47 |
| | 1.6 |
| | 3.82 |
| | 1.89 |
| | 5.02 |
| | 5.93 |
| | 3.25 |
| | 5.04 |
| | 2.67 |
| | -0.45 |
| | -0.01 |
| | 1.21 |
| | 1.75 |
| | 5.31 |
| | 3.74 |
| | 5.24 |
| | 4.65 |
| | -0.96 |
| | 3.77 |
| | 7.29 |
| | 0.63 |
| | 3.79 |
| | 3.66 |
| | 1.63 |
| | 2.94 |
| | 2.12 |
| | 5.43 |
| | 4.68 |
| | 0.84 |
| | 0.22 |

| |
|---|
| 3.37 |
| -0.79 |
| 4.33 |
| 2.82 |
| 4.24 |
| 2.69 |
| 3.94 |
| 5.23 |
| 8 |
| 3.97 |
| 5.15 |
| 6.94 |
| 2.53 |
| 7.63 |
| 8.5 |
| 1.93 |
| 2.32 |
| 5.06 |
| 5.57 |
| 1.58 |
| 4.52 |
| 1.84 |
| 1.03 |
| 3.12 |
| 5.27 |
| 0.34 |
| 6.96 |
| 7.09 |
| -2.28 |
| 1.11 |
| 6.83 |
| 0.27 |
| 4.71 |
| 4.82 |
| 4.4 |

| | |
|---|---|
| | 6.27 |
| | 3 |
| | 7.29 |
| | 8.09 |
| | 5.96 |
| | 0.58 |
| | -0.07 |
| | 0.59 |
| | 4.43 |
| | 7.8 |
| | 2.69 |
| | 5.87 |
| | 7.97 |
| | 2.21 |
| | 6.74 |
| | 5.9 |
| | 7.17 |
| | 2.8 |
| | 3.91 |
| | 6.98 |
| | 3.48 |
| | 7.6 |
| | 4.46 |
| | 5.47 |
| | 4.81 |
| | 3.68 |
| | 4.69 |
| | 6.74 |
| | 1.7 |
| | 4.6 |
| | 3.72 |
| | 5.28 |
| | 8.3 |
| | 8.07 |
| | 7.84 |

| | |
|---|---|
| | 6.62 |
| | 3.52 |
| | 5.08 |
| | 0.68 |
| | -1.12 |
| | 5.29 |
| | 5.68 |
| | 5.28 |
| | 1.92 |
| | 8.45 |
| | 7.98 |
| | 6.31 |
| | 6.32 |
| | 1.77 |
| | 5.46 |
| | 0.74 |
| | 3.85 |
| | -0.81 |
| | 1.09 |
| | 4.53 |
| | 3 |
| | 2.68 |
| | 4.75 |
| | 8.38 |
| | 7.14 |
| | 3.19 |
| | 3.65 |
| | 5.52 |
| | -0.15 |
| | 0.45 |
| | 3.81 |
| | 1.29 |
| | 4.96 |
| | 0.11 |
| | -0.74 |

| |
|---|
| -1.82 |
| -1.51 |
| -1.06 |
| 4.8 |
| 2.79 |
| 5.42 |
| 1 |
| 0.74 |
| -3.38 |
| -1.79 |
| -2.53 |
| 7.22 |
| 3.97 |
| 3.26 |
| 2.57 |
| 4.58 |
| 3.87 |
| 2.84 |
| 2.66 |
| 1.53 |
| 3.38 |
| 2.51 |
| 0.22 |
| -0.75 |
| 2.94 |
| 4.07 |
| 2.98 |
| 5.64 |
| 3.58 |
| 1.38 |
| 3.46 |
| 5.64 |
| 5.05 |
| -0.09 |
| 4.56 |

| | |
|---|---|
| | 3.78 |
| | 1.85 |
| | 5.04 |
| | -0.47 |
| | 6.46 |
| | 1.35 |
| | 5.98 |
| | -0.22 |
| | 8.83 |
| | 6.72 |
| | 2.19 |
| | 3.43 |
| | 4.48 |
| | 0.92 |
| | 4.78 |
| | 1.62 |
| | 5.74 |
| | 7.26 |
| | 5.26 |
| | 1.46 |
| | 7.21 |
| | 8.79 |
| | 8.95 |
| | 4.04 |
| | 4.57 |
| | 5.5 |
| | 8.89 |
| | 3.69 |
| | 2.28 |
| | 7.44 |
| | 2.06 |
| | 1.96 |
| | -1.41 |
| | -0.08 |
| | 4.1 |

| |
|---|
| 3.52 |
| 0.93 |
| 4.62 |
| -2.87 |
| -3.79 |
| -1.86 |
| -0.25 |
| 4.18 |
| 2.36 |
| -0.12 |
| -2.91 |
| 1.17 |
| -4.94 |
| 4.22 |
| 2.71 |
| 3.68 |
| 4.37 |
| -0.55 |
| 1.6 |
| -2.36 |
| -2.7 |
| 4.36 |
| -0.04 |
| -3.94 |
| 1.07 |
| -0.49 |
| -2.68 |
| -0.73 |
| -1.15 |
| 0.5 |
| 2.39 |
| 4.48 |
| 0.41 |
| 4.44 |
| -0.98 |

| |
|---|
| -1.18 |
| -1.7 |
| -2.95 |
| -3.82 |
| -2.28 |
| 3.67 |
| 6.17 |
| 3.7 |
| 6.4 |
| -1.08 |
| 2.05 |
| 1.21 |
| 3.26 |
| -0.35 |
| 6.38 |
| -4.66 |
| 1.64 |
| -0.94 |
| -2.25 |
| 2.79 |
| -2.16 |
| -5.01 |
| 0.35 |
| -2.71 |
| -3.26 |
| -2.63 |
| -2.24 |
| 4.36 |
| 4.16 |
| 1.13 |
| 4.37 |
| 1.21 |
| 2.58 |
| -2.34 |
| 6.13 |

| | |
|---|---|
| | 0.54 |
| | 1.83 |
| | 6.11 |
| | 2.72 |
| | 2.98 |
| | 1.37 |
| | 2.23 |
| | 3.59 |
| | 3.96 |
| | 2.52 |
| | 2.06 |
| | -3.8 |
| | 1.64 |
| | -1.8 |
| | -3.59 |
| | -1.56 |
| | -2.32 |
| | -1.78 |
| | -2.08 |
| | -1.07 |
| | -1.27 |
| | -2.38 |
| | -2.24 |
| | 3.05 |
| | -2.17 |
| | -2.68 |
| | -2.16 |
| | -2.5 |
| | 0.86 |
| | -0.36 |
| | -3.97 |
| | -3 |
| | 2.03 |
| | 1.38 |
| | 2.39 |

| | |
|---|---|
| | -0.98 |
| | -2.44 |
| | -0.33 |
| | 3.03 |
| | 2.5 |
| | 5.13 |
| | 1.37 |
| | 3.48 |
| | 1.55 |
| | 4.95 |
| | 6.16 |
| | 4.41 |
| | 5.2 |
| | 2.75 |
| | 1.64 |
| | 7.03 |
| | 4.58 |
| | -1 |
| | -0.04 |
| | 0.79 |
| | 0.11 |
| | 0.73 |
| | 3.77 |
| | 3.02 |
| | 0.51 |
| | 2.04 |
| | 1.75 |
| | -1.83 |
| | 2.67 |
| | -0.52 |
| | 3.24 |
| | 2.69 |
| | -1.23 |
| | -2.03 |
| | 3.3 |

| | |
|---|---|
| | 1.22 |
| | 1.65 |
| | 0.61 |
| | 1.56 |
| | 3.37 |
| | 5.23 |
| | 3.51 |
| | -0.26 |
| | 1.3 |
| | -0.23 |
| | -1.82 |
| | -2.22 |
| | -1.7 |
| | -2.39 |
| | 10.24 |

| |
| ---: |
| Log2 Fold Change: (SM_pos_12022024_Newanalyis) / (Water_pos_12022024_Newanalysis) |
| 0.35 |
| 1.86 |
| 0.41 |
| 5.02 |
| -0.87 |
| 0.79 |
| 1.83 |
| 0.27 |
| 2.05 |
| 2.01 |
| 2.62 |
| 3.36 |
| 1.76 |
| 4.39 |
| 0.2 |
| 0.15 |
| 0.63 |
| 0.68 |
| -2.5 |
| 1.21 |
| -0.88 |
| 0.8 |
| 2.48 |
| 0.46 |
| 1.35 |
| 1.37 |
| -0.47 |
| -0.66 |
| -1.24 |
| 2.24 |
| -1.41 |
| -2.32 |
| 2.24 |

| | |
|---|---|
| | 4.38 |
| | 2.79 |
| | 1.31 |
| | -1.39 |
| | -1.29 |
| | -2.53 |
| | -1.9 |
| | 3.45 |
| | 0.46 |
| | 2.71 |
| | 2.13 |
| | 2.01 |
| | 0.92 |
| | 1.19 |
| | -0.42 |
| | -1.76 |
| | 0.26 |
| | -1.66 |
| | 0.26 |
| | 0.87 |
| | 0.11 |
| | -2.97 |
| | 1.25 |
| | 1.99 |
| | 1.12 |
| | 0.94 |
| | -0.38 |
| | 0.28 |
| | 3.9 |
| | -0.39 |
| | 2.99 |
| | -1.56 |
| | 2.01 |
| | -0.47 |
| | 9.44 |

| | |
|---|---|
| | 3.26 |
| | 0.42 |
| | 0.85 |
| | 3.17 |
| | 2.52 |
| | 9.25 |
| | 1.79 |
| | -1.84 |
| | 2.5 |
| | 0.85 |
| | -2.24 |
| | -4.31 |
| | -2.22 |
| | 1.57 |
| | 1.57 |
| | 2.63 |
| | 1.66 |
| | 1.31 |
| | -2.02 |
| | -2.67 |
| | 3.9 |
| | -1.47 |
| | -7.15 |
| | -2.2 |
| | 0.92 |
| | 1.43 |
| | -0.19 |
| | 0.91 |
| | -0.97 |
| | 0.29 |
| | -3.69 |
| | -0.91 |
| | 0.96 |
| | 0.77 |
| | 4.99 |

| | |
|---|---|
| | -1.8 |
| | -0.63 |
| | 0.92 |
| | 2.06 |
| | 1.13 |
| | -0.62 |
| | -0.26 |
| | 2.86 |
| | 0.02 |
| | -2.14 |
| | 4.06 |
| | 1.3 |
| | 1.28 |
| | 1.63 |
| | -0.74 |
| | -2.67 |
| | -3.05 |
| | 2.17 |
| | 0.11 |
| | -2.93 |
| | -1.34 |
| | 2.31 |
| | 0.44 |
| | 1.83 |
| | -3.25 |
| | 0.93 |
| | -1.75 |
| | -1.73 |
| | 1.53 |
| | -1.99 |
| | -0.03 |
| | -2.43 |
| | -1.4 |
| | -1.56 |
| | 2.29 |

| | |
|---|---|
| | 2.82 |
| | 0.55 |
| | -1.77 |
| | 0.96 |
| | -1.87 |
| | -1.97 |
| | -1.87 |
| | -1.06 |
| | -1.07 |
| | -0.84 |
| | -0.16 |
| | 1.4 |
| | 1.98 |
| | -1.75 |
| | 0.09 |
| | 2.41 |
| | 2.36 |
| | 1.35 |
| | 0.01 |
| | -0.15 |
| | 1.28 |
| | 1.15 |
| | 0.53 |
| | 1.35 |
| | 4.44 |
| | -5.35 |
| | -1.51 |
| | -0.93 |
| | 0.95 |
| | -0.27 |
| | 3.91 |
| | 0.31 |
| | -1.51 |
| | 3.04 |
| | 3.49 |

| | |
|---|---|
| | 3.83 |
| | -0.35 |
| | -2.31 |
| | 3.16 |
| | 2.19 |
| | 0.41 |
| | 3.99 |
| | 1 |
| | 5.61 |
| | 4.06 |
| | -1.31 |
| | 1.86 |
| | 2.36 |
| | 3.75 |
| | 0.75 |
| | 4.28 |
| | 0.57 |
| | 4.97 |
| | 3.66 |
| | 0.17 |
| | 1.77 |
| | 1.11 |
| | 0.28 |
| | 0.59 |
| | -0.78 |
| | -0.27 |
| | -1.52 |
| | 3.57 |
| | 0.13 |
| | -4.35 |
| | 3.59 |
| | -3.5 |
| | 3.28 |
| | -0.17 |
| | 9.57 |

| | |
|---|---|
| | 0.29 |
| | 0.86 |
| | 1.03 |
| | 0.92 |
| | 0.35 |
| | 4.85 |
| | 4.79 |
| | 2.56 |
| | 3.23 |
| | 3.35 |
| | -0.46 |
| | 0.33 |
| | 2.18 |
| | 2.53 |
| | -0.45 |
| | -1.32 |
| | -1.54 |
| | -2.63 |
| | 0.2 |
| | -2.98 |
| | 2.4 |
| | -1.6 |
| | -3.85 |
| | 3.32 |
| | 1.01 |
| | -2.6 |
| | 1.12 |
| | -1.81 |
| | -0.99 |
| | -3.17 |
| | 0.86 |
| | 0.65 |
| | -0.26 |
| | -1.24 |
| | -0.2 |

| | |
|---|---|
| | 2.23 |
| | 1.96 |
| | 0.26 |
| | 0.47 |
| | -1.33 |
| | 2.13 |
| | 2.45 |
| | -0.36 |
| | 1.28 |
| | 3.27 |
| | 2.71 |
| | 1.78 |
| | -0.41 |
| | 2.76 |
| | 3.12 |
| | 3.02 |
| | -2.11 |
| | 2.13 |
| | -0.2 |
| | -0.16 |
| | 1.06 |
| | 5.64 |
| | -0.19 |
| | 4.58 |
| | -2.48 |
| | 0.01 |
| | 1.88 |
| | 0.4 |
| | 2.67 |
| | 1.04 |
| | 2.44 |
| | 0.62 |
| | 0.85 |
| | -0.14 |
| | 4.01 |

| |
|---|
| 3.13 |
| -0.49 |
| 0.59 |
| 4.2 |
| 5.91 |
| 2 |
| 1.97 |
| -0.6 |
| 1 |
| -0.09 |
| 2.51 |
| -1.71 |
| -1.79 |
| 0.37 |
| 0.79 |
| 1.23 |
| -1.02 |
| -1.69 |
| 2.3 |
| -0.35 |
| 0.24 |
| -1.27 |
| -0.35 |
| 1.21 |
| -1.12 |
| -1.38 |
| 0.38 |
| 0.5 |
| 0.84 |
| 5 |
| -1.77 |
| 3.96 |
| 3.28 |
| 10.19 |
| 0.6 |

| | |
|---|---|
| | 3.24 |
| | 0.5 |
| | 0.7 |
| | 0.8 |
| | 4.26 |
| | 1.66 |
| | 5.22 |
| | 1.22 |
| | 8.86 |
| | 2.18 |
| | 2.39 |
| | 3.38 |
| | 4.12 |
| | 2.05 |
| | 4.08 |
| | 4.7 |
| | 1.86 |
| | 0.91 |
| | 1.4 |
| | 2.69 |
| | 5.8 |
| | 3.64 |
| | 3.05 |
| | 1.41 |
| | 0.53 |
| | 4.68 |
| | 1.01 |
| | 1.27 |
| | 2.5 |
| | 2.81 |
| | 2.86 |
| | 1.28 |
| | 2.04 |
| | 1.38 |
| | 0.09 |

| |
|---|
| -1.77 |
| -1.56 |
| -1.18 |
| 1.71 |
| -1.13 |
| -0.82 |
| -1.76 |
| 0.65 |
| -0.5 |
| 3.11 |
| 2.33 |
| 0.41 |
| 1.91 |
| 0.08 |
| 0.35 |
| -5.56 |
| -0.81 |
| -1.25 |
| 3.31 |
| 1.27 |
| 2.58 |
| 2.25 |
| 1.71 |
| -0.17 |
| 4.4 |
| 1 |
| 1.33 |
| 0.16 |
| -3.21 |
| 2.31 |
| -0.35 |
| -1.4 |
| 0.41 |
| -2.65 |
| 3.78 |

| | |
|---|---|
| | 3.31 |
| | 2.77 |
| | 0.34 |
| | 0.46 |
| | -0.9 |
| | -0.95 |
| | 0.65 |
| | 1.76 |
| | 1.18 |
| | 0.83 |
| | 2.39 |
| | 1.29 |
| | -1.6 |
| | 1.18 |
| | 7.71 |
| | -0.43 |
| | -0.6 |
| | 3.51 |
| | 2.41 |
| | 2.6 |
| | 5.74 |
| | 2.05 |
| | -1.66 |
| | -0.59 |
| | 6.21 |
| | 0.96 |
| | -3.7 |
| | 9.57 |
| | -2.01 |
| | -2.03 |
| | -0.58 |
| | 0.91 |
| | 3.3 |
| | 3.88 |
| | 6.42 |

| | |
|---|---|
| | 3.14 |
| | 6.63 |
| | 7.67 |
| | 8.95 |
| | 8.96 |
| | 1 |
| | 5.14 |
| | -1.51 |
| | 1.14 |
| | 4.04 |
| | 0.4 |
| | 3.29 |
| | 4.43 |
| | 4.07 |
| | 4.46 |
| | 6.17 |
| | 1.11 |
| | -0.39 |
| | 4.05 |
| | 6.46 |
| | 1.04 |
| | 4.58 |
| | 3.06 |
| | 5.94 |
| | 6.27 |
| | 6.37 |
| | 2.54 |
| | -0.42 |
| | 1.46 |
| | -1.53 |
| | 2.64 |
| | 3.72 |
| | 6.03 |
| | 3.93 |
| | 5.06 |

| |
|---|
| -1.09 |
| -1.95 |
| 1.59 |
| 2.99 |
| 0.42 |
| 4.72 |
| 4.86 |
| 5.13 |
| -1.14 |
| 4.9 |
| 6.66 |
| 9.35 |
| 2.29 |
| 3.58 |
| 2.8 |
| 0.9 |
| 1.56 |
| 1.93 |
| 0.38 |
| 1.87 |
| -1.54 |
| 2.18 |
| 0.62 |
| 4.85 |
| 3.88 |
| 0.63 |
| 4.5 |
| 1.84 |
| -1.34 |
| -0.59 |
| 0.82 |
| -1.28 |
| 2.15 |
| 0.08 |
| 1.32 |

| | |
|---|---|
| | 0.07 |
| | -3.39 |
| | 0.68 |
| | 2.24 |
| | 1.77 |
| | 2.63 |
| | 3.49 |
| | -1.68 |
| | 0.82 |
| | 1.42 |
| | 0.74 |
| | 5.13 |
| | -0.84 |
| | 4.5 |
| | 0.87 |
| | 2.1 |
| | 1.89 |
| | 2.57 |
| | -0.64 |
| | 3.39 |
| | 2.5 |
| | 4.69 |
| | 5.31 |
| | -0.95 |
| | 2.16 |
| | 0.18 |
| | 2.7 |
| | -1.33 |
| | 0.27 |
| | -1.31 |
| | 2.61 |
| | 2.7 |
| | 0.92 |
| | -1.04 |
| | 1.6 |

| | |
|---|---|
| | 4.81 |
| | 1.95 |
| | 4.59 |
| | 1.1 |
| | 8.65 |
| | -1.51 |
| | 1.7 |
| | -0.2 |
| | 1.22 |
| | 3.36 |
| | 0.93 |
| | 1.91 |
| | 3.36 |
| | -0.2 |
| | 0.54 |
| | 2.08 |
| | 3.61 |
| | 9.45 |
| | 2.62 |
| | 2.53 |
| | 5.62 |
| | -1.29 |
| | 12.16 |
| | 0.28 |
| | 6.62 |
| | 2.06 |
| | 1.31 |
| | 4.24 |
| | 6.81 |
| | 9.2 |
| | 0.99 |
| | -0.59 |
| | 0.41 |
| | -1.41 |
| | -0.02 |

| |
|---|
| -1.66 |
| -0.45 |
| 1.28 |
| -2.97 |
| -2.23 |
| -2.05 |
| 0.96 |
| 3.33 |
| 0.44 |
| 2.54 |
| -3.74 |
| 0.16 |
| -6.96 |
| 0.22 |
| 1.55 |
| 1.4 |
| -1.39 |
| 1.51 |
| 2.38 |
| -1.61 |
| -5.8 |
| 1.01 |
| 0.76 |
| -0.09 |
| -0.64 |
| -0.88 |
| 2.43 |
| -0.18 |
| 0.17 |
| 4.25 |
| 1.72 |
| 3.64 |
| -1.95 |
| 4.97 |
| 0.82 |

| | |
|---|---|
| | 2.14 |
| | -2.55 |
| | -1.69 |
| | -2.5 |
| | -4.13 |
| | 0.73 |
| | 0.87 |
| | 0.09 |
| | 3.2 |
| | 4.03 |
| | -1.37 |
| | -0.96 |
| | -1.41 |
| | 0.3 |
| | 2.86 |
| | -2.22 |
| | 0.05 |
| | -2.16 |
| | 0.86 |
| | 1.77 |
| | -1.67 |
| | -2.78 |
| | 0.05 |
| | 1.15 |
| | 0.61 |
| | -2.66 |
| | 1.64 |
| | -0.02 |
| | 3.59 |
| | 1.26 |
| | 0.57 |
| | 1.34 |
| | -0.54 |
| | 2.29 |
| | 2.35 |

| | |
|---|---|
| | 3.76 |
| | -3.26 |
| | -0.63 |
| | 2 |
| | -5.26 |
| | 3.51 |
| | 2.89 |
| | 7.4 |
| | 4.89 |
| | 0.26 |
| | -3.54 |
| | -3.89 |
| | 0.67 |
| | -4.38 |
| | 0.25 |
| | 0.77 |
| | -3.26 |
| | -0.41 |
| | -3.13 |
| | -0.48 |
| | -3.19 |
| | -2.02 |
| | -1.99 |
| | 1.71 |
| | -1.66 |
| | -1.92 |
| | -1.45 |
| | 1.19 |
| | 1.28 |
| | 2.78 |
| | -0.65 |
| | -0.95 |
| | 1.15 |
| | -4.38 |
| | 1.93 |

| | |
|---|---|
| | 1.44 |
| | -1.91 |
| | 6.09 |
| | 2.38 |
| | -1.31 |
| | 0.61 |
| | 0.93 |
| | -3.32 |
| | 1.11 |
| | 1.98 |
| | -3.19 |
| | -3.48 |
| | -1.29 |
| | -1.58 |
| | -2.04 |
| | 4.49 |
| | 0.3 |
| | 1.54 |
| | 1.93 |
| | -0.51 |
| | -1.12 |
| | 0.06 |
| | 1.05 |
| | 0.14 |
| | -2.05 |
| | 0.35 |
| | -2.53 |
| | -1.37 |
| | 3.25 |
| | 1.19 |
| | -3.79 |
| | 1.37 |
| | -1.15 |
| | -0.5 |
| | 1.84 |

| |
|---|
| -0.26 |
| 2.48 |
| 0.01 |
| 2.81 |
| 0.87 |
| 1.95 |
| 0 |
| 0.5 |
| 3.21 |
| 0.13 |
| -1.29 |
| -0.36 |
| -2.3 |
| -1.79 |
| 12.8 |

| |
|---|
| P-value: (Bb_pos_12022024_Newanalysis) / (CM_pos_12022024_Newanalysis) |
| 0.870987968 |
| 0.673670258 |
| 0.999998164 |
| 0.009318927 |
| 0.96212471 |
| 0.994816111 |
| 0.849455964 |
| 0.004531536 |
| 0.943333222 |
| 0.989866628 |
| 0.988272649 |
| 0.855251059 |
| 0.419125466 |
| 0.563841336 |
| 0.999989085 |
| 0.156986726 |
| 0.772913959 |
| 4.71845E-14 |
| 0.485065133 |
| 0.000709477 |
| 0.607548181 |
| 3.67363E-11 |
| 6.40848E-05 |
| 4.70179E-13 |
| 4.60743E-14 |
| 0.884564236 |
| 0.984889361 |
| 0.979086464 |
| 0.188841878 |
| 0.996201791 |
| 0.997488276 |
| 2.96199E-08 |
| 0.429894226 |

| | |
|---|---|
| | 0.507478462 |
| | 2.42362E-13 |
| | 0.101088564 |
| | 3.58497E-05 |
| | 0.000667007 |
| | 0.00032676 |
| | 1.03781E-08 |
| | 0.998049356 |
| | 0.627619596 |
| | 0.001879044 |
| | 0.032503894 |
| | 0.992503921 |
| | 1.23283E-10 |
| | 1.43879E-08 |
| | 4.00463E-08 |
| | 0.002318725 |
| | 0.000246192 |
| | 0.00022244 |
| | 3.71578E-06 |
| | 1.46271E-10 |
| | 0.000266485 |
| | 0.851270356 |
| | 0.639220761 |
| | 0.006793918 |
| | 0.637707033 |
| | 0.913801392 |
| | 0.040300262 |
| | 0.828251964 |
| | 0.999918356 |
| | 0.207105717 |
| | 1.11378E-08 |
| | 0.000810236 |
| | 0.998490004 |
| | 6.44007E-12 |
| | 0.565078953 |

| |
|---:|
| 0.833279451 |
| 0.784408921 |
| 4.99158E-07 |
| 0.995339239 |
| 0.000407144 |
| 0.045137496 |
| 0.9981661 |
| 0.545321157 |
| 7.78266E-14 |
| 0.984647755 |
| 5.74619E-11 |
| 4.94946E-08 |
| 5.3242E-05 |
| 3.12972E-13 |
| 0.999974478 |
| 4.77282E-05 |
| 0.089385422 |
| 0.027773786 |
| 0.976396797 |
| 1.38698E-07 |
| 0.056155692 |
| 0.105133494 |
| 1.46044E-05 |
| 4.76214E-09 |
| 8.80987E-06 |
| 1.33435E-05 |
| 0.000104856 |
| 0.020783888 |
| 2.78496E-09 |
| 0.00012872 |
| 0.999977404 |
| 0.25503844 |
| 0.028820526 |
| 0.541969219 |
| 0.000405695 |

| | |
|---|---|
| | 0.994536567 |
| | 0.00381531 |
| | 0.957445303 |
| | 0.124058537 |
| | 0.998019743 |
| | 0.931528917 |
| | 0.99999999 |
| | 0.06474324 |
| | 0.696928917 |
| | 4.64774E-05 |
| | 0.035671212 |
| | 0.99333678 |
| | 0.269585443 |
| | 0.384683842 |
| | 4.60743E-14 |
| | 0.000638573 |
| | 0.947331917 |
| | 0.972071404 |
| | 0.999998123 |
| | 0.999707728 |
| | 0.186245904 |
| | 0.992988703 |
| | 0.998679763 |
| | 0.127250904 |
| | 0.9441896 |
| | 0.999936783 |
| | 0.999755914 |
| | 0.968553793 |
| | 0.181303446 |
| | 8.31944E-09 |
| | 0.01009382 |
| | 0.004971323 |
| | 0.004864235 |
| | 0.002554648 |
| | 0.985304371 |

| |
|---|
| 0.999939715 |
| 0.744779982 |
| 0.489161973 |
| 0.926041137 |
| 0.996303722 |
| 0.999976276 |
| 0.864037057 |
| 0.963938423 |
| 0.001782239 |
| 0.999999588 |
| 0.436379439 |
| 0.990828228 |
| 5.64869E-07 |
| 0.655017869 |
| 0.4884298 |
| 9.14293E-09 |
| 0.999999606 |
| 0.677642686 |
| 0.063393824 |
| 0.581223728 |
| 0.477129378 |
| 0.433294823 |
| 0.337714634 |
| 1.94356E-05 |
| 0.999999998 |
| 0.000633301 |
| 0.200577542 |
| 0.56711836 |
| 0.641805252 |
| 0.961309087 |
| 0.180235498 |
| 0.151694086 |
| 0.2109265 |
| 0.799896374 |
| 0.075512229 |

| |
|---|
| 0.048259076 |
| 0.7876729 |
| 0.001388471 |
| 3.2824E-06 |
| 0.000279122 |
| 0.538364888 |
| 0.061517287 |
| 0.948463169 |
| 0.031067541 |
| 0.844505259 |
| 0.316864092 |
| 0.82122231 |
| 5.72168E-11 |
| 0.000670394 |
| 0.239492694 |
| 0.363416123 |
| 0.997976184 |
| 0.146659298 |
| 1.68421E-13 |
| 0.004614732 |
| 0.021301741 |
| 0.007487853 |
| 0.118990061 |
| 0.843810523 |
| 0.041234387 |
| 0.991992728 |
| 0.124969316 |
| 0.367072781 |
| 0.000954265 |
| 9.46315E-05 |
| 5.01638E-05 |
| 0.003953654 |
| 0.020326828 |
| 0.194271454 |
| 0.14698541 |

| |
|---|
| 7.86702E-06 |
| 0.998972467 |
| 0.65780379 |
| 0.596947818 |
| 0.227129397 |
| 0.999976634 |
| 0.515323112 |
| 0.999904348 |
| 0.998945549 |
| 0.726352584 |
| 0.233623729 |
| 0.627812524 |
| 0.132337498 |
| 0.994637061 |
| 0.026512218 |
| 0.000428371 |
| 9.25548E-05 |
| 0.014686535 |
| 2.51365E-05 |
| 0.986710556 |
| 0.003349943 |
| 0.911160902 |
| 0.981106035 |
| 0.725926129 |
| 3.97735E-05 |
| 0.089508101 |
| 0.999999994 |
| 0.989322188 |
| 0.050812752 |
| 1 |
| 0.728139945 |
| 0.972829654 |
| 0.541532707 |
| 0.998534732 |
| 8.37117E-07 |

| | |
|---|---|
| | 0.127760199 |
| | 0.999882891 |
| | 0.99999719 |
| | 0.863564262 |
| | 0.999311173 |
| | 0.009435034 |
| | 0.119884029 |
| | 0.999999924 |
| | 0.524582566 |
| | 1.76528E-07 |
| | 0.011246097 |
| | 0.773969961 |
| | 0.829197582 |
| | 0.003205183 |
| | 5.0248E-05 |
| | 0.005383935 |
| | 0.000737979 |
| | 0.003856581 |
| | 0.999153539 |
| | 6.90004E-13 |
| | 0.471483691 |
| | 0.007499311 |
| | 0.325467992 |
| | 0.174201535 |
| | 0.999718452 |
| | 0.989028051 |
| | 0.782311375 |
| | 0.565945662 |
| | 0.013021094 |
| | 0.977981907 |
| | 0.131381446 |
| | 4.50267E-08 |
| | 0.241674566 |
| | 0.993156357 |
| | 8.75855E-13 |

| | |
|---|---|
| | 0.768823688 |
| | 2.96073E-07 |
| | 0.12706962 |
| | 0.969065697 |
| | 0.999994735 |
| | 0.717751782 |
| | 9.92021E-06 |
| | 0.999901693 |
| | 0.764750418 |
| | 0.975777935 |
| | 0.025763191 |
| | 0.987990757 |
| | 0.984067206 |
| | 0.070561144 |
| | 0.920595527 |
| | 0.868625777 |
| | 0.001335885 |
| | 0.90686434 |
| | 0.000789874 |
| | 0.999991291 |
| | 4.07082E-10 |
| | 6.04893E-05 |
| | 0.999993269 |
| | 0.025266074 |
| | 0.998724427 |
| | 0.999999999 |
| | 0.880460797 |
| | 6.0416E-06 |
| | 0.000530669 |
| | 5.07156E-05 |
| | 0.18320368 |
| | 0.01334276 |
| | 0.00689299 |
| | 0.086588566 |
| | 0.099380601 |

| | |
|---|---|
| | 0.034338204 |
| | 0.62752221 |
| | 3.56189E-05 |
| | 0.999934531 |
| | 0.984365474 |
| | 0.817093076 |
| | 0.013709214 |
| | 2.08487E-07 |
| | 0.999756955 |
| | 0.001889467 |
| | 0.427594242 |
| | 0.518154644 |
| | 0.006202138 |
| | 0.005528228 |
| | 0.261423746 |
| | 0.992402498 |
| | 0.999963112 |
| | 6.19632E-06 |
| | 1.03464E-08 |
| | 0.001726381 |
| | 0.040609746 |
| | 0.004728126 |
| | 0.225622194 |
| | 1.33107E-05 |
| | 0.200554495 |
| | 0.346742372 |
| | 0.052895714 |
| | 8.19632E-08 |
| | 1.37658E-09 |
| | 1.54869E-05 |
| | 7.77349E-05 |
| | 0.206499679 |
| | 2.73358E-05 |
| | 3.74112E-12 |
| | 0.000271288 |

| | |
|---|---|
| | 9.83853E-08 |
| | 0.999999814 |
| | 1.91291E-09 |
| | 1.60659E-06 |
| | 0.999886817 |
| | 0.999526196 |
| | 0.983708366 |
| | 0.443579617 |
| | 0.999564111 |
| | 0.021692333 |
| | 0.001082534 |
| | 0.303882037 |
| | 0.999986378 |
| | 0.005071336 |
| | 0.999933989 |
| | 0.052921794 |
| | 0.009600308 |
| | 0.994038917 |
| | 0.012219634 |
| | 0.402622421 |
| | 3.8024E-12 |
| | 0.025439351 |
| | 0.759451533 |
| | 0.006266264 |
| | 1.63539E-05 |
| | 0.99596561 |
| | 0.156519549 |
| | 0.18966754 |
| | 0.773625313 |
| | 0.025150262 |
| | 0.995663219 |
| | 0.502428345 |
| | 0.933036933 |
| | 6.27716E-05 |
| | 0.988278012 |

| |
|---:|
| 0.965688214 |
| 0.985725443 |
| 0.14507869 |
| 0.998475266 |
| 0.489257551 |
| 0.999999968 |
| 0.599629076 |
| 0.961413906 |
| 0.01819431 |
| 0.069067457 |
| 0.967275284 |
| 0.995382107 |
| 0.999991902 |
| 0.006574016 |
| 0.008550453 |
| 0.030225244 |
| 0.023540306 |
| 2.22807E-08 |
| 5.09827E-08 |
| 0.118889413 |
| 0.106704731 |
| 0.005401275 |
| 0.959618364 |
| 0.999135061 |
| 0.114591842 |
| 0.755907829 |
| 0.847452406 |
| 0.014689759 |
| 0.994440689 |
| 0.165564545 |
| 1.2064E-07 |
| 6.80611E-10 |
| 3.46694E-08 |
| 4.13102E-05 |
| 0.084539598 |

| | |
|---|---|
| | 0.001770335 |
| | 0.124702316 |
| | 0.999917878 |
| | 0.036256954 |
| | 0.944767257 |
| | 0.038791027 |
| | 8.11533E-05 |
| | 0.9943873 |
| | 0.983573523 |
| | 0.971238217 |
| | 0.997016219 |
| | 4.12117E-05 |
| | 3.88175E-06 |
| | 1.65609E-05 |
| | 0.289624467 |
| | 0.971632743 |
| | 0.008906016 |
| | 0.884374353 |
| | 9.75505E-06 |
| | 1.30673E-13 |
| | 0.85494871 |
| | 1.27528E-09 |
| | 0.002435459 |
| | 0.00017248 |
| | 0.998815803 |
| | 1.12855E-06 |
| | 0.999999992 |
| | 2.88857E-06 |
| | 0.186212706 |
| | 0.253500203 |
| | 0.123310585 |
| | 0.999853028 |
| | 0.105471806 |
| | 1.75984E-06 |
| | 6.54888E-08 |

| |
|---|
| 6.24589E-06 |
| 0.181317001 |
| 0.305062935 |
| 0.274311495 |
| 0.104430002 |
| 0.987138964 |
| 1.28978E-05 |
| 0.013186514 |
| 0.796778952 |
| 0.025771901 |
| 0.999999996 |
| 0.005469365 |
| 5.4512E-14 |
| 1.25111E-07 |
| 1.00203E-06 |
| 4.60743E-14 |
| 4.60743E-14 |
| 6.16174E-14 |
| 0.473711297 |
| 0.9997656 |
| 0.998393621 |
| 0.999929812 |
| 0.004872987 |
| 0.260918936 |
| 6.82626E-07 |
| 7.27196E-14 |
| 1.6727E-10 |
| 0.973087979 |
| 0.736680436 |
| 0.99997877 |
| 0.652330355 |
| 4.82947E-14 |
| 3.7048E-09 |
| 4.60743E-14 |
| 0.000465397 |

| | |
|---|---|
| | 0.99882651 |
| | 0.000272472 |
| | 0.416532675 |
| | 0.645650238 |
| | 2.85521E-05 |
| | 0.999831389 |
| | 0.691464842 |
| | 0.792367993 |
| | 0.810987879 |
| | 0.399223284 |
| | 0.999889685 |
| | 0.999556711 |
| | 0.99999999 |
| | 0.991236986 |
| | 0.943294748 |
| | 3.02796E-05 |
| | 7.78559E-09 |
| | 0.24267812 |
| | 2.3288E-05 |
| | 0.995957275 |
| | 0.35336586 |
| | 0.913191155 |
| | 0.983599876 |
| | 0.908431664 |
| | 0.9655201 |
| | 0.97161148 |
| | 0.389084396 |
| | 0.985021911 |
| | 0.986448329 |
| | 0.998849675 |
| | 0.007328597 |
| | 0.00062727 |
| | 0.999758144 |
| | 0.000748616 |
| | 0.023229999 |

| | |
|---|---|
| | 0.582626352 |
| | 0.944021469 |
| | 0.999999243 |
| | 0.121706918 |
| | 0.013304093 |
| | 0.999996536 |
| | 0.005395651 |
| | 0.766101141 |
| | 2.26519E-12 |
| | 0.000317901 |
| | 0.983698907 |
| | 0.855544348 |
| | 0.704302606 |
| | 0.803201547 |
| | 0.999999997 |
| | 0.001440086 |
| | 0.445547711 |
| | 7.27396E-05 |
| | 0.288788754 |
| | 0.184573836 |
| | 4.28723E-08 |
| | 1.01828E-06 |
| | 1.13079E-07 |
| | 0.984019754 |
| | 0.799762496 |
| | 0.999910662 |
| | 9.72185E-08 |
| | 0.979283395 |
| | 0.996794958 |
| | 2.87228E-06 |
| | 0.963703213 |
| | 0.505009274 |
| | 0.034008677 |
| | 5.09925E-13 |
| | 0.928968763 |

| |
|---|
| 4.60743E-14 |
| 0.997619728 |
| 9.08748E-11 |
| 0.152180177 |
| 0.635543825 |
| 0.502652873 |
| 0.44369533 |
| 0.625197563 |
| 0.972331024 |
| 0.347524188 |
| 0.999982883 |
| 0.667733005 |
| 0.631126572 |
| 0.009679894 |
| 0.34819619 |
| 0.012226774 |
| 0.935649835 |
| 0.001606209 |
| 0.058824469 |
| 0.997732105 |
| 0.063522776 |
| 0.000376236 |
| 0.470102153 |
| 0.992039145 |
| 1.25808E-07 |
| 1.618E-07 |
| 0.290395778 |
| 0.720470975 |
| 4.10984E-05 |
| 0.999676879 |
| 0.651221541 |
| 0.954741784 |
| 0.999276254 |
| 0.999953465 |
| 0.984008866 |

| | |
|---|---|
| | 0.611713031 |
| | 0.999999998 |
| | 0.994219489 |
| | 0.994339915 |
| | 0.999988933 |
| | 0.302450192 |
| | 1.55542E-13 |
| | 0.000834711 |
| | 0.00047366 |
| | 0.132706947 |
| | 0.847900348 |
| | 8.82218E-11 |
| | 0.010825114 |
| | 0.19305682 |
| | 3.08342E-12 |
| | 0.074117199 |
| | 4.59641E-10 |
| | 0.997656487 |
| | 0.002083494 |
| | 2.9135E-09 |
| | 0.99992009 |
| | 0.43901143 |
| | 0.999078506 |
| | 0.998985324 |
| | 0.968240492 |
| | 0.968016788 |
| | 0.196732435 |
| | 0.000293172 |
| | 0.435648272 |
| | 0.999999921 |
| | 7.1896E-06 |
| | 0.918273113 |
| | 0.999807671 |
| | 0.998348433 |
| | 0.999724806 |

| | |
|---|---|
| | 0.932832883 |
| | 3.2362E-08 |
| | 1.97844E-08 |
| | 0.001184974 |
| | 2.88924E-07 |
| | 7.25452E-06 |
| | 2.32455E-05 |
| | 9.77645E-08 |
| | 2.46912E-08 |
| | 0.005176715 |
| | 0.51314465 |
| | 0.989813136 |
| | 0.06429772 |
| | 0.996843245 |
| | 0.907205051 |
| | 0.99999869 |
| | 0.999025886 |
| | 0.103027812 |
| | 0.62209998 |
| | 0.825045718 |
| | 0.760222076 |
| | 0.999310593 |
| | 0.999857215 |
| | 0.999996542 |
| | 0.999982201 |
| | 0.989043732 |
| | 0.999735276 |
| | 0.995210644 |
| | 0.918312011 |
| | 0.934855313 |
| | 1 |
| | 0.99962959 |
| | 0.0096281 |
| | 0.170278694 |
| | 0.015592794 |

| | |
|---|---|
| | 0.999818636 |
| | 0.993996684 |
| | 0.999201862 |
| | 0.999963697 |
| | 0.575349968 |
| | 0.989145892 |
| | 0.601119525 |
| | 3.89142E-07 |
| | 0.655459395 |
| | 5.49255E-07 |
| | 7.56062E-14 |
| | 2.36004E-10 |
| | 2.88961E-08 |
| | 0.575141647 |
| | 0.999999818 |
| | 0.93634668 |
| | 0.987706892 |
| | 0.963298241 |
| | 0.999987726 |
| | 0.999858153 |
| | 0.027019487 |
| | 0.453164461 |
| | 0.007212798 |
| | 0.999482578 |
| | 0.861732565 |
| | 0.689565004 |
| | 1.6287E-13 |
| | 0.040995685 |
| | 0.999999662 |
| | 0.916809271 |
| | 0.932850878 |
| | 0.22027696 |
| | 0.230597758 |
| | 0.983709616 |
| | 0.06110457 |

| |
|---|
| 0.6718452 |
| 0.814088817 |
| 0.008603546 |
| 0.872452035 |
| 0.598366185 |
| 5.50105E-05 |
| 0.004344165 |
| 0.969411432 |
| 0.001605771 |
| 0.771655252 |
| 0.999823199 |
| 0.660516025 |
| 0.417533447 |
| 0.007870032 |
| 0.032790251 |

| |
|---|
| P-value: (Bb_pos_12022024_Newanalysis) / (Rabbitserum_12022024_Newanalysis) |
| 0.065877671 |
| 0.981996314 |
| 3.6749E-09 |
| 8.25374E-05 |
| 0.390766477 |
| 0.819548848 |
| 0.005111945 |
| 6.51486E-06 |
| 0.012759474 |
| 0.392537089 |
| 0.999596808 |
| 0.078165869 |
| 0.003064989 |
| 0.311945226 |
| 0.092522974 |
| 0.001234768 |
| 0.695671648 |
| 4.60743E-14 |
| 9.07714E-07 |
| 1.54732E-12 |
| 0.991914434 |
| 5.05151E-14 |
| 7.25959E-06 |
| 4.60743E-14 |
| 4.60743E-14 |
| 0.998770214 |
| 0.957793863 |
| 0.66611357 |
| 0.006219958 |
| 0.994522885 |
| 0.312977617 |
| 1.21808E-07 |
| 0.027679794 |

| |
|---|
| 0.054004405 |
| 4.65183E-14 |
| 3.83828E-07 |
| 1.63319E-11 |
| 0.001978994 |
| 0.002046197 |
| 1.05961E-10 |
| 0.968584303 |
| 8.36808E-05 |
| 0.000141834 |
| 9.53972E-05 |
| 0.535972259 |
| 1.06026E-13 |
| 1.36583E-11 |
| 5.44081E-11 |
| 3.71761E-05 |
| 3.46942E-06 |
| 1.02108E-05 |
| 1.9347E-07 |
| 8.51874E-13 |
| 2.94949E-07 |
| 0.960640088 |
| 0.386715717 |
| 1.19951E-06 |
| 0.406984781 |
| 0.00211238 |
| 0.429674431 |
| 0.050850271 |
| 0.999375872 |
| 0.053437149 |
| 7.34758E-11 |
| 1.04863E-05 |
| 0.999431119 |
| 7.93829E-09 |
| 0.007785186 |

| |
|---|
| 1.19541E-05 |
| 0.074602221 |
| 0.000290924 |
| 0.999999709 |
| 8.78223E-07 |
| 5.90919E-05 |
| 4.37044E-05 |
| 0.713951784 |
| 4.60743E-14 |
| 0.000194327 |
| 6.31717E-14 |
| 4.62963E-14 |
| 1.20343E-07 |
| 6.37143E-09 |
| 0.343299528 |
| 7.59676E-08 |
| 1.47293E-12 |
| 0.01121857 |
| 0.253669664 |
| 2.08158E-06 |
| 0.034391365 |
| 0.001982084 |
| 7.41403E-07 |
| 9.37139E-13 |
| 4.72313E-10 |
| 6.79444E-09 |
| 2.93656E-09 |
| 5.94175E-06 |
| 1.60937E-11 |
| 2.89081E-08 |
| 0.960922367 |
| 0.268828447 |
| 1.98313E-10 |
| 0.839689867 |
| 5.97357E-06 |

| | |
|---|---|
| | 0.837644143 |
| | 1.48566E-06 |
| | 0.013303322 |
| | 0.018305777 |
| | 0.698238546 |
| | 0.537738045 |
| | 0.744651081 |
| | 0.001437204 |
| | 5.56669E-06 |
| | 0.0214066 |
| | 0.000355967 |
| | 0.795808753 |
| | 2.11125E-05 |
| | 0.86236355 |
| | 4.60743E-14 |
| | 9.39125E-05 |
| | 0.93049689 |
| | 0.999954158 |
| | 4.60743E-14 |
| | 0.05404105 |
| | 0.000116537 |
| | 0.205855729 |
| | 0.613068034 |
| | 0.014905083 |
| | 0.009134146 |
| | 0.0002723 |
| | 0.79422474 |
| | 0.99451725 |
| | 1.61051E-06 |
| | 5.54001E-14 |
| | 1.4246E-08 |
| | 0.269487135 |
| | 0.017000947 |
| | 1.35668E-05 |
| | 0.264272463 |

| | |
|---|---|
| | 0.983765899 |
| | 0.312702508 |
| | 0.018287936 |
| | 0.935705905 |
| | 0.788661131 |
| | 0.598316634 |
| | 0.71144142 |
| | 0.991265333 |
| | 7.35302E-06 |
| | 0.511148622 |
| | 0.415752668 |
| | 1.77895E-06 |
| | 4.60743E-14 |
| | 0.026628649 |
| | 1.55098E-13 |
| | 4.60743E-14 |
| | 0.998131139 |
| | 0.000159245 |
| | 1.86979E-05 |
| | 0.003137654 |
| | 8.01636E-06 |
| | 6.05254E-09 |
| | 5.4956E-14 |
| | 2.61503E-10 |
| | 0.132693425 |
| | 8.03789E-06 |
| | 0.957515767 |
| | 0.334790977 |
| | 0.000982072 |
| | 0.225451653 |
| | 1.34052E-07 |
| | 0.003645925 |
| | 0.298945252 |
| | 0.001501693 |
| | 1.56195E-08 |

| | |
|---|---|
| | 0.9985602 |
| | 7.66911E-07 |
| | 6.77377E-07 |
| | 4.60743E-14 |
| | 7.93506E-09 |
| | 4.93142E-07 |
| | 0.000655213 |
| | 9.05888E-09 |
| | 3.09977E-05 |
| | 2.81069E-08 |
| | 0.131086051 |
| | 0.99999922 |
| | 4.60743E-14 |
| | 2.3576E-07 |
| | 0.000662184 |
| | 6.30266E-09 |
| | 3.28105E-07 |
| | 2.78437E-05 |
| | 4.60743E-14 |
| | 8.32612E-12 |
| | 5.14893E-08 |
| | 1.8613E-10 |
| | 0.989457943 |
| | 0.999999709 |
| | 0.000424621 |
| | 0.243361755 |
| | 0.007427755 |
| | 0.364105669 |
| | 6.67405E-07 |
| | 1.38814E-09 |
| | 4.17348E-08 |
| | 9.04214E-06 |
| | 2.14865E-08 |
| | 7.17496E-07 |
| | 8.22197E-06 |

| | |
|---|---|
| | 1.34959E-07 |
| | 0.000729788 |
| | 0.001629137 |
| | 0.778130494 |
| | 0.030444447 |
| | 0.431376765 |
| | 0.000328695 |
| | 0.00171335 |
| | 0.350880135 |
| | 0.93017753 |
| | 0.000295109 |
| | 0.541224822 |
| | 0.001066147 |
| | 0.999999998 |
| | 0.051589495 |
| | 0.000564065 |
| | 0.001877474 |
| | 2.15383E-10 |
| | 5.91764E-09 |
| | 0.99999981 |
| | 6.04191E-09 |
| | 0.087356798 |
| | 0.996346284 |
| | 5.49645E-07 |
| | 4.71845E-14 |
| | 4.19884E-07 |
| | 0.999640794 |
| | 0.480159677 |
| | 0.007841664 |
| | 0.979086413 |
| | 0.229474672 |
| | 0.953593386 |
| | 0.058130734 |
| | 0.114027946 |
| | 5.9841E-14 |

| |
|---|
| 0.996412188 |
| 0.047032509 |
| 1 |
| 0.968808886 |
| 0.08854588 |
| 1.34018E-10 |
| 1.20008E-07 |
| 0.899370142 |
| 0.002545663 |
| 4.65183E-14 |
| 0.98538981 |
| 0.301425895 |
| 0.095179033 |
| 9.28924E-13 |
| 0.20089004 |
| 0.313540784 |
| 0.999995586 |
| 0.954869935 |
| 0.999533383 |
| 4.60743E-14 |
| 1.23159E-06 |
| 0.316590111 |
| 0.550539374 |
| 0.736248225 |
| 0.603519754 |
| 1 |
| 0.011513511 |
| 0.99079808 |
| 0.080372866 |
| 0.008449061 |
| 9.99777E-07 |
| 2.32248E-12 |
| 0.004480363 |
| 0.923478963 |
| 4.60743E-14 |

| | |
|---|---|
| | 8.75741E-11 |
| | 4.02678E-13 |
| | 9.40609E-08 |
| | 0.121296597 |
| | 0.001375374 |
| | 0.02828652 |
| | 5.73985E-14 |
| | 0.799545258 |
| | 0.181763806 |
| | 4.55744E-09 |
| | 0.969664114 |
| | 4.17744E-05 |
| | 0.990643298 |
| | 1.15688E-06 |
| | 0.965077304 |
| | 0.698775301 |
| | 6.03884E-12 |
| | 0.000587734 |
| | 2.47625E-05 |
| | 0.001596835 |
| | 8.26933E-11 |
| | 0.001293065 |
| | 0.913192391 |
| | 4.76286E-14 |
| | 2.63339E-05 |
| | 9.43996E-08 |
| | 0.001660835 |
| | 5.25135E-14 |
| | 0.671376878 |
| | 3.72468E-11 |
| | 8.58219E-05 |
| | 1.26342E-06 |
| | 0.630469944 |
| | 0.001252932 |
| | 0.001880991 |

| |
|---|
| 1.51335E-12 |
| 0.988649787 |
| 5.12456E-09 |
| 0.008489569 |
| 0.002813478 |
| 0.999953111 |
| 1.75634E-07 |
| 4.62963E-14 |
| 0.877111375 |
| 3.02991E-12 |
| 0.020322715 |
| 0.000198974 |
| 0.000110703 |
| 0.00040381 |
| 0.997614521 |
| 0.980963919 |
| 1.10699E-05 |
| 1.28674E-10 |
| 5.00711E-14 |
| 1.81388E-05 |
| 0.006479179 |
| 0.979882942 |
| 0.002671642 |
| 1.1503E-12 |
| 8.7582E-07 |
| 0.137196161 |
| 0.00035214 |
| 0.288315289 |
| 0.000752567 |
| 0.103102048 |
| 0.708652714 |
| 3.15545E-05 |
| 0.730337499 |
| 1.07542E-08 |
| 0.012830296 |

| | |
|---|---|
| | 0.00049612 |
| | 0.489122493 |
| | 1.00486E-12 |
| | 2.00883E-07 |
| | 0.000222942 |
| | 0.357135333 |
| | 5.7216E-06 |
| | 5.90206E-12 |
| | 2.92944E-05 |
| | 7.71915E-09 |
| | 0.251358203 |
| | 2.318E-11 |
| | 2.17904E-07 |
| | 0.001635096 |
| | 0.936147625 |
| | 0.000665147 |
| | 0.999977749 |
| | 1.66458E-05 |
| | 0.000215406 |
| | 0.000158719 |
| | 4.60743E-14 |
| | 3.02723E-11 |
| | 0.871327968 |
| | 2.88617E-10 |
| | 2.82901E-11 |
| | 0.949501453 |
| | 0.001709987 |
| | 5.21672E-06 |
| | 0.000124801 |
| | 2.08474E-08 |
| | 0.344677552 |
| | 0.000122925 |
| | 9.17726E-07 |
| | 5.65753E-06 |
| | 0.699805466 |

| | |
|---|---|
| | 0.244091268 |
| | 0.15921725 |
| | 5.72213E-05 |
| | 1.32366E-08 |
| | 0.021788613 |
| | 4.14859E-08 |
| | 6.83596E-09 |
| | 2.27195E-07 |
| | 2.5676E-09 |
| | 0.999219842 |
| | 0.000138573 |
| | 1.41703E-11 |
| | 1.12022E-07 |
| | 1.43751E-06 |
| | 8.22433E-08 |
| | 3.1156E-05 |
| | 2.30888E-05 |
| | 2.24631E-12 |
| | 1.23346E-13 |
| | 0.00130325 |
| | 0.000826732 |
| | 5.51591E-07 |
| | 0.000110892 |
| | 6.56311E-10 |
| | 0.767476598 |
| | 0.996133696 |
| | 0.518532024 |
| | 0.00089353 |
| | 0.917008985 |
| | 0.004141089 |
| | 3.90465E-13 |
| | 4.96159E-13 |
| | 4.65183E-14 |
| | 1.24359E-08 |
| | 0.000489545 |

| |
|---|
| 1.18597E-08 |
| 0.001007104 |
| 0.00120828 |
| 0.000324118 |
| 0.717553499 |
| 1 |
| 0.02860374 |
| 0.002872553 |
| 1.71313E-06 |
| 9.5449E-06 |
| 0.023630671 |
| 4.10283E-12 |
| 7.9977E-11 |
| 3.53162E-06 |
| 1.67858E-06 |
| 5.54066E-05 |
| 0.99949746 |
| 0.382574392 |
| 4.33908E-12 |
| 4.60743E-14 |
| 4.84699E-05 |
| 6.36158E-14 |
| 1.64026E-07 |
| 7.54951E-07 |
| 0.12538788 |
| 5.4016E-11 |
| 1.93775E-08 |
| 9.97091E-13 |
| 0.005383558 |
| 0.736671747 |
| 0.006221774 |
| 0.86586387 |
| 9.21936E-07 |
| 3.52031E-11 |
| 2.37699E-13 |

| | |
|---|---|
| | 3.42985E-11 |
| | 1.16422E-06 |
| | 7.87245E-07 |
| | 0.087332211 |
| | 0.798445906 |
| | 0.693338904 |
| | 7.45519E-10 |
| | 7.01433E-09 |
| | 0.127516722 |
| | 0.001747308 |
| | 0.004338289 |
| | 0.70408137 |
| | 1.72881E-05 |
| | 8.96477E-05 |
| | 0.00932422 |
| | 4.60743E-14 |
| | 8.37885E-13 |
| | 1.8463E-13 |
| | 0.202979362 |
| | 8.84224E-09 |
| | 0.733857253 |
| | 0.01577027 |
| | 0.006538681 |
| | 7.43092E-05 |
| | 0.836725187 |
| | 4.33406E-08 |
| | 4.72901E-06 |
| | 1.06248E-13 |
| | 0.563351919 |
| | 0.00047602 |
| | 6.07463E-05 |
| | 4.18627E-08 |
| | 0.035508676 |
| | 4.60743E-14 |
| | 7.03086E-05 |

| |
|---|
| 0.955355 |
| 0.000429046 |
| 0.073028154 |
| 0.039516505 |
| 0.999799203 |
| 6.11733E-14 |
| 0.001028331 |
| 0.972437488 |
| 0.012514406 |
| 0.477639404 |
| 0.037274318 |
| 0.000323515 |
| 0.005244789 |
| 0.020195709 |
| 0.000134769 |
| 1.86778E-10 |
| 0.895406912 |
| 4.01585E-07 |
| 0.015521945 |
| 4.85004E-06 |
| 2.78488E-07 |
| 0.175293135 |
| 0.341730542 |
| 0.999998679 |
| 0.001972919 |
| 7.76146E-12 |
| 0.938126239 |
| 3.35166E-07 |
| 3.01801E-07 |
| 0.239734636 |
| 1.25929E-07 |
| 3.6453E-08 |
| 2.8806E-10 |
| 0.001071506 |
| 2.76412E-12 |

| |
|---|
| 0.003264282 |
| 0.064025507 |
| 1.36515E-08 |
| 2.90846E-06 |
| 5.09592E-14 |
| 0.011026978 |
| 7.2403E-10 |
| 0.315954809 |
| 4.60743E-14 |
| 6.40599E-14 |
| 0.000808603 |
| 3.61266E-06 |
| 0.032072909 |
| 0.012862313 |
| 6.72886E-08 |
| 4.55002E-06 |
| 3.35078E-06 |
| 2.50133E-13 |
| 9.12923E-10 |
| 0.135292574 |
| 2.3348E-13 |
| 4.60743E-14 |
| 5.73985E-14 |
| 0.00603586 |
| 2.10685E-09 |
| 0.006636175 |
| 5.9619E-14 |
| 0.085229761 |
| 3.76024E-07 |
| 2.5796E-12 |
| 0.012382986 |
| 0.000476542 |
| 0.000105435 |
| 1.86406E-13 |
| 0.467126088 |

| | |
|---|---|
| | 0.001156998 |
| | 0.952174563 |
| | 0.671064153 |
| | 0.021551117 |
| | 0.024343232 |
| | 0.135243319 |
| | 0.50684359 |
| | 0.017464116 |
| | 3.24399E-07 |
| | 0.99928215 |
| | 0.714828119 |
| | 0.836001718 |
| | 4.08496E-07 |
| | 4.57657E-11 |
| | 5.31258E-05 |
| | 8.83736E-05 |
| | 0.946259989 |
| | 0.012969646 |
| | 0.568125977 |
| | 0.403409119 |
| | 3.53808E-06 |
| | 6.76694E-06 |
| | 0.766686076 |
| | 0.000142558 |
| | 6.67438E-05 |
| | 5.5225E-07 |
| | 0.000171631 |
| | 0.002492351 |
| | 7.85629E-08 |
| | 0.929333997 |
| | 0.751934018 |
| | 4.80795E-05 |
| | 0.036372962 |
| | 0.981016456 |
| | 0.999995203 |

| |
|---|
| 6.20479E-05 |
| 0.779988549 |
| 0.634223884 |
| 0.488485585 |
| 0.671911363 |
| 0.066750658 |
| 4.60743E-14 |
| 5.99448E-11 |
| 4.76404E-09 |
| 0.324991673 |
| 0.001993074 |
| 7.01257E-06 |
| 3.69149E-11 |
| 0.005668919 |
| 4.60743E-14 |
| 3.96755E-06 |
| 6.75507E-05 |
| 0.999821911 |
| 0.011217411 |
| 0.321162755 |
| 0.936527083 |
| 7.84246E-07 |
| 0.628247226 |
| 0.857223871 |
| 0.013621112 |
| 0.618828763 |
| 0.016406228 |
| 1.29619E-12 |
| 0.209491352 |
| 0.8572666 |
| 1.49991E-13 |
| 0.000132531 |
| 2.92663E-05 |
| 0.085291221 |
| 1.35304E-05 |

| |
|---|
| 0.437664273 |
| 3.91047E-09 |
| 1.91812E-11 |
| 1.04658E-06 |
| 6.5338E-09 |
| 1.14865E-07 |
| 8.85466E-08 |
| 4.85167E-14 |
| 8.9548E-11 |
| 1.60988E-05 |
| 7.95566E-07 |
| 0.200807613 |
| 0.007867298 |
| 0.980529852 |
| 0.017053644 |
| 0.713771452 |
| 0.781529383 |
| 0.000206749 |
| 4.49017E-06 |
| 0.000104174 |
| 0.856508261 |
| 0.900201537 |
| 0.914312862 |
| 0.437915399 |
| 0.898824625 |
| 0.525985381 |
| 0.912661484 |
| 0.703298172 |
| 0.806598382 |
| 0.999124853 |
| 0.62918124 |
| 0.981023461 |
| 0.000210243 |
| 1.72377E-11 |
| 0.000200326 |

| | |
|---|---|
| | 0.095615898 |
| | 0.918996231 |
| | 0.999985598 |
| | 3.21102E-05 |
| | 1.1033E-05 |
| | 7.64355E-09 |
| | 0.016343817 |
| | 2.75399E-11 |
| | 0.269693742 |
| | 2.56934E-10 |
| | 4.60743E-14 |
| | 5.91749E-14 |
| | 4.15334E-13 |
| | 8.14993E-07 |
| | 9.92298E-05 |
| | 0.046188868 |
| | 9.01102E-06 |
| | 0.953281725 |
| | 0.99982961 |
| | 0.997414272 |
| | 0.003464245 |
| | 0.712599613 |
| | 7.99437E-07 |
| | 0.580393729 |
| | 0.302717598 |
| | 1.06542E-06 |
| | 4.56683E-06 |
| | 0.006442156 |
| | 0.001104335 |
| | 0.79124925 |
| | 0.001220565 |
| | 0.0001124 |
| | 0.129412998 |
| | 0.882561072 |
| | 7.62601E-07 |

| |
|---|
| 0.878361204 |
| 0.20251868 |
| 0.867105584 |
| 0.147663125 |
| 0.012299458 |
| 7.27632E-11 |
| 3.11802E-07 |
| 0.815232639 |
| 0.161183101 |
| 0.961561321 |
| 0.997463762 |
| 0.51708754 |
| 0.709891453 |
| 0.001776495 |
| 5.9773E-07 |

| |
|---|
| P-value: (CM_pos_12022024_Newanalysis) / (Rabbitserum_12022024_Newanalysis) |
| 0.479404778 |
| 0.965056787 |
| 2.86548E-09 |
| 0.515686566 |
| 0.090392472 |
| 0.979820698 |
| 0.084934827 |
| 0.194269777 |
| 0.001189448 |
| 0.140046528 |
| 0.935132943 |
| 0.55033034 |
| 0.243627994 |
| 0.997718755 |
| 0.125251095 |
| 0.365979984 |
| 0.999993161 |
| 1.40815E-06 |
| 0.000124669 |
| 6.30141E-08 |
| 0.907529175 |
| 0.000582307 |
| 0.967545932 |
| 1.55098E-13 |
| 0.000108273 |
| 0.686133826 |
| 0.999972933 |
| 0.967121345 |
| 0.67163531 |
| 0.908086329 |
| 0.570778769 |
| 0.99331741 |
| 0.721243749 |

| |
|---|
| 0.808181952 |
| 0.008797338 |
| 4.57242E-10 |
| 2.59222E-05 |
| 0.998550106 |
| 0.983862013 |
| 0.3911877 |
| 0.999163211 |
| 1.14335E-06 |
| 0.932656996 |
| 1.71759E-08 |
| 0.858949404 |
| 0.007658036 |
| 0.050950416 |
| 0.085960856 |
| 0.666621602 |
| 0.640915796 |
| 0.87276624 |
| 0.875380215 |
| 0.158780359 |
| 0.15917358 |
| 0.373693056 |
| 0.998219885 |
| 9.15942E-11 |
| 0.998883341 |
| 0.027542961 |
| 0.000292088 |
| 0.466674654 |
| 0.993048888 |
| 0.984539197 |
| 0.291241692 |
| 0.623260583 |
| 0.976289661 |
| 0.038490478 |
| 0.294665576 |

| | |
|---|---|
| | 0.00033312 |
| | 0.627077609 |
| | 0.216577733 |
| | 0.99770644 |
| | 0.253246754 |
| | 0.152905894 |
| | 0.000140207 |
| | 0.999769876 |
| | 0.00141397 |
| | 3.08032E-05 |
| | 0.003022553 |
| | 2.4995E-09 |
| | 0.24453992 |
| | 0.000740439 |
| | 0.440522319 |
| | 0.191258471 |
| | 5.43942E-10 |
| | 0.99902888 |
| | 0.666771027 |
| | 0.907667975 |
| | 0.999932055 |
| | 0.59020104 |
| | 0.87927569 |
| | 0.005596044 |
| | 0.006676689 |
| | 0.069634065 |
| | 0.004903077 |
| | 0.053078784 |
| | 0.231913657 |
| | 0.041014312 |
| | 0.985785397 |
| | 0.999999995 |
| | 6.42491E-07 |
| | 0.070506224 |
| | 0.652200207 |

| | |
|---|---|
| | 0.530058412 |
| | 0.074575626 |
| | 0.093783537 |
| | 0.955720739 |
| | 0.906924091 |
| | 0.115811927 |
| | 0.723596396 |
| | 0.657561741 |
| | 0.000308368 |
| | 0.232458482 |
| | 0.503823806 |
| | 0.977517855 |
| | 0.00842769 |
| | 0.959609232 |
| | 0.00101035 |
| | 0.98141854 |
| | 0.461476651 |
| | 0.992691221 |
| | 4.60743E-14 |
| | 0.027729228 |
| | 0.061846087 |
| | 0.485023083 |
| | 0.835683378 |
| | 0.931697854 |
| | 0.07728499 |
| | 0.000486957 |
| | 0.913822125 |
| | 0.776800904 |
| | 0.001266573 |
| | 3.22108E-06 |
| | 0.000289721 |
| | 0.49217144 |
| | 0.996186736 |
| | 0.421236897 |
| | 0.635872443 |

| | |
|---|---|
| | 0.997063081 |
| | 0.018015686 |
| | 0.555152321 |
| | 0.999999993 |
| | 0.964872237 |
| | 0.704397893 |
| | 0.999673418 |
| | 0.999917834 |
| | 0.371646915 |
| | 0.464666168 |
| | 0.999999991 |
| | 3.65632E-07 |
| | 2.72338E-13 |
| | 0.000495352 |
| | 3.44091E-12 |
| | 8.9738E-10 |
| | 0.995908675 |
| | 0.008761149 |
| | 0.046143723 |
| | 3.47593E-05 |
| | 0.001180938 |
| | 7.46721E-07 |
| | 4.1267E-13 |
| | 0.001569389 |
| | 0.139812021 |
| | 0.6189929 |
| | 0.03426357 |
| | 0.998559894 |
| | 0.050131419 |
| | 0.041699029 |
| | 9.6101E-05 |
| | 0.61787628 |
| | 0.001331828 |
| | 0.038286111 |
| | 3.05543E-05 |

| | |
|---|---|
| | 0.01872685 |
| | 2.67928E-08 |
| | 0.089255306 |
| | 5.4512E-14 |
| | 0.00567648 |
| | 5.1901E-05 |
| | 0.493170518 |
| | 8.32395E-08 |
| | 0.132789353 |
| | 5.94471E-07 |
| | 0.000739847 |
| | 0.861485299 |
| | 1.93512E-13 |
| | 0.067124808 |
| | 0.168540094 |
| | 1.09011E-06 |
| | 1.07127E-07 |
| | 0.025088586 |
| | 1.05419E-11 |
| | 9.25058E-08 |
| | 0.000517893 |
| | 2.55522E-06 |
| | 0.350999474 |
| | 0.874917663 |
| | 0.504805986 |
| | 0.079737078 |
| | 0.829200089 |
| | 1 |
| | 0.117311853 |
| | 0.002348401 |
| | 0.117610079 |
| | 0.262044633 |
| | 0.000210243 |
| | 0.00050094 |
| | 0.008307622 |

| | |
|---|---|
| | 0.656210995 |
| | 0.002006815 |
| | 0.071587474 |
| | 0.999612672 |
| | 0.927040438 |
| | 0.535411881 |
| | 0.031895026 |
| | 0.000919821 |
| | 0.187655969 |
| | 0.206533018 |
| | 0.096826853 |
| | 0.999992033 |
| | 0.384401458 |
| | 0.995815066 |
| | 0.999716857 |
| | 0.999998442 |
| | 0.880096378 |
| | 1.96176E-13 |
| | 0.036481341 |
| | 0.979545719 |
| | 0.000357019 |
| | 0.007933915 |
| | 0.842578865 |
| | 2.49347E-05 |
| | 5.21496E-11 |
| | 0.000847083 |
| | 0.999402654 |
| | 0.838429048 |
| | 0.971011523 |
| | 0.979396634 |
| | 0.010500874 |
| | 0.999998258 |
| | 0.000757658 |
| | 0.047997472 |
| | 5.14215E-08 |

| | |
|---|---|
| | 0.045176238 |
| | 0.079769877 |
| | 0.99999462 |
| | 0.999199873 |
| | 0.175364283 |
| | 1.34847E-06 |
| | 0.000153172 |
| | 0.878455597 |
| | 0.156010264 |
| | 1.06038E-09 |
| | 0.002014503 |
| | 0.019358196 |
| | 0.005255737 |
| | 7.89221E-09 |
| | 0.027949713 |
| | 1.69553E-05 |
| | 0.001033387 |
| | 0.032990476 |
| | 0.983588259 |
| | 4.2404E-05 |
| | 0.000181934 |
| | 2.4544E-05 |
| | 0.008623121 |
| | 0.007364782 |
| | 0.7734068 |
| | 0.98981135 |
| | 0.206182503 |
| | 0.243968558 |
| | 5.94094E-06 |
| | 0.001251511 |
| | 0.001234823 |
| | 0.001800962 |
| | 0.507456705 |
| | 0.99824027 |
| | 4.58356E-12 |

| |
|---|
| 1.8349E-09 |
| 1.72313E-05 |
| 0.000108789 |
| 0.021105631 |
| 0.000971894 |
| 0.438660549 |
| 3.98743E-09 |
| 0.668051333 |
| 0.88289873 |
| 2.81745E-08 |
| 0.141793223 |
| 7.36864E-06 |
| 0.999999639 |
| 8.06702E-10 |
| 0.999971404 |
| 0.999458504 |
| 2.05242E-07 |
| 3.45868E-05 |
| 0.808676954 |
| 0.001088456 |
| 0.979566581 |
| 5.67862E-10 |
| 0.950921249 |
| 6.30052E-13 |
| 8.99549E-06 |
| 8.85994E-08 |
| 0.027673552 |
| 2.77467E-09 |
| 0.026521627 |
| 5.3451E-05 |
| 0.049283275 |
| 0.021166816 |
| 0.228456128 |
| 0.542298263 |
| 0.595064204 |

| |
|---|
| 1.52999E-09 |
| 0.274652968 |
| 0.023497544 |
| 0.004897014 |
| 0.015766169 |
| 0.708191301 |
| 0.003053862 |
| 1.94485E-10 |
| 0.739683509 |
| 6.07127E-08 |
| 0.646543614 |
| 0.020394572 |
| 0.67935788 |
| 0.924302391 |
| 0.499003328 |
| 0.999994564 |
| 1.86355E-05 |
| 0.002057138 |
| 6.19132E-07 |
| 2.67686E-10 |
| 0.974735158 |
| 0.000707618 |
| 0.415032639 |
| 1.83616E-06 |
| 0.00058611 |
| 0.99445353 |
| 0.400308323 |
| 2.67948E-05 |
| 0.000299847 |
| 0.022982698 |
| 0.003908094 |
| 0.017919899 |
| 6.12657E-07 |
| 0.014955242 |
| 0.702889777 |

| | |
|---|---|
| | 0.04028375 |
| | 0.529212975 |
| | 0.016186662 |
| | 0.968531033 |
| | 0.000115562 |
| | 0.212504991 |
| | 9.2495E-07 |
| | 3.33844E-10 |
| | 6.95826E-05 |
| | 6.2134E-05 |
| | 2.26644E-06 |
| | 2.84995E-09 |
| | 3.26965E-07 |
| | 0.998061634 |
| | 0.860708665 |
| | 0.538038796 |
| | 0.014761643 |
| | 3.78869E-06 |
| | 0.664246811 |
| | 7.86816E-07 |
| | 4.82754E-09 |
| | 7.87875E-08 |
| | 0.999906945 |
| | 5.22396E-06 |
| | 0.000116587 |
| | 0.998862811 |
| | 0.434299724 |
| | 0.00378708 |
| | 0.004535692 |
| | 0.000159428 |
| | 0.644214528 |
| | 0.014143411 |
| | 1.18105E-05 |
| | 0.950547481 |
| | 0.32923254 |

| | |
|---|---|
| | 0.048836497 |
| | 0.458432405 |
| | 6.17009E-08 |
| | 4.90542E-09 |
| | 0.599793166 |
| | 3.69376E-08 |
| | 4.13953E-07 |
| | 2.02495E-06 |
| | 2.15521E-05 |
| | 0.143071326 |
| | 0.001219598 |
| | 4.94349E-12 |
| | 7.82405E-08 |
| | 0.045695798 |
| | 0.002296746 |
| | 0.136352807 |
| | 0.135291787 |
| | 0.003618753 |
| | 1.15777E-05 |
| | 0.457610343 |
| | 0.391945827 |
| | 0.023484333 |
| | 0.001094099 |
| | 2.87432E-10 |
| | 0.767061894 |
| | 0.953570238 |
| | 0.067347687 |
| | 0.892509309 |
| | 0.99713583 |
| | 0.619335082 |
| | 4.24405E-05 |
| | 0.018164804 |
| | 4.62109E-09 |
| | 0.048397592 |
| | 0.348960781 |

| | |
|---|---|
| | 0.001452865 |
| | 0.389365747 |
| | 0.002201646 |
| | 0.478932335 |
| | 0.221314388 |
| | 0.039782082 |
| | 0.26656924 |
| | 0.000679087 |
| | 2.84128E-07 |
| | 1.1874E-06 |
| | 0.007366185 |
| | 3.57326E-06 |
| | 0.001830772 |
| | 0.992713985 |
| | 6.51495E-09 |
| | 0.00045984 |
| | 0.003859686 |
| | 0.048820053 |
| | 1.61392E-05 |
| | 6.12215E-07 |
| | 0.00117375 |
| | 7.40105E-05 |
| | 0.015862835 |
| | 0.377386593 |
| | 0.055527163 |
| | 0.003843109 |
| | 1.77536E-08 |
| | 6.82277E-06 |
| | 0.64227616 |
| | 0.955462591 |
| | 4.74118E-06 |
| | 0.739624113 |
| | 0.001527827 |
| | 0.001456114 |
| | 3.46183E-05 |

| | |
|---|---|
| | 0.000393825 |
| | 0.000911903 |
| | 3.5532E-09 |
| | 0.990005986 |
| | 0.707872193 |
| | 0.317311895 |
| | 0.007767438 |
| | 9.7493E-05 |
| | 0.006449347 |
| | 1.99505E-07 |
| | 0.004693267 |
| | 0.155615852 |
| | 1.63863E-09 |
| | 0.179711755 |
| | 0.024201353 |
| | 0.025490094 |
| | 7.22453E-07 |
| | 0.825496305 |
| | 0.993598169 |
| | 1.74744E-08 |
| | 0.483407623 |
| | 0.009169804 |
| | 1.30762E-07 |
| | 1.8169E-07 |
| | 1.75429E-05 |
| | 2.6969E-06 |
| | 0.003661308 |
| | 2.86327E-13 |
| | 0.999724972 |
| | 0.000298635 |
| | 9.339E-07 |
| | 5.5111E-08 |
| | 1.52294E-05 |
| | 0.994682538 |
| | 0.982662201 |

| | |
|---|---|
| | 0.996940781 |
| | 0.999981464 |
| | 0.926252437 |
| | 0.598839984 |
| | 5.97366E-05 |
| | 5.58442E-14 |
| | 0.043276106 |
| | 0.99477687 |
| | 0.198138348 |
| | 0.009070886 |
| | 0.063661153 |
| | 0.000765602 |
| | 0.00476963 |
| | 0.078184015 |
| | 0.001607053 |
| | 0.000675963 |
| | 1.11315E-07 |
| | 1.39365E-09 |
| | 0.188012196 |
| | 1.89753E-05 |
| | 6.86936E-05 |
| | 0.019375702 |
| | 0.100850894 |
| | 0.938031199 |
| | 0.015784719 |
| | 3.89418E-11 |
| | 0.071925114 |
| | 6.03047E-08 |
| | 1.68796E-06 |
| | 0.431015593 |
| | 0.004177168 |
| | 0.01280474 |
| | 5.43135E-10 |
| | 0.999993946 |
| | 4.77138E-09 |

| |
|---|
| 0.15604422 |
| 0.007303755 |
| 1.69317E-08 |
| 0.003955557 |
| 3.72424E-12 |
| 0.014809405 |
| 1.83545E-05 |
| 0.020104587 |
| 1.68139E-08 |
| 3.87989E-10 |
| 0.004929651 |
| 8.67893E-05 |
| 0.000767585 |
| 0.207715328 |
| 6.24451E-08 |
| 0.321360144 |
| 0.000570291 |
| 3.60236E-08 |
| 1.90215E-07 |
| 0.999982334 |
| 5.30645E-05 |
| 3.09675E-12 |
| 2.72227E-07 |
| 0.00100753 |
| 4.81736E-08 |
| 0.003671751 |
| 4.28072E-07 |
| 0.016437463 |
| 1.32495E-06 |
| 2.69813E-05 |
| 0.00147857 |
| 0.045367111 |
| 1.97194E-08 |
| 0.987066918 |
| 0.950815608 |

| |
|---|
| 5.9841E-14 |
| 0.778861286 |
| 2.85118E-09 |
| 0.947136892 |
| 0.482203986 |
| 0.963471574 |
| 0.999998018 |
| 0.411918561 |
| 2.47281E-06 |
| 0.549000318 |
| 0.616189782 |
| 0.999612883 |
| 2.82242E-05 |
| 3.57767E-07 |
| 0.013015348 |
| 0.465559818 |
| 0.999999983 |
| 0.965959361 |
| 0.776967179 |
| 0.670971593 |
| 0.010595996 |
| 0.696497966 |
| 0.039230341 |
| 0.000696302 |
| 0.218693001 |
| 0.996975778 |
| 0.047525589 |
| 0.080408028 |
| 0.215769474 |
| 0.985128734 |
| 0.999978301 |
| 0.000510988 |
| 0.015770591 |
| 0.995870069 |
| 0.966291405 |

| |
|---|
| 0.004790232 |
| 0.794270391 |
| 0.908363045 |
| 0.217523791 |
| 0.758787869 |
| 0.000280419 |
| 2.66718E-09 |
| 5.97451E-06 |
| 0.001961908 |
| 0.995594267 |
| 0.03884235 |
| 0.001149376 |
| 2.45454E-07 |
| 8.15974E-06 |
| 1.12613E-09 |
| 0.009854963 |
| 0.000904018 |
| 0.999968992 |
| 0.986691399 |
| 5.79141E-07 |
| 0.979800152 |
| 0.000132707 |
| 0.406726014 |
| 0.655294896 |
| 0.084927369 |
| 0.966594177 |
| 0.863548082 |
| 1.1342E-07 |
| 0.002627209 |
| 0.832063405 |
| 1.19849E-07 |
| 0.001999934 |
| 6.08402E-05 |
| 0.192494587 |
| 2.96533E-05 |

| | |
|---|---|
| | 0.934713095 |
| | 0.95446834 |
| | 0.054942065 |
| | 0.1399867 |
| | 0.672591461 |
| | 0.636047021 |
| | 0.327864851 |
| | 2.86948E-08 |
| | 0.196415157 |
| | 0.311690967 |
| | 9.57022E-05 |
| | 0.505575732 |
| | 0.95054694 |
| | 0.999903062 |
| | 0.164382148 |
| | 0.65550775 |
| | 0.561082489 |
| | 0.175649351 |
| | 0.000347752 |
| | 0.002932943 |
| | 0.999960212 |
| | 0.980716381 |
| | 0.973400716 |
| | 0.371720717 |
| | 0.948012775 |
| | 0.87298473 |
| | 0.977477213 |
| | 0.936725366 |
| | 0.999796823 |
| | 0.992109721 |
| | 0.629222707 |
| | 0.9152115 |
| | 0.715514136 |
| | 4.54922E-09 |
| | 0.587500582 |

| | |
|---|---|
| | 0.160996628 |
| | 0.997552867 |
| | 0.995425981 |
| | 5.41186E-05 |
| | 0.001050681 |
| | 1.73329E-09 |
| | 0.418664001 |
| | 0.004846872 |
| | 0.983262023 |
| | 0.043260225 |
| | 1.44651E-05 |
| | 0.000363212 |
| | 0.000222234 |
| | 7.43986E-05 |
| | 0.000118844 |
| | 0.290529986 |
| | 1.61167E-06 |
| | 0.561250344 |
| | 0.998388534 |
| | 0.98065155 |
| | 0.963739958 |
| | 0.02937643 |
| | 0.025125256 |
| | 0.773889059 |
| | 0.029998822 |
| | 5.87689E-05 |
| | 2.20378E-07 |
| | 0.973796292 |
| | 0.000903491 |
| | 0.999685086 |
| | 0.014407354 |
| | 0.048735002 |
| | 0.999580464 |
| | 0.998257182 |
| | 0.002508187 |

| |
|---:|
| 0.131108841 |
| 0.868873208 |
| 0.118471429 |
| 0.716875359 |
| 0.358156375 |
| 0.000115615 |
| 0.016976547 |
| 0.997145151 |
| 0.412009367 |
| 0.99576665 |
| 0.979465791 |
| 0.999901013 |
| 0.996576803 |
| 0.992708084 |
| 0.004051719 |

| |
|---|
| P-value: (SM_pos_12022024_Newanalyis) / (Rabbitserum_12022024_Newanalysis) |
| 0.967056208 |
| 0.019299354 |
| 2.51051E-08 |
| 0.804859534 |
| 5.32024E-06 |
| 0.993204626 |
| 0.309653397 |
| 0.000229072 |
| 0.358241223 |
| 0.002438739 |
| 0.001840469 |
| 0.942760159 |
| 0.99016905 |
| 0.160198455 |
| 0.017327786 |
| 0.980209689 |
| 0.495638453 |
| 4.60743E-14 |
| 1.37134E-09 |
| 2.91434E-13 |
| 0.216411128 |
| 4.51117E-12 |
| 0.049266449 |
| 4.60743E-14 |
| 4.69624E-14 |
| 0.209127162 |
| 0.63320257 |
| 0.331715684 |
| 0.003000052 |
| 0.001288384 |
| 0.68393626 |
| 0.999999992 |
| 6.75682E-05 |

| | |
|---|---|
| | 0.000743738 |
| | 1.7292E-08 |
| | 3.99258E-08 |
| | 7.65772E-11 |
| | 0.993095021 |
| | 0.803210857 |
| | 8.35128E-10 |
| | 0.000598051 |
| | 0.786875555 |
| | 0.760307682 |
| | 0.138711232 |
| | 2.34414E-06 |
| | 3.36881E-09 |
| | 2.8584E-06 |
| | 1.50323E-08 |
| | 8.42242E-06 |
| | 0.003279346 |
| | 2.26692E-06 |
| | 0.000151342 |
| | 4.39306E-08 |
| | 0.00039481 |
| | 7.89963E-06 |
| | 0.158677231 |
| | 0.360592761 |
| | 0.237632112 |
| | 0.16737018 |
| | 9.69815E-09 |
| | 1 |
| | 0.002316307 |
| | 0.00095194 |
| | 6.43277E-06 |
| | 6.38624E-06 |
| | 3.60805E-05 |
| | 0.565487291 |
| | 0.000158437 |

| | |
|---|---|
| | 0.712773478 |
| | 0.999811327 |
| | 0.000498151 |
| | 4.09978E-08 |
| | 0.999999431 |
| | 0.000518099 |
| | 0.92848239 |
| | 0.000550285 |
| | 4.05652E-09 |
| | 5.6284E-08 |
| | 7.71605E-14 |
| | 4.60743E-14 |
| | 3.25251E-06 |
| | 2.79313E-09 |
| | 1.69533E-06 |
| | 0.998883851 |
| | 6.29847E-07 |
| | 1 |
| | 0.539331373 |
| | 3.71991E-05 |
| | 0.0275948 |
| | 0.967376766 |
| | 3.12272E-12 |
| | 4.15556E-13 |
| | 0.000408884 |
| | 0.002075716 |
| | 1.05373E-06 |
| | 0.734436449 |
| | 1.79389E-08 |
| | 0.000585582 |
| | 0.337184031 |
| | 0.459792137 |
| | 1.25249E-08 |
| | 1.06175E-06 |
| | 1.30076E-05 |

| | |
|---|---|
| | 0.052532732 |
| | 1.54623E-05 |
| | 4.66068E-08 |
| | 0.97158186 |
| | 0.791626429 |
| | 0.081536704 |
| | 0.171787846 |
| | 0.060433175 |
| | 3.11764E-06 |
| | 0.994263627 |
| | 0.82116809 |
| | 7.68586E-07 |
| | 0.999993926 |
| | 0.306099309 |
| | 4.60743E-14 |
| | 7.30621E-07 |
| | 0.033162229 |
| | 0.000832289 |
| | 4.65183E-14 |
| | 0.001296819 |
| | 0.849236588 |
| | 0.006748613 |
| | 0.250776112 |
| | 0.968243103 |
| | 7.4899E-12 |
| | 0.998648771 |
| | 0.973886191 |
| | 0.565857128 |
| | 0.00028065 |
| | 4.60743E-14 |
| | 5.02733E-09 |
| | 2.43083E-12 |
| | 0.19768826 |
| | 0.155429832 |
| | 0.997060646 |

| | |
|---|---|
| | 0.001029128 |
| | 0.998507718 |
| | 0.218285565 |
| | 0.998149205 |
| | 0.999973345 |
| | 0.994250004 |
| | 0.999905708 |
| | 0.712813951 |
| | 0.997523389 |
| | 0.568919099 |
| | 0.680298424 |
| | 0.001786783 |
| | 1.17436E-11 |
| | 0.000311672 |
| | 1.70565E-10 |
| | 9.07821E-08 |
| | 0.199382003 |
| | 0.846342501 |
| | 0.438699506 |
| | 0.001494319 |
| | 0.947115295 |
| | 0.001532938 |
| | 1.31389E-11 |
| | 0.000274085 |
| | 1.54022E-05 |
| | 1.15026E-07 |
| | 3.51395E-08 |
| | 0.958363237 |
| | 0.802150476 |
| | 8.03243E-05 |
| | 0.001437316 |
| | 0.658085894 |
| | 1.88471E-09 |
| | 0.998736576 |
| | 0.000155182 |

| | |
|---|---|
| | 0.999903805 |
| | 1.95023E-11 |
| | 2.23961E-05 |
| | 3.44502E-13 |
| | 0.000434417 |
| | 1.95527E-06 |
| | 0.519538803 |
| | 0.014812994 |
| | 0.003750097 |
| | 0.42507551 |
| | 2.48239E-07 |
| | 3.15115E-10 |
| | 3.052E-13 |
| | 0.046998151 |
| | 0.999628898 |
| | 0.018547595 |
| | 0.000169858 |
| | 0.372087754 |
| | 2.08708E-09 |
| | 1.91537E-08 |
| | 0.013854747 |
| | 0.000188513 |
| | 9.34794E-06 |
| | 0.000897924 |
| | 0.101555762 |
| | 0.198135888 |
| | 0.080312482 |
| | 0.390963855 |
| | 0.047269676 |
| | 7.6353E-11 |
| | 0.990852758 |
| | 2.84303E-07 |
| | 0.002701225 |
| | 8.98472E-10 |
| | 0.474578239 |

| | |
|---|---|
| | 0.003656263 |
| | 0.970564608 |
| | 0.97694005 |
| | 0.501460293 |
| | 0.864684998 |
| | 2.53257E-08 |
| | 4.83809E-06 |
| | 0.930216587 |
| | 5.61322E-05 |
| | 0.418024964 |
| | 0.682116572 |
| | 0.727039836 |
| | 0.764381612 |
| | 0.742045184 |
| | 0.991917251 |
| | 0.807414592 |
| | 0.000173265 |
| | 4.91829E-14 |
| | 3.21961E-07 |
| | 0.240141078 |
| | 0.038834922 |
| | 0.007889444 |
| | 0.787288213 |
| | 0.90380072 |
| | 1.31849E-09 |
| | 3.26666E-06 |
| | 0.147812677 |
| | 0.056015045 |
| | 0.498946109 |
| | 0.000403355 |
| | 0.578783805 |
| | 0.109559663 |
| | 0.010628988 |
| | 0.018076359 |
| | 2.15014E-11 |

| | |
|---|---|
| | 0.872736152 |
| | 0.193650822 |
| | 0.998326722 |
| | 0.889061193 |
| | 0.006451684 |
| | 0.309215684 |
| | 0.001429547 |
| | 0.999785884 |
| | 0.880149979 |
| | 1.6869E-07 |
| | 0.079828769 |
| | 0.750286559 |
| | 0.004300252 |
| | 3.95502E-10 |
| | 0.507991068 |
| | 0.095002064 |
| | 8.45731E-08 |
| | 0.994372958 |
| | 0.213290801 |
| | 4.65183E-14 |
| | 0.039429912 |
| | 0.24250738 |
| | 7.29734E-05 |
| | 0.002294956 |
| | 0.000257035 |
| | 0.994122933 |
| | 0.227826067 |
| | 0.468330113 |
| | 0.783871755 |
| | 0.028193768 |
| | 0.204329996 |
| | 5.0775E-07 |
| | 0.99140268 |
| | 0.998377657 |
| | 2.54034E-10 |

| | |
|---|---|
| | 2.33636E-06 |
| | 2.04045E-11 |
| | 0.074196419 |
| | 0.98718284 |
| | 0.061812513 |
| | 0.706549724 |
| | 2.08136E-08 |
| | 0.999756094 |
| | 0.229256032 |
| | 1.01583E-06 |
| | 0.034618508 |
| | 7.96567E-06 |
| | 0.999533777 |
| | 0.007863534 |
| | 0.467844157 |
| | 0.055566383 |
| | 3.59736E-11 |
| | 1.54505E-05 |
| | 0.626652115 |
| | 0.000611604 |
| | 1.50716E-08 |
| | 1.33438E-12 |
| | 0.377897658 |
| | 8.37329E-10 |
| | 3.49654E-05 |
| | 3.71835E-08 |
| | 0.034925232 |
| | 3.77823E-11 |
| | 0.000456781 |
| | 7.71849E-05 |
| | 4.82107E-05 |
| | 0.000416279 |
| | 0.927445945 |
| | 0.001908157 |
| | 0.986986893 |

| | |
|---|---|
| | 1.54402E-06 |
| | 0.002704047 |
| | 2.62153E-06 |
| | 1.10875E-05 |
| | 0.0147723 |
| | 0.45439703 |
| | 0.271369707 |
| | 1.51557E-12 |
| | 1.05863E-06 |
| | 3.33342E-06 |
| | 0.999715561 |
| | 0.000934471 |
| | 0.330419457 |
| | 0.963001566 |
| | 5.03904E-05 |
| | 0.999739761 |
| | 0.000333602 |
| | 1.3001E-08 |
| | 1.06878E-11 |
| | 1.10298E-05 |
| | 8.46147E-06 |
| | 0.209639794 |
| | 0.999770439 |
| | 1.42881E-08 |
| | 0.000116196 |
| | 7.79816E-06 |
| | 0.00085365 |
| | 1.88849E-05 |
| | 0.014644557 |
| | 0.0162973 |
| | 0.195025685 |
| | 0.961440262 |
| | 0.000167333 |
| | 0.929975739 |
| | 0.990462849 |

| | |
|---|---|
| | 0.000783509 |
| | 0.558631008 |
| | 0.97404968 |
| | 0.000634075 |
| | 0.00031292 |
| | 0.743436531 |
| | 9.2569E-08 |
| | 1.02586E-06 |
| | 0.000711462 |
| | 0.000206688 |
| | 1.33882E-07 |
| | 2.04966E-08 |
| | 1.09847E-05 |
| | 0.000143329 |
| | 0.432761064 |
| | 6.44006E-08 |
| | 0.005682451 |
| | 3.01096E-05 |
| | 0.266870546 |
| | 0.000364338 |
| | 4.79517E-09 |
| | 0.002123417 |
| | 0.631675199 |
| | 7.47553E-09 |
| | 2.34189E-06 |
| | 0.999449387 |
| | 0.352126175 |
| | 0.000181349 |
| | 0.000105075 |
| | 0.93037302 |
| | 0.867022778 |
| | 2.04807E-08 |
| | 5.46804E-08 |
| | 0.000129129 |
| | 7.25882E-10 |

| | |
|---|---|
| | 0.997012199 |
| | 0.000411636 |
| | 4.53648E-12 |
| | 1.72828E-08 |
| | 3.73438E-06 |
| | 5.19381E-08 |
| | 8.97768E-09 |
| | 7.81232E-06 |
| | 2.55462E-13 |
| | 0.000724981 |
| | 0.003443064 |
| | 1.38157E-11 |
| | 1.36773E-08 |
| | 8.19151E-06 |
| | 0.747149222 |
| | 9.36114E-05 |
| | 0.004043817 |
| | 0.000546595 |
| | 4.55674E-07 |
| | 0.41372951 |
| | 0.049451096 |
| | 0.990825861 |
| | 2.00211E-05 |
| | 1.49939E-08 |
| | 0.99999923 |
| | 0.99999998 |
| | 6.33047E-11 |
| | 0.027119995 |
| | 0.878467578 |
| | 0.006066745 |
| | 5.86897E-12 |
| | 0.717888322 |
| | 0.001966256 |
| | 0.787999628 |
| | 0.999991862 |

| |
|---|
| 0.000820318 |
| 6.4963E-05 |
| 0.999999327 |
| 0.991876731 |
| 0.255768436 |
| 0.998642897 |
| 1.18289E-09 |
| 9.17921E-07 |
| 5.29358E-06 |
| 6.92556E-08 |
| 0.000803212 |
| 8.03658E-06 |
| 1.48299E-05 |
| 0.00050284 |
| 9.46225E-06 |
| 0.753431535 |
| 2.84118E-10 |
| 1.22712E-05 |
| 0.734930244 |
| 0.420785917 |
| 0.000265456 |
| 0.00059708 |
| 0.07818371 |
| 0.746770851 |
| 0.250793215 |
| 5.29571E-07 |
| 0.000416729 |
| 8.66318E-12 |
| 0.980290961 |
| 0.001508101 |
| 0.076231141 |
| 0.864405801 |
| 0.0033756 |
| 1.65679E-05 |
| 0.000560462 |

| | |
|---|---|
| | 1.39708E-10 |
| | 8.83491E-08 |
| | 1.30754E-06 |
| | 0.440676365 |
| | 0.008962609 |
| | 0.959773171 |
| | 0.911367424 |
| | 0.999747863 |
| | 0.001528437 |
| | 0.000367079 |
| | 0.441930304 |
| | 0.002222039 |
| | 1.31987E-11 |
| | 0.99960481 |
| | 0.011973835 |
| | 0.999989959 |
| | 4.38491E-05 |
| | 0.005230673 |
| | 0.980636608 |
| | 3.05229E-07 |
| | 0.003836403 |
| | 0.845250844 |
| | 1.08118E-08 |
| | 1.56172E-07 |
| | 3.90922E-05 |
| | 4.6242E-07 |
| | 0.992062904 |
| | 1.13798E-13 |
| | 0.005286236 |
| | 0.002119634 |
| | 3.16773E-07 |
| | 2.68212E-09 |
| | 5.68122E-05 |
| | 0.999870787 |
| | 0.018668771 |

| | |
|---|---|
| | 0.616387566 |
| | 0.004582862 |
| | 0.985653 |
| | 0.29407094 |
| | 0.030196438 |
| | 1.19926E-09 |
| | 0.079024259 |
| | 0.223537809 |
| | 1.44983E-06 |
| | 0.06862658 |
| | 0.000266315 |
| | 0.562899229 |
| | 0.00745557 |
| | 0.999038733 |
| | 0.424705873 |
| | 1.68017E-05 |
| | 0.001160743 |
| | 0.987798033 |
| | 0.000277438 |
| | 0.026355875 |
| | 0.223313096 |
| | 0.047224218 |
| | 0.000113411 |
| | 0.977427291 |
| | 0.697667525 |
| | 7.43878E-09 |
| | 0.880882172 |
| | 2.70195E-12 |
| | 8.80839E-06 |
| | 0.581564307 |
| | 0.169624618 |
| | 0.00031745 |
| | 9.43512E-08 |
| | 0.022876591 |
| | 2.79658E-08 |

| |
|---|
| 0.904305983 |
| 0.9945672 |
| 0.807857041 |
| 0.02851193 |
| 0.000469403 |
| 0.005687549 |
| 9.88934E-08 |
| 0.834794223 |
| 4.76286E-14 |
| 3.44309E-07 |
| 0.504085355 |
| 0.360645462 |
| 0.871058051 |
| 0.188601784 |
| 5.56127E-08 |
| 0.225843683 |
| 0.034600333 |
| 0.00880144 |
| 0.000196471 |
| 0.993304285 |
| 0.129817912 |
| 4.60743E-14 |
| 3.88524E-05 |
| 0.000143929 |
| 0.000310157 |
| 3.36323E-05 |
| 2.13263E-12 |
| 0.992710948 |
| 3.06812E-10 |
| 0.280102573 |
| 0.380451162 |
| 0.000911717 |
| 1.69777E-08 |
| 0.002064598 |
| 0.000471234 |

| |
|---|
| 4.80727E-14 |
| 0.349023632 |
| 4.84548E-07 |
| 0.999738835 |
| 0.999411207 |
| 0.998795014 |
| 0.978237012 |
| 0.951819229 |
| 0.001399395 |
| 0.079059435 |
| 0.984041164 |
| 0.868866952 |
| 0.09760254 |
| 1.32219E-09 |
| 0.10075001 |
| 0.122186405 |
| 1.72951E-10 |
| 0.049726728 |
| 0.320659696 |
| 0.891148161 |
| 2.51592E-06 |
| 0.005219202 |
| 0.955810559 |
| 0.001040585 |
| 0.015162648 |
| 0.999973483 |
| 8.51079E-09 |
| 0.999999852 |
| 0.070940442 |
| 0.212384035 |
| 0.351819651 |
| 0.06796718 |
| 0.002555207 |
| 0.998304216 |
| 0.292926341 |

| | |
|---|---|
| | 0.000120166 |
| | 0.978468337 |
| | 0.941136024 |
| | 0.999769901 |
| | 0.460488331 |
| | 4.01696E-08 |
| | 4.67404E-14 |
| | 3.6421E-08 |
| | 0.006105905 |
| | 2.48314E-11 |
| | 6.83281E-06 |
| | 0.093535651 |
| | 1.55446E-10 |
| | 0.439906375 |
| | 4.88229E-11 |
| | 0.046773155 |
| | 0.007243188 |
| | 0.994799759 |
| | 2.03416E-06 |
| | 0.002282007 |
| | 0.996806967 |
| | 0.006143713 |
| | 0.92298541 |
| | 0.001330836 |
| | 0.000112869 |
| | 0.999841805 |
| | 1.06989E-05 |
| | 3.38228E-11 |
| | 0.347310311 |
| | 0.970755982 |
| | 2.68438E-10 |
| | 0.608868322 |
| | 0.000986337 |
| | 5.05595E-06 |
| | 6.58923E-09 |

| | |
|---|---|
| | 0.005217201 |
| | 1.13605E-10 |
| | 2.24387E-10 |
| | 0.907289008 |
| | 1.49495E-11 |
| | 0.000105012 |
| | 0.437401489 |
| | 2.32813E-11 |
| | 0.885560527 |
| | 0.016475436 |
| | 4.62963E-14 |
| | 1 |
| | 0.906724101 |
| | 3.76166E-05 |
| | 0.002022493 |
| | 0.003887669 |
| | 0.556472135 |
| | 0.662937427 |
| | 0.70536396 |
| | 0.6721199 |
| | 0.001313314 |
| | 0.999042294 |
| | 0.999953126 |
| | 0.608038266 |
| | 0.947901546 |
| | 0.953160848 |
| | 0.979401198 |
| | 0.020570555 |
| | 0.774159379 |
| | 0.0091755 |
| | 0.002159999 |
| | 0.045532296 |
| | 0.405691699 |
| | 4.71845E-14 |
| | 0.976357174 |

| |
|---|
| 0.006496167 |
| 0.966139484 |
| 2.17874E-08 |
| 0.241597058 |
| 3.73128E-06 |
| 3.68593E-11 |
| 0.995451197 |
| 4.32998E-12 |
| 0.982661236 |
| 9.26655E-06 |
| 4.60743E-14 |
| 4.65183E-14 |
| 1.34526E-12 |
| 3.76877E-07 |
| 2.36876E-06 |
| 0.963072575 |
| 1.28687E-07 |
| 0.039172869 |
| 0.259911442 |
| 0.033339454 |
| 0.688930755 |
| 0.685487429 |
| 3.20131E-05 |
| 0.003696859 |
| 0.000830744 |
| 0.003527485 |
| 6.32406E-09 |
| 0.999999936 |
| 0.999830926 |
| 0.649788758 |
| 2.94083E-07 |
| 0.19457261 |
| 0.999978939 |
| 0.005770707 |
| 0.033040818 |

| |
|---|
| 0.905734031 |
| 0.927700993 |
| 0.942659933 |
| 0.742537394 |
| 0.037442327 |
| 3.08229E-06 |
| 8.572E-05 |
| 0.912391723 |
| 0.005591869 |
| 0.789372834 |
| 0.99393126 |
| 0.298115804 |
| 0.999996011 |
| 0.283689931 |
| 0.250612245 |

| |
|---|
| P-value: (Bb_pos_12022024_Newanalysis) / (SM_pos_12022024_Newanalyis) |
| 0.300741659 |
| 0.003304043 |
| 0.969053469 |
| 0.002612076 |
| 0.001177524 |
| 0.491276694 |
| 1.57049E-05 |
| 0.79031936 |
| 5.47958E-05 |
| 0.227901492 |
| 0.004174453 |
| 0.400728421 |
| 0.014518949 |
| 0.000954698 |
| 0.976951091 |
| 0.007979153 |
| 0.999479602 |
| 0.059472499 |
| 0.136157033 |
| 0.931228793 |
| 0.068727059 |
| 0.008512018 |
| 0.026861986 |
| 0.001867856 |
| 2.15001E-07 |
| 0.389757304 |
| 0.978431925 |
| 0.992638168 |
| 7.74604E-08 |
| 0.005298721 |
| 0.987746691 |
| 1.11254E-07 |
| 0.244088151 |

| |
|---|
| 0.557805245 |
| 7.11728E-09 |
| 0.950153584 |
| 0.978485613 |
| 0.000433232 |
| 6.35307E-05 |
| 0.942000663 |
| 0.004920466 |
| 0.002902179 |
| 0.005453497 |
| 0.074154996 |
| 0.000263052 |
| 0.000161402 |
| 0.000262712 |
| 0.186447915 |
| 0.994119501 |
| 0.158741509 |
| 0.993416367 |
| 0.172756845 |
| 0.000437287 |
| 0.120439772 |
| 7.73238E-05 |
| 0.001390064 |
| 0.000303449 |
| 0.999295335 |
| 0.459747804 |
| 1.27036E-06 |
| 0.052685923 |
| 0.000933068 |
| 2.52053E-07 |
| 0.000998984 |
| 0.999970424 |
| 1.4506E-05 |
| 5.62379E-07 |
| 6.48209E-09 |

| | |
|---|---|
| | 2.56423E-07 |
| | 0.128825119 |
| | 9.42325E-10 |
| | 4.91666E-08 |
| | 1.09311E-06 |
| | 2.58838E-10 |
| | 3.15394E-06 |
| | 1.07167E-05 |
| | 5.0726E-10 |
| | 0.053193864 |
| | 0.999435483 |
| | 0.000316416 |
| | 0.814497696 |
| | 4.60743E-14 |
| | 0.000473645 |
| | 2.89679E-08 |
| | 5.77264E-05 |
| | 0.011021141 |
| | 0.994642916 |
| | 0.896999059 |
| | 4.88961E-06 |
| | 0.015449407 |
| | 0.000142367 |
| | 0.997130859 |
| | 0.000155659 |
| | 0.000667209 |
| | 0.222737181 |
| | 0.000276087 |
| | 0.049772214 |
| | 0.010820426 |
| | 0.817949454 |
| | 0.999100425 |
| | 0.511448079 |
| | 2.72204E-05 |
| | 2.60203E-12 |

| | |
|---|---|
| | 0.463487623 |
| | 0.953664177 |
| | 0.000784393 |
| | 0.002548244 |
| | 0.0982952 |
| | 0.872371907 |
| | 0.885084622 |
| | 4.34363E-07 |
| | 0.999933819 |
| | 0.0736549 |
| | 1.19202E-05 |
| | 2.49106E-05 |
| | 1.46894E-05 |
| | 0.030607419 |
| | 0.793355163 |
| | 0.497750608 |
| | 0.236965294 |
| | 0.001424468 |
| | 0.019772941 |
| | 0.683972585 |
| | 4.64382E-06 |
| | 1.08171E-05 |
| | 0.98584471 |
| | 0.091650528 |
| | 4.13542E-08 |
| | 9.13112E-05 |
| | 0.99449791 |
| | 0.863670233 |
| | 0.437733264 |
| | 0.000845551 |
| | 0.998090077 |
| | 2.59922E-10 |
| | 2.78174E-05 |
| | 0.012261536 |
| | 0.11143748 |

| | |
|---|---|
| | 0.00016145 |
| | 0.541496425 |
| | 3.5199E-05 |
| | 0.75541798 |
| | 0.687838353 |
| | 0.887097275 |
| | 0.570999855 |
| | 0.957796666 |
| | 2.52322E-05 |
| | 0.999998912 |
| | 0.99788477 |
| | 0.152906647 |
| | 3.52429E-09 |
| | 0.55039281 |
| | 0.012426201 |
| | 1.12474E-10 |
| | 0.394606614 |
| | 6.20821E-06 |
| | 1.21869E-07 |
| | 0.999759184 |
| | 9.29933E-05 |
| | 0.000797437 |
| | 0.00695272 |
| | 0.000113539 |
| | 1.66013E-08 |
| | 0.613135109 |
| | 3.07944E-07 |
| | 0.069351164 |
| | 0.026190223 |
| | 0.035754444 |
| | 0.021212668 |
| | 0.133653551 |
| | 4.03066E-07 |
| | 0.004244202 |
| | 0.019447491 |

| |
|---|
| 0.999978086 |
| 0.001592817 |
| 0.791148452 |
| 1.83823E-08 |
| 0.003726452 |
| 0.99522643 |
| 0.05590139 |
| 0.000114411 |
| 0.513696355 |
| 4.12345E-06 |
| 0.000292939 |
| 3.84478E-10 |
| 2.96346E-11 |
| 0.001036461 |
| 0.001507958 |
| 5.89818E-05 |
| 0.230539546 |
| 1.3037E-07 |
| 4.80727E-14 |
| 0.022246732 |
| 0.000835083 |
| 0.000108844 |
| 1.76875E-06 |
| 0.001091112 |
| 2.5382E-07 |
| 0.999996634 |
| 0.915766947 |
| 0.005227194 |
| 0.002976887 |
| 0.798480126 |
| 1.91819E-07 |
| 0.791660139 |
| 0.001808911 |
| 0.113287517 |
| 6.60201E-08 |

| |
|---|
| 0.008935075 |
| 0.005767116 |
| 0.000221075 |
| 0.046378769 |
| 0.001669565 |
| 2.77556E-10 |
| 2.43937E-11 |
| 0.020082761 |
| 2.27255E-07 |
| 0.075581851 |
| 0.015096638 |
| 0.043636431 |
| 2.66331E-05 |
| 0.726334349 |
| 0.012982046 |
| 1.75165E-05 |
| 0.951284693 |
| 3.19612E-05 |
| 0.622208417 |
| 0.268253834 |
| 2.3894E-05 |
| 0.910423238 |
| 0.495053285 |
| 3.76956E-08 |
| 8.14729E-07 |
| 0.971573683 |
| 0.08012005 |
| 0.838034907 |
| 6.34774E-05 |
| 0.002798478 |
| 0.986153597 |
| 0.468071016 |
| 0.979522806 |
| 0.963187396 |
| 0.007317427 |

| |
|---|
| 0.987969482 |
| 0.982458233 |
| 0.998000202 |
| 0.999690755 |
| 0.879214691 |
| 1.99199E-08 |
| 0.019204238 |
| 0.775909464 |
| 0.000126842 |
| 1.10466E-09 |
| 0.279239511 |
| 0.970897699 |
| 0.790623069 |
| 0.071127105 |
| 0.003403071 |
| 0.986350669 |
| 6.16459E-08 |
| 0.735910374 |
| 0.357709263 |
| 0.667644982 |
| 0.006686657 |
| 0.001823271 |
| 0.00719722 |
| 5.08859E-05 |
| 0.018284677 |
| 0.99461248 |
| 0.751292212 |
| 0.819504104 |
| 0.649146212 |
| 0.996355121 |
| 0.000652521 |
| 0.0001393 |
| 0.000933459 |
| 0.738141557 |
| 7.24976E-14 |

| | |
|---|---|
| | 0.003369282 |
| | 0.355917509 |
| | 0.000222821 |
| | 0.029688932 |
| | 4.27473E-07 |
| | 0.4498249 |
| | 1.59334E-06 |
| | 0.917150807 |
| | 0.000690425 |
| | 0.308747442 |
| | 0.179683851 |
| | 0.990261105 |
| | 0.940808327 |
| | 1.00773E-10 |
| | 0.122734259 |
| | 0.6375709 |
| | 0.955464464 |
| | 0.775657837 |
| | 3.53752E-07 |
| | 0.999216619 |
| | 0.260319068 |
| | 2.96174E-08 |
| | 0.057285299 |
| | 2.44584E-06 |
| | 0.999998247 |
| | 0.999073819 |
| | 0.837699778 |
| | 0.001099155 |
| | 0.023259229 |
| | 3.51327E-05 |
| | 0.999940833 |
| | 0.311854648 |
| | 0.990167585 |
| | 1.10996E-08 |
| | 0.010103966 |

| |
|---|
| 2.37423E-05 |
| 0.013548942 |
| 0.183513584 |
| 0.177839941 |
| 0.986930389 |
| 0.576055413 |
| 6.71302E-05 |
| 0.000112587 |
| 5.88938E-08 |
| 2.90576E-05 |
| 0.009953724 |
| 0.992818509 |
| 3.89757E-07 |
| 4.24107E-05 |
| 0.0001727 |
| 0.921768345 |
| 0.819502243 |
| 0.389089381 |
| 0.002424045 |
| 0.999970606 |
| 4.446E-10 |
| 0.582346837 |
| 0.005505328 |
| 0.002287031 |
| 0.492353614 |
| 0.008656563 |
| 0.999490364 |
| 5.92513E-08 |
| 4.94534E-08 |
| 1.05241E-05 |
| 0.007634764 |
| 0.000307527 |
| 0.007302118 |
| 1.08821E-09 |
| 0.002715779 |

| |
|---|
| 2.19879E-09 |
| 0.999997184 |
| 2.77445E-13 |
| 0.061343437 |
| 0.999995702 |
| 0.986407342 |
| 0.637979342 |
| 0.000244315 |
| 0.855685358 |
| 0.007306698 |
| 5.70591E-05 |
| 0.064913356 |
| 0.694099558 |
| 1.43851E-09 |
| 0.083532168 |
| 0.021118804 |
| 0.003629466 |
| 0.999931035 |
| 0.064111531 |
| 0.999638576 |
| 3.8235E-12 |
| 1.00039E-06 |
| 0.111090841 |
| 0.740902079 |
| 0.000827964 |
| 0.993531944 |
| 0.205876223 |
| 0.791963506 |
| 0.999999857 |
| 2.37694E-07 |
| 0.93961294 |
| 0.030932688 |
| 0.886739645 |
| 0.864748996 |
| 2.85804E-11 |

| | |
|---|---|
| | 0.487452256 |
| | 4.47034E-07 |
| | 2.96848E-06 |
| | 0.999997797 |
| | 0.034323082 |
| | 0.999999162 |
| | 0.999997372 |
| | 0.774437229 |
| | 0.001447989 |
| | 0.000275188 |
| | 0.845127373 |
| | 1 |
| | 0.959825 |
| | 0.987016135 |
| | 3.06726E-06 |
| | 0.998607116 |
| | 0.432639006 |
| | 1.32465E-07 |
| | 1.16902E-06 |
| | 6.58826E-06 |
| | 0.606342714 |
| | 1.16593E-07 |
| | 0.988925347 |
| | 0.782837374 |
| | 0.718086609 |
| | 0.997461729 |
| | 3.46733E-09 |
| | 1.10952E-07 |
| | 0.319638721 |
| | 0.999989799 |
| | 0.704004093 |
| | 7.51121E-12 |
| | 7.41296E-13 |
| | 3.38899E-07 |
| | 0.000717774 |

| |
|---|
| 0.003107178 |
| 4.87441E-10 |
| 0.000959924 |
| 6.54857E-05 |
| 0.011773221 |
| 0.998813539 |
| 5.25494E-06 |
| 0.063665427 |
| 0.998265117 |
| 0.453277572 |
| 0.785981375 |
| 1.81424E-05 |
| 0.000482742 |
| 2.64996E-11 |
| 0.987468342 |
| 1.35045E-06 |
| 5.93865E-10 |
| 0.002784522 |
| 8.14114E-11 |
| 4.60743E-14 |
| 0.989275605 |
| 2.23297E-10 |
| 0.000377079 |
| 2.18926E-08 |
| 0.000469139 |
| 4.60743E-14 |
| 0.009883066 |
| 0.879105322 |
| 0.03122497 |
| 3.3103E-05 |
| 0.900618469 |
| 1 |
| 0.055835152 |
| 0.000152599 |
| 5.31085E-09 |

| | |
|---|---|
| | 0.987001767 |
| | 0.921218084 |
| | 0.999962698 |
| | 0.000802516 |
| | 0.162458295 |
| | 0.988009855 |
| | 8.35503E-09 |
| | 1.39581E-08 |
| | 0.471871238 |
| | 0.992266742 |
| | 0.284056892 |
| | 0.07785063 |
| | 4.62963E-14 |
| | 0.000209362 |
| | 4.69018E-07 |
| | 4.60743E-14 |
| | 4.60743E-14 |
| | 4.90963E-10 |
| | 0.567273815 |
| | 0.73348173 |
| | 0.107634766 |
| | 0.205913598 |
| | 0.000339817 |
| | 0.236380754 |
| | 1.48307E-06 |
| | 5.42899E-14 |
| | 2.29227E-05 |
| | 0.999999818 |
| | 0.230644096 |
| | 0.993577127 |
| | 0.404336328 |
| | 4.62963E-14 |
| | 1.18556E-08 |
| | 4.60743E-14 |
| | 0.325522252 |

| | |
|---|---|
| | 0.976308928 |
| | 9.51822E-09 |
| | 0.015819933 |
| | 0.913562139 |
| | 0.015809156 |
| | 6.33958E-05 |
| | 0.523856015 |
| | 0.638003236 |
| | 0.025484322 |
| | 0.879810372 |
| | 0.429788387 |
| | 0.026213527 |
| | 0.999992938 |
| | 0.008031244 |
| | 0.021174656 |
| | 0.001209352 |
| | 6.3122E-05 |
| | 2.17281E-06 |
| | 2.09962E-08 |
| | 1.01881E-09 |
| | 0.000150315 |
| | 7.6263E-05 |
| | 4.22801E-07 |
| | 0.988053159 |
| | 0.072415315 |
| | 0.050386272 |
| | 0.999977268 |
| | 0.000266338 |
| | 0.808318634 |
| | 0.988044194 |
| | 9.83684E-05 |
| | 0.023566939 |
| | 0.194468337 |
| | 1.09798E-07 |
| | 0.003769266 |

| |
|---:|
| 0.000196307 |
| 0.018579564 |
| 3.38287E-07 |
| 0.021083664 |
| 4.60743E-14 |
| 0.99982982 |
| 0.365502693 |
| 0.943548722 |
| 0.001000366 |
| 1.77162E-07 |
| 0.070021061 |
| 0.000930173 |
| 0.305739715 |
| 1.90062E-05 |
| 0.999999641 |
| 1.11797E-08 |
| 0.019606003 |
| 4.65183E-14 |
| 0.000708538 |
| 0.354578567 |
| 3.37822E-11 |
| 1.74698E-10 |
| 4.60743E-14 |
| 0.741351251 |
| 1.10467E-13 |
| 0.402003665 |
| 0.104428636 |
| 0.02356555 |
| 4.62963E-14 |
| 8.84848E-14 |
| 0.574218604 |
| 0.999887895 |
| 0.029517055 |
| 4.60743E-14 |
| 3.06291E-06 |

| | |
|---|---|
| | 4.60743E-14 |
| | 0.848289545 |
| | 1.05077E-08 |
| | 0.010714976 |
| | 0.052857869 |
| | 0.060271081 |
| | 0.163725904 |
| | 0.123671351 |
| | 0.048192566 |
| | 0.159500289 |
| | 0.319598678 |
| | 0.999999661 |
| | 4.64316E-10 |
| | 0.677081644 |
| | 0.066150493 |
| | 0.080200638 |
| | 1.34308E-09 |
| | 0.992891308 |
| | 0.997972776 |
| | 0.951513458 |
| | 5.89417E-13 |
| | 0.180498545 |
| | 0.27778785 |
| | 0.977873941 |
| | 5.75979E-09 |
| | 3.45937E-07 |
| | 0.00961043 |
| | 0.002108331 |
| | 0.000194402 |
| | 0.028325705 |
| | 0.983886978 |
| | 0.09107753 |
| | 0.897551884 |
| | 0.999727696 |
| | 0.237718033 |

| | |
|---|---|
| | 0.999884082 |
| | 0.354651675 |
| | 0.986563917 |
| | 0.658328083 |
| | 0.025727777 |
| | 0.00010137 |
| | 8.73419E-06 |
| | 0.102033356 |
| | 0.000144787 |
| | 2.77808E-09 |
| | 0.335663509 |
| | 5.37383E-09 |
| | 0.985718138 |
| | 3.41328E-05 |
| | 5.59029E-11 |
| | 0.016574878 |
| | 2.87738E-09 |
| | 0.968353228 |
| | 0.038004035 |
| | 7.25193E-06 |
| | 0.99731096 |
| | 0.028645112 |
| | 0.990970637 |
| | 0.025972777 |
| | 0.492830299 |
| | 0.769538614 |
| | 0.103806566 |
| | 0.604502875 |
| | 0.999598876 |
| | 0.998870307 |
| | 0.006860642 |
| | 0.009833086 |
| | 0.797943814 |
| | 0.010457428 |
| | 0.067007288 |

| | |
|---|---|
| | 3.09252E-05 |
| | 0.652416856 |
| | 0.868703469 |
| | 1.69486E-05 |
| | 0.111444927 |
| | 6.6136E-13 |
| | 8.82977E-10 |
| | 4.60743E-14 |
| | 8.85481E-12 |
| | 0.139395467 |
| | 1.50151E-10 |
| | 0.206355915 |
| | 0.089167272 |
| | 5.45531E-06 |
| | 0.961344454 |
| | 7.91902E-05 |
| | 0.057503123 |
| | 0.011835446 |
| | 9.79895E-08 |
| | 0.006043025 |
| | 5.48966E-05 |
| | 0.983487603 |
| | 0.964951437 |
| | 0.999754574 |
| | 0.99998245 |
| | 0.951509275 |
| | 0.999660849 |
| | 0.377771972 |
| | 0.136295093 |
| | 0.003591243 |
| | 0.098705333 |
| | 0.1929786 |
| | 0.033675425 |
| | 0.000188447 |
| | 0.001496519 |

| | |
|---|---|
| | 3.53502E-06 |
| | 0.999961716 |
| | 1.47468E-08 |
| | 0.014579806 |
| | 0.998619898 |
| | 0.220948948 |
| | 0.054584577 |
| | 0.946307263 |
| | 0.658780607 |
| | 0.003160933 |
| | 8.56567E-06 |
| | 0.327774945 |
| | 0.985558186 |
| | 0.999693223 |
| | 0.752384077 |
| | 0.239756525 |
| | 0.614196126 |
| | 0.229860363 |
| | 0.164010302 |
| | 0.091460863 |
| | 6.2504E-05 |
| | 0.99999997 |
| | 1.16784E-12 |
| | 0.171714452 |
| | 0.150697748 |
| | 0.060687747 |
| | 5.58442E-14 |
| | 0.005616393 |
| | 0.000546369 |
| | 0.082983267 |
| | 0.049648106 |
| | 0.056969997 |
| | 0.091653217 |
| | 0.07938733 |
| | 0.00510865 |

| |
|---|
| 0.999999682 |
| 0.724902475 |
| 0.999917855 |
| 0.006050278 |
| 0.997230245 |
| 0.002048892 |
| 0.334255767 |
| 0.999898748 |
| 6.16701E-06 |
| 0.310831154 |
| 0.999998978 |
| 0.998665575 |
| 0.636525002 |
| 0.267945233 |
| 2.08062E-09 |

| |
|---|
| P-value: (CM_pos_12022024_Newanalysis) / (SM_pos_12022024_Newanalyis) |
| 0.910239641 |
| 0.117943683 |
| 0.947943121 |
| 0.996343532 |
| 0.010225575 |
| 0.803716856 |
| 0.000397609 |
| 0.099329698 |
| 4.56388E-06 |
| 0.549251559 |
| 0.020540129 |
| 0.96808641 |
| 0.570848224 |
| 0.064763036 |
| 0.949609067 |
| 0.784066546 |
| 0.579034582 |
| 2.33924E-13 |
| 0.001773192 |
| 5.17679E-05 |
| 0.780989468 |
| 3.126E-07 |
| 0.241370539 |
| 5.03921E-09 |
| 1.63397E-11 |
| 0.949812368 |
| 0.739790527 |
| 0.79570249 |
| 4.96898E-05 |
| 0.018408844 |
| 0.999968712 |
| 0.995067267 |
| 0.003230313 |

| | |
|---|---|
| | 0.019890694 |
| | 0.00041673 |
| | 0.453729748 |
| | 0.000261679 |
| | 0.999985292 |
| | 0.990944343 |
| | 1.02604E-07 |
| | 0.001582518 |
| | 3.9378E-05 |
| | 0.998527922 |
| | 1.5029E-05 |
| | 5.46018E-05 |
| | 7.63153E-05 |
| | 0.011154251 |
| | 2.4762E-05 |
| | 0.000536985 |
| | 0.130486758 |
| | 4.82139E-05 |
| | 0.003113077 |
| | 3.4731E-05 |
| | 0.180642966 |
| | 0.00189298 |
| | 0.067268969 |
| | 1.01345E-08 |
| | 0.426689916 |
| | 0.958748909 |
| | 0.00672328 |
| | 0.476112792 |
| | 0.00050797 |
| | 0.000152208 |
| | 0.002395027 |
| | 0.000493503 |
| | 4.7936E-06 |
| | 0.000512294 |
| | 4.52368E-07 |

| |
|---|
| 6.4643E-06 |
| 0.781737886 |
| 0.154447628 |
| 1.3571E-08 |
| 0.289871987 |
| 5.27441E-07 |
| 9.95E-06 |
| 0.001163759 |
| 0.000550946 |
| 0.205966443 |
| 1.18721E-10 |
| 7.51288E-13 |
| 0.001622979 |
| 0.00069102 |
| 0.000291938 |
| 0.090622761 |
| 0.079922908 |
| 0.998936272 |
| 0.999944793 |
| 2.25748E-06 |
| 0.016398475 |
| 0.958463982 |
| 3.02339E-11 |
| 1.57412E-09 |
| 0.901743224 |
| 0.719502184 |
| 0.045372905 |
| 0.586995336 |
| 7.34544E-06 |
| 0.578758951 |
| 0.725155861 |
| 0.441174703 |
| 0.649456098 |
| 0.002932709 |
| 2.13683E-07 |

| |
|---|
| 0.782691822 |
| 0.033098449 |
| 7.67901E-05 |
| 0.597776649 |
| 0.223490636 |
| 0.999979052 |
| 0.898877234 |
| 0.001320737 |
| 0.565619985 |
| 0.08207035 |
| 0.05648625 |
| 5.44313E-06 |
| 0.00600176 |
| 0.789396346 |
| 4.60743E-14 |
| 4.75124E-06 |
| 0.734308376 |
| 0.000175873 |
| 0.01530454 |
| 0.840547103 |
| 0.003473347 |
| 5.06988E-05 |
| 0.899342465 |
| 0.999983685 |
| 4.44829E-09 |
| 0.000163801 |
| 0.999843456 |
| 0.999218756 |
| 0.993530899 |
| 4.06009E-13 |
| 0.003351197 |
| 1.26454E-13 |
| 0.430960234 |
| 0.99010381 |
| 0.358605839 |

| | |
|---|---|
| | 0.000287693 |
| | 0.046878234 |
| | 0.004703691 |
| | 0.998756436 |
| | 0.921039401 |
| | 0.942668686 |
| | 0.994863486 |
| | 0.576381084 |
| | 0.640724544 |
| | 0.999978372 |
| | 0.701747085 |
| | 0.043480696 |
| | 0.367058344 |
| | 0.999979385 |
| | 0.460806523 |
| | 0.435791896 |
| | 0.438024302 |
| | 0.000376041 |
| | 0.000358703 |
| | 0.748646368 |
| | 0.012199136 |
| | 0.089256554 |
| | 0.484205261 |
| | 0.987279787 |
| | 1.77392E-08 |
| | 7.83325E-06 |
| | 0.000197532 |
| | 0.816599622 |
| | 0.494523769 |
| | 0.20041868 |
| | 0.920347526 |
| | 0.999999819 |
| | 0.000242464 |
| | 0.090744084 |
| | 0.991285929 |

| | |
|---|---|
| | 0.032539615 |
| | 0.042501094 |
| | 0.03721651 |
| | 0.377042112 |
| | 0.929198093 |
| | 0.836831924 |
| | 0.999999977 |
| | 0.001294036 |
| | 0.666622049 |
| | 0.000106287 |
| | 0.064765197 |
| | 2.54329E-11 |
| | 0.999617144 |
| | 0.999984137 |
| | 0.286368014 |
| | 0.013309995 |
| | 0.10157634 |
| | 0.000126121 |
| | 0.200723522 |
| | 0.988718103 |
| | 0.815355596 |
| | 0.630883869 |
| | 0.002501132 |
| | 4.20215E-05 |
| | 0.001306583 |
| | 0.997221424 |
| | 0.593439771 |
| | 0.387898921 |
| | 0.998134928 |
| | 2.84916E-06 |
| | 0.338248787 |
| | 0.000118583 |
| | 0.93453702 |
| | 0.000273451 |
| | 6.04356E-05 |

| | |
|---|---|
| | 0.13474444 |
| | 0.014913789 |
| | 0.012846588 |
| | 0.689971097 |
| | 0.319187257 |
| | 4.11401E-10 |
| | 1.22727E-09 |
| | 0.011357367 |
| | 8.59669E-08 |
| | 0.002370611 |
| | 0.801810621 |
| | 0.643490344 |
| | 0.027379644 |
| | 0.949869024 |
| | 0.99969163 |
| | 0.85506122 |
| | 8.37421E-06 |
| | 0.239695124 |
| | 0.001924259 |
| | 0.632394172 |
| | 0.482274701 |
| | 1 |
| | 0.164813303 |
| | 1.48308E-06 |
| | 0.713717086 |
| | 0.359185167 |
| | 0.074627333 |
| | 0.479651102 |
| | 0.145347129 |
| | 0.002780448 |
| | 0.341795762 |
| | 0.134534344 |
| | 0.916454178 |
| | 0.998341075 |
| | 0.025839812 |

| |
|---|
| 0.379755319 |
| 0.997570479 |
| 0.993996245 |
| 0.71271597 |
| 0.704239321 |
| 0.000451689 |
| 0.963160452 |
| 0.74560183 |
| 0.013295656 |
| 0.343007236 |
| 0.675088757 |
| 0.322376039 |
| 0.999999591 |
| 0.803836931 |
| 0.644989032 |
| 0.027346779 |
| 0.018458714 |
| 0.107286073 |
| 0.568538899 |
| 1.31702E-11 |
| 0.340809331 |
| 5.78984E-08 |
| 0.507055681 |
| 7.07988E-08 |
| 0.008930681 |
| 0.859460284 |
| 0.99999991 |
| 0.997854711 |
| 0.000215816 |
| 0.831087484 |
| 0.294147253 |
| 0.057218187 |
| 0.210098168 |
| 0.960505558 |
| 0.440921089 |

| |
|---|
| 0.085345593 |
| 7.23047E-05 |
| 0.151991532 |
| 0.004159992 |
| 3.05219E-07 |
| 0.997785854 |
| 0.98390218 |
| 0.823073084 |
| 0.022728357 |
| 0.740717112 |
| 3.86159E-05 |
| 0.999999997 |
| 0.999875157 |
| 3.23186E-13 |
| 0.57837738 |
| 0.111783204 |
| 0.012481653 |
| 0.999687694 |
| 0.082344576 |
| 0.999936568 |
| 9.0677E-08 |
| 0.076979077 |
| 0.076883872 |
| 0.020405914 |
| 0.996129228 |
| 0.999339759 |
| 0.999998569 |
| 0.431602104 |
| 5.77663E-08 |
| 0.999993685 |
| 0.122735647 |
| 0.674491595 |
| 0.030785476 |
| 1.74949E-05 |
| 0.921860827 |

| | |
|---|---|
| | 0.100245003 |
| | 0.353799372 |
| | 0.023363166 |
| | 0.258915493 |
| | 0.999999998 |
| | 0.998307714 |
| | 0.382314193 |
| | 0.220866873 |
| | 2.91562E-08 |
| | 0.658804443 |
| | 0.469615742 |
| | 0.843669285 |
| | 0.014939563 |
| | 0.49253158 |
| | 4.04883E-07 |
| | 0.998381672 |
| | 0.900522908 |
| | 0.001381127 |
| | 0.000909661 |
| | 0.002801835 |
| | 1.12184E-06 |
| | 0.202693379 |
| | 0.580957122 |
| | 0.441229955 |
| | 0.991321654 |
| | 3.37598E-05 |
| | 0.105998323 |
| | 0.999994964 |
| | 0.692944874 |
| | 0.999991596 |
| | 0.550474577 |
| | 0.115302482 |
| | 0.341516174 |
| | 0.129115382 |
| | 0.956665221 |

| | |
|---|---|
| | 0.649721812 |
| | 0.999999963 |
| | 0.002327353 |
| | 0.005206988 |
| | 0.9991319 |
| | 0.926119127 |
| | 0.949682146 |
| | 0.032782392 |
| | 0.957471126 |
| | 0.997875412 |
| | 0.8913747 |
| | 0.964656211 |
| | 0.782295706 |
| | 4.40741E-05 |
| | 0.052410049 |
| | 6.02758E-06 |
| | 0.998951545 |
| | 0.973212777 |
| | 0.980881576 |
| | 0.252878591 |
| | 1 |
| | 0.008829676 |
| | 0.766654374 |
| | 0.152399583 |
| | 0.716538405 |
| | 0.999999935 |
| | 0.999989075 |
| | 0.871809982 |
| | 0.738987154 |
| | 0.002150227 |
| | 0.998328036 |
| | 0.000299267 |
| | 0.357591019 |
| | 0.001422034 |
| | 7.68263E-12 |

| | |
|---|---|
| | 0.131815795 |
| | 2.57691E-06 |
| | 0.003188351 |
| | 0.995354101 |
| | 0.000318474 |
| | 0.99999328 |
| | 0.66841289 |
| | 0.995993691 |
| | 1.14765E-07 |
| | 0.291797681 |
| | 0.998774149 |
| | 0.995872888 |
| | 0.981787924 |
| | 0.032263604 |
| | 0.068088589 |
| | 0.074464715 |
| | 0.678245559 |
| | 0.980790525 |
| | 0.836060503 |
| | 0.008891644 |
| | 0.87989922 |
| | 0.005259675 |
| | 0.699433642 |
| | 0.568197092 |
| | 0.812779372 |
| | 0.943579084 |
| | 6.37643E-08 |
| | 0.001765288 |
| | 0.630794228 |
| | 0.216225491 |
| | 5.62886E-06 |
| | 0.337773557 |
| | 0.000456534 |
| | 0.501607419 |
| | 0.425865815 |

| |
|---|
| 0.999937764 |
| 3.14644E-07 |
| 0.001753883 |
| 0.195622812 |
| 0.001107455 |
| 0.015395638 |
| 9.00069E-12 |
| 0.187796545 |
| 0.882218688 |
| 0.885284422 |
| 0.95970005 |
| 0.999660356 |
| 0.514733897 |
| 0.000105776 |
| 0.087768939 |
| 1.08145E-05 |
| 8.50797E-06 |
| 0.042481391 |
| 0.000749442 |
| 0.000111887 |
| 0.993949779 |
| 0.973100026 |
| 0.982425394 |
| 0.02461883 |
| 0.000162503 |
| 2.9258E-11 |
| 0.009038197 |
| 5.91702E-05 |
| 0.957806326 |
| 0.013964696 |
| 0.011253392 |
| 0.999862655 |
| 0.999664724 |
| 0.592484592 |
| 0.912848996 |

| | |
|---|---|
| | 3.64508E-05 |
| | 0.021524942 |
| | 0.222270065 |
| | 0.165241155 |
| | 0.000187672 |
| | 0.800921191 |
| | 0.085732765 |
| | 0.000211331 |
| | 0.993901029 |
| | 0.093602339 |
| | 0.298099565 |
| | 2.27031E-06 |
| | 0.287436563 |
| | 0.098361508 |
| | 0.999720487 |
| | 0.035984652 |
| | 0.665479435 |
| | 0.000191776 |
| | 0.999987493 |
| | 0.871841793 |
| | 0.233716237 |
| | 0.137403905 |
| | 0.919857557 |
| | 0.999999896 |
| | 0.99969719 |
| | 0.98533061 |
| | 0.000778459 |
| | 0.981841667 |
| | 0.010983432 |
| | 0.978423978 |
| | 0.99842054 |
| | 0.823993383 |
| | 0.996693458 |
| | 0.999729825 |
| | 0.089568891 |

| | |
|---|---|
| | 0.871638151 |
| | 0.00703556 |
| | 0.595234308 |
| | 0.994370058 |
| | 0.211940228 |
| | 3.10878E-05 |
| | 0.999782685 |
| | 0.080032784 |
| | 0.000998228 |
| | 0.956313479 |
| | 0.303840705 |
| | 0.054298847 |
| | 0.999977042 |
| | 0.034174761 |
| | 0.154599416 |
| | 0.764400235 |
| | 0.021576907 |
| | 6.34605E-09 |
| | 0.118724683 |
| | 3.31511E-09 |
| | 0.031719072 |
| | 4.77091E-06 |
| | 7.29661E-08 |
| | 0.57692387 |
| | 0.326237154 |
| | 0.235466582 |
| | 0.489063144 |
| | 0.001626715 |
| | 0.989792151 |
| | 0.999861542 |
| | 0.613374177 |
| | 0.736618952 |
| | 0.311174396 |
| | 0.031331612 |
| | 0.978759363 |

| | |
|---|---|
| | 0.015611338 |
| | 0.001804238 |
| | 4.25119E-07 |
| | 0.968707031 |
| | 4.67404E-14 |
| | 0.99893861 |
| | 0.396277675 |
| | 0.254204139 |
| | 7.57527E-08 |
| | 0.094901874 |
| | 0.259474112 |
| | 0.019043506 |
| | 0.014635735 |
| | 0.000625785 |
| | 0.99999997 |
| | 0.001674596 |
| | 0.618360088 |
| | 6.50202E-12 |
| | 0.142528192 |
| | 0.998709763 |
| | 0.050010389 |
| | 4.65183E-14 |
| | 6.29385E-13 |
| | 0.979856468 |
| | 7.27307E-13 |
| | 0.536842935 |
| | 0.00014746 |
| | 0.003870107 |
| | 4.67404E-14 |
| | 7.88093E-08 |
| | 0.169283719 |
| | 0.652065252 |
| | 0.999999879 |
| | 0.000360672 |
| | 4.22243E-05 |

| |
|---|
| 0.849694487 |
| 0.977315937 |
| 0.350957922 |
| 0.84917732 |
| 0.68738726 |
| 0.836102784 |
| 0.989230925 |
| 0.896263807 |
| 0.225778993 |
| 0.001148907 |
| 0.24539571 |
| 0.712009262 |
| 1.9668E-08 |
| 0.251708134 |
| 0.947708088 |
| 0.965368188 |
| 1.57855E-10 |
| 0.007090968 |
| 0.021599116 |
| 0.997982799 |
| 2.49668E-10 |
| 0.15407073 |
| 0.00474106 |
| 0.999989384 |
| 0.821686178 |
| 0.999695138 |
| 2.75961E-05 |
| 0.070266521 |
| 0.992908585 |
| 0.055874222 |
| 0.269197751 |
| 0.412873838 |
| 0.980246428 |
| 0.999999291 |
| 0.062937005 |

| |
|---|
| 0.754608534 |
| 0.36902758 |
| 0.999997643 |
| 0.341201697 |
| 0.036555373 |
| 0.028774998 |
| 1.08842E-07 |
| 0.406109932 |
| 0.997982428 |
| 8.58547E-12 |
| 0.032598806 |
| 0.500107209 |
| 0.052185013 |
| 5.64658E-08 |
| 0.735105289 |
| 0.986793349 |
| 0.968967569 |
| 0.999323959 |
| 3.64526E-07 |
| 0.052483869 |
| 0.999892776 |
| 0.720495786 |
| 0.928849828 |
| 0.061309886 |
| 0.137994018 |
| 0.994008302 |
| 0.000247693 |
| 0.020467015 |
| 0.27536404 |
| 0.997841298 |
| 0.154345338 |
| 0.099521233 |
| 0.911823981 |
| 0.003596177 |
| 0.035165902 |

| | |
|---|---|
| | 0.000419117 |
| | 7.95556E-10 |
| | 3.54864E-07 |
| | 0.6448746 |
| | 3.9809E-10 |
| | 1.47764E-06 |
| | 0.005390609 |
| | 4.60743E-14 |
| | 0.01868978 |
| | 0.725147319 |
| | 3.65008E-12 |
| | 0.514887362 |
| | 0.999987635 |
| | 1.99213E-05 |
| | 0.455797728 |
| | 6.05726E-05 |
| | 0.024414714 |
| | 0.935116531 |
| | 6.51848E-06 |
| | 0.108232107 |
| | 0.00220413 |
| | 0.999408809 |
| | 0.99323963 |
| | 0.998638587 |
| | 1 |
| | 0.999847671 |
| | 1 |
| | 0.158506554 |
| | 0.615846736 |
| | 0.037413084 |
| | 0.098691642 |
| | 0.321534509 |
| | 0.995433922 |
| | 2.33193E-07 |
| | 0.944447621 |

| | |
|---|---|
| | 7.20975E-06 |
| | 0.999238187 |
| | 6.17209E-09 |
| | 0.023165542 |
| | 0.343390475 |
| | 0.543993219 |
| | 0.72857127 |
| | 3.88427E-08 |
| | 1 |
| | 0.037909031 |
| | 4.60743E-14 |
| | 2.84628E-12 |
| | 1.54772E-07 |
| | 0.400299865 |
| | 0.714949545 |
| | 0.76220727 |
| | 0.928705384 |
| | 0.680165316 |
| | 0.122076882 |
| | 0.150874581 |
| | 0.236665747 |
| | 0.480352339 |
| | 4.99205E-09 |
| | 0.090121046 |
| | 0.737691616 |
| | 0.671338172 |
| | 0.726195333 |
| | 0.964964168 |
| | 0.000446166 |
| | 0.46916585 |
| | 0.004874147 |
| | 0.983891228 |
| | 0.996605362 |
| | 0.016616484 |
| | 0.909567554 |

| | |
|---|---|
| | 0.627022098 |
| | 0.999981812 |
| | 0.015082065 |
| | 0.087031605 |
| | 0.855063361 |
| | 0.775558518 |
| | 0.383498535 |
| | 0.99370843 |
| | 0.357931543 |
| | 0.96768136 |
| | 0.999979617 |
| | 0.419934246 |
| | 0.999145421 |
| | 0.604105303 |
| | 8.62467E-06 |

| |
|---|
| P-value: (Bb_pos_12022024_Newanalysis) / (Water_pos_12022024_Newanalysis) |
| 0.11840703 |
| 0.012247997 |
| 0.999971637 |
| 0.859254047 |
| 0.452725191 |
| 0.19573896 |
| 5.94605E-07 |
| 0.28811344 |
| 0.616250343 |
| 0.897366228 |
| 0.999654706 |
| 9.46455E-11 |
| 0.388836719 |
| 0.864964333 |
| 0.999391637 |
| 0.03617144 |
| 0.999999999 |
| 0.998957292 |
| 0.991873322 |
| 0.019574989 |
| 0.361743789 |
| 0.028738396 |
| 0.99999977 |
| 0.05727081 |
| 0.001274598 |
| 0.979530049 |
| 0.983272604 |
| 0.828682418 |
| 0.00084606 |
| 0.013890989 |
| 0.999881352 |
| 0.016835495 |
| 0.89348488 |

| | |
|---|---|
| | 0.195434398 |
| | 0.30621409 |
| | 0.809456516 |
| | 0.001062888 |
| | 0.11797765 |
| | 0.162769765 |
| | 0.012209083 |
| | 0.553574999 |
| | 0.002339081 |
| | 0.526504635 |
| | 0.054378332 |
| | 0.003514362 |
| | 0.002732423 |
| | 0.000570569 |
| | 0.004851838 |
| | 0.043353202 |
| | 0.01264193 |
| | 0.087039406 |
| | 0.014873768 |
| | 0.000920153 |
| | 0.005832327 |
| | 0.686236483 |
| | 0.017832229 |
| | 0.636447808 |
| | 0.999911558 |
| | 0.480410179 |
| | 1.23098E-05 |
| | 0.03460508 |
| | 0.717584109 |
| | 3.68888E-07 |
| | 0.327470324 |
| | 0.320292183 |
| | 0.00370211 |
| | 3.26574E-09 |
| | 0.027282643 |

| | |
|---|---|
| | 0.003228629 |
| | 0.003910906 |
| | 2.9175E-08 |
| | 0.000678415 |
| | 0.001944402 |
| | 0.015229727 |
| | 0.000166837 |
| | 0.627404044 |
| | 0.017938524 |
| | 0.001657818 |
| | 1.28295E-05 |
| | 0.000405166 |
| | 0.019570079 |
| | 4.85945E-13 |
| | 0.009798716 |
| | 0.000392787 |
| | 0.007933055 |
| | 0.026871545 |
| | 0.054075923 |
| | 0.001043072 |
| | 0.011458376 |
| | 0.000153191 |
| | 0.376806793 |
| | 0.000395527 |
| | 0.001353377 |
| | 0.035149849 |
| | 0.342876701 |
| | 0.006552034 |
| | 7.40992E-05 |
| | 0.002205868 |
| | 0.000852111 |
| | 0.07021431 |
| | 0.969858701 |
| | 0.000292358 |
| | 0.002734768 |

| | |
|---|---|
| | 1.67289E-07 |
| | 0.663251887 |
| | 0.004786757 |
| | 0.003006358 |
| | 0.029683354 |
| | 0.998072286 |
| | 0.159860382 |
| | 0.016600114 |
| | 0.999507918 |
| | 0.002230607 |
| | 0.010487263 |
| | 0.001089925 |
| | 0.007027816 |
| | 0.139870683 |
| | 0.000681986 |
| | 0.738740199 |
| | 0.139002781 |
| | 0.032713362 |
| | 0.008659663 |
| | 0.415537007 |
| | 0.736512217 |
| | 0.001545757 |
| | 0.999823701 |
| | 0.008920405 |
| | 0.041862886 |
| | 0.000179923 |
| | 0.122282071 |
| | 0.110355977 |
| | 0.999389704 |
| | 0.46196173 |
| | 0.999998554 |
| | 0.000685764 |
| | 0.039557939 |
| | 0.002811158 |
| | 0.788289195 |

| | |
|---|---|
| | 0.184109038 |
| | 0.495514418 |
| | 0.080253407 |
| | 0.244056846 |
| | 0.998527926 |
| | 0.999880217 |
| | 0.95956 |
| | 0.999574267 |
| | 0.001832699 |
| | 0.747002004 |
| | 0.999993006 |
| | 0.90585789 |
| | 0.004472702 |
| | 0.957318715 |
| | 0.010311128 |
| | 0.035426098 |
| | 0.943758551 |
| | 0.419657455 |
| | 7.91902E-08 |
| | 0.973113524 |
| | 0.005638077 |
| | 0.053299429 |
| | 0.546072435 |
| | 0.588004095 |
| | 0.999997497 |
| | 0.004926384 |
| | 1.58706E-06 |
| | 0.107548974 |
| | 0.023417844 |
| | 0.039724982 |
| | 0.483033068 |
| | 0.021037232 |
| | 1.78021E-06 |
| | 0.999999464 |
| | 0.983928745 |

| | |
|---|---|
| | 0.000913501 |
| | 0.024700466 |
| | 0.278714483 |
| | 0.072119822 |
| | 0.965139865 |
| | 0.998103955 |
| | 0.997518462 |
| | 0.039185447 |
| | 1.06692E-13 |
| | 0.95183198 |
| | 0.01130029 |
| | 0.001438908 |
| | 0.00353075 |
| | 0.131947763 |
| | 0.002714105 |
| | 0.001747099 |
| | 0.09909997 |
| | 0.496765121 |
| | 0.213936069 |
| | 0.015554191 |
| | 0.897974773 |
| | 0.434868106 |
| | 7.1564E-06 |
| | 0.137421917 |
| | 3.83155E-06 |
| | 0.994046786 |
| | 0.863434238 |
| | 0.974131574 |
| | 0.006775634 |
| | 0.458554112 |
| | 0.005349503 |
| | 0.092180137 |
| | 0.58563661 |
| | 0.299757395 |
| | 0.834486178 |

| | |
|---|---|
| | 0.002113312 |
| | 0.032034392 |
| | 0.024776083 |
| | 0.460298716 |
| | 0.000140664 |
| | 0.081702431 |
| | 0.845898635 |
| | 0.917053692 |
| | 0.254999043 |
| | 0.023374499 |
| | 0.794975348 |
| | 0.193213758 |
| | 0.443356411 |
| | 0.799170933 |
| | 0.040244784 |
| | 0.00025929 |
| | 0.091645986 |
| | 0.879887279 |
| | 0.999976926 |
| | 0.975024862 |
| | 0.867037442 |
| | 0.978948405 |
| | 0.192680043 |
| | 0.288815403 |
| | 0.0072054 |
| | 0.613551906 |
| | 0.034694535 |
| | 0.421275353 |
| | 0.030686258 |
| | 0.714903993 |
| | 0.87436983 |
| | 0.845828882 |
| | 0.999963645 |
| | 0.608481939 |
| | 0.006881337 |

| |
|---|
| 0.00103089 |
| 0.031285802 |
| 0.999999692 |
| 0.999992715 |
| 0.974560073 |
| 0.154728233 |
| 0.781180789 |
| 0.961823221 |
| 0.110259503 |
| 0.730370803 |
| 2.81234E-09 |
| 0.057828452 |
| 0.999999571 |
| 3.52047E-06 |
| 1.79367E-07 |
| 5.32474E-05 |
| 0.001248576 |
| 0.03398447 |
| 0.746340755 |
| 0.966677681 |
| 0.043320355 |
| 0.041544379 |
| 0.081931851 |
| 0.911706152 |
| 0.514110152 |
| 0.64409141 |
| 0.904978575 |
| 0.999468609 |
| 0.00275846 |
| 0.977450744 |
| 0.999941578 |
| 0.006700548 |
| 0.000154884 |
| 0.998858145 |
| 0.885549655 |

| | |
|---|---|
| | 0.000131306 |
| | 0.266392722 |
| | 0.006627046 |
| | 0.981155975 |
| | 0.999995883 |
| | 0.973723379 |
| | 0.599461261 |
| | 0.508411351 |
| | 0.001798199 |
| | 0.017375359 |
| | 0.102489532 |
| | 0.04819147 |
| | 0.131977132 |
| | 4.50347E-08 |
| | 0.089591404 |
| | 0.999726057 |
| | 0.089863759 |
| | 0.013947526 |
| | 0.458352765 |
| | 0.765090136 |
| | 0.803024086 |
| | 0.000525257 |
| | 0.003152647 |
| | 0.006442221 |
| | 0.17260323 |
| | 0.025031402 |
| | 0.993857797 |
| | 0.369761936 |
| | 0.000224562 |
| | 8.88467E-07 |
| | 0.999999828 |
| | 0.72143609 |
| | 0.000161592 |
| | 0.000341678 |
| | 0.449545805 |

| | |
|---|---|
| | 0.021399736 |
| | 0.000874849 |
| | 0.975694644 |
| | 0.008295314 |
| | 1.29356E-07 |
| | 0.030866808 |
| | 0.016817275 |
| | 0.013376924 |
| | 0.001036492 |
| | 0.427990918 |
| | 0.685952203 |
| | 3.5322E-08 |
| | 0.993184297 |
| | 0.548909506 |
| | 0.984220702 |
| | 0.006166919 |
| | 0.024374605 |
| | 0.99999988 |
| | 0.709060291 |
| | 7.46471E-06 |
| | 0.056250898 |
| | 6.28438E-08 |
| | 0.999981239 |
| | 0.595167486 |
| | 0.711194641 |
| | 0.013726757 |
| | 0.050222885 |
| | 6.25441E-11 |
| | 0.987232404 |
| | 9.96328E-08 |
| | 2.12736E-05 |
| | 0.013337612 |
| | 4.34477E-07 |
| | 4.96115E-11 |
| | 0.000383576 |

| | |
|---|---|
| | 0.002024064 |
| | 0.90431215 |
| | 4.84629E-11 |
| | 0.000137472 |
| | 0.304796589 |
| | 0.414845022 |
| | 0.013443764 |
| | 0.012338334 |
| | 0.032094782 |
| | 0.224758515 |
| | 2.34297E-09 |
| | 0.321725019 |
| | 0.154013122 |
| | 1.2708E-09 |
| | 0.013902028 |
| | 0.0722624 |
| | 0.071271669 |
| | 0.19220092 |
| | 0.997759144 |
| | 0.068601768 |
| | 0.086944565 |
| | 0.038510633 |
| | 0.980383097 |
| | 0.403906727 |
| | 0.484146315 |
| | 0.616723026 |
| | 0.767265489 |
| | 0.98889624 |
| | 0.023911425 |
| | 0.000263172 |
| | 0.430814096 |
| | 0.827996605 |
| | 0.571569439 |
| | 0.025273167 |
| | 0.748503103 |

| | |
|---|---|
| | 0.348511237 |
| | 0.000106903 |
| | 1.15429E-08 |
| | 0.465429529 |
| | 0.491923944 |
| | 0.025633515 |
| | 0.602432687 |
| | 0.796016894 |
| | 3.30823E-07 |
| | 1.01589E-05 |
| | 0.353140281 |
| | 0.081454285 |
| | 0.025241329 |
| | 0.975764782 |
| | 0.999818954 |
| | 0.981608289 |
| | 0.461673317 |
| | 0.990654664 |
| | 0.221549342 |
| | 0.223598952 |
| | 4.85167E-14 |
| | 0.157114055 |
| | 0.068440459 |
| | 0.016838253 |
| | 0.003871322 |
| | 0.996899692 |
| | 1.80732E-06 |
| | 0.113167037 |
| | 0.999975068 |
| | 0.198325195 |
| | 0.055150378 |
| | 2.67453E-13 |
| | 0.309983002 |
| | 0.980519161 |
| | 0.92085665 |

| | |
|---|---|
| | 0.416907817 |
| | 0.106418237 |
| | 0.031548725 |
| | 0.945493417 |
| | 0.016704901 |
| | 0.877831588 |
| | 0.001838743 |
| | 0.028221323 |
| | 0.542514286 |
| | 3.23543E-10 |
| | 0.060976132 |
| | 0.996641394 |
| | 0.999995299 |
| | 0.836356137 |
| | 1.10965E-10 |
| | 0.000108026 |
| | 3.46512E-12 |
| | 0.028009441 |
| | 0.713479745 |
| | 0.239547527 |
| | 0.383511974 |
| | 0.152536018 |
| | 0.999665369 |
| | 0.902258728 |
| | 0.099786602 |
| | 0.983553747 |
| | 0.619809023 |
| | 0.999992153 |
| | 0.854876749 |
| | 2.17338E-05 |
| | 0.013190119 |
| | 0.000551359 |
| | 0.04509763 |
| | 0.99998928 |
| | 0.656254076 |

| | |
|---|---|
| | 0.999751658 |
| | 0.998215482 |
| | 0.040812107 |
| | 0.9610184 |
| | 0.941925552 |
| | 1.40784E-06 |
| | 0.999706961 |
| | 0.999991908 |
| | 0.994863074 |
| | 2.31871E-07 |
| | 0.014305554 |
| | 9.08303E-09 |
| | 4.60743E-14 |
| | 1.27865E-08 |
| | 1.30373E-10 |
| | 4.60743E-14 |
| | 4.60743E-14 |
| | 6.28086E-12 |
| | 0.148365416 |
| | 0.00035558 |
| | 0.743649516 |
| | 0.999983348 |
| | 3.16349E-05 |
| | 5.30798E-13 |
| | 5.89528E-14 |
| | 5.76206E-14 |
| | 4.68836E-12 |
| | 4.37624E-07 |
| | 0.378279957 |
| | 0.003600318 |
| | 0.069565027 |
| | 2.21345E-12 |
| | 2.85438E-13 |
| | 4.62963E-14 |
| | 0.004298348 |

| | |
|---|---|
| | 0.446695259 |
| | 0.789291354 |
| | 0.174421207 |
| | 0.999428648 |
| | 8.70348E-05 |
| | 0.952404702 |
| | 7.11398E-05 |
| | 0.999999986 |
| | 0.957453818 |
| | 0.94497764 |
| | 0.999992933 |
| | 0.582847347 |
| | 0.883195245 |
| | 0.999155944 |
| | 0.956438969 |
| | 0.999998115 |
| | 2.17138E-10 |
| | 0.038123799 |
| | 7.10585E-08 |
| | 0.693412632 |
| | 0.190481291 |
| | 0.847777172 |
| | 0.992368572 |
| | 0.952968152 |
| | 0.999999742 |
| | 0.020771906 |
| | 0.000897168 |
| | 0.027635617 |
| | 0.999512573 |
| | 0.999820536 |
| | 0.741462716 |
| | 0.843995054 |
| | 0.39598439 |
| | 0.000376326 |
| | 0.647279345 |

| | |
|---|---|
| | 0.991489737 |
| | 0.932900232 |
| | 0.861556569 |
| | 0.000212884 |
| | 0.672212756 |
| | 0.823152528 |
| | 0.941826784 |
| | 0.244945905 |
| | 0.078913979 |
| | 0.999985058 |
| | 0.753261539 |
| | 0.579772368 |
| | 0.010858137 |
| | 1.13999E-05 |
| | 0.943300463 |
| | 0.096738986 |
| | 0.999994096 |
| | 0.946942374 |
| | 0.65824949 |
| | 0.00072487 |
| | 0.03889719 |
| | 1.022E-06 |
| | 0.898022775 |
| | 0.660646087 |
| | 0.999999904 |
| | 0.000249257 |
| | 0.996493715 |
| | 0.15700725 |
| | 0.197446251 |
| | 0.115261784 |
| | 0.997716372 |
| | 0.397154463 |
| | 0.4719131 |
| | 8.3944E-13 |
| | 1.66604E-06 |

| | |
|---|---|
| | 7.16094E-14 |
| | 0.962027457 |
| | 5.07079E-10 |
| | 0.999992448 |
| | 0.258732188 |
| | 0.944044444 |
| | 0.6017807 |
| | 0.962621935 |
| | 0.444199331 |
| | 0.838441842 |
| | 0.083146005 |
| | 0.999985689 |
| | 0.950832432 |
| | 0.461055872 |
| | 0.999390409 |
| | 0.021280378 |
| | 8.52415E-08 |
| | 0.057439153 |
| | 0.012807565 |
| | 0.07735793 |
| | 0.550752838 |
| | 0.031703624 |
| | 0.991494949 |
| | 0.78695487 |
| | 1.28441E-07 |
| | 1.64269E-05 |
| | 0.998895904 |
| | 0.037033573 |
| | 8.98823E-05 |
| | 0.907362483 |
| | 0.136483797 |
| | 0.139894741 |
| | 0.935112907 |
| | 1.99377E-09 |
| | 0.864328551 |

| | |
|---|---|
| | 0.128264578 |
| | 0.999745463 |
| | 0.999999996 |
| | 0.999196899 |
| | 0.999696362 |
| | 0.000144434 |
| | 0.001038037 |
| | 0.081197311 |
| | 0.969941863 |
| | 0.994919745 |
| | 0.99999973 |
| | 1.54375E-08 |
| | 0.240560824 |
| | 0.001693863 |
| | 0.213926872 |
| | 0.290726571 |
| | 2.62909E-09 |
| | 0.152479538 |
| | 0.805035332 |
| | 1.05658E-08 |
| | 0.131900387 |
| | 0.890789402 |
| | 0.999150584 |
| | 0.400688321 |
| | 1 |
| | 0.156285538 |
| | 0.970060351 |
| | 0.924197209 |
| | 0.001316811 |
| | 0.78070318 |
| | 0.394553061 |
| | 0.712908539 |
| | 0.999999915 |
| | 0.99999643 |
| | 3.92302E-06 |

| | |
|---|---|
| | 0.010147287 |
| | 0.0001357 |
| | 0.002995682 |
| | 0.008518225 |
| | 0.020516041 |
| | 9.95703E-05 |
| | 0.000862603 |
| | 2.61769E-07 |
| | 0.00043195 |
| | 0.013372055 |
| | 0.972630454 |
| | 0.027174246 |
| | 0.002491765 |
| | 0.000309056 |
| | 0.685646275 |
| | 1.82824E-05 |
| | 0.085160374 |
| | 0.44282237 |
| | 0.035268624 |
| | 0.024289352 |
| | 0.685412056 |
| | 0.09958539 |
| | 0.079950643 |
| | 0.982427547 |
| | 0.122107888 |
| | 0.07403587 |
| | 0.094587404 |
| | 0.182616537 |
| | 0.195097451 |
| | 0.581988598 |
| | 0.000533369 |
| | 2.28213E-05 |
| | 0.004900648 |
| | 1.34467E-07 |
| | 0.00427582 |

| | |
|---|---|
| | 1.46352E-05 |
| | 0.019694258 |
| | 0.229710708 |
| | 0.999478239 |
| | 0.097734229 |
| | 0.005417782 |
| | 0.006901034 |
| | 0.002035143 |
| | 0.999993912 |
| | 0.091426609 |
| | 0.097564545 |
| | 0.001009233 |
| | 0.001235144 |
| | 0.326169815 |
| | 0.916830587 |
| | 0.523524269 |
| | 0.132827197 |
| | 0.991890138 |
| | 0.998820865 |
| | 0.57086259 |
| | 0.006093579 |
| | 0.956022096 |
| | 5.71765E-14 |
| | 0.044634504 |
| | 0.507160833 |
| | 0.185999263 |
| | 1.59465E-09 |
| | 0.958677555 |
| | 0.999976687 |
| | 0.631209657 |
| | 0.999051013 |
| | 0.966955258 |
| | 0.998919375 |
| | 0.031965068 |
| | 0.411636429 |

| |
|---|
| 0.788575425 |
| 0.000516594 |
| 0.999987395 |
| 0.84713454 |
| 0.524145731 |
| 0.661111866 |
| 0.296673819 |
| 0.951008153 |
| 0.401183641 |
| 0.259399161 |
| 0.557885655 |
| 0.999826743 |
| 0.998907594 |
| 0.537500264 |
| 0.999997325 |

| |
|---|
| P-value: (CM_pos_12022024_Newanalysis) / (Water_pos_12022024_Newanalysis) |
| 0.651628913 |
| 0.296072658 |
| 0.999999608 |
| 0.130794982 |
| 0.90400668 |
| 0.447232801 |
| 1.40893E-05 |
| 0.44653527 |
| 0.161520721 |
| 0.571435565 |
| 0.937759245 |
| 1.32264E-09 |
| 0.99999993 |
| 0.085865147 |
| 0.996481581 |
| 0.982310834 |
| 0.785591748 |
| 4.80727E-14 |
| 0.825316802 |
| 6.31705E-08 |
| 0.998318752 |
| 8.70984E-08 |
| 5.3042E-05 |
| 2.05822E-10 |
| 8.16014E-14 |
| 0.998938754 |
| 0.762422455 |
| 0.40959597 |
| 0.251330904 |
| 0.045064043 |
| 0.982026794 |
| 0.000369892 |
| 0.062117167 |

| |
|---|
| 0.988362374 |
| 1.28709E-11 |
| 0.685590631 |
| 0.821108677 |
| 0.324271305 |
| 0.155277362 |
| 0.000165133 |
| 0.310851807 |
| 3.14868E-05 |
| 0.124260091 |
| 0.999915521 |
| 0.000760824 |
| 4.40096E-06 |
| 0.005409841 |
| 0.00193808 |
| 0.85187323 |
| 0.685380285 |
| 0.213458252 |
| 0.048900153 |
| 1.64081E-05 |
| 0.860977933 |
| 0.121102788 |
| 0.403903475 |
| 0.222448826 |
| 0.497582495 |
| 0.965441705 |
| 0.051543709 |
| 0.371144873 |
| 0.581630522 |
| 0.000227311 |
| 2.45312E-06 |
| 0.138061505 |
| 0.001254042 |
| 0.086787687 |
| 0.58135463 |

| |
|---|
| 0.06279326 |
| 0.090454922 |
| 0.879241547 |
| 0.000165219 |
| 0.992134812 |
| 0.997355469 |
| 0.000532614 |
| 0.999993899 |
| 2.69521E-11 |
| 0.009535106 |
| 0.000368386 |
| 0.02545749 |
| 0.268841438 |
| 0.999805008 |
| 0.00623993 |
| 0.972172499 |
| 0.90744261 |
| 1 |
| 0.00922299 |
| 0.029092002 |
| 0.984696876 |
| 0.140193752 |
| 0.003387635 |
| 0.002337693 |
| 0.467623209 |
| 0.062739887 |
| 0.024405237 |
| 0.997238781 |
| 0.006414479 |
| 0.902383065 |
| 0.001357936 |
| 0.984905098 |
| 0.154899155 |
| 0.026021643 |
| 0.980435167 |

| | |
|---|---|
| | 6.75179E-07 |
| | 0.135976709 |
| | 0.000494792 |
| | 0.636377055 |
| | 0.078190825 |
| | 0.994943448 |
| | 0.171296325 |
| | 0.991706856 |
| | 0.864621525 |
| | 0.72347187 |
| | 0.995801585 |
| | 0.00023854 |
| | 0.574369278 |
| | 0.990858083 |
| | 4.76286E-14 |
| | 0.023727542 |
| | 0.019267654 |
| | 0.004862516 |
| | 0.006627339 |
| | 0.26837449 |
| | 0.90641312 |
| | 0.006805931 |
| | 0.98589632 |
| | 0.857821468 |
| | 0.257744823 |
| | 0.000322294 |
| | 0.068346496 |
| | 0.018689692 |
| | 0.093596239 |
| | 1.13976E-10 |
| | 0.0129473 |
| | 1.56849E-08 |
| | 0.957134174 |
| | 0.999999991 |
| | 0.987582177 |

| |
|---|
| 0.265575766 |
| 0.039487831 |
| 0.900117908 |
| 0.787276275 |
| 0.946662995 |
| 0.99999979 |
| 0.999622663 |
| 0.870291168 |
| 1 |
| 0.702448864 |
| 0.360729746 |
| 0.594479826 |
| 0.028523239 |
| 0.202714995 |
| 0.417638209 |
| 4.25499E-05 |
| 0.922390118 |
| 0.998133178 |
| 0.000226023 |
| 0.19138944 |
| 0.303358045 |
| 0.863537791 |
| 0.999127321 |
| 0.00173576 |
| 0.99999937 |
| 0.971945184 |
| 0.001073861 |
| 0.9049229 |
| 0.466461561 |
| 0.217286717 |
| 0.00260601 |
| 0.945107879 |
| 0.001117174 |
| 0.756400839 |
| 0.015765332 |

| |
|---|
| 0.635765728 |
| 0.338590875 |
| 0.23383449 |
| 2.05133E-09 |
| 0.002490344 |
| 0.791258721 |
| 0.155547937 |
| 0.238349033 |
| 3.6218E-11 |
| 0.999570109 |
| 0.62760315 |
| 4.88659E-05 |
| 1.32034E-06 |
| 5.53624E-07 |
| 0.398895113 |
| 6.90996E-06 |
| 0.038458894 |
| 0.972318904 |
| 1.17976E-11 |
| 0.996732323 |
| 0.203864827 |
| 0.397733806 |
| 0.009593982 |
| 0.009098143 |
| 0.018389654 |
| 0.872078359 |
| 0.67897426 |
| 0.095582608 |
| 0.976062462 |
| 0.013397487 |
| 0.537547238 |
| 0.781806724 |
| 0.000267607 |
| 0.00118193 |
| 0.009441931 |

| | |
|---|---|
| | 0.351019138 |
| | 0.07450583 |
| | 0.464903807 |
| | 0.999918453 |
| | 0.056221577 |
| | 0.116919014 |
| | 0.99224535 |
| | 0.971684939 |
| | 0.127868426 |
| | 0.385117997 |
| | 0.913380507 |
| | 0.960901659 |
| | 0.977648845 |
| | 0.483590818 |
| | 0.99997324 |
| | 0.999969505 |
| | 0.111282457 |
| | 0.000817433 |
| | 4.04581E-05 |
| | 0.999998536 |
| | 0.054798667 |
| | 0.999655501 |
| | 0.546925275 |
| | 0.973397925 |
| | 0.4183243 |
| | 0.838126579 |
| | 0.032101895 |
| | 0.788640349 |
| | 0.999925736 |
| | 0.71632883 |
| | 0.999694072 |
| | 0.405244546 |
| | 0.427773425 |
| | 0.368810985 |
| | 0.027355379 |

| | |
|---|---|
| | 0.387382545 |
| | 0.017567162 |
| | 0.999966918 |
| | 0.798797544 |
| | 0.88291663 |
| | 0.821501742 |
| | 0.005512128 |
| | 0.950101109 |
| | 0.930684751 |
| | 7.43392E-06 |
| | 4.08177E-05 |
| | 0.566792277 |
| | 0.79022139 |
| | 1.1739E-10 |
| | 0.324868014 |
| | 0.5493959 |
| | 0.999959246 |
| | 0.952051034 |
| | 0.911448132 |
| | 3.1094E-12 |
| | 0.788621447 |
| | 0.980968422 |
| | 0.975154473 |
| | 0.706387778 |
| | 0.34967219 |
| | 0.289069089 |
| | 0.214551679 |
| | 0.762175251 |
| | 0.990460295 |
| | 1 |
| | 0.085851477 |
| | 0.001571483 |
| | 0.056077459 |
| | 0.999974562 |
| | 1.54432E-13 |

| | |
|---|---|
| | 3.37641E-06 |
| | 0.000120426 |
| | 0.803107166 |
| | 0.999999343 |
| | 1 |
| | 0.283314732 |
| | 0.00085105 |
| | 0.374079023 |
| | 0.051860207 |
| | 0.094377431 |
| | 0.989167817 |
| | 0.010542483 |
| | 0.030441595 |
| | 4.92583E-11 |
| | 0.483800122 |
| | 0.957827416 |
| | 0.546061806 |
| | 0.000940985 |
| | 0.08093905 |
| | 0.683046016 |
| | 8.1716E-09 |
| | 0.968881515 |
| | 0.004494573 |
| | 0.993755502 |
| | 0.078734828 |
| | 0.023588421 |
| | 0.993432759 |
| | 0.00148222 |
| | 7.99067E-10 |
| | 1.72473E-12 |
| | 0.162427762 |
| | 0.000308966 |
| | 0.735942882 |
| | 1.6929E-07 |
| | 0.949884532 |

| | |
|---|---|
| | 3.66174E-06 |
| | 0.047980036 |
| | 0.000275046 |
| | 0.014198858 |
| | 7.47965E-07 |
| | 0.357383645 |
| | 1.01149E-06 |
| | 0.003730803 |
| | 0.002192473 |
| | 0.171375726 |
| | 0.023771565 |
| | 3.45611E-06 |
| | 0.001415926 |
| | 0.24720316 |
| | 0.638402521 |
| | 0.025685351 |
| | 0.015341296 |
| | 7.3011E-06 |
| | 4.03175E-07 |
| | 0.381077781 |
| | 1.3827E-05 |
| | 0.003193286 |
| | 0.297406657 |
| | 0.00116031 |
| | 0.932913234 |
| | 0.639100422 |
| | 0.999999999 |
| | 0.053537258 |
| | 6.36402E-09 |
| | 0.433070946 |
| | 0.996949588 |
| | 0.812209169 |
| | 0.652690643 |
| | 0.81687763 |
| | 0.999995187 |

| | |
|---|---|
| | 0.011505752 |
| | 0.925662463 |
| | 0.596940448 |
| | 0.594902775 |
| | 0.204009474 |
| | 0.254764567 |
| | 0.002332808 |
| | 0.504467362 |
| | 0.06543458 |
| | 4.47226E-05 |
| | 0.000385222 |
| | 0.99999999 |
| | 0.11347137 |
| | 3.81023E-05 |
| | 0.008109829 |
| | 0.999991432 |
| | 0.957349272 |
| | 0.064471685 |
| | 0.003939014 |
| | 0.92497309 |
| | 1.7477E-09 |
| | 0.999974553 |
| | 0.986783821 |
| | 0.389439116 |
| | 1.3095E-07 |
| | 0.883325441 |
| | 0.007336981 |
| | 0.493493985 |
| | 0.34516662 |
| | 0.527165372 |
| | 0.737728038 |
| | 0.057987262 |
| | 0.130495506 |
| | 0.248875519 |
| | 0.373144106 |

| |
|---|
| 0.816367969 |
| 0.000650515 |
| 9.08273E-06 |
| 0.713874913 |
| 0.014192055 |
| 0.022952049 |
| 0.033695171 |
| 0.316691971 |
| 7.4361E-11 |
| 0.025035946 |
| 0.082748813 |
| 0.221413699 |
| 0.017984044 |
| 0.000919108 |
| 0.016502117 |
| 0.137021668 |
| 0.648087064 |
| 1.01301E-07 |
| 2.46965E-05 |
| 0.999390073 |
| 3.02092E-13 |
| 0.698236058 |
| 0.328605985 |
| 0.00677379 |
| 2.62344E-06 |
| 0.948363137 |
| 4.49207E-05 |
| 1.06314E-05 |
| 0.99916265 |
| 0.999998622 |
| 0.000421756 |
| 0.006974961 |
| 9.32566E-06 |
| 5.98226E-06 |
| 0.46678797 |

| | |
|---|---|
| | 9.12588E-06 |
| | 0.999999542 |
| | 0.018459081 |
| | 0.003857167 |
| | 0.126539832 |
| | 0.338038255 |
| | 0.863657079 |
| | 0.007495593 |
| | 0.905355196 |
| | 1.93054E-09 |
| | 0.15969374 |
| | 0.000154901 |
| | 5.45727E-06 |
| | 0.000453376 |
| | 1.78575E-08 |
| | 1.30223E-05 |
| | 1.62761E-08 |
| | 0.264134552 |
| | 0.000502008 |
| | 6.7909E-12 |
| | 0.041716906 |
| | 7.15314E-07 |
| | 0.005325443 |
| | 9.79182E-06 |
| | 0.209628242 |
| | 7.03898E-06 |
| | 0.64155508 |
| | 4.209E-06 |
| | 0.807174514 |
| | 0.009510832 |
| | 0.921783869 |
| | 0.001092626 |
| | 3.51892E-05 |
| | 2.62162E-06 |
| | 1.54779E-09 |

| | |
|---|---|
| | 2.92524E-06 |
| | 0.07843088 |
| | 0.910528953 |
| | 0.748381939 |
| | 0.012827742 |
| | 7.98954E-06 |
| | 2.8549E-05 |
| | 0.009270416 |
| | 0.483739823 |
| | 0.002039808 |
| | 0.013273582 |
| | 0.000337945 |
| | 5.41809E-08 |
| | 0.943961625 |
| | 0.012104989 |
| | 0.995443124 |
| | 0.098408646 |
| | 0.036852031 |
| | 0.97863554 |
| | 0.0001662 |
| | 0.926155409 |
| | 0.999999927 |
| | 0.457869747 |
| | 4.42558E-11 |
| | 5.73056E-08 |
| | 0.996614133 |
| | 0.560541976 |
| | 6.13506E-08 |
| | 0.99063534 |
| | 0.002294815 |
| | 0.744147794 |
| | 0.007150684 |
| | 0.00115957 |
| | 0.999226693 |
| | 0.961261171 |

| | |
|---|---|
| | 0.250281304 |
| | 0.008737621 |
| | 0.994350842 |
| | 0.442662698 |
| | 0.998515963 |
| | 0.989765017 |
| | 0.003875848 |
| | 0.771622756 |
| | 0.321547317 |
| | 0.07919571 |
| | 0.999998414 |
| | 0.394469857 |
| | 0.868405973 |
| | 0.931867552 |
| | 0.999999912 |
| | 4.03782E-05 |
| | 0.656288931 |
| | 0.942949516 |
| | 0.286047399 |
| | 0.926754038 |
| | 0.998991335 |
| | 0.999980635 |
| | 0.812961191 |
| | 0.434543003 |
| | 0.951131839 |
| | 0.116029417 |
| | 0.114621884 |
| | 0.11947828 |
| | 0.999455852 |
| | 0.999995988 |
| | 0.170645035 |
| | 0.014206303 |
| | 0.561734078 |
| | 0.999852424 |
| | 0.459185325 |

| | |
|---|---|
| | 0.895585975 |
| | 0.458501623 |
| | 0.821553752 |
| | 0.153484417 |
| | 0.31312359 |
| | 0.875897798 |
| | 0.050296075 |
| | 0.013680455 |
| | 1.04798E-09 |
| | 0.000490766 |
| | 0.982710792 |
| | 0.996234182 |
| | 0.25167098 |
| | 0.000375704 |
| | 0.950408899 |
| | 0.541421265 |
| | 0.371562105 |
| | 0.000843938 |
| | 0.987001666 |
| | 0.233053976 |
| | 0.00021118 |
| | 1.68421E-13 |
| | 1.81053E-06 |
| | 0.957237394 |
| | 0.828350994 |
| | 0.000465099 |
| | 3.41471E-07 |
| | 0.488486132 |
| | 0.420305813 |
| | 0.004195885 |
| | 0.809818086 |
| | 0.999968509 |
| | 0.730432045 |
| | 0.999747674 |
| | 2.26537E-05 |

| | |
|---|---|
| | 0.000446385 |
| | 0.803413034 |
| | 0.972080843 |
| | 0.116044975 |
| | 0.984055793 |
| | 0.951592303 |
| | 0.999829613 |
| | 0.194830942 |
| | 0.876408398 |
| | 0.956262973 |
| | 0.116385528 |
| | 0.570767949 |
| | 0.981922968 |
| | 0.429539717 |
| | 0.542672838 |
| | 0.999915149 |
| | 8.19904E-09 |
| | 0.71486304 |
| | 0.986966396 |
| | 0.028485001 |
| | 0.809693811 |
| | 0.547218395 |
| | 0.188444974 |
| | 0.44160984 |
| | 1 |
| | 0.533113888 |
| | 0.497746962 |
| | 0.504409706 |
| | 0.999733919 |
| | 0.976942983 |
| | 0.003640892 |
| | 0.021007836 |
| | 0.794670574 |
| | 3.20281E-09 |
| | 0.480265277 |

| | |
|---|---|
| | 0.910383696 |
| | 0.999845194 |
| | 0.992579484 |
| | 0.99997122 |
| | 0.999991202 |
| | 0.038851601 |
| | 1.50028E-09 |
| | 0.469524603 |
| | 5.54631E-05 |
| | 0.043333097 |
| | 0.814292284 |
| | 0.269003843 |
| | 0.719220685 |
| | 2.36113E-06 |
| | 4.65673E-10 |
| | 0.978585255 |
| | 0.974909942 |
| | 0.060836727 |
| | 6.53697E-05 |
| | 0.994567066 |
| | 0.083765455 |
| | 0.96456028 |
| | 1 |
| | 0.626156802 |
| | 0.965329336 |
| | 0.028398362 |
| | 0.039065624 |
| | 0.00408162 |
| | 0.129079474 |
| | 0.809384306 |
| | 0.001557761 |
| | 0.997804384 |
| | 0.99950877 |
| | 0.999679824 |
| | 1.81542E-06 |

| | |
|---|---|
| | 0.092338795 |
| | 0.043514336 |
| | 0.00149011 |
| | 0.974715484 |
| | 0.0032821 |
| | 0.930898919 |
| | 0.779899942 |
| | 0.998889636 |
| | 0.012214638 |
| | 0.999002725 |
| | 0.917430547 |
| | 0.005943735 |
| | 0.778418777 |
| | 8.37735E-05 |
| | 0.159155493 |
| | 1.40058E-05 |
| | 0.177680674 |
| | 0.956351414 |
| | 0.592073281 |
| | 0.298693182 |
| | 0.083165769 |
| | 0.047134814 |
| | 0.046102699 |
| | 0.965122895 |
| | 0.087223619 |
| | 0.017654303 |
| | 0.051194943 |
| | 0.064038281 |
| | 0.730594395 |
| | 0.979993636 |
| | 0.000533271 |
| | 5.2652E-05 |
| | 0.999817895 |
| | 0.000104923 |
| | 0.99564123 |

| |
|---|
| 3.00941E-05 |
| 0.004989612 |
| 0.118021752 |
| 0.995325337 |
| 0.882726763 |
| 0.025469672 |
| 0.247951819 |
| 0.041072367 |
| 0.573656679 |
| 0.001086592 |
| 5.33695E-12 |
| 2.68837E-05 |
| 0.005352992 |
| 0.997967668 |
| 0.936256719 |
| 0.114466597 |
| 0.39225049 |
| 0.99989167 |
| 0.99989857 |
| 0.722898728 |
| 0.9906965 |
| 0.106477736 |
| 1.64609E-11 |
| 0.0207552 |
| 0.069396571 |
| 0.931468178 |
| 0.00106267 |
| 0.227694448 |
| 0.999998548 |
| 0.992553125 |
| 0.99204313 |
| 0.653147015 |
| 0.415228598 |
| 0.005968057 |
| 0.902374145 |

| |
|---|
| 0.999947138 |
| 0.013911121 |
| 0.012660601 |
| 0.232338711 |
| 0.024867209 |
| 0.003461794 |
| 0.426542122 |
| 0.999998946 |
| 0.167125199 |
| 0.944026869 |
| 0.717390178 |
| 0.500031791 |
| 0.231874978 |
| 7.58027E-05 |
| 0.042513634 |

| |
|---|
| P-value: (Rabbitserum_12022024_Newanalysis) / (Water_pos_12022024_Newanalysis) |
| 0.999752526 |
| 0.062918716 |
| 2.38945E-09 |
| 0.001921105 |
| 0.007841047 |
| 0.856024227 |
| 0.026446179 |
| 0.002468685 |
| 0.350736766 |
| 0.942757214 |
| 1 |
| 8.5445E-08 |
| 0.266709456 |
| 0.031943621 |
| 0.044361446 |
| 0.774556209 |
| 0.709690464 |
| 4.60743E-14 |
| 4.31794E-06 |
| 4.94049E-14 |
| 0.707839108 |
| 1.71585E-12 |
| 6.02297E-06 |
| 4.60743E-14 |
| 4.60743E-14 |
| 0.879947306 |
| 0.658020342 |
| 0.102537849 |
| 0.974457665 |
| 0.003538393 |
| 0.209383651 |
| 0.001677782 |
| 0.001819128 |

| |
|---|
| 0.988026899 |
| 5.65104E-14 |
| 1.54982E-08 |
| 9.12746E-07 |
| 0.554507748 |
| 0.461473047 |
| 7.74022E-07 |
| 0.167395691 |
| 0.827830851 |
| 0.014590673 |
| 3.00845E-08 |
| 3.20013E-05 |
| 2.98482E-10 |
| 1.33663E-06 |
| 8.98142E-07 |
| 0.113762483 |
| 0.053497793 |
| 0.019366571 |
| 0.003089977 |
| 2.21521E-08 |
| 0.012161766 |
| 0.986562976 |
| 0.658196671 |
| 2.12675E-08 |
| 0.289853378 |
| 0.156780192 |
| 7.91912E-08 |
| 0.999979996 |
| 0.885431373 |
| 0.001411109 |
| 9.27067E-09 |
| 0.003280789 |
| 0.008737301 |
| 0.999063341 |
| 0.99567219 |

| | |
|---|---|
| | 0.347665192 |
| | 0.830197756 |
| | 0.020541022 |
| | 0.000557592 |
| | 0.08422655 |
| | 0.335519784 |
| | 0.996452028 |
| | 0.999990959 |
| | 5.20695E-14 |
| | 1.88902E-09 |
| | 5.65216E-09 |
| | 1.76581E-12 |
| | 0.001404126 |
| | 0.001522635 |
| | 3.79151E-05 |
| | 0.038846941 |
| | 6.16003E-09 |
| | 0.999192755 |
| | 0.000160237 |
| | 0.243856833 |
| | 0.997459121 |
| | 0.934888884 |
| | 0.000169765 |
| | 5.49579E-08 |
| | 4.62645E-05 |
| | 3.07174E-05 |
| | 5.21594E-07 |
| | 0.139596863 |
| | 1.23434E-05 |
| | 0.003039498 |
| | 0.00766702 |
| | 0.981444301 |
| | 1.18189E-09 |
| | 1.08975E-05 |
| | 0.256677065 |

| | |
|---|---|
| | 8.17151E-09 |
| | 9.34255E-05 |
| | 2.65602E-07 |
| | 0.980324875 |
| | 0.470198768 |
| | 0.29905993 |
| | 0.006800535 |
| | 0.933761829 |
| | 1.33992E-05 |
| | 0.949152604 |
| | 0.802404526 |
| | 3.20261E-05 |
| | 0.302293118 |
| | 0.71452894 |
| | 4.60743E-14 |
| | 0.004081932 |
| | 0.597150679 |
| | 0.02036357 |
| | 4.67404E-14 |
| | 0.87889566 |
| | 0.00505904 |
| | 0.33280476 |
| | 0.453308356 |
| | 0.999947024 |
| | 0.988429942 |
| | 0.999988391 |
| | 0.006030178 |
| | 0.292139588 |
| | 6.57326E-07 |
| | 4.65183E-14 |
| | 1.82092E-08 |
| | 1.61459E-06 |
| | 0.999216225 |
| | 0.400851026 |
| | 0.938883471 |

| | |
|---|---|
| | 0.517022608 |
| | 0.999445743 |
| | 0.987059814 |
| | 0.769205911 |
| | 0.548859236 |
| | 0.743387911 |
| | 0.990500919 |
| | 0.944582414 |
| | 0.366060676 |
| | 0.998775051 |
| | 0.341982211 |
| | 2.95685E-05 |
| | 4.67404E-14 |
| | 0.165096106 |
| | 2.0138E-10 |
| | 5.56222E-14 |
| | 0.996609162 |
| | 0.025023367 |
| | 0.346012104 |
| | 0.021633329 |
| | 0.191524363 |
| | 1.63325E-05 |
| | 2.3237E-13 |
| | 1.24991E-08 |
| | 0.164164161 |
| | 0.210595451 |
| | 1.70729E-07 |
| | 0.987938592 |
| | 0.825379751 |
| | 9.10654E-05 |
| | 1.54196E-09 |
| | 0.982212588 |
| | 7.21142E-09 |
| | 0.001206331 |
| | 3.02404E-09 |

| |
|---|
| 0.000304688 |
| 1.99225E-10 |
| 0.00026545 |
| 4.60743E-14 |
| 5.91989E-08 |
| 1.52819E-06 |
| 0.002210453 |
| 3.73509E-05 |
| 1.46735E-08 |
| 2.61283E-07 |
| 9.75678E-06 |
| 0.001136092 |
| 4.60743E-14 |
| 4.04158E-10 |
| 0.994955656 |
| 5.25247E-13 |
| 0.000573872 |
| 0.003641163 |
| 4.60743E-14 |
| 2.76056E-08 |
| 7.95636E-07 |
| 1.60332E-08 |
| 3.86009E-05 |
| 0.119210775 |
| 0.540600387 |
| 0.531707399 |
| 0.107330977 |
| 0.094484801 |
| 0.022618227 |
| 1.32777E-07 |
| 0.001831317 |
| 0.016239593 |
| 4.25855E-10 |
| 3.18193E-09 |
| 3.25095E-07 |

| |
|---|
| 0.01501394 |
| 0.693593356 |
| 0.895462721 |
| 0.994685906 |
| 0.349894178 |
| 0.933926373 |
| 0.007749773 |
| 0.000111837 |
| 0.999950631 |
| 0.002026234 |
| 0.009150385 |
| 0.982353901 |
| 0.107451806 |
| 0.784839781 |
| 0.999997649 |
| 0.999735169 |
| 0.61884196 |
| 2.76064E-09 |
| 9.01698E-09 |
| 0.964117127 |
| 1.0243E-07 |
| 0.016808638 |
| 0.072715131 |
| 0.000201022 |
| 1.0637E-12 |
| 3.14108E-05 |
| 0.068511717 |
| 0.999998466 |
| 0.993469789 |
| 0.979411766 |
| 0.021618 |
| 0.34949185 |
| 0.08754073 |
| 0.891238219 |
| 2.26142E-11 |

| |
|---|
| 0.000272606 |
| 8.18266E-06 |
| 0.999999909 |
| 0.939236845 |
| 0.015787232 |
| 5.40471E-08 |
| 4.54965E-09 |
| 0.999912415 |
| 0.633886883 |
| 4.9849E-14 |
| 5.97855E-10 |
| 0.000232968 |
| 0.111627776 |
| 4.60743E-14 |
| 0.000111214 |
| 1.78486E-07 |
| 0.001743042 |
| 0.20380059 |
| 0.553664011 |
| 4.62963E-14 |
| 0.006015378 |
| 0.000169604 |
| 0.001210287 |
| 0.191453279 |
| 0.024391116 |
| 0.637917972 |
| 0.000752631 |
| 0.93910708 |
| 1.17467E-06 |
| 0.001238171 |
| 5.74979E-07 |
| 1.29459E-08 |
| 0.816635489 |
| 0.990878621 |
| 4.60743E-14 |

| |
|---|
| 4.9627E-14 |
| 3.02091E-11 |
| 0.003435439 |
| 0.025869564 |
| 0.000988341 |
| 0.004256964 |
| 2.50577E-13 |
| 0.996290385 |
| 0.399482808 |
| 4.38748E-05 |
| 0.401103435 |
| 0.112795762 |
| 0.036284992 |
| 0.814278848 |
| 0.379834357 |
| 0.850096923 |
| 2.90768E-09 |
| 0.836381528 |
| 0.003852229 |
| 0.046868401 |
| 1.48099E-09 |
| 3.59206E-09 |
| 0.039370114 |
| 1.73539E-12 |
| 0.019410325 |
| 0.000825041 |
| 0.006998628 |
| 2.72338E-13 |
| 3.62733E-06 |
| 4.60743E-14 |
| 7.17972E-05 |
| 3.17951E-08 |
| 0.010796913 |
| 2.40132E-09 |
| 0.15934331 |

| |
|---|
| 4.94049E-14 |
| 0.000157927 |
| 3.18719E-08 |
| 2.90966E-07 |
| 0.011016701 |
| 0.019131716 |
| 4.0936E-11 |
| 9.91429E-14 |
| 4.93517E-05 |
| 1.67421E-10 |
| 0.000418854 |
| 0.034217068 |
| 2.3736E-05 |
| 0.033465125 |
| 0.999913862 |
| 0.034849535 |
| 1.92091E-09 |
| 1.47094E-10 |
| 9.31477E-14 |
| 3.88567E-12 |
| 1.81397E-06 |
| 1.06447E-08 |
| 0.004137802 |
| 3.04171E-11 |
| 4.33822E-05 |
| 0.90942957 |
| 0.413309434 |
| 9.4371E-09 |
| 0.004188224 |
| 0.000154024 |
| 0.001113488 |
| 0.253951247 |
| 1.21394E-08 |
| 0.220320977 |
| 0.774500905 |

| |
|---|
| 0.995151187 |
| 0.972576175 |
| 0.420265425 |
| 0.189186434 |
| 0.054942529 |
| 0.999998045 |
| 0.075925516 |
| 2.28417E-08 |
| 0.124066159 |
| 3.79711E-11 |
| 1.52637E-11 |
| 2.61205E-09 |
| 4.52571E-10 |
| 0.000123896 |
| 0.116546625 |
| 0.452342917 |
| 0.102360993 |
| 0.011101955 |
| 6.38894E-05 |
| 6.26044E-08 |
| 4.60743E-14 |
| 5.03059E-08 |
| 0.998281221 |
| 2.98401E-08 |
| 7.76601E-13 |
| 0.97981563 |
| 4.37823E-05 |
| 2.85848E-05 |
| 0.380883016 |
| 0.016160759 |
| 0.99998569 |
| 4.32334E-06 |
| 1.27932E-08 |
| 0.042400074 |
| 0.99999936 |

| |
|---|
| 0.002180538 |
| 0.06943008 |
| 1.21347E-13 |
| 1.75807E-10 |
| 0.00019052 |
| 0.000331917 |
| 1.58497E-10 |
| 8.86728E-09 |
| 4.71845E-14 |
| 2.68138E-05 |
| 5.42525E-07 |
| 1.05138E-13 |
| 0.00097941 |
| 2.03429E-07 |
| 1.61503E-07 |
| 0.000212164 |
| 0.00355005 |
| 7.40263E-12 |
| 6.73117E-12 |
| 0.279779981 |
| 1.2233E-11 |
| 0.000521956 |
| 0.165306562 |
| 4.78874E-06 |
| 0.000102383 |
| 0.999999999 |
| 1.93814E-08 |
| 5.97083E-07 |
| 0.962018148 |
| 0.558393995 |
| 1.67027E-10 |
| 0.998884805 |
| 5.82867E-14 |
| 2.25995E-09 |
| 0.006823934 |

| | |
|---|---|
| | 1.32248E-10 |
| | 0.433879304 |
| | 1.74472E-07 |
| | 2.7266E-05 |
| | 0.000389171 |
| | 0.882186236 |
| | 0.889786694 |
| | 0.944252814 |
| | 2.03194E-08 |
| | 4.78506E-14 |
| | 8.14175E-06 |
| | 1.08819E-11 |
| | 1.05383E-10 |
| | 9.52915E-05 |
| | 4.62963E-14 |
| | 6.93462E-11 |
| | 1.84686E-12 |
| | 0.0001511 |
| | 8.82988E-11 |
| | 4.60743E-14 |
| | 2.31917E-07 |
| | 1.91924E-12 |
| | 3.55955E-07 |
| | 5.04768E-08 |
| | 0.000122118 |
| | 2.4198E-10 |
| | 4.39235E-10 |
| | 1.29174E-12 |
| | 0.086332062 |
| | 0.000997766 |
| | 3.42836E-07 |
| | 2.41741E-05 |
| | 4.17209E-10 |
| | 4.84305E-11 |
| | 5.87308E-14 |

| | |
|---|---|
| | 1.89334E-11 |
| | 3.82712E-07 |
| | 3.29956E-10 |
| | 0.384852968 |
| | 0.278152401 |
| | 3.12239E-08 |
| | 1.46408E-09 |
| | 4.99481E-09 |
| | 0.32338287 |
| | 8.14115E-12 |
| | 2.59804E-07 |
| | 3.58543E-07 |
| | 4.62963E-14 |
| | 0.025847826 |
| | 1.32075E-06 |
| | 0.081380025 |
| | 7.70403E-10 |
| | 0.388992774 |
| | 0.999979523 |
| | 2.5413E-13 |
| | 0.092934552 |
| | 0.010527554 |
| | 1.36524E-09 |
| | 4.60743E-14 |
| | 1.56208E-13 |
| | 7.4485E-07 |
| | 3.72614E-05 |
| | 4.60743E-14 |
| | 0.999569569 |
| | 0.974275417 |
| | 2.64299E-08 |
| | 0.001818993 |
| | 1.66515E-10 |
| | 0.94925891 |
| | 0.665098093 |

| | |
|---|---|
| | 0.103446428 |
| | 0.013256519 |
| | 0.997952298 |
| | 0.017938575 |
| | 0.000182198 |
| | 4.9849E-14 |
| | 0.924946152 |
| | 0.96590544 |
| | 0.001373907 |
| | 0.940569939 |
| | 0.050877005 |
| | 3.52281E-06 |
| | 0.073107438 |
| | 0.008234953 |
| | 0.00138032 |
| | 2.36153E-10 |
| | 2.52855E-09 |
| | 1.73904E-10 |
| | 0.001039961 |
| | 0.000272631 |
| | 0.000192232 |
| | 0.012835195 |
| | 0.680695766 |
| | 0.927767289 |
| | 0.001634767 |
| | 1.88753E-08 |
| | 7.03549E-05 |
| | 1.10016E-10 |
| | 7.02999E-07 |
| | 0.363433442 |
| | 4.9412E-06 |
| | 7.84343E-07 |
| | 3.08551E-08 |
| | 0.99884597 |
| | 6.82334E-11 |

| | |
|---|---|
| | 0.014700689 |
| | 0.3703354 |
| | 8.76409E-10 |
| | 0.633256862 |
| | 1.12688E-13 |
| | 0.172689873 |
| | 6.30275E-09 |
| | 0.999985936 |
| | 4.60743E-14 |
| | 6.9611E-14 |
| | 0.027446999 |
| | 0.000336427 |
| | 1.61594E-06 |
| | 0.132446844 |
| | 7.02129E-09 |
| | 0.00813208 |
| | 2.35026E-06 |
| | 1.15163E-12 |
| | 3.66749E-08 |
| | 0.305385092 |
| | 1.14307E-10 |
| | 4.60743E-14 |
| | 1.19571E-13 |
| | 9.87612E-05 |
| | 2.41521E-09 |
| | 8.31795E-09 |
| | 7.89369E-14 |
| | 0.000136425 |
| | 0.00024958 |
| | 8.01227E-10 |
| | 0.036106125 |
| | 0.068333419 |
| | 7.28809E-07 |
| | 0.939031809 |
| | 0.000253293 |

| |
|---|
| 2.09769E-10 |
| 0.999999965 |
| 1.84018E-08 |
| 0.015373566 |
| 0.865508478 |
| 0.557169069 |
| 0.999987202 |
| 0.002112387 |
| 5.14195E-05 |
| 0.956588039 |
| 0.002533074 |
| 0.755081262 |
| 4.24303E-06 |
| 3.04351E-09 |
| 0.000134331 |
| 0.34023675 |
| 9.0621E-09 |
| 0.988346137 |
| 0.393249982 |
| 0.000575025 |
| 0.000374471 |
| 0.039593786 |
| 0.974065832 |
| 4.01098E-06 |
| 0.221791476 |
| 0.809442931 |
| 0.000487812 |
| 0.891208337 |
| 0.126779146 |
| 0.999999757 |
| 0.005687286 |
| 5.01191E-08 |
| 0.246545769 |
| 1.08091E-08 |
| 0.9121628 |

| | |
|---|---|
| | 0.057547996 |
| | 0.905507082 |
| | 0.615715459 |
| | 0.294587538 |
| | 0.832248642 |
| | 5.55871E-08 |
| | 4.60743E-14 |
| | 4.7988E-08 |
| | 7.55382E-10 |
| | 0.128650497 |
| | 0.001648609 |
| | 0.214944866 |
| | 6.79185E-09 |
| | 0.997331162 |
| | 4.60743E-14 |
| | 0.001491878 |
| | 0.006571419 |
| | 0.090264746 |
| | 0.000388495 |
| | 2.39554E-06 |
| | 0.016272483 |
| | 1.42699E-05 |
| | 0.410176981 |
| | 0.045316897 |
| | 0.013073493 |
| | 0.003814265 |
| | 0.002206637 |
| | 8.89699E-12 |
| | 2.07484E-06 |
| | 0.171216578 |
| | 4.45344E-12 |
| | 0.006345438 |
| | 2.50794E-05 |
| | 0.10910249 |
| | 1.9803E-12 |

| |
|---|
| 0.463784925 |
| 0.005247301 |
| 4.01926E-07 |
| 0.027393682 |
| 5.48116E-05 |
| 0.157275823 |
| 0.022663436 |
| 1.11545E-08 |
| 1.90962E-05 |
| 0.163802756 |
| 6.18215E-06 |
| 4.86034E-05 |
| 0.997775032 |
| 4.4294E-05 |
| 0.000343519 |
| 0.000936489 |
| 0.003575483 |
| 0.028502355 |
| 0.024952166 |
| 0.342679582 |
| 0.123657307 |
| 0.008592889 |
| 0.007303395 |
| 0.838894887 |
| 0.010976317 |
| 0.000955835 |
| 0.008842827 |
| 0.006799176 |
| 0.866428585 |
| 0.794961401 |
| 7.0824E-06 |
| 3.3713E-06 |
| 0.85259772 |
| 0.007848532 |
| 0.867585993 |

| | |
|---|---|
| | 0.02387797 |
| | 0.001517792 |
| | 0.298238494 |
| | 1.31257E-05 |
| | 0.018121703 |
| | 1.31617E-12 |
| | 0.999337378 |
| | 9.27325E-07 |
| | 0.215266112 |
| | 2.244E-07 |
| | 4.62963E-14 |
| | 1.02456E-10 |
| | 6.13857E-09 |
| | 0.000244646 |
| | 0.001527818 |
| | 0.00052896 |
| | 1.02539E-08 |
| | 0.70606084 |
| | 0.986924824 |
| | 0.311691781 |
| | 0.999930013 |
| | 0.991797783 |
| | 3.84404E-08 |
| | 0.000652413 |
| | 0.999037046 |
| | 0.000802262 |
| | 0.043386227 |
| | 0.049789836 |
| | 0.00068986 |
| | 0.999768234 |
| | 0.003267812 |
| | 0.000998069 |
| | 0.258433114 |
| | 0.001984007 |
| | 0.000147016 |

| |
|---|
| 0.193015605 |
| 0.170085963 |
| 0.922901162 |
| 0.010181433 |
| 0.000116162 |
| 2.32854E-09 |
| 0.000105911 |
| 0.999021236 |
| 0.99358414 |
| 0.727529308 |
| 0.302495005 |
| 0.366513137 |
| 0.477652408 |
| 1.59169E-05 |
| 8.04427E-07 |

| |
|---|
| P-value: (SM_pos_12022024_Newanalyis) / (Water_pos_12022024_Newanalysis) |
| 0.995272109 |
| 0.995600677 |
| 0.927567352 |
| 0.046207042 |
| 0.11207904 |
| 0.990504483 |
| 0.831147098 |
| 0.951015655 |
| 0.004177255 |
| 0.024710287 |
| 0.001888668 |
| 8.77615E-09 |
| 0.60440149 |
| 4.18667E-05 |
| 0.998650398 |
| 0.989262511 |
| 0.999268002 |
| 0.130675659 |
| 0.039261782 |
| 0.158301635 |
| 0.945706008 |
| 0.996170584 |
| 0.0227927 |
| 0.747923292 |
| 0.036354006 |
| 0.809524149 |
| 0.999999984 |
| 0.986387553 |
| 0.020316203 |
| 0.998918181 |
| 0.950821702 |
| 0.001526276 |
| 0.835691609 |

| |
|---|
| 0.004043563 |
| 1.6845E-06 |
| 0.998916886 |
| 0.007168299 |
| 0.251401846 |
| 0.044456418 |
| 0.10021385 |
| 0.225977009 |
| 0.999999465 |
| 0.260434507 |
| 2.78606E-05 |
| 0.929401577 |
| 0.902949746 |
| 0.999737774 |
| 0.617173162 |
| 0.011799658 |
| 0.868165684 |
| 0.024863154 |
| 0.876522444 |
| 0.9997807 |
| 0.791465793 |
| 1.36907E-06 |
| 0.921511122 |
| 4.18E-06 |
| 0.999996295 |
| 0.999999992 |
| 0.958920039 |
| 0.999968753 |
| 0.035981806 |
| 0.999990169 |
| 0.155799883 |
| 0.238158771 |
| 0.357458562 |
| 0.351132236 |
| 3.91708E-05 |

| |
|---|
| 0.018635193 |
| 0.68556693 |
| 0.720785479 |
| 0.015999475 |
| 0.100003571 |
| 1.77733E-06 |
| 0.705007775 |
| 0.000812988 |
| 3.29656E-06 |
| 0.741513687 |
| 3.18445E-05 |
| 8.49003E-10 |
| 0.267611989 |
| 0.00033346 |
| 0.864003311 |
| 0.015605864 |
| 0.475437195 |
| 0.999112442 |
| 0.015462384 |
| 0.016413761 |
| 0.076915624 |
| 0.529481441 |
| 6.26045E-07 |
| 0.000110197 |
| 0.967902319 |
| 0.649650288 |
| 0.999809045 |
| 0.84675345 |
| 0.164602494 |
| 0.989700598 |
| 2.81423E-05 |
| 0.148175825 |
| 0.922010185 |
| 0.954107189 |
| 3.51231E-08 |

| | |
|---|---|
| | 2.33262E-05 |
| | 0.986110994 |
| | 0.983710573 |
| | 0.999999857 |
| | 0.99426104 |
| | 0.981887758 |
| | 0.722704847 |
| | 0.00617861 |
| | 0.994391075 |
| | 0.720074103 |
| | 0.159898027 |
| | 0.746321149 |
| | 0.243330645 |
| | 0.980990538 |
| | 0.019750778 |
| | 0.038858486 |
| | 0.000494252 |
| | 0.823859047 |
| | 0.999433813 |
| | 0.022416546 |
| | 0.000212977 |
| | 0.483043948 |
| | 0.998671143 |
| | 0.918683061 |
| | 2.77262E-11 |
| | 0.99986809 |
| | 0.038595428 |
| | 0.007703276 |
| | 0.264618755 |
| | 0.084259632 |
| | 0.9948778 |
| | 4.47468E-05 |
| | 0.099541575 |
| | 0.992525165 |
| | 0.737669301 |

| | |
|---|---|
| | 0.080970209 |
| | 0.999999639 |
| | 0.06049032 |
| | 0.942338163 |
| | 0.441298602 |
| | 0.958191312 |
| | 0.960610582 |
| | 0.994575075 |
| | 0.634308598 |
| | 0.797618395 |
| | 0.992438736 |
| | 0.675319606 |
| | 0.000142588 |
| | 0.148092908 |
| | 0.999999658 |
| | 2.61051E-07 |
| | 0.076997015 |
| | 0.001194597 |
| | 0.999979954 |
| | 0.90416312 |
| | 0.663399485 |
| | 0.577364732 |
| | 0.28839335 |
| | 0.009288135 |
| | 2.18553E-08 |
| | 6.43214E-05 |
| | 0.989105687 |
| | 0.999941782 |
| | 0.999999966 |
| | 0.99999997 |
| | 0.000177448 |
| | 0.959763434 |
| | 0.99315819 |
| | 0.003428555 |
| | 0.003488857 |

| | |
|---|---|
| | 0.000573276 |
| | 0.89282302 |
| | 0.94576759 |
| | 2.34101E-11 |
| | 0.028345193 |
| | 0.999999009 |
| | 0.143152466 |
| | 0.262145194 |
| | 1.66922E-12 |
| | 4.48263E-05 |
| | 0.748304476 |
| | 3.38999E-05 |
| | 5.85887E-07 |
| | 8.4651E-07 |
| | 0.999924095 |
| | 7.19587E-10 |
| | 0.997694377 |
| | 1.53713E-05 |
| | 1.60427E-13 |
| | 0.999989863 |
| | 0.013308612 |
| | 0.016789048 |
| | 0.995261492 |
| | 0.3785646 |
| | 0.909995939 |
| | 0.985459718 |
| | 0.99999216 |
| | 0.000700641 |
| | 0.99955763 |
| | 0.043042743 |
| | 0.008717362 |
| | 0.004152059 |
| | 2.00641E-05 |
| | 0.994362203 |
| | 3.34943E-09 |

| | |
|---|---|
| | 0.993563058 |
| | 0.982144195 |
| | 0.492304175 |
| | 0.813589652 |
| | 0.943626927 |
| | 2.81959E-07 |
| | 3.26465E-10 |
| | 0.001525728 |
| | 9.78811E-05 |
| | 1.05306E-05 |
| | 0.23900122 |
| | 0.978613297 |
| | 0.004414548 |
| | 0.111700409 |
| | 0.996944906 |
| | 0.92346794 |
| | 0.012062642 |
| | 1.59181E-06 |
| | 0.726348933 |
| | 0.69281704 |
| | 0.000540881 |
| | 0.999642953 |
| | 0.00302649 |
| | 1.15365E-05 |
| | 0.025582149 |
| | 0.960970343 |
| | 0.9989743 |
| | 0.044692915 |
| | 0.217694433 |
| | 5.64615E-05 |
| | 0.507813091 |
| | 0.986178368 |
| | 0.944475206 |
| | 0.186506648 |
| | 0.999999999 |

| | |
|---|---|
| | 0.005542593 |
| | 0.005659118 |
| | 0.999178046 |
| | 0.999986366 |
| | 0.999215881 |
| | 1.51517E-05 |
| | 0.000620554 |
| | 0.99601755 |
| | 0.116918235 |
| | 3.32107E-08 |
| | 7.03288E-07 |
| | 0.009073298 |
| | 0.748385958 |
| | 2.14823E-09 |
| | 0.012629733 |
| | 0.000319973 |
| | 0.011412825 |
| | 0.466033431 |
| | 0.986023206 |
| | 0.979723665 |
| | 0.97232504 |
| | 3.53972E-07 |
| | 0.908561292 |
| | 3.16795E-06 |
| | 0.529758937 |
| | 0.91104106 |
| | 0.193852679 |
| | 0.942460791 |
| | 4.11309E-05 |
| | 0.829195571 |
| | 0.000368836 |
| | 0.711114425 |
| | 0.985889503 |
| | 0.914279209 |
| | 5.02931E-14 |

| |
|---|
| 2.53586E-09 |
| 0.999967784 |
| 0.807239334 |
| 0.005188127 |
| 3.10202E-07 |
| 0.128083256 |
| 0.000133576 |
| 0.971552892 |
| 0.99921875 |
| 0.741026077 |
| 0.000212363 |
| 0.011328966 |
| 0.016942349 |
| 0.138668126 |
| 0.999987055 |
| 0.46224301 |
| 0.407369457 |
| 0.000422957 |
| 5.27103E-05 |
| 0.552354936 |
| 0.928408246 |
| 0.0122517 |
| 0.848316346 |
| 0.072114672 |
| 0.208186027 |
| 0.01014258 |
| 0.986125836 |
| 0.142596739 |
| 0.511773247 |
| 1.33793E-12 |
| 0.999990565 |
| 0.016231173 |
| 0.000848803 |
| 2.86993E-13 |
| 0.450766398 |

| | |
|---|---|
| | 3.27466E-09 |
| | 0.901546811 |
| | 0.556062243 |
| | 0.756173742 |
| | 6.97541E-07 |
| | 0.602784092 |
| | 6.41012E-09 |
| | 0.496683167 |
| | 1.92146E-12 |
| | 0.005137628 |
| | 0.000190006 |
| | 1.49282E-07 |
| | 9.14676E-08 |
| | 0.004350592 |
| | 2.70694E-05 |
| | 0.065974914 |
| | 0.000994391 |
| | 0.42295601 |
| | 0.081971519 |
| | 1.22577E-05 |
| | 8.40994E-13 |
| | 4.8356E-06 |
| | 0.008434626 |
| | 0.115332649 |
| | 0.999183338 |
| | 5.01592E-07 |
| | 0.101095162 |
| | 0.070843751 |
| | 2.57104E-07 |
| | 0.517387533 |
| | 0.288391607 |
| | 0.720537705 |
| | 0.014142991 |
| | 0.747624881 |
| | 0.97817312 |

| |
|---|
| 0.000188846 |
| 0.939104034 |
| 0.116196227 |
| 0.208872477 |
| 0.367476953 |
| 0.800374038 |
| 0.000212655 |
| 0.689536806 |
| 0.323518919 |
| 1.34939E-05 |
| 0.006790946 |
| 0.957460558 |
| 0.005162201 |
| 0.999999937 |
| 0.970909547 |
| 8.86761E-06 |
| 0.826735289 |
| 0.278693184 |
| 0.023215812 |
| 0.034712675 |
| 1.75923E-09 |
| 0.005612993 |
| 0.381692018 |
| 0.993123847 |
| 5.79186E-06 |
| 0.903138433 |
| 0.010710524 |
| 0.984435032 |
| 0.020568395 |
| 0.136952742 |
| 0.924050988 |
| 0.346146117 |
| 0.991861451 |
| 0.266336928 |
| 5.97766E-10 |

| |
|---|
| 0.007439043 |
| 0.364034435 |
| 0.300020565 |
| 0.403049893 |
| 0.713402303 |
| 0.031639956 |
| 0.533145276 |
| 0.131012197 |
| 0.047630403 |
| 0.837995762 |
| 0.034781956 |
| 0.08358686 |
| 0.003046628 |
| 0.741269963 |
| 6.24761E-06 |
| 0.999668402 |
| 0.999999955 |
| 6.32793E-07 |
| 0.000675833 |
| 0.003722551 |
| 8.13793E-14 |
| 0.000101329 |
| 0.016087228 |
| 0.267646301 |
| 8.04632E-05 |
| 1 |
| 0.175547692 |
| 1.76002E-10 |
| 0.413113519 |
| 0.255922674 |
| 0.630048253 |
| 0.484884232 |
| 5.28771E-11 |
| 5.49861E-08 |
| 0.009760331 |

| | |
|---|---|
| | 1.62699E-05 |
| | 3.86738E-07 |
| | 1.40037E-07 |
| | 5.56879E-06 |
| | 8.53912E-07 |
| | 0.677638374 |
| | 1.02567E-10 |
| | 1.05245E-05 |
| | 0.307222629 |
| | 2.73123E-08 |
| | 0.567726458 |
| | 6.80405E-05 |
| | 0.000680178 |
| | 3.73884E-10 |
| | 2.77766E-11 |
| | 4.00335E-12 |
| | 0.196779294 |
| | 0.942083497 |
| | 2.13698E-09 |
| | 4.65183E-14 |
| | 0.133618679 |
| | 9.87005E-08 |
| | 0.000849397 |
| | 1.74474E-09 |
| | 2.72972E-07 |
| | 4.62963E-14 |
| | 0.000139911 |
| | 0.927519718 |
| | 0.318632975 |
| | 0.999987441 |
| | 0.000843632 |
| | 0.000556562 |
| | 1.55348E-05 |
| | 0.000229503 |
| | 1.52669E-10 |

| | |
|---|---|
| | 0.93908244 |
| | 0.992805184 |
| | 0.026119536 |
| | 0.007236426 |
| | 0.621570257 |
| | 2.64635E-07 |
| | 1.70341E-08 |
| | 9.88982E-09 |
| | 0.211254647 |
| | 1.05304E-06 |
| | 4.11219E-05 |
| | 1.38366E-11 |
| | 1.72337E-05 |
| | 0.012169043 |
| | 0.024455038 |
| | 0.110275457 |
| | 0.002519108 |
| | 0.366147719 |
| | 0.95157151 |
| | 7.60184E-06 |
| | 0.773556508 |
| | 0.152670056 |
| | 0.954071815 |
| | 5.02144E-11 |
| | 2.74856E-08 |
| | 0.999971511 |
| | 7.64495E-06 |
| | 3.68525E-07 |
| | 0.002317901 |
| | 0.014788084 |
| | 0.925985751 |
| | 0.000265279 |
| | 0.000314332 |
| | 0.987527269 |
| | 0.390799803 |

| |
|---|
| 0.867053922 |
| 2.53356E-07 |
| 0.884382045 |
| 0.765460049 |
| 0.402573699 |
| 5.84076E-06 |
| 0.007852758 |
| 0.661880724 |
| 0.159266734 |
| 0.372229502 |
| 0.354571067 |
| 0.000353324 |
| 0.92969014 |
| 0.003151016 |
| 0.138642829 |
| 0.001602673 |
| 0.000391904 |
| 7.36211E-10 |
| 0.996520393 |
| 3.56655E-08 |
| 0.073605904 |
| 3.03619E-06 |
| 1.93788E-06 |
| 0.999895 |
| 0.062153866 |
| 0.998921932 |
| 0.001429961 |
| 0.505620351 |
| 0.934958483 |
| 0.999031673 |
| 0.004212463 |
| 0.27429352 |
| 0.997762857 |
| 0.055764345 |
| 0.141795166 |

| | |
|---|---|
| | 0.000976915 |
| | 0.150274721 |
| | 1.81316E-08 |
| | 0.524810015 |
| | 4.60743E-14 |
| | 0.680070218 |
| | 0.879783316 |
| | 0.75395458 |
| | 0.518013159 |
| | 2.67489E-07 |
| | 0.644230518 |
| | 0.059936401 |
| | 3.44609E-05 |
| | 0.999966414 |
| | 0.960243485 |
| | 1.52474E-05 |
| | 0.014190943 |
| | 4.74065E-14 |
| | 0.03581006 |
| | 0.111352158 |
| | 5.66729E-08 |
| | 0.016473953 |
| | 4.60743E-14 |
| | 0.999992856 |
| | 1.17573E-13 |
| | 0.03947378 |
| | 0.258139314 |
| | 2.85828E-05 |
| | 5.87308E-14 |
| | 6.80001E-12 |
| | 0.828755551 |
| | 0.538089642 |
| | 0.695116365 |
| | 0.000166656 |
| | 0.999912683 |

| |
|---|
| 1.76598E-05 |
| 0.373923913 |
| 0.799626289 |
| 0.007543053 |
| 0.965797006 |
| 0.334080517 |
| 0.950709365 |
| 0.019810575 |
| 0.835031299 |
| 0.010747886 |
| 0.000408931 |
| 0.999902962 |
| 3.61666E-09 |
| 0.999237502 |
| 0.133014199 |
| 0.991872779 |
| 0.55495791 |
| 0.181506692 |
| 0.004238672 |
| 0.009882486 |
| 1.6198E-11 |
| 0.962498868 |
| 0.608062443 |
| 0.359188877 |
| 0.817746572 |
| 0.711907039 |
| 0.003594852 |
| 0.866104755 |
| 0.999751628 |
| 0.239581434 |
| 0.422352012 |
| 0.000126135 |
| 0.377012138 |
| 3.93556E-09 |
| 0.861578771 |

| |
|---|
| 0.20158937 |
| 0.516357912 |
| 0.983563184 |
| 0.440668806 |
| 0.050618624 |
| 0.999994748 |
| 0.525204334 |
| 0.999997648 |
| 1.69312E-05 |
| 9.87496E-09 |
| 0.372789957 |
| 0.997982568 |
| 0.598659382 |
| 0.718146729 |
| 1.28335E-08 |
| 0.751580221 |
| 0.999999988 |
| 0.027668596 |
| 0.419618637 |
| 0.158586035 |
| 0.050021966 |
| 0.260973315 |
| 0.93062365 |
| 0.73547 |
| 0.502827342 |
| 0.007400892 |
| 0.40324646 |
| 0.987662465 |
| 0.000569592 |
| 0.552953678 |
| 0.418568489 |
| 0.22955349 |
| 0.76813899 |
| 0.007731723 |
| 0.010968673 |

| | |
|---|---|
| | 0.299767039 |
| | 2.0367E-06 |
| | 0.04939667 |
| | 0.233947057 |
| | 1.5118E-05 |
| | 1.2071E-07 |
| | 0.000154732 |
| | 4.60743E-14 |
| | 1.003E-06 |
| | 0.902975358 |
| | 2.85364E-11 |
| | 5.06895E-05 |
| | 0.691557748 |
| | 2.55486E-11 |
| | 0.986134602 |
| | 0.994553815 |
| | 3.56967E-05 |
| | 0.495161029 |
| | 0.000577965 |
| | 0.993305367 |
| | 9.7928E-07 |
| | 0.021520855 |
| | 0.012112586 |
| | 0.998421223 |
| | 0.087309018 |
| | 0.009382767 |
| | 0.0495265 |
| | 0.997730144 |
| | 0.999962648 |
| | 0.167411371 |
| | 0.327107084 |
| | 0.01471034 |
| | 0.97057206 |
| | 1.08769E-12 |
| | 0.998671336 |

| | |
|---|---|
| | 0.995053292 |
| | 0.012273711 |
| | 6.01181E-06 |
| | 0.0063912 |
| | 0.040876779 |
| | 0.584577459 |
| | 0.955676074 |
| | 0.000177663 |
| | 0.577036137 |
| | 0.737037559 |
| | 6.74486E-09 |
| | 3.29868E-06 |
| | 0.007048451 |
| | 0.200351965 |
| | 0.207286964 |
| | 0.00473997 |
| | 0.914932904 |
| | 0.534383088 |
| | 0.075284724 |
| | 0.872803545 |
| | 0.555792908 |
| | 0.945511867 |
| | 0.151887352 |
| | 0.98679006 |
| | 0.002248694 |
| | 0.993538822 |
| | 2.2036E-05 |
| | 0.04414801 |
| | 0.000339933 |
| | 0.806791465 |
| | 0.020939619 |
| | 0.270612602 |
| | 0.192253446 |
| | 0.998501613 |
| | 0.340169146 |

| | |
|---|---|
| | 0.74873345 |
| | 0.020852093 |
| | 0.999999738 |
| | 0.00025421 |
| | 0.273273371 |
| | 0.084958218 |
| | 0.999999598 |
| | 0.987143947 |
| | 0.001296235 |
| | 0.999997348 |
| | 0.615296825 |
| | 0.999993125 |
| | 0.406880005 |
| | 0.00597117 |
| | 2.71581E-09 |

| |
|---|
| Adj. P-value: (Bb_pos_12022024_Newanalysis) / (CM_pos_12022024_Newanalysis) |
| 1 |
| 1 |
| 1 |
| 0.087855222 |
| 1 |
| 1 |
| 1 |
| 0.049246863 |
| 1 |
| 1 |
| 1 |
| 1 |
| 1 |
| 1 |
| 1 |
| 0.730117293 |
| 1 |
| 3.41414E-11 |
| 1 |
| 0.010281558 |
| 1 |
| 6.88319E-09 |
| 0.001311584 |
| 1.64844E-10 |
| 3.41414E-11 |
| 1 |
| 1 |
| 1 |
| 0.828833146 |
| 1 |
| 1 |
| 2.00712E-06 |
| 1 |

| |
|---|
| 1 |
| 9.25133E-11 |
| 0.536991378 |
| 0.00079473 |
| 0.009774792 |
| 0.005287 |
| 8.32703E-07 |
| 1 |
| 1 |
| 0.023537311 |
| 0.232464102 |
| 1 |
| 1.93975E-08 |
| 1.09196E-06 |
| 2.5962E-06 |
| 0.028048176 |
| 0.004136976 |
| 0.003787174 |
| 0.000112637 |
| 2.236E-08 |
| 0.004432949 |
| 1 |
| 1 |
| 0.0681081 |
| 1 |
| 1 |
| 0.273228579 |
| 1 |
| 1 |
| 0.881950486 |
| 8.82681E-07 |
| 0.01154846 |
| 1 |
| 1.53302E-09 |
| 1 |

| |
|---|
| 1 |
| 1 |
| 2.11014E-05 |
| 1 |
| 0.006392027 |
| 0.297268807 |
| 1 |
| 1 |
| 3.92234E-11 |
| 1 |
| 9.99156E-09 |
| 3.09689E-06 |
| 0.001117683 |
| 1.14067E-10 |
| 1 |
| 0.001017834 |
| 0.491497917 |
| 0.205894201 |
| 1 |
| 7.13509E-06 |
| 0.349939462 |
| 0.552518265 |
| 0.000365308 |
| 4.29658E-07 |
| 0.000236408 |
| 0.000337432 |
| 0.001989496 |
| 0.165025963 |
| 2.70569E-07 |
| 0.0023725 |
| 1 |
| 1 |
| 0.211466347 |
| 1 |
| 0.006370835 |

| |
|---|
| 1 |
| 0.042737567 |
| 1 |
| 0.621593492 |
| 1 |
| 1 |
| 1 |
| 0.38891763 |
| 1 |
| 0.000995768 |
| 0.249741422 |
| 1 |
| 1 |
| 1 |
| 3.41414E-11 |
| 0.009418424 |
| 1 |
| 1 |
| 1 |
| 1 |
| 0.821126521 |
| 1 |
| 1 |
| 0.63163488 |
| 1 |
| 1 |
| 1 |
| 1 |
| 0.805721289 |
| 6.90717E-07 |
| 0.093731442 |
| 0.053079587 |
| 0.052133542 |
| 0.030462166 |
| 1 |

| | |
|---|---|
| | 1 |
| | 1 |
| | 1 |
| | 1 |
| | 1 |
| | 1 |
| | 1 |
| | 1 |
| | 0.022503865 |
| | 1 |
| | 1 |
| | 1 |
| | 2.33438E-05 |
| | 1 |
| | 1 |
| | 7.49206E-07 |
| | 1 |
| | 1 |
| | 0.382898089 |
| | 1 |
| | 1 |
| | 1 |
| | 1 |
| | 0.000467136 |
| | 1 |
| | 0.009355225 |
| | 0.863019893 |
| | 1 |
| | 1 |
| | 1 |
| | 0.802497902 |
| | 0.711952381 |
| | 0.892195167 |
| | 1 |
| | 0.435598131 |

| | |
|---|---|
| | 0.312569576 |
| | 1 |
| | 0.018170581 |
| | 0.000101412 |
| | 0.004615778 |
| | 1 |
| | 0.375021755 |
| | 1 |
| | 0.224330318 |
| | 1 |
| | 1 |
| | 1 |
| | 9.97582E-09 |
| | 0.00981996 |
| | 0.969969467 |
| | 1 |
| | 1 |
| | 0.696224249 |
| | 6.87647E-11 |
| | 0.049973545 |
| | 0.168177129 |
| | 0.073791839 |
| | 0.603320668 |
| | 1 |
| | 0.27799717 |
| | 1 |
| | 0.624459377 |
| | 1 |
| | 0.013230092 |
| | 0.001823926 |
| | 0.001063096 |
| | 0.043989344 |
| | 0.162227348 |
| | 0.844504817 |
| | 0.696972583 |

| | |
|---|---|
| | 0.000214407 |
| | 1 |
| | 1 |
| | 1 |
| | 0.935400752 |
| | 1 |
| | 1 |
| | 1 |
| | 1 |
| | 1 |
| | 0.952719308 |
| | 1 |
| | 0.649406057 |
| | 1 |
| | 0.198941865 |
| | 0.006673324 |
| | 0.001793721 |
| | 0.125937571 |
| | 0.000583293 |
| | 1 |
| | 0.038336845 |
| | 1 |
| | 1 |
| | 1 |
| | 0.000870939 |
| | 0.491837106 |
| | 1 |
| | 1 |
| | 0.324538249 |
| | 1 |
| | 1 |
| | 1 |
| | 1 |
| | 1 |
| | 3.23756E-05 |

| |
|---|
| 0.633303397 |
| 1 |
| 1 |
| 1 |
| 1 |
| 0.088686293 |
| 0.606661334 |
| 1 |
| 1 |
| 8.79366E-06 |
| 0.101915013 |
| 1 |
| 1 |
| 0.036962169 |
| 0.001064181 |
| 0.05661249 |
| 0.010645578 |
| 0.043102575 |
| 1 |
| 2.23679E-10 |
| 1 |
| 0.073893475 |
| 1 |
| 0.783500034 |
| 1 |
| 1 |
| 1 |
| 1 |
| 0.114382101 |
| 1 |
| 0.645696869 |
| 2.85332E-06 |
| 0.975615224 |
| 1 |
| 2.76968E-10 |

| |
|---|
| 1 |
| 1.35972E-05 |
| 0.631141147 |
| 1 |
| 1 |
| 1 |
| 0.000261525 |
| 1 |
| 1 |
| 1 |
| 0.194766793 |
| 1 |
| 1 |
| 0.4141779 |
| 1 |
| 1 |
| 0.017576568 |
| 1 |
| 0.011288165 |
| 1 |
| 5.29453E-08 |
| 0.001253103 |
| 1 |
| 0.19200311 |
| 1 |
| 1 |
| 1 |
| 0.000170194 |
| 0.008022841 |
| 0.00107338 |
| 0.811955802 |
| 0.116613983 |
| 0.068925519 |
| 0.480336088 |
| 0.530584385 |

| | |
|---|---|
| | 0.242464701 |
| | 1 |
| | 0.000790429 |
| | 1 |
| | 1 |
| | 1 |
| | 0.119156751 |
| | 1.00972E-05 |
| | 1 |
| | 0.023644478 |
| | 1 |
| | 1 |
| | 0.063518007 |
| | 0.057818736 |
| | 1 |
| | 1 |
| | 1 |
| | 0.000174096 |
| | 8.31194E-07 |
| | 0.02187563 |
| | 0.275008895 |
| | 0.050988199 |
| | 0.931214827 |
| | 0.000336867 |
| | 0.863000538 |
| | 1 |
| | 0.334539461 |
| | 4.6833E-06 |
| | 1.485E-07 |
| | 0.000383935 |
| | 0.001551571 |
| | 0.880275279 |
| | 0.000628452 |
| | 9.53911E-10 |
| | 0.004501236 |

| |
|---|
| 5.42889E-06 |
| 1 |
| 1.965E-07 |
| 5.52376E-05 |
| 1 |
| 1 |
| 1 |
| 1 |
| 1 |
| 0.170551174 |
| 0.014720546 |
| 1 |
| 1 |
| 0.053950483 |
| 1 |
| 0.334540413 |
| 0.089964536 |
| 1 |
| 0.108789483 |
| 1 |
| 9.65721E-10 |
| 0.193001592 |
| 1 |
| 0.064022285 |
| 0.000401595 |
| 1 |
| 0.728504769 |
| 0.830778674 |
| 1 |
| 0.191316399 |
| 1 |
| 1 |
| 1 |
| 0.001291262 |
| 1 |

| |
|---|
| 1 |
| 1 |
| 0.690820544 |
| 1 |
| 1 |
| 1 |
| 1 |
| 1 |
| 0.148891913 |
| 0.408065767 |
| 1 |
| 1 |
| 1 |
| 0.06643008 |
| 0.082081803 |
| 0.219649712 |
| 0.181583173 |
| 1.58646E-06 |
| 3.17155E-06 |
| 0.603057369 |
| 0.558410173 |
| 0.056730096 |
| 1 |
| 1 |
| 0.587297982 |
| 1 |
| 1 |
| 0.125948482 |
| 1 |
| 0.757433431 |
| 6.42648E-06 |
| 8.23757E-08 |
| 2.29387E-06 |
| 0.000899704 |
| 0.472048613 |

| | |
|---|---|
| | 0.022375455 |
| | 0.623559912 |
| | 1 |
| | 0.252803299 |
| | 1 |
| | 0.265874327 |
| | 0.001611823 |
| | 1 |
| | 1 |
| | 1 |
| | 1 |
| | 0.000898164 |
| | 0.000116742 |
| | 0.000405597 |
| | 1 |
| | 1 |
| | 0.084926402 |
| | 1 |
| | 0.000257592 |
| | 5.77379E-11 |
| | 1 |
| | 1.40389E-07 |
| | 0.029260253 |
| | 0.003050551 |
| | 1 |
| | 4.1085E-05 |
| | 1 |
| | 9.07657E-05 |
| | 0.821092391 |
| | 1 |
| | 0.618903436 |
| | 1 |
| | 0.553709813 |
| | 5.98772E-05 |
| | 3.89013E-06 |

| | |
|---|---|
| | 0.00017526 |
| | 0.805721289 |
| | 1 |
| | 1 |
| | 0.549929609 |
| | 1 |
| | 0.000328478 |
| | 0.115563377 |
| | 1 |
| | 0.194791488 |
| | 1 |
| | 0.057314613 |
| | 3.41414E-11 |
| | 6.6101E-06 |
| | 3.756E-05 |
| | 3.41414E-11 |
| | 3.41414E-11 |
| | 3.54905E-11 |
| | 1 |
| | 1 |
| | 1 |
| | 1 |
| | 0.052192656 |
| | 1 |
| | 2.73178E-05 |
| | 3.82558E-11 |
| | 2.50361E-08 |
| | 1 |
| | 1 |
| | 1 |
| | 1 |
| | 3.41414E-11 |
| | 3.47379E-07 |
| | 3.41414E-11 |
| | 0.007155086 |

| |
|---:|
| 1 |
| 0.004517387 |
| 1 |
| 1 |
| 0.000651537 |
| 1 |
| 1 |
| 1 |
| 1 |
| 1 |
| 1 |
| 1 |
| 1 |
| 1 |
| 1 |
| 0.000686103 |
| 6.53969E-07 |
| 0.978325555 |
| 0.000546647 |
| 1 |
| 1 |
| 1 |
| 1 |
| 1 |
| 1 |
| 1 |
| 1 |
| 1 |
| 1 |
| 1 |
| 0.072510399 |
| 0.009272503 |
| 1 |
| 0.010779734 |
| 0.179900032 |

| | |
|---|---|
| | 1 |
| | 1 |
| | 1 |
| | 0.613192229 |
| | 0.116404043 |
| | 1 |
| | 0.05670796 |
| | 1 |
| | 6.27156E-10 |
| | 0.005170904 |
| | 1 |
| | 1 |
| | 1 |
| | 1 |
| | 1 |
| | 0.018741165 |
| | 1 |
| | 0.00145909 |
| | 1 |
| | 0.816065992 |
| | 2.75468E-06 |
| | 3.79925E-05 |
| | 6.09293E-06 |
| | 1 |
| | 1 |
| | 1 |
| | 5.39257E-06 |
| | 1 |
| | 1 |
| | 9.02978E-05 |
| | 1 |
| | 1 |
| | 0.240567323 |
| | 1.74975E-10 |
| | 1 |

| |
|---|
| 3.41414E-11 |
| 1 |
| 1.50316E-08 |
| 0.713574391 |
| 1 |
| 1 |
| 1 |
| 1 |
| 1 |
| 1 |
| 1 |
| 1 |
| 1 |
| 0.090565618 |
| 1 |
| 0.108823009 |
| 1 |
| 0.02055068 |
| 0.362580401 |
| 1 |
| 0.383407025 |
| 0.005972183 |
| 1 |
| 1 |
| 6.63062E-06 |
| 8.16723E-06 |
| 1 |
| 1 |
| 0.000896301 |
| 1 |
| 1 |
| 1 |
| 1 |
| 1 |
| 1 |

| |
|---|
| 1 |
| 1 |
| 1 |
| 1 |
| 1 |
| 1 |
| 6.59001E-11 |
| 0.011839755 |
| 0.007264819 |
| 0.650479851 |
| 1 |
| 1.47059E-08 |
| 0.098927343 |
| 0.840774286 |
| 8.0531E-10 |
| 0.429548153 |
| 5.85997E-08 |
| 1 |
| 0.025630602 |
| 2.82632E-07 |
| 1 |
| 1 |
| 1 |
| 1 |
| 1 |
| 1 |
| 0.851816189 |
| 0.004823401 |
| 1 |
| 1 |
| 0.000198121 |
| 1 |
| 1 |
| 1 |
| 1 |

| | |
|---|---|
| | 1 |
| | 2.16564E-06 |
| | 1.43565E-06 |
| | 0.01586936 |
| | 1.33418E-05 |
| | 0.000199739 |
| | 0.000545892 |
| | 5.40582E-06 |
| | 1.72384E-06 |
| | 0.054835773 |
| | 1 |
| | 1 |
| | 0.386745542 |
| | 1 |
| | 1 |
| | 1 |
| | 1 |
| | 0.544289917 |
| | 1 |
| | 1 |
| | 1 |
| | 1 |
| | 1 |
| | 1 |
| | 1 |
| | 1 |
| | 1 |
| | 1 |
| | 1 |
| | 1 |
| | 1 |
| | 1 |
| | 0.090146407 |
| | 0.771666917 |
| | 0.13192015 |

| | |
|---|---|
| | 1 |
| | 1 |
| | 1 |
| | 1 |
| | 1 |
| | 1 |
| | 1 |
| | 1.71473E-05 |
| | 1 |
| | 2.28155E-05 |
| | 3.90188E-11 |
| | 3.39836E-08 |
| | 1.96841E-06 |
| | 1 |
| | 1 |
| | 1 |
| | 1 |
| | 1 |
| | 1 |
| | 1 |
| | 0.201739251 |
| | 1 |
| | 0.071496253 |
| | 1 |
| | 1 |
| | 1 |
| | 6.73508E-11 |
| | 0.276871397 |
| | 1 |
| | 1 |
| | 1 |
| | 0.916896804 |
| | 0.945039158 |
| | 1 |
| | 0.373211115 |

| |
|---|
| 1 |
| 1 |
| 0.082444255 |
| 1 |
| 1 |
| 0.001149572 |
| 0.047592195 |
| 1 |
| 0.020549158 |
| 1 |
| 1 |
| 1 |
| 1 |
| 0.076807227 |
| 0.234070938 |

| |
|---|
| Adj. P-value: (Bb_pos_12022024_Newanalysis) / (Rabbitserum_12022024_Newanalysis) |
| 0.179043165 |
| 1 |
| 1.09399E-07 |
| 0.000629595 |
| 0.719313609 |
| 1 |
| 0.021444375 |
| 7.21939E-05 |
| 0.046143831 |
| 0.721707973 |
| 1 |
| 0.205287635 |
| 0.013924117 |
| 0.60814707 |
| 0.235171673 |
| 0.006438322 |
| 1 |
| 8.4353E-12 |
| 1.32902E-05 |
| 1.18127E-10 |
| 1 |
| 8.4353E-12 |
| 7.91209E-05 |
| 8.4353E-12 |
| 8.4353E-12 |
| 1 |
| 1 |
| 1 |
| 0.025291428 |
| 1 |
| 0.609551854 |
| 2.34986E-06 |
| 0.08782126 |

| |
|---|
| 0.151979416 |
| 8.4353E-12 |
| 6.37014E-06 |
| 9.0591E-10 |
| 0.009644549 |
| 0.009927727 |
| 4.77345E-09 |
| 1 |
| 0.000637638 |
| 0.001001942 |
| 0.0007117 |
| 0.900085804 |
| 1.26459E-11 |
| 7.79042E-10 |
| 2.67521E-09 |
| 0.000318194 |
| 4.21254E-05 |
| 0.000105866 |
| 3.53161E-06 |
| 7.01844E-11 |
| 5.07797E-06 |
| 1 |
| 0.713792015 |
| 1.69286E-05 |
| 0.741257755 |
| 0.010196769 |
| 0.771066473 |
| 0.144783112 |
| 1 |
| 0.150705187 |
| 3.48744E-09 |
| 0.000108276 |
| 1 |
| 2.14717E-07 |
| 0.030441409 |

| | |
|---|---|
| | 0.000121061 |
| | 0.197731594 |
| | 0.001864077 |
| | 1 |
| | 1.29112E-05 |
| | 0.000473014 |
| | 0.000365204 |
| | 1 |
| | 8.4353E-12 |
| | 0.001315154 |
| | 8.71968E-12 |
| | 8.4353E-12 |
| | 2.32645E-06 |
| | 1.76631E-07 |
| | 0.653493834 |
| | 1.55775E-06 |
| | 1.13253E-10 |
| | 0.041456721 |
| | 0.519730358 |
| | 2.71388E-05 |
| | 0.105136336 |
| | 0.009657422 |
| | 1.1161E-05 |
| | 7.59483E-11 |
| | 1.76121E-08 |
| | 1.86992E-07 |
| | 9.03375E-08 |
| | 6.68822E-05 |
| | 8.95003E-10 |
| | 6.72992E-07 |
| | 1 |
| | 0.543827749 |
| | 8.24836E-09 |
| | 1 |
| | 6.71818E-05 |

| | |
|---|---|
| | 1 |
| | 2.02921E-05 |
| | 0.047778495 |
| | 0.062238099 |
| | 1 |
| | 0.901915691 |
| | 1 |
| | 0.007333812 |
| | 6.33032E-05 |
| | 0.07087194 |
| | 0.002220189 |
| | 1 |
| | 0.000196475 |
| | 1 |
| | 8.4353E-12 |
| | 0.000702429 |
| | 1 |
| | 1 |
| | 8.4353E-12 |
| | 0.152062643 |
| | 0.000844601 |
| | 0.442127881 |
| | 0.985424301 |
| | 0.052516626 |
| | 0.034856181 |
| | 0.001761012 |
| | 1 |
| | 1 |
| | 2.1758E-05 |
| | 8.48898E-12 |
| | 3.58989E-07 |
| | 0.544784402 |
| | 0.058670684 |
| | 0.000134957 |
| | 0.536748838 |

| |
|---|
| 1 |
| 0.609199976 |
| 0.062180718 |
| 1 |
| 1 |
| 0.969981929 |
| 1 |
| 1 |
| 7.99705E-05 |
| 0.870858799 |
| 0.753219325 |
| 2.37059E-05 |
| 8.4353E-12 |
| 0.085044514 |
| 1.68158E-11 |
| 8.4353E-12 |
| 1 |
| 0.001107468 |
| 0.000177383 |
| 0.014206209 |
| 8.62179E-05 |
| 1.69245E-07 |
| 8.48898E-12 |
| 1.05633E-08 |
| 0.313853215 |
| 8.64207E-05 |
| 1 |
| 0.641251958 |
| 0.005289154 |
| 0.474619663 |
| 2.55321E-06 |
| 0.016134301 |
| 0.588818162 |
| 0.0076153 |
| 3.89342E-07 |

| |
|---:|
| 1 |
| 1.14894E-05 |
| 1.03206E-05 |
| 8.4353E-12 |
| 2.14717E-07 |
| 7.88611E-06 |
| 0.003745485 |
| 2.41283E-07 |
| 0.00027262 |
| 6.55825E-07 |
| 0.310861343 |
| 1 |
| 8.4353E-12 |
| 4.18631E-06 |
| 0.003777636 |
| 1.75252E-07 |
| 5.56416E-06 |
| 0.000249333 |
| 8.4353E-12 |
| 5.06715E-10 |
| 1.1135E-06 |
| 7.81212E-09 |
| 1 |
| 1 |
| 0.002583933 |
| 0.503193006 |
| 0.029308489 |
| 0.682991593 |
| 1.02E-05 |
| 4.6088E-08 |
| 9.27105E-07 |
| 9.5593E-05 |
| 5.16043E-07 |
| 1.08296E-05 |
| 8.80826E-05 |

| | |
|---|---|
| | 2.56821E-06 |
| | 0.004102712 |
| | 0.008171016 |
| | 1 |
| | 0.094965006 |
| | 0.773325416 |
| | 0.002071525 |
| | 0.008534996 |
| | 0.664297632 |
| | 1 |
| | 0.00188714 |
| | 0.90593765 |
| | 0.005687209 |
| | 1 |
| | 0.146454777 |
| | 0.00329114 |
| | 0.009220992 |
| | 8.88955E-09 |
| | 1.65977E-07 |
| | 1 |
| | 1.69022E-07 |
| | 0.224660622 |
| | 1 |
| | 8.64398E-06 |
| | 8.4353E-12 |
| | 6.86609E-06 |
| | 1 |
| | 0.834565988 |
| | 0.030629774 |
| | 1 |
| | 0.480911282 |
| | 1 |
| | 0.161589763 |
| | 0.278117994 |
| | 8.59765E-12 |

| | |
|---|---|
| | 1 |
| | 0.135813212 |
| | 1 |
| | 1 |
| | 0.227063271 |
| | 5.88532E-09 |
| | 2.32136E-06 |
| | 1 |
| | 0.011912139 |
| | 8.4353E-12 |
| | 1 |
| | 0.592517398 |
| | 0.240397769 |
| | 7.55443E-11 |
| | 0.433613829 |
| | 0.610353843 |
| | 1 |
| | 1 |
| | 1 |
| | 8.4353E-12 |
| | 1.73058E-05 |
| | 0.6146761 |
| | 0.916878663 |
| | 1 |
| | 0.975350335 |
| | 1 |
| | 0.042364623 |
| | 1 |
| | 0.210176887 |
| | 0.032654342 |
| | 1.44577E-05 |
| | 1.67791E-10 |
| | 0.019165058 |
| | 1 |
| | 8.4353E-12 |

| | |
|---|---|
| | 4.04395E-09 |
| | 3.67422E-11 |
| | 1.86991E-06 |
| | 0.292124374 |
| | 0.007061873 |
| | 0.089444803 |
| | 8.53175E-12 |
| | 1 |
| | 0.400874636 |
| | 1.32903E-07 |
| | 1 |
| | 0.000351672 |
| | 1 |
| | 1.64023E-05 |
| | 1 |
| | 1 |
| | 3.84566E-10 |
| | 0.003412357 |
| | 0.000225212 |
| | 0.008033362 |
| | 3.86002E-09 |
| | 0.006705313 |
| | 1 |
| | 8.4353E-12 |
| | 0.000237528 |
| | 1.87549E-06 |
| | 0.00830618 |
| | 8.4353E-12 |
| | 1 |
| | 1.91001E-09 |
| | 0.000651108 |
| | 1.77028E-05 |
| | 1 |
| | 0.006524067 |
| | 0.009235311 |

| | |
|---|---|
| | 1.15808E-10 |
| | 1 |
| | 1.46536E-07 |
| | 0.032780398 |
| | 0.012964106 |
| | 1 |
| | 3.24739E-06 |
| | 8.4353E-12 |
| | 1 |
| | 2.11079E-10 |
| | 0.067824232 |
| | 0.001342937 |
| | 0.000809323 |
| | 0.002475036 |
| | 1 |
| | 1 |
| | 0.000113371 |
| | 5.67766E-09 |
| | 8.4353E-12 |
| | 0.000172918 |
| | 0.026162483 |
| | 1 |
| | 0.012410717 |
| | 9.07164E-11 |
| | 1.28876E-05 |
| | 0.322069181 |
| | 0.002199676 |
| | 0.573041849 |
| | 0.00421388 |
| | 0.256392215 |
| | 1 |
| | 0.00027646 |
| | 1 |
| | 2.81442E-07 |
| | 0.046345395 |

| |
|---|
| 0.002954646 |
| 0.845592712 |
| 8.00293E-11 |
| 3.64939E-06 |
| 0.001479173 |
| 0.673351186 |
| 6.47658E-05 |
| 3.76972E-10 |
| 0.000260121 |
| 2.09316E-07 |
| 0.516257049 |
| 1.24301E-09 |
| 3.91124E-06 |
| 0.008195171 |
| 1 |
| 0.003790181 |
| 1 |
| 0.000160703 |
| 0.001436855 |
| 0.00110429 |
| 8.4353E-12 |
| 1.57999E-09 |
| 1 |
| 1.14859E-08 |
| 1.48615E-09 |
| 1 |
| 0.008520875 |
| 5.99022E-05 |
| 0.000897336 |
| 5.02002E-07 |
| 0.655419409 |
| 0.000885431 |
| 1.34094E-05 |
| 6.41757E-05 |
| 1 |

| | |
|---|---|
| | 0.504221322 |
| | 0.361875236 |
| | 0.00046021 |
| | 3.37107E-07 |
| | 0.071938771 |
| | 9.2221E-07 |
| | 1.87974E-07 |
| | 4.06106E-06 |
| | 8.01335E-08 |
| | 1 |
| | 0.000980837 |
| | 8.0469E-10 |
| | 2.18127E-06 |
| | 1.97096E-05 |
| | 1.66893E-06 |
| | 0.000273675 |
| | 0.000212688 |
| | 1.63925E-10 |
| | 1.41333E-11 |
| | 0.006750111 |
| | 0.004561214 |
| | 8.67036E-06 |
| | 0.000810342 |
| | 2.36926E-08 |
| | 1 |
| | 1 |
| | 0.879743862 |
| | 0.004878681 |
| | 1 |
| | 0.017966351 |
| | 3.58306E-11 |
| | 4.38455E-11 |
| | 8.4353E-12 |
| | 3.20129E-07 |
| | 0.002920609 |

| | |
|---|---|
| | 3.06887E-07 |
| | 0.005406812 |
| | 0.006326286 |
| | 0.002046674 |
| | 1 |
| | 1 |
| | 0.090248815 |
| | 0.013186812 |
| | 2.29472E-05 |
| | 0.000100039 |
| | 0.077049005 |
| | 2.73423E-10 |
| | 3.74406E-09 |
| | 4.27359E-05 |
| | 2.25512E-05 |
| | 0.0004474 |
| | 1 |
| | 0.708275908 |
| | 2.86797E-10 |
| | 8.4353E-12 |
| | 0.000398707 |
| | 8.7316E-12 |
| | 3.05288E-06 |
| | 1.13445E-05 |
| | 0.299939637 |
| | 2.65998E-09 |
| | 4.71537E-07 |
| | 7.96069E-11 |
| | 0.022394462 |
| | 1 |
| | 0.025295685 |
| | 1 |
| | 1.34679E-05 |
| | 1.81683E-09 |
| | 2.35544E-11 |

| | |
|---|---|
| | 1.77576E-09 |
| | 1.64846E-05 |
| | 1.17477E-05 |
| | 0.224615299 |
| | 1 |
| | 1 |
| | 2.65416E-08 |
| | 1.92469E-07 |
| | 0.304141078 |
| | 0.00867535 |
| | 0.018668738 |
| | 1 |
| | 0.000166084 |
| | 0.000675841 |
| | 0.035430609 |
| | 8.4353E-12 |
| | 6.91202E-11 |
| | 1.92726E-11 |
| | 0.437333376 |
| | 2.35903E-07 |
| | 1 |
| | 0.055050871 |
| | 0.026369737 |
| | 0.000575265 |
| | 1 |
| | 9.55538E-07 |
| | 5.5166E-05 |
| | 1.26459E-11 |
| | 0.931077892 |
| | 0.002853528 |
| | 0.000484513 |
| | 9.28666E-07 |
| | 0.107785841 |
| | 8.4353E-12 |
| | 0.000548441 |

| |
|---|
| 1 |
| 0.002606925 |
| 0.194438326 |
| 0.117827624 |
| 1 |
| 8.61635E-12 |
| 0.005506114 |
| 1 |
| 0.045390577 |
| 0.831579573 |
| 0.112137585 |
| 0.002043858 |
| 0.021917557 |
| 0.067475926 |
| 0.000959811 |
| 7.83165E-09 |
| 1 |
| 6.61658E-06 |
| 0.054309364 |
| 5.6325E-05 |
| 4.84365E-06 |
| 0.389761836 |
| 0.65127588 |
| 1 |
| 0.009617852 |
| 4.76804E-10 |
| 1 |
| 5.66454E-06 |
| 5.1656E-06 |
| 0.497740058 |
| 2.42208E-06 |
| 8.25697E-07 |
| 1.14708E-08 |
| 0.005711218 |
| 1.94665E-10 |

| | |
|---|---|
| | 0.014676127 |
| | 0.174878712 |
| | 3.4617E-07 |
| | 3.6221E-05 |
| | 8.4353E-12 |
| | 0.040856376 |
| | 2.5845E-08 |
| | 0.613904544 |
| | 8.4353E-12 |
| | 8.7553E-12 |
| | 0.004475205 |
| | 4.35613E-05 |
| | 0.099233734 |
| | 0.046442843 |
| | 1.4098E-06 |
| | 5.32901E-05 |
| | 4.0903E-05 |
| | 2.45821E-11 |
| | 3.18683E-08 |
| | 0.318460335 |
| | 2.3172E-11 |
| | 8.4353E-12 |
| | 8.53175E-12 |
| | 0.024673552 |
| | 6.70182E-08 |
| | 0.026691168 |
| | 8.59765E-12 |
| | 0.220429455 |
| | 6.25833E-06 |
| | 1.83271E-10 |
| | 0.044999237 |
| | 0.002855709 |
| | 0.00077626 |
| | 1.93953E-11 |
| | 0.818688978 |

| | |
|---|---|
| | 0.006100364 |
| | 1 |
| | 1 |
| | 0.071269933 |
| | 0.078989078 |
| | 0.318379246 |
| | 0.865787899 |
| | 0.059954772 |
| | 5.50855E-06 |
| | 1 |
| | 1 |
| | 1 |
| | 6.70536E-06 |
| | 2.29398E-09 |
| | 0.000431188 |
| | 0.000667562 |
| | 1 |
| | 0.046778032 |
| | 0.936425131 |
| | 0.736616442 |
| | 4.2806E-05 |
| | 7.45777E-05 |
| | 1 |
| | 0.001006282 |
| | 0.000524454 |
| | 8.67648E-06 |
| | 0.001181005 |
| | 0.011700861 |
| | 1.60419E-06 |
| | 1 |
| | 1 |
| | 0.00039622 |
| | 0.109912861 |
| | 1 |
| | 1 |

| | |
|---|---|
| | 0.000492944 |
| | 1 |
| | 1 |
| | 0.844808587 |
| | 1 |
| | 0.180986089 |
| | 8.4353E-12 |
| | 2.91632E-09 |
| | 1.37789E-07 |
| | 0.626893903 |
| | 0.009701444 |
| | 7.69224E-05 |
| | 1.896E-09 |
| | 0.023400434 |
| | 8.4353E-12 |
| | 4.7214E-05 |
| | 0.000529543 |
| | 1 |
| | 0.041455772 |
| | 0.621396512 |
| | 1 |
| | 1.17165E-05 |
| | 1 |
| | 1 |
| | 0.048713712 |
| | 0.991201577 |
| | 0.056947312 |
| | 1.00865E-10 |
| | 0.448324281 |
| | 1 |
| | 1.63964E-11 |
| | 0.000946162 |
| | 0.000259907 |
| | 0.22055304 |
| | 0.000134674 |

| |
|---|
| 0.781556326 |
| 1.15771E-07 |
| 1.04332E-09 |
| 1.50434E-05 |
| 1.80757E-07 |
| 2.23257E-06 |
| 1.77782E-06 |
| 8.4353E-12 |
| 4.12036E-09 |
| 0.000156311 |
| 1.18554E-05 |
| 0.433450416 |
| 0.03070583 |
| 1 |
| 0.058811641 |
| 1 |
| 1 |
| 0.001387431 |
| 5.26944E-05 |
| 0.00076803 |
| 1 |
| 1 |
| 1 |
| 0.781942678 |
| 1 |
| 0.888424057 |
| 1 |
| 1 |
| 1 |
| 1 |
| 1 |
| 1 |
| 0.001407801 |
| 9.52663E-10 |
| 0.00135023 |

| | |
|---|---|
| | 0.24135943 |
| | 1 |
| | 1 |
| | 0.000280529 |
| | 0.000113106 |
| | 2.07353E-07 |
| | 0.056776502 |
| | 1.45384E-09 |
| | 0.545080044 |
| | 1.03917E-08 |
| | 8.4353E-12 |
| | 8.59765E-12 |
| | 3.77369E-11 |
| | 1.20807E-05 |
| | 0.000736969 |
| | 0.133760275 |
| | 9.53265E-05 |
| | 1 |
| | 1 |
| | 1 |
| | 0.015451735 |
| | 1 |
| | 1.19007E-05 |
| | 0.950115448 |
| | 0.594360611 |
| | 1.52767E-05 |
| | 5.34579E-05 |
| | 0.026035802 |
| | 0.005860049 |
| | 1 |
| | 0.006379912 |
| | 0.000819211 |
| | 0.307762705 |
| | 1 |
| | 1.14434E-05 |

| |
|---|
| 1 |
| 0.436489058 |
| 1 |
| 0.341473622 |
| 0.044748635 |
| 3.46417E-09 |
| 5.32116E-06 |
| 1 |
| 0.36527513 |
| 1 |
| 1 |
| 0.878078318 |
| 1 |
| 0.00879792 |
| 9.28474E-06 |

| |
|---|
| Adj. P-value: (CM_pos_12022024_Newanalysis) / (Rabbitserum_12022024_Newanalysis) |
| 1 |
| 1 |
| 1.02298E-07 |
| 1 |
| 0.30301488 |
| 1 |
| 0.288057708 |
| 0.540909978 |
| 0.008162035 |
| 0.423847901 |
| 1 |
| 1 |
| 0.637661456 |
| 1 |
| 0.388830912 |
| 0.847459304 |
| 1 |
| 2.19578E-05 |
| 0.001154752 |
| 1.4359E-06 |
| 1 |
| 0.004430312 |
| 1 |
| 3.21604E-11 |
| 0.001020706 |
| 1 |
| 1 |
| 1 |
| 1 |
| 1 |
| 1 |
| 1 |
| 1 |

| | |
|---|---|
| | 1 |
| | 0.044816892 |
| | 2.12972E-08 |
| | 0.00028792 |
| | 1 |
| | 1 |
| | 0.886420968 |
| | 1 |
| | 1.82522E-05 |
| | 1 |
| | 4.71096E-07 |
| | 1 |
| | 0.039946627 |
| | 0.192546222 |
| | 0.290744162 |
| | 1 |
| | 1 |
| | 1 |
| | 1 |
| | 0.466693829 |
| | 0.467547929 |
| | 0.8596726 |
| | 1 |
| | 5.54279E-09 |
| | 1 |
| | 0.11659534 |
| | 0.002427545 |
| | 0.99141528 |
| | 1 |
| | 1 |
| | 0.724035668 |
| | 1 |
| | 1 |
| | 0.153235048 |
| | 0.73018584 |

| | |
|---|---|
| | 0.002722959 |
| | 1 |
| | 0.586049899 |
| | 1 |
| | 0.656339565 |
| | 0.453286119 |
| | 0.001278768 |
| | 1 |
| | 0.009480845 |
| | 0.000335265 |
| | 0.018129698 |
| | 9.17195E-08 |
| | 0.63945157 |
| | 0.005451462 |
| | 0.955506365 |
| | 0.534648524 |
| | 2.47459E-08 |
| | 1 |
| | 1 |
| | 1 |
| | 1 |
| | 1 |
| | 1 |
| | 0.030627031 |
| | 0.035540326 |
| | 0.246859016 |
| | 0.027361375 |
| | 0.199098869 |
| | 0.615111835 |
| | 0.161360814 |
| | 1 |
| | 1 |
| | 1.09388E-05 |
| | 0.249073185 |
| | 1 |

| | |
|---|---|
| | 1 |
| | 0.260145657 |
| | 0.312015264 |
| | 1 |
| | 1 |
| | 0.366622212 |
| | 1 |
| | 1 |
| | 0.002548327 |
| | 0.616202197 |
| | 1 |
| | 1 |
| | 0.043261738 |
| | 1 |
| | 0.00710771 |
| | 1 |
| | 0.984420447 |
| | 1 |
| | 1.78878E-11 |
| | 0.117190177 |
| | 0.22468272 |
| | 1 |
| | 1 |
| | 1 |
| | 0.267528156 |
| | 0.003792084 |
| | 1 |
| | 1 |
| | 0.008615208 |
| | 4.54787E-05 |
| | 0.002410356 |
| | 1 |
| | 1 |
| | 0.929088902 |
| | 1 |

| |
|---|
| 1 |
| 0.081954159 |
| 1 |
| 1 |
| 1 |
| 1 |
| 1 |
| 1 |
| 0.856615068 |
| 0.988719077 |
| 1 |
| 6.60513E-06 |
| 4.8881E-11 |
| 0.003850019 |
| 3.5687E-10 |
| 3.85163E-08 |
| 1 |
| 0.044674867 |
| 0.177482205 |
| 0.00037285 |
| 0.008114859 |
| 1.25087E-05 |
| 6.70563E-11 |
| 0.010344465 |
| 0.423380437 |
| 1 |
| 0.139524234 |
| 1 |
| 0.190178056 |
| 0.163639526 |
| 0.000918441 |
| 1 |
| 0.008997405 |
| 0.152534401 |
| 0.000333118 |

| | |
|---|---|
| | 0.08471304 |
| | 6.87785E-07 |
| | 0.299826067 |
| | 1.78878E-11 |
| | 0.030996251 |
| | 0.00053218 |
| | 1 |
| | 1.81045E-06 |
| | 0.407024667 |
| | 1.02074E-05 |
| | 0.005447723 |
| | 1 |
| | 3.83509E-11 |
| | 0.239808461 |
| | 0.487666359 |
| | 1.74977E-05 |
| | 2.25548E-06 |
| | 0.107929905 |
| | 8.99548E-10 |
| | 1.9806E-06 |
| | 0.00400159 |
| | 3.71423E-05 |
| | 0.824371176 |
| | 1 |
| | 1 |
| | 0.273976377 |
| | 1 |
| | 1 |
| | 0.370244014 |
| | 0.014590711 |
| | 0.370840521 |
| | 0.672708794 |
| | 0.001824428 |
| | 0.003887365 |
| | 0.042742462 |

| | |
|---|---|
| | 1 |
| | 0.01278614 |
| | 0.252132652 |
| | 1 |
| | 1 |
| | 1 |
| | 0.131598855 |
| | 0.006563902 |
| | 0.52773384 |
| | 0.565679319 |
| | 0.319503849 |
| | 1 |
| | 0.876152283 |
| | 1 |
| | 1 |
| | 1 |
| | 1 |
| | 3.85834E-11 |
| | 0.14687707 |
| | 1 |
| | 0.002887573 |
| | 0.041156066 |
| | 1 |
| | 0.000278245 |
| | 3.47538E-09 |
| | 0.006115871 |
| | 1 |
| | 1 |
| | 1 |
| | 1 |
| | 0.052100553 |
| | 1 |
| | 0.005556678 |
| | 0.183594244 |
| | 1.20713E-06 |

| | |
|---|---|
| | 0.174614684 |
| | 0.274012416 |
| | 1 |
| | 1 |
| | 0.502074821 |
| | 2.11603E-05 |
| | 0.001382954 |
| | 1 |
| | 0.460430479 |
| | 4.43326E-08 |
| | 0.012824985 |
| | 0.087060598 |
| | 0.029013144 |
| | 2.4187E-07 |
| | 0.117968854 |
| | 0.000198509 |
| | 0.007249214 |
| | 0.135245017 |
| | 1 |
| | 0.000445131 |
| | 0.001607966 |
| | 0.000274503 |
| | 0.044100011 |
| | 0.038664981 |
| | 1 |
| | 1 |
| | 0.564959149 |
| | 0.638267973 |
| | 7.85993E-05 |
| | 0.00852624 |
| | 0.008424112 |
| | 0.011635461 |
| | 1 |
| | 1 |
| | 4.5701E-10 |

| |
|---|
| 7.04159E-08 |
| 0.000201339 |
| 0.001025267 |
| 0.093620591 |
| 0.00687692 |
| 0.952954774 |
| 1.35885E-07 |
| 1 |
| 1 |
| 7.18679E-07 |
| 0.427932454 |
| 9.54137E-05 |
| 1 |
| 3.49969E-08 |
| 1 |
| 1 |
| 3.95702E-06 |
| 0.000371277 |
| 1 |
| 0.007576211 |
| 1 |
| 2.56353E-08 |
| 1 |
| 9.00168E-11 |
| 0.000113695 |
| 1.91283E-06 |
| 0.117022343 |
| 9.97737E-08 |
| 0.11297126 |
| 0.000545934 |
| 0.187592847 |
| 0.093854705 |
| 0.608618824 |
| 1 |
| 1 |

| | |
|---|---|
| | 6.06263E-08 |
| | 0.695199834 |
| | 0.102310109 |
| | 0.027332267 |
| | 0.073397961 |
| | 1 |
| | 0.018290654 |
| | 1.0386E-08 |
| | 1 |
| | 1.3933E-06 |
| | 1 |
| | 0.090910298 |
| | 1 |
| | 1 |
| | 1 |
| | 1 |
| | 0.000215792 |
| | 0.013056472 |
| | 1.05858E-05 |
| | 1.35758E-08 |
| | 1 |
| | 0.005238515 |
| | 0.920313335 |
| | 2.77143E-05 |
| | 0.004456095 |
| | 1 |
| | 0.899485698 |
| | 0.00029654 |
| | 0.002484346 |
| | 0.100542448 |
| | 0.022546156 |
| | 0.081598977 |
| | 1.04834E-05 |
| | 0.070154353 |
| | 1 |

| | |
|---|---|
| | 0.158968927 |
| | 1 |
| | 0.075025258 |
| | 1 |
| | 0.001081829 |
| | 0.577707047 |
| | 1.51366E-05 |
| | 1.62661E-08 |
| | 0.000689943 |
| | 0.000624437 |
| | 3.34113E-05 |
| | 1.019E-07 |
| | 5.97859E-06 |
| | 1 |
| | 1 |
| | 1 |
| | 0.069392524 |
| | 5.24031E-05 |
| | 1 |
| | 1.31122E-05 |
| | 1.59951E-07 |
| | 1.72525E-06 |
| | 1 |
| | 7.01057E-05 |
| | 0.00109 |
| | 1 |
| | 0.947309391 |
| | 0.021944179 |
| | 0.025628228 |
| | 0.001432407 |
| | 1 |
| | 0.066939943 |
| | 0.000144518 |
| | 1 |
| | 0.78974501 |

| | |
|---|---|
| | 0.186163353 |
| | 0.980065433 |
| | 1.41146E-06 |
| | 1.62198E-07 |
| | 1 |
| | 9.12268E-07 |
| | 7.38498E-06 |
| | 3.02243E-05 |
| | 0.000244603 |
| | 0.430743046 |
| | 0.008337088 |
| | 4.8806E-10 |
| | 1.71573E-06 |
| | 0.176095334 |
| | 0.014319427 |
| | 0.415353522 |
| | 0.41282384 |
| | 0.021099578 |
| | 0.000142155 |
| | 0.978859448 |
| | 0.887577678 |
| | 0.102273295 |
| | 0.007609735 |
| | 1.44298E-08 |
| | 1 |
| | 1 |
| | 0.240395059 |
| | 1 |
| | 1 |
| | 1 |
| | 0.000445394 |
| | 0.082527749 |
| | 1.5422E-07 |
| | 0.184839777 |
| | 0.821562264 |

| | |
|---|---|
| | 0.009699299 |
| | 0.883533867 |
| | 0.013822695 |
| | 1 |
| | 0.595614066 |
| | 0.157368989 |
| | 0.680721308 |
| | 0.005058066 |
| | 5.29436E-06 |
| | 1.88667E-05 |
| | 0.038664981 |
| | 4.97863E-05 |
| | 0.011803226 |
| | 1 |
| | 2.05694E-07 |
| | 0.003604843 |
| | 0.022298795 |
| | 0.186122667 |
| | 0.000190023 |
| | 1.04814E-05 |
| | 0.008075695 |
| | 0.00072903 |
| | 0.073778766 |
| | 0.865484345 |
| | 0.206268414 |
| | 0.022226912 |
| | 4.8488E-07 |
| | 8.88448E-05 |
| | 1 |
| | 1 |
| | 6.41741E-05 |
| | 1 |
| | 0.010115993 |
| | 0.009716969 |
| | 0.000371461 |

| | |
|---|---|
| | 0.003146214 |
| | 0.006516768 |
| | 1.23103E-07 |
| | 1 |
| | 1 |
| | 0.769424018 |
| | 0.040425769 |
| | 0.000930228 |
| | 0.03450957 |
| | 3.85908E-06 |
| | 0.02637535 |
| | 0.459503759 |
| | 6.41431E-08 |
| | 0.511029062 |
| | 0.104794553 |
| | 0.109340115 |
| | 1.21813E-05 |
| | 1 |
| | 1 |
| | 4.78673E-07 |
| | 1 |
| | 0.046377425 |
| | 2.67708E-06 |
| | 3.5604E-06 |
| | 0.000204314 |
| | 3.89036E-05 |
| | 0.021295731 |
| | 5.06053E-11 |
| | 1 |
| | 0.002476471 |
| | 1.52637E-05 |
| | 1.28162E-06 |
| | 0.000180763 |
| | 1 |
| | 1 |

| |
|---|
| 1 |
| 1 |
| 1 |
| 1 |
| 0.000603352 |
| 1.78878E-11 |
| 0.168502027 |
| 1 |
| 0.548744464 |
| 0.045974454 |
| 0.229930072 |
| 0.005606016 |
| 0.026726973 |
| 0.269959365 |
| 0.010559277 |
| 0.005036538 |
| 2.33234E-06 |
| 5.60154E-08 |
| 0.528252036 |
| 0.000219294 |
| 0.000682178 |
| 0.087127183 |
| 0.329462255 |
| 1 |
| 0.073468414 |
| 2.71583E-09 |
| 0.253089462 |
| 1.38659E-06 |
| 2.57618E-05 |
| 0.942791804 |
| 0.02387852 |
| 0.061547856 |
| 2.47267E-08 |
| 1 |
| 1.58334E-07 |

| |
|---|
| 0.460490037 |
| 0.038399772 |
| 4.65785E-07 |
| 0.022785336 |
| 3.80746E-10 |
| 0.069571417 |
| 0.000212806 |
| 0.089878512 |
| 4.63137E-07 |
| 1.85127E-08 |
| 0.027471645 |
| 0.000839271 |
| 0.005617347 |
| 0.568109775 |
| 1.42445E-06 |
| 0.776037441 |
| 0.004351721 |
| 8.94489E-07 |
| 3.70159E-06 |
| 1 |
| 0.00054233 |
| 3.28571E-10 |
| 5.09617E-06 |
| 0.007090188 |
| 1.14295E-06 |
| 0.021346852 |
| 7.61372E-06 |
| 0.075969391 |
| 2.08368E-05 |
| 0.000298246 |
| 0.009836296 |
| 0.175176136 |
| 5.2859E-07 |
| 1 |
| 1 |

| | |
|---|---|
| | 1.80046E-11 |
| | 1 |
| | 1.019E-07 |
| | 1 |
| | 1 |
| | 1 |
| | 1 |
| | 0.915980125 |
| | 3.60255E-05 |
| | 1 |
| | 1 |
| | 1 |
| | 0.000310019 |
| | 6.47392E-06 |
| | 0.062374272 |
| | 0.989857416 |
| | 1 |
| | 1 |
| | 1 |
| | 1 |
| | 0.0524837 |
| | 1 |
| | 0.155566098 |
| | 0.005168387 |
| | 0.590164631 |
| | 1 |
| | 0.18212402 |
| | 0.275587409 |
| | 0.584524787 |
| | 1 |
| | 1 |
| | 0.003958668 |
| | 0.073413252 |
| | 1 |
| | 1 |

| | |
|---|---|
| | 0.026816792 |
| | 1 |
| | 1 |
| | 0.587912437 |
| | 1 |
| | 0.002343549 |
| | 9.6392E-08 |
| | 7.89787E-05 |
| | 0.012533443 |
| | 1 |
| | 0.154369146 |
| | 0.00793178 |
| | 4.65575E-06 |
| | 0.000104139 |
| | 4.66881E-08 |
| | 0.049405012 |
| | 0.006474043 |
| | 1 |
| | 1 |
| | 9.96011E-06 |
| | 1 |
| | 0.00121951 |
| | 0.908671695 |
| | 1 |
| | 0.28804756 |
| | 1 |
| | 1 |
| | 2.36988E-06 |
| | 0.01606839 |
| | 1 |
| | 2.4828E-06 |
| | 0.012748588 |
| | 0.000612771 |
| | 0.537149658 |
| | 0.000324116 |

| |
|---|
| 1 |
| 1 |
| 0.204672439 |
| 0.423730387 |
| 1 |
| 1 |
| 0.787261281 |
| 7.29068E-07 |
| 0.545611754 |
| 0.760167262 |
| 0.000915443 |
| 1 |
| 1 |
| 1 |
| 0.478770706 |
| 1 |
| 1 |
| 0.50268485 |
| 0.002822272 |
| 0.017669944 |
| 1 |
| 1 |
| 1 |
| 0.856632138 |
| 1 |
| 1 |
| 1 |
| 1 |
| 1 |
| 1 |
| 1 |
| 1 |
| 1 |
| 1.52215E-07 |
| 1 |

| | |
|---|---|
| | 0.471571579 |
| | 1 |
| | 1 |
| | 0.000552054 |
| | 0.007358532 |
| | 6.748E-08 |
| | 0.925550309 |
| | 0.027096953 |
| | 1 |
| | 0.168474764 |
| | 0.000173093 |
| | 0.002929953 |
| | 0.001913804 |
| | 0.000732182 |
| | 0.001107033 |
| | 0.722722437 |
| | 2.47427E-05 |
| | 1 |
| | 1 |
| | 1 |
| | 1 |
| | 0.122849313 |
| | 0.108040943 |
| | 1 |
| | 0.12514382 |
| | 0.000594788 |
| | 4.22609E-06 |
| | 1 |
| | 0.006470988 |
| | 1 |
| | 0.067964783 |
| | 0.185853332 |
| | 1 |
| | 1 |
| | 0.015431945 |

| |
|---|
| 0.402888171 |
| 1 |
| 0.372738552 |
| 1 |
| 0.835490987 |
| 0.00108201 |
| 0.077991527 |
| 1 |
| 0.916128646 |
| 1 |
| 1 |
| 1 |
| 1 |
| 1 |
| 0.023260335 |

| |
|---|
| Adj. P-value: (SM_pos_12022024_Newanalyis) / (Rabbitserum_12022024_Newanalysis) |
| 1 |
| 0.057207246 |
| 3.21527E-07 |
| 1 |
| 3.75341E-05 |
| 1 |
| 0.605480799 |
| 0.001104524 |
| 0.678572431 |
| 0.009155739 |
| 0.007131322 |
| 1 |
| 1 |
| 0.354075524 |
| 0.052042248 |
| 1 |
| 0.86623599 |
| 6.47189E-12 |
| 2.49127E-08 |
| 1.90867E-11 |
| 0.453203711 |
| 1.78209E-10 |
| 0.129689814 |
| 6.47189E-12 |
| 6.47189E-12 |
| 0.441051346 |
| 1 |
| 0.638889914 |
| 0.010991217 |
| 0.005198503 |
| 1 |
| 1 |
| 0.000369817 |

| | |
|---|---|
| | 0.003174444 |
| | 2.3124E-07 |
| | 4.81423E-07 |
| | 2.03125E-09 |
| | 1 |
| | 1 |
| | 1.61784E-08 |
| | 0.002609242 |
| | 1 |
| | 1 |
| | 0.313819933 |
| | 1.80174E-05 |
| | 5.52E-08 |
| | 2.15064E-05 |
| | 2.0437E-07 |
| | 5.6738E-05 |
| | 0.011897978 |
| | 1.75003E-05 |
| | 0.000764078 |
| | 5.22775E-07 |
| | 0.001798544 |
| | 5.35637E-05 |
| | 0.351182522 |
| | 0.682085358 |
| | 0.488843635 |
| | 0.367184162 |
| | 1.39369E-07 |
| | 1 |
| | 0.008740858 |
| | 0.003965239 |
| | 4.45239E-05 |
| | 4.42485E-05 |
| | 0.000210524 |
| | 0.951202972 |
| | 0.000795823 |

| |
|---|
| 1 |
| 1 |
| 0.002214424 |
| 4.92875E-07 |
| 1 |
| 0.002292986 |
| 1 |
| 0.002419995 |
| 6.48861E-08 |
| 6.49716E-07 |
| 7.22442E-12 |
| 6.47189E-12 |
| 2.41672E-05 |
| 4.6777E-08 |
| 1.34786E-05 |
| 1 |
| 5.59584E-06 |
| 1 |
| 0.920673905 |
| 0.000216541 |
| 0.078261697 |
| 1 |
| 1.31493E-10 |
| 2.55067E-11 |
| 0.001855975 |
| 0.007936833 |
| 8.83381E-06 |
| 1 |
| 2.38655E-07 |
| 0.002559342 |
| 0.647264635 |
| 0.819829145 |
| 1.74172E-07 |
| 8.88831E-06 |
| 8.38111E-05 |

| | |
|---|---|
| | 0.137163013 |
| | 9.80416E-05 |
| | 5.50807E-07 |
| | 1 |
| | 1 |
| | 0.200347863 |
| | 0.375072973 |
| | 0.154618232 |
| | 2.32804E-05 |
| | 1 |
| | 1 |
| | 6.68664E-06 |
| | 1 |
| | 0.599959481 |
| | 6.47189E-12 |
| | 6.39169E-06 |
| | 0.091929673 |
| | 0.003508527 |
| | 6.47189E-12 |
| | 0.005230247 |
| | 1 |
| | 0.02255293 |
| | 0.510937137 |
| | 1 |
| | 2.73443E-10 |
| | 1 |
| | 1 |
| | 0.951552144 |
| | 0.001323253 |
| | 6.47189E-12 |
| | 7.8242E-08 |
| | 1.06531E-10 |
| | 0.421331759 |
| | 0.344965886 |
| | 1 |

| |
|---|
| 0.004249694 |
| 1 |
| 0.456276384 |
| 1 |
| 1 |
| 1 |
| 1 |
| 1 |
| 1 |
| 0.954859275 |
| 1 |
| 0.006944115 |
| 4.01261E-10 |
| 0.001454651 |
| 4.02636E-09 |
| 9.97166E-07 |
| 0.424352787 |
| 1 |
| 0.79237611 |
| 0.005927109 |
| 1 |
| 0.00605987 |
| 4.41455E-10 |
| 0.001296371 |
| 9.77084E-05 |
| 1.22935E-06 |
| 4.30198E-07 |
| 1 |
| 1 |
| 0.000431846 |
| 0.005722121 |
| 1 |
| 3.30658E-08 |
| 1 |
| 0.000781542 |

| |
|---|
| 1 |
| 6.2251E-10 |
| 0.000136867 |
| 2.19386E-11 |
| 0.001961518 |
| 1.5318E-05 |
| 0.896446577 |
| 0.045254131 |
| 0.013392314 |
| 0.774031704 |
| 2.43948E-06 |
| 6.88854E-09 |
| 1.97675E-11 |
| 0.124393825 |
| 1 |
| 0.055269985 |
| 0.000846336 |
| 0.699093666 |
| 3.62027E-08 |
| 2.53094E-07 |
| 0.042670185 |
| 0.000928886 |
| 6.22988E-05 |
| 0.003758928 |
| 0.241617602 |
| 0.422188147 |
| 0.197640886 |
| 0.726853173 |
| 0.12502529 |
| 2.02614E-09 |
| 1 |
| 2.75298E-06 |
| 0.010019322 |
| 1.72347E-08 |
| 0.838893404 |

| | |
|---|---|
| | 0.013091277 |
| | 1 |
| | 1 |
| | 0.873428755 |
| | 1 |
| | 3.23575E-07 |
| | 3.44183E-05 |
| | 1 |
| | 0.000312837 |
| | 0.764473151 |
| | 1 |
| | 1 |
| | 1 |
| | 1 |
| | 1 |
| | 1 |
| | 0.00086132 |
| | 6.47189E-12 |
| | 3.07199E-06 |
| | 0.493077247 |
| | 0.105412809 |
| | 0.025899345 |
| | 1 |
| | 1 |
| | 2.40747E-08 |
| | 2.42611E-05 |
| | 0.330802976 |
| | 0.144900575 |
| | 0.87020151 |
| | 0.001833071 |
| | 0.965562945 |
| | 0.257362678 |
| | 0.033715692 |
| | 0.054013892 |
| | 6.73981E-10 |

| | |
|---|---|
| | 1 |
| | 0.414397086 |
| | 1 |
| | 1 |
| | 0.021665767 |
| | 0.604818041 |
| | 0.005696465 |
| | 1 |
| | 1 |
| | 1.7309E-06 |
| | 0.196613259 |
| | 1 |
| | 0.01512839 |
| | 8.3927E-09 |
| | 0.881330279 |
| | 0.228278137 |
| | 9.37278E-07 |
| | 1 |
| | 0.447911375 |
| | 6.47189E-12 |
| | 0.106812112 |
| | 0.497065774 |
| | 0.000395955 |
| | 0.008672462 |
| | 0.001223714 |
| | 1 |
| | 0.472437544 |
| | 0.83100384 |
| | 1 |
| | 0.079765795 |
| | 0.432836484 |
| | 4.61988E-06 |
| | 1 |
| | 1 |
| | 5.70405E-09 |

| | |
|---|---|
| | 1.79684E-05 |
| | 6.45874E-10 |
| | 0.184568349 |
| | 1 |
| | 0.157734383 |
| | 1 |
| | 2.72375E-07 |
| | 1 |
| | 0.474914313 |
| | 8.54498E-06 |
| | 0.095417215 |
| | 5.39491E-05 |
| | 1 |
| | 0.025824803 |
| | 0.83032427 |
| | 0.143866888 |
| | 1.05388E-09 |
| | 9.79763E-05 |
| | 1 |
| | 0.002662431 |
| | 2.04818E-07 |
| | 6.50156E-11 |
| | 0.707605127 |
| | 1.62113E-08 |
| | 0.000204554 |
| | 4.51904E-07 |
| | 0.096135294 |
| | 1.09889E-09 |
| | 0.002050871 |
| | 0.000416495 |
| | 0.000273461 |
| | 0.001887285 |
| | 1 |
| | 0.007357751 |
| | 1 |

| |
|---|
| 1.23948E-05 |
| 0.010028635 |
| 1.99122E-05 |
| 7.26149E-05 |
| 0.045142239 |
| 0.812561398 |
| 0.545292168 |
| 7.25829E-11 |
| 8.8691E-06 |
| 2.46972E-05 |
| 1 |
| 0.003898514 |
| 0.636963876 |
| 1 |
| 0.000284449 |
| 1 |
| 0.001545248 |
| 1.80287E-07 |
| 3.70392E-10 |
| 7.22799E-05 |
| 5.69483E-05 |
| 0.441884108 |
| 1 |
| 1.9549E-07 |
| 0.000602024 |
| 5.29369E-05 |
| 0.003590126 |
| 0.000117423 |
| 0.044803204 |
| 0.049291521 |
| 0.416758544 |
| 1 |
| 0.000835113 |
| 1 |
| 1 |

| | |
|---|---|
| | 0.003324786 |
| | 0.943186932 |
| | 1 |
| | 0.002749492 |
| | 0.001459825 |
| | 1 |
| | 1.0142E-06 |
| | 8.62033E-06 |
| | 0.003050203 |
| | 0.001008503 |
| | 1.40925E-06 |
| | 2.68533E-07 |
| | 7.20361E-05 |
| | 0.000727814 |
| | 0.784197086 |
| | 7.34397E-07 |
| | 0.019352493 |
| | 0.000178444 |
| | 0.537780372 |
| | 0.001672928 |
| | 7.50452E-08 |
| | 0.008097277 |
| | 1 |
| | 1.10843E-07 |
| | 1.80066E-05 |
| | 1 |
| | 0.669979636 |
| | 0.000896944 |
| | 0.000549834 |
| | 1 |
| | 1 |
| | 2.68429E-07 |
| | 6.33405E-07 |
| | 0.000662331 |
| | 1.42895E-08 |

| |
|---|
| 1 |
| 0.001867282 |
| 1.7899E-10 |
| 2.31167E-07 |
| 2.73569E-05 |
| 6.05883E-07 |
| 1.30263E-07 |
| 5.30217E-05 |
| 1.72384E-11 |
| 0.003100949 |
| 0.012425155 |
| 4.62028E-10 |
| 1.88619E-07 |
| 5.53045E-05 |
| 1 |
| 0.000495924 |
| 0.014316027 |
| 0.002405569 |
| 4.19397E-06 |
| 0.758747177 |
| 0.130136918 |
| 1 |
| 0.000123689 |
| 2.03933E-07 |
| 1 |
| 1 |
| 1.72457E-09 |
| 0.07707436 |
| 1 |
| 0.02050968 |
| 2.23104E-10 |
| 1 |
| 0.007558286 |
| 1 |
| 1 |

| | |
|---|---|
| | 0.003463265 |
| | 0.000356934 |
| | 1 |
| | 1 |
| | 0.51931329 |
| | 1 |
| | 2.19214E-08 |
| | 7.82337E-06 |
| | 3.73743E-05 |
| | 7.84079E-07 |
| | 0.003399278 |
| | 5.43609E-05 |
| | 9.44583E-05 |
| | 0.002232652 |
| | 6.29223E-05 |
| | 1 |
| | 6.28118E-09 |
| | 7.95435E-05 |
| | 1 |
| | 0.768150418 |
| | 0.001259716 |
| | 0.002605183 |
| | 0.193087215 |
| | 1 |
| | 0.510948406 |
| | 4.79543E-06 |
| | 0.001888831 |
| | 3.09276E-10 |
| | 1 |
| | 0.00597296 |
| | 0.188821644 |
| | 1 |
| | 0.012212438 |
| | 0.000104303 |
| | 0.002460219 |

| | |
|---|---|
| | 3.39711E-09 |
| | 9.7409E-07 |
| | 1.07015E-05 |
| | 0.794589606 |
| | 0.028974087 |
| | 1 |
| | 1 |
| | 1 |
| | 0.006044967 |
| | 0.001683989 |
| | 0.796205214 |
| | 0.008427706 |
| | 4.43127E-10 |
| | 1 |
| | 0.037510104 |
| | 1 |
| | 0.000250995 |
| | 0.017994384 |
| | 1 |
| | 2.93098E-06 |
| | 0.013666485 |
| | 1 |
| | 1.53189E-07 |
| | 1.61246E-06 |
| | 0.000226275 |
| | 4.24757E-06 |
| | 1 |
| | 9.30431E-12 |
| | 0.018159384 |
| | 0.00808524 |
| | 3.02831E-06 |
| | 4.51994E-08 |
| | 0.000316053 |
| | 1 |
| | 0.055585822 |

| |
|---|
| 1 |
| 0.016005654 |
| 1 |
| 0.580901992 |
| 0.084668674 |
| 2.21865E-08 |
| 0.194945116 |
| 0.465235001 |
| 1.17174E-05 |
| 0.172663833 |
| 0.001263233 |
| 0.948308505 |
| 0.024631713 |
| 1 |
| 0.773467785 |
| 0.000105652 |
| 0.004730229 |
| 1 |
| 0.001309736 |
| 0.075208645 |
| 0.464887306 |
| 0.12492042 |
| 0.000589059 |
| 1 |
| 1 |
| 1.10392E-07 |
| 1 |
| 1.16202E-10 |
| 5.90429E-05 |
| 0.968973434 |
| 0.371145546 |
| 0.001477645 |
| 1.03163E-06 |
| 0.066440163 |
| 3.51397E-07 |

| |
|---|
| 1 |
| 1 |
| 1 |
| 0.080537767 |
| 0.002100673 |
| 0.019365766 |
| 1.07473E-06 |
| 1 |
| 6.47189E-12 |
| 3.26254E-06 |
| 0.876581563 |
| 0.682145041 |
| 1 |
| 0.405462094 |
| 6.42589E-07 |
| 0.469080653 |
| 0.095371196 |
| 0.028510213 |
| 0.000963755 |
| 1 |
| 0.296934732 |
| 6.47189E-12 |
| 0.000225089 |
| 0.000730632 |
| 0.001448507 |
| 0.000197418 |
| 9.58716E-11 |
| 1 |
| 6.72754E-09 |
| 0.559009312 |
| 0.711264837 |
| 0.003812217 |
| 2.27794E-07 |
| 0.007899599 |
| 0.002107787 |

| | |
|---|---|
| | 6.47189E-12 |
| | 0.665155525 |
| | 4.42688E-06 |
| | 1 |
| | 1 |
| | 1 |
| | 1 |
| | 1 |
| | 0.005592551 |
| | 0.195011056 |
| | 1 |
| | 1 |
| | 0.23356109 |
| | 2.41354E-08 |
| | 0.240024489 |
| | 0.282305254 |
| | 4.07489E-09 |
| | 0.130741408 |
| | 0.622315192 |
| | 1 |
| | 1.91869E-05 |
| | 0.017958756 |
| | 1 |
| | 0.004291707 |
| | 0.046186724 |
| | 1 |
| | 1.24243E-07 |
| | 1 |
| | 0.177647259 |
| | 0.446309749 |
| | 0.669455657 |
| | 0.171205107 |
| | 0.00953702 |
| | 1 |
| | 0.579180587 |

| | |
|---|---|
| | 0.000620402 |
| | 1 |
| | 1 |
| | 1 |
| | 0.820859991 |
| | 4.84001E-07 |
| | 6.47189E-12 |
| | 4.43778E-07 |
| | 0.020620456 |
| | 7.62796E-10 |
| | 4.69719E-05 |
| | 0.225199107 |
| | 3.71951E-09 |
| | 0.79373368 |
| | 1.37837E-09 |
| | 0.123856431 |
| | 0.023995177 |
| | 1 |
| | 1.5879E-05 |
| | 0.008630607 |
| | 1 |
| | 0.020736197 |
| | 1 |
| | 0.005353051 |
| | 0.000586483 |
| | 1 |
| | 7.03266E-05 |
| | 9.99501E-10 |
| | 0.662620342 |
| | 1 |
| | 5.98167E-09 |
| | 0.998629499 |
| | 0.004093186 |
| | 3.58377E-05 |
| | 9.9038E-08 |

| | |
|---|---|
| | 0.017952828 |
| | 2.84049E-09 |
| | 5.12034E-09 |
| | 1 |
| | 4.94813E-10 |
| | 0.000549549 |
| | 0.790518576 |
| | 7.20752E-10 |
| | 1 |
| | 0.049769628 |
| | 6.47189E-12 |
| | 1 |
| | 1 |
| | 0.000218709 |
| | 0.007755053 |
| | 0.013826968 |
| | 0.940596836 |
| | 1 |
| | 1 |
| | 1 |
| | 0.005289169 |
| | 1 |
| | 1 |
| | 0.997648563 |
| | 1 |
| | 1 |
| | 1 |
| | 0.060535857 |
| | 1 |
| | 0.029589658 |
| | 0.008217831 |
| | 0.120965671 |
| | 0.747617485 |
| | 6.47189E-12 |
| | 1 |

| | |
|---|---|
| | 0.021795795 |
| | 1 |
| | 2.83711E-07 |
| | 0.49553053 |
| | 2.73399E-05 |
| | 1.07641E-09 |
| | 1 |
| | 1.72318E-10 |
| | 1 |
| | 6.17947E-05 |
| | 6.47189E-12 |
| | 6.47189E-12 |
| | 6.53975E-11 |
| | 3.53377E-06 |
| | 1.81937E-05 |
| | 1 |
| | 1.35981E-06 |
| | 0.106236506 |
| | 0.525838523 |
| | 0.09235746 |
| | 1 |
| | 1 |
| | 0.00018872 |
| | 0.013225621 |
| | 0.003503385 |
| | 0.012684397 |
| | 9.55646E-08 |
| | 1 |
| | 1 |
| | 1 |
| | 2.83408E-06 |
| | 0.415984815 |
| | 1 |
| | 0.019615783 |
| | 0.091624605 |

| |
|---|
| 1 |
| 1 |
| 1 |
| 1 |
| 0.102161507 |
| 2.30457E-05 |
| 0.000457764 |
| 1 |
| 0.019076229 |
| 1 |
| 1 |
| 0.587363342 |
| 1 |
| 0.564492211 |
| 0.510692608 |

| Adj. P-value: (Bb_pos_12022024_Newanalysis) / (SM_pos_12022024_Newanalyis) |
|---:|
| 0.581839143 |
| 0.013131087 |
| 1 |
| 0.010730755 |
| 0.005421576 |
| 0.848748247 |
| 0.000134473 |
| 1 |
| 0.00039272 |
| 0.465162478 |
| 0.016021774 |
| 0.728689549 |
| 0.046334706 |
| 0.004538509 |
| 1 |
| 0.027870223 |
| 1 |
| 0.152559682 |
| 0.305448957 |
| 1 |
| 0.172153945 |
| 0.02944133 |
| 0.077933027 |
| 0.00802714 |
| 3.41871E-06 |
| 0.713789742 |
| 1 |
| 1 |
| 1.44737E-06 |
| 0.019656152 |
| 1 |
| 1.96362E-06 |
| 0.491953195 |

| |
|---|
| 0.929942821 |
| 1.90321E-07 |
| 1 |
| 1 |
| 0.002311836 |
| 0.000444653 |
| 1 |
| 0.018448171 |
| 0.01175338 |
| 0.020139977 |
| 0.183397439 |
| 0.001502392 |
| 0.00098811 |
| 0.001500849 |
| 0.395052058 |
| 1 |
| 0.346310947 |
| 1 |
| 0.371178266 |
| 0.002328694 |
| 0.276035445 |
| 0.00052651 |
| 0.006242901 |
| 0.001699556 |
| 1 |
| 0.808679778 |
| 1.55004E-05 |
| 0.137814001 |
| 0.004453402 |
| 3.91712E-06 |
| 0.004716367 |
| 1 |
| 0.000125575 |
| 7.71403E-06 |
| 1.7616E-07 |

| | |
|---|---|
| | 3.97193E-06 |
| | 0.291965585 |
| | 3.47932E-08 |
| | 9.80141E-07 |
| | 1.36528E-05 |
| | 1.17589E-08 |
| | 3.37806E-05 |
| | 9.70603E-05 |
| | 2.06979E-08 |
| | 0.138945466 |
| | 1 |
| | 0.001763454 |
| | 1 |
| | 1.40652E-11 |
| | 0.002495699 |
| | 6.20836E-07 |
| | 0.000410125 |
| | 0.036701104 |
| | 1 |
| | 1 |
| | 4.92472E-05 |
| | 0.048874128 |
| | 0.000887608 |
| | 1 |
| | 0.000958804 |
| | 0.003341224 |
| | 0.456717064 |
| | 0.001566987 |
| | 0.131369646 |
| | 0.036140918 |
| | 1 |
| | 1 |
| | 0.873874587 |
| | 0.000216281 |
| | 2.59038E-10 |

| |
|---|
| 0.813837789 |
| 1 |
| 0.003841866 |
| 0.01050599 |
| 0.232672488 |
| 1 |
| 1 |
| 6.20615E-06 |
| 1 |
| 0.182335378 |
| 0.000106447 |
| 0.000200097 |
| 0.000126856 |
| 0.087050726 |
| 1 |
| 0.856857867 |
| 0.480047453 |
| 0.006374336 |
| 0.060181762 |
| 1 |
| 4.7105E-05 |
| 9.78564E-05 |
| 1 |
| 0.219373293 |
| 8.44228E-07 |
| 0.00060827 |
| 1 |
| 1 |
| 0.779285044 |
| 0.004094777 |
| 1 |
| 1.17915E-08 |
| 0.000220212 |
| 0.040190623 |
| 0.258433039 |

| | |
|---|---|
| | 0.00098811 |
| | 0.910800574 |
| | 0.00026888 |
| | 1 |
| | 1 |
| | 1 |
| | 0.945067665 |
| | 1 |
| | 0.000202368 |
| | 1 |
| | 1 |
| | 0.335967568 |
| | 1.05353E-07 |
| | 0.921573396 |
| | 0.040662179 |
| | 5.95704E-09 |
| | 0.720220149 |
| | 6.07082E-05 |
| | 2.11679E-06 |
| | 1 |
| | 0.000618071 |
| | 0.003896285 |
| | 0.024779671 |
| | 0.000730977 |
| | 3.87522E-07 |
| | 0.993767579 |
| | 4.6567E-06 |
| | 0.173505666 |
| | 0.076300919 |
| | 0.099235005 |
| | 0.063840878 |
| | 0.300942079 |
| | 5.82086E-06 |
| | 0.016244049 |
| | 0.059342399 |

| |
|---|
| 1 |
| 0.007000928 |
| 1 |
| 4.23061E-07 |
| 0.014541977 |
| 1 |
| 0.144792366 |
| 0.000735491 |
| 0.876820444 |
| 4.25198E-05 |
| 0.001648565 |
| 1.64147E-08 |
| 1.95074E-09 |
| 0.004866589 |
| 0.006683503 |
| 0.000417801 |
| 0.469477432 |
| 2.24152E-06 |
| 1.40652E-11 |
| 0.066423061 |
| 0.004050467 |
| 0.000705541 |
| 2.0596E-05 |
| 0.005082506 |
| 3.93793E-06 |
| 1 |
| 1 |
| 0.019436642 |
| 0.012011093 |
| 1 |
| 3.10442E-06 |
| 1 |
| 0.007808676 |
| 0.262054566 |
| 1.2593E-06 |

| | |
|---|---|
| | 0.030674349 |
| | 0.021140847 |
| | 0.001294035 |
| | 0.12377521 |
| | 0.007293041 |
| | 1.24428E-08 |
| | 1.67587E-09 |
| | 0.060983755 |
| | 3.5888E-06 |
| | 0.186440241 |
| | 0.047891677 |
| | 0.117575231 |
| | 0.000212165 |
| | 1 |
| | 0.042175141 |
| | 0.000147718 |
| | 1 |
| | 0.000247755 |
| | 1 |
| | 0.5309602 |
| | 0.000192988 |
| | 1 |
| | 0.853489963 |
| | 7.80907E-07 |
| | 1.06157E-05 |
| | 1 |
| | 0.195867228 |
| | 1 |
| | 0.000444425 |
| | 0.011381994 |
| | 1 |
| | 0.819594915 |
| | 1 |
| | 1 |
| | 0.025865601 |

| |
|---|
| 1 |
| 1 |
| 1 |
| 1 |
| 1 |
| 4.52352E-07 |
| 0.058725134 |
| 1 |
| 0.000804661 |
| 3.94148E-08 |
| 0.548597298 |
| 1 |
| 1 |
| 0.177158671 |
| 0.013458321 |
| 1 |
| 1.18852E-06 |
| 1 |
| 0.668264011 |
| 1 |
| 0.023979391 |
| 0.007860144 |
| 0.025503581 |
| 0.000368671 |
| 0.056372801 |
| 1 |
| 1 |
| 1 |
| 1 |
| 1 |
| 0.003279219 |
| 0.000871903 |
| 0.004454941 |
| 1 |
| 1.58001E-11 |

| | |
|---|---|
| | 0.013350063 |
| | 0.665475581 |
| | 0.001302372 |
| | 0.084864987 |
| | 6.11719E-06 |
| | 0.795542393 |
| | 1.88322E-05 |
| | 1 |
| | 0.003438267 |
| | 0.594439726 |
| | 0.383301169 |
| | 1 |
| | 1 |
| | 5.43098E-09 |
| | 0.280566516 |
| | 1 |
| | 1 |
| | 1 |
| | 5.22568E-06 |
| | 1 |
| | 0.518196164 |
| | 6.33074E-07 |
| | 0.147813074 |
| | 2.72083E-05 |
| | 1 |
| | 1 |
| | 1 |
| | 0.005114801 |
| | 0.069030773 |
| | 0.000268485 |
| | 1 |
| | 0.599188186 |
| | 1 |
| | 2.76782E-07 |
| | 0.03407606 |

| |
|---|
| 0.00019198 |
| 0.043701771 |
| 0.39002607 |
| 0.380083971 |
| 1 |
| 0.951342345 |
| 0.000466054 |
| 0.000726154 |
| 1.14573E-06 |
| 0.00022857 |
| 0.033644994 |
| 1 |
| 5.66035E-06 |
| 0.000315416 |
| 0.00104629 |
| 1 |
| 1 |
| 0.712819767 |
| 0.010060807 |
| 1 |
| 1.85879E-08 |
| 0.958787057 |
| 0.020297692 |
| 0.009573444 |
| 0.850130662 |
| 0.029861231 |
| 1 |
| 1.15093E-06 |
| 9.85249E-07 |
| 9.55131E-05 |
| 0.026821252 |
| 0.001720303 |
| 0.02582139 |
| 3.89139E-08 |
| 0.0110932 |

| |
|---|
| 7.04289E-08 |
| 1 |
| 4.09564E-11 |
| 0.156599333 |
| 1 |
| 1 |
| 1 |
| 0.001408969 |
| 1 |
| 0.025836171 |
| 0.000406099 |
| 0.164115088 |
| 1 |
| 4.94926E-08 |
| 0.20289394 |
| 0.063596958 |
| 0.014219413 |
| 1 |
| 0.1624189 |
| 1 |
| 3.58508E-10 |
| 1.26614E-05 |
| 0.257842346 |
| 1 |
| 0.004019867 |
| 1 |
| 0.428601515 |
| 1 |
| 1 |
| 3.73172E-06 |
| 1 |
| 0.087820954 |
| 1 |
| 1 |
| 1.89417E-09 |

| |
|---|
| 0.843949142 |
| 6.35425E-06 |
| 3.21843E-05 |
| 1 |
| 0.09583959 |
| 1 |
| 1 |
| 1 |
| 0.006463896 |
| 0.001562626 |
| 1 |
| 1 |
| 1 |
| 1 |
| 3.30443E-05 |
| 1 |
| 0.772710825 |
| 2.27088E-06 |
| 1.44112E-05 |
| 6.38918E-05 |
| 0.986249816 |
| 2.03888E-06 |
| 1 |
| 1 |
| 1 |
| 1 |
| 1.03891E-07 |
| 1.96043E-06 |
| 0.611140897 |
| 1 |
| 1 |
| 6.25224E-10 |
| 8.98891E-11 |
| 5.04439E-06 |
| 0.003557162 |

| | |
|---|---|
| | 0.012462325 |
| | 2.00797E-08 |
| | 0.004559711 |
| | 0.000456158 |
| | 0.038844646 |
| | 1 |
| | 5.24762E-05 |
| | 0.161472644 |
| | 1 |
| | 0.800310266 |
| | 1 |
| | 0.000152154 |
| | 0.002536587 |
| | 1.78817E-09 |
| | 1 |
| | 1.63221E-05 |
| | 2.37215E-08 |
| | 0.011336687 |
| | 4.52356E-09 |
| | 1.40652E-11 |
| | 1 |
| | 1.04383E-08 |
| | 0.002051215 |
| | 4.88851E-07 |
| | 0.002474382 |
| | 1.40652E-11 |
| | 0.033444713 |
| | 1 |
| | 0.088514428 |
| | 0.000255348 |
| | 1 |
| | 1 |
| | 0.144655649 |
| | 0.000943015 |
| | 1.49347E-07 |

| |
|---|
| 1 |
| 1 |
| 1 |
| 0.003916948 |
| 0.353022691 |
| 1 |
| 2.17507E-07 |
| 3.34617E-07 |
| 0.824250996 |
| 1 |
| 0.55604643 |
| 0.191152291 |
| 1.40652E-11 |
| 0.001233534 |
| 6.61486E-06 |
| 1.40652E-11 |
| 1.40652E-11 |
| 2.01861E-08 |
| 0.940668685 |
| 1 |
| 0.251003822 |
| 0.428651657 |
| 0.001877654 |
| 0.479088958 |
| 1.76584E-05 |
| 1.40652E-11 |
| 0.000186193 |
| 1 |
| 0.469618283 |
| 1 |
| 0.73359592 |
| 1.40652E-11 |
| 2.918E-07 |
| 1.40652E-11 |
| 0.620495833 |

| | |
|---|---|
| | 1 |
| | 2.42888E-07 |
| | 0.049799632 |
| | 1 |
| | 0.049772996 |
| | 0.000443949 |
| | 0.888963606 |
| | 1 |
| | 0.074554153 |
| | 1 |
| | 0.769065764 |
| | 0.076365363 |
| | 1 |
| | 0.028026376 |
| | 0.06374432 |
| | 0.005549136 |
| | 0.000442272 |
| | 2.46082E-05 |
| | 4.72763E-07 |
| | 3.70691E-08 |
| | 0.000930594 |
| | 0.000520386 |
| | 6.05839E-06 |
| | 1 |
| | 0.179865635 |
| | 0.132664723 |
| | 1 |
| | 0.001518467 |
| | 1 |
| | 1 |
| | 0.000648054 |
| | 0.069789903 |
| | 0.409140578 |
| | 1.94429E-06 |
| | 0.014673547 |

| | |
|---|---|
| | 0.001168243 |
| | 0.057129061 |
| | 5.03877E-06 |
| | 0.06350596 |
| | 1.40652E-11 |
| | 1 |
| | 0.679194347 |
| | 1 |
| | 0.004722203 |
| | 2.89921E-06 |
| | 0.17491708 |
| | 0.004443086 |
| | 0.589881236 |
| | 0.00015841 |
| | 1 |
| | 2.78207E-07 |
| | 0.059738506 |
| | 1.40652E-11 |
| | 0.003517067 |
| | 0.663375869 |
| | 2.17929E-09 |
| | 8.53772E-09 |
| | 1.40652E-11 |
| | 1 |
| | 2.03027E-11 |
| | 0.730431963 |
| | 0.244712533 |
| | 0.069788993 |
| | 1.40652E-11 |
| | 1.78827E-11 |
| | 0.949062083 |
| | 1 |
| | 0.084456011 |
| | 1.40652E-11 |
| | 3.3025E-05 |

| |
|---|
| 1.40652E-11 |
| 1 |
| 2.63961E-07 |
| 0.035846244 |
| 0.138179728 |
| 0.15436832 |
| 0.355298485 |
| 0.282348488 |
| 0.127838415 |
| 0.347648872 |
| 0.611085927 |
| 1 |
| 1.9337E-08 |
| 1 |
| 0.166720125 |
| 0.196027097 |
| 4.65566E-08 |
| 1 |
| 1 |
| 1 |
| 7.45555E-11 |
| 0.384670008 |
| 0.546161172 |
| 1 |
| 1.60019E-07 |
| 5.13208E-06 |
| 0.032640248 |
| 0.008923852 |
| 0.001158833 |
| 0.081539101 |
| 1 |
| 0.218189672 |
| 1 |
| 1 |
| 0.481239888 |

| | |
|---|---|
| | 1 |
| | 0.663454922 |
| | 1 |
| | 1 |
| | 0.075191361 |
| | 0.000664418 |
| | 8.13753E-05 |
| | 0.240198948 |
| | 0.000900956 |
| | 8.5954E-08 |
| | 0.635563633 |
| | 1.50856E-07 |
| | 1 |
| | 0.000261852 |
| | 3.3055E-09 |
| | 0.051817075 |
| | 8.84757E-08 |
| | 1 |
| | 0.104500615 |
| | 6.94201E-05 |
| | 1 |
| | 0.082315299 |
| | 1 |
| | 0.075780365 |
| | 0.850641528 |
| | 1 |
| | 0.243475915 |
| | 0.984303613 |
| | 1 |
| | 1 |
| | 0.02450609 |
| | 0.033298795 |
| | 1 |
| | 0.035117577 |
| | 0.168537125 |

| | |
|---|---|
| | 0.000240766 |
| | 1 |
| | 1 |
| | 0.000143692 |
| | 0.258441018 |
| | 8.23636E-11 |
| | 3.29442E-08 |
| | 1.40652E-11 |
| | 7.16717E-10 |
| | 0.311646167 |
| | 7.55555E-09 |
| | 0.429350752 |
| | 0.214433578 |
| | 5.41835E-05 |
| | 1 |
| | 0.00053746 |
| | 0.148280227 |
| | 0.039024003 |
| | 1.77018E-06 |
| | 0.022028341 |
| | 0.000393181 |
| | 1 |
| | 1 |
| | 1 |
| | 1 |
| | 1 |
| | 1 |
| | 0.696566338 |
| | 0.305726779 |
| | 0.014085919 |
| | 0.233574742 |
| | 0.406552676 |
| | 0.094300545 |
| | 0.001128864 |
| | 0.00664256 |

| | |
|---|---|
| | 3.72865E-05 |
| | 1 |
| | 3.51427E-07 |
| | 0.046492502 |
| | 1 |
| | 0.45368484 |
| | 0.14193125 |
| | 1 |
| | 1 |
| | 0.012648046 |
| | 8.0048E-05 |
| | 0.623849699 |
| | 1 |
| | 1 |
| | 1 |
| | 0.484727762 |
| | 0.994907825 |
| | 0.468346386 |
| | 0.355831862 |
| | 0.218977889 |
| | 0.000438609 |
| | 1 |
| | 1.3194E-10 |
| | 0.369329487 |
| | 0.331928152 |
| | 0.15522646 |
| | 1.41411E-11 |
| | 0.020646996 |
| | 0.002816097 |
| | 0.201765813 |
| | 0.131079615 |
| | 0.147134858 |
| | 0.219373293 |
| | 0.194363225 |
| | 0.019051856 |

| |
|---:|
| 1 |
| 1 |
| 1 |
| 0.022047304 |
| 1 |
| 0.008714581 |
| 0.633642067 |
| 1 |
| 6.03602E-05 |
| 0.597755784 |
| 1 |
| 1 |
| 1 |
| 0.530574977 |
| 6.74139E-08 |

| |
| --- |
| Adj. P-value: (CM_pos_12022024_Newanalysis) / (SM_pos_12022024_Newanalyis) |
| 1 |
| 0.380085273 |
| 1 |
| 1 |
| 0.05454373 |
| 1 |
| 0.003873774 |
| 0.333199459 |
| 9.78775E-05 |
| 1 |
| 0.095416123 |
| 1 |
| 1 |
| 0.238383708 |
| 1 |
| 1 |
| 1 |
| 7.6992E-11 |
| 0.013126995 |
| 0.000732679 |
| 1 |
| 1.04491E-05 |
| 0.650274105 |
| 3.09895E-07 |
| 2.52171E-09 |
| 1 |
| 1 |
| 1 |
| 0.000708552 |
| 0.087281273 |
| 1 |
| 1 |
| 0.021412605 |

| | |
|---|---|
| | 0.092961577 |
| | 0.004021433 |
| | 1 |
| | 0.002750229 |
| | 1 |
| | 1 |
| | 4.14816E-06 |
| | 0.011975162 |
| | 0.000586804 |
| | 1 |
| | 0.000266133 |
| | 0.000766564 |
| | 0.001000021 |
| | 0.058548471 |
| | 0.000401457 |
| | 0.004934026 |
| | 0.410259323 |
| | 0.000693021 |
| | 0.020787141 |
| | 0.000529542 |
| | 0.525514215 |
| | 0.013859509 |
| | 0.245698176 |
| | 5.6022E-07 |
| | 0.965808191 |
| | 1 |
| | 0.03883933 |
| | 1 |
| | 0.004718383 |
| | 0.001761881 |
| | 0.016793826 |
| | 0.004609884 |
| | 0.000101856 |
| | 0.004747605 |
| | 1.42361E-05 |

| |
|---|
| 0.000130316 |
| 1 |
| 0.46675801 |
| 7.22926E-07 |
| 0.740403939 |
| 1.61717E-05 |
| 0.000188233 |
| 0.009323658 |
| 0.005040436 |
| 0.578722735 |
| 1.3025E-08 |
| 1.92324E-10 |
| 0.012218122 |
| 0.006064997 |
| 0.00301038 |
| 0.31071349 |
| 0.281893208 |
| 1 |
| 1 |
| 5.49754E-05 |
| 0.079514855 |
| 1 |
| 4.31502E-09 |
| 1.15002E-07 |
| 1 |
| 1 |
| 0.180541903 |
| 1 |
| 0.00014531 |
| 1 |
| 1 |
| 0.987377882 |
| 1 |
| 0.019760714 |
| 7.62006E-06 |

| |
|---|
| 1 |
| 0.1403155 |
| 0.001004813 |
| 1 |
| 0.614682623 |
| 1 |
| 1 |
| 0.010334879 |
| 1 |
| 0.287987285 |
| 0.214550637 |
| 0.000112869 |
| 0.03541977 |
| 1 |
| 3.21958E-11 |
| 0.000101123 |
| 1 |
| 0.001984187 |
| 0.075113807 |
| 1 |
| 0.022752396 |
| 0.00072055 |
| 1 |
| 1 |
| 2.78331E-07 |
| 0.001869454 |
| 1 |
| 1 |
| 1 |
| 1.21359E-10 |
| 0.022048518 |
| 4.9916E-11 |
| 0.971942391 |
| 1 |
| 0.859412388 |

| | |
|---|---|
| | 0.002972305 |
| | 0.185165007 |
| | 0.029117655 |
| | 1 |
| | 1 |
| | 1 |
| | 1 |
| | 1 |
| | 1 |
| | 1 |
| | 1 |
| | 0.174458247 |
| | 0.873471311 |
| | 1 |
| | 1 |
| | 0.979305927 |
| | 0.982781988 |
| | 0.003704701 |
| | 0.003566022 |
| | 1 |
| | 0.062806566 |
| | 0.307221229 |
| | 1 |
| | 1 |
| | 9.09105E-07 |
| | 0.000153454 |
| | 0.002180119 |
| | 1 |
| | 1 |
| | 0.567012062 |
| | 1 |
| | 1 |
| | 0.002578532 |
| | 0.31103742 |
| | 1 |

| | |
|---|---|
| | 0.13850162 |
| | 0.171252067 |
| | 0.153889946 |
| | 0.889420655 |
| | 1 |
| | 1 |
| | 1 |
| | 0.010160454 |
| | 1 |
| | 0.001315284 |
| | 0.238383708 |
| | 3.71193E-09 |
| | 1 |
| | 1 |
| | 0.734303226 |
| | 0.06725895 |
| | 0.338885482 |
| | 0.001508227 |
| | 0.567725115 |
| | 1 |
| | 1 |
| | 1 |
| | 0.017401083 |
| | 0.000618524 |
| | 0.010244006 |
| | 1 |
| | 1 |
| | 0.906150982 |
| | 1 |
| | 6.63775E-05 |
| | 0.826374901 |
| | 0.001437114 |
| | 1 |
| | 0.002851188 |
| | 0.000830041 |

| |
|---|
| 0.420469396 |
| 0.073559795 |
| 0.065414272 |
| 1 |
| 0.793417462 |
| 3.74852E-08 |
| 9.34722E-08 |
| 0.059377888 |
| 3.5618E-06 |
| 0.016666098 |
| 1 |
| 1 |
| 0.120284719 |
| 1 |
| 1 |
| 1 |
| 0.000162034 |
| 0.647138713 |
| 0.014029604 |
| 1 |
| 1 |
| 1 |
| 0.490207303 |
| 3.86403E-05 |
| 1 |
| 0.860545017 |
| 0.266783293 |
| 1 |
| 0.445516645 |
| 0.018968559 |
| 0.832581074 |
| 0.419976118 |
| 1 |
| 1 |
| 0.114833721 |

| | |
|---|---|
| | 0.893468602 |
| | 1 |
| | 1 |
| | 1 |
| | 1 |
| | 0.00430088 |
| | 1 |
| | 1 |
| | 0.067212818 |
| | 0.834430611 |
| | 1 |
| | 0.798789256 |
| | 1 |
| | 1 |
| | 1 |
| | 0.120156703 |
| | 0.087473125 |
| | 0.352897439 |
| | 1 |
| | 2.09823E-09 |
| | 0.830921902 |
| | 2.55296E-06 |
| | 1 |
| | 3.02867E-06 |
| | 0.048894017 |
| | 1 |
| | 1 |
| | 1 |
| | 0.002342246 |
| | 1 |
| | 0.748242873 |
| | 0.216730976 |
| | 0.587288015 |
| | 1 |
| | 0.986971698 |

| | |
|---|---|
| | 0.29673624 |
| | 0.000955815 |
| | 0.460958685 |
| | 0.026346074 |
| | 1.02555E-05 |
| | 1 |
| | 1 |
| | 1 |
| | 0.103648122 |
| | 1 |
| | 0.00057718 |
| | 1 |
| | 1 |
| | 9.88091E-11 |
| | 1 |
| | 0.364364332 |
| | 0.063970082 |
| | 1 |
| | 0.288620596 |
| | 1 |
| | 3.73298E-06 |
| | 0.273546688 |
| | 0.273281094 |
| | 0.094929365 |
| | 1 |
| | 1 |
| | 1 |
| | 0.973008972 |
| | 2.54891E-06 |
| | 1 |
| | 0.391886587 |
| | 1 |
| | 0.132371597 |
| | 0.000301603 |
| | 1 |

| |
|---|
| 0.335645202 |
| 0.852454037 |
| 0.106023215 |
| 0.684338037 |
| 1 |
| 1 |
| 0.897557631 |
| 0.609362842 |
| 1.4005E-06 |
| 1 |
| 1 |
| 1 |
| 0.073664388 |
| 1 |
| 1.3053E-05 |
| 1 |
| 1 |
| 0.010719022 |
| 0.007622057 |
| 0.019092252 |
| 3.04458E-05 |
| 0.571867479 |
| 1 |
| 0.987467282 |
| 1 |
| 0.000517179 |
| 0.349695806 |
| 1 |
| 1 |
| 1 |
| 1 |
| 0.373533025 |
| 0.832088543 |
| 0.407138201 |
| 1 |

| | |
|---|---|
| | 1 |
| | 1 |
| | 0.016417447 |
| | 0.031637743 |
| | 1 |
| | 1 |
| | 1 |
| | 0.139296014 |
| | 1 |
| | 1 |
| | 1 |
| | 1 |
| | 1 |
| | 0.000641955 |
| | 0.202332273 |
| | 0.000122973 |
| | 1 |
| | 1 |
| | 1 |
| | 0.672625981 |
| | 1 |
| | 0.048476798 |
| | 1 |
| | 0.461845123 |
| | 1 |
| | 1 |
| | 1 |
| | 1 |
| | 1 |
| | 0.015373062 |
| | 1 |
| | 0.00307318 |
| | 0.85815285 |
| | 0.010967885 |
| | 1.35043E-09 |

| |
|---|
| 0.413510841 |
| 6.10646E-05 |
| 0.021197057 |
| 1 |
| 0.003229801 |
| 1 |
| 1 |
| 1 |
| 4.57006E-06 |
| 0.743938205 |
| 1 |
| 1 |
| 1 |
| 0.137553705 |
| 0.248003775 |
| 0.266356837 |
| 1 |
| 1 |
| 1 |
| 0.048729924 |
| 1 |
| 0.031880263 |
| 1 |
| 1 |
| 1 |
| 1 |
| 2.77373E-06 |
| 0.013081994 |
| 1 |
| 0.599892759 |
| 0.000116159 |
| 0.82546396 |
| 0.004334859 |
| 1 |
| 0.964729561 |

| | |
|---|---|
| | 1 |
| | 1.05115E-05 |
| | 0.013012418 |
| | 0.557182551 |
| | 0.008955992 |
| | 0.075446111 |
| | 1.54424E-09 |
| | 0.541006346 |
| | 1 |
| | 1 |
| | 1 |
| | 1 |
| | 1 |
| | 0.001310473 |
| | 0.303221017 |
| | 0.000202098 |
| | 0.000164031 |
| | 0.171194063 |
| | 0.006486849 |
| | 0.001371426 |
| | 1 |
| | 1 |
| | 1 |
| | 0.110596149 |
| | 0.001857048 |
| | 4.20364E-09 |
| | 0.049361166 |
| | 0.000816137 |
| | 1 |
| | 0.06983972 |
| | 0.058963312 |
| | 1 |
| | 1 |
| | 1 |
| | 1 |

| | |
|---|---|
| | 0.000551205 |
| | 0.099198027 |
| | 0.612082381 |
| | 0.491185463 |
| | 0.002089524 |
| | 1 |
| | 0.297873681 |
| | 0.002305198 |
| | 1 |
| | 0.318323945 |
| | 0.755615046 |
| | 5.52046E-05 |
| | 0.736106894 |
| | 0.330742814 |
| | 1 |
| | 0.149756139 |
| | 1 |
| | 0.002126778 |
| | 1 |
| | 1 |
| | 0.635149016 |
| | 0.426942186 |
| | 1 |
| | 1 |
| | 1 |
| | 1 |
| | 0.006690429 |
| | 1 |
| | 0.057844821 |
| | 1 |
| | 1 |
| | 1 |
| | 1 |
| | 1 |
| | 0.308033329 |

| | |
|---|---|
| | 1 |
| | 0.040305904 |
| | 1 |
| | 1 |
| | 0.591181912 |
| | 0.000482433 |
| | 1 |
| | 0.282234454 |
| | 0.008238955 |
| | 1 |
| | 0.766553143 |
| | 0.208265569 |
| | 1 |
| | 0.144027442 |
| | 0.46708544 |
| | 1 |
| | 0.099380711 |
| | 3.75926E-07 |
| | 0.382007008 |
| | 2.14716E-07 |
| | 0.135644504 |
| | 0.000101508 |
| | 3.10653E-06 |
| | 1 |
| | 0.805313846 |
| | 0.638581965 |
| | 1 |
| | 0.012240684 |
| | 1 |
| | 1 |
| | 1 |
| | 1 |
| | 0.779053064 |
| | 0.134281312 |
| | 1 |

| | |
|---|---|
| | 0.07634077 |
| | 0.013312527 |
| | 1.3563E-05 |
| | 1 |
| | 3.21958E-11 |
| | 1 |
| | 0.919200058 |
| | 0.675215378 |
| | 3.20446E-06 |
| | 0.32189494 |
| | 0.685505567 |
| | 0.089677828 |
| | 0.072498754 |
| | 0.005579733 |
| | 1 |
| | 0.012533726 |
| | 1 |
| | 1.17492E-09 |
| | 0.438836086 |
| | 1 |
| | 0.195029029 |
| | 3.21958E-11 |
| | 1.70595E-10 |
| | 1 |
| | 1.87674E-10 |
| | 1 |
| | 0.001717704 |
| | 0.024819627 |
| | 3.21958E-11 |
| | 3.30559E-06 |
| | 0.500228699 |
| | 1 |
| | 1 |
| | 0.003582289 |
| | 0.000620209 |

| |
|---|
| 1 |
| 1 |
| 0.847665062 |
| 1 |
| 1 |
| 1 |
| 1 |
| 1 |
| 0.619155028 |
| 0.009222997 |
| 0.657739623 |
| 1 |
| 9.92014E-07 |
| 0.670429374 |
| 1 |
| 1 |
| 1.67081E-08 |
| 0.040578225 |
| 0.099447504 |
| 1 |
| 2.47474E-08 |
| 0.465823942 |
| 0.029301185 |
| 1 |
| 1 |
| 1 |
| 0.000438458 |
| 0.254214194 |
| 1 |
| 0.21270188 |
| 0.703473503 |
| 0.945222915 |
| 1 |
| 1 |
| 0.23349817 |

| |
|---|
| 1 |
| 0.876218087 |
| 1 |
| 0.831573594 |
| 0.151720201 |
| 0.125254732 |
| 4.35569E-06 |
| 0.934880338 |
| 1 |
| 1.48088E-09 |
| 0.138689505 |
| 1 |
| 0.201697644 |
| 2.50008E-06 |
| 1 |
| 1 |
| 1 |
| 1 |
| 1.19429E-05 |
| 0.202532414 |
| 1 |
| 1 |
| 1 |
| 0.228936139 |
| 0.428270251 |
| 1 |
| 0.002623757 |
| 0.095158714 |
| 0.714924707 |
| 1 |
| 0.466531955 |
| 0.333720489 |
| 1 |
| 0.023369535 |
| 0.147193895 |

| | |
|---|---|
| | 0.004041927 |
| | 6.54609E-08 |
| | 1.16917E-05 |
| | 1 |
| | 3.66868E-08 |
| | 3.85297E-05 |
| | 0.032515024 |
| | 3.21958E-11 |
| | 0.08836688 |
| | 1 |
| | 7.0499E-10 |
| | 1 |
| | 1 |
| | 0.000335542 |
| | 1 |
| | 0.000831567 |
| | 0.109908808 |
| | 1 |
| | 0.000131244 |
| | 0.355411211 |
| | 0.015706223 |
| | 1 |
| | 1 |
| | 1 |
| | 1 |
| | 1 |
| | 1 |
| | 0.476246015 |
| | 1 |
| | 0.154544282 |
| | 0.33162818 |
| | 0.7977459 |
| | 1 |
| | 8.21587E-06 |
| | 1 |

| | |
|---|---|
| | 0.00014302 |
| | 1 |
| | 3.66631E-07 |
| | 0.105321664 |
| | 0.835109687 |
| | 1 |
| | 1 |
| | 1.80399E-06 |
| | 1 |
| | 0.156252738 |
| | 3.21958E-11 |
| | 5.77401E-10 |
| | 5.81838E-06 |
| | 0.925833368 |
| | 1 |
| | 1 |
| | 1 |
| | 1 |
| | 0.390362826 |
| | 0.458451211 |
| | 0.641140163 |
| | 1 |
| | 3.07581E-07 |
| | 0.309453807 |
| | 1 |
| | 1 |
| | 1 |
| | 1 |
| | 0.004258245 |
| | 1 |
| | 0.029971518 |
| | 1 |
| | 1 |
| | 0.080366574 |
| | 1 |

| |
|---:|
| 1 |
| 1 |
| 0.074214077 |
| 0.301299175 |
| 1 |
| 1 |
| 0.899421599 |
| 1 |
| 0.858634941 |
| 1 |
| 1 |
| 0.95620606 |
| 1 |
| 1 |
| 0.000165885 |

| |
|---|
| Adj. P-value: (Bb_pos_12022024_Newanalysis) / (Water_pos_12022024_Newanalysis) |
| 0.316108156 |
| 0.061679804 |
| 1 |
| 1 |
| 0.834461044 |
| 0.458187388 |
| 1.90457E-05 |
| 0.607035013 |
| 1 |
| 1 |
| 1 |
| 1.10209E-08 |
| 0.750249948 |
| 1 |
| 1 |
| 0.133789322 |
| 1 |
| 1 |
| 1 |
| 0.086367727 |
| 0.71353285 |
| 0.113486261 |
| 1 |
| 0.18631824 |
| 0.011310085 |
| 1 |
| 1 |
| 1 |
| 0.00817435 |
| 0.067417069 |
| 1 |
| 0.077597725 |
| 1 |

| | |
|---|---|
| | 0.457671362 |
| | 0.633908958 |
| | 1 |
| | 0.009779899 |
| | 0.315235813 |
| | 0.400254536 |
| | 0.061531868 |
| | 0.953175652 |
| | 0.018276904 |
| | 0.921554537 |
| | 0.179793255 |
| | 0.02500678 |
| | 0.020625875 |
| | 0.005925213 |
| | 0.031650525 |
| | 0.152359721 |
| | 0.063092285 |
| | 0.252568472 |
| | 0.070803006 |
| | 0.00873056 |
| | 0.036285408 |
| | 1 |
| | 0.080845953 |
| | 1 |
| | 1 |
| | 0.869027554 |
| | 0.000243069 |
| | 0.129758992 |
| | 1 |
| | 1.26579E-05 |
| | 0.664743947 |
| | 0.654587319 |
| | 0.026004262 |
| | 2.23882E-07 |
| | 0.109500612 |

| | |
|---|---|
| | 0.023489216 |
| | 0.02703907 |
| | 1.4949E-06 |
| | 0.006810545 |
| | 0.015869608 |
| | 0.07205212 |
| | 0.002151239 |
| | 1 |
| | 0.081156756 |
| | 0.013963417 |
| | 0.000251465 |
| | 0.004461039 |
| | 0.086367727 |
| | 1.20383E-10 |
| | 0.052553637 |
| | 0.004351243 |
| | 0.045184655 |
| | 0.10835337 |
| | 0.179087108 |
| | 0.009638179 |
| | 0.05880559 |
| | 0.00200698 |
| | 0.734463566 |
| | 0.004375144 |
| | 0.011869647 |
| | 0.131222033 |
| | 0.686732776 |
| | 0.039520497 |
| | 0.001091356 |
| | 0.017477345 |
| | 0.00822161 |
| | 0.215959208 |
| | 1 |
| | 0.003417284 |
| | 0.020641144 |

| | |
|---|---|
| | 6.55241E-06 |
| | 1 |
| | 0.031333707 |
| | 0.022268937 |
| | 0.11626431 |
| | 1 |
| | 0.394770634 |
| | 0.076853261 |
| | 1 |
| | 0.017617098 |
| | 0.055162004 |
| | 0.009974618 |
| | 0.04151689 |
| | 0.357950595 |
| | 0.006838942 |
| | 1 |
| | 0.356220301 |
| | 0.124702414 |
| | 0.04817479 |
| | 0.785877816 |
| | 1 |
| | 0.01321107 |
| | 1 |
| | 0.049180011 |
| | 0.148644581 |
| | 0.002291342 |
| | 0.323558373 |
| | 0.300088541 |
| | 1 |
| | 0.845926685 |
| | 1 |
| | 0.006872559 |
| | 0.14266689 |
| | 0.021091855 |
| | 1 |

| | |
|---|---|
| | 0.438293381 |
| | 0.886284834 |
| | 0.238168362 |
| | 0.53785278 |
| | 1 |
| | 1 |
| | 1 |
| | 1 |
| | 0.015143768 |
| | 1 |
| | 1 |
| | 1 |
| | 0.029828802 |
| | 1 |
| | 0.05444188 |
| | 0.131915706 |
| | 1 |
| | 0.79116554 |
| | 3.54023E-06 |
| | 1 |
| | 0.035363379 |
| | 0.177155431 |
| | 0.944325884 |
| | 0.989992322 |
| | 1 |
| | 0.032026708 |
| | 4.3382E-05 |
| | 0.294257647 |
| | 0.098216167 |
| | 0.143101328 |
| | 0.871897182 |
| | 0.090983608 |
| | 4.79304E-05 |
| | 1 |
| | 1 |

| |
|---|
| 0.008678928 |
| 0.102136214 |
| 0.592690905 |
| 0.220005188 |
| 1 |
| 1 |
| 1 |
| 0.141656105 |
| 3.42424E-11 |
| 1 |
| 0.058318536 |
| 0.012471308 |
| 0.025098479 |
| 0.342588353 |
| 0.020519728 |
| 0.014587803 |
| 0.277146532 |
| 0.887832608 |
| 0.488946922 |
| 0.073224905 |
| 1 |
| 0.811353005 |
| 0.00015307 |
| 0.353162612 |
| 9.04734E-05 |
| 1 |
| 1 |
| 1 |
| 0.040535673 |
| 0.841541963 |
| 0.03404332 |
| 0.263080559 |
| 0.98750171 |
| 0.624470372 |
| 1 |

| | |
|---|---|
| | 0.016908067 |
| | 0.122774397 |
| | 0.102363593 |
| | 0.843856428 |
| | 0.001867506 |
| | 0.241229521 |
| | 1 |
| | 1 |
| | 0.555517637 |
| | 0.098112367 |
| | 1 |
| | 0.454020309 |
| | 0.822409124 |
| | 1 |
| | 0.144368059 |
| | 0.00309813 |
| | 0.261944286 |
| | 1 |
| | 1 |
| | 1 |
| | 1 |
| | 1 |
| | 0.452997178 |
| | 0.608077078 |
| | 0.042267924 |
| | 1 |
| | 0.1300264 |
| | 0.793270869 |
| | 0.11907191 |
| | 1 |
| | 1 |
| | 1 |
| | 1 |
| | 1 |
| | 0.040951571 |

| | |
|---|---|
| | 0.009556079 |
| | 0.120694121 |
| | 1 |
| | 1 |
| | 1 |
| | 0.385580342 |
| | 1 |
| | 1 |
| | 0.299892088 |
| | 1 |
| | 1.98277E-07 |
| | 0.187642143 |
| | 1 |
| | 8.42379E-05 |
| | 6.97046E-06 |
| | 0.000823936 |
| | 0.011129695 |
| | 0.128087062 |
| | 1 |
| | 1 |
| | 0.152269171 |
| | 0.14781159 |
| | 0.241663558 |
| | 1 |
| | 0.907732592 |
| | 1 |
| | 1 |
| | 1 |
| | 0.020778639 |
| | 1 |
| | 1 |
| | 0.040210108 |
| | 0.002025449 |
| | 1 |
| | 1 |

| | |
|---|---|
| | 0.00176691 |
| | 0.57327628 |
| | 0.039846281 |
| | 1 |
| | 1 |
| | 1 |
| | 1 |
| | 0.901132439 |
| | 0.014914094 |
| | 0.079320953 |
| | 0.28384492 |
| | 0.164715043 |
| | 0.342650815 |
| | 2.18271E-06 |
| | 0.257831078 |
| | 1 |
| | 0.258338983 |
| | 0.067625323 |
| | 0.841252329 |
| | 1 |
| | 1 |
| | 0.005549347 |
| | 0.02307958 |
| | 0.039022316 |
| | 0.418060913 |
| | 0.103167376 |
| | 1 |
| | 0.724827326 |
| | 0.002744699 |
| | 2.65347E-05 |
| | 1 |
| | 1 |
| | 0.002096606 |
| | 0.003880569 |
| | 0.830095028 |

| |
|---|
| 0.09205768 |
| 0.008392045 |
| 1 |
| 0.046719984 |
| 5.25489E-06 |
| 0.119635772 |
| 0.077546989 |
| 0.065653177 |
| 0.009595881 |
| 0.802677388 |
| 1 |
| 1.77739E-06 |
| 1 |
| 0.947674473 |
| 1 |
| 0.037773256 |
| 0.101145362 |
| 1 |
| 1 |
| 0.000158456 |
| 0.184157281 |
| 2.90613E-06 |
| 1 |
| 0.997408805 |
| 1 |
| 0.066841267 |
| 0.169564994 |
| 7.74418E-09 |
| 1 |
| 4.25932E-06 |
| 0.000385062 |
| 0.065520164 |
| 1.4545E-05 |
| 6.4525E-09 |
| 0.004267765 |

| | |
|---|---|
| | 0.016356299 |
| | 1 |
| | 6.33701E-09 |
| | 0.001834195 |
| | 0.631887006 |
| | 0.784916991 |
| | 0.065890988 |
| | 0.061975076 |
| | 0.12297662 |
| | 0.506804566 |
| | 1.69255E-07 |
| | 0.656423262 |
| | 0.384213871 |
| | 9.92485E-08 |
| | 0.067445275 |
| | 0.220263062 |
| | 0.218264199 |
| | 0.452200348 |
| | 1 |
| | 0.212324182 |
| | 0.252352826 |
| | 0.139875059 |
| | 1 |
| | 0.770315787 |
| | 0.872865258 |
| | 1 |
| | 1 |
| | 1 |
| | 0.09972941 |
| | 0.003134646 |
| | 0.806269345 |
| | 1 |
| | 0.973385801 |
| | 0.103746231 |
| | 1 |

| |
|---|
| 0.694326484 |
| 0.001485314 |
| 6.69033E-07 |
| 0.850487739 |
| 0.882137478 |
| 0.104758825 |
| 1 |
| 1 |
| 1.15172E-05 |
| 0.000206475 |
| 0.701064149 |
| 0.240706181 |
| 0.103667187 |
| 1 |
| 1 |
| 1 |
| 0.845518558 |
| 1 |
| 0.501514074 |
| 0.504931157 |
| 2.45358E-11 |
| 0.389757965 |
| 0.211949796 |
| 0.077604893 |
| 0.026859431 |
| 1 |
| 4.85388E-05 |
| 0.305706481 |
| 1 |
| 0.462653653 |
| 0.181684755 |
| 7.18891E-11 |
| 0.639453935 |
| 1 |
| 1 |

| | |
|---|---|
| | 0.787662126 |
| | 0.292005125 |
| | 0.121473604 |
| | 1 |
| | 0.077183295 |
| | 1 |
| | 0.015176214 |
| | 0.112054997 |
| | 0.940290076 |
| | 3.11055E-08 |
| | 0.194885566 |
| | 1 |
| | 1 |
| | 1 |
| | 1.26472E-08 |
| | 0.001498877 |
| | 6.67268E-10 |
| | 0.111426309 |
| | 1 |
| | 0.530691677 |
| | 0.742845844 |
| | 0.381555343 |
| | 1 |
| | 1 |
| | 0.278415021 |
| | 1 |
| | 1 |
| | 1 |
| | 1 |
| | 0.000392269 |
| | 0.065024895 |
| | 0.005770405 |
| | 0.156782071 |
| | 1 |
| | 1 |

| |
|---|
| 1 |
| 1 |
| 0.14586897 |
| 1 |
| 1 |
| 3.93842E-05 |
| 1 |
| 1 |
| 1 |
| 8.62737E-06 |
| 0.068884923 |
| 5.43046E-07 |
| 2.45358E-11 |
| 7.33761E-07 |
| 1.45257E-08 |
| 2.45358E-11 |
| 2.45358E-11 |
| 1.11008E-09 |
| 0.373854654 |
| 0.004011627 |
| 1 |
| 1 |
| 0.000535353 |
| 1.29214E-10 |
| 2.45358E-11 |
| 2.45358E-11 |
| 8.66608E-10 |
| 1.46275E-05 |
| 0.736111732 |
| 0.025480696 |
| 0.214483578 |
| 4.50441E-10 |
| 7.60894E-11 |
| 2.45358E-11 |
| 0.029013961 |

| |
|---|
| 0.826179396 |
| 1 |
| 0.421292287 |
| 1 |
| 0.001248966 |
| 1 |
| 0.001054026 |
| 1 |
| 1 |
| 1 |
| 1 |
| 0.984516704 |
| 1 |
| 1 |
| 1 |
| 1 |
| 2.21639E-08 |
| 0.13884919 |
| 3.22589E-06 |
| 1 |
| 0.44925227 |
| 1 |
| 1 |
| 1 |
| 1 |
| 0.090095855 |
| 0.008555252 |
| 0.110434314 |
| 1 |
| 1 |
| 1 |
| 1 |
| 0.759814189 |
| 0.004205956 |
| 1 |

| |
|---|
| 1 |
| 1 |
| 1 |
| 0.002627342 |
| 1 |
| 1 |
| 1 |
| 0.539372621 |
| 0.235159866 |
| 1 |
| 1 |
| 0.981836965 |
| 0.056648478 |
| 0.000228105 |
| 1 |
| 0.27235018 |
| 1 |
| 1 |
| 1 |
| 0.007186317 |
| 0.140977454 |
| 2.99133E-05 |
| 1 |
| 1 |
| 1 |
| 0.003004406 |
| 1 |
| 0.389634071 |
| 0.46106051 |
| 0.309851357 |
| 1 |
| 0.761470439 |
| 0.858205229 |
| 1.95495E-10 |
| 4.52152E-05 |

| | |
|---|---|
| | 2.63287E-11 |
| | 1 |
| | 4.6008E-08 |
| | 1 |
| | 0.561469823 |
| | 1 |
| | 1 |
| | 1 |
| | 0.823404466 |
| | 1 |
| | 0.244071624 |
| | 1 |
| | 1 |
| | 0.844771481 |
| | 1 |
| | 0.091690969 |
| | 3.75097E-06 |
| | 0.186707637 |
| | 0.063682443 |
| | 0.231818649 |
| | 0.949656563 |
| | 0.121833357 |
| | 1 |
| | 1 |
| | 5.22757E-06 |
| | 0.000309401 |
| | 1 |
| | 0.13605834 |
| | 0.001283663 |
| | 1 |
| | 0.351340677 |
| | 0.357977379 |
| | 1 |
| | 1.4716E-07 |
| | 1 |

| |
|---|
| 0.335455604 |
| 1 |
| 1 |
| 1 |
| 1 |
| 0.00191166 |
| 0.009606046 |
| 0.240144807 |
| 1 |
| 1 |
| 1 |
| 8.59997E-07 |
| 0.532201372 |
| 0.014220601 |
| 0.488943224 |
| 0.61101232 |
| 1.87406E-07 |
| 0.381479737 |
| 1 |
| 6.19075E-07 |
| 0.342520488 |
| 1 |
| 1 |
| 0.76606021 |
| 1 |
| 0.388257799 |
| 1 |
| 1 |
| 0.011600093 |
| 1 |
| 0.757631134 |
| 1 |
| 1 |
| 1 |
| 9.22281E-05 |

| | |
|---|---|
| | 0.053810233 |
| | 0.001815425 |
| | 0.022200049 |
| | 0.04759729 |
| | 0.089280906 |
| | 0.001398188 |
| | 0.008298021 |
| | 9.51902E-06 |
| | 0.004706143 |
| | 0.065634276 |
| | 1 |
| | 0.109170795 |
| | 0.019195574 |
| | 0.003571821 |
| | 1 |
| | 0.00033813 |
| | 0.248628519 |
| | 0.82163113 |
| | 0.13149257 |
| | 0.100888939 |
| | 1 |
| | 0.27804603 |
| | 0.237513977 |
| | 1 |
| | 0.323272454 |
| | 0.224312133 |
| | 0.268047508 |
| | 0.43549565 |
| | 0.457147818 |
| | 0.98365406 |
| | 0.005619613 |
| | 0.000407586 |
| | 0.031892232 |
| | 5.42153E-06 |
| | 0.028901212 |

| |
|---:|
| 0.000280654 |
| 0.086721951 |
| 0.514553901 |
| 1 |
| 0.274469335 |
| 0.034342253 |
| 0.041034195 |
| 0.01641704 |
| 1 |
| 0.261572226 |
| 0.274136017 |
| 0.009413765 |
| 0.011038161 |
| 0.662945108 |
| 1 |
| 0.918281325 |
| 0.344123794 |
| 1 |
| 1 |
| 0.97251664 |
| 0.037427094 |
| 1 |
| 2.45358E-11 |
| 0.155654021 |
| 0.899768607 |
| 0.441506143 |
| 1.21167E-07 |
| 1 |
| 1 |
| 1 |
| 1 |
| 1 |
| 1 |
| 0.122603397 |
| 0.780578882 |

| |
|---|
| 1 |
| 0.005473726 |
| 1 |
| 1 |
| 0.918946627 |
| 1 |
| 0.619767749 |
| 1 |
| 0.766848104 |
| 0.562595477 |
| 0.958000469 |
| 1 |
| 1 |
| 0.934915391 |
| 1 |

| |
|---|
| Adj. P-value: (CM_pos_12022024_Newanalysis) / (Water_pos_12022024_Newanalysis) |
| 1 |
| 0.702632056 |
| 1 |
| 0.38345684 |
| 1 |
| 0.929807213 |
| 0.000415422 |
| 0.928806032 |
| 0.449687183 |
| 1 |
| 1 |
| 2.17109E-07 |
| 1 |
| 0.279290104 |
| 1 |
| 1 |
| 1 |
| 7.0709E-11 |
| 1 |
| 5.25146E-06 |
| 1 |
| 6.84375E-06 |
| 0.001177473 |
| 4.48568E-08 |
| 8.09862E-11 |
| 1 |
| 1 |
| 0.876938568 |
| 0.624166793 |
| 0.17201911 |
| 1 |
| 0.005211109 |
| 0.218768272 |

| |
|---|
| 1 |
| 4.48813E-09 |
| 1 |
| 1 |
| 0.74907763 |
| 0.436845839 |
| 0.002821911 |
| 0.72738756 |
| 0.000786681 |
| 0.368789957 |
| 1 |
| 0.008982566 |
| 0.000164583 |
| 0.037683708 |
| 0.018059441 |
| 1 |
| 1 |
| 0.553686846 |
| 0.18292542 |
| 0.000469395 |
| 1 |
| 0.361682446 |
| 0.868700846 |
| 0.570379026 |
| 0.994055518 |
| 1 |
| 0.190200475 |
| 0.821836259 |
| 1 |
| 0.003602909 |
| 0.000104491 |
| 0.399516023 |
| 0.0130762 |
| 0.281467684 |
| 1 |

| | |
|---|---|
| | 0.220499329 |
| | 0.290469762 |
| | 1 |
| | 0.002822643 |
| | 1 |
| | 1 |
| | 0.006878668 |
| | 1 |
| | 8.4402E-09 |
| | 0.056189796 |
| | 0.005195583 |
| | 0.113627807 |
| | 0.655415939 |
| | 1 |
| | 0.041653341 |
| | 1 |
| | 1 |
| | 1 |
| | 0.054927538 |
| | 0.124954373 |
| | 1 |
| | 0.404250801 |
| | 0.027117053 |
| | 0.020720606 |
| | 0.956023743 |
| | 0.220359891 |
| | 0.110127436 |
| | 1 |
| | 0.042466533 |
| | 1 |
| | 0.013876197 |
| | 1 |
| | 0.436012938 |
| | 0.115482677 |
| | 1 |

| | |
|---|---|
| | 3.75481E-05 |
| | 0.395039743 |
| | 0.006507443 |
| | 1 |
| | 0.260152153 |
| | 1 |
| | 0.469866738 |
| | 1 |
| | 1 |
| | 1 |
| | 1 |
| | 0.00374048 |
| | 1 |
| | 1 |
| | 7.0709E-11 |
| | 0.107876786 |
| | 0.092796173 |
| | 0.034911622 |
| | 0.043395215 |
| | 0.654425943 |
| | 1 |
| | 0.044241293 |
| | 1 |
| | 1 |
| | 0.635909226 |
| | 0.00468839 |
| | 0.234985473 |
| | 0.090754387 |
| | 0.298093971 |
| | 2.81036E-08 |
| | 0.069614731 |
| | 1.69203E-06 |
| | 1 |
| | 1 |
| | 1 |

| | |
|---|---|
| | 0.649368633 |
| | 0.15614564 |
| | 1 |
| | 1 |
| | 1 |
| | 1 |
| | 1 |
| | 1 |
| | 1 |
| | 1 |
| | 0.80652535 |
| | 1 |
| | 0.123235828 |
| | 0.533174978 |
| | 0.888759018 |
| | 0.000994858 |
| | 1 |
| | 1 |
| | 0.003590687 |
| | 0.511053138 |
| | 0.714689485 |
| | 1 |
| | 1 |
| | 0.016640493 |
| | 1 |
| | 1 |
| | 0.011674215 |
| | 1 |
| | 0.954712758 |
| | 0.560522265 |
| | 0.022439098 |
| | 1 |
| | 0.011997487 |
| | 1 |
| | 0.080238386 |

| |
|---|
| 1 |
| 0.771567606 |
| 0.591764267 |
| 3.18104E-07 |
| 0.02173429 |
| 1 |
| 0.437362045 |
| 0.600592126 |
| 1.07525E-08 |
| 1 |
| 1 |
| 0.001107249 |
| 6.32804E-05 |
| 3.19478E-05 |
| 0.861484003 |
| 0.000235106 |
| 0.153260642 |
| 1 |
| 4.20476E-09 |
| 1 |
| 0.535455395 |
| 0.859922744 |
| 0.05639984 |
| 0.05438366 |
| 0.089749223 |
| 1 |
| 1 |
| 0.302820649 |
| 1 |
| 0.071368444 |
| 1 |
| 1 |
| 0.00407442 |
| 0.01250616 |
| 0.055810473 |

| | |
|---|---|
| | 0.791535394 |
| | 0.251037871 |
| | 0.952698368 |
| | 1 |
| | 0.203322506 |
| | 0.35231902 |
| | 1 |
| | 1 |
| | 0.377156607 |
| | 0.841597627 |
| | 1 |
| | 1 |
| | 1 |
| | 0.977362072 |
| | 1 |
| | 1 |
| | 0.339328385 |
| | 0.009455372 |
| | 0.000960246 |
| | 1 |
| | 0.19919209 |
| | 1 |
| | 1 |
| | 1 |
| | 0.889690776 |
| | 1 |
| | 0.134255639 |
| | 1 |
| | 1 |
| | 1 |
| | 1 |
| | 0.870830302 |
| | 0.903117675 |
| | 0.818212966 |
| | 0.119664711 |

| |
|---|
| 0.844629329 |
| 0.08684632 |
| 1 |
| 1 |
| 1 |
| 1 |
| 0.038140875 |
| 1 |
| 1 |
| 0.000249355 |
| 0.000967004 |
| 1 |
| 1 |
| 2.87941E-08 |
| 0.749999485 |
| 1 |
| 1 |
| 1 |
| 1 |
| 1.38336E-09 |
| 1 |
| 1 |
| 1 |
| 1 |
| 0.78933956 |
| 0.690430855 |
| 0.555607114 |
| 1 |
| 1 |
| 1 |
| 0.279287887 |
| 0.015467859 |
| 0.202967247 |
| 1 |
| 1.27723E-10 |

| | |
|---|---|
| | 0.000133627 |
| | 0.002210771 |
| | 1 |
| | 1 |
| | 1 |
| | 0.680830672 |
| | 0.009748088 |
| | 0.825977759 |
| | 0.191052848 |
| | 0.300017378 |
| | 1 |
| | 0.060269436 |
| | 0.129145175 |
| | 1.40604E-08 |
| | 0.977540823 |
| | 1 |
| | 1 |
| | 0.010540535 |
| | 0.267078164 |
| | 1 |
| | 9.85327E-07 |
| | 1 |
| | 0.033034622 |
| | 1 |
| | 0.261610882 |
| | 0.107425983 |
| | 1 |
| | 0.014810716 |
| | 1.44797E-07 |
| | 8.69238E-10 |
| | 0.451685121 |
| | 0.004550552 |
| | 1 |
| | 1.18182E-05 |
| | 1 |

| | |
|---|---|
| | 0.000142129 |
| | 0.180267451 |
| | 0.004165069 |
| | 0.074438713 |
| | 4.07872E-05 |
| | 0.801432855 |
| | 5.16395E-05 |
| | 0.028941717 |
| | 0.019731647 |
| | 0.470024578 |
| | 0.108023643 |
| | 0.000135864 |
| | 0.014345677 |
| | 0.616717297 |
| | 1 |
| | 0.114431076 |
| | 0.078669872 |
| | 0.000246465 |
| | 2.4583E-05 |
| | 0.835798037 |
| | 0.000409416 |
| | 0.026016531 |
| | 0.704967974 |
| | 0.012339527 |
| | 1 |
| | 1 |
| | 1 |
| | 0.195679878 |
| | 8.03411E-07 |
| | 0.910339428 |
| | 1 |
| | 1 |
| | 1 |
| | 1 |
| | 1 |

| | |
|---|---|
| | 0.064196163 |
| | 1 |
| | 1 |
| | 1 |
| | 0.535804465 |
| | 0.630340284 |
| | 0.020697213 |
| | 1 |
| | 0.227294101 |
| | 0.001033394 |
| | 0.005381706 |
| | 1 |
| | 0.344407552 |
| | 0.000915788 |
| | 0.050102298 |
| | 1 |
| | 1 |
| | 0.224887446 |
| | 0.030103754 |
| | 1 |
| | 2.76333E-07 |
| | 1 |
| | 1 |
| | 0.847881113 |
| | 9.56694E-06 |
| | 1 |
| | 0.046677384 |
| | 0.988919513 |
| | 0.782248328 |
| | 1 |
| | 1 |
| | 0.207866098 |
| | 0.382905639 |
| | 0.619547723 |
| | 0.824875824 |

| | |
|---|---|
| | 1 |
| | 0.007997854 |
| | 0.000291651 |
| | 1 |
| | 0.074415142 |
| | 0.105196211 |
| | 0.138981809 |
| | 0.737058917 |
| | 1.99047E-08 |
| | 0.112297932 |
| | 0.271743328 |
| | 0.568324338 |
| | 0.088298249 |
| | 0.01034072 |
| | 0.083012445 |
| | 0.397132005 |
| | 1 |
| | 7.72452E-06 |
| | 0.000654035 |
| | 1 |
| | 2.08538E-10 |
| | 1 |
| | 0.755600501 |
| | 0.044099021 |
| | 0.000109752 |
| | 1 |
| | 0.001036791 |
| | 0.000331478 |
| | 1 |
| | 1 |
| | 0.00577564 |
| | 0.045054816 |
| | 0.000297259 |
| | 0.000209965 |
| | 0.95504256 |

| | |
|---|---|
| | 0.000292891 |
| | 1 |
| | 0.089959605 |
| | 0.029629177 |
| | 0.373870319 |
| | 0.770723397 |
| | 1 |
| | 0.047424552 |
| | 1 |
| | 3.02279E-07 |
| | 0.445891267 |
| | 0.002690537 |
| | 0.000195041 |
| | 0.006086845 |
| | 1.88233E-06 |
| | 0.000391459 |
| | 1.74116E-06 |
| | 0.646530828 |
| | 0.006579544 |
| | 2.5922E-09 |
| | 0.162607709 |
| | 3.94064E-05 |
| | 0.037252691 |
| | 0.000309188 |
| | 0.546361908 |
| | 0.000238418 |
| | 1 |
| | 0.000158785 |
| | 1 |
| | 0.056082613 |
| | 1 |
| | 0.011821264 |
| | 0.000855334 |
| | 0.000109747 |
| | 2.49619E-07 |

| | |
|---|---|
| | 0.00011936 |
| | 0.260816333 |
| | 1 |
| | 1 |
| | 0.069184653 |
| | 0.000264582 |
| | 0.000734037 |
| | 0.055098079 |
| | 0.977480218 |
| | 0.01874206 |
| | 0.07089574 |
| | 0.004857461 |
| | 4.64171E-06 |
| | 1 |
| | 0.066464624 |
| | 1 |
| | 0.309433697 |
| | 0.148647817 |
| | 1 |
| | 0.002836511 |
| | 1 |
| | 1 |
| | 0.943649641 |
| | 1.28024E-08 |
| | 4.85142E-06 |
| | 1 |
| | 1 |
| | 5.12945E-06 |
| | 1 |
| | 0.02043602 |
| | 1 |
| | 0.045835712 |
| | 0.012333695 |
| | 1 |
| | 1 |

| | |
|---|---|
| | 0.622278846 |
| | 0.052896389 |
| | 1 |
| | 0.92343069 |
| | 1 |
| | 1 |
| | 0.029737265 |
| | 1 |
| | 0.745026126 |
| | 0.262696176 |
| | 1 |
| | 0.85483296 |
| | 1 |
| | 1 |
| | 1 |
| | 0.000959411 |
| | 1 |
| | 1 |
| | 0.685455559 |
| | 1 |
| | 1 |
| | 1 |
| | 1 |
| | 0.912515641 |
| | 1 |
| | 0.350225092 |
| | 0.347001629 |
| | 0.357924392 |
| | 1 |
| | 1 |
| | 0.46845871 |
| | 0.074459589 |
| | 1 |
| | 1 |
| | 0.945273808 |

| |
|---:|
| 1 |
| 0.944529449 |
| 1 |
| 0.432880676 |
| 0.730810521 |
| 1 |
| 0.186804065 |
| 0.07246294 |
| 1.80762E-07 |
| 0.006465037 |
| 1 |
| 1 |
| 0.624722752 |
| 0.00527343 |
| 1 |
| 1 |
| 0.822506052 |
| 0.009682094 |
| 1 |
| 0.590066799 |
| 0.00340665 |
| 1.35018E-10 |
| 8.17337E-05 |
| 1 |
| 1 |
| 0.006203484 |
| 2.13453E-05 |
| 0.982914547 |
| 0.892353288 |
| 0.031510659 |
| 1 |
| 1 |
| 1 |
| 1 |
| 0.000610693 |

| | |
|---|---|
| | 0.006013011 |
| | 1 |
| | 1 |
| | 0.350225092 |
| | 1 |
| | 1 |
| | 1 |
| | 0.517988781 |
| | 1 |
| | 1 |
| | 0.351055802 |
| | 1 |
| | 1 |
| | 0.905701282 |
| | 1 |
| | 1 |
| | 9.86791E-07 |
| | 1 |
| | 1 |
| | 0.123144846 |
| | 1 |
| | 1 |
| | 0.504988379 |
| | 0.9219797 |
| | 1 |
| | 1 |
| | 0.994322501 |
| | 1 |
| | 1 |
| | 1 |
| | 0.028445437 |
| | 0.098742604 |
| | 1 |
| | 4.54094E-07 |
| | 0.972897254 |

| |
|---|
| 1 |
| 1 |
| 1 |
| 1 |
| 1 |
| 0.154367786 |
| 2.42565E-07 |
| 0.958422685 |
| 0.001217139 |
| 0.167187739 |
| 1 |
| 0.655638428 |
| 1 |
| 0.000101461 |
| 8.96733E-08 |
| 1 |
| 1 |
| 0.215340317 |
| 0.001376306 |
| 1 |
| 0.274188628 |
| 1 |
| 1 |
| 1 |
| 1 |
| 0.122902074 |
| 0.154932186 |
| 0.030889875 |
| 0.379773643 |
| 1 |
| 0.015377378 |
| 1 |
| 1 |
| 1 |
| 8.18972E-05 |

| |
|---|
| 0.29515674 |
| 0.167707549 |
| 0.01487343 |
| 1 |
| 0.02652572 |
| 1 |
| 1 |
| 1 |
| 0.0668845 |
| 1 |
| 1 |
| 0.040246703 |
| 1 |
| 0.001672618 |
| 0.444881605 |
| 0.000413697 |
| 0.482801073 |
| 1 |
| 1 |
| 0.706996544 |
| 0.272668415 |
| 0.177889597 |
| 0.174977062 |
| 1 |
| 0.282611534 |
| 0.087216963 |
| 0.189281622 |
| 0.223787082 |
| 1 |
| 1 |
| 0.006885771 |
| 0.001172043 |
| 1 |
| 0.001987837 |
| 1 |

| | |
|---|---|
| | 0.000762218 |
| | 0.035551122 |
| | 0.354813274 |
| | 1 |
| | 1 |
| | 0.113666448 |
| | 0.617939805 |
| | 0.160697379 |
| | 1 |
| | 0.011770961 |
| | 2.0957E-09 |
| | 0.000699951 |
| | 0.037393364 |
| | 1 |
| | 1 |
| | 0.346677943 |
| | 0.851989196 |
| | 1 |
| | 1 |
| | 1 |
| | 1 |
| | 0.328042349 |
| | 5.46494E-09 |
| | 0.097917068 |
| | 0.237671099 |
| | 1 |
| | 0.011572048 |
| | 0.580484068 |
| | 1 |
| | 1 |
| | 1 |
| | 1 |
| | 0.88506185 |
| | 0.040352097 |
| | 1 |

| |
|---:|
| 1 |
| 0.073347327 |
| 0.068565731 |
| 0.58874457 |
| 0.111735298 |
| 0.027503962 |
| 0.901540028 |
| 1 |
| 0.46164454 |
| 1 |
| 1 |
| 0.997219243 |
| 0.587883665 |
| 0.00154993 |
| 0.164906169 |

| |
|---|
| Adj. P-value: (Rabbitserum_12022024_Newanalysis) / (Water_pos_12022024_Newanalysis) |
| 1 |
| 0.147274541 |
| 8.78311E-08 |
| 0.008004295 |
| 0.025733921 |
| 1 |
| 0.07138554 |
| 0.00985366 |
| 0.601836115 |
| 1 |
| 1 |
| 1.90795E-06 |
| 0.482552995 |
| 0.08349607 |
| 0.110008338 |
| 1 |
| 1 |
| 1.25336E-11 |
| 5.29765E-05 |
| 1.25336E-11 |
| 1 |
| 1.73767E-10 |
| 7.00916E-05 |
| 1.25336E-11 |
| 1.25336E-11 |
| 1 |
| 0.966064913 |
| 0.220652367 |
| 1 |
| 0.013271801 |
| 0.397204592 |
| 0.007167795 |
| 0.00765555 |

| | |
|---|---|
| | 1 |
| | 1.26177E-11 |
| | 4.39022E-07 |
| | 1.43159E-05 |
| | 0.855396772 |
| | 0.744129037 |
| | 1.24178E-05 |
| | 0.329841963 |
| | 1 |
| | 0.043398725 |
| | 7.79106E-07 |
| | 0.000277922 |
| | 1.46093E-08 |
| | 1.97359E-05 |
| | 1.41349E-05 |
| | 0.240317565 |
| | 0.128687546 |
| | 0.055073286 |
| | 0.011850023 |
| | 5.94192E-07 |
| | 0.037200357 |
| | 1 |
| | 0.966191854 |
| | 5.75971E-07 |
| | 0.516826508 |
| | 0.31259126 |
| | 1.78748E-06 |
| | 1 |
| | 1 |
| | 0.006228576 |
| | 2.84245E-07 |
| | 0.012472812 |
| | 0.028156824 |
| | 1 |
| | 1 |

| | |
|---|---|
| | 0.597789901 |
| | 1 |
| | 0.057824286 |
| | 0.002921322 |
| | 0.18764521 |
| | 0.581041617 |
| | 1 |
| | 1 |
| | 1.25336E-11 |
| | 7.11388E-08 |
| | 1.86221E-07 |
| | 1.78267E-10 |
| | 0.006203694 |
| | 0.006635718 |
| | 0.000320061 |
| | 0.098325269 |
| | 1.99891E-07 |
| | 1 |
| | 0.001057696 |
| | 0.449270435 |
| | 1 |
| | 1 |
| | 0.001109802 |
| | 1.29542E-06 |
| | 0.000377263 |
| | 0.000268908 |
| | 8.91102E-06 |
| | 0.284189397 |
| | 0.000126977 |
| | 0.011693582 |
| | 0.025261734 |
| | 1 |
| | 4.76224E-08 |
| | 0.000114569 |
| | 0.467931878 |

| | |
|---|---|
| | 2.54905E-07 |
| | 0.000676417 |
| | 4.99825E-06 |
| | 1 |
| | 0.755312233 |
| | 0.530009782 |
| | 0.022837194 |
| | 1 |
| | 0.00013597 |
| | 1 |
| | 1 |
| | 0.00027809 |
| | 0.534652877 |
| | 1 |
| | 1.25336E-11 |
| | 0.014943842 |
| | 0.902624075 |
| | 0.057424983 |
| | 1.25336E-11 |
| | 1 |
| | 0.017858202 |
| | 0.577410688 |
| | 0.733990664 |
| | 1 |
| | 1 |
| | 1 |
| | 0.020674254 |
| | 0.520275097 |
| | 1.0845E-05 |
| | 1.25336E-11 |
| | 5.02856E-07 |
| | 2.30539E-05 |
| | 1 |
| | 0.667964241 |
| | 1 |

| | |
|---|---|
| | 0.811788163 |
| | 1 |
| | 1 |
| | 1 |
| | 0.848821933 |
| | 1 |
| | 1 |
| | 1 |
| | 0.622336914 |
| | 1 |
| | 0.589952465 |
| | 0.000260904 |
| | 1.25336E-11 |
| | 0.326147597 |
| | 1.04936E-08 |
| | 1.25901E-11 |
| | 1 |
| | 0.068212864 |
| | 0.595439508 |
| | 0.060346192 |
| | 0.369055399 |
| | 0.000160244 |
| | 3.17588E-11 |
| | 3.65771E-07 |
| | 0.324505287 |
| | 0.399209912 |
| | 3.45133E-06 |
| | 1 |
| | 1 |
| | 0.000661557 |
| | 5.99952E-08 |
| | 1 |
| | 2.2928E-07 |
| | 0.005477611 |
| | 1.07839E-07 |

| |
|---|
| 0.001785388 |
| 1.04377E-08 |
| 0.001599195 |
| 1.25336E-11 |
| 1.38179E-06 |
| 2.20173E-05 |
| 0.0089954 |
| 0.000316167 |
| 4.18474E-07 |
| 4.9256E-06 |
| 0.000104588 |
| 0.005210014 |
| 1.25336E-11 |
| 1.89892E-08 |
| 1 |
| 6.50358E-11 |
| 0.002989871 |
| 0.013584607 |
| 1.25336E-11 |
| 7.2333E-07 |
| 1.27014E-05 |
| 4.51265E-07 |
| 0.000324957 |
| 0.249887476 |
| 0.839430279 |
| 0.829174084 |
| 0.229104033 |
| 0.206232939 |
| 0.062665428 |
| 2.78009E-06 |
| 0.007701323 |
| 0.047508783 |
| 1.98067E-08 |
| 1.12846E-07 |
| 5.98341E-06 |

| | |
|---|---|
| | 0.044453335 |
| | 1 |
| | 1 |
| | 1 |
| | 0.600741847 |
| | 1 |
| | 0.025478536 |
| | 0.000785563 |
| | 1 |
| | 0.008362917 |
| | 0.029239713 |
| | 1 |
| | 0.229281646 |
| | 1 |
| | 1 |
| | 1 |
| | 0.925396104 |
| | 9.9658E-08 |
| | 2.77125E-07 |
| | 1 |
| | 2.22708E-06 |
| | 0.048845679 |
| | 0.166289238 |
| | 0.001272986 |
| | 1.15716E-10 |
| | 0.000273974 |
| | 0.158094536 |
| | 1 |
| | 1 |
| | 1 |
| | 0.060308647 |
| | 0.600338682 |
| | 0.193783972 |
| | 1 |
| | 1.58055E-09 |

| | |
|---|---|
| | 0.001634977 |
| | 9.05737E-05 |
| | 1 |
| | 1 |
| | 0.046389466 |
| | 1.27901E-06 |
| | 1.54717E-07 |
| | 1 |
| | 0.941150379 |
| | 1.25336E-11 |
| | 2.62722E-08 |
| | 0.001438384 |
| | 0.236551733 |
| | 1.25336E-11 |
| | 0.000782081 |
| | 3.58584E-06 |
| | 0.007391773 |
| | 0.388326326 |
| | 0.854511349 |
| | 1.25336E-11 |
| | 0.020633382 |
| | 0.001109012 |
| | 0.00549171 |
| | 0.368962509 |
| | 0.066765853 |
| | 0.945459918 |
| | 0.003728362 |
| | 1 |
| | 1.76935E-05 |
| | 0.005596188 |
| | 9.67951E-06 |
| | 3.76189E-07 |
| | 1 |
| | 1 |
| | 1.25336E-11 |

| | |
|---|---|
| | 1.25336E-11 |
| | 2.02156E-09 |
| | 0.012951157 |
| | 0.070075398 |
| | 0.004648432 |
| | 0.015459567 |
| | 3.41028E-11 |
| | 1 |
| | 0.666217774 |
| | 0.000361615 |
| | 0.668215924 |
| | 0.238603574 |
| | 0.092916197 |
| | 1 |
| | 0.640654545 |
| | 1 |
| | 1.04498E-07 |
| | 1 |
| | 0.014233764 |
| | 0.115184599 |
| | 5.79725E-08 |
| | 1.25392E-07 |
| | 0.099434894 |
| | 1.7547E-10 |
| | 0.055173389 |
| | 0.004015345 |
| | 0.023406164 |
| | 3.62237E-11 |
| | 4.58282E-05 |
| | 1.25336E-11 |
| | 0.000545412 |
| | 8.15872E-07 |
| | 0.033644519 |
| | 8.81668E-08 |
| | 0.316712993 |

| | |
|---|---|
| | 1.25336E-11 |
| | 0.001045342 |
| | 8.17193E-07 |
| | 5.42973E-06 |
| | 0.034212073 |
| | 0.054515968 |
| | 2.61982E-09 |
| | 1.67427E-11 |
| | 0.000398507 |
| | 8.95548E-09 |
| | 0.002315161 |
| | 0.088495491 |
| | 0.000217625 |
| | 0.086878152 |
| | 1 |
| | 0.08981796 |
| | 7.2171E-08 |
| | 8.03473E-09 |
| | 1.59813E-11 |
| | 3.5009E-10 |
| | 2.54943E-05 |
| | 3.20434E-07 |
| | 0.015104662 |
| | 2.03338E-09 |
| | 0.000358026 |
| | 1 |
| | 0.683979421 |
| | 2.88525E-07 |
| | 0.015258504 |
| | 0.001023708 |
| | 0.005121923 |
| | 0.463998497 |
| | 3.57748E-07 |
| | 0.413839248 |
| | 1 |

| |
|---|
| 1 |
| 1 |
| 0.692834903 |
| 0.365336074 |
| 0.131510613 |
| 1 |
| 0.172390364 |
| 6.10659E-07 |
| 0.258127714 |
| 2.45935E-09 |
| 1.11893E-09 |
| 9.48244E-08 |
| 2.08687E-08 |
| 0.000855737 |
| 0.24517129 |
| 0.732832104 |
| 0.220382059 |
| 0.034428762 |
| 0.000493298 |
| 1.45274E-06 |
| 1.25336E-11 |
| 1.20372E-06 |
| 1 |
| 7.73709E-07 |
| 8.8985E-11 |
| 1 |
| 0.000360949 |
| 0.000253506 |
| 0.641919779 |
| 0.047323222 |
| 1 |
| 5.30225E-05 |
| 3.72422E-07 |
| 0.10587709 |
| 1 |

| |
|---|
| 0.0088945 |
| 0.159944809 |
| 1.93287E-11 |
| 9.3344E-09 |
| 0.001218687 |
| 0.001910589 |
| 8.56337E-09 |
| 2.73436E-07 |
| 1.25336E-11 |
| 0.00024088 |
| 9.20038E-06 |
| 1.75711E-11 |
| 0.004615108 |
| 3.99124E-06 |
| 3.28661E-06 |
| 0.001331777 |
| 0.01330856 |
| 6.05252E-10 |
| 5.57619E-10 |
| 0.501740425 |
| 9.29506E-10 |
| 0.00276812 |
| 0.32646338 |
| 5.78288E-05 |
| 0.000729565 |
| 1 |
| 5.31589E-07 |
| 9.98907E-06 |
| 1 |
| 0.859905145 |
| 8.94181E-09 |
| 1 |
| 1.26177E-11 |
| 8.35949E-08 |
| 0.022907209 |

| | |
|---|---|
| | 7.30423E-09 |
| | 0.710184054 |
| | 3.51945E-06 |
| | 0.00024416 |
| | 0.002177966 |
| | 1 |
| | 1 |
| | 1 |
| | 5.52596E-07 |
| | 1.25336E-11 |
| | 9.02301E-05 |
| | 8.45769E-10 |
| | 6.02148E-09 |
| | 0.000687381 |
| | 1.25336E-11 |
| | 4.14215E-09 |
| | 1.84428E-10 |
| | 0.001008429 |
| | 5.16424E-09 |
| | 1.25336E-11 |
| | 4.44869E-06 |
| | 1.90771E-10 |
| | 6.46518E-06 |
| | 1.20647E-06 |
| | 0.000845623 |
| | 1.22433E-08 |
| | 2.03703E-08 |
| | 1.36161E-10 |
| | 0.191509279 |
| | 0.00468286 |
| | 6.26198E-06 |
| | 0.00022092 |
| | 1.94326E-08 |
| | 3.01606E-09 |
| | 1.26713E-11 |

| | |
|---|---|
| | 1.3511E-09 |
| | 6.88473E-06 |
| | 1.58847E-08 |
| | 0.647122767 |
| | 0.499307437 |
| | 8.04094E-07 |
| | 5.74149E-08 |
| | 1.67298E-07 |
| | 0.56409683 |
| | 6.58953E-10 |
| | 4.90774E-06 |
| | 6.50334E-06 |
| | 1.25336E-11 |
| | 0.070025334 |
| | 1.95506E-05 |
| | 0.182253964 |
| | 3.29349E-08 |
| | 0.652654785 |
| | 1 |
| | 3.43688E-11 |
| | 0.203510232 |
| | 0.032943608 |
| | 5.37351E-08 |
| | 1.25336E-11 |
| | 2.32725E-11 |
| | 1.20216E-05 |
| | 0.000315658 |
| | 1.25336E-11 |
| | 1 |
| | 1 |
| | 6.95063E-07 |
| | 0.007655482 |
| | 8.92268E-09 |
| | 1 |
| | 0.973531448 |

| | |
|---|---|
| | 0.222281295 |
| | 0.040025182 |
| | 1 |
| | 0.05163844 |
| | 0.0011753 |
| | 1.25336E-11 |
| | 1 |
| | 1 |
| | 0.006095228 |
| | 1 |
| | 0.12339094 |
| | 4.46476E-05 |
| | 0.167020606 |
| | 0.026800364 |
| | 0.006119893 |
| | 1.20253E-08 |
| | 9.22607E-08 |
| | 9.26387E-09 |
| | 0.004844241 |
| | 0.001634977 |
| | 0.001227687 |
| | 0.038949968 |
| | 0.988511412 |
| | 1 |
| | 0.007018891 |
| | 5.20138E-07 |
| | 0.000536033 |
| | 6.22555E-09 |
| | 1.1455E-05 |
| | 0.618750926 |
| | 5.92862E-05 |
| | 1.25522E-05 |
| | 7.96504E-07 |
| | 1 |
| | 4.08704E-09 |

| | |
|---|---|
| | 0.04366816 |
| | 0.627809853 |
| | 3.66684E-08 |
| | 0.940503965 |
| | 1.83471E-11 |
| | 0.33869098 |
| | 2.03805E-07 |
| | 1 |
| | 1.25336E-11 |
| | 1.35668E-11 |
| | 0.07361766 |
| | 0.001931379 |
| | 2.3068E-05 |
| | 0.272227934 |
| | 2.2383E-07 |
| | 0.0265164 |
| | 3.17253E-05 |
| | 1.2382E-10 |
| | 9.2454E-07 |
| | 0.539027916 |
| | 6.42892E-09 |
| | 1.25336E-11 |
| | 1.91879E-11 |
| | 0.00070782 |
| | 8.84754E-08 |
| | 2.58973E-07 |
| | 1.45894E-11 |
| | 0.00092679 |
| | 0.001519564 |
| | 3.40271E-08 |
| | 0.092520602 |
| | 0.157722597 |
| | 1.17868E-05 |
| | 1 |
| | 0.001537732 |

| | |
|---|---|
| | 1.08518E-08 |
| | 1 |
| | 5.07959E-07 |
| | 0.045368194 |
| | 1 |
| | 0.858483249 |
| | 1 |
| | 0.008661466 |
| | 0.000412566 |
| | 1 |
| | 0.010059628 |
| | 1 |
| | 5.22232E-05 |
| | 1.08413E-07 |
| | 0.000915256 |
| | 0.587444069 |
| | 2.78379E-07 |
| | 1 |
| | 0.658325063 |
| | 0.002994175 |
| | 0.002109425 |
| | 0.099928394 |
| | 1 |
| | 4.98264E-05 |
| | 0.416117034 |
| | 1 |
| | 0.002617596 |
| | 1 |
| | 0.262950014 |
| | 1 |
| | 0.019706494 |
| | 1.20103E-06 |
| | 0.453163164 |
| | 3.23728E-07 |
| | 1 |

| | |
|---|---|
| | 0.136591531 |
| | 1 |
| | 0.922065238 |
| | 0.523622795 |
| | 1 |
| | 1.30778E-06 |
| | 1.25336E-11 |
| | 1.16075E-06 |
| | 3.23785E-08 |
| | 0.266095212 |
| | 0.00706581 |
| | 0.405690433 |
| | 2.16902E-07 |
| | 1 |
| | 1.25336E-11 |
| | 0.006522166 |
| | 0.022205345 |
| | 0.198797541 |
| | 0.002174749 |
| | 3.22287E-05 |
| | 0.047579109 |
| | 0.000143432 |
| | 0.679799534 |
| | 0.111973074 |
| | 0.039539194 |
| | 0.01411616 |
| | 0.008982783 |
| | 7.08336E-10 |
| | 2.84844E-05 |
| | 0.336160842 |
| | 3.91405E-10 |
| | 0.021576813 |
| | 0.000228094 |
| | 0.232092387 |
| | 1.95935E-10 |

| | |
|---|---|
| | 0.746904799 |
| | 0.018415941 |
| | 7.1625E-06 |
| | 0.073512476 |
| | 0.000433799 |
| | 0.31337606 |
| | 0.062761032 |
| | 3.32521E-07 |
| | 0.000182557 |
| | 0.323949715 |
| | 7.15869E-05 |
| | 0.000393056 |
| | 1 |
| | 0.000364653 |
| | 0.00196423 |
| | 0.004454242 |
| | 0.013384462 |
| | 0.075950552 |
| | 0.068067668 |
| | 0.590886435 |
| | 0.257501303 |
| | 0.027754091 |
| | 0.024270659 |
| | 1 |
| | 0.034112935 |
| | 0.004527564 |
| | 0.028431556 |
| | 0.022836156 |
| | 1 |
| | 1 |
| | 8.02539E-05 |
| | 4.30573E-05 |
| | 1 |
| | 0.025751934 |
| | 1 |

| | |
|---|---|
| | 0.065581056 |
| | 0.006617518 |
| | 0.528830027 |
| | 0.000133693 |
| | 0.052095859 |
| | 1.38129E-10 |
| | 1 |
| | 1.45058E-05 |
| | 0.406201682 |
| | 4.32766E-06 |
| | 1.25336E-11 |
| | 5.86981E-09 |
| | 1.99496E-07 |
| | 0.001494945 |
| | 0.006654229 |
| | 0.002798027 |
| | 3.10407E-07 |
| | 1 |
| | 1 |
| | 0.547776632 |
| | 1 |
| | 1 |
| | 9.60787E-07 |
| | 0.003316039 |
| | 1 |
| | 0.003926699 |
| | 0.107928024 |
| | 0.12115965 |
| | 0.003473455 |
| | 1 |
| | 0.012434762 |
| | 0.004683937 |
| | 0.470521567 |
| | 0.008226526 |
| | 0.000985738 |

| |
|---|
| 0.37147399 |
| 0.334336202 |
| 1 |
| 0.032039232 |
| 0.000810822 |
| 8.58365E-08 |
| 0.00075072 |
| 1 |
| 1 |
| 1 |
| 0.534936615 |
| 0.62284351 |
| 0.764656348 |
| 0.00015708 |
| 1.28227E-05 |

| Adj. P-value: (SM_pos_12022024_Newanalyis) / (Water_pos_12022024_Newanalysis) |
|---|
| 1 |
| 1 |
| 1 |
| 0.136715876 |
| 0.277466377 |
| 1 |
| 1 |
| 1 |
| 0.020620959 |
| 0.082998055 |
| 0.011067123 |
| 6.74791E-07 |
| 1 |
| 0.00059086 |
| 1 |
| 1 |
| 1 |
| 0.313227317 |
| 0.120032448 |
| 0.364937229 |
| 1 |
| 1 |
| 0.077879083 |
| 1 |
| 0.112950679 |
| 1 |
| 1 |
| 1 |
| 0.071008196 |
| 1 |
| 1 |
| 0.009353986 |
| 1 |

| |
|---|
| 0.020083945 |
| 4.70828E-05 |
| 1 |
| 0.031404209 |
| 0.525522324 |
| 0.132618891 |
| 0.253889779 |
| 0.483444214 |
| 1 |
| 0.540116116 |
| 0.000429059 |
| 1 |
| 1 |
| 1 |
| 1 |
| 0.046323996 |
| 1 |
| 0.083441995 |
| 1 |
| 1 |
| 1 |
| 3.98909E-05 |
| 1 |
| 9.7032E-05 |
| 1 |
| 1 |
| 1 |
| 1 |
| 0.112030938 |
| 1 |
| 0.360445075 |
| 0.504033755 |
| 0.687351302 |
| 0.678302427 |
| 0.000562035 |

| |
|---|
| 0.066413805 |
| 1 |
| 1 |
| 0.058867627 |
| 0.253439106 |
| 4.91239E-05 |
| 1 |
| 0.005791384 |
| 8.03708E-05 |
| 1 |
| 0.000476633 |
| 1.05732E-07 |
| 0.551307704 |
| 0.002907605 |
| 1 |
| 0.057682594 |
| 0.850115124 |
| 1 |
| 0.057270391 |
| 0.060027827 |
| 0.205808075 |
| 0.917429032 |
| 2.17481E-05 |
| 0.001244542 |
| 1 |
| 1 |
| 1 |
| 1 |
| 0.376502745 |
| 1 |
| 0.000432252 |
| 0.346876086 |
| 1 |
| 1 |
| 2.073E-06 |

| |
|---|
| 0.00037256 |
| 1 |
| 1 |
| 1 |
| 1 |
| 1 |
| 1 |
| 0.027970674 |
| 1 |
| 1 |
| 0.367854987 |
| 1 |
| 0.512479919 |
| 1 |
| 0.069510239 |
| 0.119106885 |
| 0.003949971 |
| 1 |
| 1 |
| 0.076843079 |
| 0.002061078 |
| 0.859784374 |
| 1 |
| 1 |
| 7.43513E-09 |
| 1 |
| 0.118476851 |
| 0.03322891 |
| 0.546747675 |
| 0.221138683 |
| 1 |
| 0.000622652 |
| 0.252573436 |
| 1 |
| 1 |

| |
|---|
| 0.2143984 |
| 1 |
| 0.169566355 |
| 1 |
| 0.804514262 |
| 1 |
| 1 |
| 1 |
| 1 |
| 1 |
| 1 |
| 1 |
| 0.001513512 |
| 0.346730791 |
| 1 |
| 1.06585E-05 |
| 0.205974597 |
| 0.007770059 |
| 1 |
| 1 |
| 1 |
| 0.974102909 |
| 0.583809426 |
| 0.038406358 |
| 1.40427E-06 |
| 0.000826404 |
| 1 |
| 1 |
| 1 |
| 1 |
| 0.001786111 |
| 1 |
| 1 |
| 0.017620784 |
| 0.017855309 |

| |
|---|
| 0.004434301 |
| 1 |
| 1 |
| 6.56601E-09 |
| 0.092791456 |
| 1 |
| 0.337441649 |
| 0.542732002 |
| 8.09635E-10 |
| 0.000623624 |
| 1 |
| 0.000499859 |
| 2.06596E-05 |
| 2.74001E-05 |
| 1 |
| 9.35898E-08 |
| 1 |
| 0.000268145 |
| 1.49988E-10 |
| 1 |
| 0.050900488 |
| 0.061100162 |
| 1 |
| 0.717764001 |
| 1 |
| 1 |
| 1 |
| 0.005166119 |
| 1 |
| 0.129238417 |
| 0.036557046 |
| 0.02051542 |
| 0.000330947 |
| 1 |
| 3.09559E-07 |

| |
|---|
| 1 |
| 1 |
| 0.87148187 |
| 1 |
| 1 |
| 1.12906E-05 |
| 5.05042E-08 |
| 0.009351515 |
| 0.001140384 |
| 0.000199392 |
| 0.505457454 |
| 1 |
| 0.021519002 |
| 0.276703837 |
| 1 |
| 1 |
| 0.047115096 |
| 4.50669E-05 |
| 1 |
| 1 |
| 0.004236554 |
| 1 |
| 0.016015003 |
| 0.000214659 |
| 0.085367304 |
| 1 |
| 1 |
| 0.133158134 |
| 0.469524168 |
| 0.000747144 |
| 0.89086839 |
| 1 |
| 1 |
| 0.415866077 |
| 1 |

| |
|---|
| 0.025702875 |
| 0.026096815 |
| 1 |
| 1 |
| 1 |
| 0.000265824 |
| 0.004711109 |
| 1 |
| 0.286797891 |
| 1.97095E-06 |
| 2.37014E-05 |
| 0.037745434 |
| 1 |
| 2.16034E-07 |
| 0.0488163 |
| 0.002811941 |
| 0.045133662 |
| 0.837767828 |
| 1 |
| 1 |
| 1 |
| 1.36582E-05 |
| 1 |
| 7.81508E-05 |
| 0.917787867 |
| 1 |
| 0.428873221 |
| 1 |
| 0.000583155 |
| 1 |
| 0.003142733 |
| 1 |
| 1 |
| 1 |
| 7.74934E-11 |

| |
|---|
| 2.45628E-07 |
| 1 |
| 1 |
| 0.024440345 |
| 1.22843E-05 |
| 0.308232971 |
| 0.0014439 |
| 1 |
| 1 |
| 1 |
| 0.002056992 |
| 0.044899648 |
| 0.061564294 |
| 0.3286484 |
| 1 |
| 0.83266214 |
| 0.758937582 |
| 0.003500484 |
| 0.000707815 |
| 0.944738291 |
| 1 |
| 0.047706715 |
| 1 |
| 0.195388416 |
| 0.45367296 |
| 0.04114822 |
| 1 |
| 0.336389805 |
| 0.895624008 |
| 6.79605E-10 |
| 1 |
| 0.059499542 |
| 0.005988876 |
| 2.12803E-10 |
| 0.817444138 |

| |
|---|
| 3.04831E-07 |
| 1 |
| 0.949008579 |
| 1 |
| 2.3646E-05 |
| 1 |
| 5.24103E-07 |
| 0.877019519 |
| 8.66908E-10 |
| 0.024255592 |
| 0.001885448 |
| 6.71093E-06 |
| 4.47691E-06 |
| 0.021276377 |
| 0.000419475 |
| 0.18185881 |
| 0.006743919 |
| 0.779902593 |
| 0.216489915 |
| 0.00022527 |
| 4.80674E-10 |
| 0.000108427 |
| 0.035619121 |
| 0.283670473 |
| 1 |
| 1.82065E-05 |
| 0.255660705 |
| 0.192588723 |
| 1.05373E-05 |
| 0.902486811 |
| 0.583809426 |
| 1 |
| 0.053298537 |
| 1 |
| 1 |

| |
|---|
| 0.00187567 |
| 1 |
| 0.285413647 |
| 0.454830835 |
| 0.701753596 |
| 1 |
| 0.002058886 |
| 1 |
| 0.637836253 |
| 0.000242999 |
| 0.030110932 |
| 1 |
| 0.024346656 |
| 1 |
| 1 |
| 0.000173268 |
| 1 |
| 0.568669849 |
| 0.07902343 |
| 0.108836678 |
| 1.86352E-07 |
| 0.02592869 |
| 0.72225015 |
| 1 |
| 0.000124544 |
| 1 |
| 0.042979341 |
| 1 |
| 0.071672185 |
| 0.325323909 |
| 1 |
| 0.671151634 |
| 1 |
| 0.549277341 |
| 8.10124E-08 |

| |
|---|
| 0.032302954 |
| 0.696933209 |
| 0.601645269 |
| 0.753361011 |
| 1 |
| 0.101330168 |
| 0.921921454 |
| 0.313835754 |
| 0.14010475 |
| 1 |
| 0.109006218 |
| 0.219798526 |
| 0.016088571 |
| 1 |
| 0.000132185 |
| 1 |
| 1 |
| 2.19188E-05 |
| 0.005028604 |
| 0.018796318 |
| 9.38605E-11 |
| 0.001171383 |
| 0.059119693 |
| 0.551325569 |
| 0.000979841 |
| 1 |
| 0.396144458 |
| 3.19828E-08 |
| 0.767059641 |
| 0.532687113 |
| 1 |
| 0.862162087 |
| 1.24949E-08 |
| 2.94614E-06 |
| 0.039936508 |

| |
|---|
| 0.000280859 |
| 1.47102E-05 |
| 6.34843E-06 |
| 0.000121202 |
| 2.75525E-05 |
| 1 |
| 2.08069E-08 |
| 0.000199335 |
| 0.612659799 |
| 1.68443E-06 |
| 0.962751606 |
| 0.000861997 |
| 0.005049866 |
| 5.50669E-08 |
| 7.43513E-09 |
| 1.58915E-09 |
| 0.434035364 |
| 1 |
| 2.154E-07 |
| 7.74934E-11 |
| 0.318812774 |
| 4.73395E-06 |
| 0.005991753 |
| 1.85121E-07 |
| 1.10196E-05 |
| 7.74934E-11 |
| 0.001492502 |
| 1 |
| 0.630463859 |
| 1 |
| 0.005958788 |
| 0.00433404 |
| 0.000270413 |
| 0.0021856 |
| 2.89667E-08 |

| |
|---|
| 1 |
| 1 |
| 0.086789756 |
| 0.031621071 |
| 1 |
| 1.07504E-05 |
| 1.15134E-06 |
| 7.44448E-07 |
| 0.458609411 |
| 3.23023E-05 |
| 0.000583155 |
| 4.333E-09 |
| 0.000293105 |
| 0.047436398 |
| 0.08229926 |
| 0.273989745 |
| 0.013887167 |
| 0.699774527 |
| 1 |
| 0.000153729 |
| 1 |
| 0.354935322 |
| 1 |
| 1.19532E-08 |
| 1.69189E-06 |
| 1 |
| 0.000154455 |
| 1.41341E-05 |
| 0.013009206 |
| 0.055223113 |
| 1 |
| 0.002438814 |
| 0.002775087 |
| 1 |
| 0.735657228 |

| |
|---|
| 1 |
| 1.04102E-05 |
| 1 |
| 1 |
| 0.752710838 |
| 0.000125262 |
| 0.033745068 |
| 1 |
| 0.366638011 |
| 0.708633438 |
| 0.683174922 |
| 0.003038501 |
| 1 |
| 0.016516783 |
| 0.328624664 |
| 0.009735256 |
| 0.003296611 |
| 9.51763E-08 |
| 1 |
| 2.09454E-06 |
| 0.198682658 |
| 7.54778E-05 |
| 5.28372E-05 |
| 1 |
| 0.173363277 |
| 1 |
| 0.008892116 |
| 0.888253828 |
| 1 |
| 1 |
| 0.020755061 |
| 0.561893436 |
| 1 |
| 0.15916194 |
| 0.334804625 |

| |
|---|
| 0.006648466 |
| 0.350630145 |
| 1.20709E-06 |
| 0.911445086 |
| 7.74934E-11 |
| 1 |
| 1 |
| 1 |
| 0.903114126 |
| 1.0839E-05 |
| 1 |
| 0.168335462 |
| 0.000506857 |
| 1 |
| 1 |
| 0.000266259 |
| 0.05343047 |
| 7.74934E-11 |
| 0.111599371 |
| 0.275995225 |
| 3.02146E-06 |
| 0.060201367 |
| 7.74934E-11 |
| 1 |
| 1.2641E-10 |
| 0.120536473 |
| 0.53634911 |
| 0.000437041 |
| 7.74934E-11 |
| 2.41735E-09 |
| 1 |
| 0.92774682 |
| 1 |
| 0.001697075 |
| 1 |

| |
|---|
| 0.00029873 |
| 0.711243745 |
| 1 |
| 0.032655682 |
| 1 |
| 0.653492652 |
| 1 |
| 0.069678872 |
| 1 |
| 0.04309174 |
| 0.003410046 |
| 1 |
| 3.28144E-07 |
| 1 |
| 0.317652462 |
| 1 |
| 0.947750695 |
| 0.40690147 |
| 0.020849158 |
| 0.040321243 |
| 4.83767E-09 |
| 1 |
| 1 |
| 0.690052546 |
| 1 |
| 1 |
| 0.01827886 |
| 1 |
| 1 |
| 0.506388306 |
| 0.779067891 |
| 0.001379383 |
| 0.715556593 |
| 3.4922E-07 |
| 1 |

| |
|---|
| 0.442361054 |
| 0.901153796 |
| 1 |
| 0.803810006 |
| 0.147189877 |
| 1 |
| 0.911954447 |
| 1 |
| 0.000289214 |
| 7.44198E-07 |
| 0.709379668 |
| 1 |
| 0.997585222 |
| 1 |
| 9.26053E-07 |
| 1 |
| 1 |
| 0.090968916 |
| 0.775833261 |
| 0.365396996 |
| 0.145757127 |
| 0.540861264 |
| 1 |
| 1 |
| 0.884575724 |
| 0.03218061 |
| 0.753531926 |
| 1 |
| 0.004410561 |
| 0.945429148 |
| 0.774379914 |
| 0.489617974 |
| 1 |
| 0.033315974 |
| 0.043776525 |

| |
|---|
| 0.601211617 |
| 5.48973E-05 |
| 0.144267437 |
| 0.496754195 |
| 0.000265449 |
| 5.58609E-06 |
| 0.00160582 |
| 7.74934E-11 |
| 3.11226E-05 |
| 1 |
| 7.57565E-09 |
| 0.000687982 |
| 1 |
| 7.01335E-09 |
| 1 |
| 1 |
| 0.000520553 |
| 0.874757626 |
| 0.004463072 |
| 1 |
| 3.05776E-05 |
| 0.074364473 |
| 0.047259777 |
| 1 |
| 0.227430033 |
| 0.038700915 |
| 0.144581162 |
| 1 |
| 1 |
| 0.38145336 |
| 0.643293508 |
| 0.054999654 |
| 1 |
| 5.75136E-10 |
| 1 |

| |
|---|
| 1 |
| 0.047757831 |
| 0.000128163 |
| 0.028723442 |
| 0.123899874 |
| 0.982217783 |
| 1 |
| 0.001787713 |
| 0.973701313 |
| 1 |
| 5.43209E-07 |
| 8.0392E-05 |
| 0.030984367 |
| 0.440154791 |
| 0.452202565 |
| 0.022762355 |
| 1 |
| 0.923393593 |
| 0.202167398 |
| 1 |
| 0.948692853 |
| 1 |
| 0.353523347 |
| 1 |
| 0.012717039 |
| 1 |
| 0.000356454 |
| 0.131872689 |
| 0.002946355 |
| 1 |
| 0.072757528 |
| 0.5559978 |
| 0.425960633 |
| 1 |
| 0.662529788 |

| |
|---|
| 1 |
| 0.072488469 |
| 1 |
| 0.002360261 |
| 0.560162218 |
| 0.222646029 |
| 1 |
| 1 |
| 0.008266939 |
| 1 |
| 1 |
| 1 |
| 0.758344768 |
| 0.027205324 |
| 2.59551E-07 |
