## Supplementary file 3 for "A lipid compendium of a metabolically compromised bacterium provides insights into lipid acquisition, biosynthesis, and metabolism"

### Supplementary spreadsheet 3: Untargeted lipidomics analysis of ESI-neagitive mode data obtained via

| Name | Formula | Annot. DeltaV | Calc. MW | m/z | RT [min] | MS Depth | Reference Ion |
| --- | --- | --- | --- | --- | --- | --- | --- |
| WE(47:0) | C47 H94 O2 | -3.69 | 690.72283 | 689.71555 | 19.813 | 1 | [M-H]-1 |
| WE(45:0) | C45 H90 O2 | -1.97 | 662.69278 | 661.6855 | 19.245 | 2 | [M-H]-1 |
| WE(43:0) | C43 H86 O2 | -1.55 | 634.6618 | 633.65452 | 18.836 | 2 | [M-H]-1 |
| WE(26:0) | C26 H52 O2 | -1.92 | 396.39597 | 395.38869 | 12.146 | 2 | [M-H]-1 |
| WE(25:0) | C25 H50 O2 | -3.16 | 382.37987 | 381.37259 | 8.634 | 2 | [M-H]-1 |
| WE(23:0) | C23 H46 O2 | -3.09 | 354.34868 | 353.34141 | 9.742 | 2 | [M-H]-1 |
| ST(d42:2) | C48 H91 N O1 | -3.61 | 889.62807 | 888.6208 | 9.727 | 2 | [M-H]-1 |
| ST(d42:1) | C48 H93 N O1 | -3.66 | 891.64367 | 890.63639 | 10.702 | 2 | [M-H]-1 |
| ST(d41:1) | C47 H91 N O1 | -3.73 | 877.62801 | 876.62073 | 10.209 | 2 | [M-H]-1 |
| ST(d40:1) | C46 H89 N O1 | -3.63 | 863.6125 | 862.60522 | 9.746 | 2 | [M-H]-1 |
| ST(d34:1) | C40 H77 N O1 | -4.77 | 779.51801 | 778.51074 | 6.895 | 2 | [M-H]-1 |
| SQMG(16:0) | C25 H48 O11 | -2.69 | 556.29024 | 555.28296 | 1.865 | 1 | [M-H]-1 |
| SQDG(16:0_18:3) | C43 H76 O12 | -2.54 | 816.50367 | 815.4964 | 6.605 | 2 | [M-H]-1 |
| SQDG(16:0_18:2) | C43 H78 O12 | -1.46 | 818.5202 | 817.51292 | 7.218 | 1 | [M-H]-1 |
| SQDG(16:0_16:0) | C41 H78 O12 | -0.97 | 794.52063 | 793.51335 | 7.795 | 1 | [M-H]-1 |
| SM(d42:2) | C47 H93 N2 C | -0.12 | 812.67703 | 871.69086 | 16.531 | 2 | [M-H+HAc]-1 |
| SM(d41:2) | C46 H91 N2 C | -2.2 | 798.65972 | 797.65041 | 15.978 | 2 | [M-H]-1 |
| PS(18:1e_20:4) | C44 H78 N O9 | -1.77 | 795.54001 | 794.53274 | 8.762 | 2 | [M-H]-1 |
| PS(18:1e_18:2) | C42 H78 N O9 | -1.63 | 771.54016 | 770.53289 | 8.928 | 2 | [M-H]-1 |
| PI(18:1_20:4) | C47 H81 O13 | -2.06 | 884.53965 | 883.53238 | 7.63 | 2 | [M-H]-1 |
| PI(18:1_18:2) | C45 H81 O13 | -2.31 | 860.53949 | 859.53222 | 7.792 | 2 | [M-H]-1 |
| PI(18:0_22:6) | C49 H83 O13 | -0.19 | 910.55696 | 909.54968 | 8.169 | 2 | [M-H]-1 |
| PI(18:0_22:5) | C49 H85 O13 | -1.18 | 912.57171 | 911.56443 | 8.463 | 2 | [M-H]-1 |
| PI(18:0_20:4) | C47 H83 O13 | -2.39 | 886.55501 | 885.54773 | 8.102 | 1 | [M-H]-1 |
| PI(18:0_20:3) | C47 H85 O13 | -2.23 | 888.5708 | 887.56352 | 9.179 | 2 | [M-H]-1 |
| PI(18:0_20:2) | C47 H87 O13 | -3.64 | 890.58518 | 889.57791 | 9.475 | 2 | [M-H]-1 |
| PI(18:0_18:2) | C45 H83 O13 | -2.41 | 862.55505 | 861.54778 | 8.565 | 2 | [M-H]-1 |
| PI(18:0_18:1) | C45 H85 O13 | -1.64 | 864.57136 | 863.56408 | 9.36 | 2 | [M-H]-1 |
| PI(16:0_22:6) | C47 H79 O13 | -1.9 | 882.52415 | 881.51687 | 6.791 | 1 | [M-H]-1 |
| PI(16:0_20:4) | C45 H79 O13 | -1.43 | 858.5246 | 857.51733 | 7.391 | 2 | [M-H]-1 |
| PI(16:0_18:2) | C43 H79 O13 | -1.78 | 834.52435 | 833.51707 | 7.517 | 2 | [M-H]-1 |
| PI(16:0_18:1) | C43 H81 O13 | -1.01 | 836.54063 | 835.53335 | 8.309 | 2 | [M-H]-1 |
| PG(20:0e_16:0) | C42 H85 O9 P | 1.71 | 764.59443 | 763.58715 | 13.549 | 2 | [M-H]-1 |
| PG(18:1_18:2) | C42 H77 O10 | -1.1 | 772.52459 | 771.51731 | 7.859 | 2 | [M-H]-1 |
| PG(18:1_18:1) | C42 H79 O10 | -1.62 | 774.53983 | 773.53256 | 8.375 | 1 | [M-H]-1 |
| PG(18:0_18:1) | C42 H81 O10 | -1.23 | 776.55578 | 775.54851 | 9.648 | 2 | [M-H]-1 |
| PG(18:0_16:0) | C40 H79 O10 | -1.26 | 750.54014 | 749.53284 | 9.648 | 2 | [M-H]-1 |
| PG(16:1_18:2) | C40 H73 O10 | -1.01 | 744.49338 | 743.48611 | 7.139 | 1 | [M-H]-1 |
| PG(16:0_18:2) | C40 H75 O10 | -1.66 | 746.50854 | 745.50127 | 7.819 | 2 | [M-H]-1 |
| PG(16:0_18:1) | C40 H77 O10 | -1.1 | 748.52461 | 747.51733 | 8.602 | 2 | [M-H]-1 |
| PG(16:0_17:0) | C39 H77 O10 | -1.54 | 736.5243 | 735.51702 | 8.832 | 1 | [M-H]-1 |
| PG(16:0_16:1) | C38 H73 O10 | -1.26 | 720.49323 | 719.48595 | 7.599 | 2 | [M-H]-1 |
| PG(16:0_16:0) | C38 H75 O10 | -1.52 | 722.50868 | 721.50141 | 8.477 | 2 | [M-H]-1 |
| PG(16:0_14:0) | C36 H71 O10 | -1.57 | 694.47739 | 693.47011 | 7.452 | 2 | [M-H]-1 |
| PG(15:0_16:0) | C37 H73 O10 | -1.65 | 708.49297 | 707.48569 | 7.961 | 2 | [M-H]-1 |
| PEt(34:2e) | C39 H75 O7 P | 6.22 | 686.52931 | 685.52203 | 4.137 | 1 | [M-H]-1 |
| PE(24:0_14:4) | C43 H78 N O8 | -1.35 | 767.54547 | 766.53825 | 9.155 | 2 | [M-H]-1 |
| PE(22:0_14:4) | C41 H74 N O8 | -0.93 | 739.51452 | 738.50724 | 7.425 | 1 | [M-H]-1 |
| PE(20:1e_22:6) | C47 H82 N O7 | -1.03 | 803.58206 | 802.57479 | 13.334 | 2 | [M-H]-1 |
| PE(20:1e_18:2) | C43 H82 N O7 | -1.66 | 755.58163 | 754.57437 | 14.127 | 2 | [M-H]-1 |
| PE(18:2e_22:6) | C45 H76 N O7 | -1.96 | 773.53442 | 772.52714 | 10.824 | 2 | [M-H]-1 |
| PE(18:2e_22:5) | C45 H78 N O7 | -2.36 | 775.54976 | 774.54248 | 11.857 | 2 | [M-H]-1 |
| PE(18:2e_20:5) | C43 H74 N O7 | -2.01 | 747.51879 | 746.51151 | 9.999 | 2 | [M-H]-1 |
| PE(18:2e_20:4) | C43 H76 N O7 | 0.02 | 749.53595 | 748.52867 | 11.386 | 2 | [M-H]-1 |
| PE(18:2e_18:2) | C41 H76 N O7 | -2.4 | 725.5342 | 724.52692 | 11.495 | 2 | [M-H]-1 |
| PE(18:2_20:4) | C43 H74 N O8 | -0.74 | 763.51464 | 762.50737 | 9.354 | 2 | [M-H]-1 |
| PE(18:2_18:2) | C41 H74 N O8 | -1.17 | 739.51434 | 738.50707 | 9.612 | 2 | [M-H]-1 |
| PE(18:1e_22:5) | C45 H80 N O7 | -2.31 | 777.56544 | 776.55817 | 12 | 1 | [M-H]-1 |
| PE(18:1e_20:4) | C43 H78 N O7 | -1.51 | 751.55046 | 750.54318 | 12.293 | 2 | [M-H]-1 |
| PE(18:1e_20:3) | C43 H80 N O7 | -1.49 | 753.56612 | 752.55884 | 12.999 | 2 | [M-H]-1 |
| PE(18:1e_18:2) | C41 H78 N O7 | -2.61 | 727.54969 | 726.54242 | 12.4 | 2 | [M-H]-1 |
| PE(18:1e_18:1) | C41 H80 N O7 | -1.38 | 729.56623 | 728.55897 | 13.745 | 2 | [M-H]-1 |
| PE(18:1_22:6) | C45 H76 N O8 | -0.18 | 789.53072 | 788.52344 | 9.987 | 2 | [M-H]-1 |
| PE(18:1_20:4) | C43 H76 N O8 | -1.86 | 765.52943 | 746.5116 | 10.515 | 2 | [M-H-H2O]-1 |
| PE(18:1_18:2) | C41 H76 N O8 | -1.13 | 741.53002 | 740.52274 | 10.587 | 2 | [M-H]-1 |
| PE(18:1_18:1) | C41 H78 N O8 | -1.47 | 743.54541 | 742.53847 | 11.303 | 2 | [M-H]-1 |
| PE(18:0_22:6) | C45 H78 N O8 | 0.43 | 791.54685 | 772.52899 | 10.797 | 2 | [M-H-H2O]-1 |
| PE(18:0_22:5) | C45 H80 N O8 | -0.02 | 793.56214 | 792.55207 | 11.867 | 2 | [M-H]-1 |
| PE(18:0_20:4) | C43 H78 N O8 | -1.57 | 767.5453 | 766.53802 | 12.061 | 1 | [M-H]-1 |
| PE(18:0_20:3) | C43 H80 N O8 | -2.53 | 769.56021 | 768.55296 | 11.878 | 2 | [M-H]-1 |
| PE(18:0_18:2) | C41 H78 N O8 | -1.23 | 743.54559 | 742.53831 | 11.817 | 2 | [M-H]-1 |
| PE(18:0_18:1) | C41 H80 N O8 | -0.62 | 745.56169 | 744.55206 | 12.571 | 2 | [M-H]-1 |
| PE(17:1_20:4) | C42 H74 N O8 | -1.13 | 751.51436 | 750.50699 | 8.146 | 2 | [M-H]-1 |
| PE(17:1_16:0) | C38 H74 N O8 | -0.83 | 703.51462 | 702.50734 | 10.957 | 1 | [M-H]-1 |
| PE(17:0_20:4) | C42 H76 N O8 | -1.14 | 753.52999 | 752.52272 | 10.865 | 2 | [M-H]-1 |
| PE(17:0_18:2) | C40 H76 N O8 | -0.27 | 729.53066 | 728.52338 | 11.112 | 2 | [M-H]-1 |
| PE(16:1e_22:6) | C43 H74 N O7 | -0.51 | 747.51991 | 746.51263 | 10.148 | 2 | [M-H]-1 |
| PE(16:1e_22:4) | C43 H78 N O7 | -2.38 | 751.5498 | 750.54253 | 11.258 | 2 | [M-H]-1 |
| PE(16:1e_20:5) | C41 H72 N O7 | -1.06 | 721.50388 | 720.4966 | 9.993 | 2 | [M-H]-1 |
| PE(16:1e_20:4) | C41 H74 N O7 | -1.1 | 723.51949 | 722.51221 | 10.873 | 2 | [M-H]-1 |
| PE(16:1e_18:2) | C39 H74 N O7 | -0.99 | 699.5196 | 698.51232 | 11.192 | 2 | [M-H]-1 |
| PE(16:1e_18:1) | C39 H76 N O7 | -1.49 | 701.53489 | 700.52762 | 12.277 | 2 | [M-H]-1 |
| PE(16:0e_18:2) | C39 H76 N O7 | -1.29 | 701.53503 | 700.52776 | 11.44 | 2 | [M-H]-1 |
| PE(16:0e_18:1) | C39 H78 N O7 | -0.92 | 703.55094 | 702.54366 | 12.537 | 2 | [M-H]-1 |
| PE(16:0_22:6) | C43 H74 N O8 | -1.83 | 763.51381 | 762.50653 | 9.897 | 2 | [M-H]-1 |
| PE(16:0_22:4) | C43 H78 N O8 | -2 | 767.54497 | 766.53832 | 10.979 | 2 | [M-H]-1 |
| PE(16:0_20:5) | C41 H72 N O8 | -0.74 | 737.49901 | 736.49173 | 9.378 | 2 | [M-H]-1 |
| PE(16:0_20:4) | C41 H74 N O8 | -1.68 | 739.51396 | 738.50682 | 9.898 | 2 | [M-H]-1 |
| PE(16:0_18:2) | C39 H74 N O8 | -0.81 | 715.51463 | 714.50735 | 10.468 | 2 | [M-H]-1 |
| PE(16:0_18:1) | C39 H76 N O8 | -1.31 | 717.52991 | 716.52263 | 11.484 | 2 | [M-H]-1 |
| PC(44:6e) | C52 H94 N O7 | -4.91 | 875.67249 | 874.66521 | 15.987 | 1 | [M-H]-1 |
| PC(44:5e) | C52 H96 N O7 | -4.5 | 877.68849 | 876.68121 | 18.302 | 1 | [M-H]-1 |
| PC(36:6e) | C44 H78 N O7 | -3.37 | 763.54901 | 808.54755 | 9.406 | 2 | [M+FA-H]-1 |
| PC(30:1_20:4) | C58 H106 N O | -1.06 | 975.76457 | 974.75729 | 14.702 | 1 | [M-H]-1 |
| PC(20:5_18:2) | C46 H78 N O8 | -1.02 | 803.54569 | 802.53816 | 8.971 | 2 | [M-H]-1 |
| PC(18:1_20:4) | C46 H82 N O8 | 0.11 | 807.57789 | 806.57062 | 9.499 | 1 | [M-H]-1 |
| PC(17:1_20:4) | C45 H80 N O8 | -0.91 | 793.56143 | 792.55415 | 9.313 | 2 | [M-H]-1 |
| PC(17:0_20:3) | C45 H84 N O8 | -2.8 | 797.59122 | 856.6084 | 12.311 | 1 | [M-H+HAc]-1 |
| PC(17:0_18:2) | C43 H82 N O8 | -2 | 771.57626 | 816.57449 | 11.347 | 2 | [M+FA-H]-1 |
| PC(16:0_20:4) | C44 H80 N O8 | -0.5 | 781.56177 | 780.55449 | 11.834 | 1 | [M-H]-1 |
| PC(16:0_18:2) | C42 H80 N O8 | -0.42 | 757.56183 | 756.5543 | 12.251 | 1 | [M-H]-1 |
| PC(16:0_18:1) | C42 H82 N O8 | -0.85 | 759.57716 | 758.56989 | 13.623 | 1 | [M-H]-1 |
| PC(15:0_22:6) | C45 H78 N O8 | -0.53 | 791.54609 | 790.53873 | 9.34 | 2 | [M-H]-1 |
| PC(15:0_20:4) | C43 H78 N O8 | -3.5 | 767.54382 | 766.53224 | 10.243 | 2 | [M-H]-1 |
| Palmitic acid_putative) |  |  | 256.16702 | 255.15975 | 2.334 | 2 | [M-H]-1 |
| PA(36:1e) | C39 H77 O7 P | -3.61 | 688.53821 | 687.53093 | 4.28 | 1 | [M-H]-1 |
| PA(36:0e) | C39 H79 O7 P | 9.33 | 690.56279 | 689.55551 | 6.258 | 2 | [M-H]-1 |
| PA(35:0) | C38 H75 O8 P | 1.7 | 690.52113 | 689.51385 | 13.315 | 1 | [M-H]-1 |
| OAHFA(44:9) | C44 H68 O4 | 7.11 | 660.51646 | 659.50918 | 3.577 | 2 | [M-H]-1 |
| OAHFA(42:10) | C42 H62 O4 | -6.16 | 630.46093 | 629.45365 | 3.654 | 2 | [M-H]-1 |
| OAHFA(40:6) | C40 H66 O4 | -0.95 | 610.49553 | 609.48825 | 3.88 | 2 | [M-H]-1 |
| OAHFA(38:6) | C38 H62 O4 | -6.82 | 582.46084 | 581.45356 | 3.874 | 2 | [M-H]-1 |
| OAHFA(38:4) | C38 H66 O4 | -5.97 | 586.49261 | 585.48533 | 4.754 | 2 | [M-H]-1 |
| OAHFA(36:4) | C36 H62 O4 | -1.59 | 558.46392 | 557.45665 | 3.643 | 2 | [M-H]-1 |
| OAHFA(36:3) | C36 H64 O4 | -0.72 | 560.48005 | 559.47278 | 4.1 | 1 | [M-H]-1 |
| OAHFA(36:1) | C36 H68 O4 | -2.93 | 564.51011 | 563.50283 | 4.962 | 2 | [M-H]-1 |
| MGDG(42:4) | C51 H90 O10 | -1.29 | 862.65228 | 861.64501 | 13.969 | 2 | [M-H]-1 |
| MGDG(18:1_18:2) | C45 H80 O10 | -1.62 | 780.57388 | 839.58768 | 10.866 | 2 | [M-H+HAc]-1 |
| MGDG(18:1_18:1) | C45 H82 O10 | 0.09 | 782.59087 | 841.60468 | 11.884 | 2 | [M-H+HAc]-1 |
| MGDG(18:1_15:0) | C42 H78 O10 | 3.69 | 742.56224 | 741.55496 | 10.919 | 1 | [M-H]-1 |
| MGDG(18:0_18:2) | C45 H82 O10 | 6.31 | 782.59574 | 781.58846 | 13.328 | 1 | [M-H]-1 |
| MGDG(16:0_18:2) | C43 H78 O10 | -1.29 | 754.55853 | 753.55104 | 10.707 | 2 | [M-H]-1 |
| MGDG(16:0_18:1) | C43 H80 O10 | -1.67 | 756.57389 | 815.5877 | 11.726 | 2 | [M-H+HAc]-1 |
| LPI(18:0) | C27 H53 O12 | -3.16 | 600.32557 | 599.31829 | 2.881 | 2 | [M-H]-1 |
| LPE(22:6) | C27 H44 N O7 | -1.55 | 525.28472 | 524.27745 | 2.67 | 2 | [M-H]-1 |
| LPE(22:5) | C27 H46 N O7 | -1.06 | 527.30063 | 526.29335 | 3.16 | 2 | [M-H]-1 |
| LPE(22:4) | C27 H48 N O7 | -1.12 | 529.31625 | 528.30897 | 3.464 | 2 | [M-H]-1 |
| LPE(20:5) | C25 H42 N O7 | -1.71 | 499.26903 | 498.26176 | 2.351 | 2 | [M-H]-1 |
| LPE(20:4) | C25 H44 N O7 | -2.35 | 501.28436 | 500.27708 | 3.151 | 1 | [M-H]-1 |
| LPE(20:3) | C25 H46 N O7 | 0.13 | 503.30125 | 502.29398 | 3.159 | 2 | [M-H]-1 |
| LPE(18:2e) | C23 H46 N O6 | -0.96 | 463.30583 | 462.29855 | 3.909 | 2 | [M-H]-1 |
| LPE(18:2) | C23 H44 N O7 | -1.33 | 477.2849 | 476.27763 | 2.942 | 2 | [M-H]-1 |
| LPE(18:1e) | C23 H48 N O6 | -1.37 | 465.32129 | 464.31401 | 4.956 | 2 | [M-H]-1 |
| LPE(18:1) | C23 H46 N O7 | -1.25 | 479.30059 | 478.29331 | 3.544 | 2 | [M-H]-1 |
| LPE(18:0) | C23 H48 N O7 | -1.84 | 481.31595 | 480.30868 | 4.472 | 2 | [M-H]-1 |
| LPE(16:1e) | C21 H44 N O6 | -1.15 | 437.29012 | 436.28284 | 3.768 | 2 | [M-H]-1 |
| LPE(16:0) | C21 H44 N O7 | -1.01 | 453.28508 | 452.27781 | 3.41 | 2 | [M-H]-1 |
| LPC(16:1) | C24 H48 N O7 | 0.95 | 493.31731 | 492.31 | 3.41 | 1 | [M-H]-1 |
| LPC(16:0) | C24 H50 N O7 | -0.44 | 495.33227 | 494.32504 | 4.605 | 1 | [M-H]-1 |
| LPC(15:0) | C23 H48 N O7 | 2.34 | 481.31797 | 480.31049 | 2.819 | 1 | [M-H]-1 |
| LdMePE(18:2) | C25 H48 N O7 | -1.16 | 505.31625 | 504.30863 | 3.137 | 2 | [M-H]-1 |
| LdMePE(18:1) | C25 H50 N O7 | -2.32 | 507.33131 | 506.32404 | 3.463 | 1 | [M-H]-1 |
| LdMePE(18:0) | C25 H52 N O7 | -2.72 | 509.34675 | 508.33948 | 4.613 | 1 | [M-H]-1 |
| LdMePE(16:0) | C23 H48 N O7 | -0.96 | 481.31638 | 480.30885 | 3.653 | 2 | [M-H]-1 |
| Hex1Cer(d18:1_24:1) | C48 H91 N O8 | -2.84 | 809.67217 | 868.68616 | 13.866 | 2 | [M-H+HAc]-1 |
| FA(24:5) | C24 H38 O2 | -1.65 | 358.28659 | 357.27931 | 4.92 | 2 | [M-H]-1 |
| FA(24:4) | C24 H40 O2 | -1.8 | 360.30218 | 359.29498 | 5.841 | 2 | [M-H]-1 |
| FA(22:6) | C22 H32 O2 | -2.6 | 328.23938 | 327.2321 | 3.57 | 2 | [M-H]-1 |
| FA(22:5) | C22 H34 O2 | -1.94 | 330.25524 | 329.24796 | 3.921 | 2 | [M-H]-1 |
| FA(22:4) | C22 H36 O2 | -4.21 | 332.27013 | 331.26325 | 5.376 | 1 | [M-H]-1 |
| FA(20:5) | C20 H30 O2 | -2.12 | 302.22394 | 301.21663 | 1.875 | 2 | [M-H]-1 |
| FA(20:4) | C20 H32 O2 | -2.45 | 304.23948 | 303.23221 | 4.546 | 2 | [M-H]-1 |
| FA(18:4) | C18 H28 O2 | -1.41 | 276.20854 | 275.20126 | 2.744 | 2 | [M-H]-1 |
| FA(18:2) | C18 H32 O2 | -2.28 | 280.23959 | 279.23231 | 3.873 | 2 | [M-H]-1 |
| FA(16:1) | C16 H30 O2 | -6.41 | 254.22295 | 253.21567 | 4.148 | 2 | [M-H]-1 |
| FA(14:1) | C14 H26 O2 | -3 | 226.1926 | 225.18532 | 3.085 | 2 | [M-H]-1 |
| dMePE(18:2_20:4) | C45 H78 N O8 | -2.37 | 791.54463 | 790.53735 | 8.378 | 2 | [M-H]-1 |
| dMePE(18:0_18:2) | C43 H82 N O8 | -2.2 | 771.57611 | 770.56668 | 13.074 | 2 | [M-H]-1 |
| dMePE(17:1_20:4) | C44 H78 N O8 | -0.77 | 779.54591 | 778.53931 | 9.333 | 2 | [M-H]-1 |
| dMePE(17:1_18:2) | C42 H78 N O8 | -1.69 | 755.54523 | 754.53795 | 9.424 | 2 | [M-H]-1 |
| dMePE(16:1_18:2) | C41 H76 N O8 | -1.86 | 741.52947 | 740.52233 | 8.887 | 1 | [M-H]-1 |
| dMePE(16:0e_18:2) | C41 H80 N O7 | -1.57 | 729.5661 | 728.55882 | 12.891 | 2 | [M-H]-1 |
| Cer(t18:0_23:0) | C41 H83 N O4 | -1.4 | 653.63129 | 712.64525 | 15.789 | 2 | [M-H+HAc]-1 |
| Cer(d19:1_24:1) | C43 H83 N O3 | -2.05 | 661.63594 | 720.64978 | 16.223 | 2 | [M-H+HAc]-1 |
| Cer(d19:1_24:0) | C43 H85 N O3 | -1.73 | 663.6518 | 722.66565 | 17.277 | 2 | [M-H+HAc]-1 |
| Cer(d18:2_23:0) | C41 H79 N O3 | -0.15 | 633.6059 | 668.57472 | 15.348 | 2 | [M+Cl]-1 |
| Cer(d18:2_22:0) | C40 H77 N O3 | -2.16 | 619.58901 | 618.58037 | 14.602 | 1 | [M-H]-1 |
| Cer(d18:1_24:2) | C42 H79 N O3 | -1.68 | 645.60491 | 704.61872 | 14.569 | 2 | [M-H+HAc]-1 |
| Cer(d18:1_24:1) | C42 H81 N O3 | -1.6 | 647.62061 | 706.63445 | 15.787 | 2 | [M-H+HAc]-1 |
| Cer(d18:1_24:0) | C42 H83 N O3 | -2.13 | 649.63591 | 708.64976 | 16.983 | 2 | [M-H+HAc]-1 |
| Cer(d18:1_23:0) | C41 H81 N O3 | -1.93 | 635.62042 | 694.63426 | 16.481 | 2 | [M-H+HAc]-1 |
| Cer(d18:1_22:0) | C40 H79 N O3 | -1.63 | 621.60498 | 680.61883 | 15.62 | 2 | [M-H+HAc]-1 |
| Cer(d18:1_19:0) | C37 H73 N O3 | 0.07 | 579.55908 | 624.55737 | 13.047 | 1 | [M+FA-H]-1 |
| Cer(d18:1_18:0) | C36 H71 N O3 | -2.72 | 565.54185 | 624.55565 | 12.756 | 2 | [M-H+HAc]-1 |
| Cer(d18:1_16:0) | C34 H67 N O3 | -1.67 | 537.5112 | 596.52505 | 11.32 | 3 | [M-H+HAc]-1 |
| Cer(d18:0_24:1) | C42 H83 N O3 | -1.57 | 649.63628 | 694.63446 | 16.308 | 2 | [M+FA-H]-1 |
| Cer(d18:0_18:0) | C36 H73 N O3 | -2.19 | 567.5578 | 626.57166 | 13.329 | 2 | [M-H+HAc]-1 |
| Cer(d18:0_16:0) | C34 H69 N O3 | -1.82 | 539.52676 | 598.54047 | 11.827 | 2 | [M-H+HAc]-1 |
| Cer(d17:1_22:0) | C39 H77 N O3 | -0.97 | 607.58975 | 666.60358 | 15.096 | 2 | [M-H+HAc]-1 |
| Cer(d16:1_22:0) | C38 H75 N O3 | -1.22 | 593.57397 | 628.54278 | 14.343 | 2 | [M+Cl]-1 |
| Cer(d16:1_16:0) | C32 H63 N O3 | -1.39 | 509.48009 | 568.49393 | 9.982 | 2 | [M-H+HAc]-1 |

**Orbitrap-IDX**

| Log2 Fold Change: (Bb_neg_newanalysis_12022024) / (CM_neg_newanalysis_12022024) |  |
| --- | --- |
|  | -0.64 |
|  | -0.75 |
|  | -0.03 |
|  | -0.25 |
|  | -0.51 |
|  | -0.11 |
|  | -5.44 |
|  | -3.15 |
|  | -4.89 |
|  | -4.82 |
|  | -4.57 |
|  | -0.11 |
|  | 1.66 |
|  | 0.23 |
|  | 0.87 |
|  | -3.94 |
|  | 0.36 |
|  | -1.57 |
|  | 5.54 |
|  | -0.76 |
|  | -0.63 |
|  | 0.7 |
|  | -0.52 |
|  | 0.85 |
|  | -1.88 |
|  | -3.69 |
|  | -5.52 |
|  | -3.41 |
|  | 0.34 |
|  | -1.46 |
|  | -3.38 |
|  | -2.94 |
|  | -2.74 |

|  |
| --- |
| 6.04 |
| 4.19 |
| 7.2 |
| 6.08 |
| 4.42 |
| 8.44 |
| 7.68 |
| 2.6 |
| 6.78 |
| 7.8 |
| 5.43 |
| 4.77 |
| -1.11 |
| -0.51 |
| 0.5 |
| 0.42 |
| -2.83 |
| 1.81 |
| 0.25 |
| -0.7 |
| -0.09 |
| -1.62 |
| 1.27 |
| -2.39 |
| -0.79 |
| -4.13 |
| -2 |
| -4.95 |
| -2.05 |
| 2.31 |
| 0.09 |
| -3.47 |
| 1.07 |
| 1.22 |
| 3.94 |

|  |  |
| --- | --- |
|  | 3.63 |
|  | -0.28 |
|  | -1.82 |
|  | 1.24 |
|  | 0.39 |
|  | -0.15 |
|  | 0.69 |
|  | -1.11 |
|  | 1.52 |
|  | -0.04 |
|  | 0.58 |
|  | -1.68 |
|  | -3.54 |
|  | -1.29 |
|  | -0.31 |
|  | 0.39 |
|  | 0.23 |
|  | -1.22 |
|  | 0.95 |
|  | 5.25 |
|  | -4.1 |
|  | -1.66 |
|  | 0.56 |
|  | 0.22 |
|  | 0.64 |
|  | 0.87 |
|  | 1.14 |
|  | 0.56 |
|  | -0.86 |
|  | -1.01 |
|  | 0.92 |
|  | -0.19 |
|  | 0.14 |
|  | 0.12 |
|  | 0.2 |

|  |
| --- |
| 0.08 |
| 1.03 |
| -3.56 |
| 0 |
| 0.52 |
| -5.13 |
| -0.77 |
| -3.34 |
| -6.04 |
| -4.76 |
| -0.71 |
| 0.38 |
| 0.18 |
| 5.79 |
| 11.13 |
| 8.15 |
| 1 |
| 0.59 |
| 4.54 |
| 12.22 |
| 0.98 |
| -0.29 |
| -0.96 |
| -0.56 |
| 0.26 |
| -0.76 |
| -0.73 |
| -2.57 |
| -1.91 |
| -3.63 |
| -1.68 |
| -3.26 |
| -2.21 |
| -3.8 |
| -0.39 |
| -1.06 |
| 0.39 |
| 1.28 |
| -1 |
| -1.71 |
| -2.63 |
| 0.64 |
| -0.39 |
| -1.43 |
| -1.93 |
| -8.56 |
| 0.65 |
| -1.79 |
| -0.37 |
| -2.33 |
| -4.86 |
| -1.47 |
| -3.89 |
| 0.03 |
| -0.86 |
| 0.08 |
| -1.06 |
| -1.14 |
| -2.3 |
| -2.46 |
| 0.05 |
| -3.72 |
| -0.15 |
| 0.71 |
| -3.42 |
| -5.61 |
| -5.86 |
| -5.65 |
| -1.74 |
| -3.74 |

|  |  |
| --- | --- |
|  | -3.03 |
|  | -2.96 |
|  | -1.65 |
|  | -1.09 |
|  | -0.57 |
|  | -2.03 |
|  | -2.31 |
|  | 0.66 |

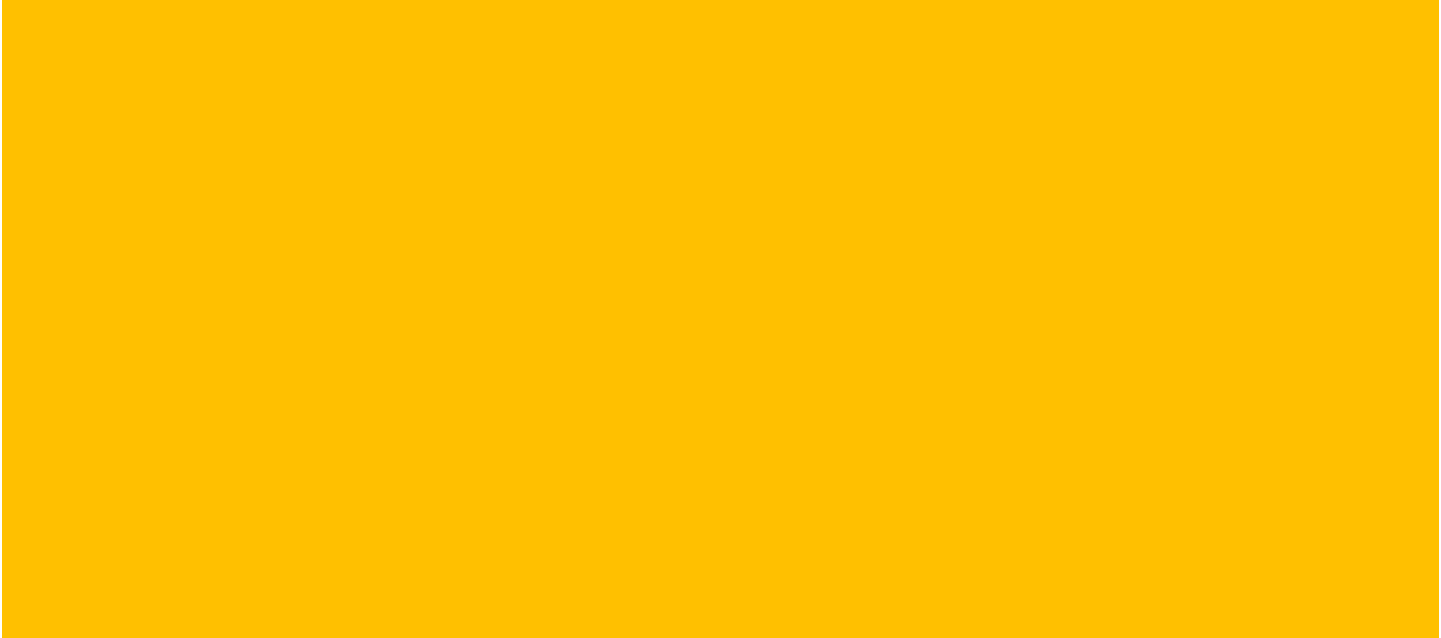

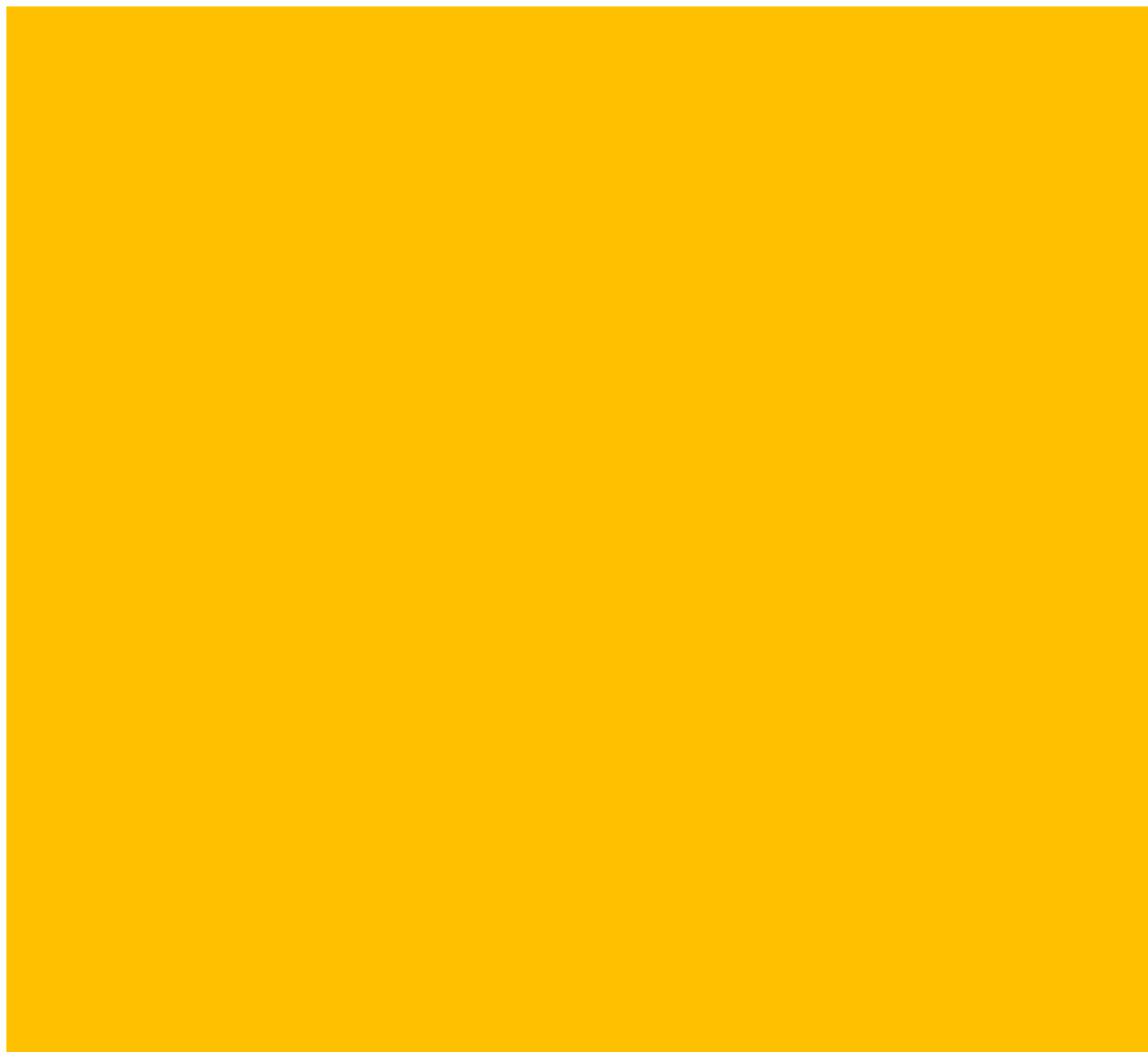

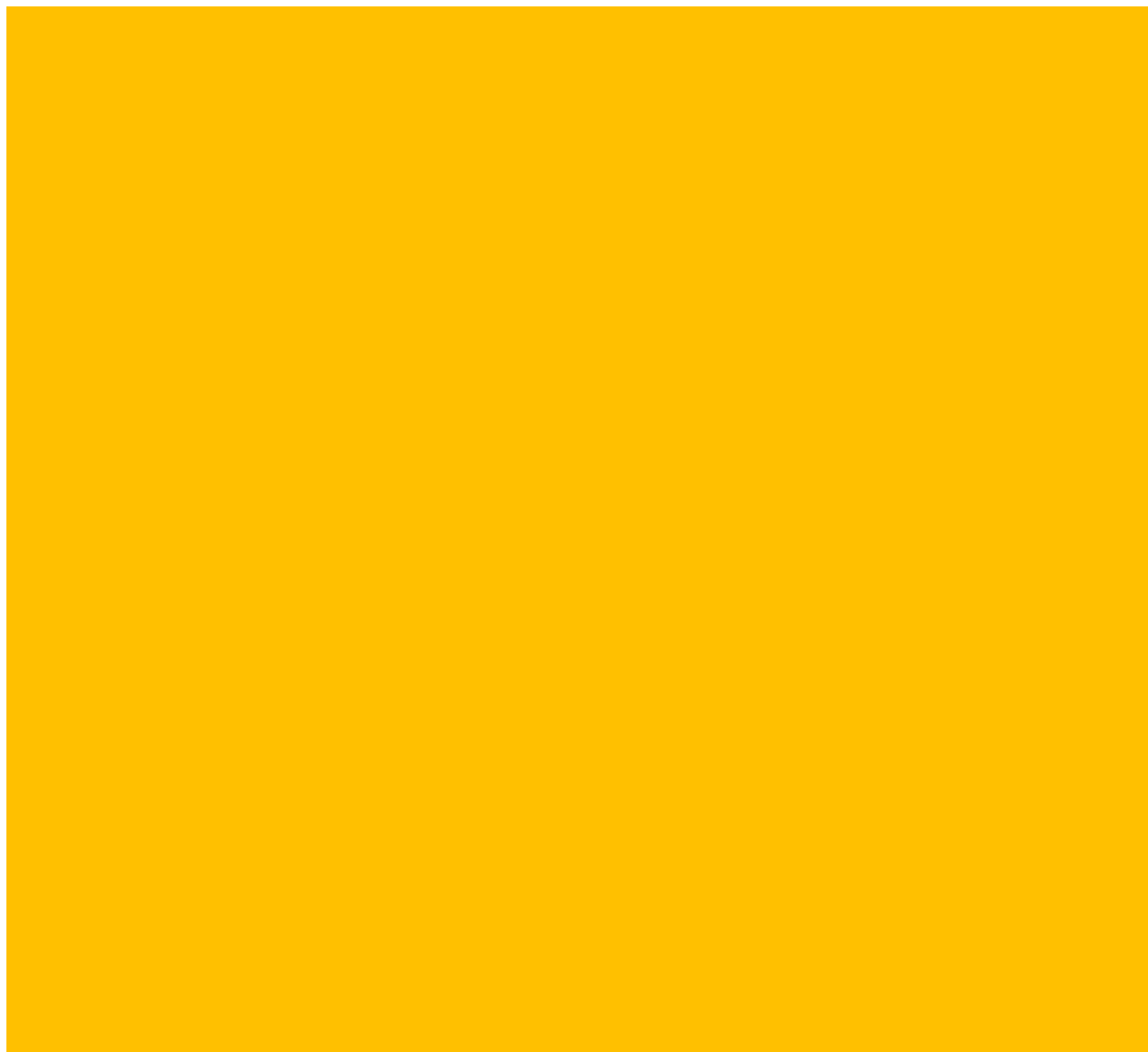

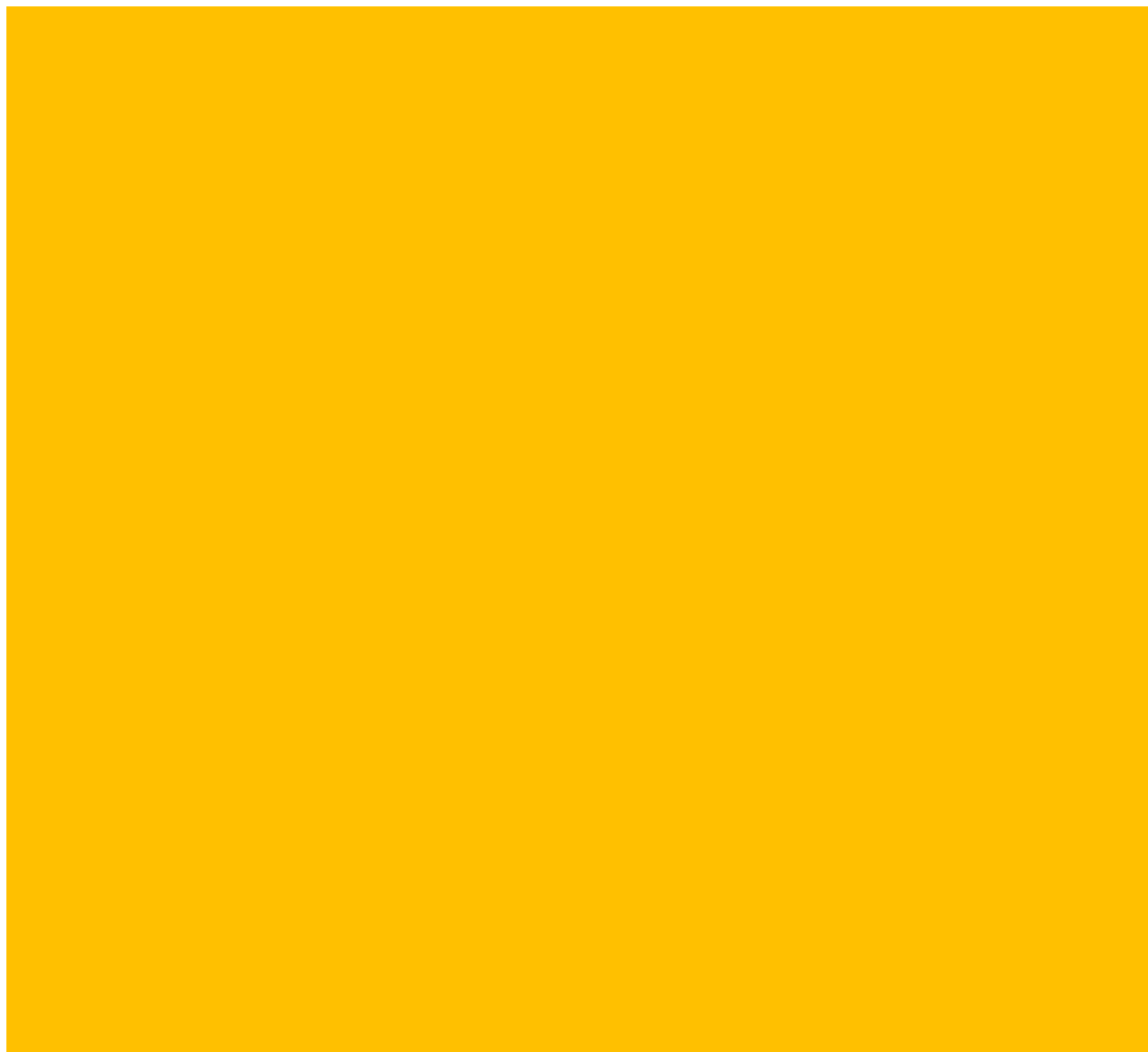

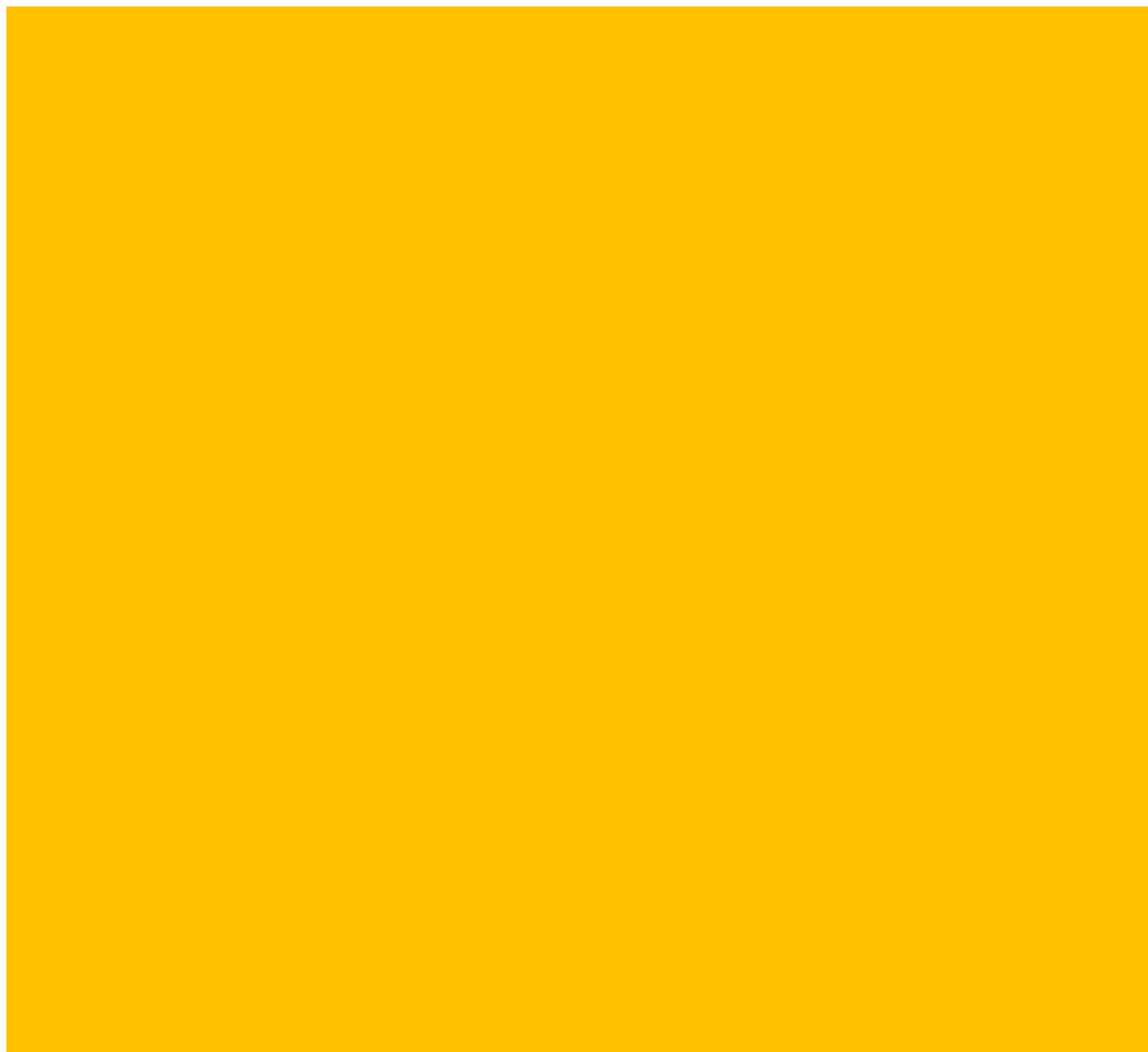

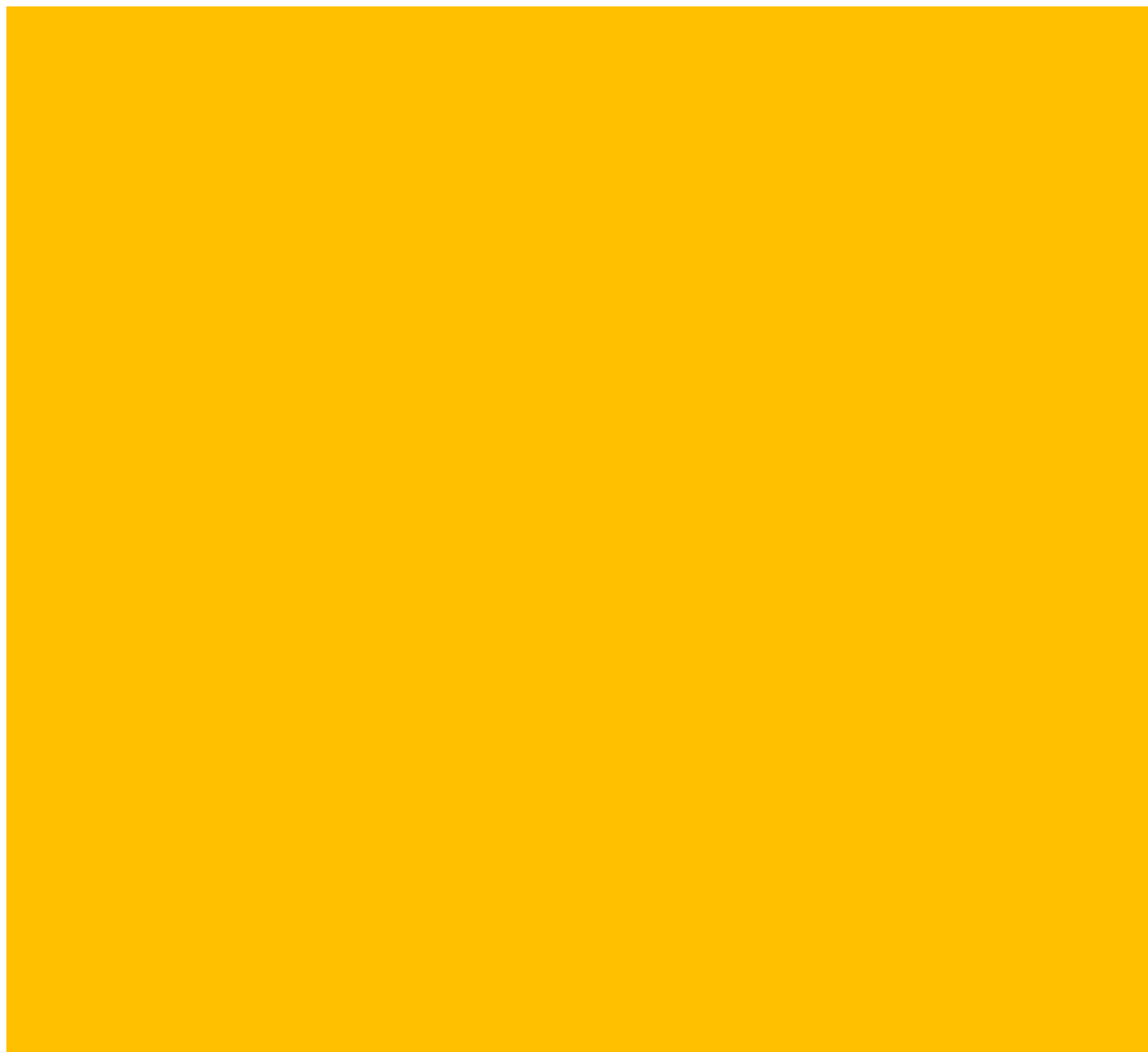

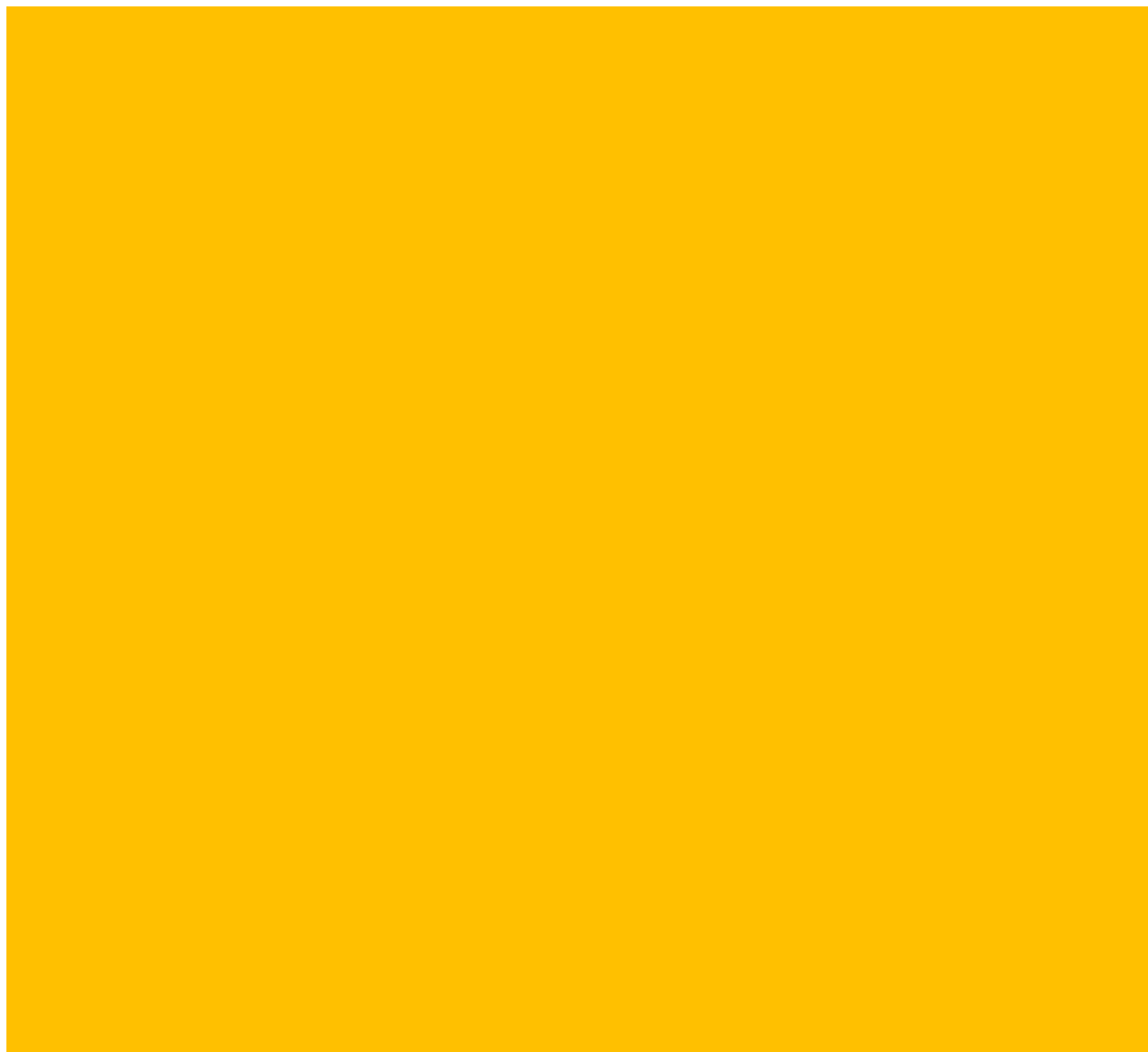

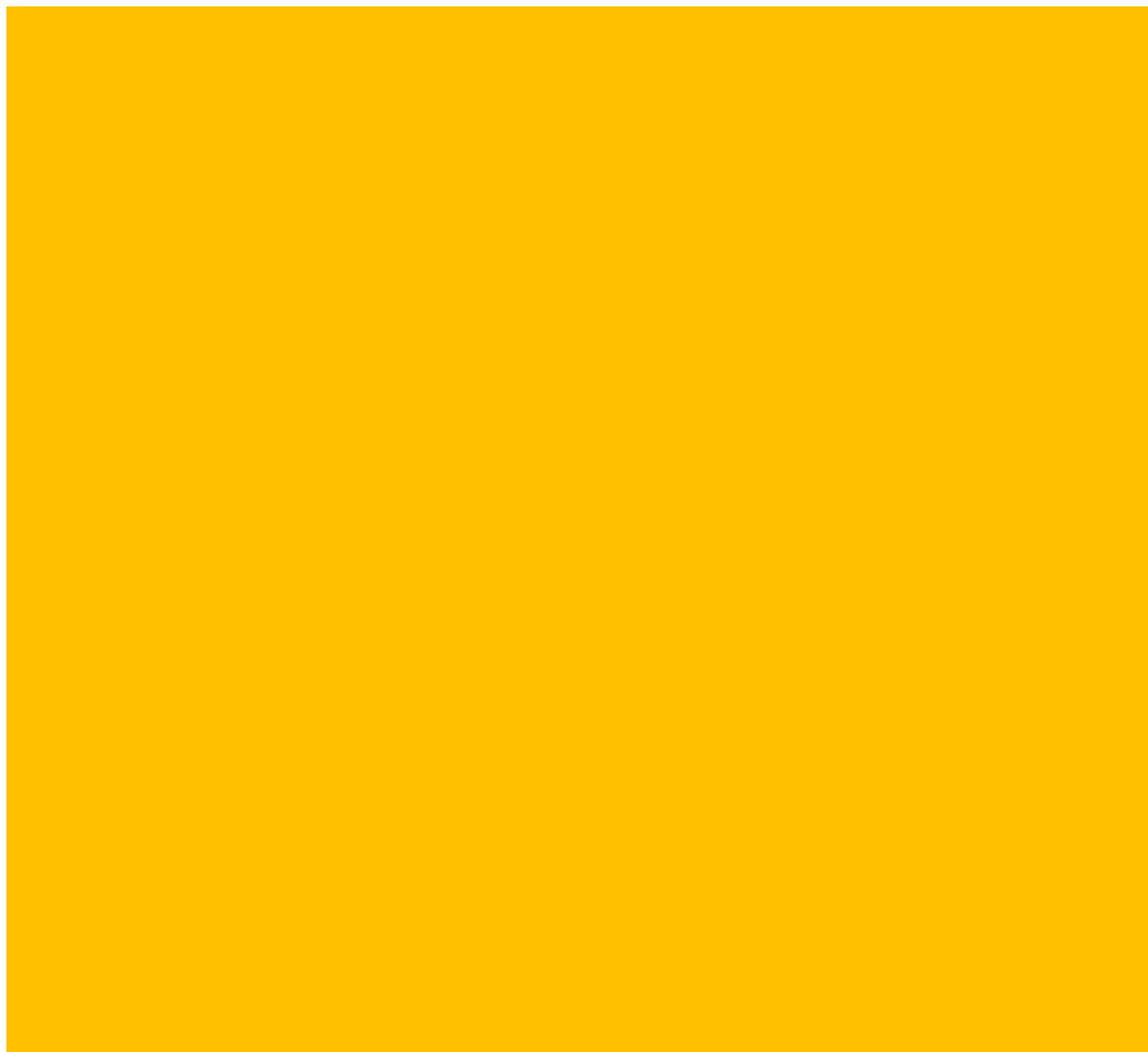

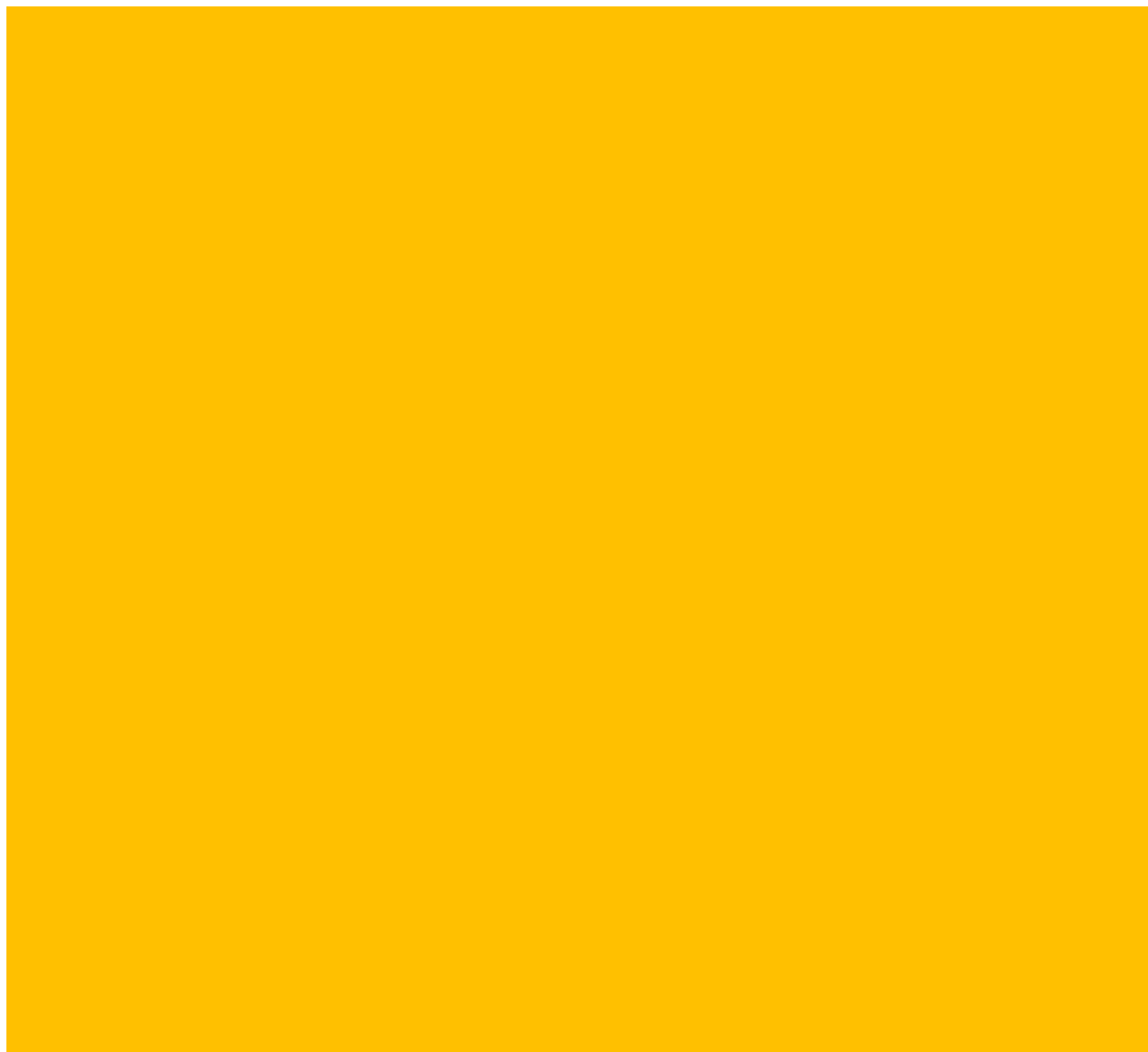

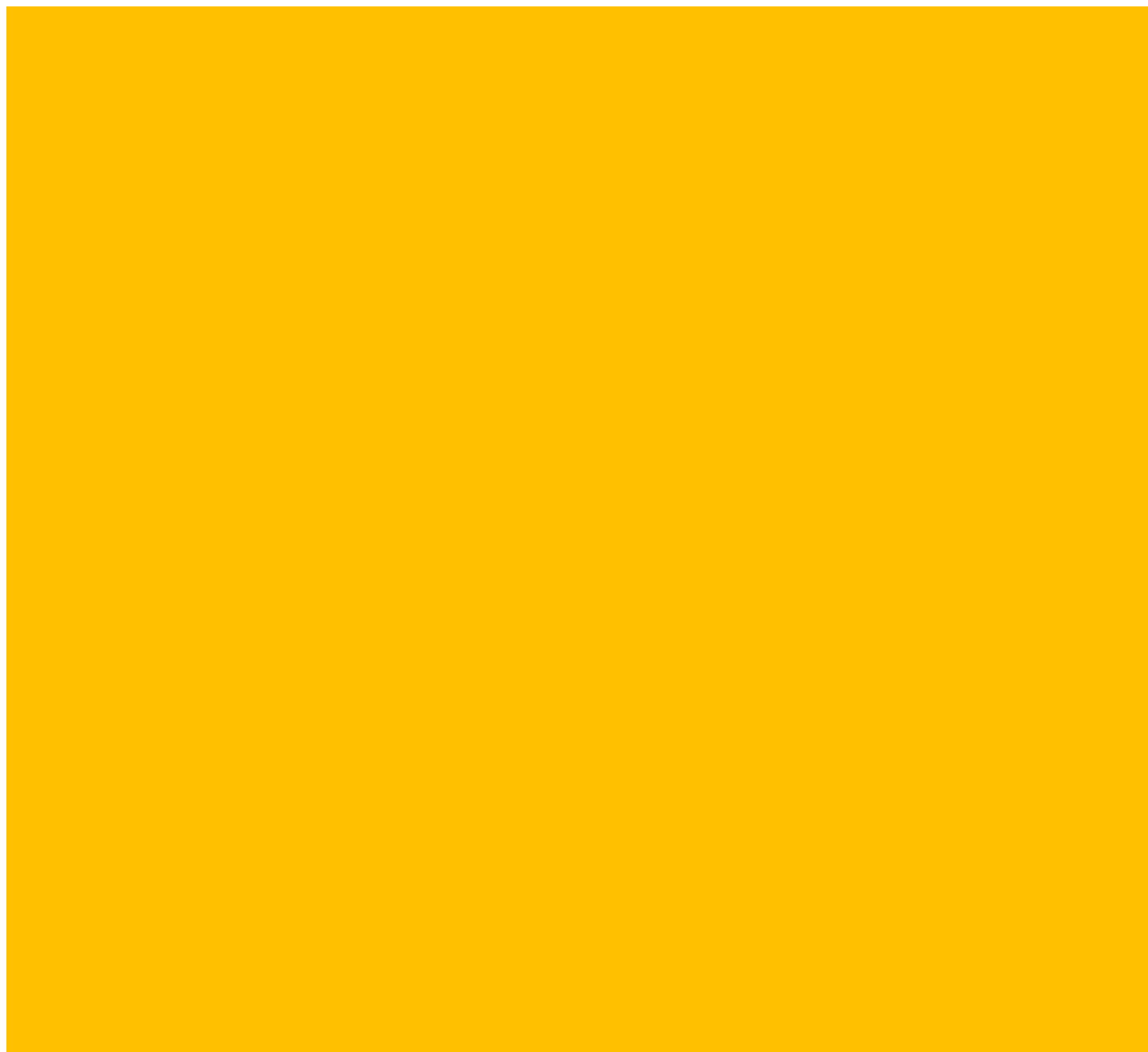

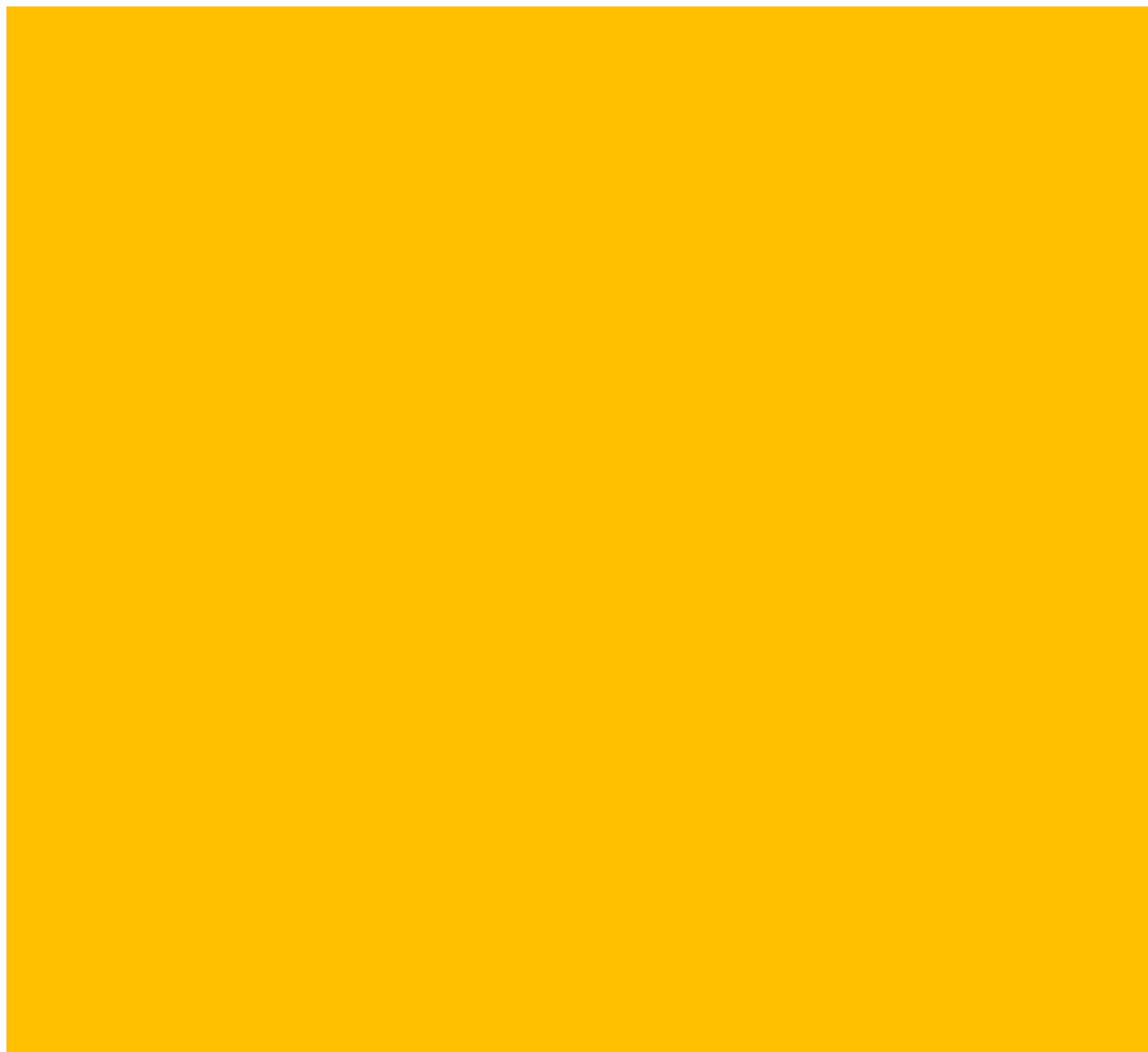

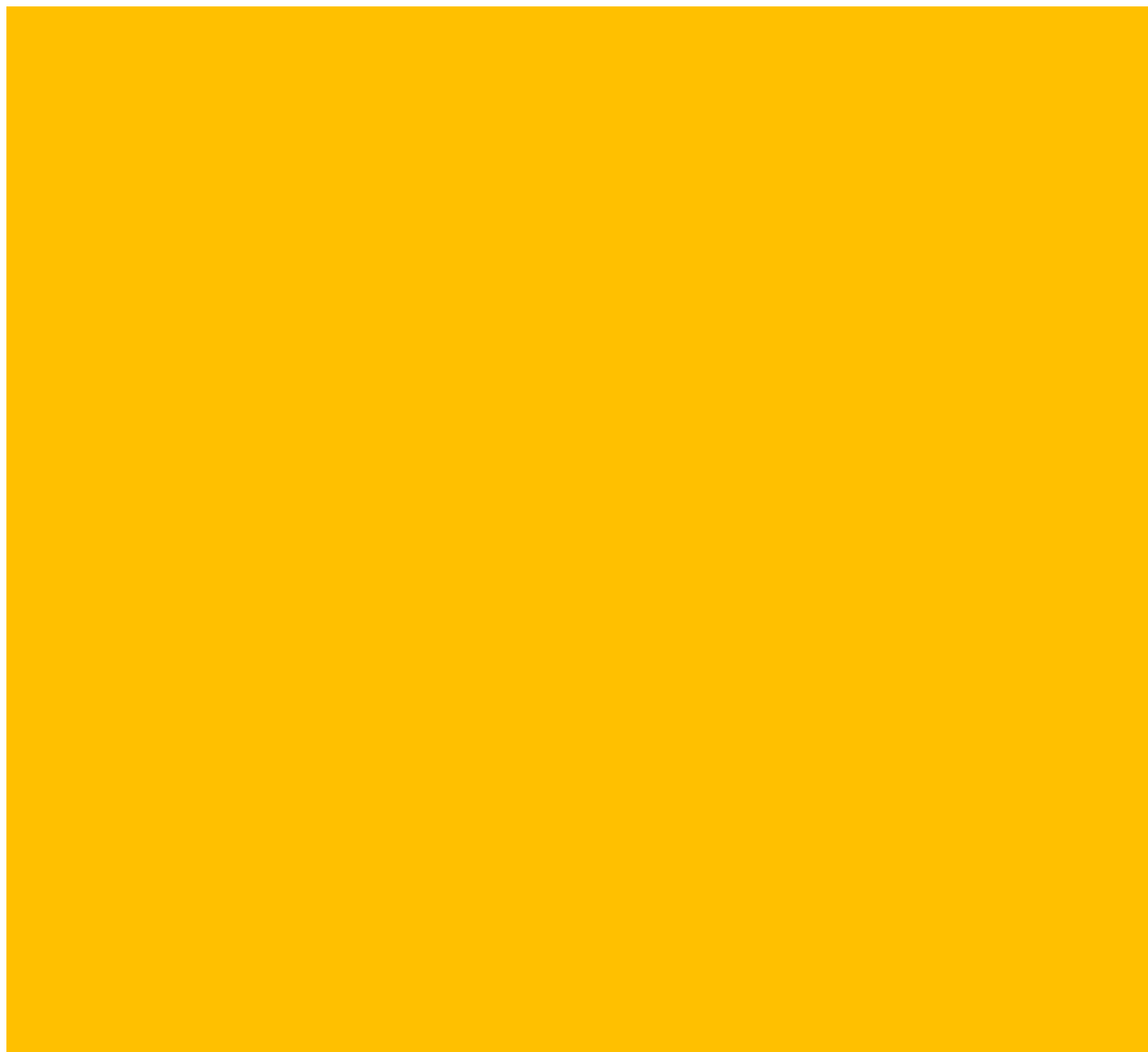

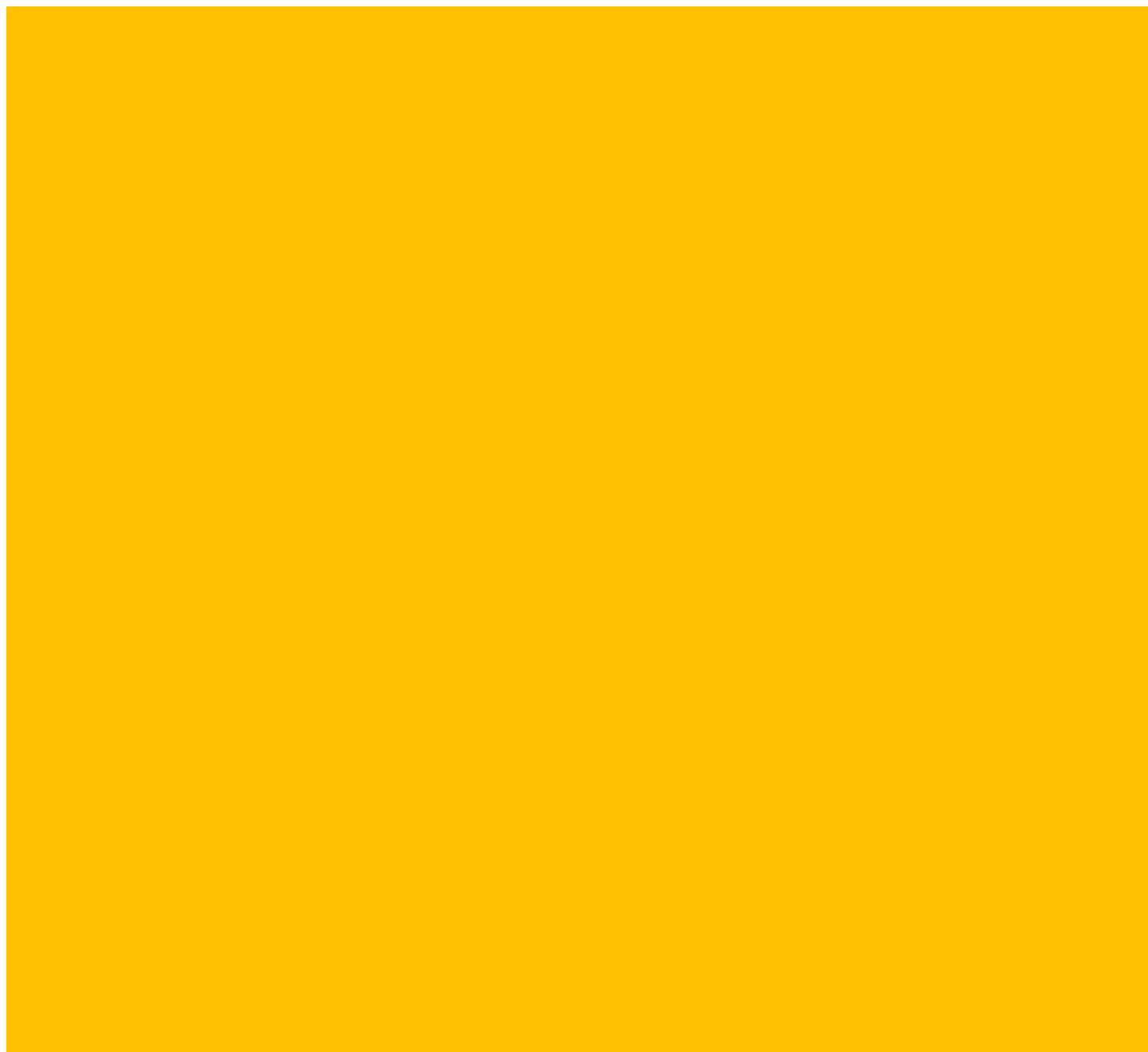

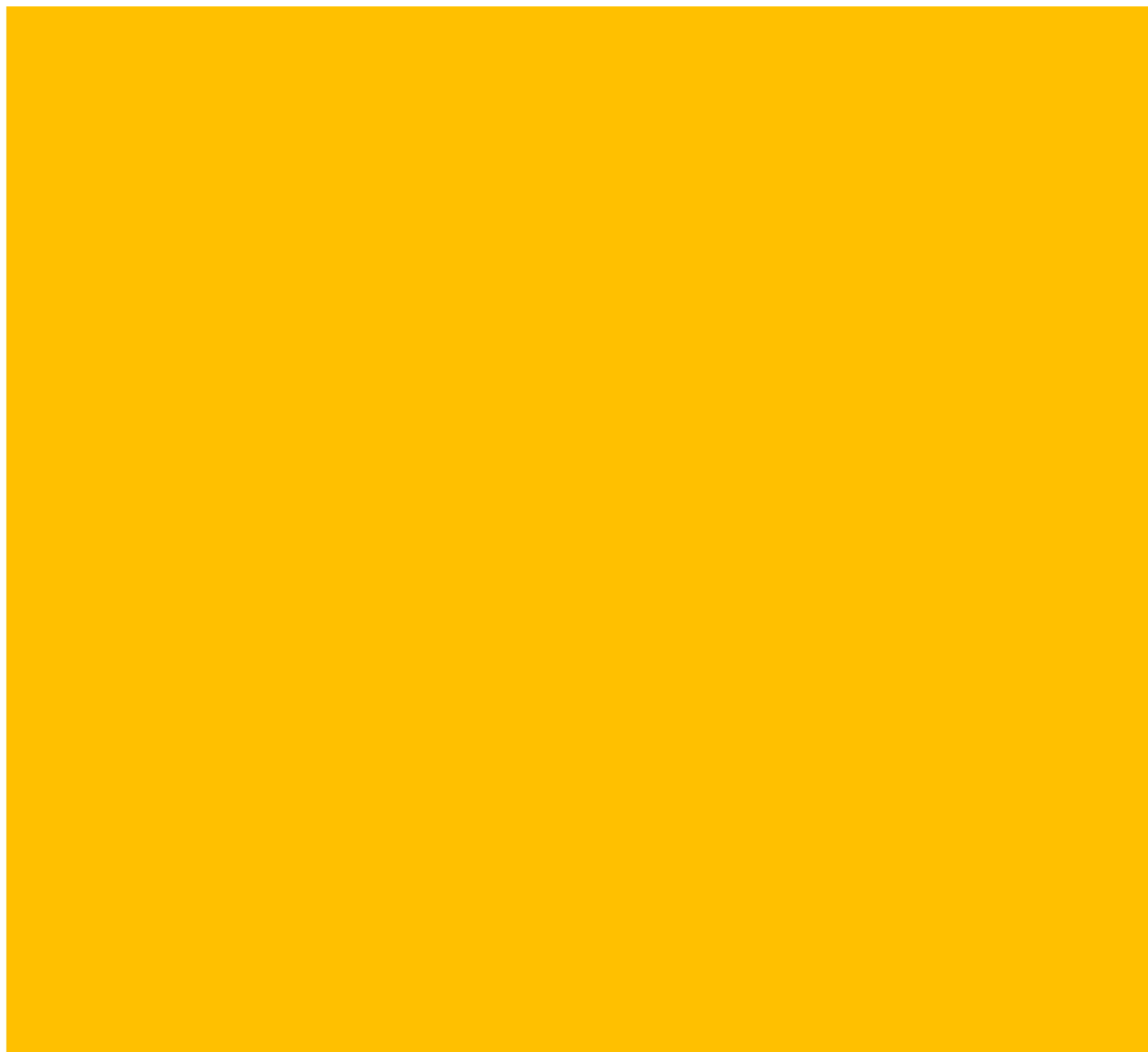

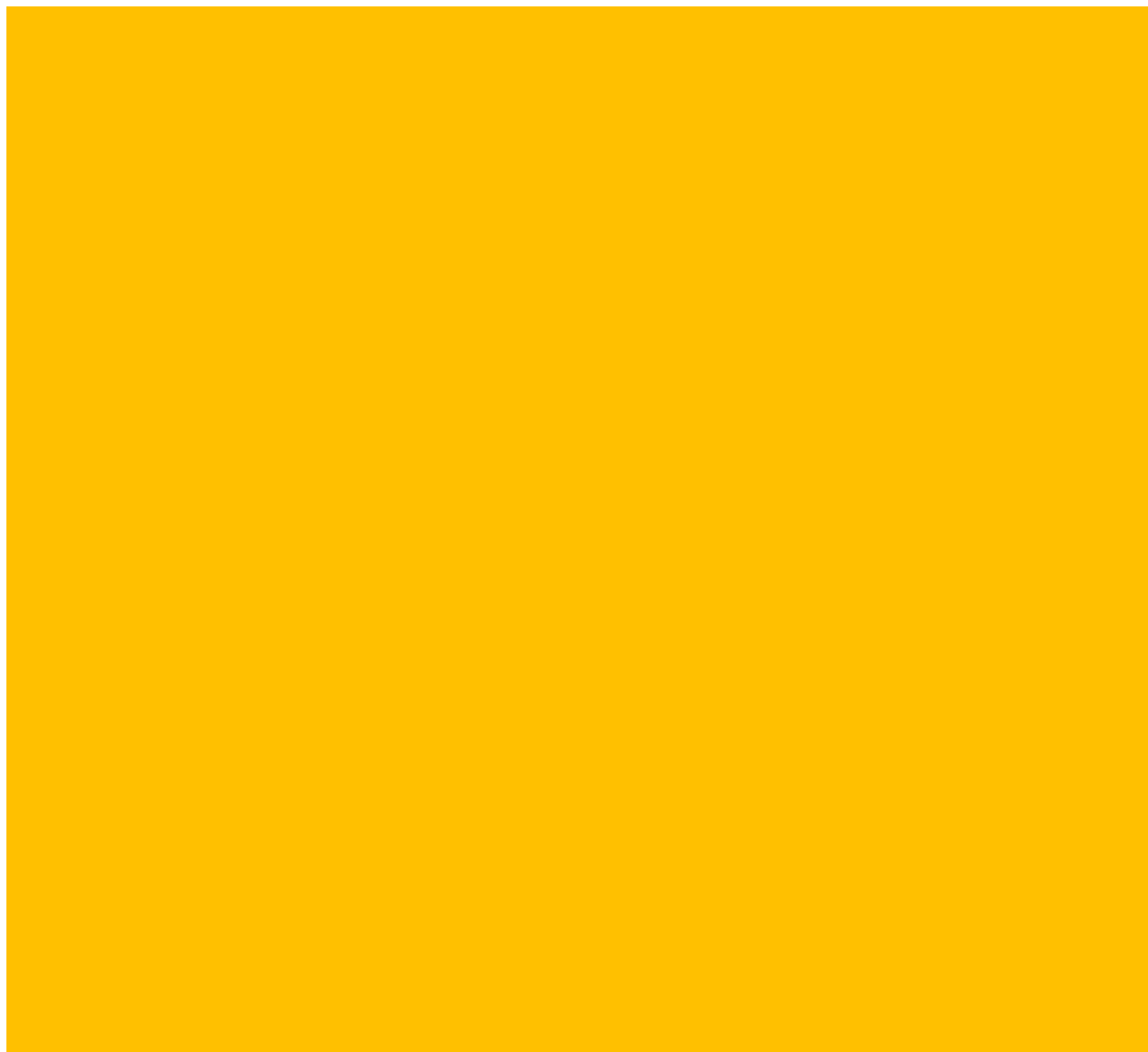

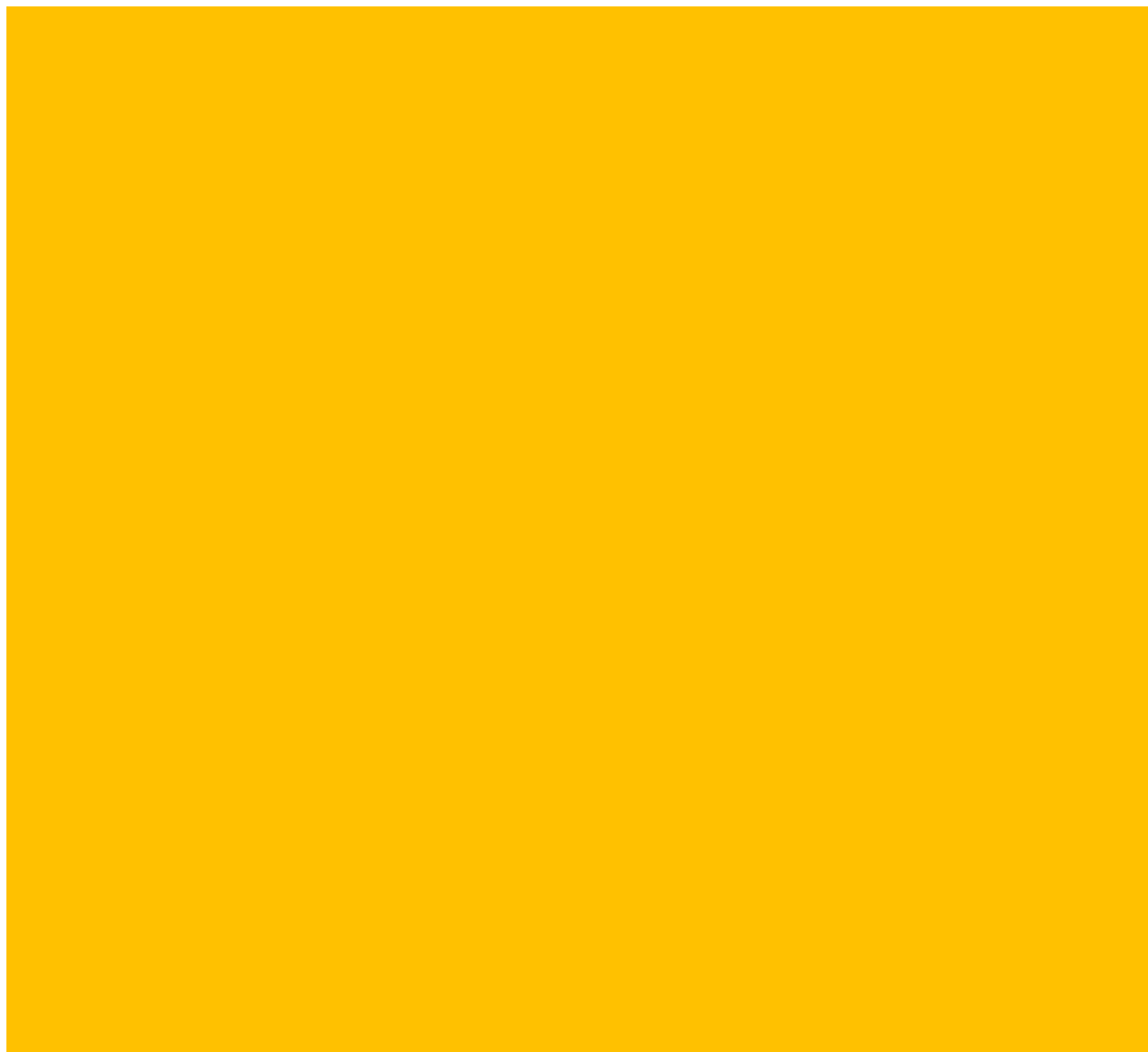

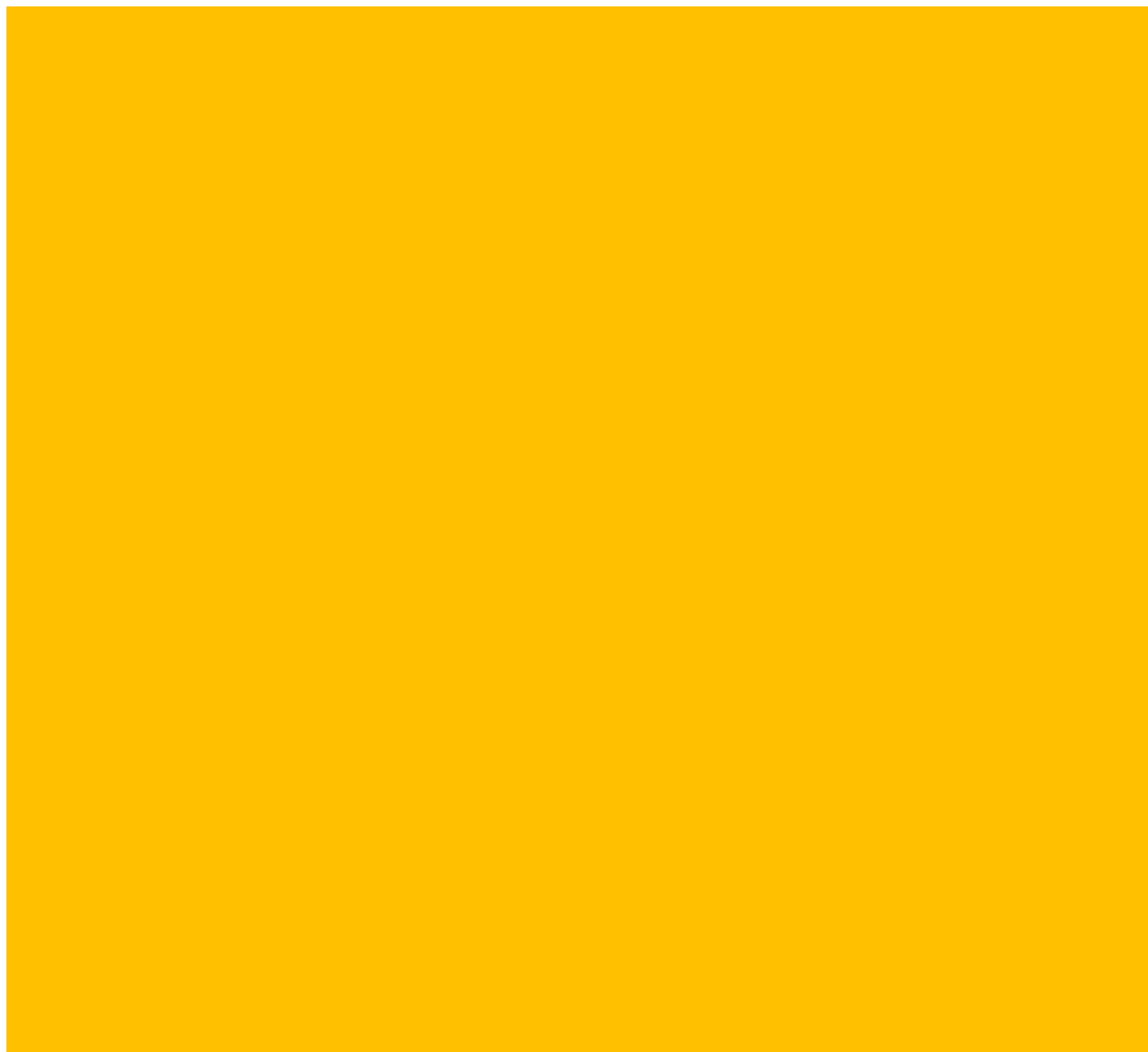

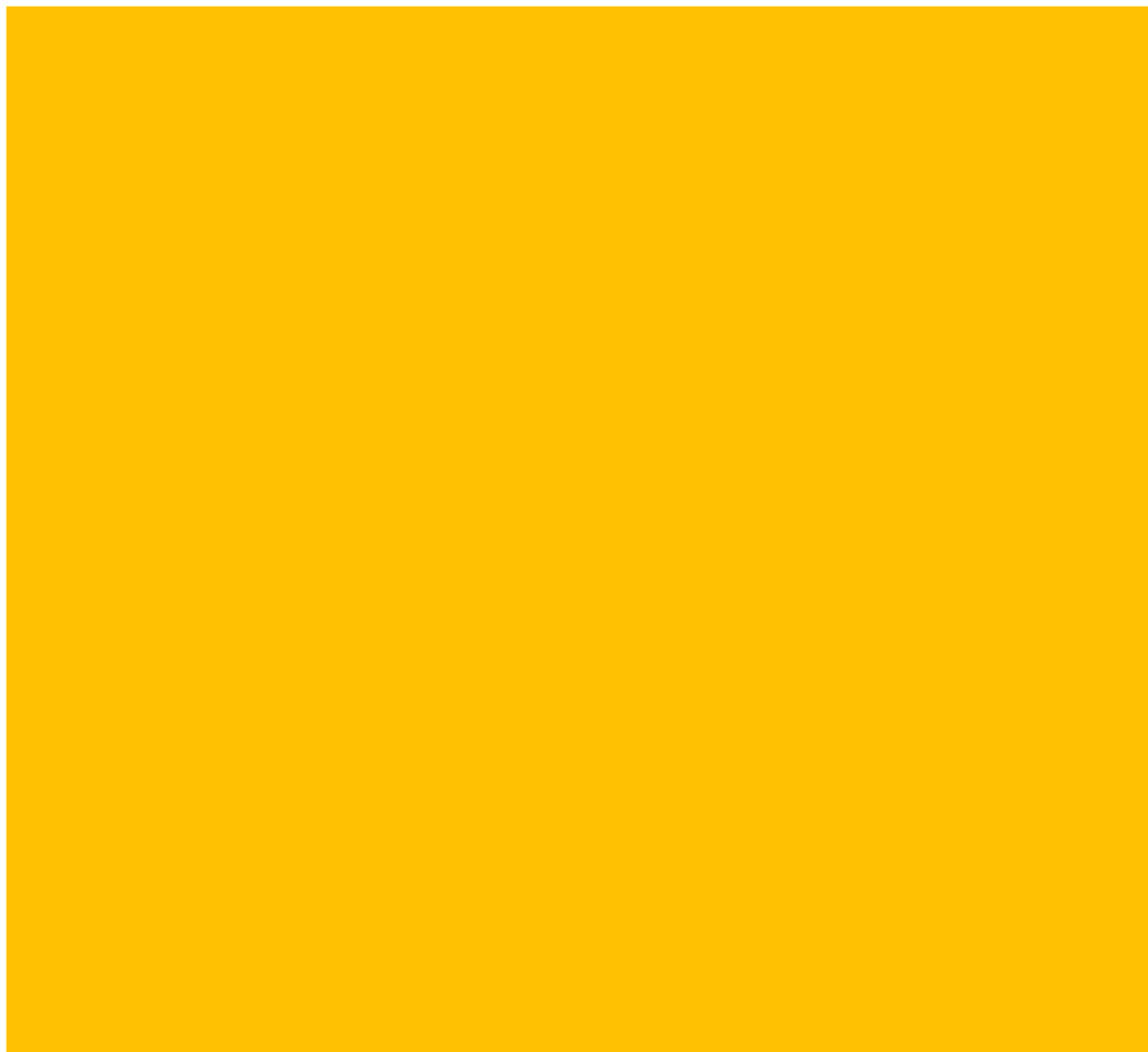

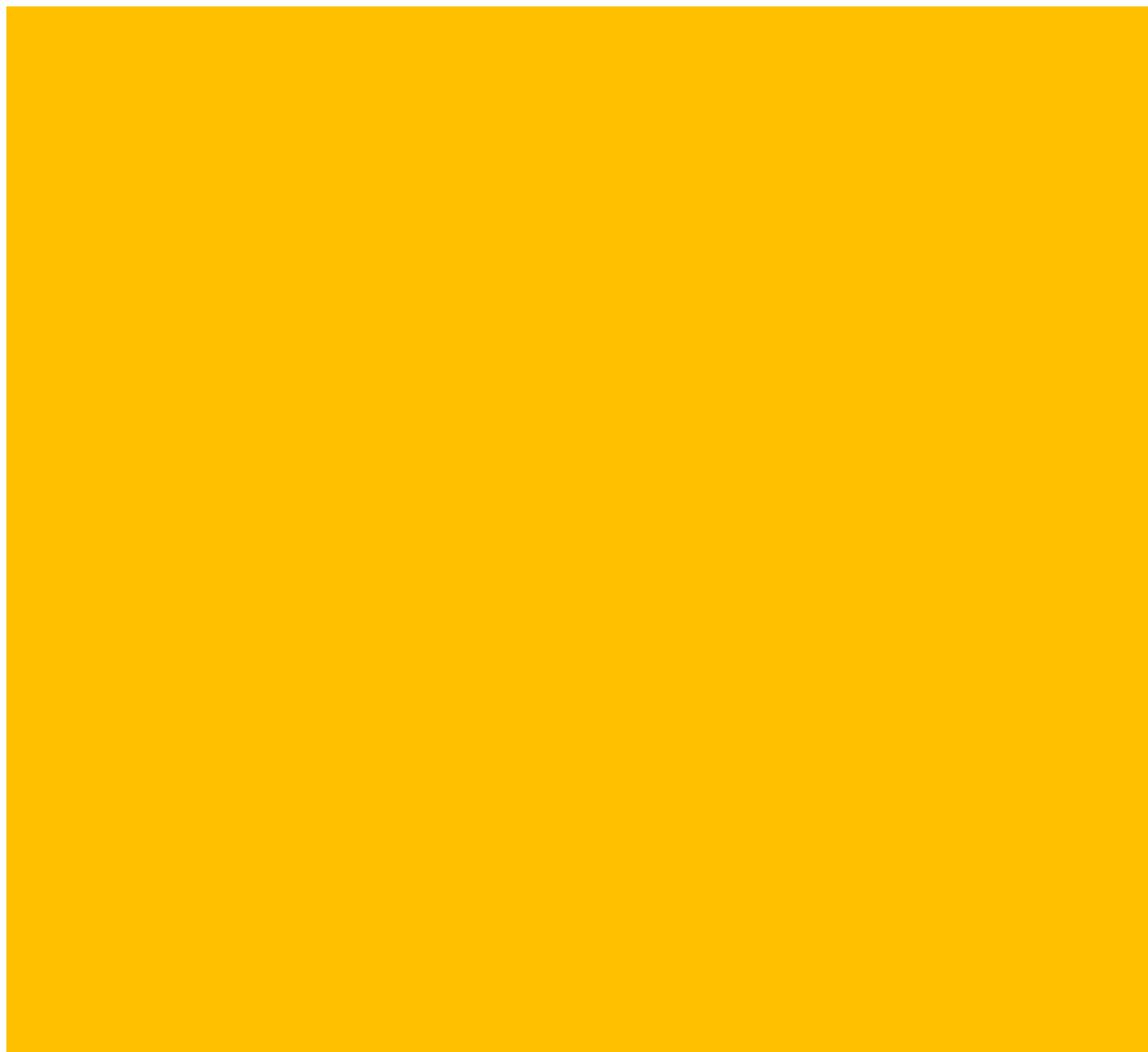

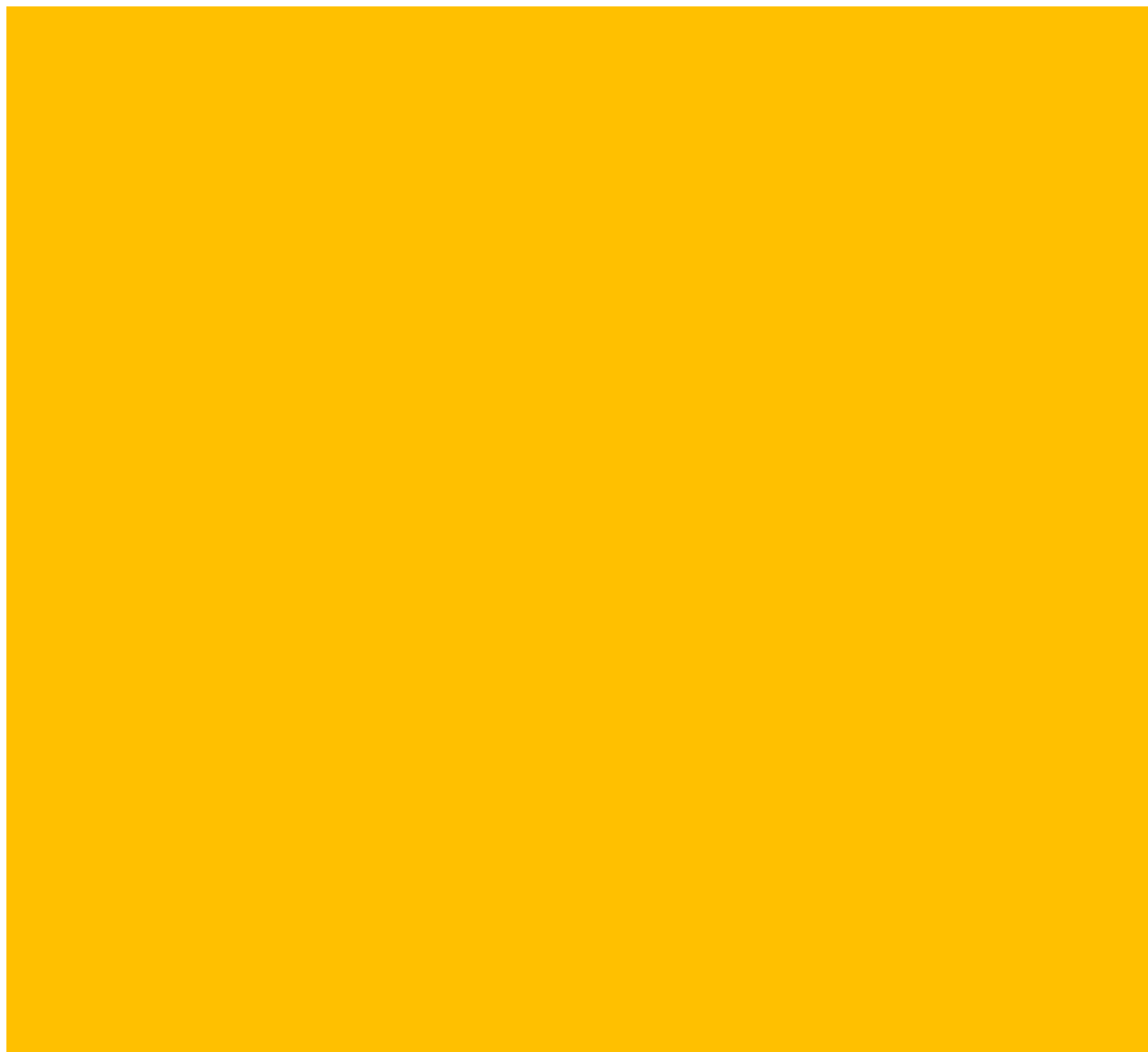

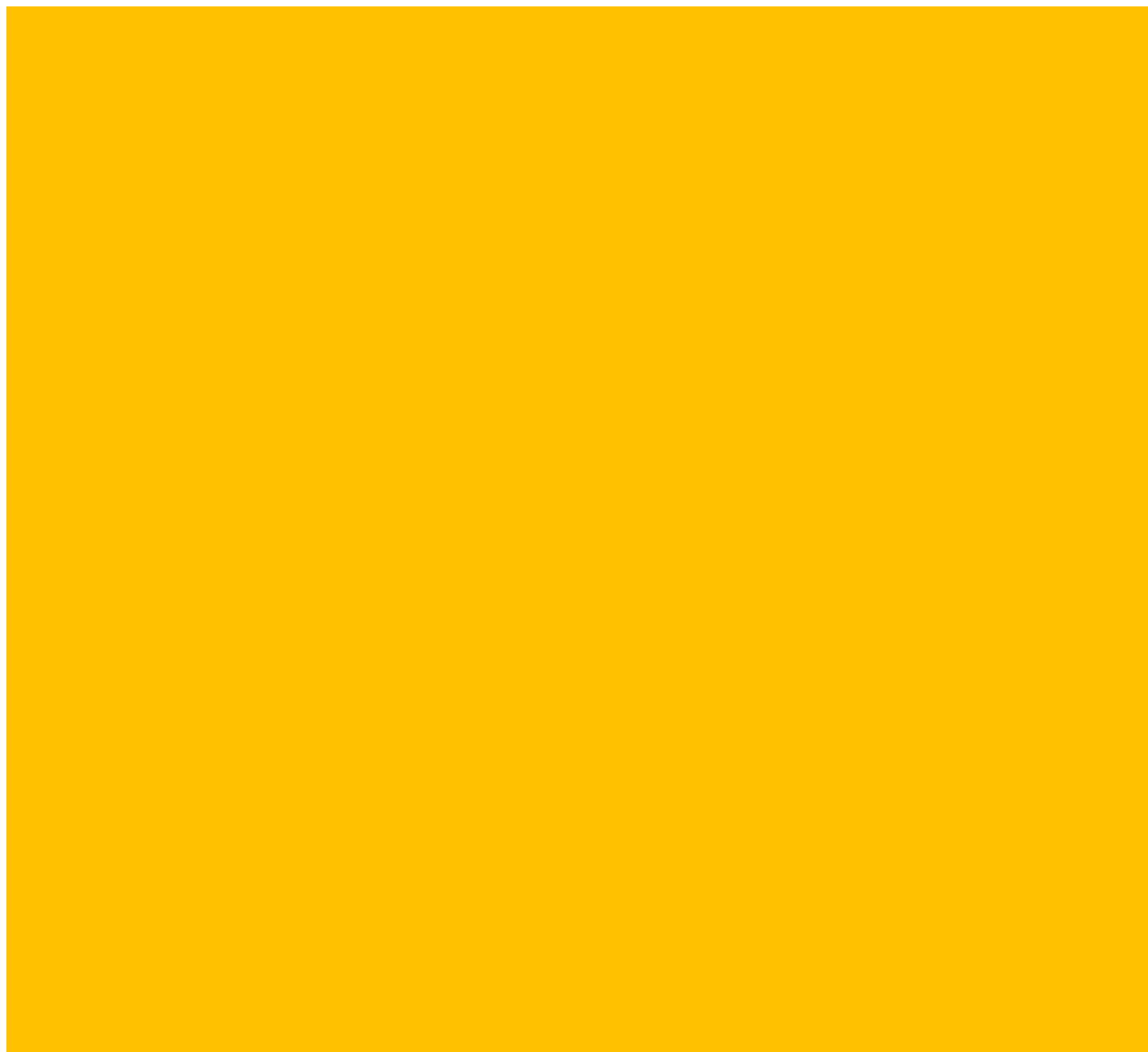

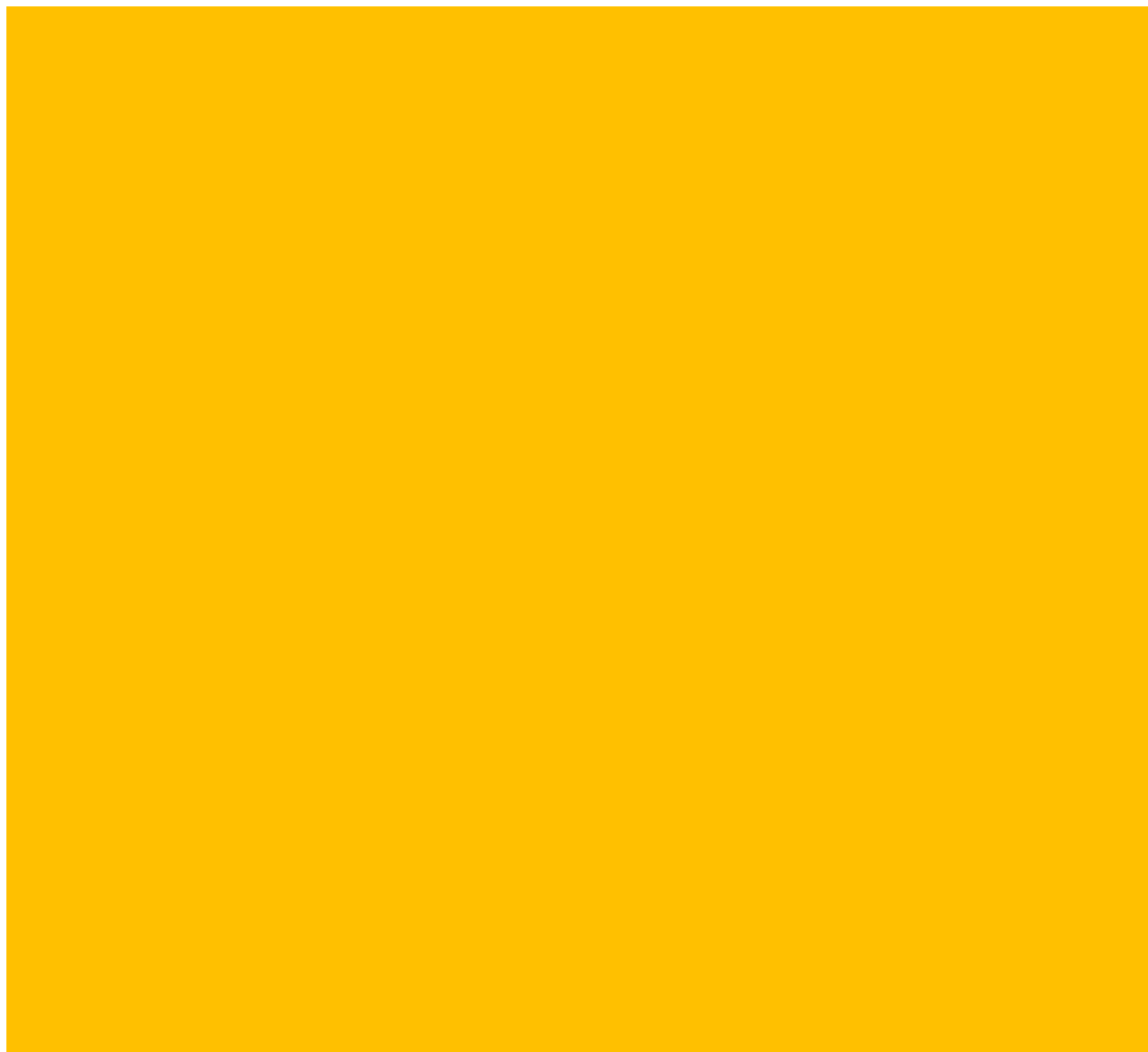

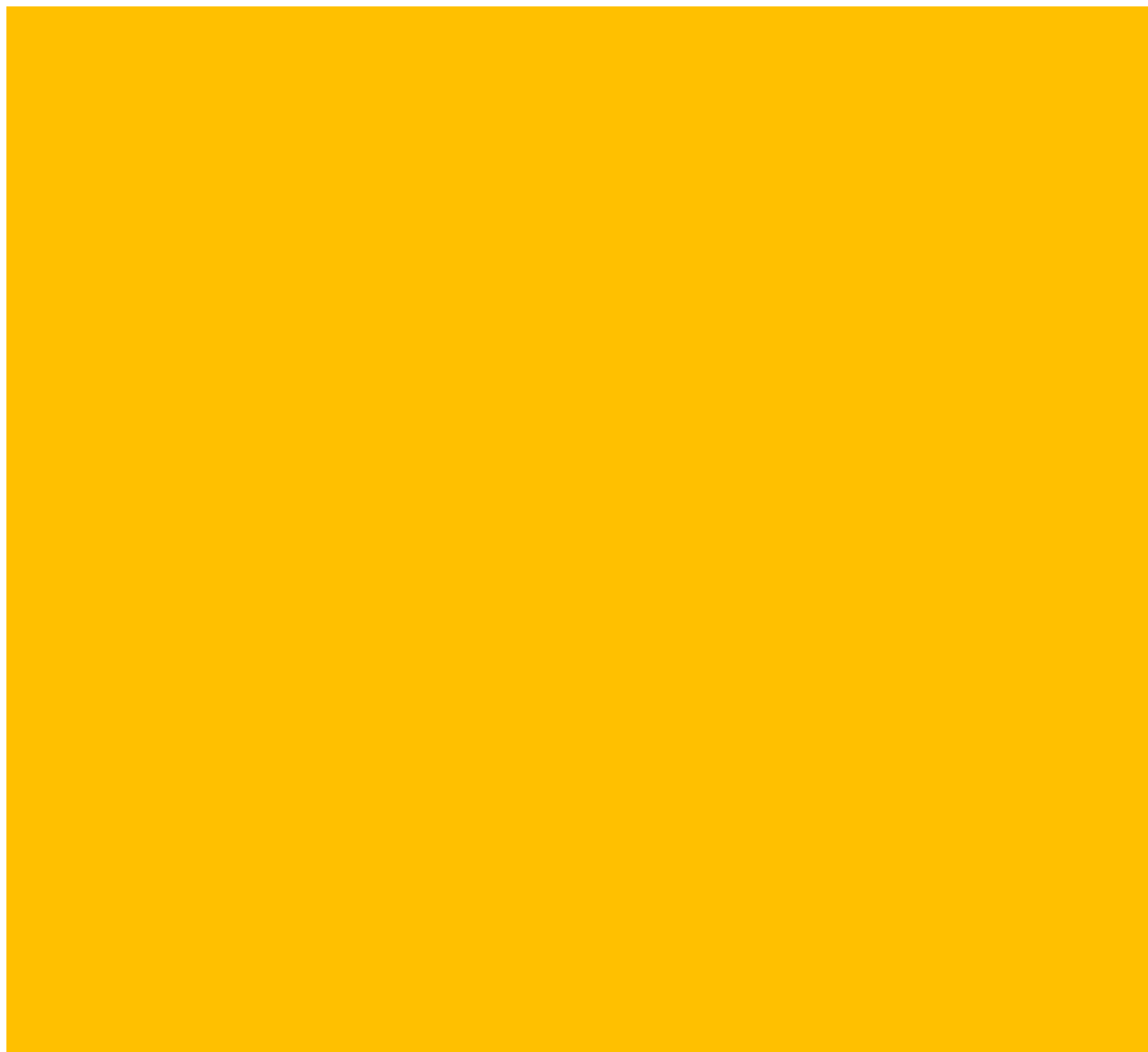

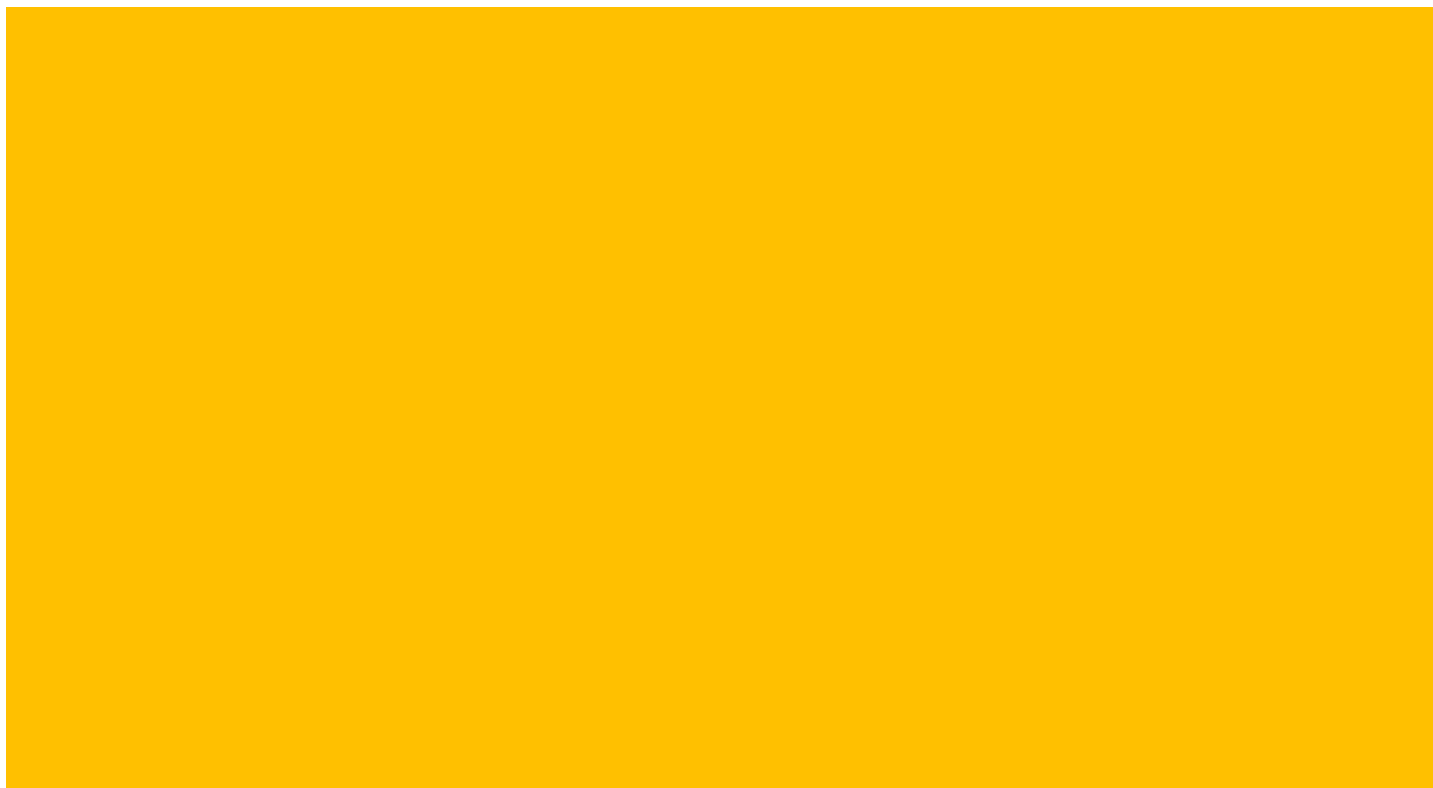

| Log2 Fold Change: (Bb_neg_newanalysis_12022024) / (RS_NEG_NEWANALYSIS_12022024) | Log2 Fold Change: (CM_neg_newanalysis_12022024) / (RS_NEG |
| --- | --- |
| 2.12 | 2.76 |
| 2.68 | 3.43 |
| 2.2 | 2.23 |
| 0.37 | 0.62 |
| 0.91 | 1.41 |
| 0.14 | 0.25 |
| -5.08 | 0.36 |
| -3.32 | -0.17 |
| -4.79 | 0.11 |
| -4.81 | 0.01 |
| -3.31 | 1.25 |
| -1.95 | -1.84 |
| 2.24 | 0.59 |
| 1.17 | 0.93 |
| 1.68 | 0.81 |
| -6.03 | -2.09 |
| -1.4 | -1.76 |
| -3.87 | -2.3 |
| 2.72 | -2.81 |
| 0.98 | 1.74 |
| -0.2 | 0.44 |
| 1.62 | 0.91 |
| 0.57 | 1.09 |
| 1.67 | 0.82 |
| -1.64 | 0.24 |
| -3.3 | 0.39 |
| -5.21 | 0.31 |
| -2.87 | 0.54 |
| 1.43 | 1.08 |
| -0.81 | 0.65 |
| -2.5 | 0.88 |
| -2.17 | 0.77 |
| -6.29 | -3.55 |
| 4.36 | -1.68 |
| 2.84 | -1.35 |
| 8.63 | 1.43 |
| 5.15 | -0.93 |
| 5.38 | 0.97 |
| 10.07 | 1.63 |
| 9.91 | 2.22 |
| 3.16 | 0.56 |
| 7.98 | 1.21 |
| 9.92 | 2.12 |
| 6.45 | 1.02 |
| 5.46 | 0.68 |
| -5.3 | -4.2 |
| -1.25 | -0.74 |
| 0.51 | 0.02 |
| -0.05 | -0.47 |
| -4.52 | -1.69 |
| 2.34 | 0.53 |
| 0.56 | 0.31 |
| -2.16 | -1.46 |
| -4.04 | -3.95 |
| -3.51 | -1.88 |
| 0.68 | -0.59 |
| -3.14 | -0.75 |
| 3.41 | 4.21 |
| -5.63 | -1.51 |
| -7.85 | -5.85 |
| -3.03 | 1.92 |
| -4.45 | -2.4 |
| 2.79 | 0.48 |
| -0.37 | -0.46 |
| -4.5 | -1.04 |
| -1.23 | -2.3 |
| -3.51 | -4.73 |
| 6.72 | 2.78 |
| 0 | -3.63 |
| -1.03 | -0.75 |
| -6.36 | -4.54 |
| 0.69 | -0.55 |
| -1.23 | -1.63 |
| 0.37 | 0.52 |
| -0.95 | -1.63 |
| -2.96 | -1.85 |
| 1.31 | -0.21 |
| -1.55 | -1.51 |
| -0.94 | -1.53 |
| -3.69 | -2.01 |
| -5.12 | -1.58 |
| -4.13 | -2.84 |
| -3.87 | -3.56 |
| -0.33 | -0.72 |
| -1.39 | -1.62 |
| -2.46 | -1.24 |
| 1.99 | 1.04 |
| 3.5 | -1.75 |
| -5.08 | -0.98 |
| -3.73 | -2.07 |
| -0.75 | -1.31 |
| 1.8 | 1.58 |
| 1.6 | 0.95 |
| -0.25 | -1.12 |
| 1.84 | 0.7 |
| 1.16 | 0.59 |
| -0.77 | 0.09 |
| -2.47 | -1.46 |
| -1.44 | -2.36 |
| 0.03 | 0.22 |
| 1.18 | 1.04 |
| 0.94 | 0.82 |
| 1.1 | 0.91 |
| -1.81 | -1.89 |
| 2.64 | 1.61 |
| -0.28 | 3.28 |
| 0.82 | 0.82 |
| -1.39 | -1.91 |
| -7.75 | -2.62 |
| -6.89 | -6.12 |
| -7.28 | -3.94 |
| -7.56 | -1.51 |
| -7.21 | -2.45 |
| -7.05 | -6.34 |
| -4.76 | -5.14 |
| -4.28 | -4.46 |
| 6.62 | 0.83 |
| 11.75 | 0.62 |
| 11.62 | 3.47 |
| -1.36 | -2.35 |
| -0.06 | -0.65 |
| 4.88 | 0.34 |
| 13.02 | 0.79 |
| -2.68 | -3.66 |
| 1.19 | 1.48 |
| -0.58 | 0.39 |
| -1.14 | -0.58 |
| 1.44 | 1.17 |
| 0.98 | 1.74 |
| 0.13 | 0.86 |
| -7.08 | -4.51 |
| -3.77 | -1.86 |
| -6.64 | -3.01 |
| -4.35 | -2.67 |
| -5.38 | -2.12 |
| -4.92 | -2.71 |
| -6.22 | -2.41 |
| -1.25 | -0.86 |
| -1.96 | -0.9 |
| 1.04 | 0.66 |
| -1.4 | -2.68 |
| -1.51 | -0.51 |
| -3.22 | -1.51 |
| -6.14 | -3.51 |
| -2.68 | -3.33 |
| -5.23 | -4.84 |
| -5.18 | -3.75 |
| -5.94 | -4.01 |
| -12.82 | -4.26 |
| -1.14 | -1.78 |
| -5.44 | -3.65 |
| -4.03 | -3.66 |
| -5.07 | -2.74 |
| -7.69 | -2.82 |
| -1.62 | -0.15 |
| -2.48 | 1.41 |
| 1.4 | 1.37 |
| -3.81 | -2.95 |
| -1.38 | -1.46 |
| -3.1 | -2.04 |
| -2.38 | -1.24 |
| -4.41 | -2.11 |
| -6.23 | -3.77 |
| -3.17 | -3.22 |
| -6.32 | -2.6 |
| -2.1 | -1.95 |
| 0.85 | 0.14 |
| -7.31 | -3.89 |
| -10.04 | -4.44 |
| -9.05 | -3.19 |
| -8.91 | -3.26 |
| -6.13 | -4.38 |
| -3.34 | 0.4 |
| -6.49 | -3.46 |
| -8.43 | -5.47 |
| -5.24 | -3.59 |
| -5.46 | -4.37 |
| -4.25 | -3.68 |
| -5.29 | -3.25 |
| -4.79 | -2.48 |
| -2.05 | -2.71 |

| Log2 Fold Change: (SM_neg_newanalysis_12022024) / (RS_NEG) | Log2 Fold Change: (Bb_neg_newanalysis_12022024) / (SM_neg_newa |
| --- | --- |
| 1.57 | 0.55 |
| 2.31 | 0.37 |
| 1.4 | 0.8 |
| 0.3 | 0.07 |
| 0.87 | 0.04 |
| 0.83 | -0.69 |
| 1.13 | -6.2 |
| 1.58 | -4.9 |
| 1.8 | -6.59 |
| 1.45 | -6.26 |
| 1.84 | -5.16 |
| -1.3 | -0.65 |
| 1.54 | 0.7 |
| 1.64 | -0.47 |
| 2.89 | -1.21 |
| -1.99 | -4.04 |
| -1.91 | 0.51 |
| -2.36 | -1.51 |
| -1.13 | 3.85 |
| 2.91 | -1.94 |
| 1.26 | -1.46 |
| 0.95 | 0.66 |
| 1.49 | -0.92 |
| 1.03 | 0.64 |
| 1.84 | -3.48 |
| 1.97 | -5.27 |
| 1.43 | -6.64 |
| 2.23 | -5.1 |
| 0.71 | 0.71 |
| 2.19 | -3 |
| 1.77 | -4.28 |
| 2.2 | -4.37 |
| -3.26 | -3.03 |
| -1.93 | 6.29 |
| -1.58 | 4.42 |
| 1.04 | 7.59 |
| -1.67 | 6.81 |
| 0.88 | 4.5 |
| 1.38 | 8.68 |
| 0.84 | 9.06 |
| 0.47 | 2.69 |
| 0.98 | 7 |
| 2.51 | 7.42 |
| 0.77 | 5.68 |
| 0.56 | 4.9 |
| -2.9 | -2.4 |
| -1.6 | 0.35 |
| 0.12 | 0.39 |
| -0.16 | 0.1 |
| -1.53 | -2.98 |
| 0.64 | 1.69 |
| -0.88 | 1.44 |
| -2.88 | 0.72 |
| -2.62 | -1.42 |
| -1.52 | -1.99 |
| -0.85 | 1.53 |
| -0.46 | -2.68 |
| 2.71 | 0.7 |
| -1.37 | -4.26 |
| -6.4 | -1.46 |
| -1.22 | -1.81 |
| -1.51 | -2.94 |
| 0.24 | 2.56 |
| -0.96 | 0.59 |
| -0.65 | -3.85 |
| -2.77 | 1.54 |
| -4.69 | 1.18 |
| 0.23 | 6.49 |
| -3.12 | 3.13 |
| -2.07 | 1.05 |
| -0.91 | -5.46 |
| -2.42 | 3.11 |
| -1.57 | 0.33 |
| -0.39 | 0.76 |
| -1.41 | 0.46 |
| -1.32 | -1.65 |
| -0.02 | 1.33 |
| -2.05 | 0.5 |
| -1.77 | 0.82 |
| -1.46 | -2.23 |
| -1.32 | -3.8 |
| -1.77 | -2.36 |
| -3.2 | -0.67 |
| -1.81 | 1.48 |
| -2.56 | 1.17 |
| -0.13 | -2.32 |
| 0.8 | 1.19 |
| -1.71 | 5.21 |
| -0.94 | -4.14 |
| -1.3 | -2.43 |
| -1.38 | 0.63 |
| 0.95 | 0.85 |
| 0.98 | 0.61 |
| -1.08 | 0.83 |
| 0.73 | 1.11 |
| 0.78 | 0.38 |
| -0.64 | -0.12 |
| -2.23 | -0.24 |
| -1.61 | 0.17 |
| -1.67 | 1.69 |
| 0.07 | 1.11 |
| 0.15 | 0.79 |
| 0.67 | 0.43 |
| -1.57 | -0.24 |
| -4.36 | 7 |
| 4.66 | -4.94 |
| 0.62 | 0.2 |
| -1.62 | 0.23 |
| -1.75 | -6 |
| -4.25 | -2.64 |
| -2.87 | -4.41 |
| -0.65 | -6.9 |
| -1.73 | -5.47 |
| -5.43 | -1.62 |
| -5.12 | 0.36 |
| -4.99 | 0.71 |
| 0.93 | 5.69 |
| 0.6 | 11.14 |
| 0.88 | 10.75 |
| -2.6 | 1.24 |
| 0.15 | -0.21 |
| 0.41 | 4.48 |
| 1.64 | 11.37 |
| -3.06 | 0.38 |
| 0.74 | 0.45 |
| -1.51 | 0.94 |
| -1.44 | 0.3 |
| 0.82 | 0.62 |
| -0.16 | 1.13 |
| -1.04 | 1.17 |
| -2.43 | -4.66 |
| -2.01 | -1.76 |
| -1.71 | -4.93 |
| -3.36 | -0.98 |
| -1.68 | -3.7 |
| -1.05 | -3.87 |
| -3.06 | -3.15 |
| -1.56 | 0.3 |
| -0.72 | -1.24 |
| -0.1 | 1.15 |
| -3.8 | 2.4 |
| 1.05 | -2.56 |
| -1.87 | -1.35 |
| -3.76 | -2.38 |
| -2.06 | -0.62 |
| -3.4 | -1.83 |
| -2.79 | -2.39 |
| -3.96 | -1.98 |
| -3.2 | -9.63 |
| -2.03 | 0.9 |
| -2.5 | -2.94 |
| -2.97 | -1.06 |
| -2.24 | -2.84 |
| -1.97 | -5.72 |
| -0.46 | -1.15 |
| 1.79 | -4.28 |
| 0.75 | 0.65 |
| -2.19 | -1.62 |
| -1.79 | 0.41 |
| -2.12 | -0.98 |
| -1.95 | -0.42 |
| -1.74 | -2.67 |
| -3.21 | -3.01 |
| -2.94 | -0.23 |
| -2.84 | -3.48 |
| -2.99 | 0.9 |
| 0.02 | 0.83 |
| -3.87 | -3.44 |
| -3.95 | -6.09 |
| -3.08 | -5.97 |
| -3.01 | -5.9 |
| -3.85 | -2.28 |
| -2.95 | -0.39 |
| -3.39 | -3.1 |
| -4.47 | -3.96 |
| -3.33 | -1.91 |
| -3.45 | -2.01 |
| -4.25 | 0 |
| -2.99 | -2.3 |
| -1.8 | -2.99 |
| -2.95 | 0.9 |

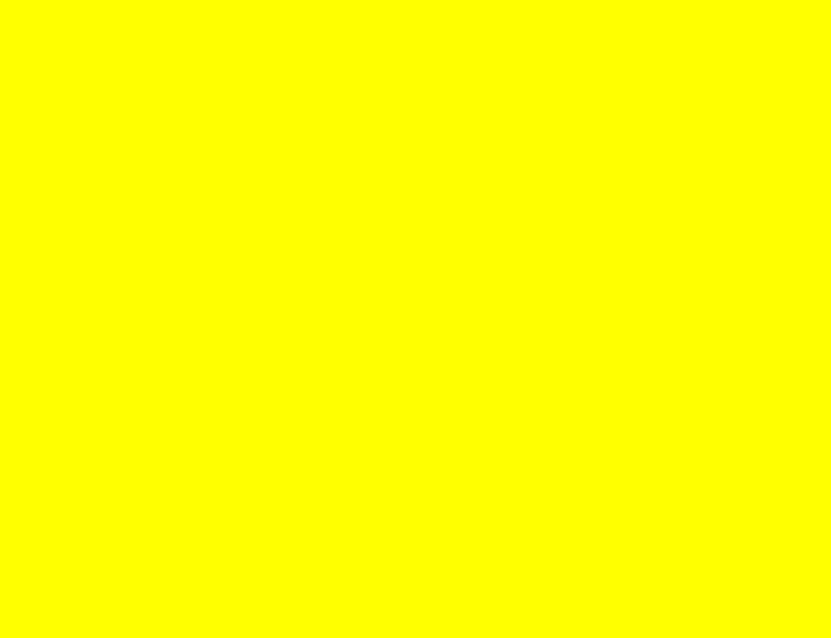

| Log2 Fold Change: (CM_neg_newanalysis_12022024) / (SM_neg_newanalysis_12022024) | Log2 Fold Change: (Bb_neg_newanalysis_12022024) / (WATER_NEG_NEWANALYSIS_12022024) |
| --- | --- |
| 1.19 | -1.26 |
| 1.12 | -1.34 |
| 0.83 | 0.24 |
| 0.33 | -1.82 |
| 0.55 | -1.54 |
| -0.58 | -0.48 |
| -0.76 | -1.46 |
| -1.75 | 0 |
| -1.69 | -1.55 |
| -1.44 | -1.42 |
| -0.59 | -1.23 |
| -0.54 | -2.05 |
| -0.96 | -1.38 |
| -0.7 | -2.64 |
| -2.08 | -3.94 |
| -0.1 | -1.9 |
| 0.15 | -1.25 |
| 0.06 | -1.44 |
| -1.68 | 5.23 |
| -1.17 | -1.32 |
| -0.82 | -1.12 |
| -0.04 | -1.18 |
| -0.4 | -1.1 |
| -0.21 | -0.98 |
| -1.6 | -1.64 |
| -1.58 | -1.58 |
| -1.12 | -2.29 |
| -1.69 | -1.36 |
| 0.37 | -1.32 |
| -1.54 | -1.3 |
| -0.89 | -1.13 |
| -1.43 | -1.67 |
| -0.29 | -1.43 |
| 0.25 | 4.68 |
| 0.23 | 2.67 |
| 0.39 | 6.1 |
| 0.73 | 4.96 |
| 0.08 | 2.34 |
| 0.24 | 7.41 |
| 1.38 | 6.8 |
| 0.09 | 0.83 |
| 0.23 | 5.55 |
| -0.38 | 7.29 |
| 0.25 | 3.76 |
| 0.12 | 2.84 |
| -1.3 | -0.69 |
| 0.86 | -1.63 |
| -0.11 | -1.29 |
| -0.31 | -1.66 |
| -0.16 | -0.86 |
| -0.12 | -0.26 |
| 1.19 | -0.24 |
| 1.42 | -1 |
| -1.33 | -0.39 |
| -0.36 | -0.82 |
| 0.26 | 0.82 |
| -0.29 | -1.57 |
| 1.49 | -1.31 |
| -0.13 | -1.22 |
| 0.55 | -1.42 |
| 3.14 | -1.25 |
| -0.89 | -1.4 |
| 0.24 | 0.49 |
| 0.5 | -1.28 |
| -0.38 | -1.25 |
| 0.47 | 2.59 |
| -0.04 | -0.76 |
| 2.55 | 5 |
| -0.51 | 1.32 |
| 1.32 | -0.1 |
| -3.64 | -1.77 |
| 1.87 | 2.28 |
| -0.06 | -1.41 |
| 0.91 | -1.01 |
| -0.22 | 0.02 |
| -0.54 | -0.96 |
| -0.19 | -0.42 |
| 0.53 | -0.72 |
| 0.24 | -1.12 |
| -0.55 | -0.04 |
| -0.26 | -1.08 |
| -1.06 | -1.22 |
| -0.35 | -0.75 |
| 1.09 | -0.42 |
| 0.94 | -0.68 |
| -1.11 | -1.16 |
| 0.24 | -0.92 |
| -0.04 | 3.53 |
| -0.04 | -0.91 |
| -0.77 | 0.2 |
| 0.07 | -1.21 |
| 0.63 | -1.19 |
| -0.03 | -1.34 |
| -0.04 | -1.1 |
| -0.03 | -0.71 |
| -0.19 | -1.49 |
| 0.74 | -1.68 |
| 0.78 | -1.3 |
| -0.75 | 3.55 |
| 1.89 | -0.21 |
| 0.98 | -0.66 |
| 0.68 | -0.95 |
| 0.23 | -1.59 |
| -0.32 | -1.4 |
| 5.97 | 5.77 |
| -1.38 | -0.99 |
| 0.2 | -1.01 |
| -0.29 | -1.45 |
| -0.87 | -1.42 |
| -1.87 | -1.17 |
| -1.07 | -1.16 |
| -0.86 | -0.83 |
| -0.71 | -0.62 |
| -0.91 | -1.17 |
| -0.02 | -1.44 |
| 0.53 | -0.78 |
| -0.1 | 3.82 |
| 0.01 | 9.2 |
| 2.59 | 9.26 |
| 0.25 | -0.85 |
| -0.8 | -1.06 |
| -0.07 | 2.52 |
| -0.85 | 11.25 |
| -0.6 | -0.99 |
| 0.74 | -1.54 |
| 1.9 | -0.81 |
| 0.86 | -1.61 |
| 0.35 | -0.5 |
| 1.89 | -0.61 |
| 1.9 | -0.57 |
| -2.08 | -1.34 |
| 0.15 | -1.13 |
| -1.3 | -1.08 |
| 0.69 | -0.21 |
| -0.44 | -0.36 |
| -1.66 | -0.86 |
| 0.65 | -1.89 |
| 0.7 | -0.82 |
| -0.18 | -1.12 |
| 0.76 | -0.9 |
| 1.12 | 1.46 |
| -1.56 | -1.61 |
| 0.36 | -1.1 |
| 0.25 | -0.75 |
| -1.27 | -1.08 |
| -1.44 | -1.51 |
| -0.96 | -0.59 |
| -0.05 | -1.7 |
| -1.06 | -3.54 |
| 0.25 | 0.51 |
| -1.15 | -1.83 |
| -0.68 | -1.32 |
| -0.5 | 0.12 |
| -0.86 | -1.57 |
| 0.32 | -1.15 |
| -0.39 | -1.83 |
| 0.62 | -1.18 |
| -0.76 | 0.21 |
| 0.33 | -1.57 |
| 0.08 | -1.45 |
| 0.71 | -1.44 |
| -0.37 | -1 |
| -0.55 | -1.15 |
| -0.29 | 0.13 |
| 0.24 | -1.71 |
| 1.05 | -1.07 |
| 0.13 | -1.27 |
| -0.02 | -1.45 |
| -0.48 | -1.26 |
| -0.1 | -2.33 |
| -0.25 | -0.85 |
| -0.54 | -1.47 |
| 3.35 | -1.3 |
| -0.07 | 0.28 |
| -1 | 0.06 |
| -0.26 | -0.52 |
| -0.92 | -1.46 |
| 0.57 | -0.58 |
| -0.26 | -1.13 |
| -0.68 | -1.38 |
| 0.24 | -1.16 |

| Log2 Fold Change: (CM_neg_newanalysis_12022024) / (WATER_NEG_NEWANALYSIS_12022024) |  |
| --- | --- |
|  | -0.62 |
|  | -0.59 |
|  | 0.28 |
|  | -1.57 |
|  | -1.03 |
|  | -0.37 |
|  | 3.98 |
|  | 3.15 |
|  | 3.34 |
|  | 3.4 |
|  | 3.33 |
|  | -1.94 |
|  | -3.04 |
|  | -2.87 |
|  | -4.81 |
|  | 2.03 |
|  | -1.61 |
|  | 0.13 |
|  | -0.3 |
|  | -0.56 |
|  | -0.48 |
|  | -1.88 |
|  | -0.58 |
|  | -1.84 |
|  | 0.24 |
|  | 2.11 |
|  | 3.22 |
|  | 2.06 |
|  | -1.67 |
|  | 0.17 |
|  | 2.25 |
|  | 1.27 |
|  | 1.31 |

|  |
| --- |
| -1.36 |
| -1.52 |
| -1.09 |
| -1.12 |
| -2.08 |
| -1.03 |
| -0.89 |
| -1.77 |
| -1.22 |
| -0.51 |
| -1.67 |
| -1.93 |
| 0.42 |
| -1.13 |
| -1.78 |
| -2.08 |
| 1.97 |
| -2.07 |
| -0.49 |
| -0.3 |
| -0.3 |
| 0.81 |
| -0.45 |
| 0.83 |
| -0.51 |
| 2.9 |
| 0.59 |
| 3.71 |
| 0.65 |
| -1.82 |
| -1.36 |
| 2.21 |
| 1.52 |
| -1.98 |
| 1.07 |
| -2.31 |
| 0.17 |
| 0.04 |
| 1.04 |
| -1.8 |
| -0.86 |
| -0.67 |
| 0.15 |
| -1.94 |
| -0.68 |
| -1.7 |
| 1.64 |
| 2.45 |
| 0.07 |
| -0.44 |
| -0.81 |
| -0.91 |
| 0.05 |
| -1.86 |
| -1.73 |
| 3.19 |
| 1.86 |
| -1.76 |
| -1.41 |
| -1.98 |
| -1.97 |
| -1.85 |
| -2.05 |
| -0.82 |
| -0.28 |
| 2.64 |
| -0.01 |
| -0.8 |
| -1.06 |
| -1.79 |
| -1.49 |
| 4.74 |
| 2.58 |
| -1.01 |
| -1.97 |
| 3.71 |
| -0.4 |
| 2.18 |
| 5.21 |
| 4.14 |
| -0.46 |
| -1.82 |
| -0.95 |
| -1.97 |
| -1.93 |
| 1.11 |
| -1.84 |
| -1.65 |
| -2.02 |
| -0.97 |
| -1.97 |
| -1.25 |
| 0.15 |
| -1.05 |
| -0.77 |
| 0.15 |
| 0.15 |
| 1.23 |
| 0.78 |
| 2.55 |
| 1.47 |
| 2.9 |
| 1.35 |
| 1.91 |
| -0.42 |
| -0.06 |
| -1.29 |
| 0.18 |
| -0.61 |
| 0.61 |
| 1.88 |
| -1.73 |
| -1.11 |
| 0.84 |
| 0.23 |
| 5.02 |
| -0.14 |
| -0.04 |
| -0.95 |
| 2.46 |
| 3.29 |
| 0.32 |
| 2.06 |
| -1.21 |
| 1.06 |
| -1.65 |
| -0.39 |
| -0.3 |
| 1.3 |
| 1.31 |
| 0.08 |
| 2.01 |
| -0.92 |
| -1.97 |
| 1.98 |
| 4.35 |
| 3.53 |
| 4.79 |
| 0.27 |
| 2.44 |

|  |  |
| --- | --- |
|  | 3.31 |
|  | 3.02 |
|  | 1.13 |
|  | -0.37 |
|  | -0.01 |
|  | 0.9 |
|  | 0.93 |
|  | -1.82 |

| Log2 Fold Change: (RS_NEG_NEWANALYSIS_12022024) / (WATER_NEG_NEV |
| --- |
| -3.38 |
| -4.02 |
| -1.96 |
| -2.19 |
| -2.44 |
| -0.62 |
| 3.61 |
| 3.32 |
| 3.24 |
| 3.38 |
| 2.08 |
| -0.1 |
| -3.62 |
| -3.8 |
| -5.62 |
| 4.13 |
| 0.15 |
| 2.43 |
| 2.51 |
| -2.3 |
| -0.92 |
| -2.8 |
| -1.67 |
| -2.66 |
| 0 |
| 1.72 |
| 2.91 |
| 1.52 |
| -2.75 |
| -0.49 |
| 1.37 |
| 0.5 |
| 4.85 |
| 0.32 |
| -0.17 |
| -2.53 |
| -0.19 |
| -3.05 |
| -2.66 |
| -3.11 |
| -2.33 |
| -2.43 |
| -2.63 |
| -2.69 |
| -2.62 |
| 4.62 |
| -0.38 |
| -1.8 |
| -1.6 |
| 3.66 |
| -2.59 |
| -0.8 |
| 1.16 |
| 3.65 |
| 2.69 |
| 0.15 |
| 1.58 |
| -4.72 |
| 4.41 |
| 6.44 |
| 1.79 |
| 3.05 |
| -2.3 |
| -0.91 |
| 3.25 |
| 3.82 |
| 2.75 |
| -1.72 |

|  |  |
| --- | --- |
|  | 1.32 |
|  | 0.92 |
|  | 4.59 |
|  | 1.59 |
|  | -0.17 |
|  | -1.38 |
|  | 0.96 |
|  | 2.01 |
|  | -1.73 |
|  | 0.84 |
|  | -0.18 |
|  | 3.65 |
|  | 4.03 |
|  | 2.91 |
|  | 3.12 |
|  | -0.09 |
|  | 0.7 |
|  | 1.29 |
|  | -2.9 |
|  | 0.02 |
|  | 4.17 |
|  | 3.92 |
|  | -0.45 |
|  | -2.99 |
|  | -2.93 |
|  | -0.85 |
|  | -2.55 |
|  | -2.65 |
|  | -0.91 |
|  | 1.18 |
|  | 4.99 |
|  | -0.24 |
|  | -1.84 |
|  | -1.89 |
|  | -2.69 |

|  |
| --- |
| 0.4 |
| 3.13 |
| -0.7 |
| -1.83 |
| -0.07 |
| 6.34 |
| 5.72 |
| 6.12 |
| 6.72 |
| 6.58 |
| 5.88 |
| 3.31 |
| 3.5 |
| -2.8 |
| -2.54 |
| -2.37 |
| 0.51 |
| -1.01 |
| -2.36 |
| -1.77 |
| 1.69 |
| -2.73 |
| -0.23 |
| -0.47 |
| -1.94 |
| -1.59 |
| -0.71 |
| 5.74 |
| 2.64 |
| 5.56 |
| 4.14 |
| 5.02 |
| 4.06 |
| 4.32 |
| 0.44 |
| 0.84 |
| -1.95 |
| 2.86 |
| -0.1 |
| 2.12 |
| 5.39 |
| 1.6 |
| 3.72 |
| 4.59 |
| 4.24 |
| 9.28 |
| 1.64 |
| 3.61 |
| 2.7 |
| 5.2 |
| 6.12 |
| 0.47 |
| 0.65 |
| -2.58 |
| 4.02 |
| -0.19 |
| 1.64 |
| 0.94 |
| 3.41 |
| 5.08 |
| 3.3 |
| 4.61 |
| 1.03 |
| -2.12 |
| 5.87 |
| 8.78 |
| 6.71 |
| 8.06 |
| 4.65 |
| 2.04 |

|  |  |
| --- | --- |
|  | 6.77 |
|  | 8.49 |
|  | 4.72 |
|  | 4 |
|  | 3.66 |
|  | 4.16 |
|  | 3.42 |
|  | 0.89 |

| Log2 Fold Change: (SM_neg_newanalysis_12022024)/(WATER_NEG_NEWANALYSIS | P-value: (Bb_neg_newanalysis_12022024)/(CM_ |
| --- | --- |
| -1.81 | 0.998887562 |
| -1.71 | 0.997207754 |
| -0.56 | 0.999929847 |
| -1.9 | 0.691706019 |
| -1.57 | 0.510642703 |
| 0.21 | 0.767955458 |
| 4.74 | 5.20695E-14 |
| 4.9 | 3.04322E-05 |
| 5.04 | 4.98923E-06 |
| 4.84 | 8.21376E-12 |
| 3.92 | 2.73576E-10 |
| -1.4 | 0.992741652 |
| -2.08 | 0.375109609 |
| -2.17 | 0.475617325 |
| -2.73 | 0.831605454 |
| 2.13 | 0.000877359 |
| -1.76 | 0.999038506 |
| 0.07 | 0.005353313 |
| 1.38 | 2.44273E-09 |
| 0.61 | 0.7392595 |
| 0.34 | 0.831348992 |
| -1.84 | 0.968518487 |
| -0.18 | 0.999933604 |
| -1.63 | 0.556453232 |
| 1.84 | 0.227919622 |
| 3.69 | 2.73407E-06 |
| 4.35 | 7.16996E-08 |
| 3.74 | 7.08531E-05 |
| -2.04 | 0.947194982 |
| 1.7 | 0.323812154 |
| 3.14 | 0.009963098 |
| 2.7 | 0.000943023 |
| 1.59 | 5.14729E-06 |
| -1.61 | 1.59573E-10 |
| -1.75 | 2.5796E-05 |
| -1.49 | 1.13191E-07 |
| -1.85 | 1.13446E-09 |
| -2.16 | 5.39934E-09 |
| -1.27 | 6.25056E-14 |
| -2.27 | 1.19348E-10 |
| -1.86 | 0.004174598 |
| -1.45 | 4.85167E-14 |
| -0.12 | 5.69544E-14 |
| -1.92 | 3.68279E-10 |
| -2.06 | 1.35299E-10 |
| 1.72 | 0.326979209 |
| -1.98 | 0.997355663 |
| -1.68 | 0.946705208 |
| -1.76 | 0.997161756 |
| 2.13 | 3.16355E-08 |
| -1.95 | 0.445698447 |
| -1.68 | 0.999999509 |
| -1.72 | 0.920473305 |
| 1.03 | 0.961663234 |
| 1.17 | 0.623966979 |
| -0.71 | 0.002489766 |
| 1.11 | 8.4713E-07 |
| -2.01 | 0.917265312 |
| 3.04 | 0.000180934 |
| 0.04 | 0.121888986 |
| 0.57 | 1.18638E-08 |
| 1.54 | 0.000170391 |
| -2.06 | 0.000545196 |
| -1.86 | 0.999991493 |
| 2.6 | 8.51129E-11 |
| 1.05 | 0.341149057 |
| -1.94 | 0.895340855 |
| -1.49 | 0.0029226 |
| -1.8 | 0.000320641 |
| -1.15 | 0.946999704 |
| 3.68 | 0.057979746 |
| -0.83 | 0.711401902 |
| -1.74 | 0.996716775 |
| -1.78 | 0.999878796 |
| -0.45 | 0.341264537 |
| 0.69 | 0.097439819 |
| -1.75 | 0.818724359 |
| -1.21 | 0.999999293 |
| -1.95 | 0.998750662 |
| 2.19 | 0.009383766 |
| 2.71 | 1.04229E-06 |
| 1.13 | 0.001117396 |
| -0.08 | 1 |
| -1.9 | 0.84340626 |
| -1.86 | 0.527507991 |
| 1.16 | 0.026000951 |
| -2.1 | 0.520688518 |
| -1.69 | 1.48643E-05 |
| 3.23 | 2.30249E-12 |
| 2.63 | 0.003310303 |
| -1.84 | 0.966538245 |
| -2.04 | 0.991905325 |
| -1.95 | 0.999966273 |
| -1.93 | 0.961245578 |
| -1.82 | 0.40046124 |
| -1.87 | 0.957663852 |
| -1.56 | 0.940578451 |
| -1.06 | 0.596793629 |
| 3.38 | 0.451875905 |
| -1.9 | 0.999982392 |
| -1.77 | 0.999512118 |
| -1.74 | 0.995697441 |
| -2.02 | 0.981697833 |
| -1.16 | 0.999722182 |
| -1.23 | 0.999996065 |
| 3.96 | 0.075957875 |
| -1.21 | 0.999825949 |
| -1.68 | 0.998209024 |
| 4.58 | 3.18655E-11 |
| 1.47 | 0.017433527 |
| 3.25 | 2.74682E-07 |
| 6.07 | 4.99419E-10 |
| 4.85 | 1.46764E-05 |
| 0.45 | 0.089361049 |
| -1.8 | 0.99999277 |
| -1.48 | 0.961490295 |
| -1.87 | 2.49856E-12 |
| -1.94 | 4.60743E-14 |
| -1.49 | 4.69624E-14 |
| -2.09 | 0.768978039 |
| -0.85 | 0.999952085 |
| -1.95 | 1.78374E-10 |
| -0.12 | 4.60743E-14 |
| -1.37 | 0.999700727 |
| -1.99 | 0.988337559 |
| -1.75 | 0.986381699 |
| -1.91 | 0.990329985 |
| -1.12 | 0.74721748 |
| -1.74 | 0.994208262 |
| -1.75 | 0.994422861 |
| 3.31 | 1.44308E-06 |
| 0.63 | 0.000546281 |
| 3.85 | 9.3584E-10 |
| 0.77 | 0.025548659 |
| 3.34 | 6.96178E-06 |
| 3.01 | 9.65226E-05 |
| 1.26 | 5.46892E-07 |
| -1.12 | 0.952202916 |
| 0.13 | 0.098813529 |
| -2.05 | 0.22406121 |
| -0.94 | 0.282886195 |
| 0.95 | 0.739411481 |
| 0.25 | 0.000522498 |
| 1.63 | 0.000165152 |
| -0.46 | 0.989201012 |
| 0.32 | 0.708841357 |
| 1.8 | 3.8228E-05 |
| 0.28 | 0.007726467 |
| 6.08 | 4.65183E-14 |
| -0.39 | 0.999962408 |
| 1.11 | 0.004883342 |
| -0.27 | 0.944601223 |
| 2.96 | 2.66814E-06 |
| 4.15 | 1.72318E-12 |
| 0 | 0.026857003 |
| 2.44 | 9.99451E-09 |
| -1.83 | 0.999999458 |
| 1.82 | 0.175825086 |
| -1.98 | 0.999927445 |
| -0.47 | 0.038912472 |
| -1.01 | 0.731532807 |
| 1.67 | 9.80003E-05 |
| 1.87 | 3.50434E-06 |
| 0.37 | 0.998999636 |
| 1.77 | 4.26398E-11 |
| -1.97 | 0.995410569 |
| -2.1 | 0.922513506 |
| 1.99 | 2.08161E-08 |
| 4.83 | 3.55252E-11 |
| 3.63 | 1.47027E-12 |
| 5.04 | 5.02931E-14 |
| 0.81 | 0.027078951 |
| -0.91 | 0.008434152 |
| 3.38 | 7.4573E-05 |
| 4.02 | 0.000909207 |
| 1.39 | 0.000335983 |
| 0.55 | 0.469399523 |
| -0.58 | 0.928580997 |
| 1.17 | 0.001959158 |
| 1.61 | 0.000289853 |
| -2.06 | 0.992221451 |

| P-value: (Bb_neg_newanalysis_12022024)/(RS | P-value: (CM_neg_newanalysis_12022024)/(RS_NEG_NEWANALYS |
| --- | --- |
| 0.000706636 | 0.000248991 |
| 6.9329E-06 | 1.98688E-06 |
| 2.13912E-05 | 1.18163E-05 |
| 0.421765486 | 0.023740456 |
| 0.072226931 | 0.000865663 |
| 0.964514708 | 0.994817208 |
| 7.89369E-14 | 0.923860827 |
| 5.15053E-05 | 0.999962202 |
| 1.62574E-05 | 0.997949172 |
| 1.55898E-10 | 0.746777337 |
| 6.98921E-07 | 0.037395632 |
| 0.304107594 | 0.630694968 |
| 0.319916943 | 0.999997931 |
| 0.031003794 | 0.703804924 |
| 0.208782356 | 0.860825572 |
| 1.91956E-06 | 0.258077779 |
| 0.029321762 | 0.011905026 |
| 1.52067E-10 | 2.84235E-06 |
| 0.000465299 | 0.000941805 |
| 0.762593733 | 0.101673468 |
| 0.999993838 | 0.888323633 |
| 0.000168095 | 0.001444941 |
| 0.223749582 | 0.318378792 |
| 0.000332181 | 0.027493259 |
| 0.453105118 | 0.997509098 |
| 1.36151E-06 | 0.999828248 |
| 3.26317E-08 | 0.999583661 |
| 0.000112789 | 0.999979676 |
| 0.008254623 | 0.069093389 |
| 0.514754408 | 0.999367838 |
| 0.00745311 | 0.999997098 |
| 0.002297793 | 0.999435595 |
| 1.10023E-13 | 8.25717E-08 |
| 2.00299E-08 | 0.34750754 |
| 0.001035541 | 0.763912709 |
| 5.94412E-10 | 0.296003382 |
| 8.31403E-09 | 0.959996621 |
| 3.17002E-12 | 0.0242915 |
| 4.65183E-14 | 0.28185812 |
| 9.08051E-13 | 0.200513106 |
| 0.00013721 | 0.808181529 |
| 4.60743E-14 | 0.006245558 |
| 4.60743E-14 | 0.01106061 |
| 3.22509E-12 | 0.265151103 |
| 8.42326E-13 | 0.167183714 |
| 9.65821E-09 | 2.10362E-06 |
| 0.034832446 | 0.095532572 |
| 0.705547682 | 0.993508948 |
| 0.999801308 | 0.999957083 |
| 9.35918E-14 | 9.00896E-06 |
| 0.385766875 | 0.99999802 |
| 0.972888904 | 0.958601374 |
| 0.000124957 | 0.001854014 |
| 8.73797E-11 | 5.57701E-10 |
| 2.71133E-05 | 0.002055552 |
| 0.463831267 | 0.185249605 |
| 7.16691E-10 | 0.081279077 |
| 8.32339E-06 | 5.68821E-07 |
| 1.22868E-08 | 0.013341167 |
| 1.70762E-09 | 1.35153E-06 |
| 3.42266E-05 | 0.054049397 |
| 3.57525E-12 | 7.47917E-07 |
| 1.84793E-06 | 0.333417804 |
| 0.983397534 | 0.962376842 |
| 1.25566E-13 | 0.017481196 |
| 0.022045548 | 9.24598E-05 |
| 0.023682062 | 0.001547404 |
| 1.88151E-07 | 0.015257985 |
| 0.99967474 | 0.000719911 |
| 0.289479649 | 0.800672274 |
| 3.59714E-08 | 0.000107202 |
| 0.951901552 | 0.992822611 |
| 0.067002924 | 0.022293447 |
| 0.971084989 | 0.912118267 |
| 0.105022291 | 0.000625016 |
| 4.06716E-06 | 0.007249773 |
| 0.962748861 | 0.998228534 |
| 0.001848524 | 0.002325038 |
| 0.27412206 | 0.135681508 |
| 1.83964E-08 | 0.000416215 |
| 2.03385E-10 | 0.019108434 |
| 2.81564E-12 | 9.07552E-08 |
| 7.84669E-10 | 7.7689E-10 |
| 0.975993871 | 0.414368699 |
| 0.724708435 | 0.041088071 |
| 2.27981E-06 | 0.018597423 |
| 0.000141035 | 0.014861373 |
| 0.000854629 | 0.689037054 |
| 4.78506E-14 | 0.005500061 |
| 1.21169E-08 | 0.000786972 |
| 0.111623533 | 0.018419975 |
| 0.000341376 | 0.001652835 |
| 0.028622663 | 0.018259043 |
| 0.995957826 | 0.999461681 |
| 9.88922E-05 | 0.017880258 |
| 0.005496312 | 0.043803624 |
| 0.998716181 | 0.994871616 |
| 1.00188E-05 | 0.000869539 |
| 0.200932583 | 0.002658071 |
| 0.992488988 | 0.998470694 |
| 0.881947574 | 0.72211388 |
| 0.477490054 | 0.7810304 |
| 0.001473517 | 0.009135895 |
| 0.154022834 | 0.086636329 |
| 0.705623435 | 0.774301604 |
| 0.88946866 | 0.491021471 |
| 0.167285342 | 0.264989798 |
| 0.115715313 | 0.046924964 |
| 4.62963E-14 | 1.25793E-06 |
| 4.67404E-14 | 3.62044E-13 |
| 4.67404E-14 | 1.82073E-09 |
| 1.91958E-13 | 0.00551079 |
| 6.77092E-12 | 1.96219E-05 |
| 4.65183E-14 | 8.49321E-14 |
| 3.51708E-06 | 2.43038E-06 |
| 0.000791858 | 8.19254E-05 |
| 5.87308E-14 | 0.080849736 |
| 4.60743E-14 | 0.247182059 |
| 4.60743E-14 | 8.20069E-05 |
| 0.024058587 | 0.000752021 |
| 0.928656322 | 0.855004806 |
| 5.05185E-12 | 0.565726265 |
| 4.60743E-14 | 0.975202587 |
| 1.79119E-05 | 8.08494E-06 |
| 0.007454847 | 0.001422779 |
| 0.991433509 | 0.819409433 |
| 0.073878162 | 0.238731674 |
| 0.010407404 | 0.215110371 |
| 0.229707902 | 0.080685918 |
| 0.968580255 | 0.775829335 |
| 4.62963E-14 | 5.43802E-11 |
| 3.61674E-08 | 0.014412251 |
| 4.60743E-14 | 3.14783E-09 |
| 8.79794E-09 | 5.96982E-05 |
| 4.47226E-11 | 0.000495231 |
| 2.26716E-10 | 0.000270707 |
| 8.76743E-13 | 3.10938E-05 |
| 0.981241366 | 0.631231664 |
| 0.000216453 | 0.187997644 |
| 0.012407228 | 0.772992448 |
| 0.657289827 | 0.010658873 |
| 0.004395507 | 0.117222998 |
| 2.98937E-10 | 6.98505E-05 |
| 2.34146E-13 | 1.53909E-08 |
| 1.05474E-06 | 2.07346E-07 |
| 1.12721E-12 | 2.0013E-11 |
| 5.20695E-14 | 4.87218E-10 |
| 7.01939E-12 | 4.50553E-08 |
| 4.60743E-14 | 4.91737E-09 |
| 0.774090568 | 0.663604306 |
| 1.65412E-12 | 1.13099E-08 |
| 3.88393E-06 | 4.58609E-05 |
| 7.4718E-14 | 4.83882E-08 |
| 4.60743E-14 | 5.72357E-08 |
| 0.002440149 | 0.933495559 |
| 1.46182E-05 | 0.09083046 |
| 0.019416048 | 0.015902958 |
| 4.24201E-07 | 0.000337056 |
| 0.920837839 | 0.835249366 |
| 7.6431E-09 | 3.10978E-05 |
| 0.000692851 | 0.026270503 |
| 2.80443E-11 | 1.92213E-05 |
| 4.60743E-14 | 5.20861E-12 |
| 1.64219E-07 | 4.34158E-07 |
| 4.60743E-14 | 2.76645E-08 |
| 0.233934038 | 0.499515657 |
| 0.356590688 | 0.89935117 |
| 4.60743E-14 | 6.11433E-12 |
| 4.60743E-14 | 9.95598E-11 |
| 4.60743E-14 | 2.95399E-07 |
| 4.60743E-14 | 1.12546E-09 |
| 4.9627E-14 | 1.37212E-12 |
| 0.001741854 | 0.990445147 |
| 9.77984E-12 | 6.37551E-06 |
| 1.50879E-13 | 1.59203E-09 |
| 6.42819E-14 | 3.7479E-10 |
| 2.20379E-13 | 6.11255E-12 |
| 6.25343E-09 | 6.83806E-08 |
| 1.95921E-12 | 3.33329E-08 |
| 8.77109E-12 | 1.46068E-06 |
| 7.03557E-05 | 1.44585E-05 |

| P-value: (SM_neg_newanalysis_12022024) / (RS_NEG_NEWANALYSIS) | P-value: (Bb_neg_newanalysis_12022024) / (SM_neg_newanalysis_12022024) |
| --- | --- |
| 0.019503181 | 0.802748177 |
| 0.000102616 | 0.922706473 |
| 0.002416389 | 0.532891428 |
| 0.623563198 | 0.999425968 |
| 0.029668344 | 0.9987094 |
| 0.763637036 | 0.295349088 |
| 0.028630613 | 4.62963E-14 |
| 0.055744794 | 1.77125E-08 |
| 0.129869386 | 1.69664E-08 |
| 0.012798897 | 1.50657E-13 |
| 0.002266859 | 2.39307E-11 |
| 0.961312008 | 0.7826884 |
| 0.631921493 | 0.994379358 |
| 0.331959252 | 0.841489661 |
| 0.014029622 | 0.819214743 |
| 0.267883788 | 0.000825405 |
| 0.000269918 | 0.493388192 |
| 8.04556E-06 | 0.00194652 |
| 0.371797932 | 1.92621E-06 |
| 0.001808263 | 0.052563374 |
| 0.0235568 | 0.017067923 |
| 0.03953496 | 0.32567323 |
| 0.008141009 | 0.677477387 |
| 0.035013584 | 0.493374431 |
| 0.019174666 | 0.000137777 |
| 0.00185842 | 3.48316E-11 |
| 0.189917372 | 1.09834E-10 |
| 0.002723801 | 1.83894E-09 |
| 0.664463131 | 0.234420049 |
| 0.00283641 | 2.32819E-05 |
| 0.072483033 | 2.86805E-06 |
| 0.003184981 | 3.14028E-08 |
| 3.94531E-07 | 1.0148E-06 |
| 0.061333346 | 2.15251E-11 |
| 0.827313829 | 3.61653E-05 |
| 0.577110691 | 3.19976E-08 |
| 0.599156227 | 1.87416E-10 |
| 0.246718562 | 3.98232E-10 |
| 0.482125355 | 5.4512E-14 |
| 0.426244182 | 4.21035E-11 |
| 0.992910143 | 0.000644707 |
| 0.119669242 | 4.65183E-14 |
| 0.0002036 | 2.00617E-13 |
| 0.676889543 | 7.28254E-11 |
| 0.68313086 | 1.57091E-11 |
| 0.000610427 | 0.003318262 |
| 0.002047957 | 0.872560377 |
| 0.999921311 | 0.569916505 |
| 0.9999439 | 0.996832024 |
| 0.000433181 | 1.04525E-09 |
| 0.998895689 | 0.20962582 |
| 0.153047672 | 0.029699125 |
| 1.9041E-05 | 0.983081887 |
| 6.04605E-07 | 0.012382411 |
| 0.045985331 | 0.085094957 |
| 0.210027428 | 0.002995125 |
| 0.430620408 | 7.24816E-08 |
| 0.002835076 | 0.305955046 |
| 0.221349967 | 5.21022E-06 |
| 3.00108E-07 | 0.33368216 |
| 0.907660097 | 0.000577995 |
| 0.003254619 | 4.47045E-08 |
| 0.999303502 | 4.69902E-06 |
| 0.167492729 | 0.488055281 |
| 0.247788957 | 6.09435E-12 |
| 6.94795E-05 | 0.290383756 |
| 0.000544227 | 0.705223813 |
| 0.915403367 | 2.68604E-06 |
| 0.007377096 | 0.003418126 |
| 0.019395546 | 0.787921944 |
| 0.826379999 | 8.54429E-07 |
| 0.02555875 | 0.002800715 |
| 0.001258408 | 0.617558699 |
| 0.949989708 | 0.999997817 |
| 0.005767702 | 0.82346189 |
| 0.111620032 | 0.006111123 |
| 0.867874939 | 0.400713807 |
| 0.000112393 | 0.908268467 |
| 0.002294695 | 0.322860734 |
| 0.02556321 | 0.000136206 |
| 0.285524628 | 3.60542E-08 |
| 0.000102275 | 8.85886E-07 |
| 2.293E-08 | 0.730875528 |
| 0.004254926 | 0.027246755 |
| 0.003618831 | 0.106256423 |
| 0.978595239 | 1.61687E-05 |
| 0.323712618 | 0.034419197 |
| 0.325204937 | 2.75701E-06 |
| 0.047168066 | 4.62186E-13 |
| 0.024852839 | 8.83177E-05 |
| 0.000451039 | 0.269687728 |
| 0.15826886 | 0.164588499 |
| 0.450876927 | 0.708431296 |
| 0.876709885 | 0.606288718 |
| 0.283889604 | 0.030933774 |
| 0.253940008 | 0.537668851 |
| 0.989725263 | 0.999926008 |
| 3.45028E-05 | 0.997513624 |
| 0.075436175 | 0.996124128 |
| 0.005898802 | 0.02456638 |
| 0.999997754 | 0.920286181 |
| 0.985759546 | 0.854836231 |
| 0.322144604 | 0.207128066 |
| 0.117795224 | 0.999992755 |
| 0.342433361 | 0.017646634 |
| 0.060594866 | 0.004371446 |
| 0.553407638 | 0.968570378 |
| 0.028734453 | 0.988031123 |
| 0.000595116 | 3.96683E-13 |
| 8.70292E-11 | 1.00594E-05 |
| 8.26187E-07 | 6.85745E-10 |
| 0.901512008 | 1.08813E-12 |
| 0.079953857 | 3.80091E-09 |
| 8.75855E-13 | 0.001464996 |
| 4.27405E-07 | 0.968176756 |
| 1.55589E-05 | 0.715504012 |
| 0.385279612 | 5.0393E-13 |
| 0.544922574 | 4.60743E-14 |
| 0.270910044 | 4.60743E-14 |
| 2.34888E-06 | 0.020676092 |
| 0.994965849 | 0.997837094 |
| 0.980202484 | 2.17718E-11 |
| 0.166137169 | 4.60743E-14 |
| 8.51052E-06 | 0.99978434 |
| 0.079897271 | 0.917010829 |
| 0.976144983 | 0.773996143 |
| 0.000927047 | 0.51967107 |
| 0.174444423 | 0.809116413 |
| 0.9399216 | 0.740795711 |
| 0.976197201 | 0.658122871 |
| 1.74431E-05 | 7.84295E-12 |
| 0.000594599 | 0.013342022 |
| 0.002528238 | 7.4496E-14 |
| 1.5696E-06 | 0.363672445 |
| 0.039451048 | 7.7809E-08 |
| 0.63359557 | 8.93052E-09 |
| 2.76238E-06 | 5.90566E-06 |
| 0.514002825 | 0.896177772 |
| 0.491422463 | 0.024409516 |
| 0.999980229 | 0.018777018 |
| 8.12349E-05 | 0.005095262 |
| 0.999890538 | 0.002351989 |
| 2.90167E-05 | 0.001242838 |
| 1.39938E-09 | 0.00247083 |
| 0.000182605 | 0.436465387 |
| 2.28539E-08 | 0.001337621 |
| 3.26078E-08 | 3.39038E-07 |
| 8.91642E-08 | 0.003852231 |
| 5.08268E-06 | 4.60743E-14 |
| 0.015514575 | 0.260251621 |
| 9.26219E-06 | 4.41899E-06 |
| 0.000181874 | 0.731598283 |
| 1.74088E-05 | 8.71377E-09 |
| 9.41719E-05 | 5.67324E-14 |
| 0.324843049 | 0.282782363 |
| 0.005503572 | 6.08485E-10 |
| 0.636177823 | 0.426498011 |
| 0.00237799 | 0.038727054 |
| 0.353607005 | 0.899251861 |
| 1.71095E-05 | 0.063490851 |
| 0.002860931 | 0.994807699 |
| 0.000969585 | 1.98078E-06 |
| 5.66636E-10 | 1.26407E-08 |
| 2.83087E-06 | 0.890067088 |
| 4.0292E-09 | 2.40348E-10 |
| 0.010757496 | 0.727229173 |
| 0.999378433 | 0.205321959 |
| 4.38954E-11 | 2.11256E-09 |
| 2.02221E-08 | 4.21108E-13 |
| 2.16124E-06 | 3.86469E-13 |
| 3.61894E-08 | 4.65183E-14 |
| 1.67431E-10 | 5.84827E-05 |
| 0.002531198 | 0.999991928 |
| 5.25732E-05 | 9.00726E-06 |
| 1.58043E-07 | 5.52197E-06 |
| 5.31237E-09 | 1.4984E-05 |
| 5.16991E-09 | 0.000503456 |
| 6.03867E-09 | 1 |
| 3.75876E-07 | 0.000149844 |
| 0.000436096 | 9.81601E-07 |
| 7.21575E-08 | 0.141349218 |

| P-value: (CM_neg_newanalysis_12022024) / (SM_neg_r | P-value: (Bb_neg_newanalysis_12022024) / (WAT |
| --- | --- |
| 0.579910706 | 0.198986235 |
| 0.707838209 | 0.356601204 |
| 0.404499993 | 0.99999637 |
| 0.487602381 | 3.4953E-05 |
| 0.748921698 | 0.000274471 |
| 0.963034864 | 0.188531745 |
| 0.221108812 | 0.029804971 |
| 0.084391303 | 0.91628519 |
| 0.283312199 | 0.806480168 |
| 0.248316942 | 0.296458941 |
| 0.875622586 | 0.480726819 |
| 0.975493052 | 0.260527052 |
| 0.696607376 | 0.966800219 |
| 0.988164118 | 0.790372209 |
| 0.177305454 | 0.050757319 |
| 0.999999999 | 0.158413236 |
| 0.719368102 | 0.356518926 |
| 0.998851532 | 0.116137228 |
| 0.126576092 | 3.99437E-10 |
| 0.579033216 | 0.5839643 |
| 0.229603771 | 0.701101347 |
| 0.785176352 | 0.110259779 |
| 0.545412828 | 0.952943233 |
| 0.99999827 | 0.679795596 |
| 0.055041961 | 0.500548017 |
| 0.000922898 | 0.399860641 |
| 0.103950049 | 0.086948523 |
| 0.004247382 | 0.430205344 |
| 0.73102677 | 0.542835376 |
| 0.0068761 | 0.661783292 |
| 0.056469856 | 0.999861835 |
| 0.007541659 | 0.380101252 |
| 0.990305967 | 0.356313389 |
| 0.93935273 | 4.07391E-09 |
| 0.999995796 | 0.00073543 |
| 0.996045917 | 1.49689E-07 |
| 0.968562308 | 6.58319E-09 |
| 0.876841697 | 8.85427E-06 |
| 0.998995031 | 1.3356E-13 |
| 0.996737506 | 1.93828E-09 |
| 0.981509659 | 0.138276795 |
| 0.806738338 | 1.39555E-13 |
| 0.675901672 | 6.71685E-14 |
| 0.977740946 | 1.20831E-07 |
| 0.917268168 | 6.44E-08 |
| 0.336897563 | 0.922017566 |
| 0.626605426 | 0.15126356 |
| 0.970710782 | 0.290259018 |
| 1 | 0.184258083 |
| 0.727866533 | 0.522692479 |
| 0.996449906 | 0.999972838 |
| 0.024561773 | 0.99999386 |
| 0.568081141 | 0.582430865 |
| 0.084516491 | 0.999986219 |
| 0.818565678 | 0.992390563 |
| 0.999999751 | 0.005325537 |
| 0.933609137 | 0.042649122 |
| 0.042959271 | 0.762600099 |
| 0.79215804 | 0.34198429 |
| 0.992594283 | 0.851691436 |
| 0.004358073 | 0.642752555 |
| 0.048268143 | 0.19996139 |
| 0.530686892 | 0.130061082 |
| 0.575019097 | 0.250516423 |
| 0.814433458 | 0.154795415 |
| 0.999998198 | 3.76834E-05 |
| 0.998824688 | 0.993776934 |
| 0.143922432 | 2.78033E-05 |
| 0.950890379 | 0.438532235 |
| 0.27821825 | 0.999980351 |
| 0.002992717 | 0.627623599 |
| 0.091152274 | 0.0177895 |
| 0.875322438 | 0.094290769 |
| 0.999993461 | 0.847923862 |
| 0.960265782 | 0.925571024 |
| 0.850876485 | 0.772200838 |
| 0.979684008 | 0.993444562 |
| 0.876222797 | 0.993986131 |
| 0.545150026 | 0.446138649 |
| 0.626442762 | 0.999994839 |
| 0.789197554 | 0.856460412 |
| 0.129202782 | 0.479086598 |
| 0.72839933 | 0.999985664 |
| 0.303033524 | 0.999954432 |
| 0.923898899 | 0.982998617 |
| 0.095768322 | 0.368668471 |
| 0.686128299 | 0.847100782 |
| 0.989100843 | 0.001481099 |
| 0.950569091 | 0.150352469 |
| 0.770691409 | 0.48254028 |
| 0.72567856 | 0.313374521 |
| 0.423628652 | 0.23739587 |
| 0.594623597 | 0.543819655 |
| 0.969438908 | 0.561965275 |
| 0.776639404 | 0.593995072 |
| 0.951073379 | 0.210260729 |
| 0.86567168 | 0.139423596 |
| 0.849322071 | 0.356663674 |
| 0.751357508 | 6.05406E-07 |
| 0.016503136 | 0.999996249 |
| 0.782484815 | 0.995849948 |
| 0.985894221 | 0.961829225 |
| 0.568762024 | 0.13554115 |
| 0.999988858 | 0.708821921 |
| 0.023680166 | 0.161112981 |
| 0.846067885 | 0.974157941 |
| 0.994802282 | 0.158860696 |
| 0.999937394 | 0.548296605 |
| 0.251578568 | 0.390591981 |
| 0.094153958 | 0.322847237 |
| 0.182962805 | 0.461468303 |
| 0.068524255 | 0.921422584 |
| 0.037557104 | 0.998857646 |
| 0.568953904 | 0.327529512 |
| 0.986172645 | 0.530538787 |
| 0.990299801 | 0.999870113 |
| 0.953196555 | 8.37813E-10 |
| 0.993399977 | 4.60743E-14 |
| 0.028199186 | 4.60743E-14 |
| 0.319044915 | 0.858132827 |
| 0.987524484 | 0.673469941 |
| 0.926863353 | 1.42553E-07 |
| 0.525079657 | 4.60743E-14 |
| 1 | 0.66808931 |
| 0.595120606 | 0.115486633 |
| 0.386718444 | 0.882362615 |
| 0.211830285 | 0.048963155 |
| 0.999996859 | 0.9936251 |
| 0.415309354 | 0.925514658 |
| 0.341904334 | 0.916803541 |
| 0.000252522 | 0.28269867 |
| 0.832168596 | 0.068728474 |
| 0.000227232 | 0.286097522 |
| 0.769490192 | 0.999807439 |
| 0.551182215 | 0.787242112 |
| 0.016999445 | 0.287997638 |
| 0.948262022 | 0.065506527 |
| 0.999963759 | 0.993390041 |
| 0.988780039 | 0.112168111 |
| 0.855616877 | 0.998040921 |
| 0.479957714 | 0.400770095 |
| 0.071506667 | 0.442989468 |
| 0.999530262 | 0.041366943 |
| 0.918446311 | 0.970570676 |
| 0.15981906 | 0.515783964 |
| 0.050489453 | 0.295830437 |
| 0.518894265 | 0.274312126 |
| 0.999797945 | 0.008063498 |
| 0.110619559 | 1.9755E-06 |
| 0.351614546 | 0.975345442 |
| 0.140800347 | 0.043272345 |
| 0.99595109 | 0.43609928 |
| 0.265644398 | 0.999971772 |
| 0.095175559 | 0.062444398 |
| 0.859970849 | 0.169471553 |
| 0.847873512 | 0.000908021 |
| 0.380868764 | 0.584065024 |
| 0.9785461 | 0.973156653 |
| 0.960086167 | 0.813688031 |
| 0.999928277 | 0.129897265 |
| 0.951107317 | 0.414087516 |
| 0.716949901 | 0.575060585 |
| 0.288511369 | 0.174308213 |
| 0.980729717 | 0.938441556 |
| 0.96897079 | 0.01994637 |
| 0.416782798 | 0.957456984 |
| 0.739189394 | 0.37670583 |
| 0.934222253 | 0.198016218 |
| 0.245871993 | 0.476610259 |
| 0.974486181 | 0.008315134 |
| 0.715195411 | 0.651355659 |
| 0.226657855 | 0.181209603 |
| 0.012014596 | 0.945571566 |
| 0.971559714 | 0.932508585 |
| 0.453153312 | 0.765861035 |
| 0.868651787 | 0.947273847 |
| 0.054222812 | 0.196174794 |
| 0.924286141 | 0.99662526 |
| 0.933871399 | 0.897315219 |
| 0.330331869 | 0.396620481 |
| 0.376386858 | 0.282023781 |

| P-value: (CM_neg_newanalysis_12022024) / (WATER_NEG_NEWANALYSIS_12022024) | P-value: (RS_NEG_NEWANALYSIS_12022024) / (WATER_NEG_NE |
| --- | --- |
| 0.37024739 | 1.03901E-06 |
| 0.630421151 | 3.32516E-08 |
| 0.99939645 | 2.96887E-05 |
| 0.001954467 | 2.03459E-07 |
| 0.02781414 | 1.10658E-07 |
| 0.888367008 | 0.607758363 |
| 2.46636E-12 | 1.79222E-11 |
| 1.9893E-06 | 3.32896E-06 |
| 0.000160361 | 0.000526326 |
| 9.14842E-10 | 2.52248E-08 |
| 2.02315E-08 | 9.69245E-05 |
| 0.571508065 | 0.999998804 |
| 0.089754991 | 0.071552107 |
| 0.044345354 | 0.001119738 |
| 0.002481696 | 0.000108636 |
| 0.299810405 | 0.001834057 |
| 0.193858869 | 0.807041003 |
| 0.783608071 | 9.07337E-08 |
| 0.970175157 | 0.000111635 |
| 0.99983626 | 0.058702982 |
| 0.999880918 | 0.776333045 |
| 0.018660641 | 1.17626E-07 |
| 0.888173652 | 0.037660858 |
| 0.038425125 | 5.54362E-06 |
| 0.994667643 | 0.999999528 |
| 0.000577386 | 0.000283518 |
| 0.000135441 | 5.74411E-05 |
| 0.011635205 | 0.017675914 |
| 0.132526693 | 8.1735E-05 |
| 0.992109814 | 0.999887996 |
| 0.018481733 | 0.013955054 |
| 0.12289431 | 0.228017152 |
| 0.001353861 | 3.00948E-12 |
| 0.733379876 | 0.986337091 |
| 0.828871909 | 0.999995212 |
| 0.999997738 | 0.247713511 |
| 0.976243751 | 0.999998779 |
| 0.078423862 | 1.15336E-05 |
| 0.933782145 | 0.043326484 |
| 0.828980248 | 0.013943402 |
| 0.678831324 | 0.098257629 |
| 0.311531331 | 1.96218E-05 |
| 0.99877277 | 0.004069459 |
| 0.198727657 | 0.000996326 |
| 0.136568371 | 0.000283022 |
| 0.878008845 | 1.14106E-07 |
| 0.332650483 | 0.9826782 |
| 0.050517678 | 0.013564906 |
| 0.072865645 | 0.109848994 |
| 3.00903E-06 | 1.2389E-12 |
| 0.553943668 | 0.489223036 |
| 0.999999938 | 0.946814815 |
| 0.985485725 | 0.010374944 |
| 0.984531771 | 1.23213E-10 |
| 0.913869226 | 0.00013131 |
| 0.999708557 | 0.303504045 |
| 0.004248339 | 1.84733E-06 |
| 0.99927585 | 2.28092E-07 |
| 0.039042088 | 2.52921E-06 |
| 0.68761357 | 2.93377E-08 |
| 6.23899E-07 | 0.00237518 |
| 0.076607689 | 6.03286E-10 |
| 0.266519994 | 0.002318798 |
| 0.195679188 | 0.627471728 |
| 3.21353E-08 | 1.03956E-11 |
| 0.009847291 | 5.05064E-09 |
| 0.6061542 | 0.082212355 |
| 0.547538311 | 0.452545941 |
| 0.042064206 | 0.618941298 |
| 0.896088357 | 0.218465081 |
| 0.719514495 | 2.18759E-06 |
| 0.339082446 | 0.125445892 |
| 0.032650822 | 0.999984118 |
| 0.935889733 | 0.401617476 |
| 0.884534296 | 0.011149218 |
| 0.717238027 | 0.000156015 |
| 0.492879904 | 0.74758734 |
| 0.997313431 | 0.007704095 |
| 0.247759753 | 0.999449134 |
| 0.012948206 | 2.53024E-08 |
| 2.41095E-05 | 2.96866E-09 |
| 0.098383983 | 1.26955E-10 |
| 0.999983604 | 1.13086E-09 |
| 0.740841076 | 0.994121264 |
| 0.898595974 | 0.323225193 |
| 0.768397413 | 0.000561149 |
| 0.067750331 | 5.5323E-06 |
| 0.563970509 | 0.99994849 |
| 5.12536E-10 | 1.94844E-13 |
| 2.36281E-05 | 1.72978E-10 |
| 0.069297734 | 0.992414194 |
| 0.07707358 | 6.69427E-07 |
| 0.659300544 | 0.000329061 |
| 0.159235293 | 0.283570352 |
| 0.014670799 | 1.16276E-06 |
| 0.036498078 | 8.99475E-06 |
| 0.575268277 | 0.281436165 |
| 0.998463133 | 0.002599831 |
| 9.53926E-05 | 1.55936E-09 |
| 0.999734128 | 0.982213917 |
| 0.999971067 | 0.6126808 |
| 0.771451906 | 0.122472107 |
| 0.029885559 | 1.24231E-06 |
| 0.533258319 | 0.886702906 |
| 0.201388503 | 0.89761948 |
| 0.311319133 | 0.9994822 |
| 0.094653762 | 0.000351477 |
| 0.310398218 | 0.926424078 |
| 2.72881E-09 | 4.87388E-14 |
| 0.725342724 | 7.02771E-14 |
| 3.97977E-05 | 6.13953E-14 |
| 5.07494E-09 | 9.73333E-13 |
| 4.20569E-05 | 1.49166E-11 |
| 0.980109807 | 5.71765E-14 |
| 0.447988383 | 0.000407158 |
| 0.991856516 | 0.000405333 |
| 0.107226507 | 7.46922E-05 |
| 0.214508611 | 0.000998975 |
| 0.561911821 | 0.007653239 |
| 0.165000108 | 0.2640655 |
| 0.549763236 | 0.994226287 |
| 0.093839725 | 0.001538551 |
| 0.850166001 | 0.419705699 |
| 0.488999785 | 0.001129108 |
| 0.350468881 | 5.25E-06 |
| 0.997648262 | 0.560539196 |
| 0.171004382 | 0.999966416 |
| 0.415351102 | 0.002488708 |
| 0.997961905 | 0.030545565 |
| 0.997127924 | 0.496756961 |
| 0.000564634 | 5.20695E-14 |
| 0.423881079 | 8.70831E-05 |
| 1.98527E-07 | 4.69624E-14 |
| 0.013352235 | 4.61493E-09 |
| 0.000248928 | 8.21923E-10 |
| 0.029669214 | 4.01814E-08 |
| 0.001647149 | 4.01381E-10 |
| 0.718027801 | 0.999990712 |
| 0.999999861 | 0.16764027 |
| 0.098790012 | 0.004133184 |
| 0.999903566 | 0.018904079 |
| 0.996340322 | 0.285526543 |
| 0.550551414 | 6.90511E-07 |
| 0.001373837 | 8.54317E-13 |
| 0.204003098 | 0.000126695 |
| 0.97953537 | 9.29771E-11 |
| 0.014128075 | 3.30513E-13 |
| 1 | 4.31938E-08 |
| 1.06762E-10 | 4.60743E-14 |
| 0.993486646 | 0.337178856 |
| 0.948731311 | 1.34917E-09 |
| 0.920864903 | 0.000691747 |
| 1.64405E-06 | 6.69464E-14 |
| 9.68136E-10 | 4.62963E-14 |
| 0.955106626 | 0.488126677 |
| 0.002338828 | 0.671932482 |
| 0.634532376 | 0.000252332 |
| 0.035449297 | 5.95987E-08 |
| 0.693204875 | 0.999818535 |
| 0.993089066 | 6.6975E-06 |
| 0.99487795 | 0.085916538 |
| 0.008550682 | 1.14123E-09 |
| 0.002902598 | 4.87388E-14 |
| 0.788482295 | 1.57349E-08 |
| 1.53969E-07 | 4.94049E-14 |
| 0.999420978 | 0.703747193 |
| 0.060812553 | 0.004707089 |
| 1.10471E-05 | 4.62963E-14 |
| 2.16105E-09 | 4.60743E-14 |
| 5.87812E-09 | 4.78506E-14 |
| 1.0425E-13 | 4.60743E-14 |
| 0.947647726 | 2.91211E-13 |
| 0.071389583 | 0.017719819 |
| 5.53876E-06 | 1.49269E-12 |
| 2.28297E-05 | 5.60663E-14 |
| 0.003722185 | 1.34448E-13 |
| 0.99294296 | 1.95831E-11 |
| 0.996625795 | 2.03822E-08 |
| 0.028839153 | 1.65047E-11 |
| 0.045756892 | 6.28697E-10 |
| 0.096825419 | 0.023283173 |

|  |  |
| --- | --- |
| P-value: (SM_neg_newanalysis_12022024) / (WATER_NEG_ |  |
|  | 0.012085947 |
|  | 0.056029499 |
|  | 0.606824443 |
|  | 1.40312E-05 |
|  | 0.000803873 |
|  | 0.999819076 |
|  | 7.78266E-14 |
|  | 1.57373E-09 |
|  | 4.29105E-07 |
|  | 6.58795E-12 |
|  | 1.37329E-09 |
|  | 0.93941126 |
|  | 0.769791375 |
|  | 0.165900825 |
|  | 0.477319499 |
|  | 0.289249264 |
|  | 0.007933212 |
|  | 0.555523918 |
|  | 0.022588252 |
|  | 0.734733158 |
|  | 0.339381916 |
|  | 0.000613404 |
|  | 0.98849014 |
|  | 0.030229389 |
|  | 0.015788933 |
|  | 2.89617E-09 |
|  | 8.93165E-08 |
|  | 2.05074E-07 |
|  | 0.004970232 |
|  | 0.001502199 |
|  | 5.62029E-06 |
|  | 5.78272E-06 |
|  | 0.000261159 |

|  |
| --- |
| 0.223739397 |
| 0.882505382 |
| 0.990148871 |
| 0.658261293 |
| 0.005478112 |
| 0.779235832 |
| 0.553026642 |
| 0.280029087 |
| 0.023696918 |
| 0.87776844 |
| 0.044538328 |
| 0.014477416 |
| 0.038630725 |
| 0.012184141 |
| 0.00775287 |
| 0.071170784 |
| 7.40805E-08 |
| 0.283879055 |
| 0.021636968 |
| 0.220034252 |
| 0.01820996 |
| 0.253554322 |
| 0.999924902 |
| 0.000334059 |
| 0.018827484 |
| 0.001475759 |
| 0.943202256 |
| 0.031908234 |
| 2.51023E-05 |
| 0.005754485 |
| 0.004356029 |
| 1.54427E-09 |
| 0.012777345 |
| 0.376540515 |
| 0.955143787 |
| 0.246797116 |
| 0.867547305 |
| 6.18814E-05 |
| 0.978417185 |
| 0.001936554 |
| 0.892428678 |
| 0.999833091 |
| 0.134093792 |
| 0.160146282 |
| 0.631202687 |
| 0.005460563 |
| 0.000193777 |
| 7.20123E-07 |
| 0.000124939 |
| 0.815402656 |
| 0.01686267 |
| 0.357871889 |
| 0.003886478 |
| 0.001725354 |
| 0.233435284 |
| 7.19229E-11 |
| 6.44367E-07 |
| 0.002122883 |
| 0.000630831 |
| 0.04095288 |
| 0.029650661 |
| 0.000449462 |
| 0.00413267 |
| 0.089565357 |
| 0.623597053 |
| 2.26808E-06 |
| 0.032649649 |
| 0.679036985 |
| 0.381340351 |
| 0.000386346 |
| 0.62585504 |
| 0.915422073 |
| 0.028775548 |
| 0.029205856 |
| 0.218424762 |
| 1.74741E-11 |
| 0.003109077 |
| 6.03686E-08 |
| 7.47169E-12 |
| 9.62385E-09 |
| 0.202397686 |
| 0.155311165 |
| 0.844738132 |
| 0.014889722 |
| 0.071769524 |
| 0.593173447 |
| 0.001035942 |
| 0.895058724 |
| 0.009821914 |
| 0.992637391 |
| 0.500393481 |
| 0.011701817 |
| 0.187042059 |
| 0.000557895 |
| 0.483231446 |
| 0.210008234 |
| 0.154211653 |
| 9.29993E-10 |
| 0.981274601 |
| 1.6408E-12 |
| 0.238372138 |
| 2.37294E-06 |
| 2.39889E-06 |
| 0.01638269 |
| 0.603103448 |
| 0.98283767 |
| 0.006391103 |
| 0.349632675 |
| 0.190041129 |
| 0.743749899 |
| 0.018069471 |
| 0.999993826 |
| 0.214004736 |
| 0.00013225 |
| 0.999728328 |
| 4.81726E-13 |
| 0.680530193 |
| 0.019861394 |
| 0.996404393 |
| 5.62188E-09 |
| 2.51899E-12 |
| 0.999688797 |
| 9.4218E-05 |
| 0.015798545 |
| 0.005978109 |
| 0.231961488 |
| 0.99932905 |
| 0.732416072 |
| 0.000186125 |
| 7.71099E-06 |
| 0.372756346 |
| 1.21242E-06 |
| 0.250505345 |
| 0.001927243 |
| 8.69765E-07 |
| 1.38359E-11 |
| 9.88814E-10 |
| 5.09592E-14 |
| 0.036338726 |
| 0.9732553 |
| 7.01898E-07 |
| 1.56687E-07 |
| 0.000174403 |
| 0.172732342 |
| 0.996122471 |
| 0.002652085 |
| 0.00020561 |
| 0.00067285 |

| Adj. P-value: (Bb_neg_newanalysis_12022024)/(CM_neg_newanalysis_12022024) |  |
| --- | --- |
|  | 1 |
|  | 1 |
|  | 1 |
|  | 1 |
|  | 1 |
|  | 1 |
|  | 1 |
|  | 1.40686E-11 |
|  | 0.000674715 |
|  | 0.000130363 |
|  | 8.59889E-10 |
|  | 1.81009E-08 |
|  | 1 |
|  | 1 |
|  | 1 |
|  | 1 |
|  | 0.013387352 |
|  | 1 |
|  | 0.064797216 |
|  | 1.29146E-07 |
|  | 1 |
|  | 1 |
|  | 1 |
|  | 1 |
|  | 1 |
|  | 1 |
|  | 1 |
|  | 7.56922E-05 |
|  | 2.69864E-06 |
|  | 0.0014378 |
|  | 1 |
|  | 1 |
|  | 0.109299501 |
|  | 0.014320181 |
|  | 0.00013383 |

|  |
| --- |
| 1.15184E-08 |
| 0.000579238 |
| 4.07592E-06 |
| 6.45091E-08 |
| 2.61818E-07 |
| 1.45745E-11 |
| 8.85754E-09 |
| 0.052085271 |
| 1.40686E-11 |
| 1.4291E-11 |
| 2.35698E-08 |
| 9.90187E-09 |
| 1 |
| 1 |
| 1 |
| 1 |
| 1.29558E-06 |
| 1 |
| 1 |
| 1 |
| 1 |
| 1 |
| 0.033354426 |
| 2.60028E-05 |
| 1 |
| 0.003318108 |
| 0.781354863 |
| 5.29782E-07 |
| 0.003154026 |
| 0.008803333 |
| 1 |
| 6.69386E-09 |
| 1 |
| 1 |
| 0.038372476 |

[illegible]

|  |
| --- |
| 1 |
| 1 |
| 0.554859454 |
| 1 |
| 1 |
| 2.87846E-09 |
| 0.173544836 |
| 9.16072E-06 |
| 3.04819E-08 |
| 0.000347832 |
| 0.624776408 |
| 1 |
| 1 |
| 3.01505E-10 |
| 1.40686E-11 |
| 1.40686E-11 |
| 1 |
| 1 |
| 1.25484E-08 |
| 1.40686E-11 |
| 1 |
| 1 |
| 1 |
| 1 |
| 1 |
| 1 |
| 1 |
| 4.19343E-05 |
| 0.008811852 |
| 5.41896E-08 |
| 0.238695747 |
| 0.000178655 |
| 0.00191687 |
| 1.73949E-05 |
| 1 |
| 0.670981216 |
| 1 |
| 1 |
| 1 |
| 0.008497581 |
| 0.003067825 |
| 1 |
| 1 |
| 0.000831236 |
| 0.088187715 |
| 1.40686E-11 |
| 1 |
| 0.059841765 |
| 1 |
| 7.39964E-05 |
| 2.21463E-10 |
| 0.248278075 |
| 4.54004E-07 |
| 1 |
| 0.974901073 |
| 1 |
| 0.335037231 |
| 1 |
| 0.001943774 |
| 9.43724E-05 |
| 1 |
| 3.62392E-09 |
| 1 |
| 1 |
| 8.82124E-07 |
| 3.10267E-09 |
| 1.96966E-10 |
| 1.40686E-11 |
| 0.250037412 |
| 0.095097771 |

|  |  |
| --- | --- |
|  | 0.001509411 |
|  | 0.013846576 |
|  | 0.005773065 |
|  | 1 |
|  | 1 |
|  | 0.026884008 |
|  | 0.005068582 |
|  | 1 |

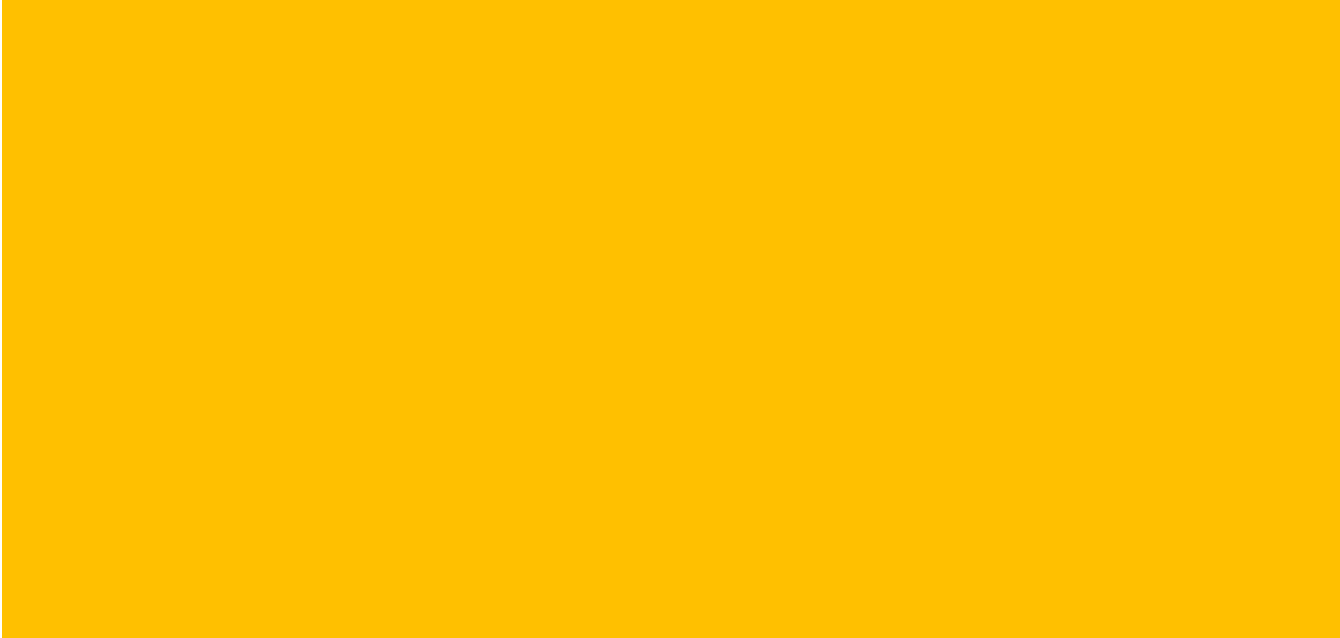

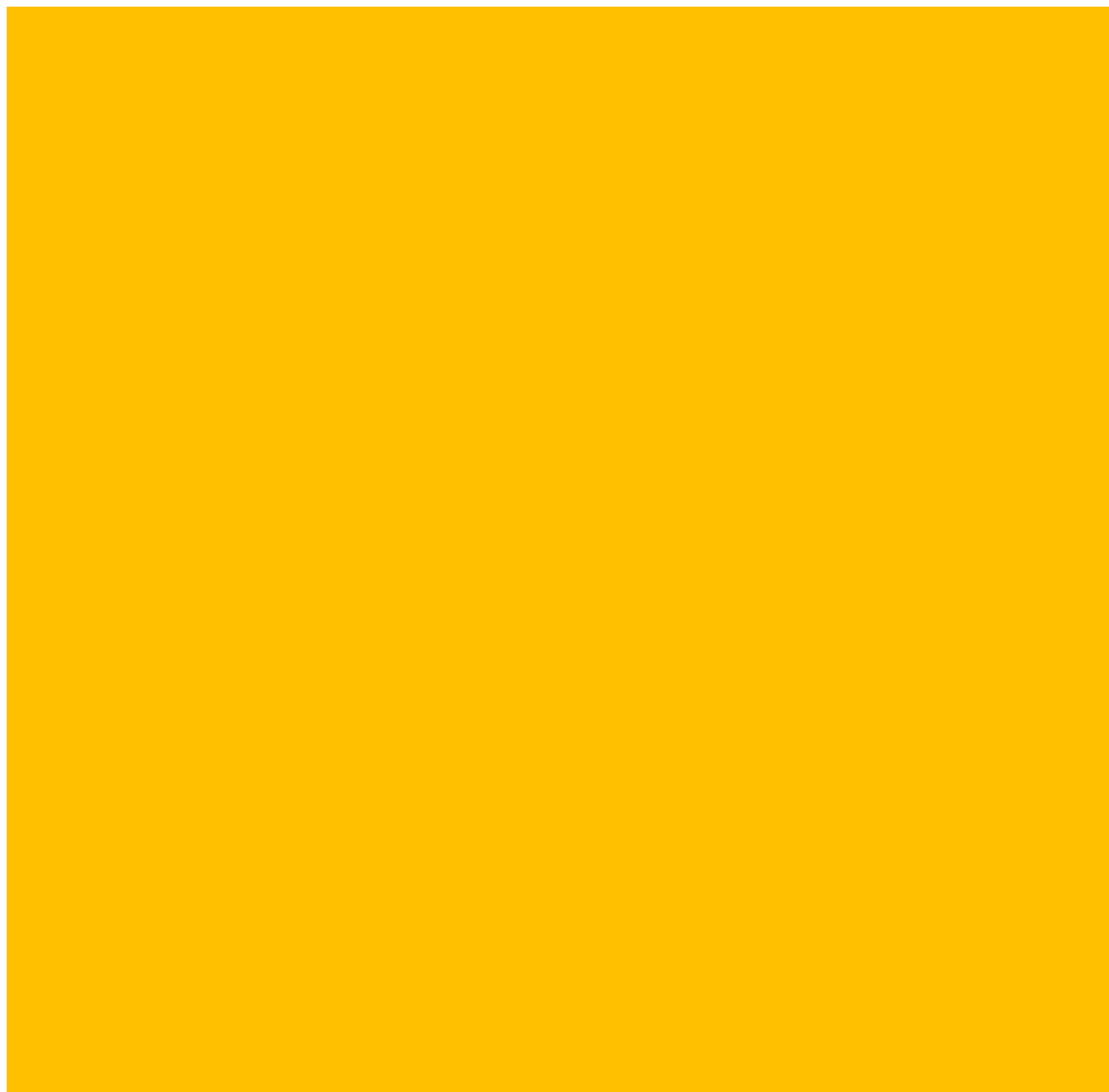

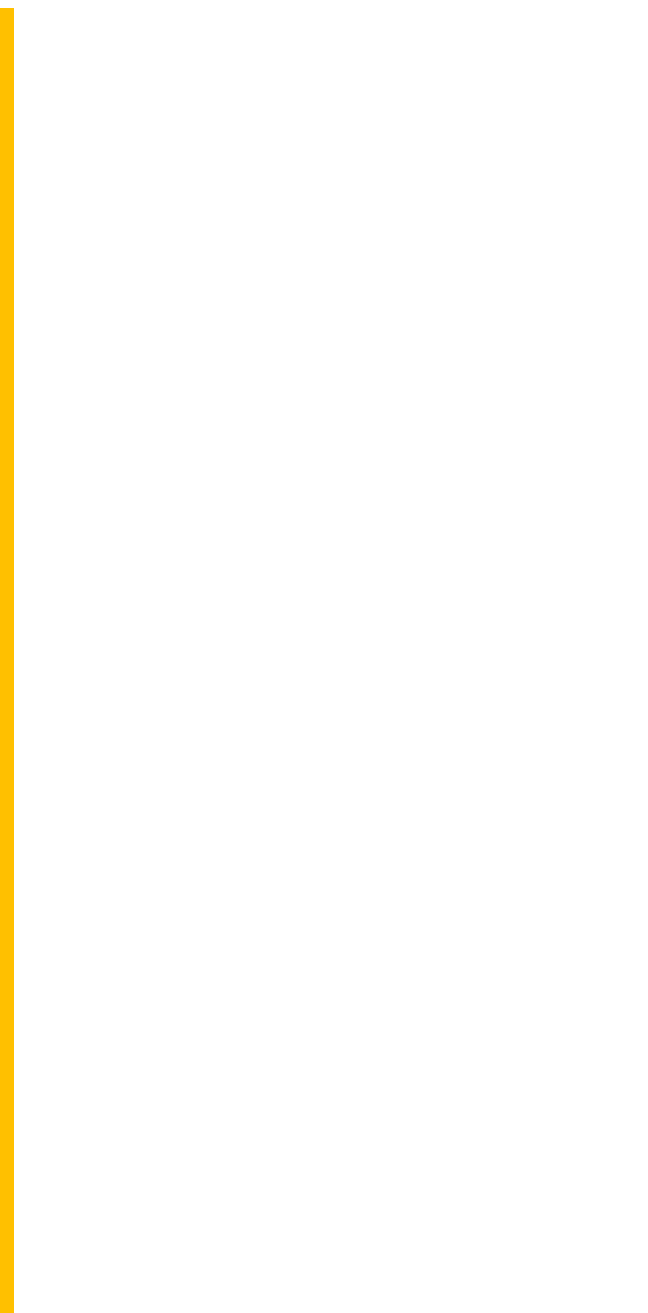

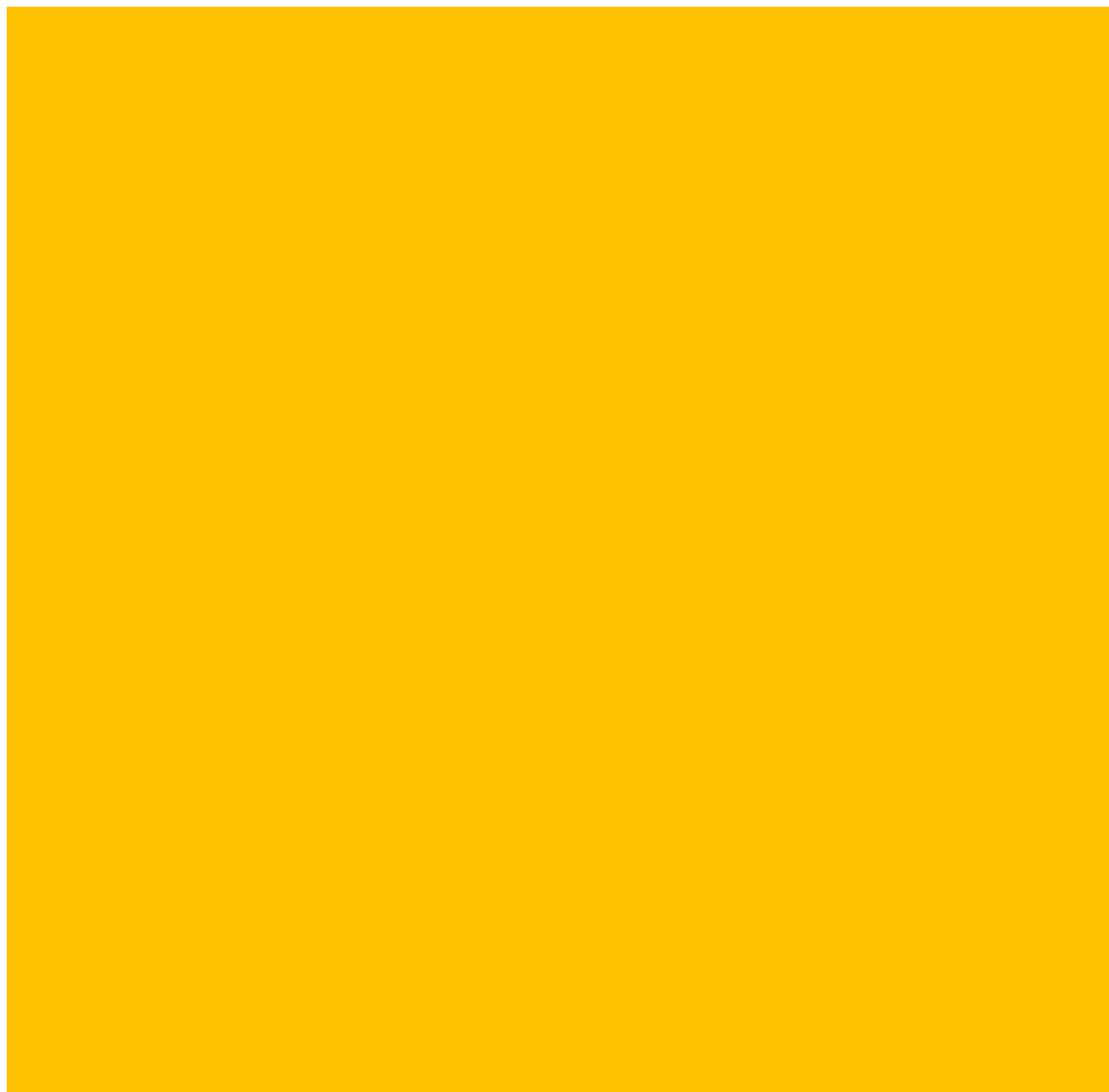

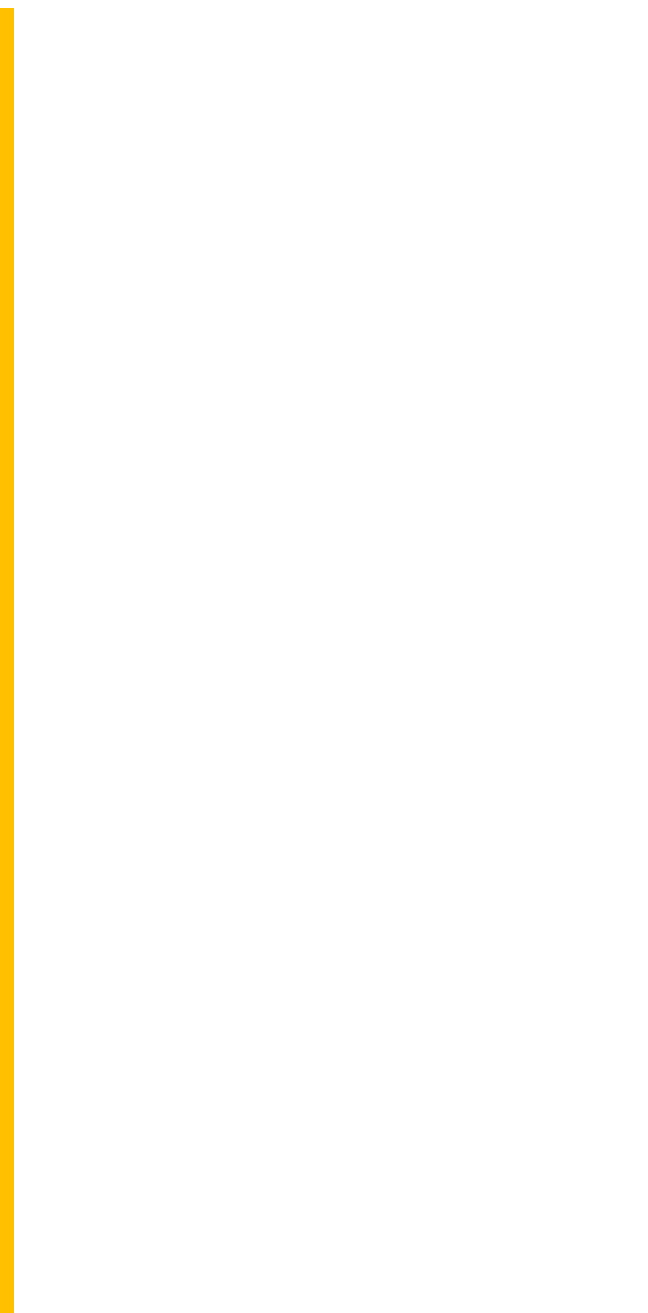

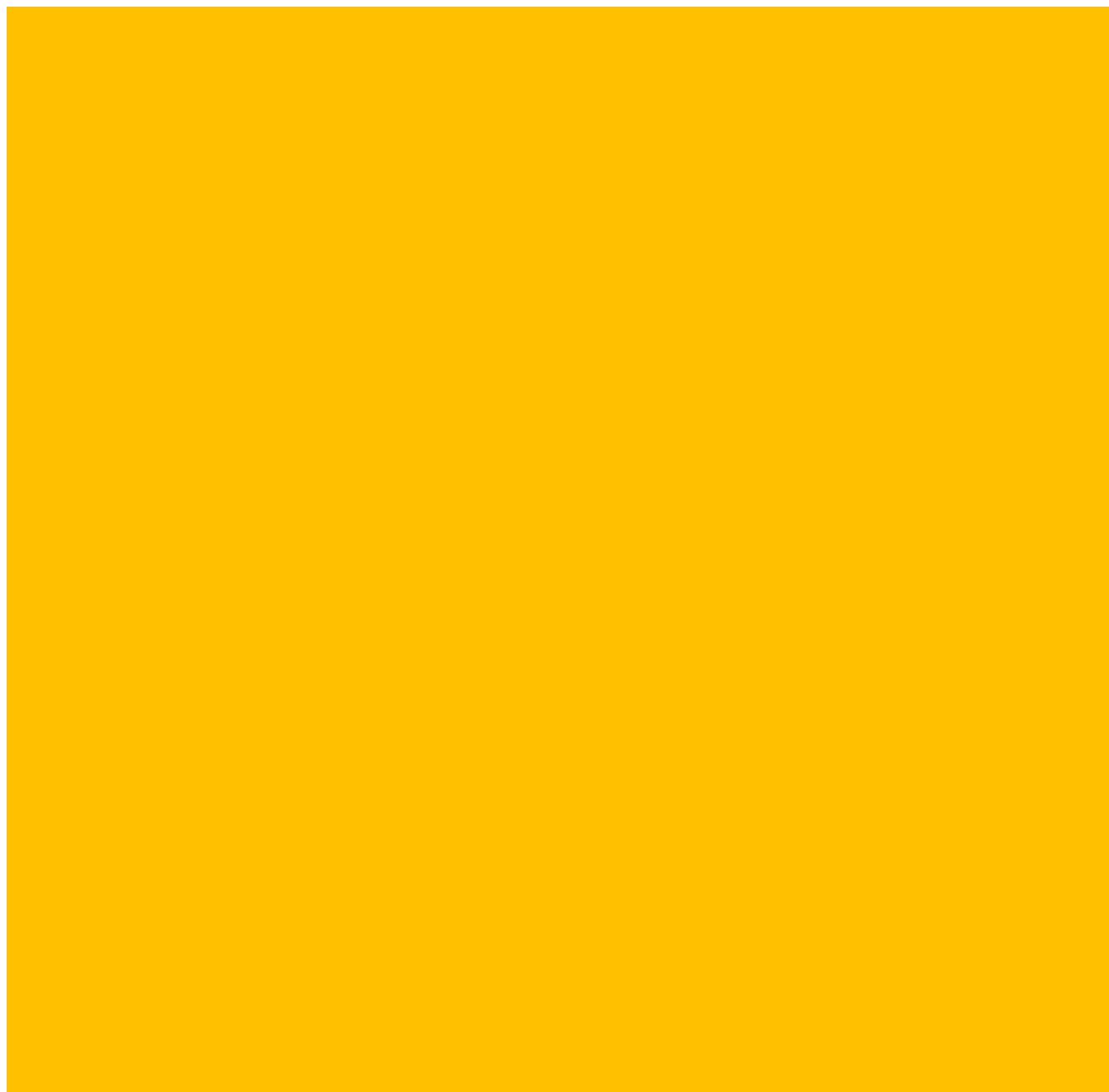

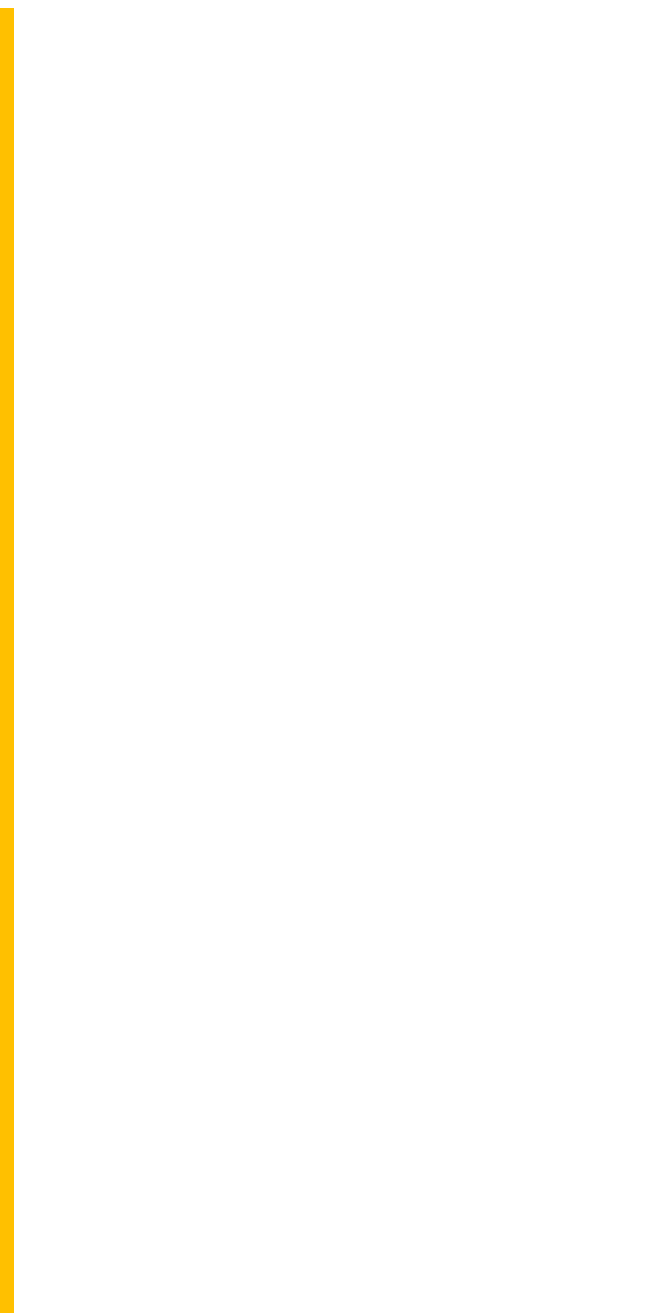

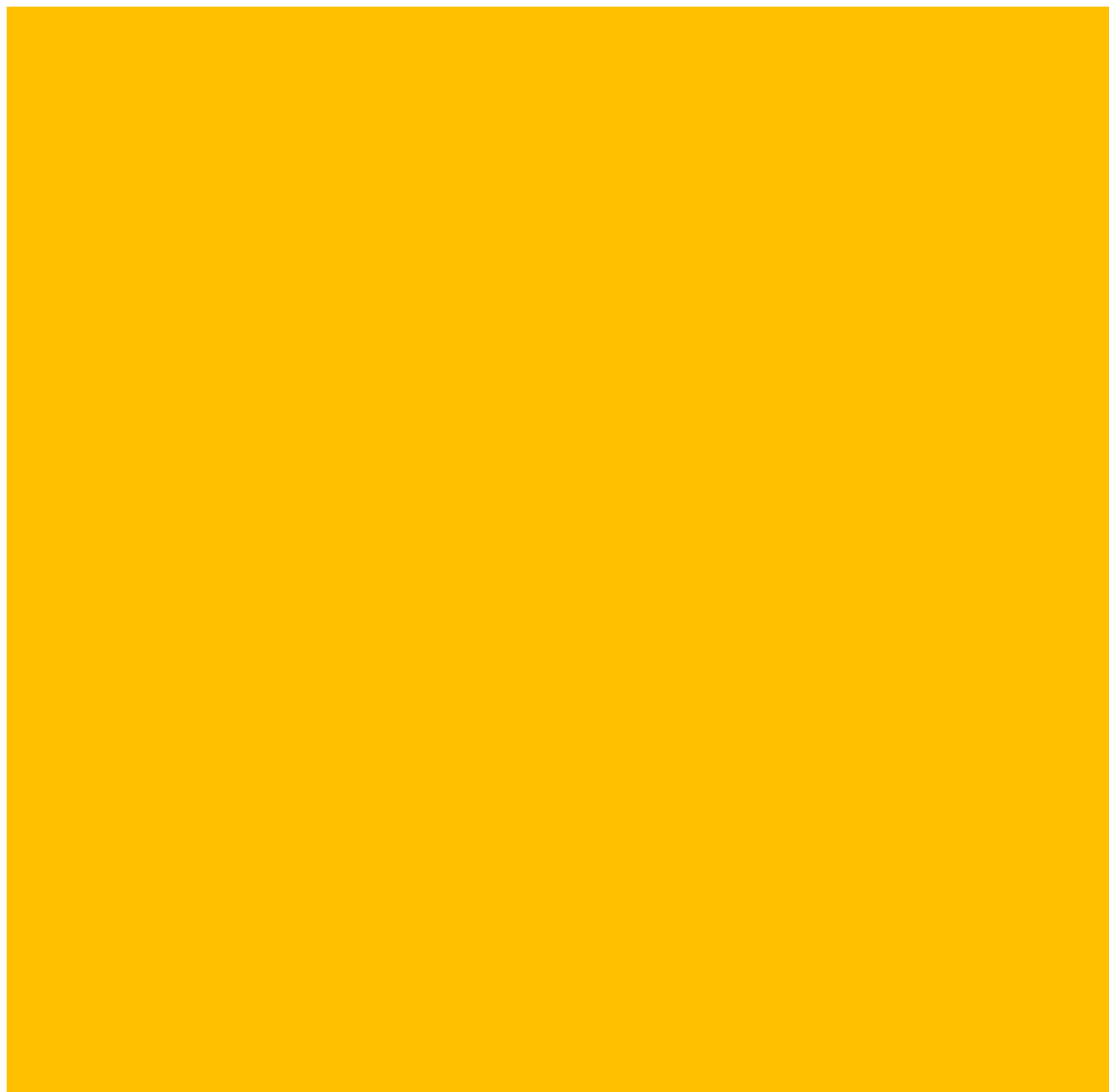

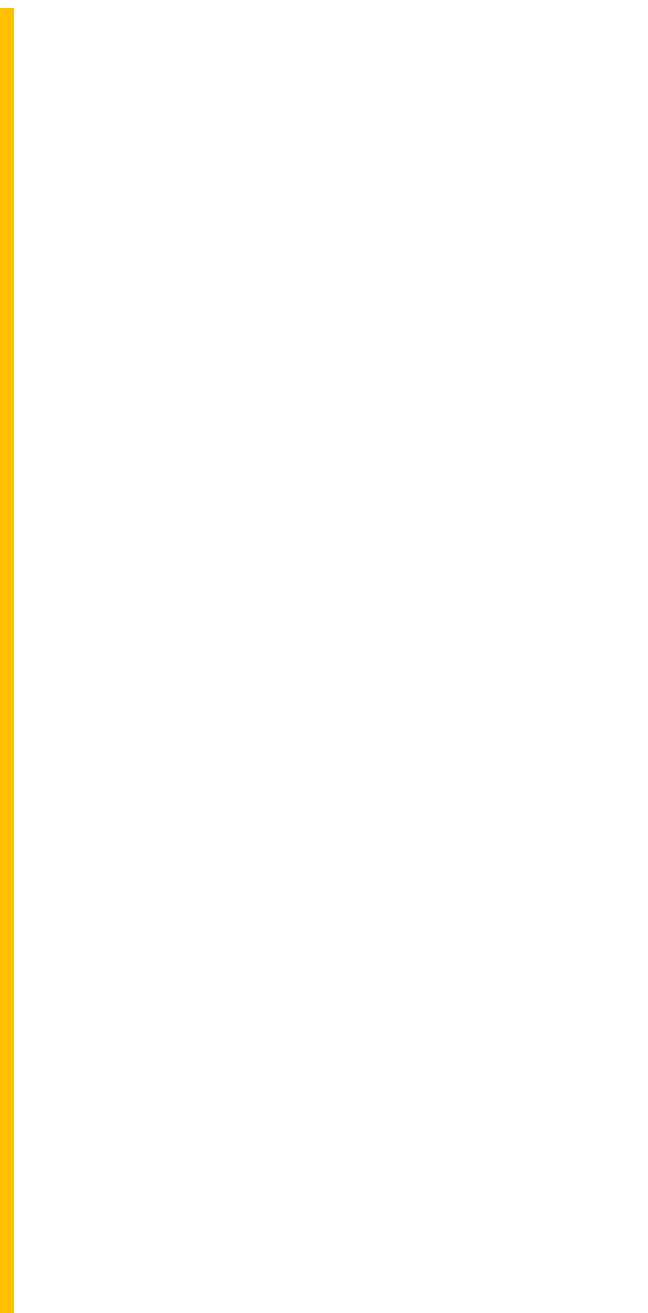

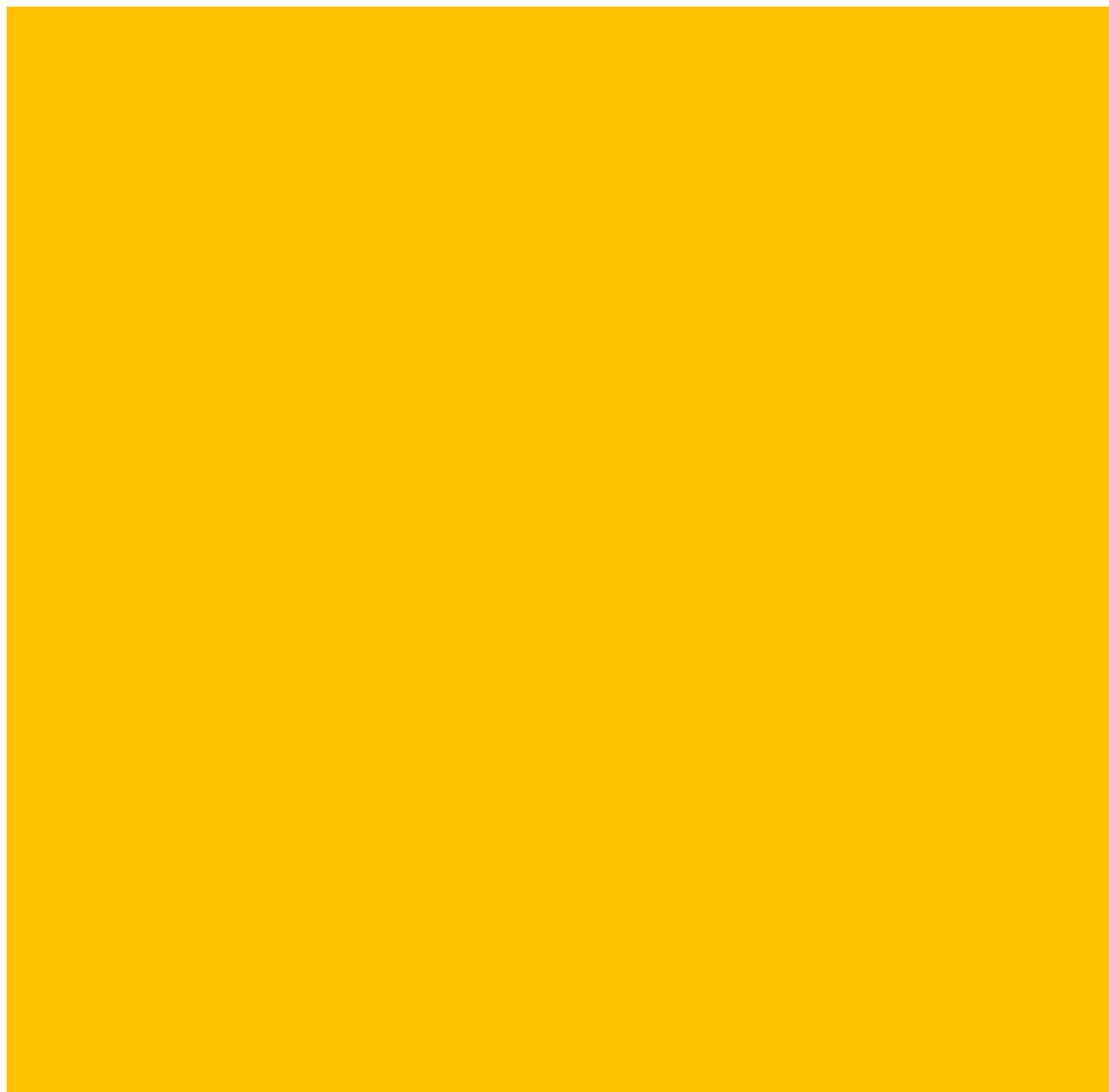

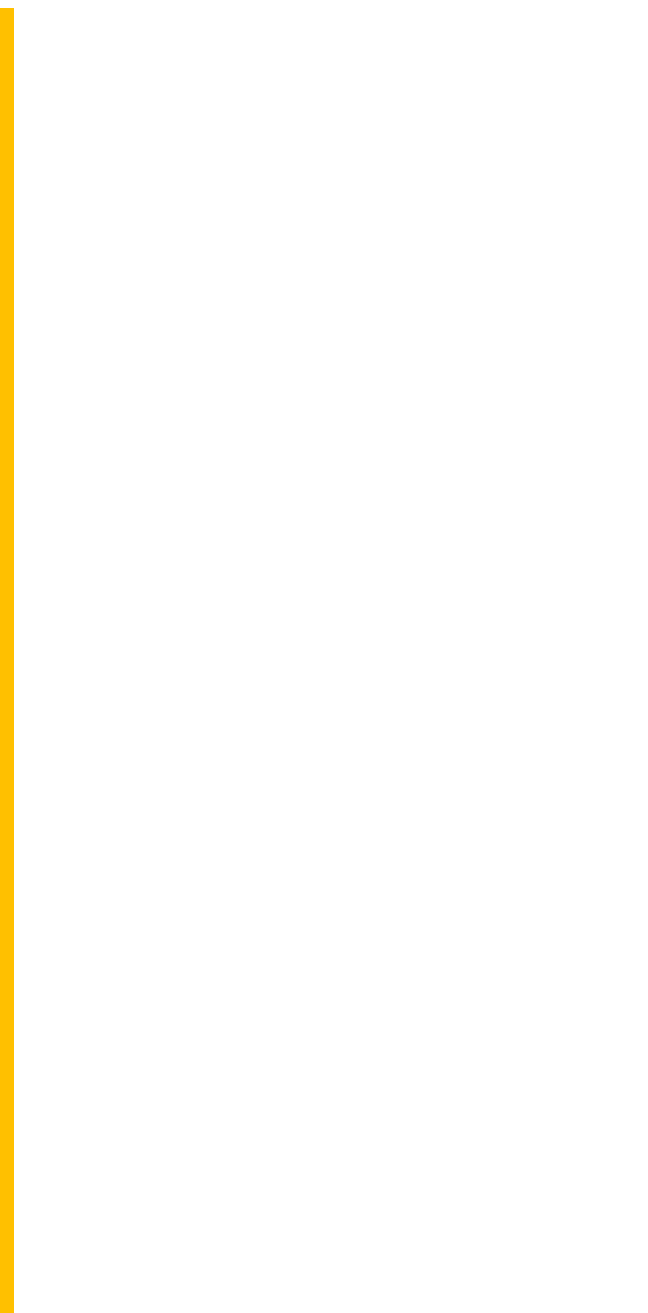

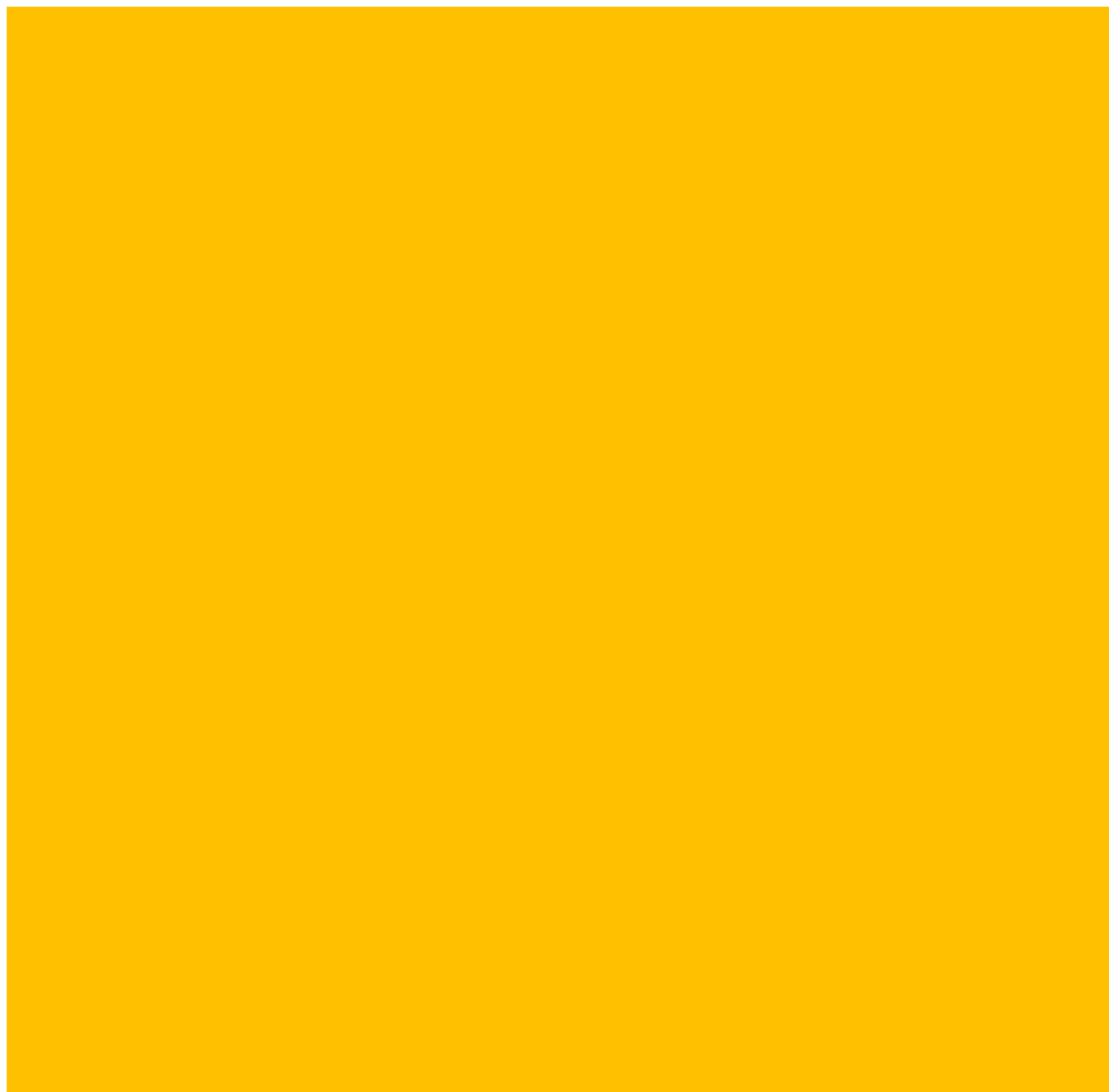

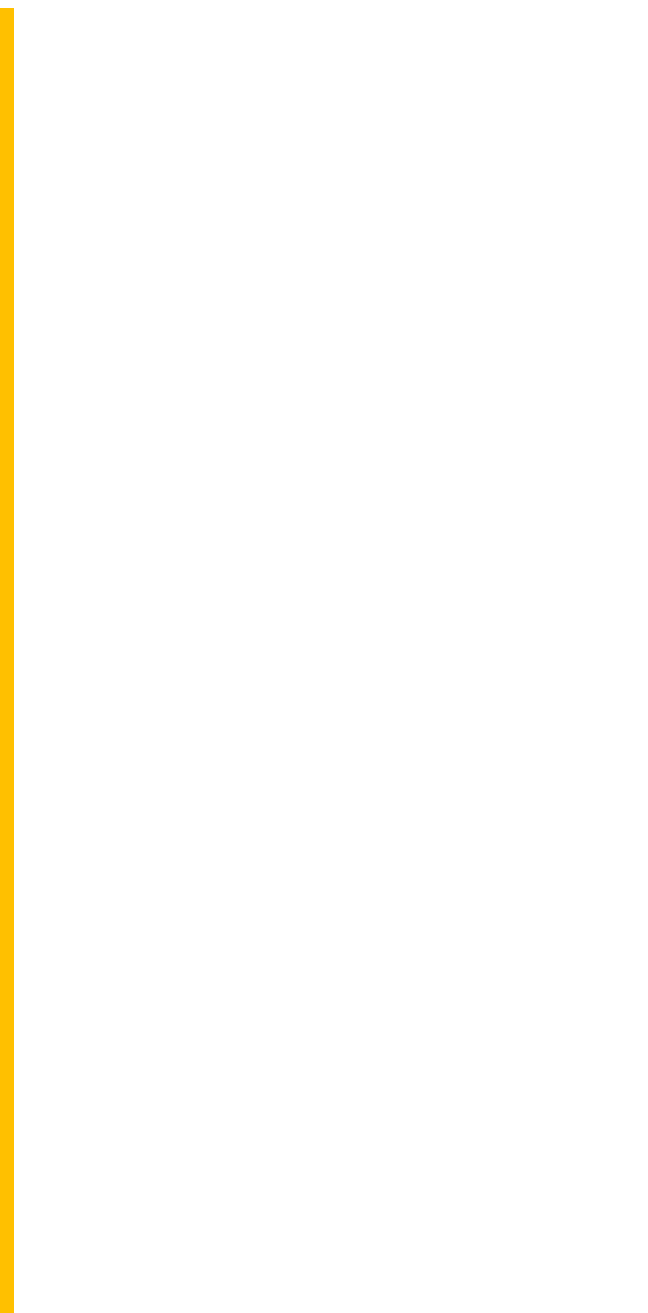

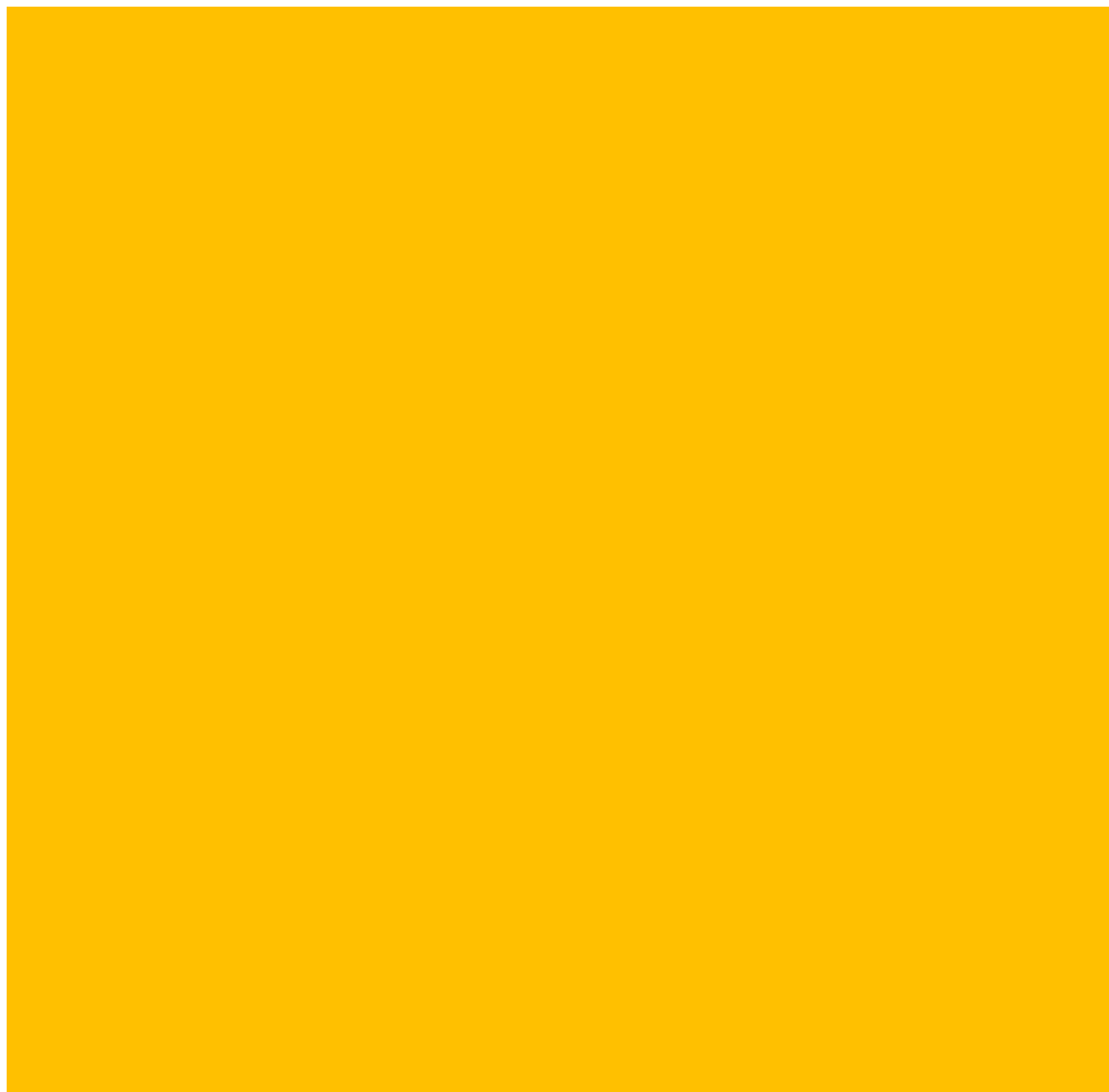

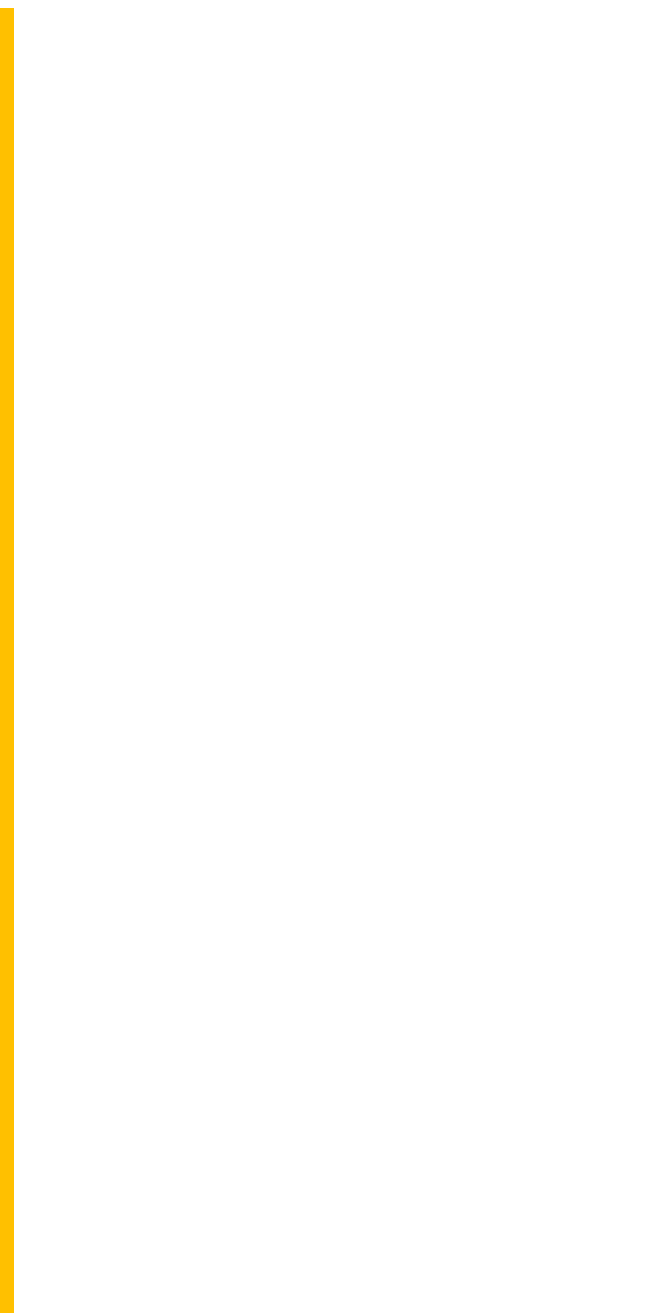

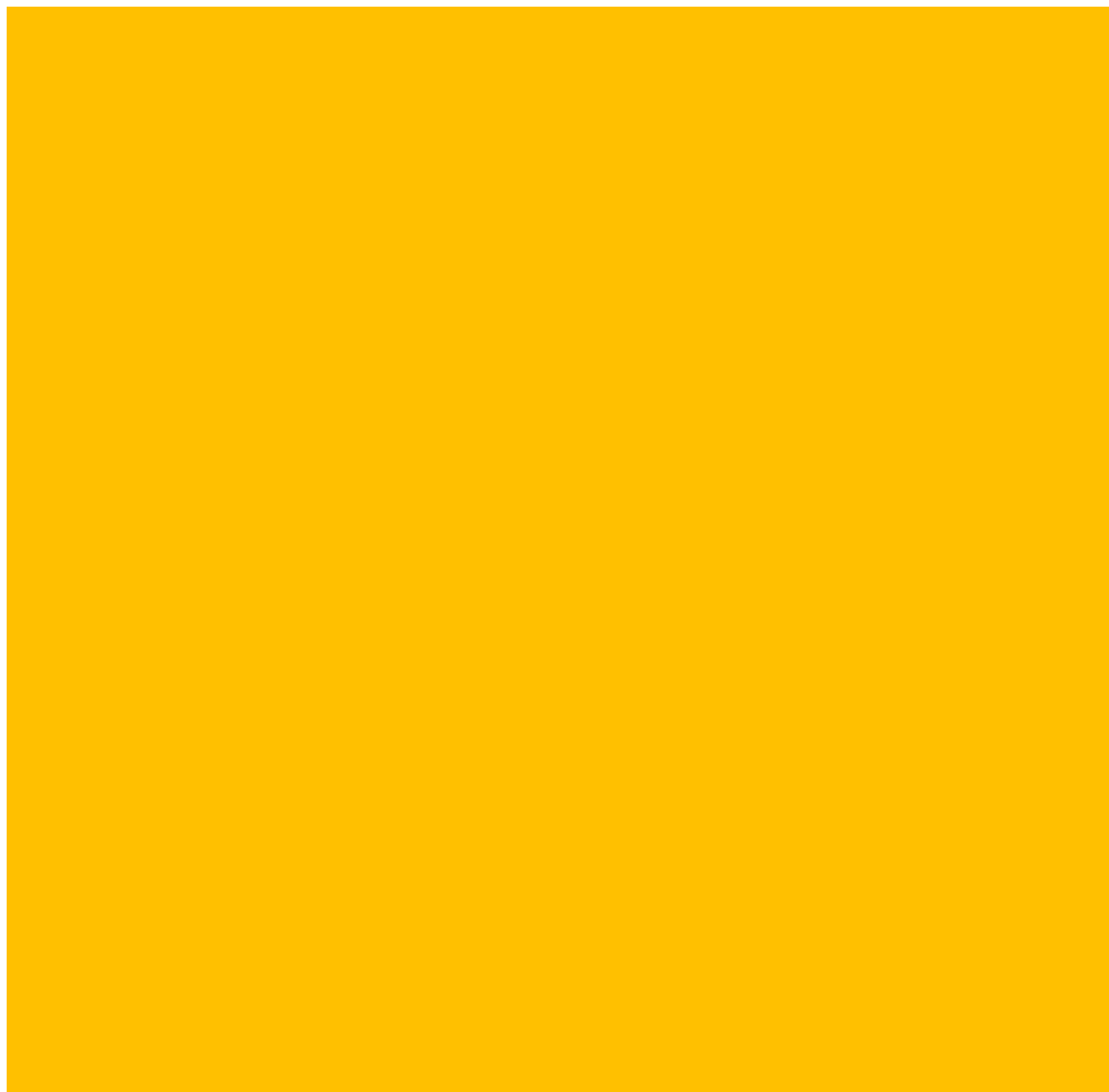

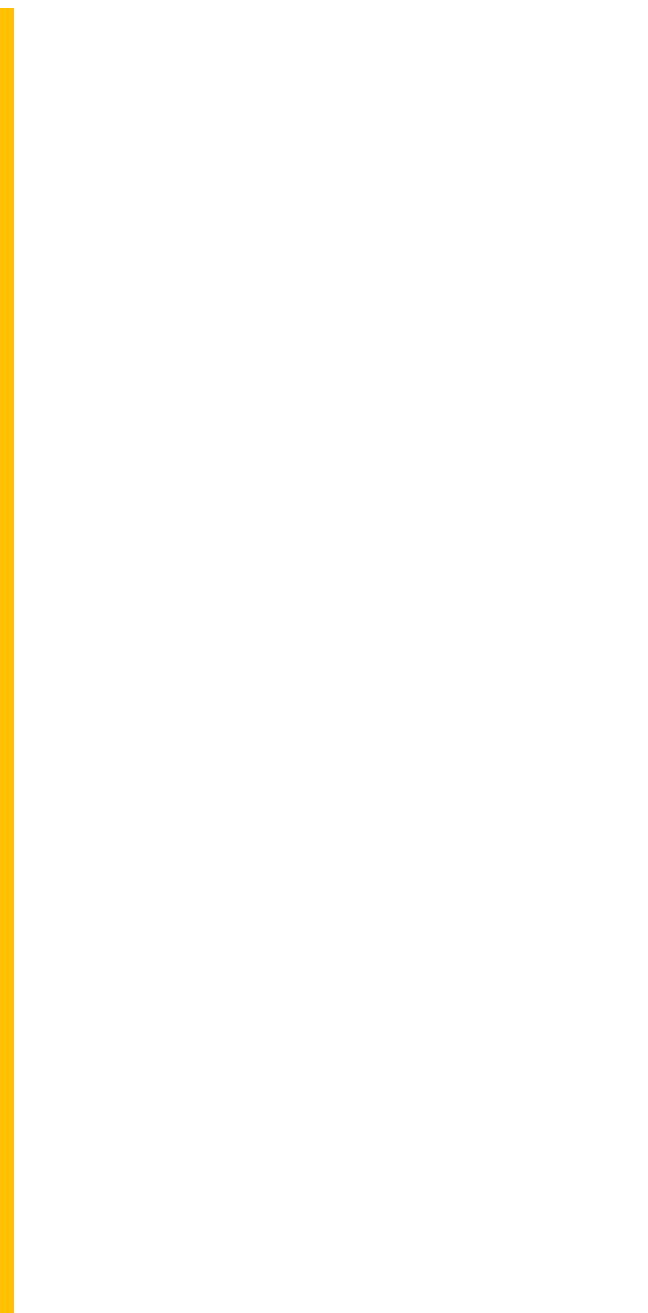

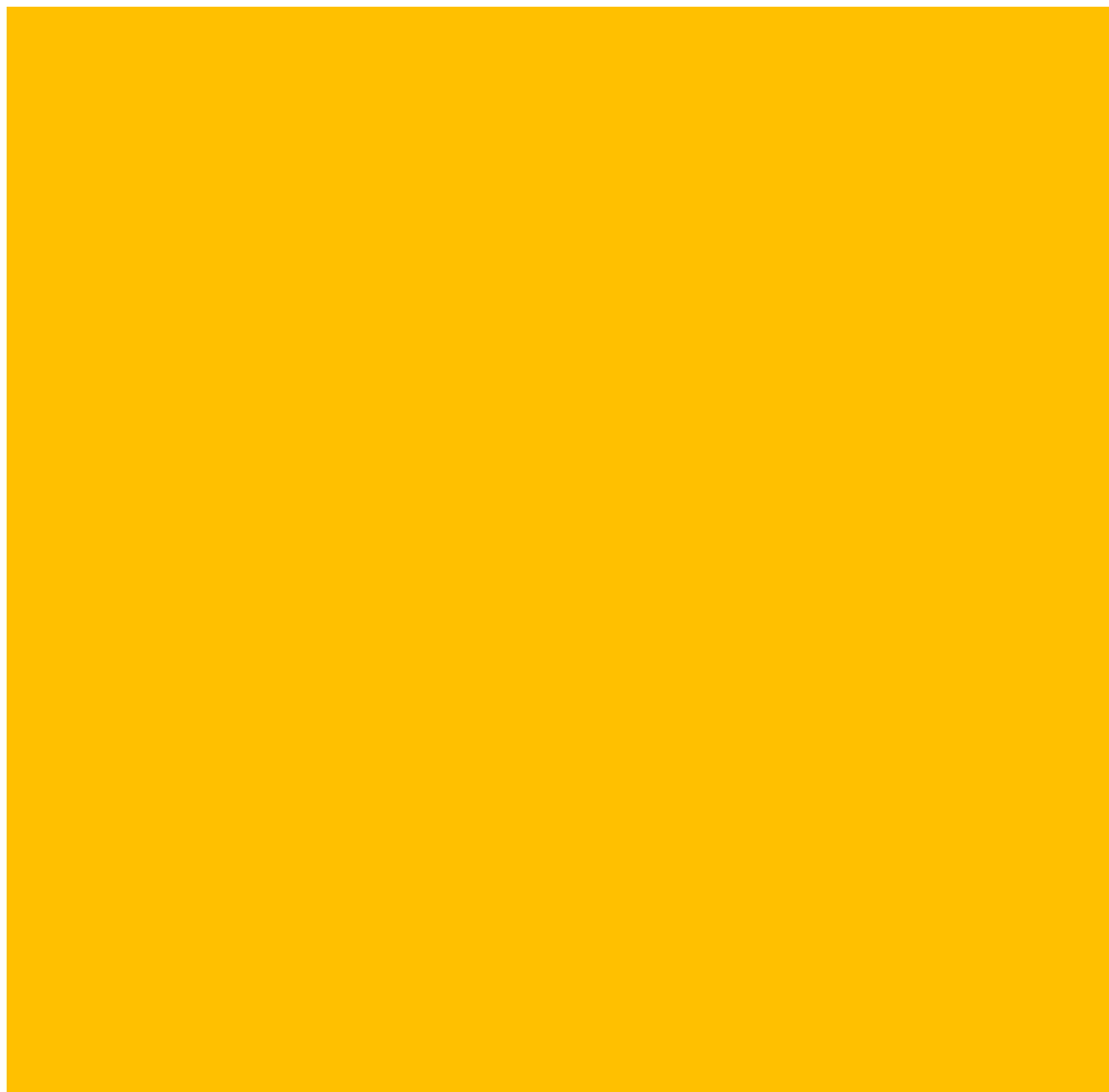

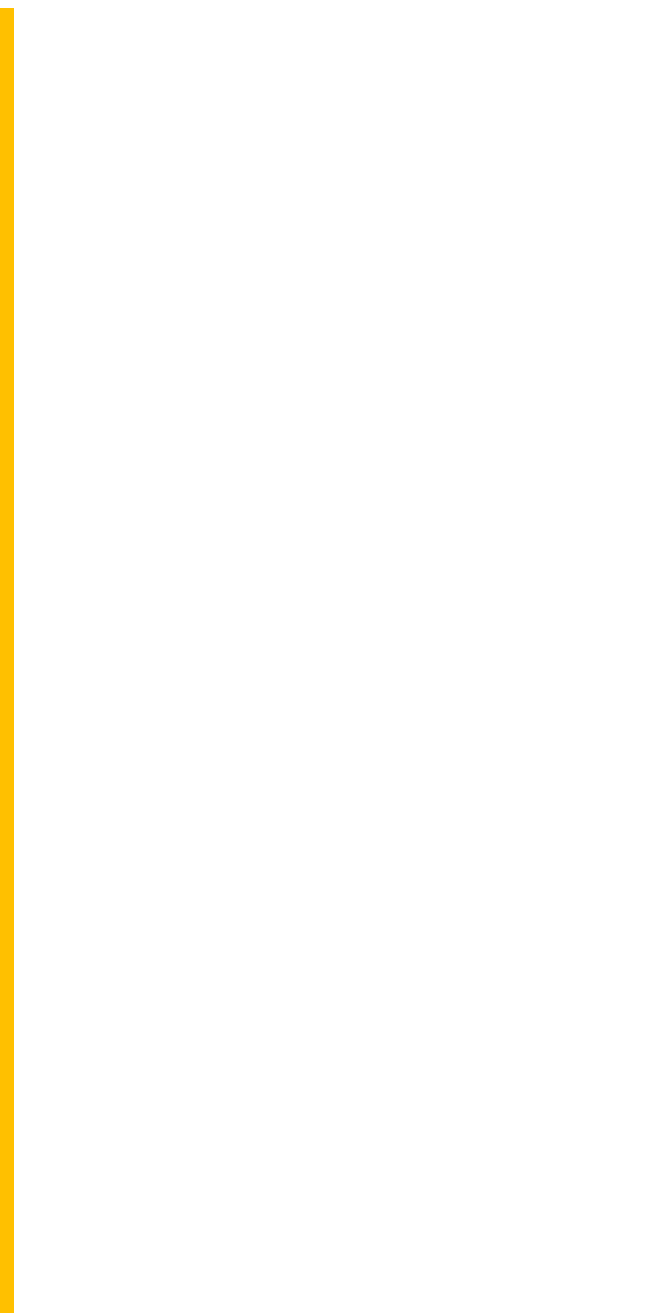

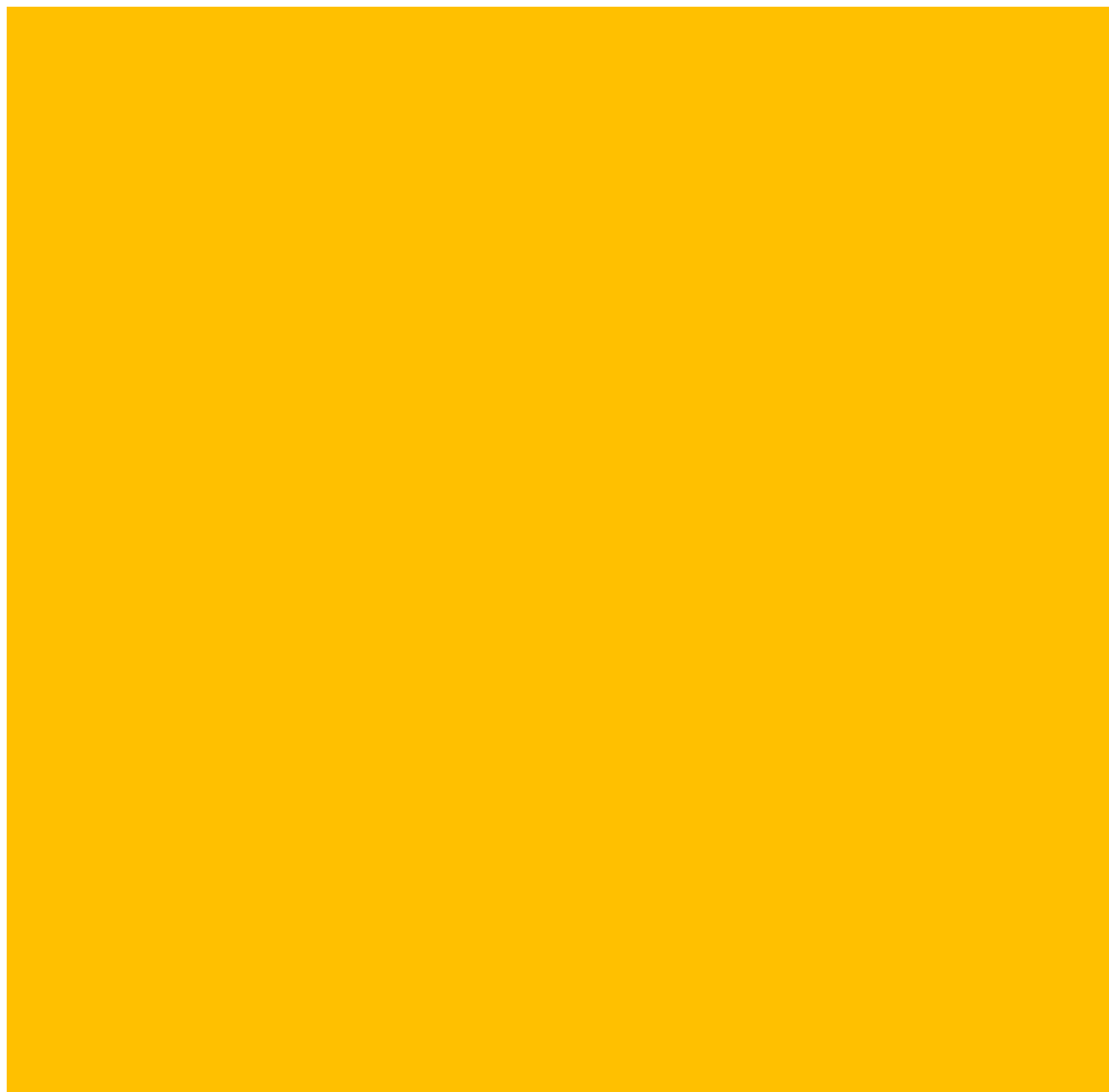

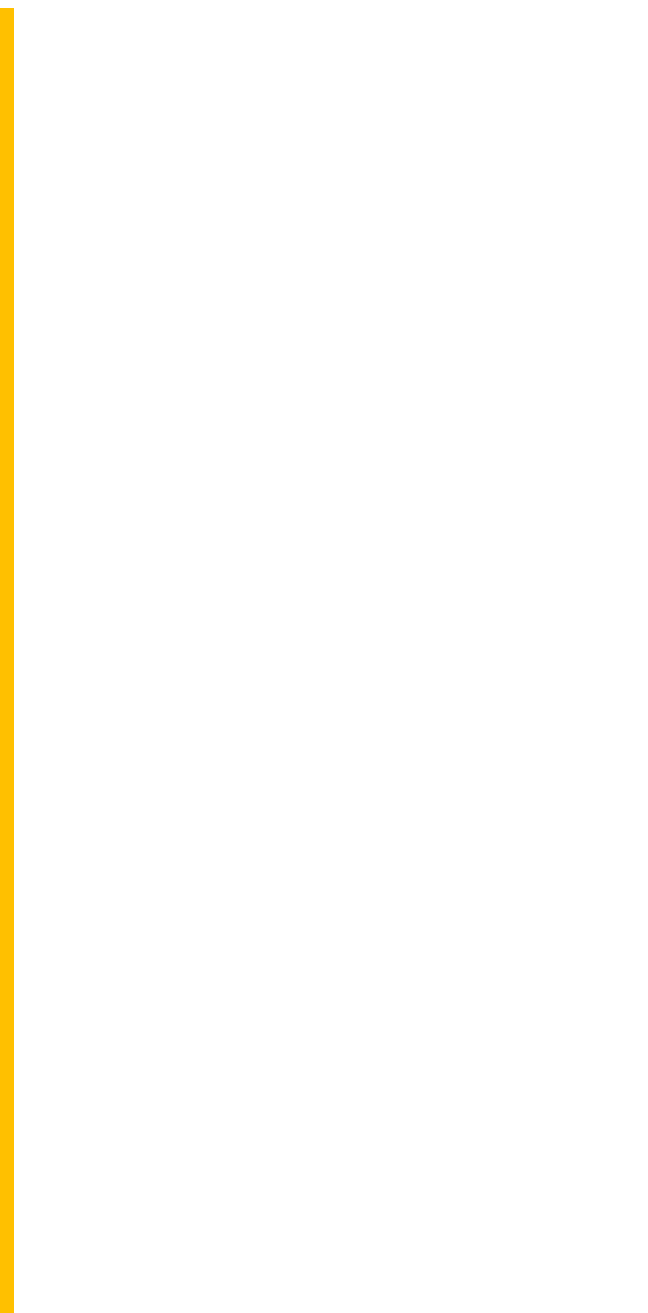

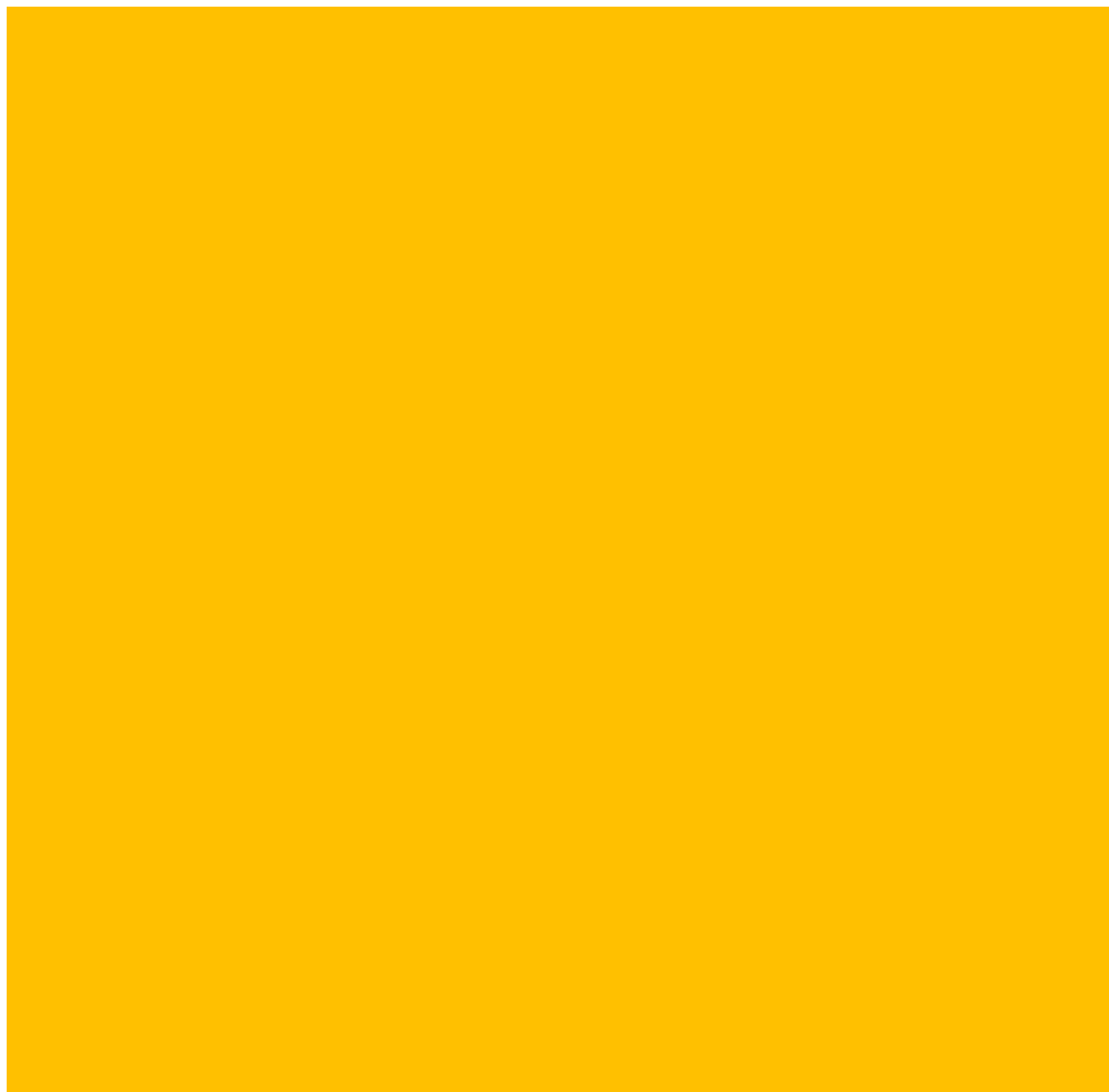

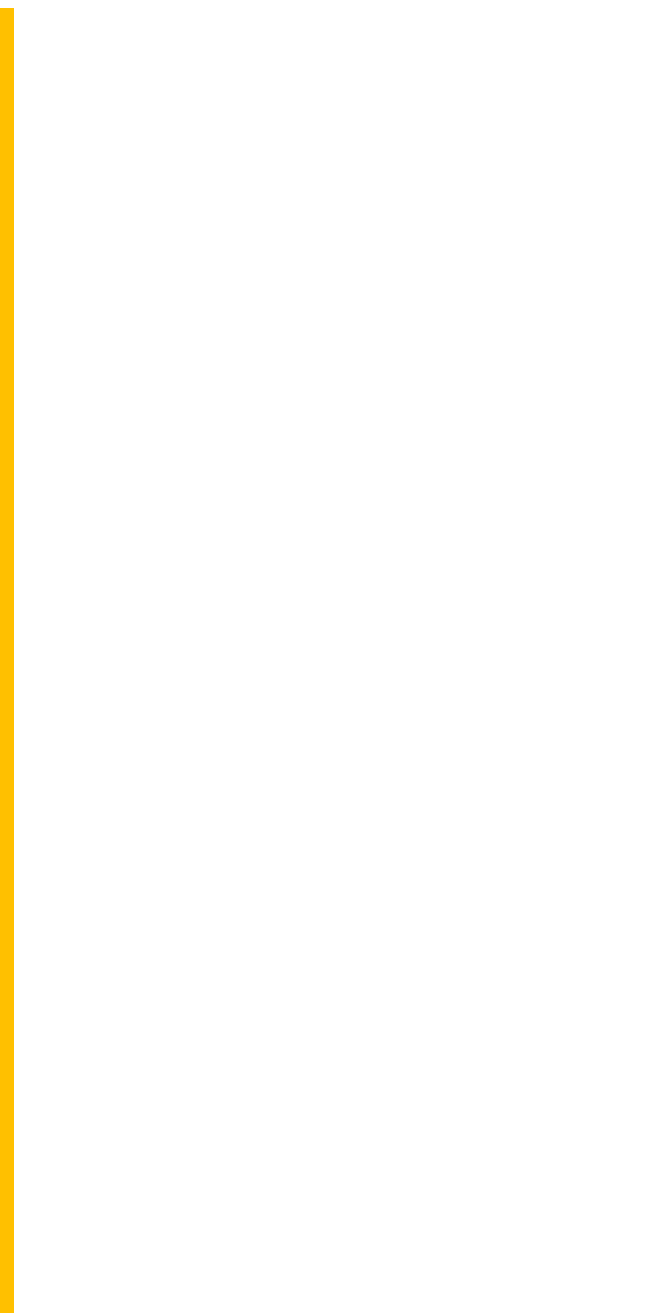

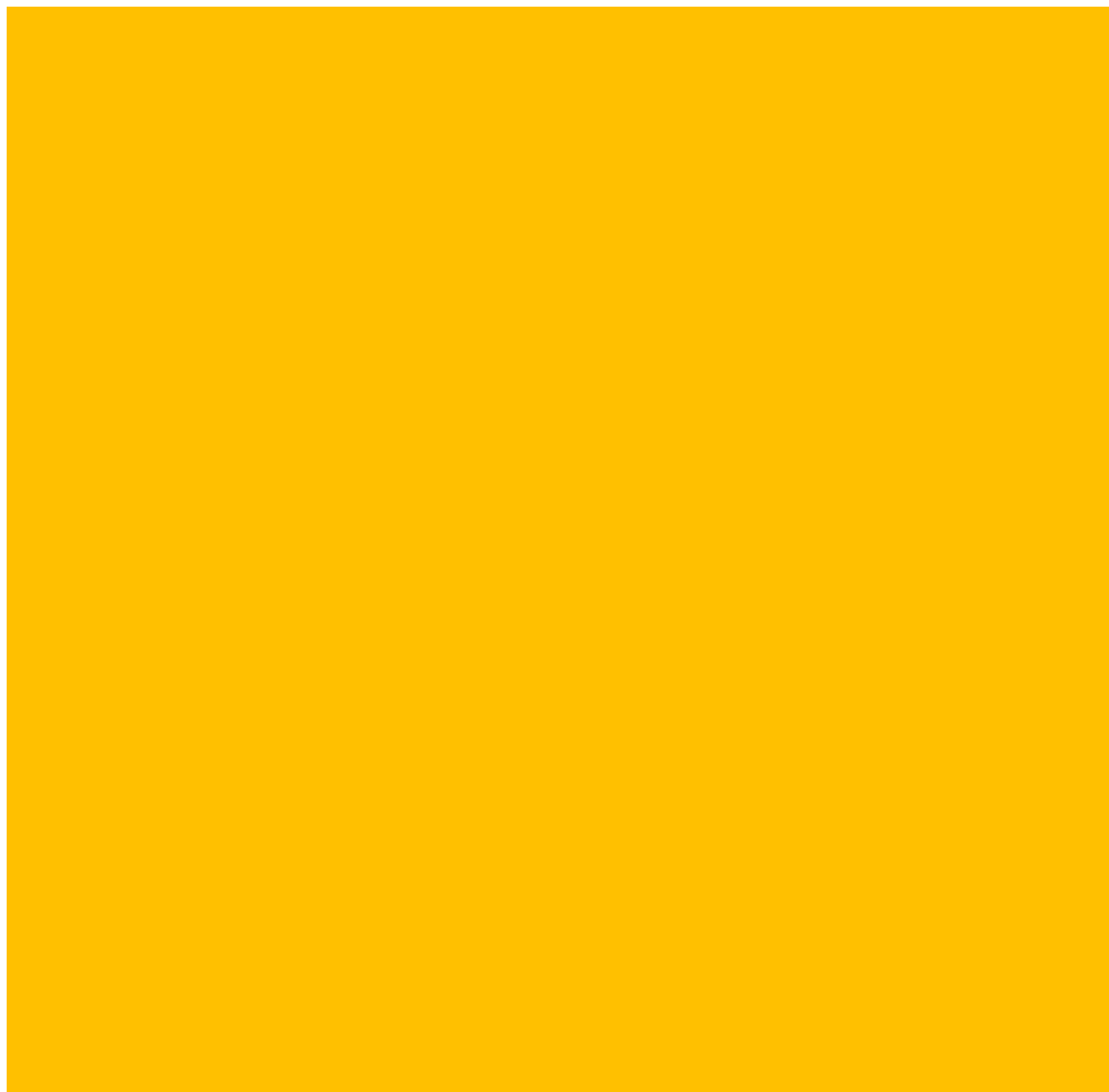

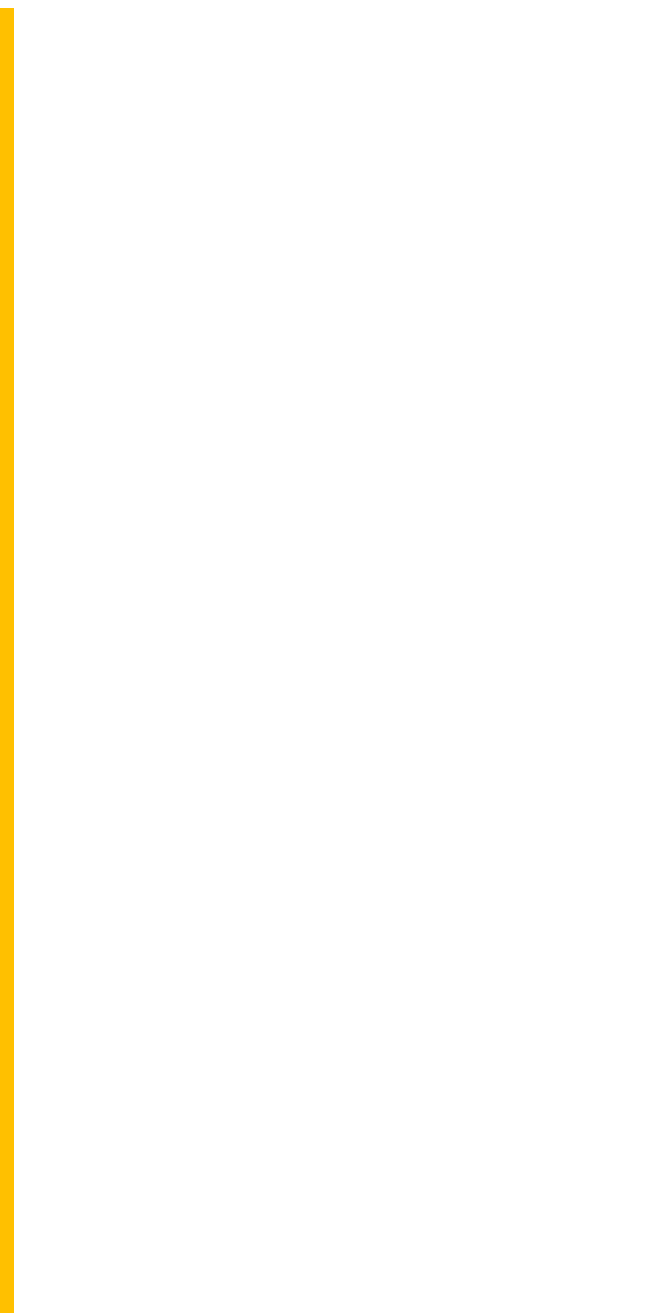

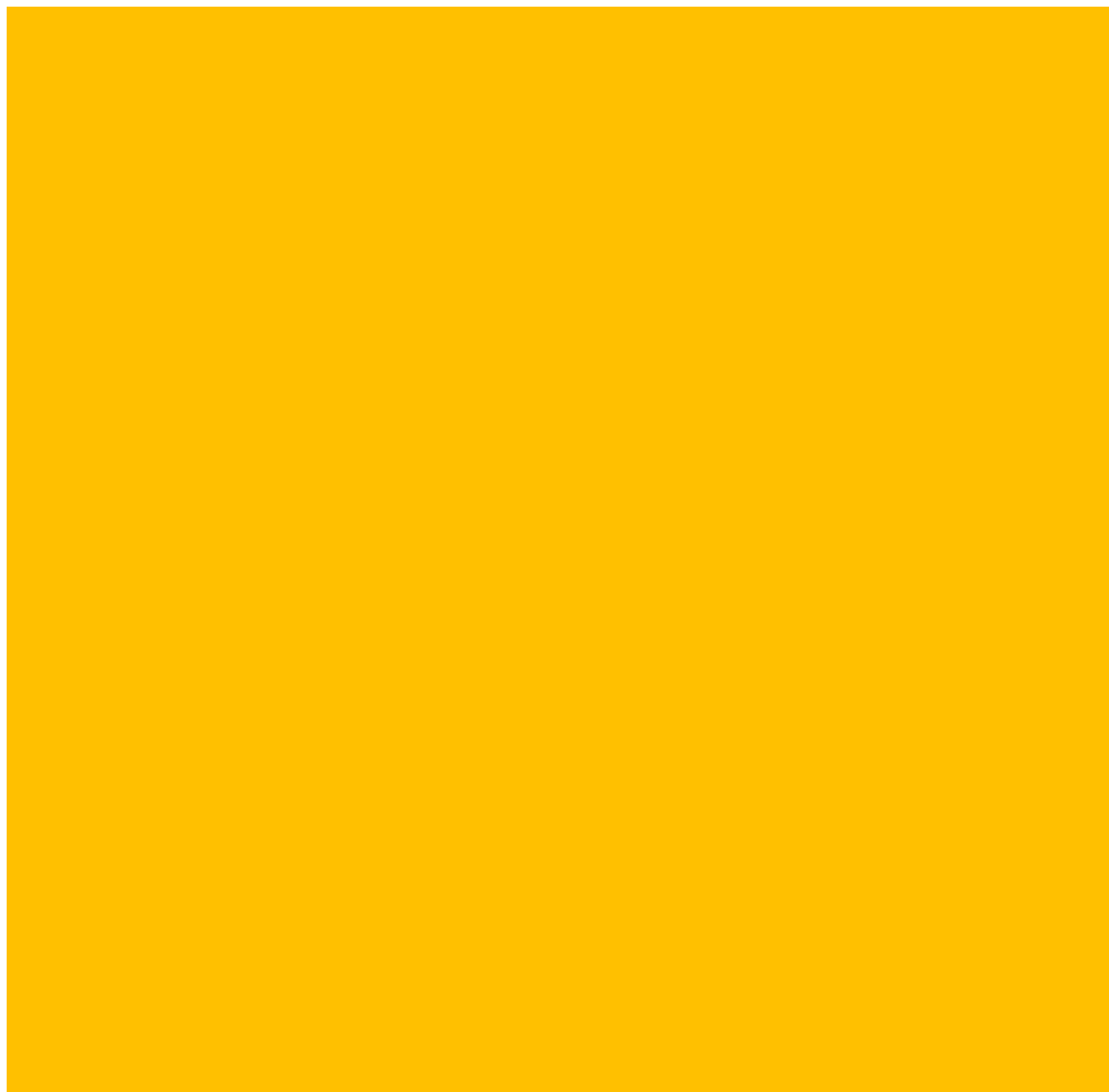

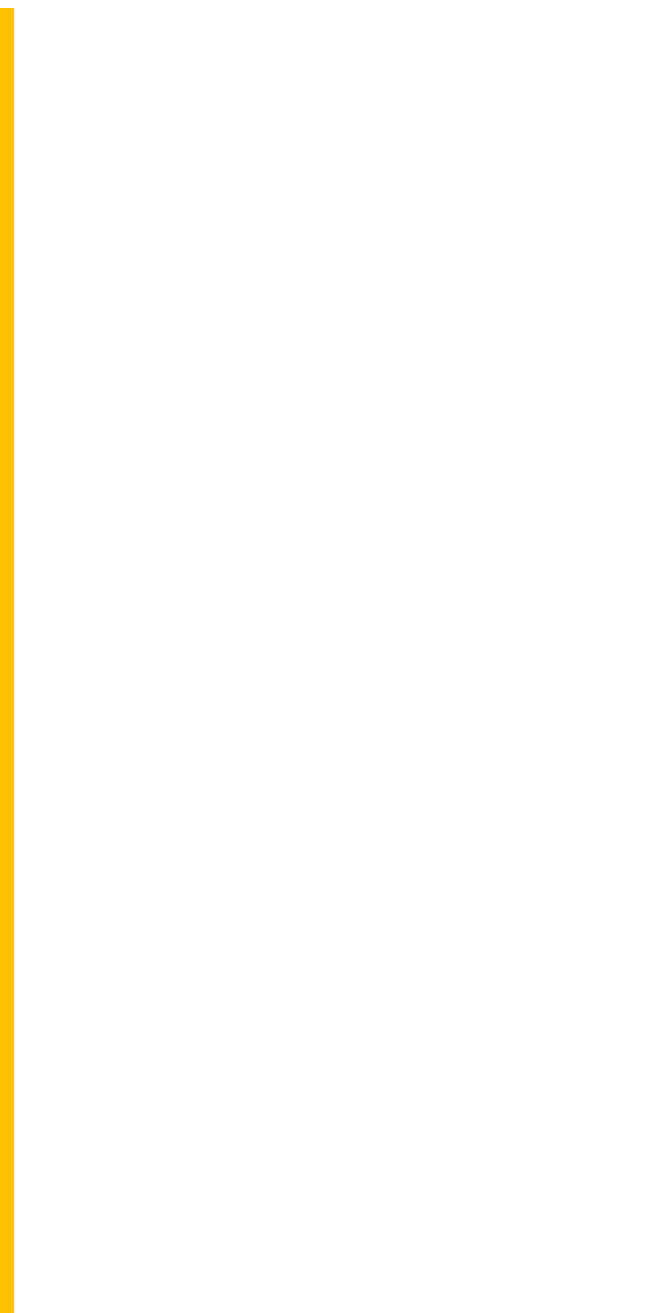

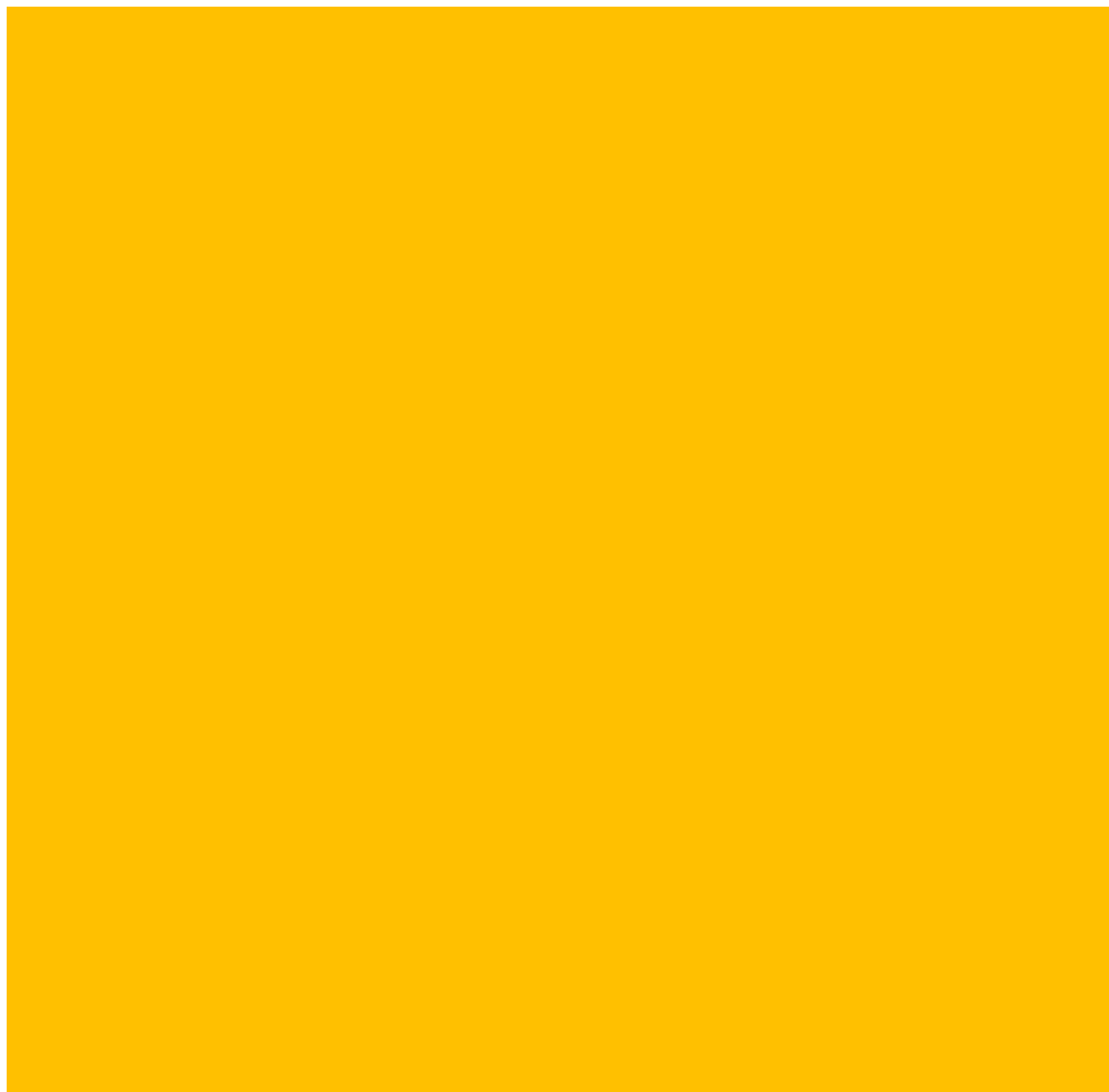

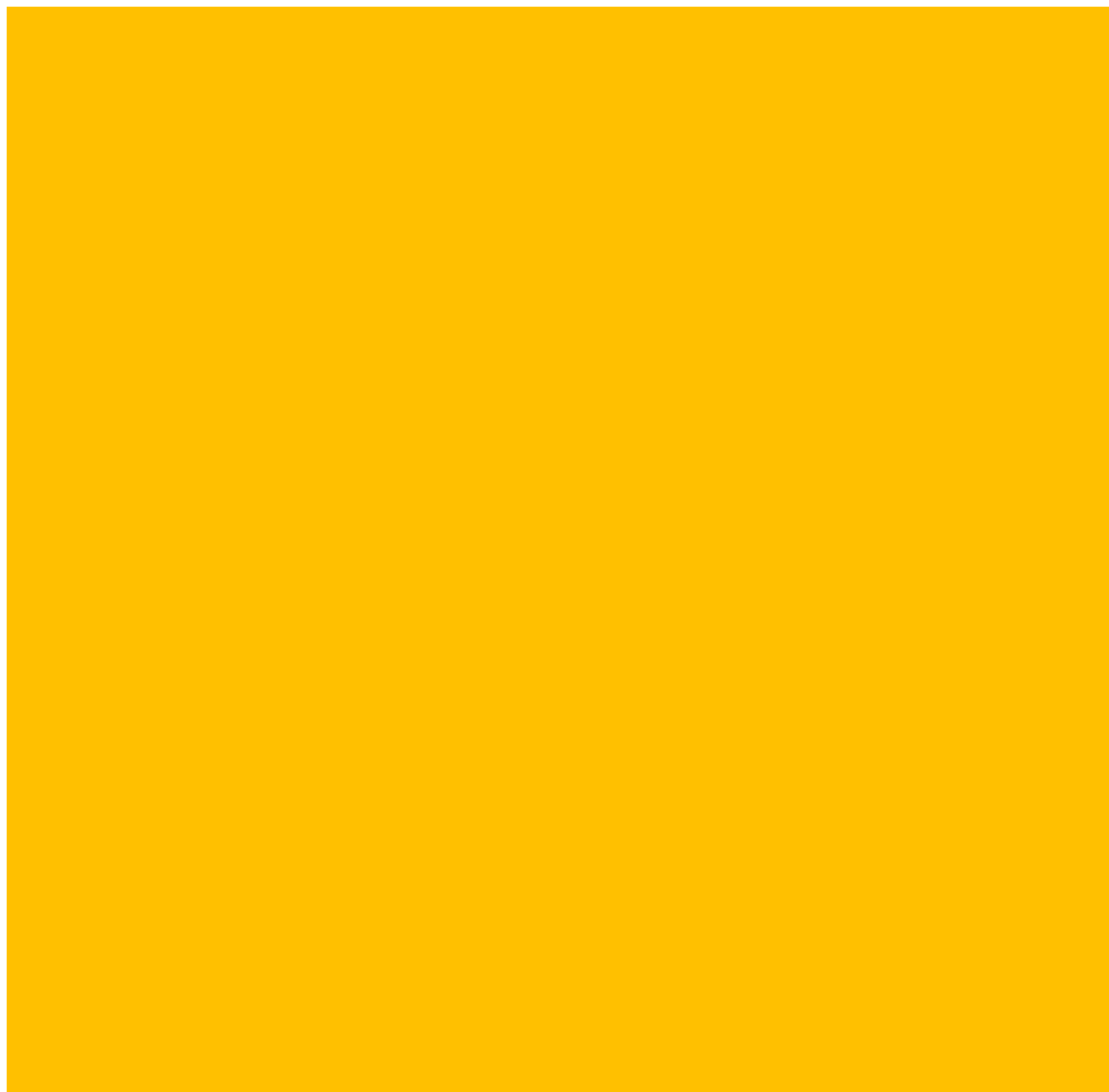

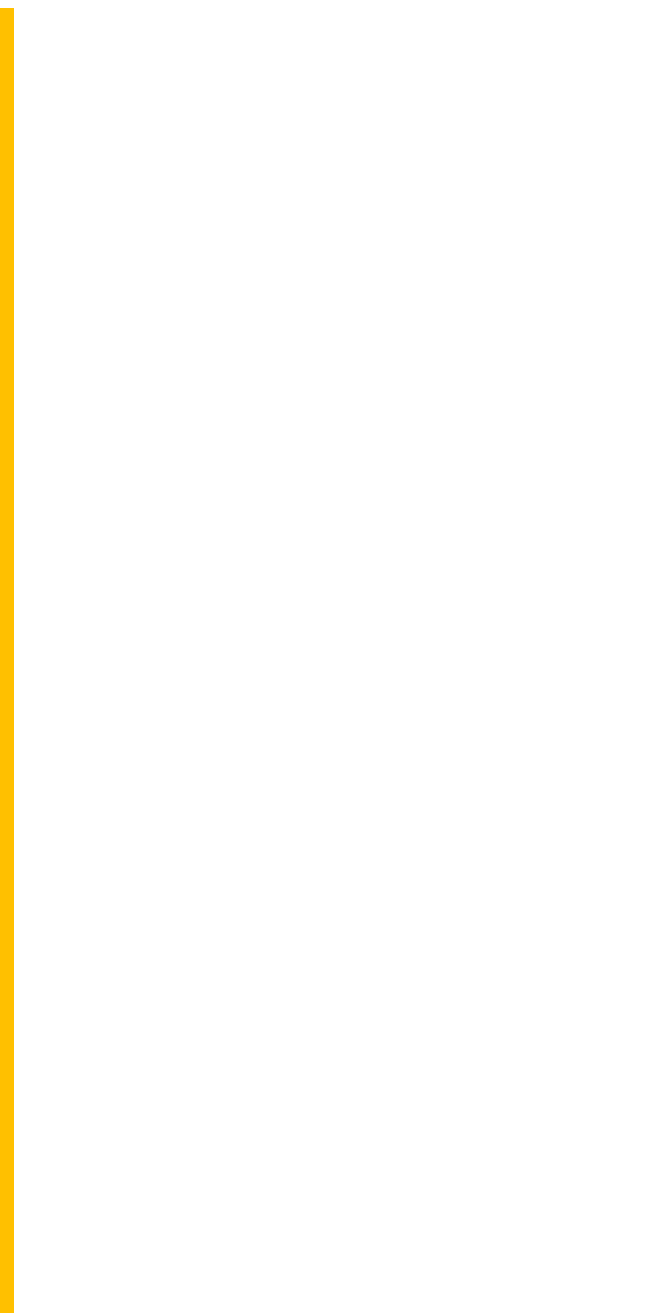

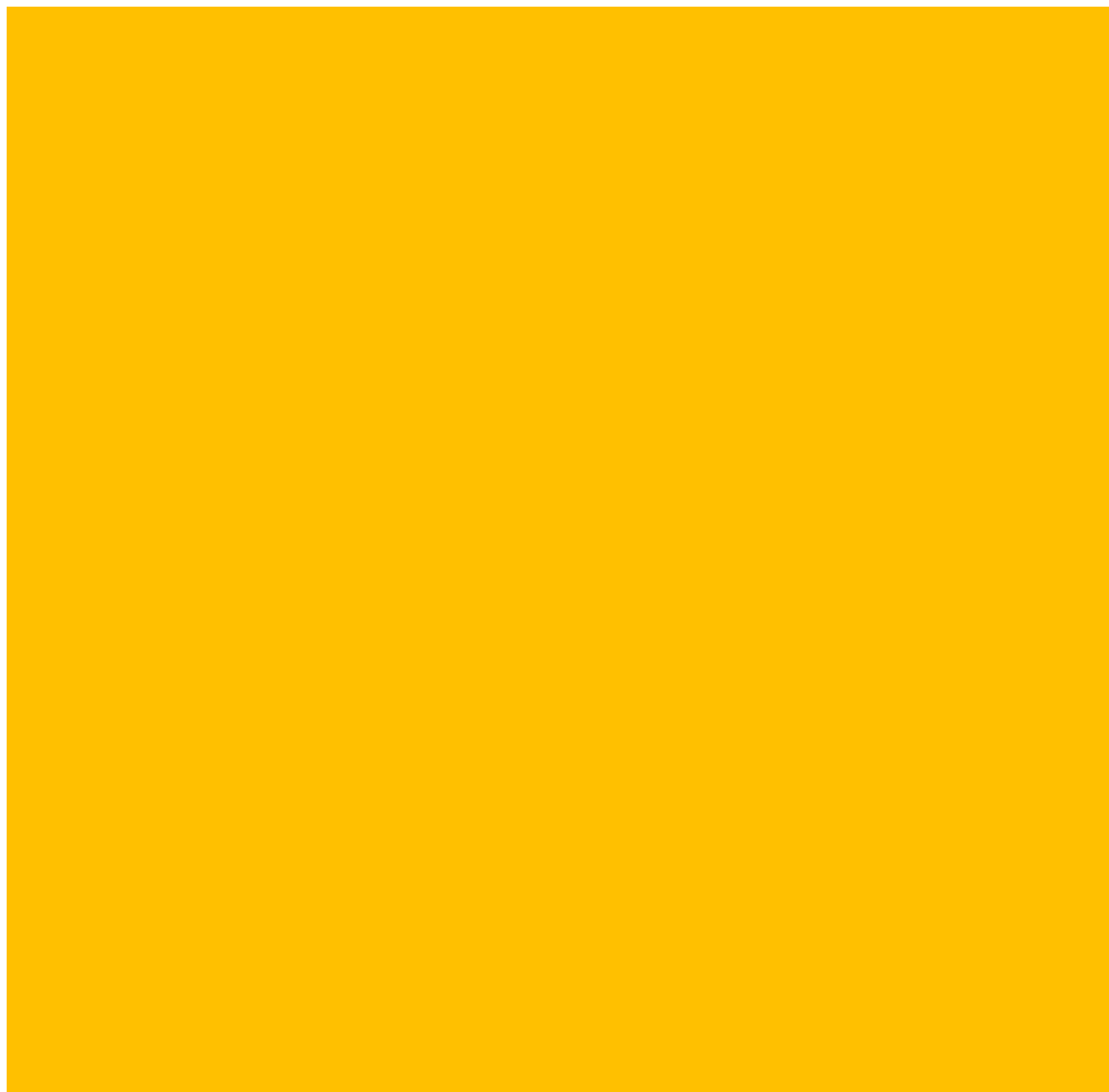

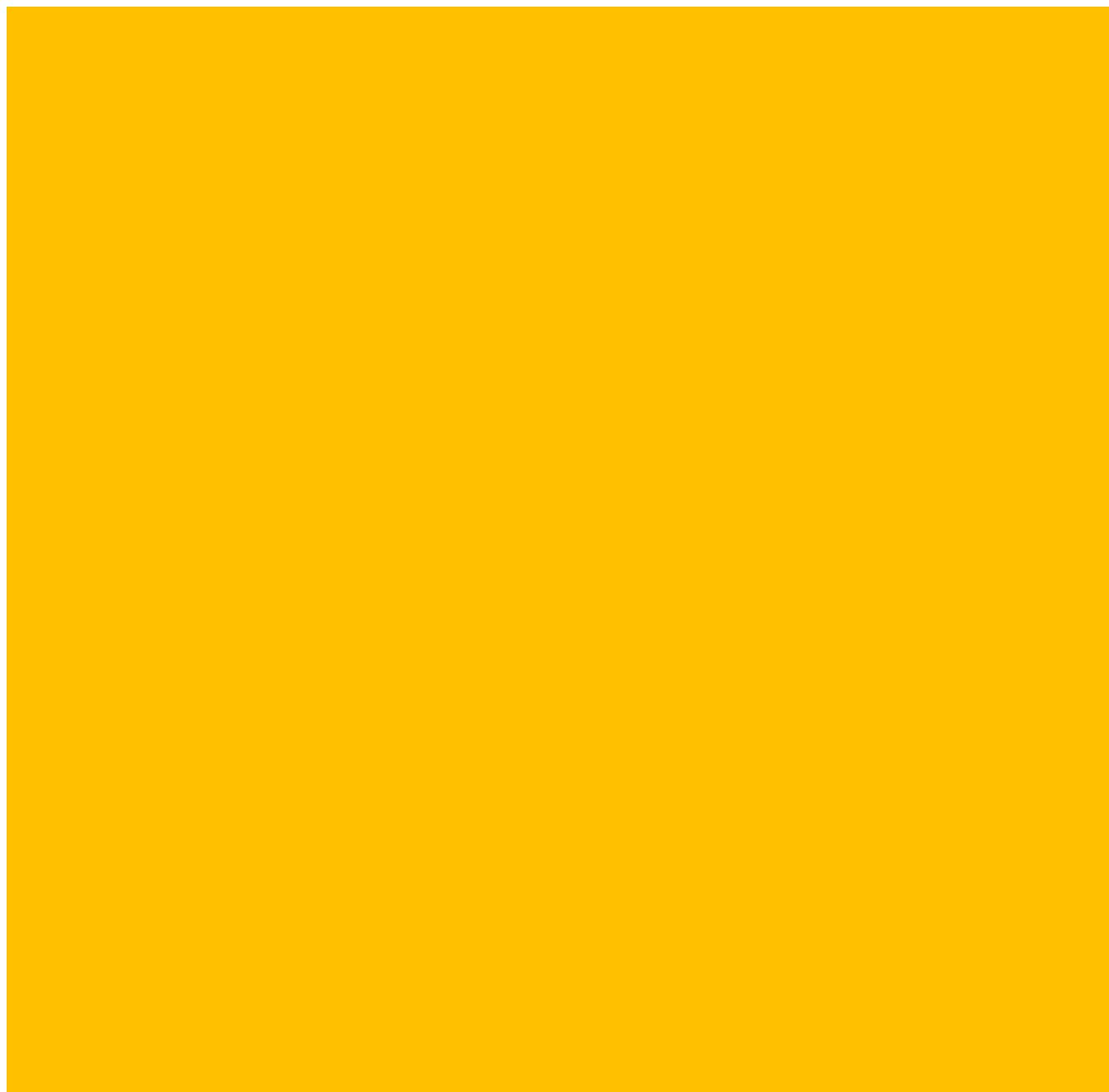

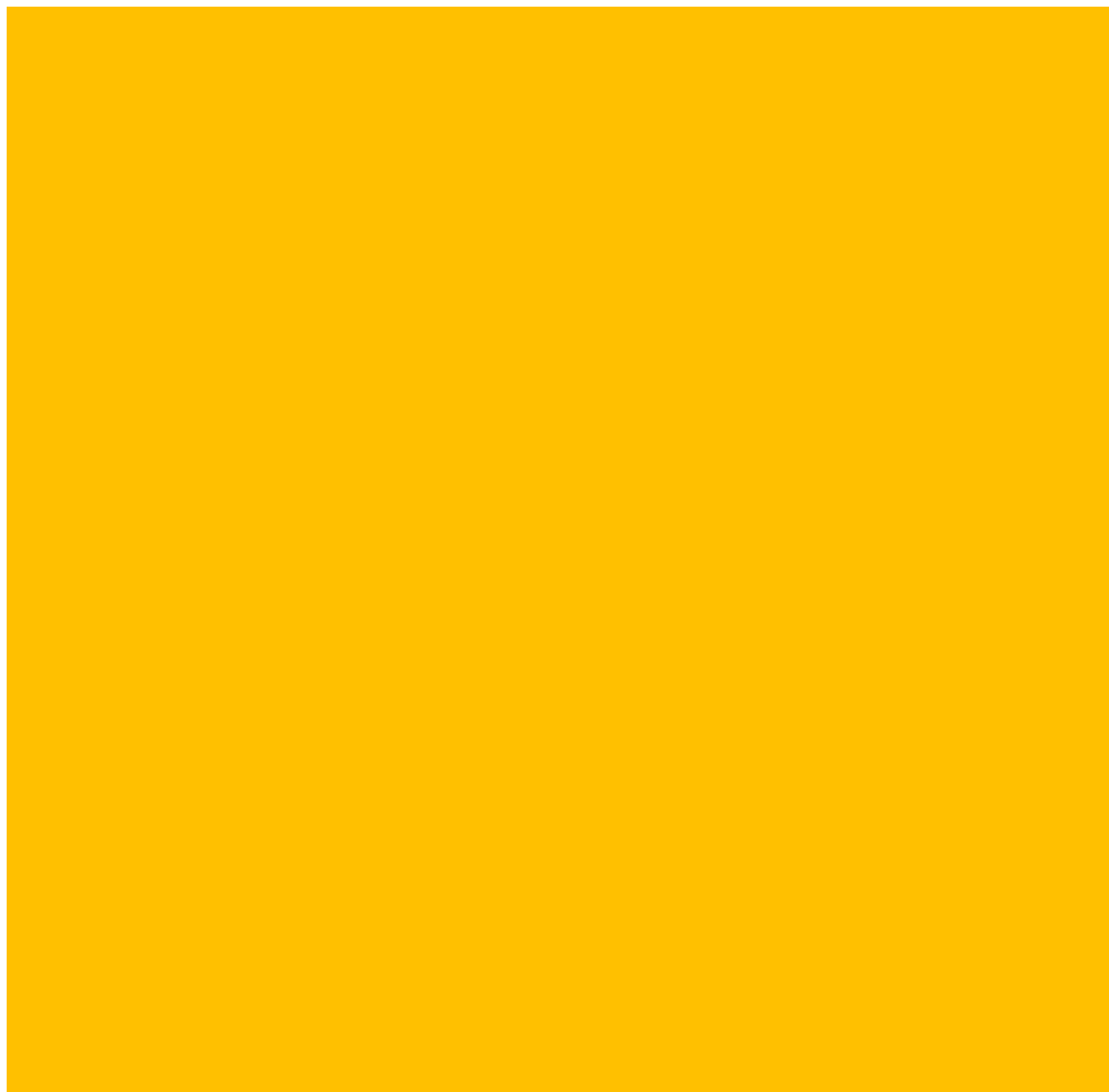

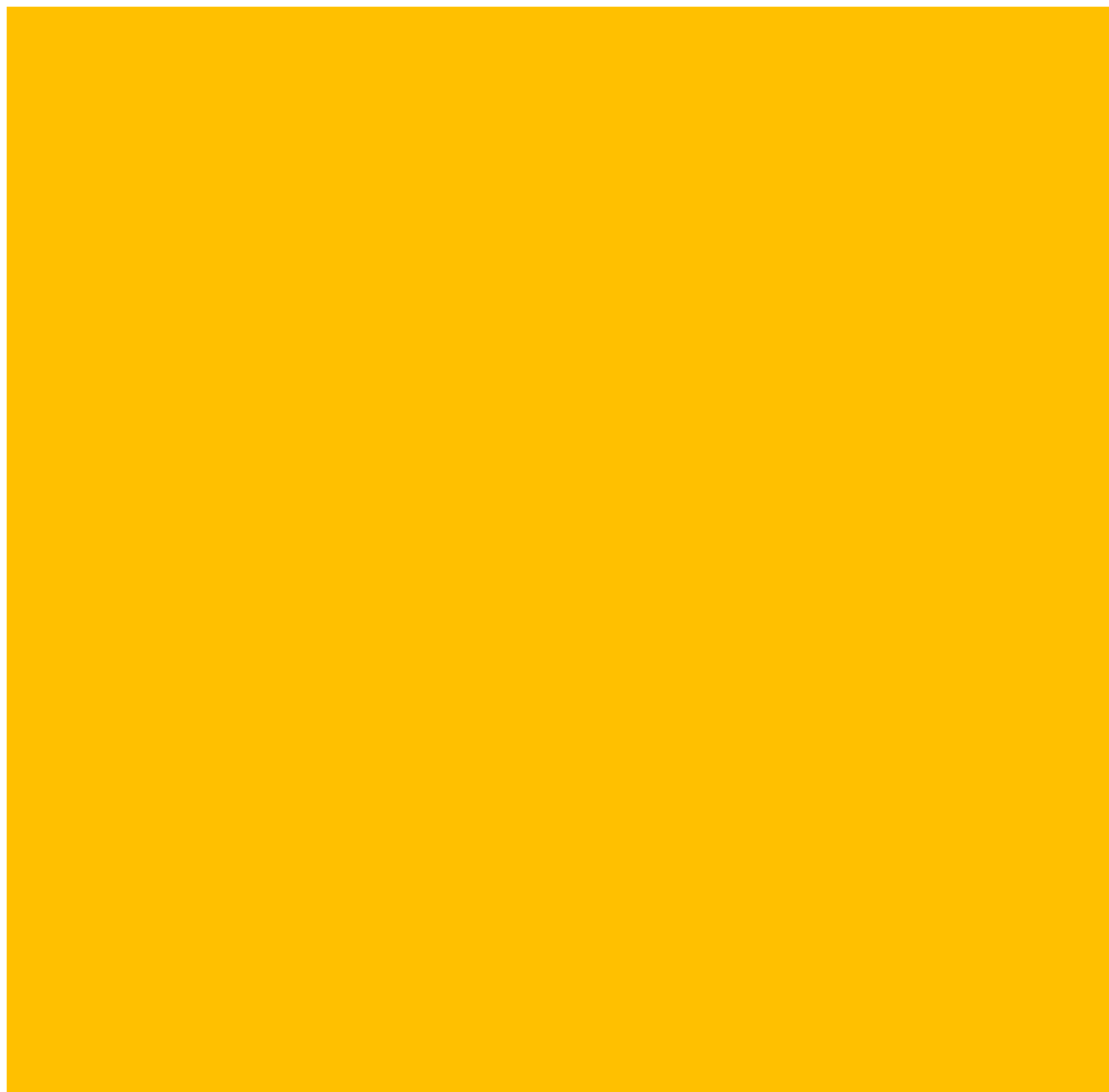

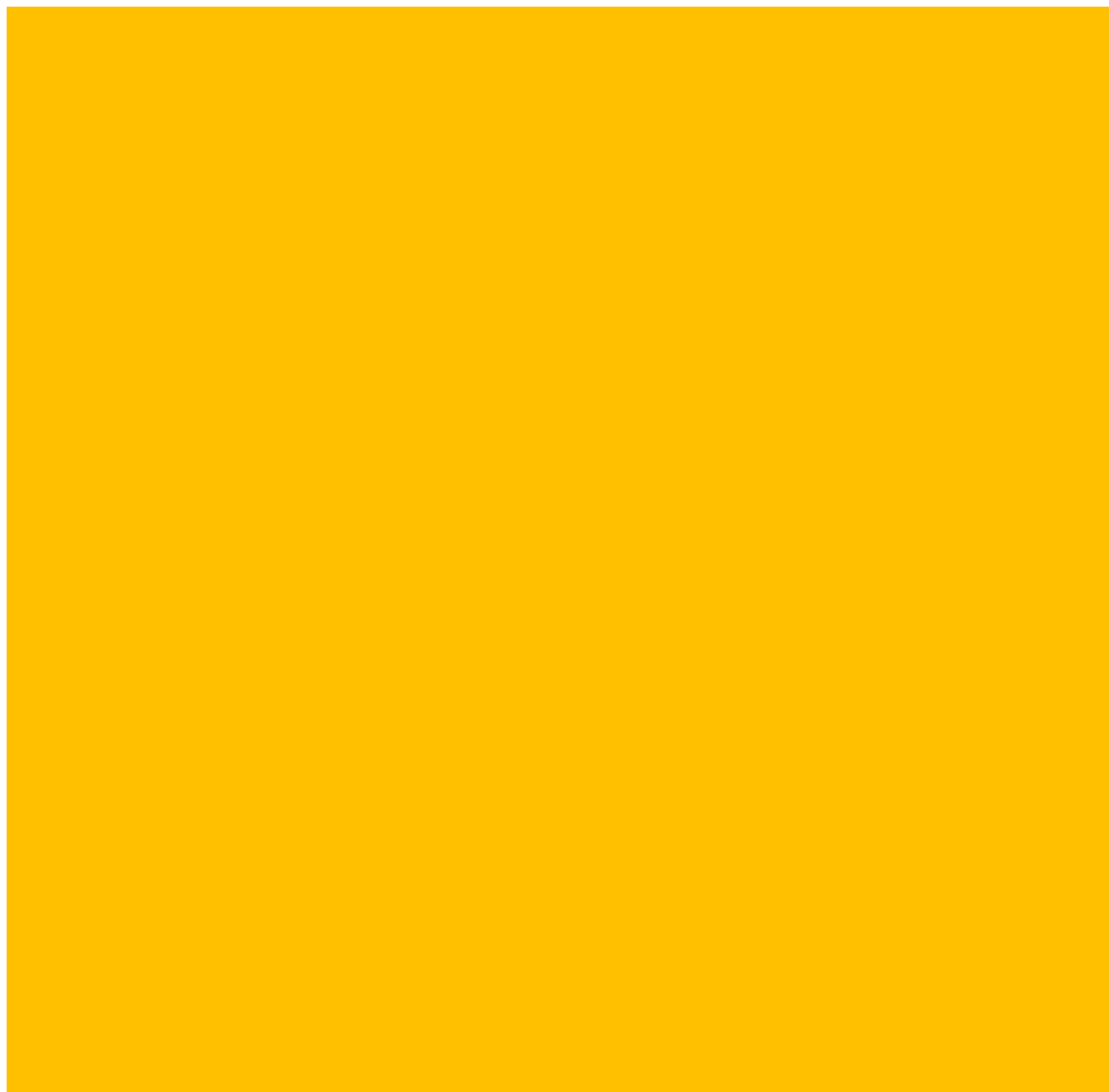

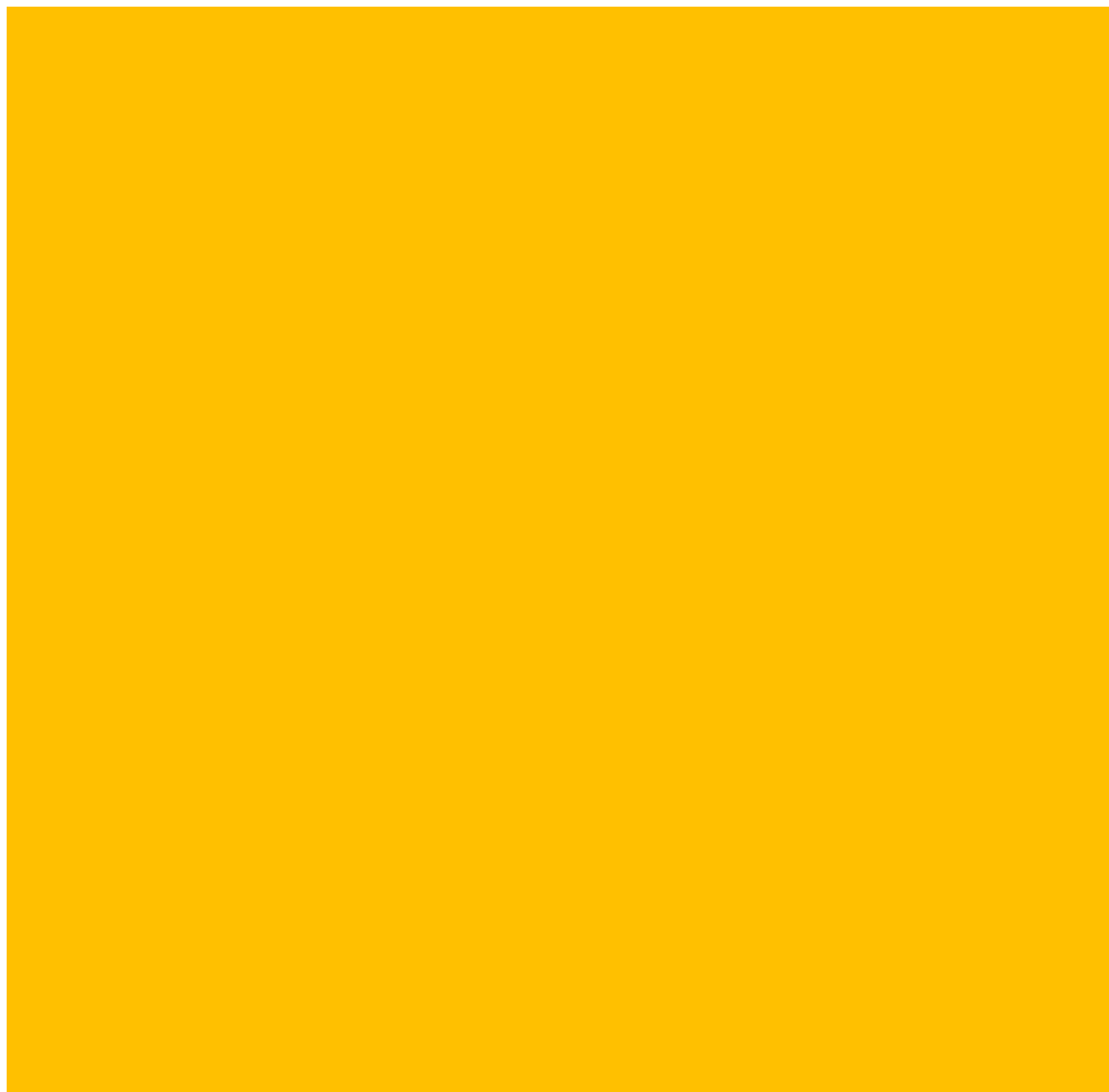

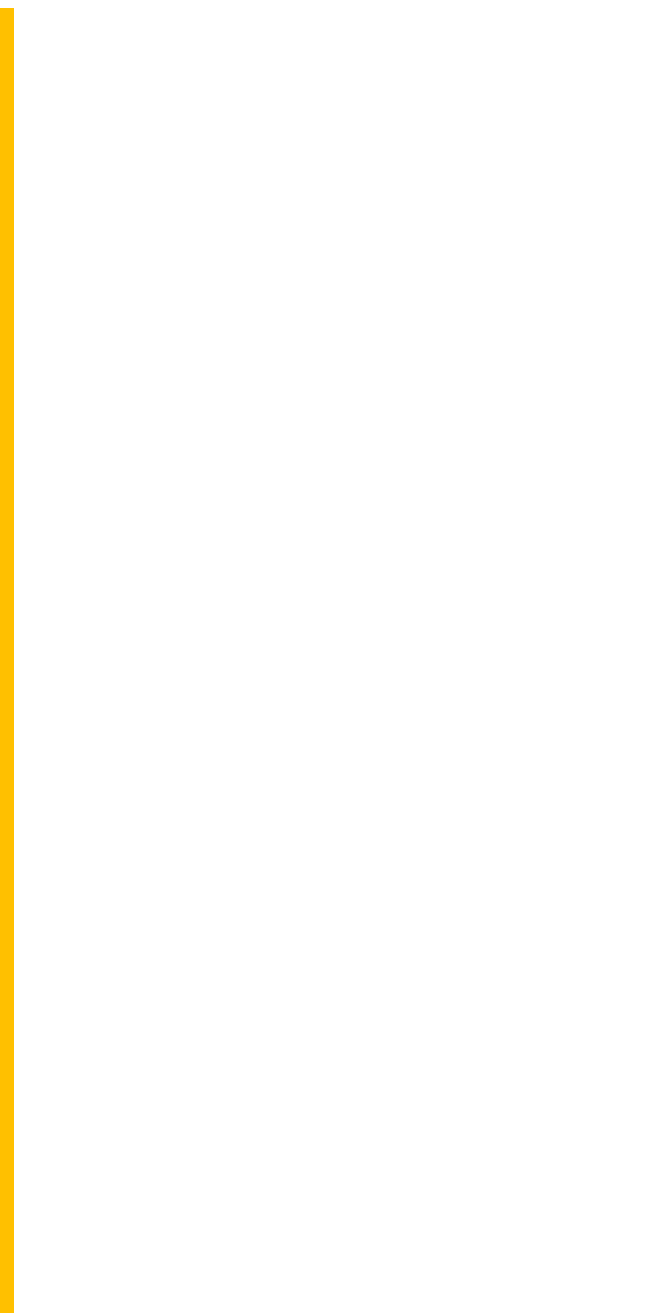

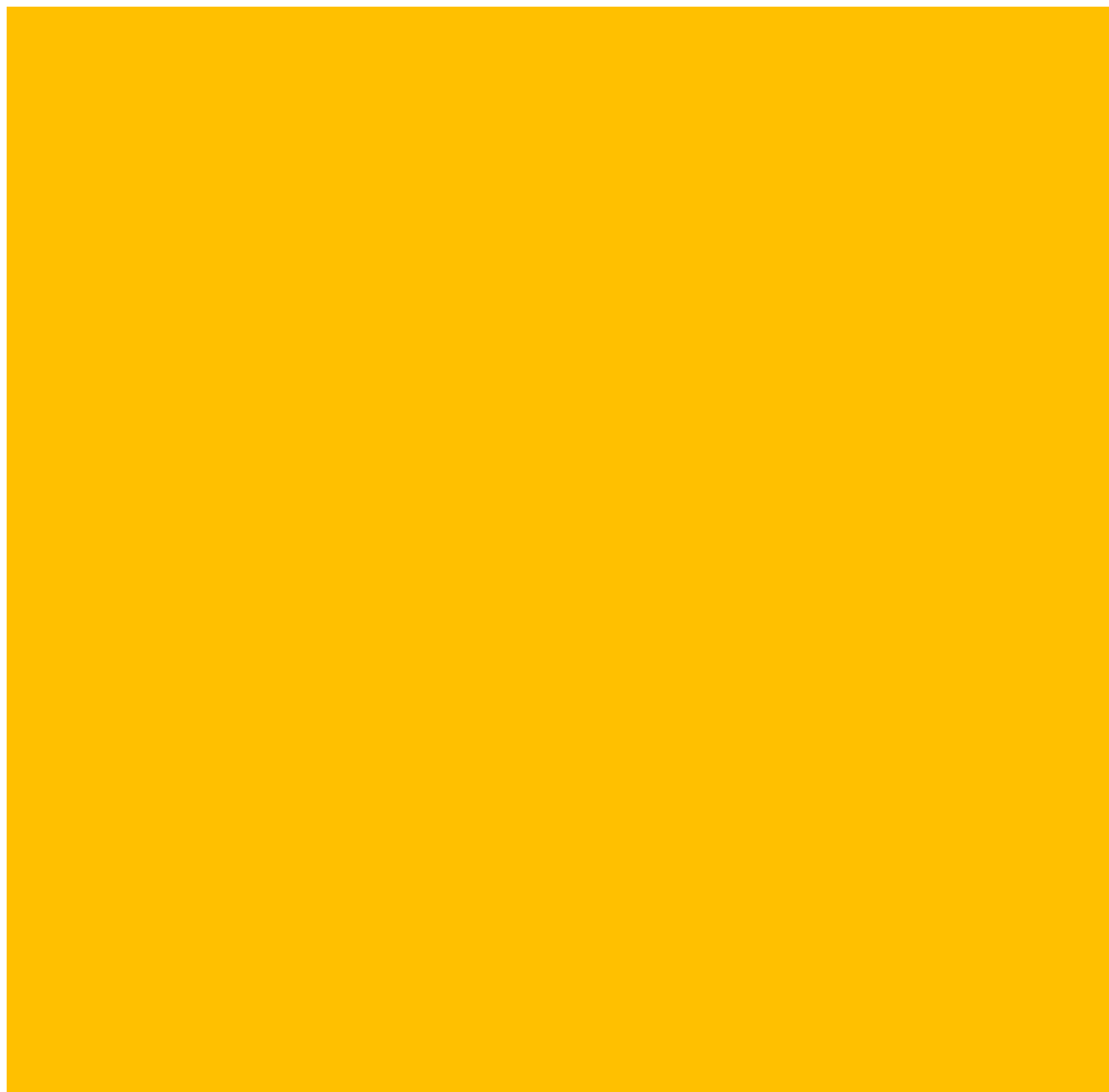

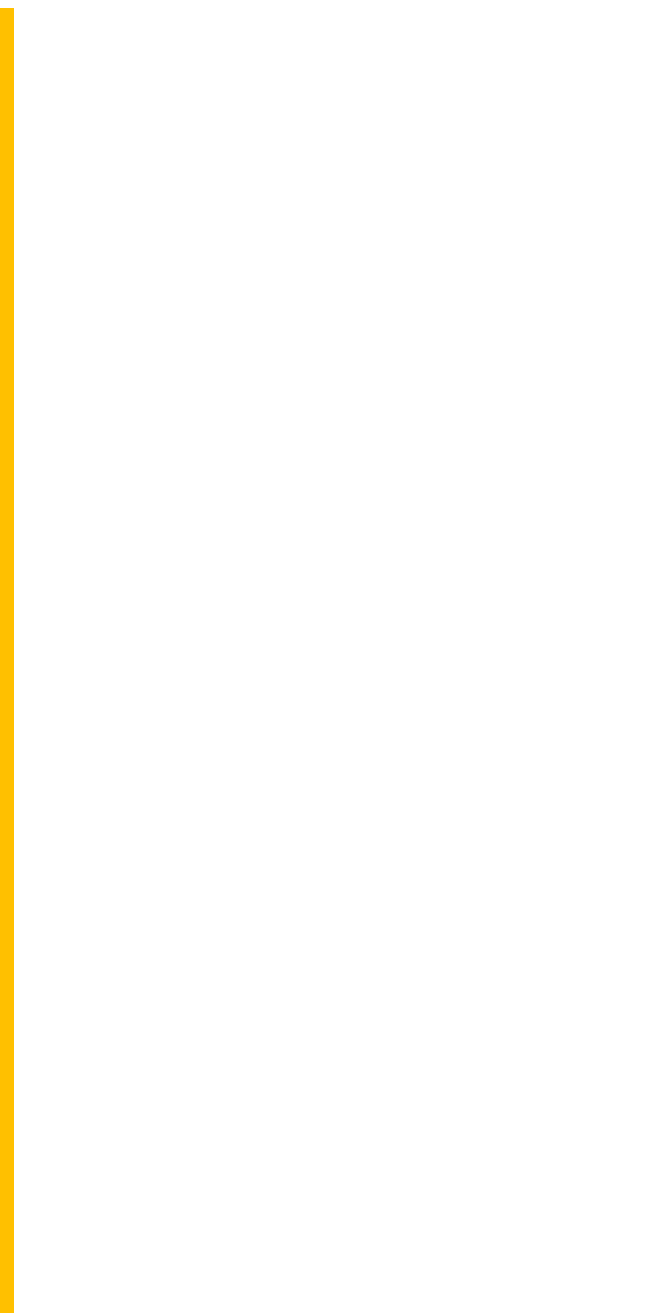

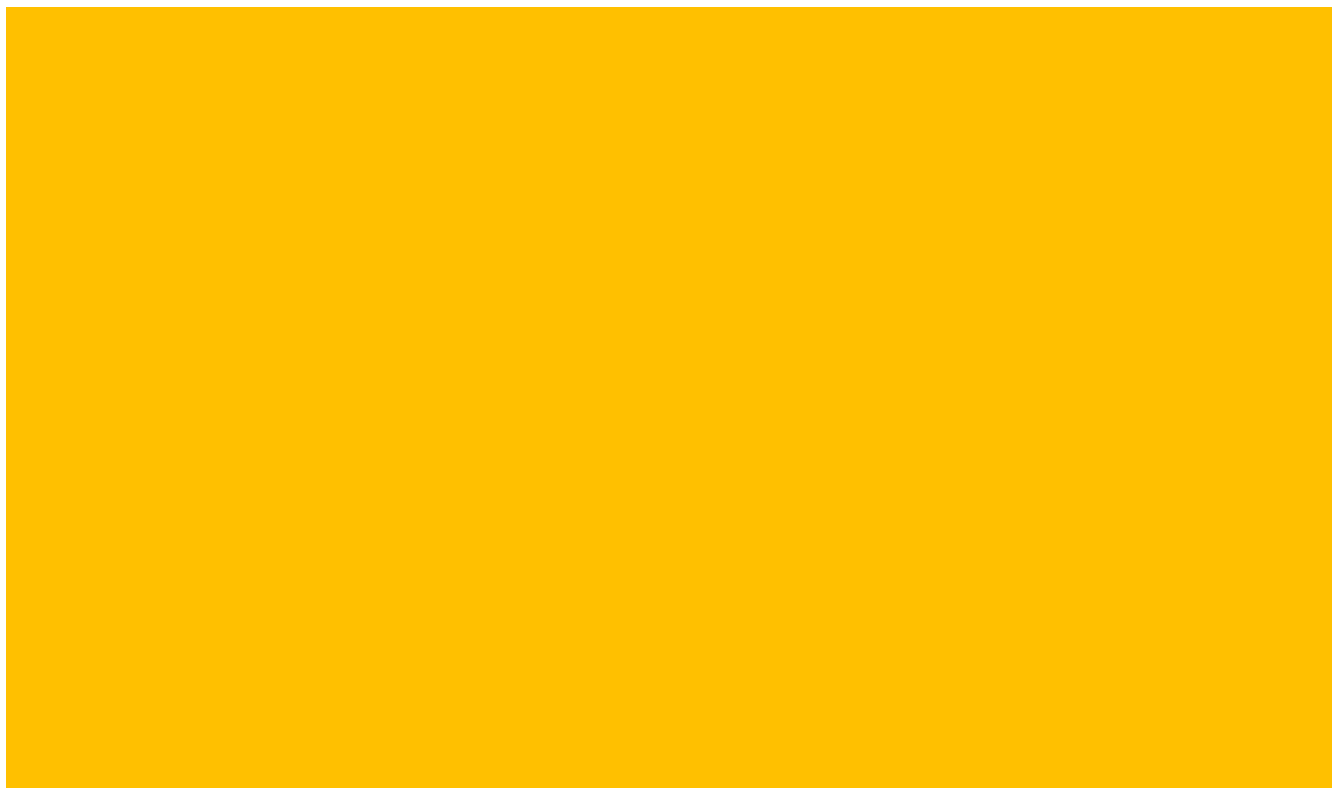

| Adj. P-value: (Bb_neg_newanalysis_12022024)/(RS_NEG_1 | Adj. P-value: (CM_neg_newanalysis_12022024)/(RS_NEG_1 |
| --- | --- |
| 0.002803841 | 0.001930382 |
| 5.72E-05 | 3.3882E-05 |
| 0.00014989 | 0.000152235 |
| 0.639853052 | 0.072296115 |
| 0.143545804 | 0.005287634 |
| 1 | 1 |
| 5.0932E-12 | 1 |
| 0.000321436 | 1 |
| 0.000117889 | 1 |
| 3.77401E-09 | 1 |
| 7.72086E-06 | 0.104259285 |
| 0.488252371 | 0.942613789 |
| 0.509392328 | 1 |
| 0.068828525 | 1 |
| 0.356726274 | 1 |
| 1.87542E-05 | 0.486024961 |
| 0.06562628 | 0.041741493 |
| 3.68692E-09 | 4.53858E-05 |
| 0.001986348 | 0.005658775 |
| 1 | 0.233341199 |
| 1 | 1 |
| 0.000869802 | 0.00797836 |
| 0.378054318 | 0.569787382 |
| 0.001513727 | 0.081236157 |
| 0.678348869 | 1 |
| 1.38767E-05 | 1 |
| 4.88024E-07 | 1 |
| 0.000620251 | 1 |
| 0.022105552 | 0.170314719 |
| 0.753036126 | 1 |
| 0.020324093 | 1 |
| 0.007529749 | 1 |
| 6.6894E-12 | 2.35188E-06 |
| 3.11953E-07 | 0.609564935 |
| 0.003835481 | 1 |
| 1.26297E-08 | 0.539640349 |
| 1.3893E-07 | 1 |
| 1.15199E-10 | 0.073602468 |
| 4.24286E-12 | 0.519120722 |
| 3.90067E-11 | 0.399157685 |
| 0.00073253 | 1 |
| 4.24286E-12 | 0.025071044 |
| 4.24286E-12 | 0.039415265 |
| 1.16664E-10 | 0.496565411 |
| 3.67831E-11 | 0.345153473 |
| 1.58708E-07 | 3.54143E-05 |
| 0.075970102 | 0.221693907 |
| 0.954088773 | 1 |
| 1 | 1 |
| 5.87103E-12 | 0.000121269 |
| 0.595236971 | 1 |
| 1 | 1 |
| 0.000675783 | 0.009762907 |
| 2.24967E-09 | 4.02563E-08 |
| 0.000185304 | 0.010625952 |
| 0.69191702 | 0.374431115 |
| 1.48876E-08 | 0.193911809 |
| 6.6756E-05 | 1.15133E-05 |
| 1.98194E-07 | 0.045589531 |
| 3.28795E-08 | 2.44171E-05 |
| 0.000226572 | 0.139884229 |
| 1.27292E-10 | 1.45964E-05 |
| 1.81216E-05 | 0.590022237 |
| 1 | 1 |
| 7.50363E-12 | 0.05651181 |
| 0.051408176 | 0.000856743 |
| 0.054739879 | 0.008449521 |
| 2.36618E-06 | 0.050702602 |
| 1 | 0.004550323 |
| 0.46910244 | 1 |
| 5.33429E-07 | 0.000965061 |
| 1 | 1 |
| 0.134499331 | 0.068563192 |
| 1 | 1 |
| 0.198231926 | 0.004056347 |
| 3.60189E-05 | 0.028061805 |
| 1 | 1 |
| 0.006270488 | 0.011746311 |
| 0.448073779 | 0.293413582 |
| 2.8879E-07 | 0.00291001 |
| 4.82025E-09 | 0.060680219 |
| 1.05723E-10 | 2.53026E-06 |
| 1.61933E-08 | 5.11711E-08 |
| 1 | 0.696030219 |
| 0.971934414 | 0.112276616 |
| 2.17892E-05 | 0.059355554 |
| 0.000749155 | 0.04975441 |
| 0.003278062 | 0.996579855 |
| 4.24286E-12 | 0.022766421 |
| 1.95653E-07 | 0.00489589 |
| 0.20906929 | 0.058872417 |
| 0.001547591 | 0.008935712 |
| 0.064298288 | 0.058499989 |
| 1 | 1 |
| 0.000555342 | 0.057542979 |
| 0.015672023 | 0.118004036 |
| 1 | 1 |
| 7.80951E-05 | 0.005307211 |
| 0.345592674 | 0.013008914 |
| 1 | 1 |
| 1 | 1 |
| 0.708614606 | 1 |
| 0.005159104 | 0.03371941 |
| 0.276429718 | 0.204501582 |
| 0.954109594 | 1 |
| 1 | 0.790434563 |
| 0.296130648 | 0.496380936 |
| 0.215812608 | 0.124754427 |
| 4.24286E-12 | 2.29624E-05 |
| 4.24286E-12 | 1.5061E-10 |
| 4.24286E-12 | 9.8967E-08 |
| 1.0573E-11 | 0.022781006 |
| 2.26289E-10 | 0.00023322 |
| 4.24286E-12 | 5.11562E-11 |
| 3.17159E-05 | 4.01037E-05 |
| 0.003079652 | 0.000777343 |
| 4.30496E-12 | 0.193238091 |
| 4.24286E-12 | 0.469928321 |
| 4.24286E-12 | 0.000777343 |
| 0.055399584 | 0.004721187 |
| 1 | 1 |
| 1.75516E-10 | 0.876163495 |
| 4.24286E-12 | 1 |
| 0.000128588 | 0.000111136 |
| 0.020325322 | 0.007880621 |
| 1 | 1 |
| 0.146385808 | 0.457106383 |
| 0.026913175 | 0.421789226 |
| 0.386504051 | 0.192914729 |
| 1 | 1 |
| 4.24286E-12 | 7.10448E-09 |
| 5.3565E-07 | 0.048525849 |
| 4.24286E-12 | 1.54537E-07 |
| 1.45784E-07 | 0.000594343 |
| 1.21892E-09 | 0.003352021 |
| 5.32529E-09 | 0.002066317 |
| 3.79653E-11 | 0.000342813 |
| 1 | 0.942881049 |
| 0.001070651 | 0.379016293 |
| 0.03140145 | 1 |
| 0.906353593 | 0.038294422 |
| 0.013033985 | 0.261172607 |
| 6.81904E-09 | 0.000677421 |
| 1.25708E-11 | 5.82057E-07 |
| 1.10934E-05 | 4.91412E-06 |
| 4.72651E-11 | 3.47279E-09 |
| 4.24286E-12 | 3.59904E-08 |
| 2.32626E-10 | 1.41316E-06 |
| 4.24286E-12 | 2.24664E-07 |
| 1 | 0.974840337 |
| 6.58649E-11 | 4.48089E-07 |
| 3.459E-05 | 0.000472908 |
| 4.90101E-12 | 1.49107E-06 |
| 4.24286E-12 | 1.71391E-06 |
| 0.007907724 | 1 |
| 0.000107883 | 0.212749727 |
| 0.045940562 | 0.052473106 |
| 4.93797E-06 | 0.002470179 |
| 1 | 1 |
| 1.28946E-07 | 0.000342813 |
| 0.002755642 | 0.078237398 |
| 8.07514E-10 | 0.000229148 |
| 4.24286E-12 | 1.21085E-09 |
| 2.0986E-06 | 9.16311E-06 |
| 4.24286E-12 | 9.34446E-07 |
| 0.391982723 | 0.801496499 |
| 0.55817265 | 1 |
| 4.24286E-12 | 1.38079E-09 |
| 4.24286E-12 | 1.05627E-08 |
| 4.24286E-12 | 6.58627E-06 |
| 4.24286E-12 | 6.65562E-08 |
| 4.24286E-12 | 4.33811E-10 |
| 0.00596517 | 1 |
| 3.09819E-10 | 9.03083E-05 |
| 8.70474E-12 | 8.83044E-08 |
| 4.41419E-12 | 2.87606E-08 |
| 1.19307E-11 | 1.38079E-09 |
| 1.0745E-07 | 2.0073E-06 |
| 7.62837E-11 | 1.10236E-06 |
| 2.83545E-10 | 2.61499E-05 |
| 0.000418586 | 0.000179263 |

| Adj. P-value: (SM_neg_newanalysis_12022024) / (RS_NEG_NEWANALYSIS_12022024) | Adj. P-value: (Bb_neg_newanalysis_12022024) / (SM_neg_r |
| --- | --- |
| 0.086809259 | 1 |
| 0.001055402 | 1 |
| 0.015863071 | 0.896981247 |
| 1 | 1 |
| 0.121627898 | 1 |
| 1 | 0.654546248 |
| 0.118324897 | 9.72174E-12 |
| 0.194700331 | 5.77317E-07 |
| 0.356543114 | 5.57599E-07 |
| 0.062499525 | 2.05955E-11 |
| 0.015050192 | 1.5504E-09 |
| 1 | 1 |
| 1 | 1 |
| 0.679152447 | 1 |
| 0.067287701 | 1 |
| 0.587745582 | 0.0087161 |
| 0.002446594 | 0.86061647 |
| 0.000117546 | 0.017209248 |
| 0.732657904 | 4.04914E-05 |
| 0.012476789 | 0.216950867 |
| 0.101274379 | 0.093586447 |
| 0.150594857 | 0.690988331 |
| 0.043127707 | 1 |
| 0.137793118 | 0.86061647 |
| 0.085625175 | 0.001903821 |
| 0.012745297 | 2.15928E-09 |
| 0.461417177 | 5.86571E-09 |
| 0.017453525 | 7.304E-08 |
| 1 | 0.573462107 |
| 0.018057981 | 0.000388228 |
| 0.236591324 | 5.81259E-05 |
| 0.019908334 | 9.81058E-07 |
| 8.68627E-06 | 2.23425E-05 |
| 0.208148889 | 1.42372E-09 |
| 1 | 0.000571638 |
| 0.980226261 | 9.97669E-07 |
| 1 | 9.3875E-09 |
| 0.555573652 | 1.78843E-08 |
| 0.870351558 | 9.72174E-12 |
| 0.805218455 | 2.5599E-09 |
| 1 | 0.007102113 |
| 0.337080261 | 9.72174E-12 |
| 0.001929558 | 2.51695E-11 |
| 1 | 4.11151E-09 |
| 1 | 1.08916E-09 |
| 0.004925796 | 0.026162141 |
| 0.01382918 | 1 |
| 1 | 0.927067309 |
| 1 | 1 |
| 0.003659926 | 4.29176E-08 |
| 1 | 0.539855587 |
| 0.398908095 | 0.142397259 |
| 0.000246318 | 1 |
| 1.25428E-05 | 0.07290173 |
| 0.1688988 | 0.304064439 |
| 0.496354251 | 0.024122449 |
| 0.80990451 | 2.09086E-06 |
| 0.018056763 | 0.666684921 |
| 0.515207136 | 9.94723E-05 |
| 6.80646E-06 | 0.699953236 |
| 1 | 0.006457209 |
| 0.02023958 | 1.35641E-06 |
| 1 | 9.08095E-05 |
| 0.423365057 | 0.856480671 |
| 0.557269574 | 4.69948E-10 |
| 0.000759566 | 0.648176562 |
| 0.004469166 | 1 |
| 1 | 5.493E-05 |
| 0.039761868 | 0.026762622 |
| 0.086440594 | 1 |
| 1 | 1.91831E-05 |
| 0.108077888 | 0.022875009 |
| 0.009213946 | 0.965901645 |
| 1 | 1 |
| 0.032446914 | 1 |
| 0.321987128 | 0.041639925 |
| 1 | 0.772644093 |
| 0.001142579 | 1 |
| 0.015196707 | 0.688026756 |
| 0.108077888 | 0.001888205 |
| 0.61384106 | 1.111E-06 |
| 0.001052582 | 1.97519E-05 |
| 7.2933E-07 | 1 |
| 0.025324502 | 0.133555566 |
| 0.022106824 | 0.353621377 |
| 1 | 0.000278729 |
| 0.668222651 | 0.1587219 |
| 0.67007816 | 5.61634E-05 |
| 0.172320959 | 4.87082E-11 |
| 0.105724886 | 0.001276167 |
| 0.003788532 | 0.620646907 |
| 0.408542478 | 0.469811302 |
| 0.834108201 | 1 |
| 1 | 0.957081292 |
| 0.611740303 | 0.146759032 |
| 0.566726681 | 0.900775927 |
| 1 | 1 |
| 0.000410399 | 1 |
| 0.24436374 | 1 |
| 0.033043323 | 0.123546784 |
| 1 | 1 |
| 1 | 1 |
| 0.666413336 | 0.536135575 |
| 0.334476393 | 1 |
| 0.693602582 | 0.095894807 |
| 0.206484942 | 0.032111437 |
| 0.954321801 | 1 |
| 0.11862999 | 1 |
| 0.004831841 | 4.35469E-11 |
| 7.31786E-09 | 0.000180089 |
| 1.65531E-05 | 2.93774E-08 |
| 1 | 1.00007E-10 |
| 0.254615346 | 1.41376E-07 |
| 1.73069E-10 | 0.013846179 |
| 9.3192E-06 | 1 |
| 0.000208614 | 1 |
| 0.749600013 | 5.20662E-11 |
| 0.944843265 | 9.72174E-12 |
| 0.592648116 | 9.72174E-12 |
| 4.10288E-05 | 0.108227703 |
| 1 | 1 |
| 1 | 1.42809E-09 |
| 0.421075826 | 9.72174E-12 |
| 0.000123313 | 1 |
| 0.254486412 | 1 |
| 1 | 1 |
| 0.007076179 | 0.884091062 |
| 0.43500357 | 1 |
| 1 | 1 |
| 1 | 0.99689597 |
| 0.000229119 | 5.79352E-10 |
| 0.004830125 | 0.07725666 |
| 0.016440307 | 1.18953E-11 |
| 2.88514E-05 | 0.73508552 |
| 0.150311442 | 2.22424E-06 |
| 1 | 3.0824E-07 |
| 4.72084E-05 | 0.000111007 |
| 0.909486978 | 1 |
| 0.881564938 | 0.122926293 |
| 1 | 0.100755976 |
| 0.000869019 | 0.036200405 |
| 1 | 0.019995579 |
| 0.000355854 | 0.012187832 |
| 7.22921E-08 | 0.02086479 |
| 0.001749462 | 0.810776629 |
| 7.28375E-07 | 0.012932791 |
| 9.82309E-07 | 8.44018E-06 |
| 2.36494E-06 | 0.029201008 |
| 7.93943E-05 | 9.72174E-12 |
| 0.072775786 | 0.608677136 |
| 0.000131907 | 8.59231E-05 |
| 0.001744574 | 1 |
| 0.00022895 | 3.01416E-07 |
| 0.000983919 | 9.8938E-12 |
| 0.669768999 | 0.638985357 |
| 0.031182962 | 2.65716E-08 |
| 1 | 0.799354546 |
| 0.015649984 | 0.173123883 |
| 0.708322051 | 1 |
| 0.000225389 | 0.247146854 |
| 0.018170186 | 1 |
| 0.007346941 | 4.15833E-05 |
| 3.38015E-08 | 4.26974E-07 |
| 4.81704E-05 | 1 |
| 1.71681E-07 | 1.16547E-08 |
| 0.054209275 | 1 |
| 1 | 0.533135599 |
| 3.98793E-09 | 8.30729E-08 |
| 6.59116E-07 | 4.5595E-11 |
| 3.8088E-05 | 4.27224E-11 |
| 1.0794E-06 | 9.72174E-12 |
| 1.24847E-08 | 0.00087713 |
| 0.016452785 | 1 |
| 0.000595755 | 0.000163288 |
| 3.92821E-06 | 0.000104917 |
| 2.14745E-07 | 0.000260867 |
| 2.09889E-07 | 0.005788093 |
| 2.38485E-07 | 1 |
| 8.31567E-06 | 0.002042015 |
| 0.003678654 | 2.17217E-05 |
| 1.97006E-06 | 0.42715512 |

| Adj. P-value: (CM_neg_newanalysis_12022024) / (SM_neg_newanalysis_12022024) | Adj. P-value: (Bb_neg_newanalysis_12022024) / (WATER_NEG_NEV |
| --- | --- |
| 1 | 0.840837853 |
| 1 | 1 |
| 1 | 1 |
| 1 | 0.001898751 |
| 1 | 0.012222059 |
| 1 | 0.827988817 |
| 1 | 0.445749271 |
| 1 | 1 |
| 1 | 1 |
| 1 | 0.947517779 |
| 1 | 1 |
| 1 | 0.907731863 |
| 1 | 1 |
| 1 | 1 |
| 1 | 0.589237952 |
| 1 | 0.79266544 |
| 1 | 1 |
| 1 | 0.747822983 |
| 1 | 6.64662E-08 |
| 1 | 1 |
| 1 | 1 |
| 1 | 0.747822983 |
| 1 | 1 |
| 1 | 1 |
| 1 | 1 |
| 0.124693722 | 1 |
| 1 | 0.717954028 |
| 0.349701127 | 1 |
| 1 | 1 |
| 0.431338875 | 1 |
| 1 | 1 |
| 0.4450931 | 1 |
| 1 | 1 |

|  |  |  |
| --- | --- | --- |
|  | 1 | 5.19357E-07 |
|  | 1 | 0.027358822 |
|  | 1 | 1.40017E-05 |
|  | 1 | 8.00516E-07 |
|  | 1 | 0.000566673 |
|  | 1 | 5.27828E-11 |
|  | 1 | 2.6414E-07 |
|  | 1 | 0.772989589 |
|  | 1 | 5.3807E-11 |
|  | 1 | 2.94944E-11 |
|  | 1 | 1.17011E-05 |
|  | 1 | 6.6976E-06 |
|  | 1 | 1 |
|  | 1 | 0.782873861 |
|  | 1 | 0.9392865 |
|  | 1 | 0.824036842 |
|  | 1 | 1 |
|  | 1 | 1 |
| 0.803813347 | 1 | 1 |
|  | 1 | 1 |
|  | 1 | 1 |
|  | 1 | 1 |
|  | 1 | 0.130117606 |
|  | 1 | 0.540222214 |
|  | 1 | 1 |
|  | 1 | 0.993766112 |
|  | 1 | 1 |
| 0.349865373 | 1 | 1 |
|  | 1 | 0.842009346 |
|  | 1 | 0.762383042 |
|  | 1 | 0.897224176 |
|  | 1 | 0.788180989 |
|  | 1 | 0.002023419 |
|  | 1 | 1 |
|  | 1 | 0.001553052 |
|  | 1 | 1 |
|  | 1 | 1 |
| 0.270336405 |  | 1 |
|  | 1 | 0.31956411 |
|  | 1 | 0.725642831 |
|  | 1 | 1 |
|  | 1 | 1 |
|  | 1 | 1 |
|  | 1 | 1 |
|  | 1 | 1 |
|  | 1 | 1 |
|  | 1 | 1 |
|  | 1 | 1 |
|  | 1 | 1 |
|  | 1 | 1 |
|  | 1 | 1 |
|  | 1 | 1 |
|  | 1 | 1 |
|  | 1 | 1 |
|  | 1 | 1 |
|  | 1 | 1 |
|  | 1 | 1 |
|  | 1 | 1 |
|  | 1 | 0.04768475 |
|  | 1 | 0.782260932 |
|  | 1 | 1 |
|  | 1 | 0.967237017 |
|  | 1 | 0.884185576 |
|  | 1 | 1 |
|  | 1 | 1 |
|  | 1 | 1 |
|  | 1 | 0.852321508 |
|  | 1 | 0.77316027 |
|  | 1 | 1 |
|  | 1 | 4.98451E-05 |
| 0.660459677 |  | 1 |
|  | 1 | 1 |
|  | 1 | 1 |
|  | 1 | 0.770170684 |

|  |  |
| --- | --- |
| 1 | 1 |
| 0.791408165 | 0.795568956 |
| 1 | 1 |
| 1 | 0.793181028 |
| 1 | 1 |
| 1 | 1 |
| 1 | 0.976105918 |
| 1 | 1 |
| 1 | 1 |
| 1 | 1 |
| 1 | 0.980279903 |
| 1 | 1 |
| 1 | 1 |
| 1 | 1.24945E-07 |
| 1 | 2.65677E-11 |
| 0.857255267 | 2.65677E-11 |
| 1 | 1 |
| 1 | 1 |
| 1 | 1.35317E-05 |
| 1 | 2.65677E-11 |
| 1 | 1 |
| 1 | 0.747822983 |
| 1 | 1 |
| 1 | 0.579782433 |
| 1 | 1 |
| 1 | 1 |
| 1 | 1 |
| 0.049282221 | 0.93082703 |
| 1 | 0.672629034 |
| 0.046052278 | 0.935393927 |
| 1 | 1 |
| 1 | 1 |
| 0.671243329 | 0.937040814 |
| 1 | 0.657822441 |
| 1 | 1 |

|  |  |  |
| --- | --- | --- |
|  | 1 | 0.747822983 |
|  | 1 | 1 |
|  | 1 | 1 |
|  | 1 | 1 |
|  | 1 | 0.531791267 |
|  | 1 | 1 |
|  | 1 | 1 |
|  | 1 | 0.945891494 |
|  | 1 | 0.921869934 |
|  | 1 | 0.179027766 |
|  | 1 | 0.000148004 |
|  | 1 | 1 |
|  | 1 | 0.546297177 |
|  | 1 | 1 |
|  | 1 | 1 |
|  | 1 | 0.640427919 |
|  | 1 | 0.806861921 |
|  | 1 | 0.032475097 |
|  | 1 | 1 |
|  | 1 | 1 |
|  | 1 | 1 |
|  | 1 | 0.762383042 |
|  | 1 | 1 |
|  | 1 | 1 |
|  | 1 | 0.815170383 |
|  | 1 | 1 |
|  | 1 | 0.348026736 |
|  | 1 | 1 |
|  | 1 | 1 |
|  | 1 | 0.839202174 |
|  | 1 | 1 |
|  | 1 | 0.183840061 |
|  | 1 | 1 |
|  | 1 | 0.819913047 |
| 0.552112614 |  | 1 |

|  |  |
| --- | --- |
| 1 | 1 |
| 1 | 1 |
| 1 | 1 |
| 1 | 0.837991924 |
| 1 | 1 |
| 1 | 1 |
| 1 | 1 |
| 1 | 0.929867706 |

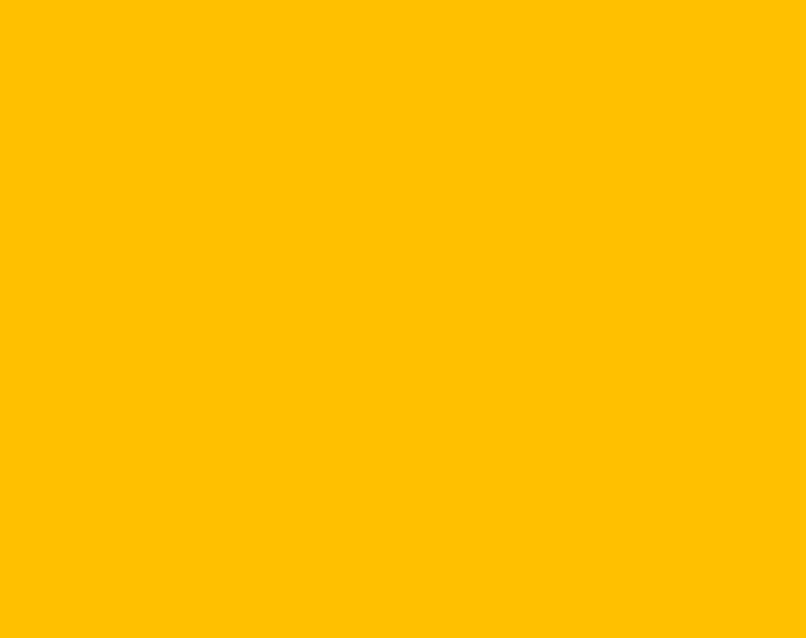

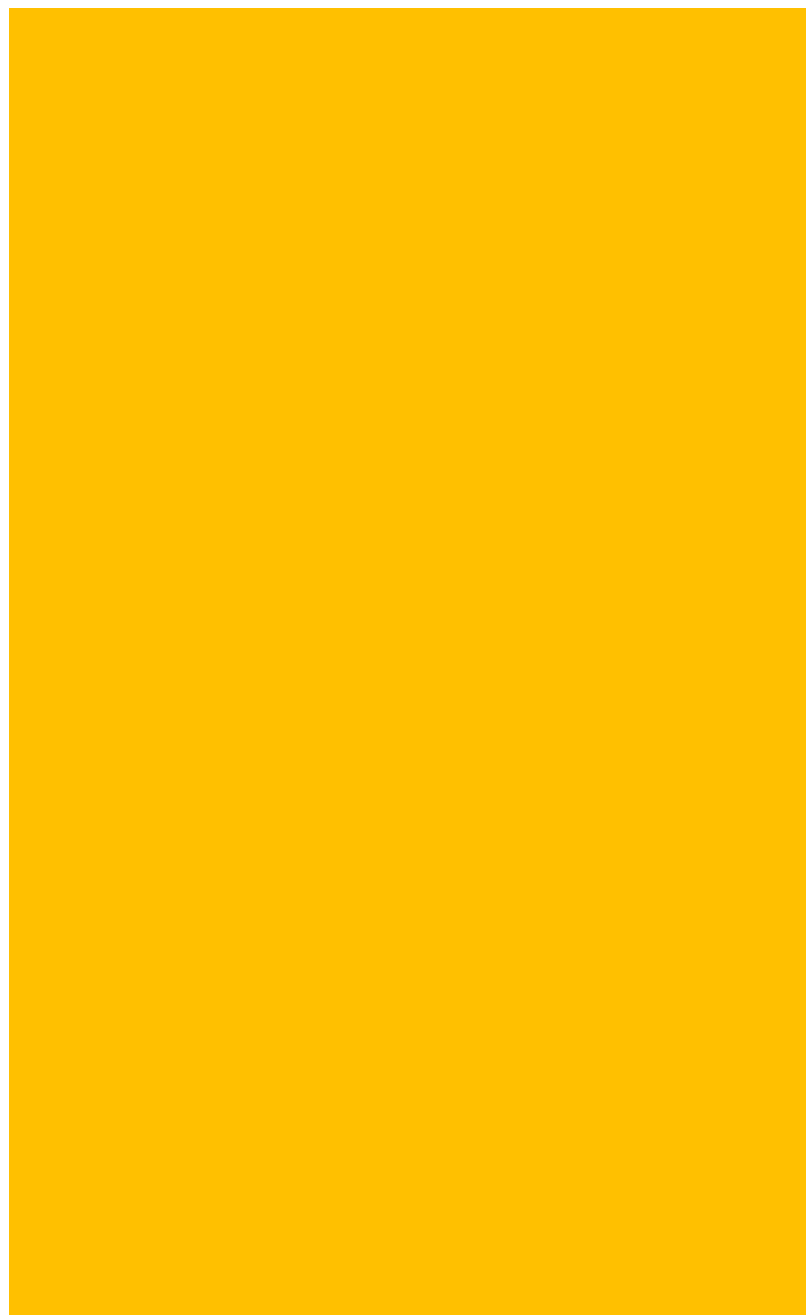

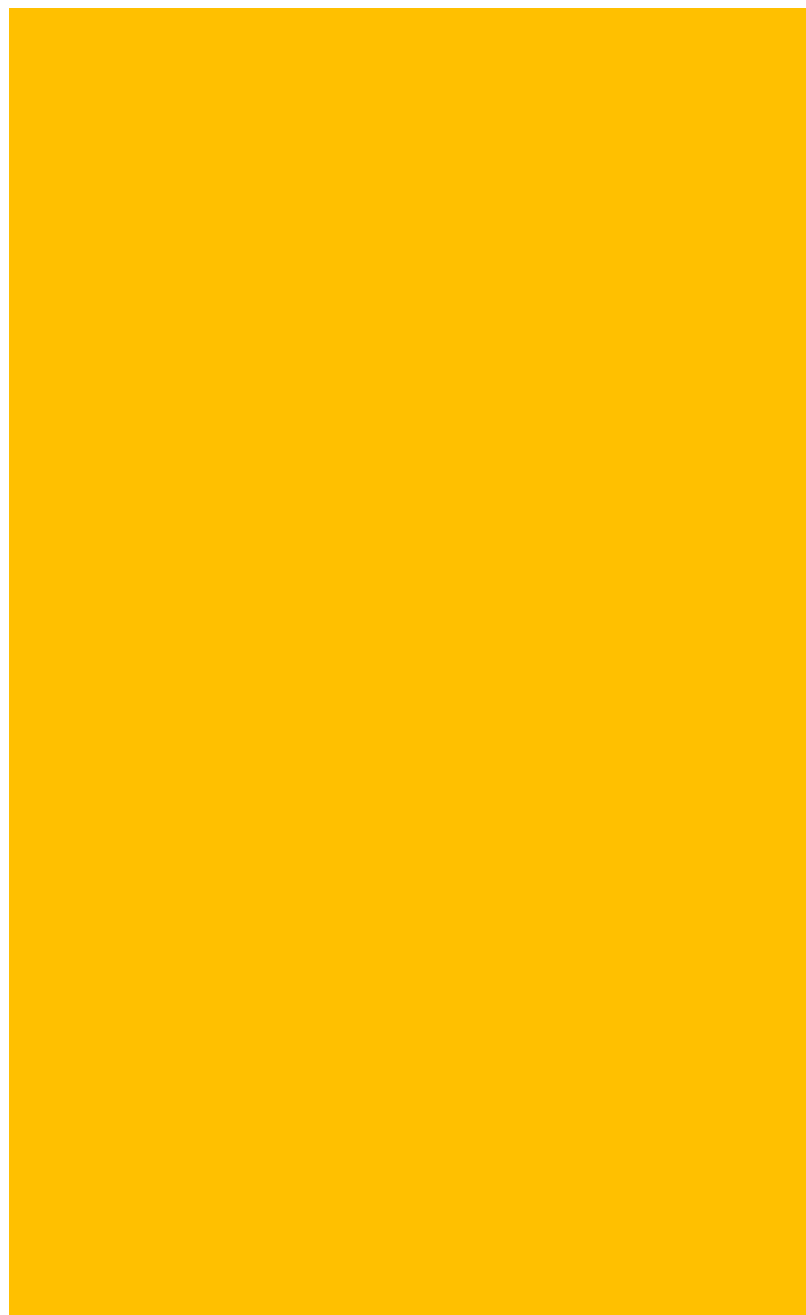

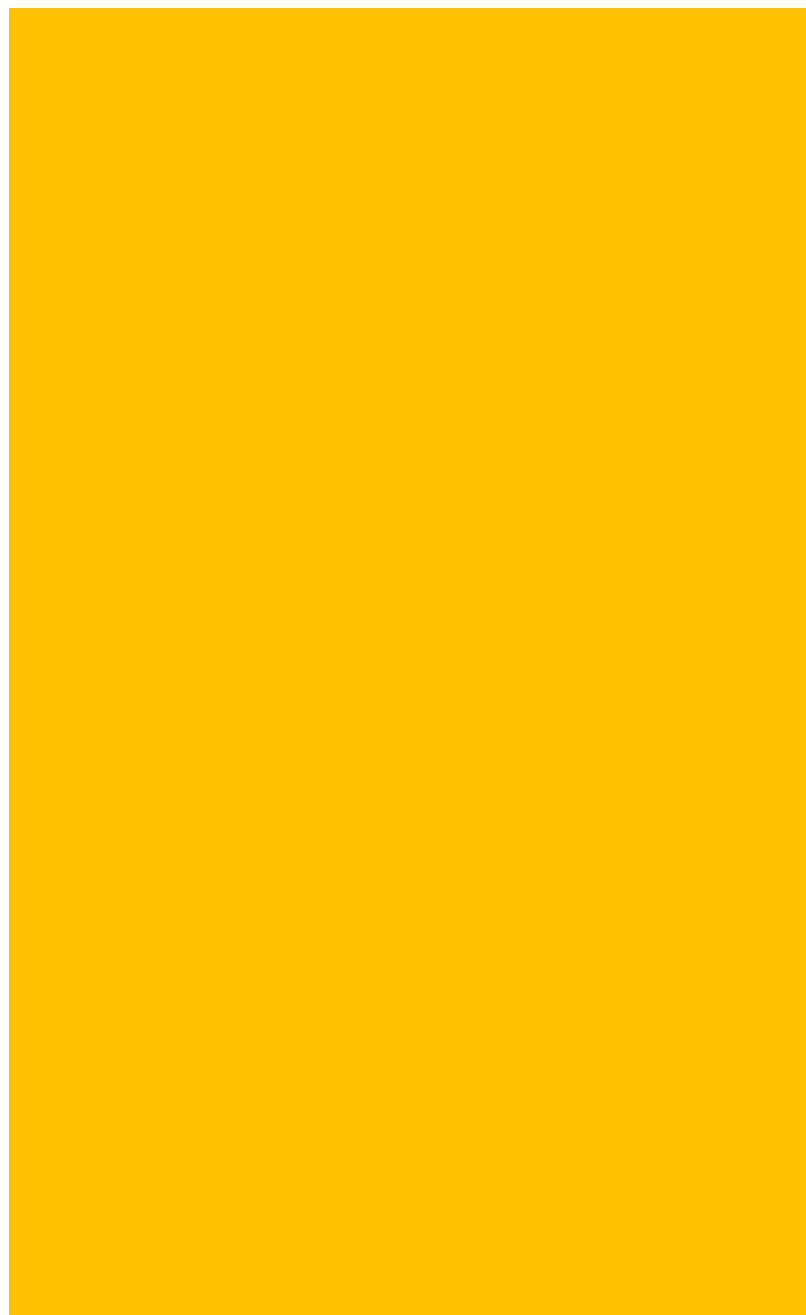

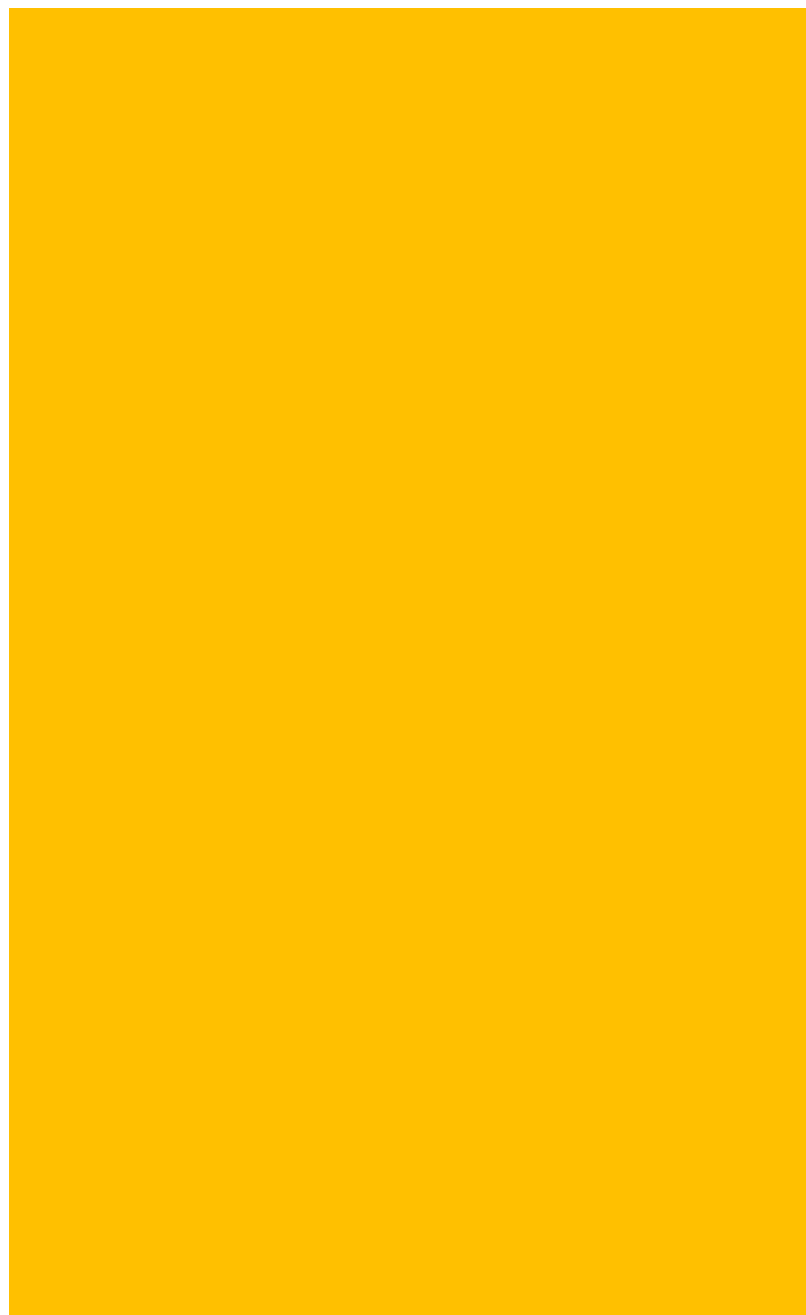

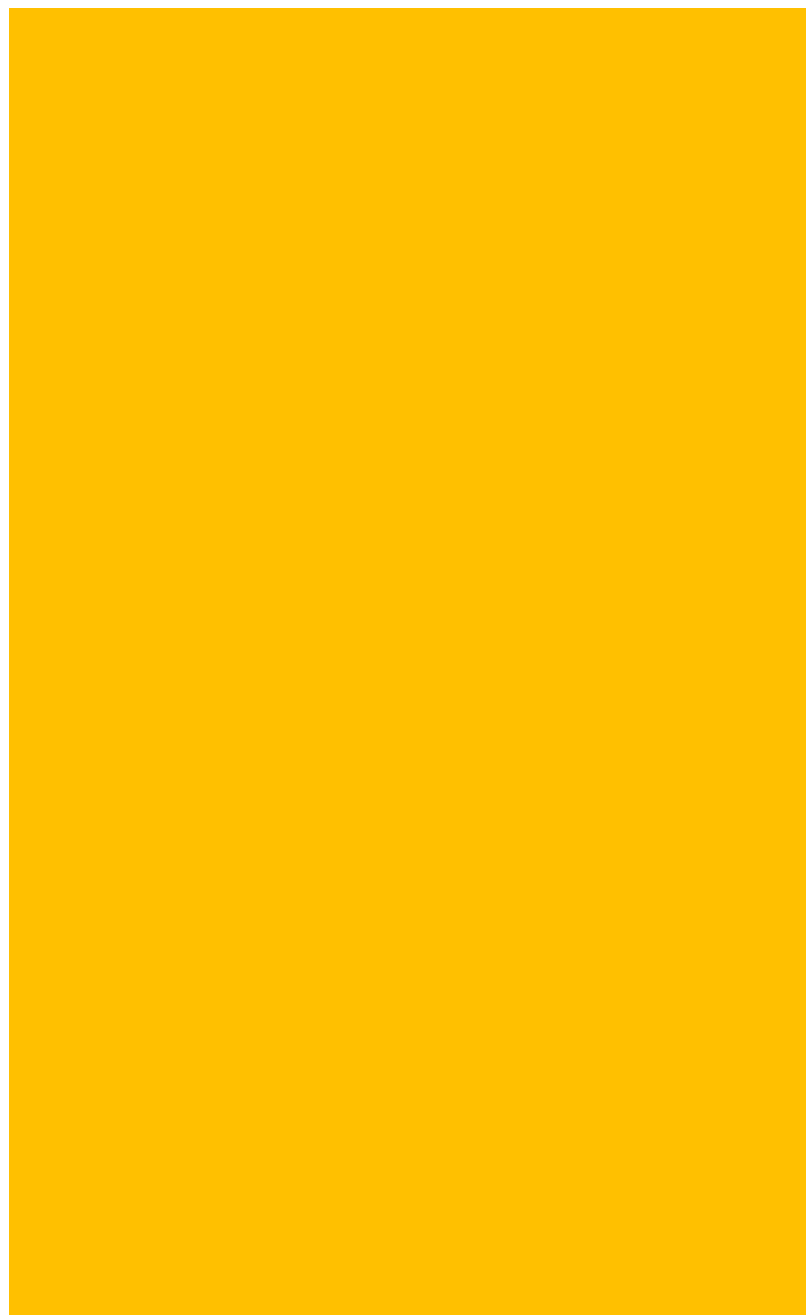

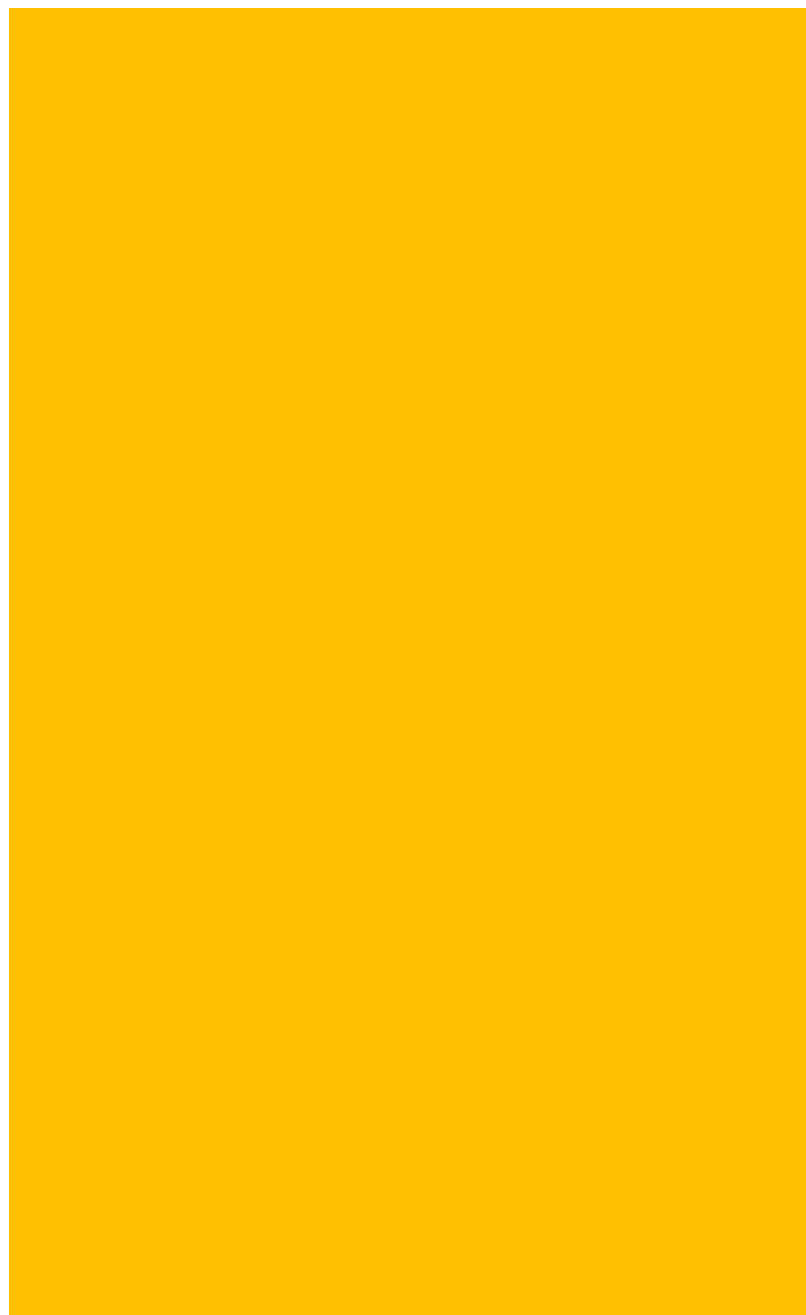

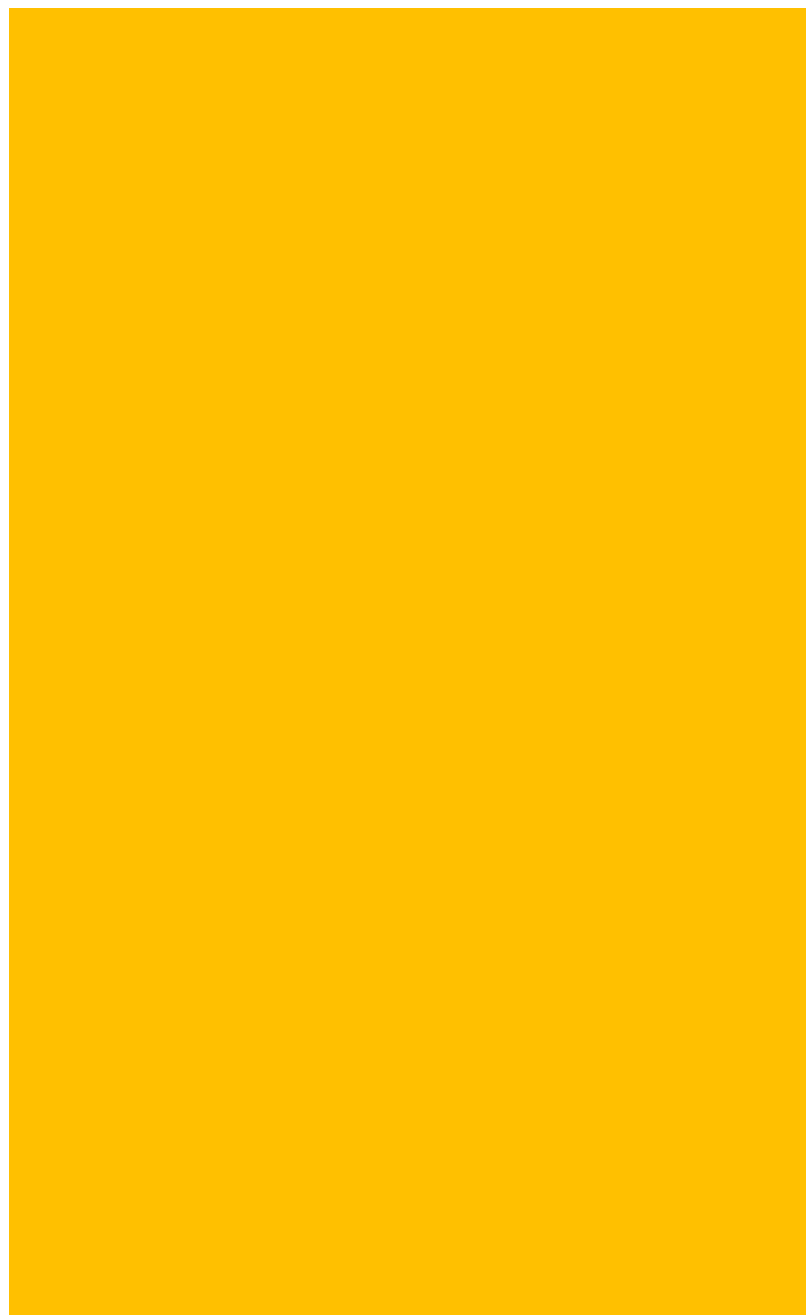

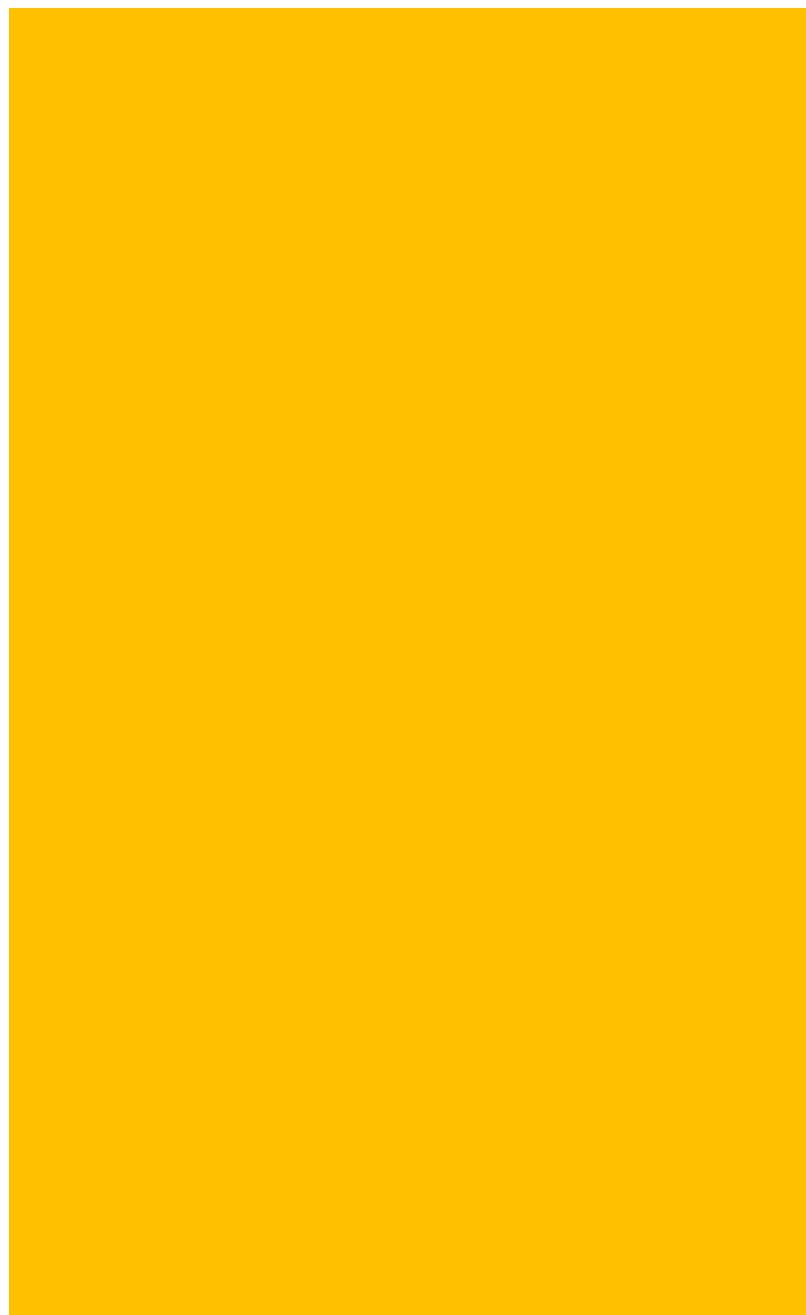

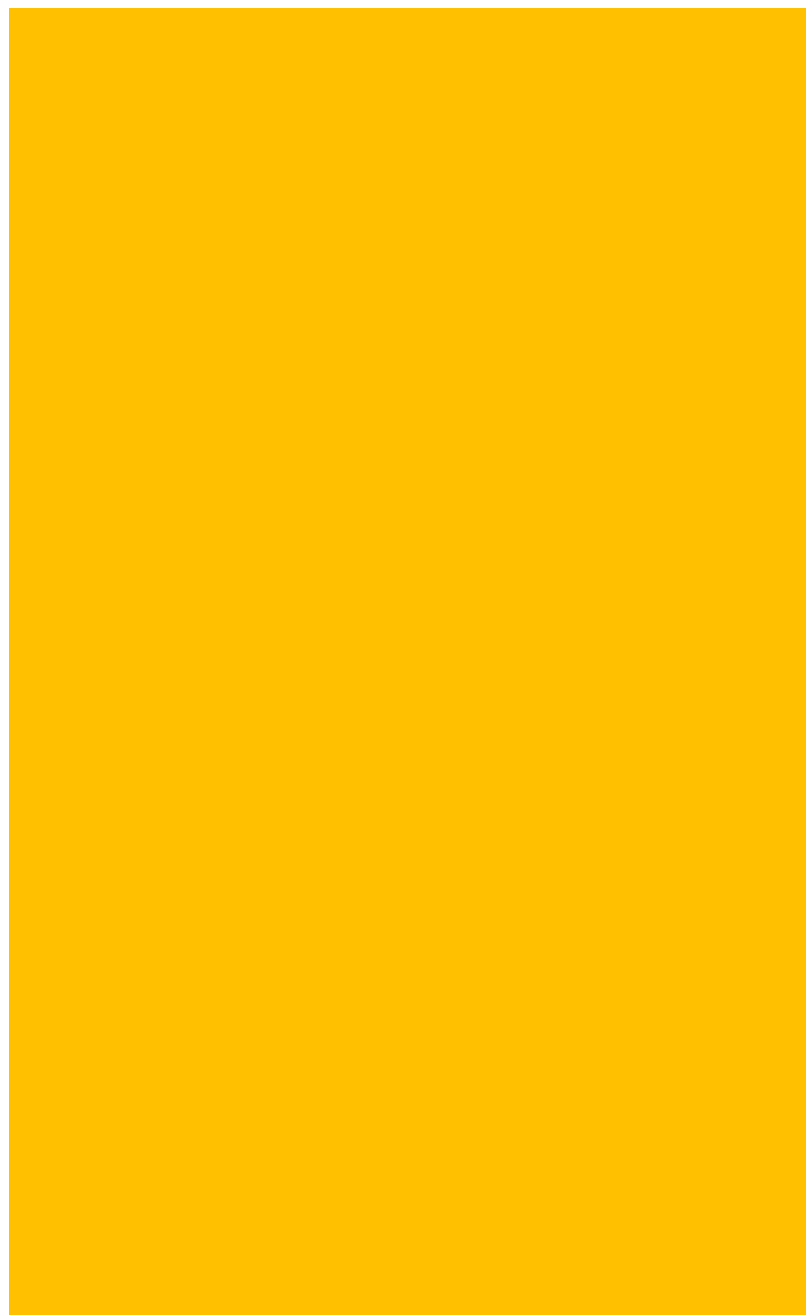

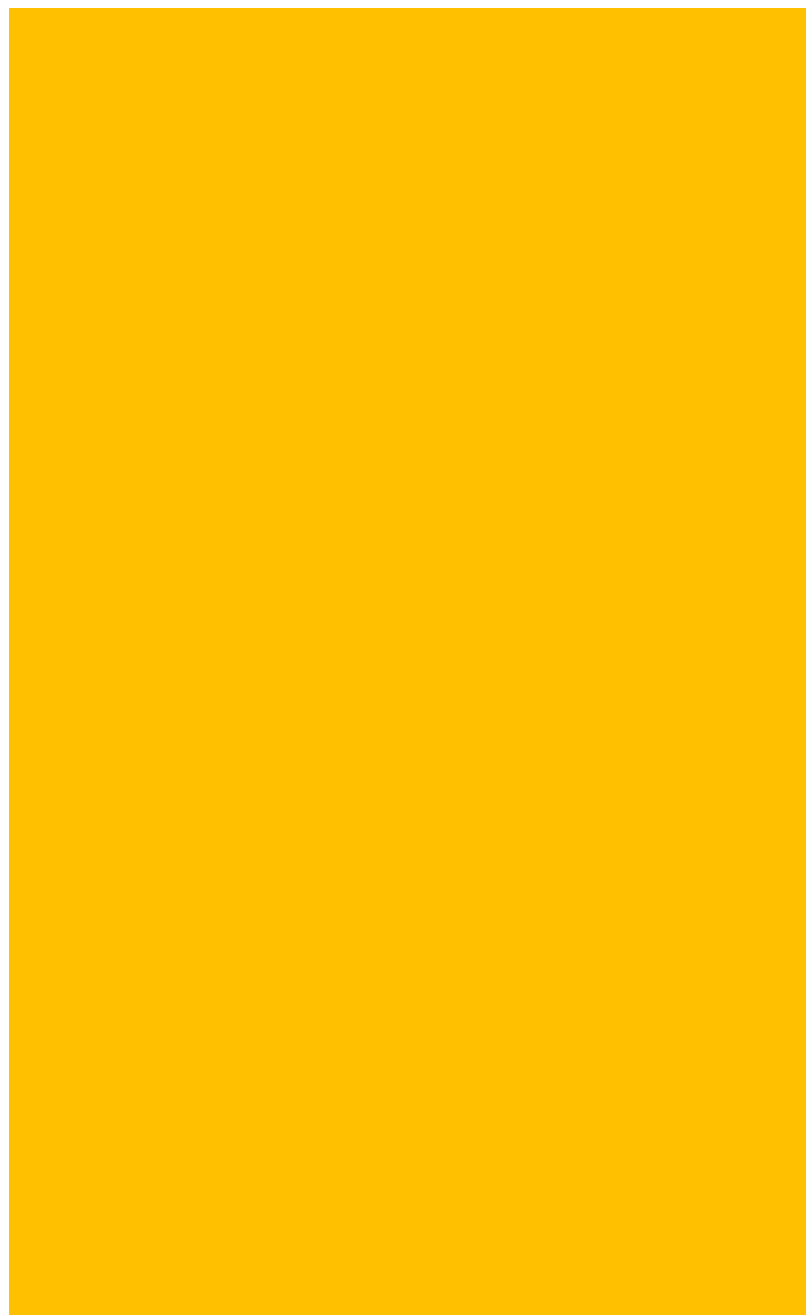

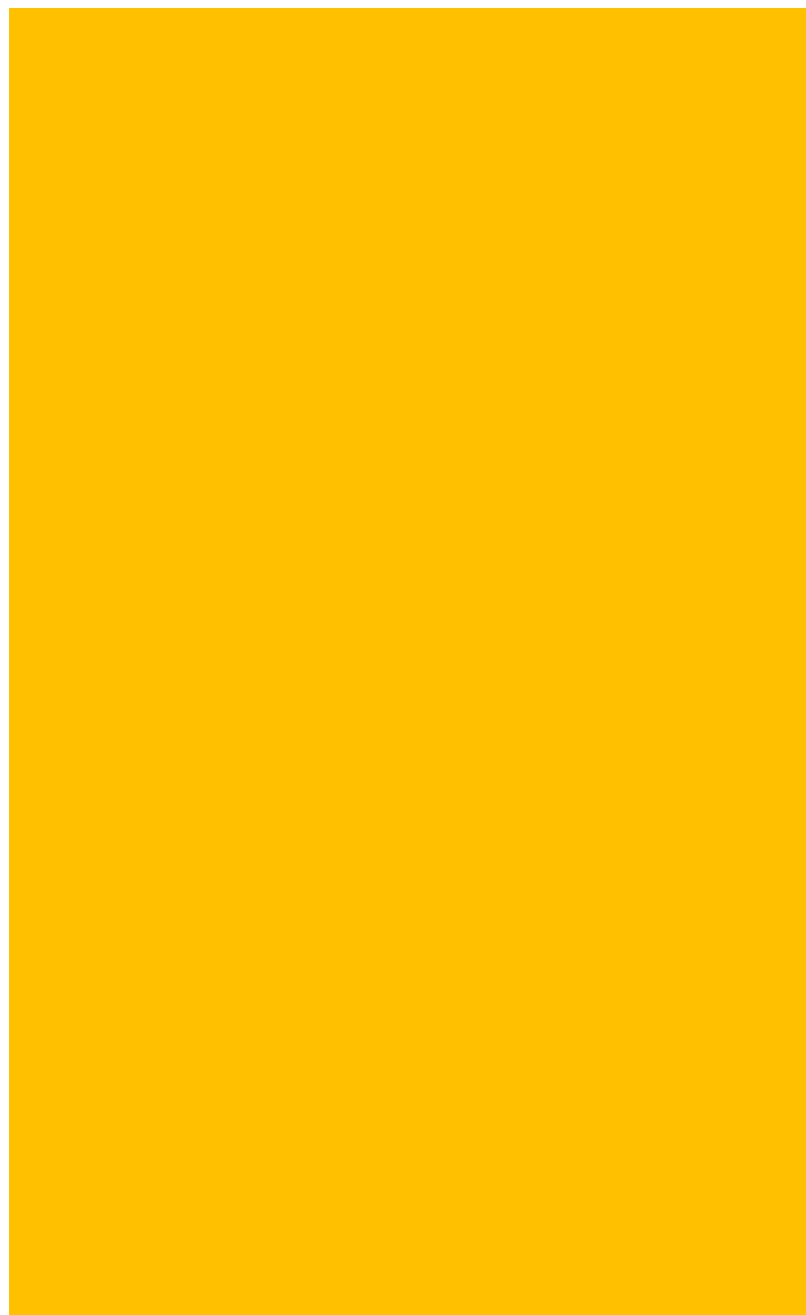

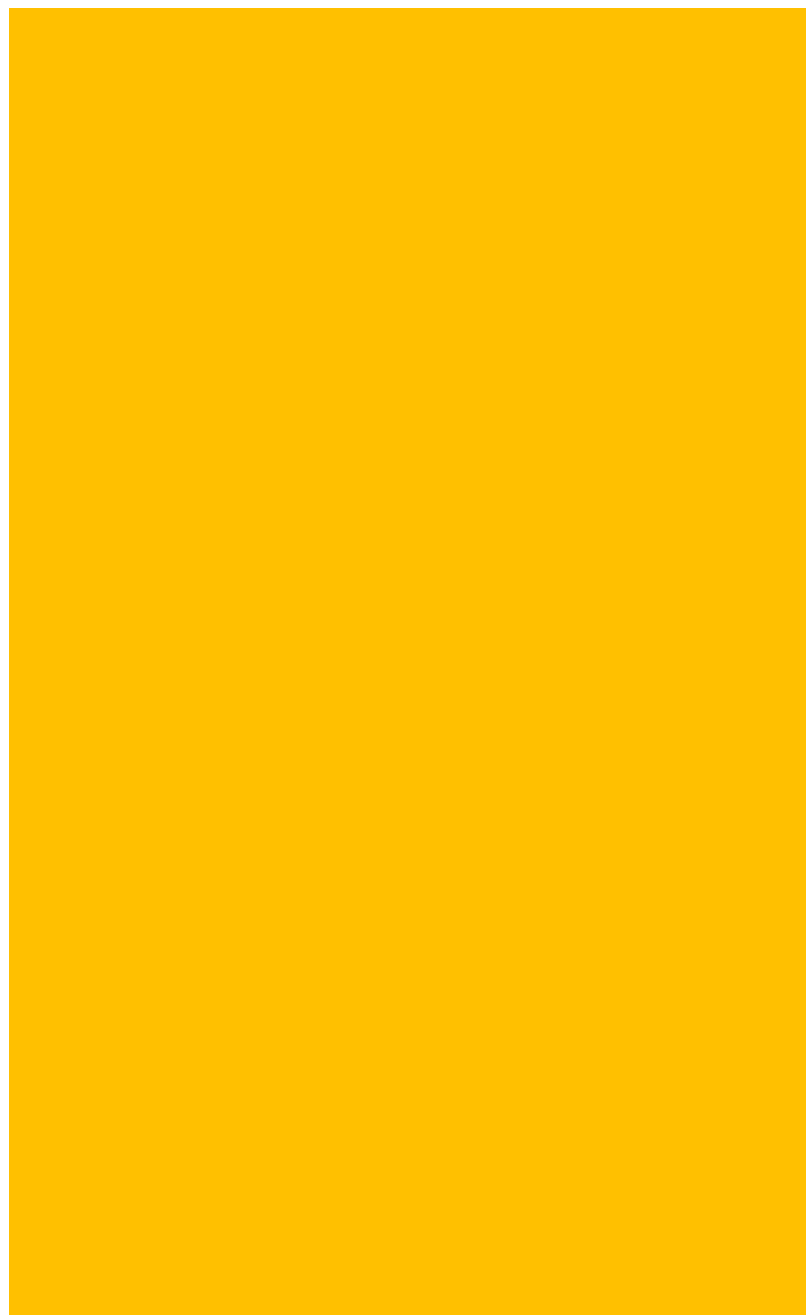

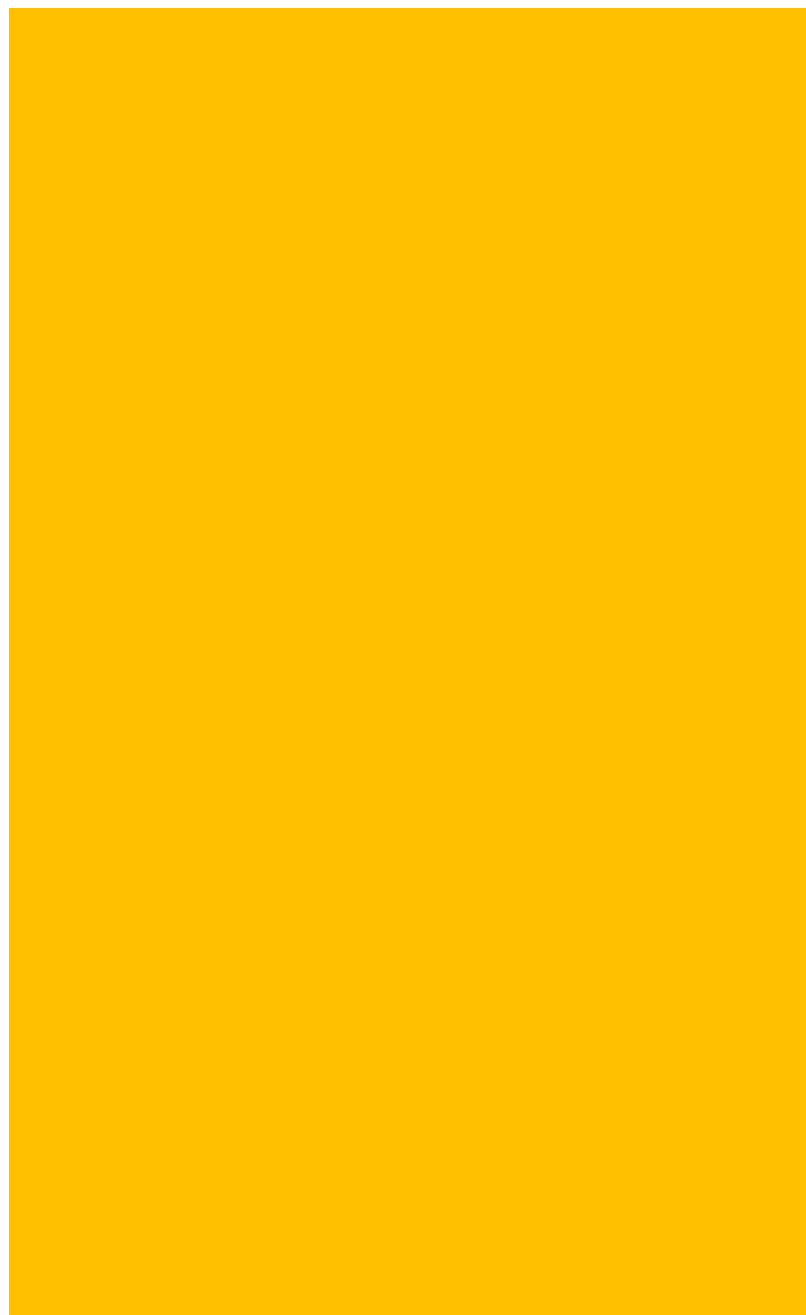

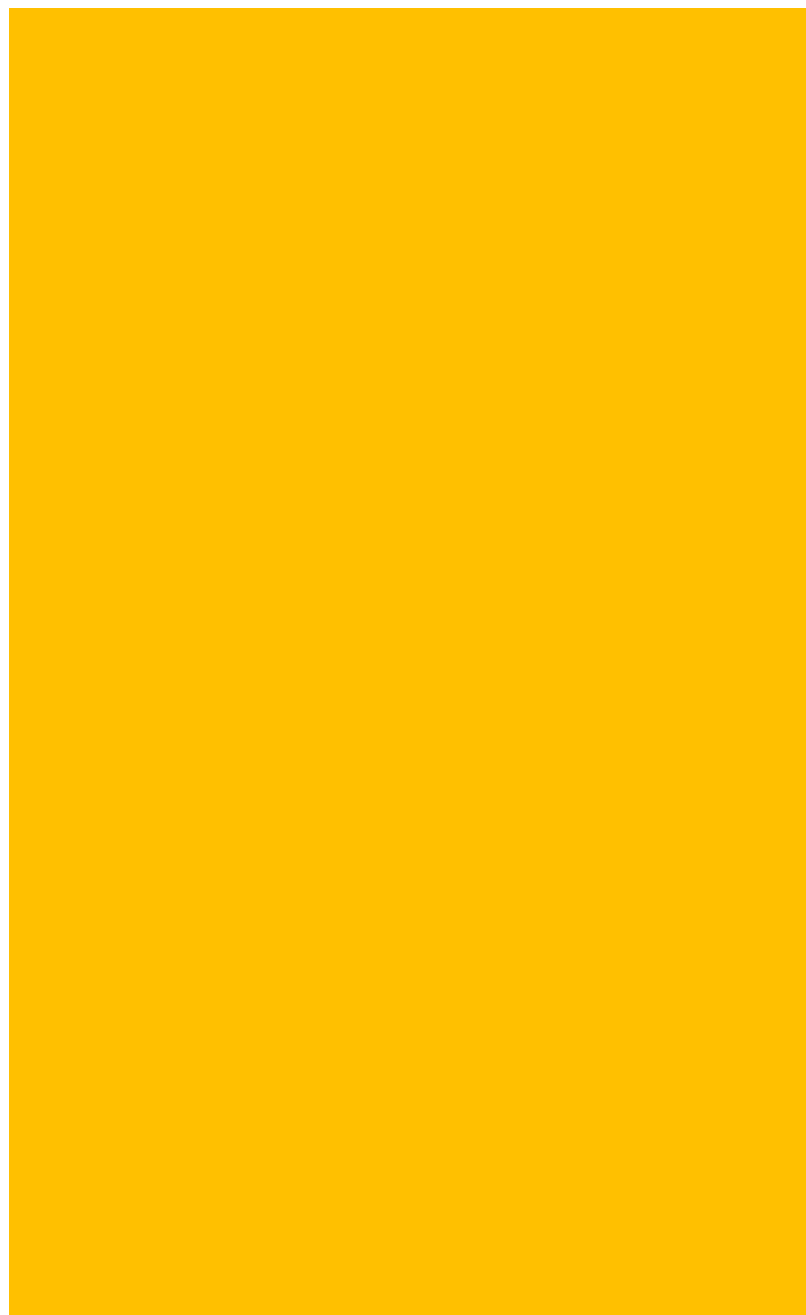

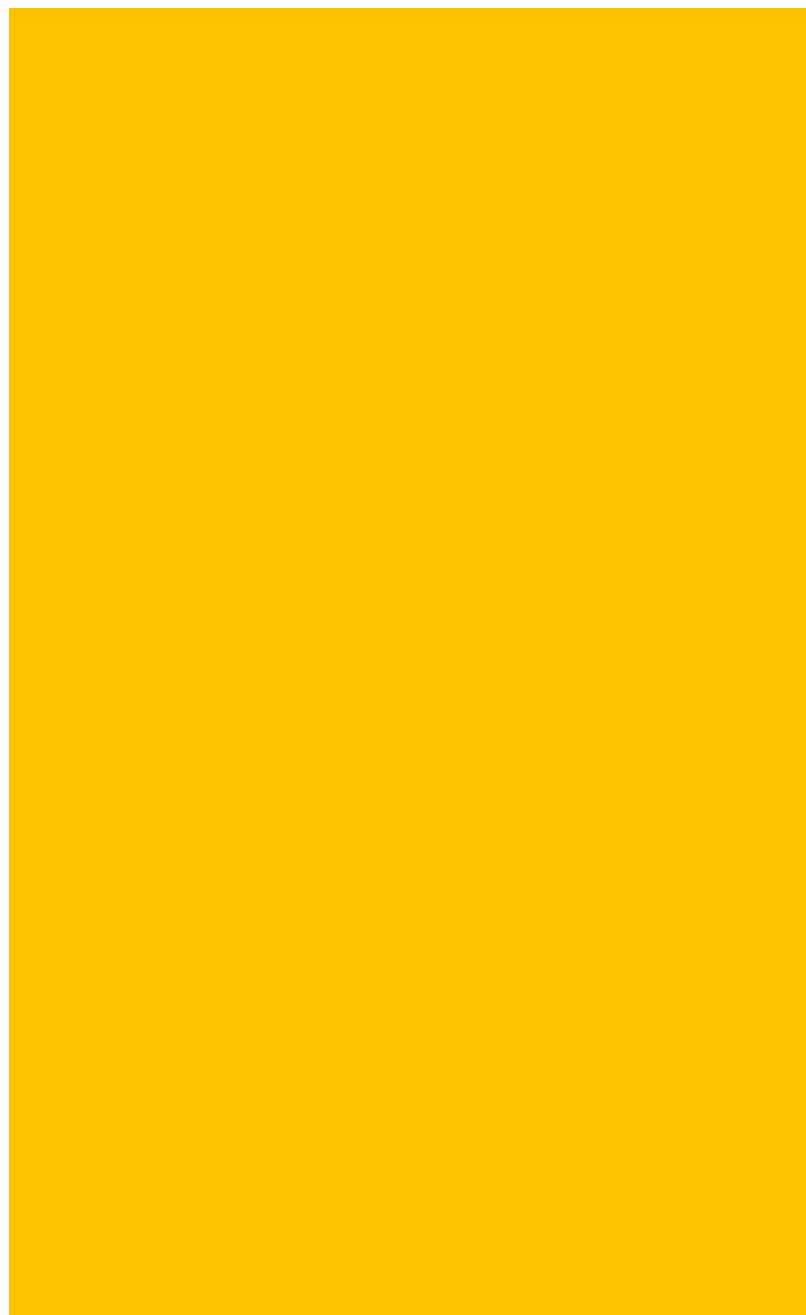

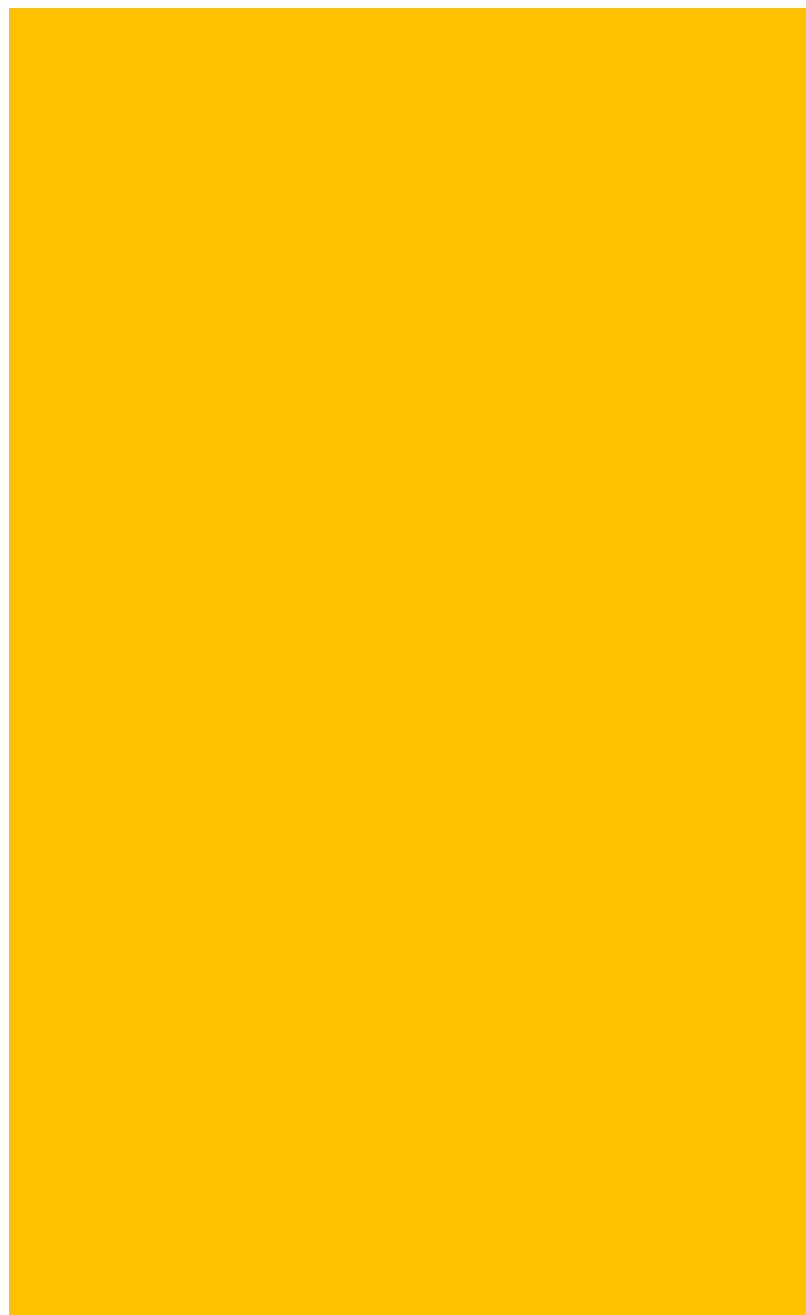

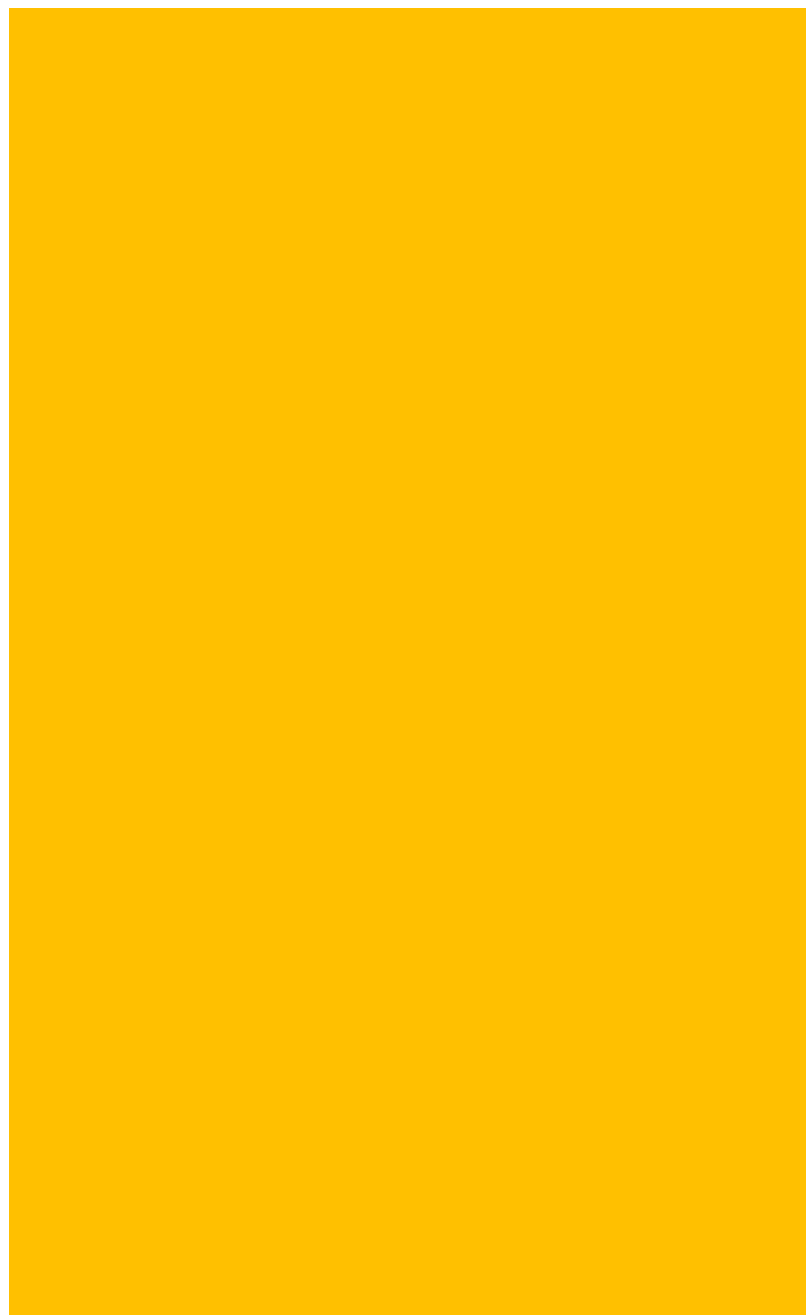

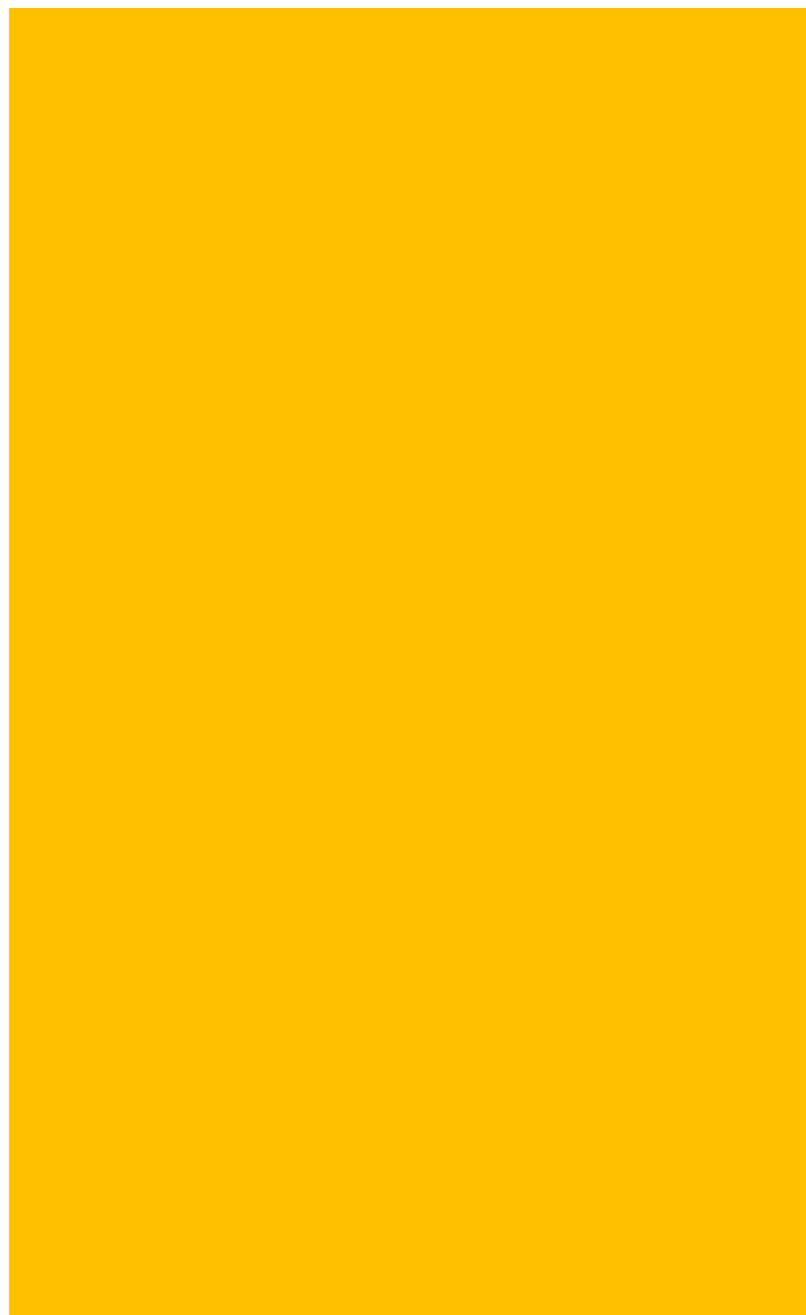

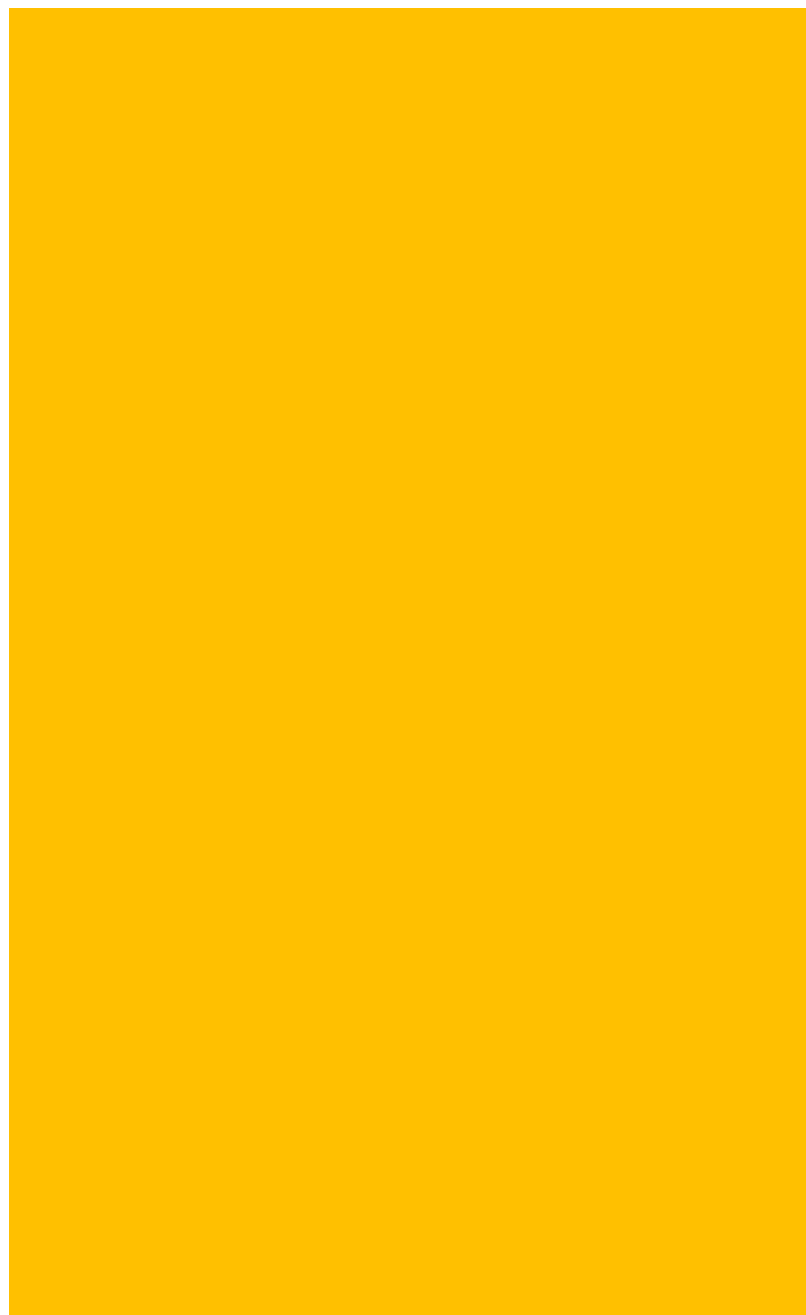

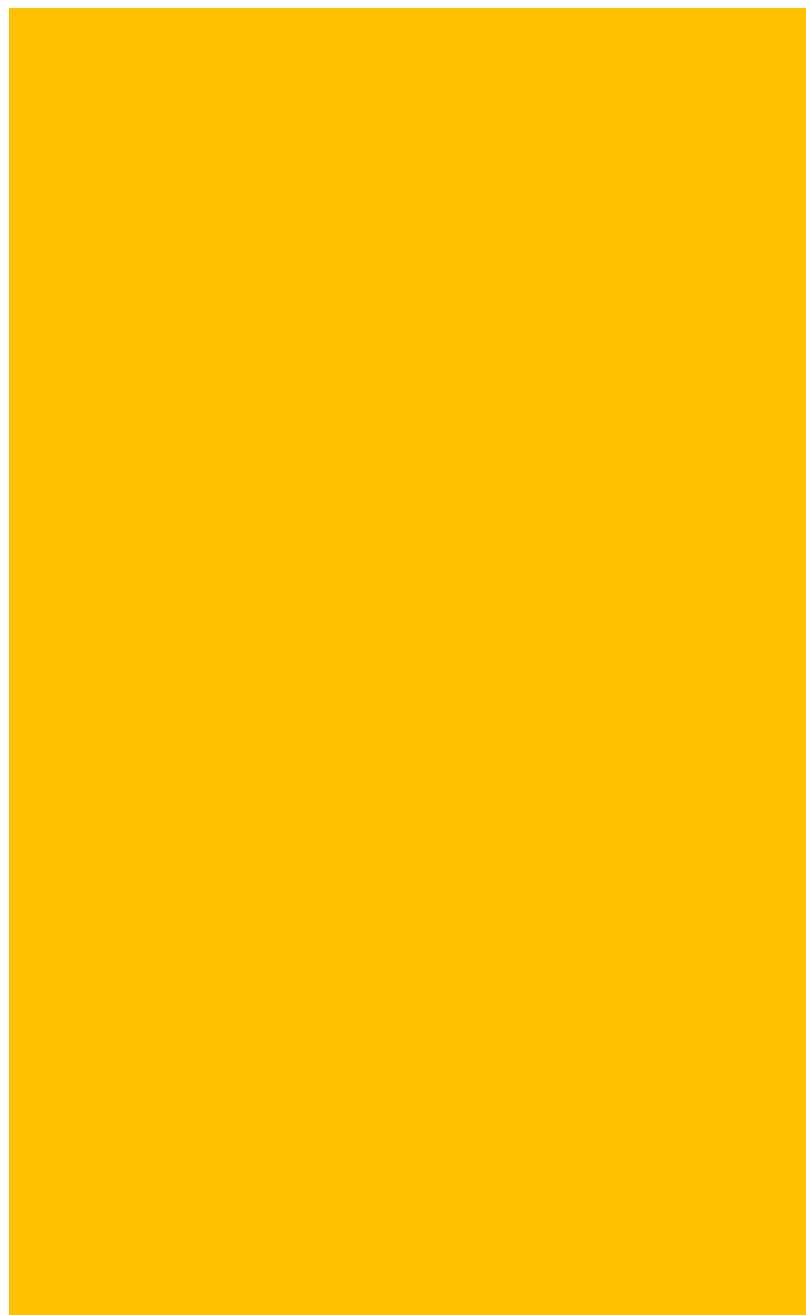

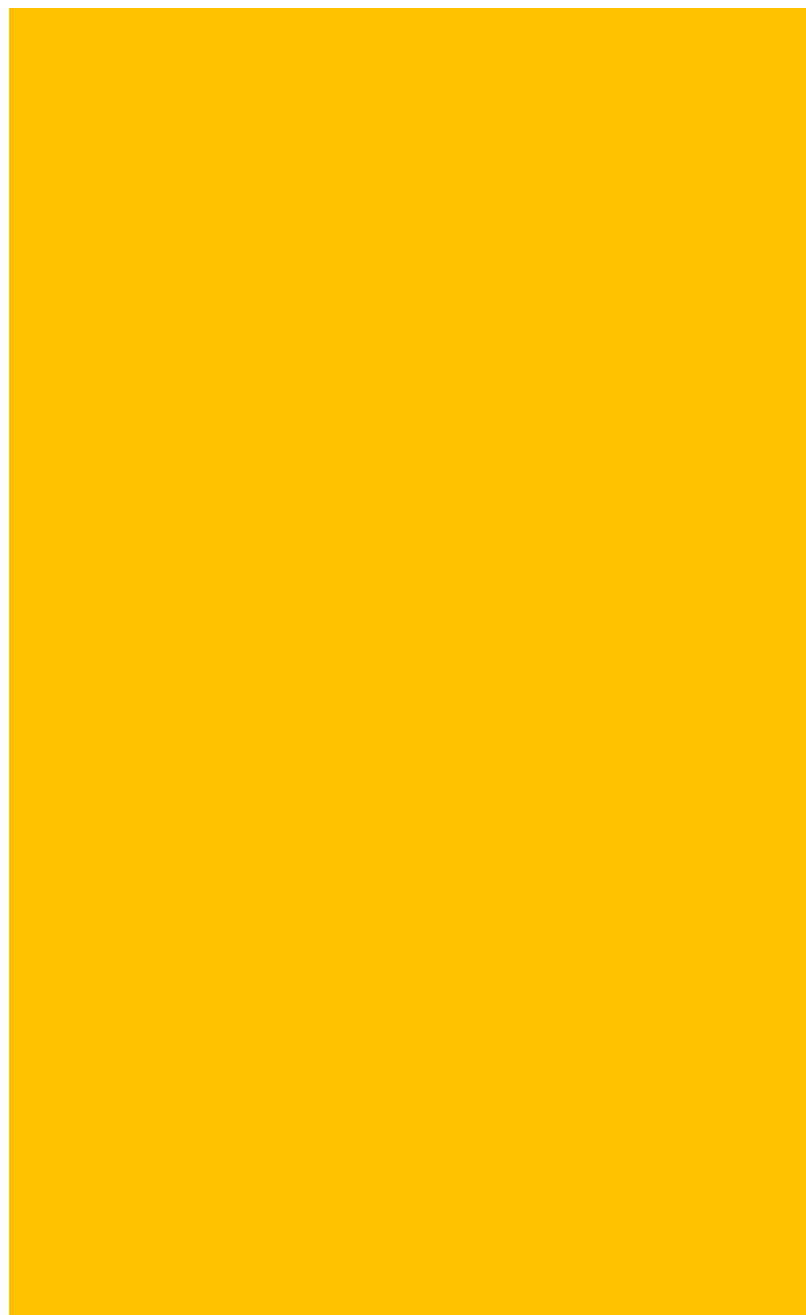

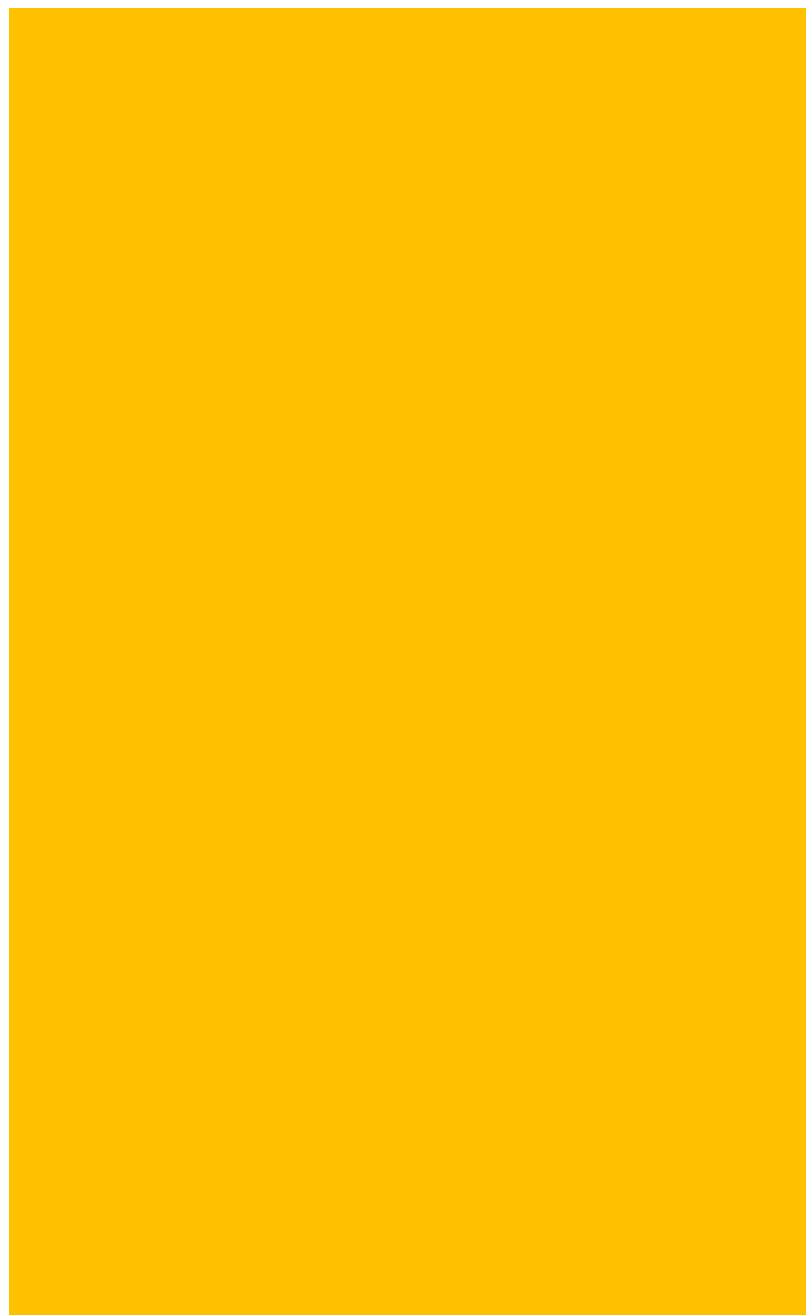

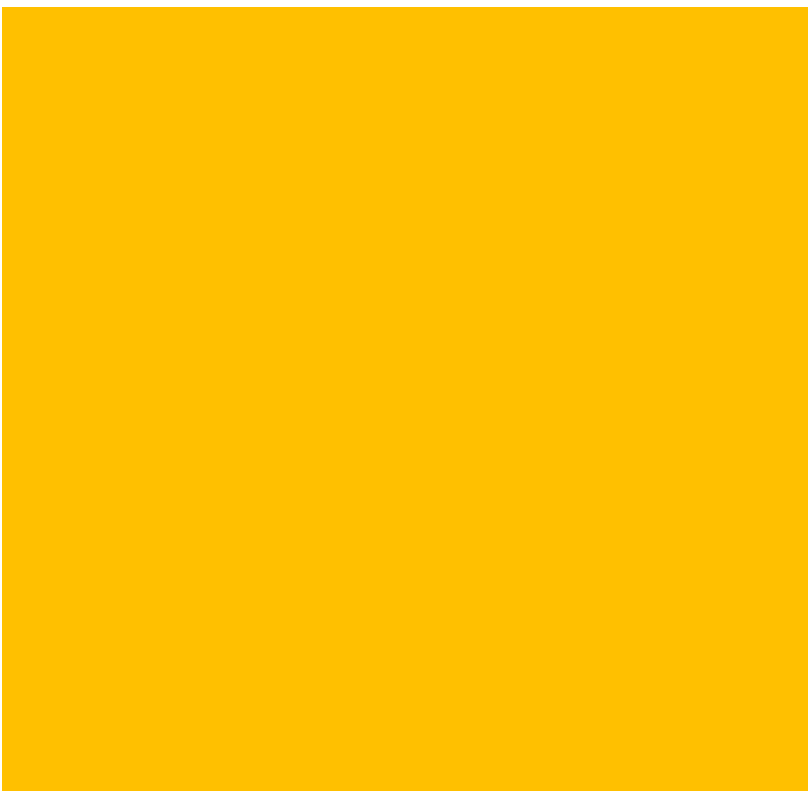

| Adj. P-value: (CM_neg_newanalysis_12022024) / (WATER_NEG_NEWANALYSIS_12022024) | Adj. P-value: (RS_NEG_NEWANALYSIS_12022024) / (WATER_NEG_NEWANALYSIS_12022024) |
| --- | --- |
| 0.735100571 | 7.35869E-06 |
| 1 | 4.86255E-07 |
| 1 | 0.000122891 |
| 0.04000326 | 2.07636E-06 |
| 0.181213954 | 1.2948E-06 |
| 1 | 0.802425808 |
| 2.29343E-09 | 9.25864E-10 |
| 0.000142293 | 1.88888E-05 |
| 0.005756991 | 0.001524677 |
| 1.9543E-07 | 3.93637E-07 |
| 2.5002E-06 | 0.00034499 |
| 0.985320045 | 1 |
| 0.310503681 | 0.125999299 |
| 0.222520365 | 0.002999122 |
| 0.047322862 | 0.000380444 |
| 0.638561423 | 0.004649258 |
| 0.479956303 | 0.991193629 |
| 1 | 1.10163E-06 |
| 1 | 0.000389735 |
| 1 | 0.105547855 |
| 1 | 0.962804149 |
| 0.154419344 | 1.36058E-06 |
| 1 | 0.070571698 |
| 0.208665194 | 2.91098E-05 |
| 1 | 1 |
| 0.015763938 | 0.000876388 |
| 0.004979188 | 0.000218644 |
| 0.127016152 | 0.035823185 |
| 0.386153686 | 0.0002973 |
| 1 | 1 |
| 0.154419344 | 0.029041798 |
| 0.369466983 | 0.351075791 |
| 0.030314213 | 2.0506E-10 |
| 1 | 1 |
| 1 | 1 |
| 1 | 0.377547438 |
| 1 | 1 |
| 0.29069512 | 5.45879E-05 |
| 1 | 0.080030973 |
| 1 | 0.029021368 |
| 1 | 0.1672094 |
| 0.654704505 | 8.62021E-05 |
| 1 | 0.009572918 |
| 0.487657064 | 0.002701993 |
| 0.392408075 | 0.000875197 |
| 1 | 1.32826E-06 |
| 0.68468713 | 1 |
| 0.235534834 | 0.02829398 |
| 0.280804514 | 0.184615448 |
| 0.000205917 | 9.99209E-11 |
| 0.966634453 | 0.675286262 |
| 1 | 1 |
| 1 | 0.022238253 |
| 1 | 4.75061E-09 |
| 1 | 0.000448812 |
| 1 | 0.449736778 |
| 0.069665705 | 1.17185E-05 |
| 1 | 2.26488E-06 |
| 0.21013211 | 1.51217E-05 |
| 1 | 4.41266E-07 |
| 5.27412E-05 | 0.005868528 |
| 0.287925427 | 1.86629E-08 |
| 0.590159415 | 0.005743584 |
| 0.48254113 | 0.821863707 |
| 3.6028E-06 | 5.93261E-10 |
| 0.117507281 | 1.06454E-07 |
| 1 | 0.142532673 |
| 0.958950323 | 0.632940528 |
| 0.218303009 | 0.813724554 |
| 1 | 0.338202443 |
| 1 | 1.34663E-05 |
| 0.693430181 | 0.207649074 |
| 0.19455115 | 1 |
| 1 | 0.572532306 |
| 1 | 0.023746543 |
| 1 | 0.000523184 |
| 0.89381264 | 0.935864609 |
| 1 | 0.017033055 |
| 0.564085455 | 1 |
| 0.132054994 | 3.94458E-07 |
| 0.001213766 | 6.9627E-08 |
| 0.32413078 | 4.88001E-09 |
| 1 | 3.1584E-08 |
| 1 | 1 |
| 1 | 0.474515933 |
| 1 | 0.001615489 |
| 0.273353045 | 2.90643E-05 |
| 0.976880318 | 1 |
| 1.2276E-07 | 2.0953E-11 |
| 0.001193332 | 6.35914E-09 |
| 0.275795211 | 1 |
| 0.28860693 | 5.1621E-06 |
| 1 | 0.000999577 |
| 0.427474292 | 0.4247778 |
| 0.139205281 | 8.02661E-06 |
| 0.203512793 | 4.39805E-05 |
| 0.989428889 | 0.4220608 |
| 1 | 0.006372792 |
| 0.003714204 | 4.06101E-08 |
| 1 | 1 |
| 1 | 0.80723938 |
| 1 | 0.203237357 |
| 0.186216363 | 8.46846E-06 |
| 0.943136045 | 1 |
| 0.491591947 | 1 |
| 0.65443256 | 1 |
| 0.318058771 | 0.001059042 |
| 0.653383498 | 1 |
| 4.68881E-07 | 8.54557E-12 |
| 1 | 9.18133E-12 |
| 0.001742721 | 8.58883E-12 |
| 7.86516E-07 | 8.09813E-11 |
| 0.00181154 | 7.99329E-10 |
| 1 | 8.54557E-12 |
| 0.838380532 | 0.001207503 |
| 1 | 0.001202383 |
| 0.341672369 | 0.000275387 |
| 0.512063429 | 0.002708249 |
| 0.97537082 | 0.016927719 |
| 0.43734435 | 0.399420862 |
| 0.961781455 | 1 |
| 0.316563887 | 0.003975385 |
| 1 | 0.594507858 |
| 0.889745466 | 0.00302217 |
| 0.70828587 | 2.7803E-05 |
| 1 | 0.753614868 |
| 0.44785243 | 1 |
| 0.796249118 | 0.006121286 |
| 1 | 0.058479385 |
| 1 | 0.683797809 |
| 0.015523013 | 8.54557E-12 |
| 0.807606617 | 0.000314223 |
| 1.87923E-05 | 8.54557E-12 |
| 0.13342107 | 9.92556E-08 |
| 0.008180991 | 2.39281E-08 |
| 0.185837406 | 5.64166E-07 |
| 0.035044582 | 1.30556E-08 |
| 1 | 1 |
| 1 | 0.267984254 |
| 0.32467204 | 0.00969829 |
| 1 | 0.038082794 |
| 1 | 0.426979813 |
| 0.962734154 | 5.29626E-06 |
| 0.030595297 | 7.22194E-11 |
| 0.496670411 | 0.000435706 |
| 1 | 3.71157E-09 |
| 0.137599611 | 3.16652E-11 |
| 1 | 5.96684E-07 |
| 3.37539E-08 | 8.54557E-12 |
| 1 | 0.492435639 |
| 1 | 3.62431E-08 |
| 1 | 0.001948528 |
| 0.000123171 | 9.02805E-12 |
| 1.98757E-07 | 8.54557E-12 |
| 1 | 0.67408985 |
| 0.045531556 | 0.865676608 |
| 1 | 0.000791912 |
| 0.201277563 | 7.737E-07 |
| 1 | 1 |
| 1 | 3.39885E-05 |
| 1 | 0.148329451 |
| 0.109358472 | 3.18175E-08 |
| 0.053249789 | 8.54557E-12 |
| 1 | 2.70661E-07 |
| 1.50243E-05 | 8.54557E-12 |
| 1 | 0.89579158 |
| 0.259187068 | 0.010904113 |
| 0.000621466 | 8.54557E-12 |
| 3.88203E-07 | 8.54557E-12 |
| 8.68424E-07 | 8.54557E-12 |
| 1.83109E-10 | 8.54557E-12 |
| 1 | 2.84165E-11 |
| 0.278304937 | 0.03589836 |
| 0.000353051 | 1.1579E-10 |
| 0.001160427 | 8.54557E-12 |
| 0.06333724 | 1.55135E-11 |
| 1 | 1.00184E-09 |
| 1 | 3.33888E-07 |
| 0.184282302 | 8.66798E-10 |
| 0.225844471 | 1.92951E-08 |
| 0.321558031 | 0.045812845 |

|  |  |
| --- | --- |
| Adj. P-value: (SM_neg_newanalysis_12022024)/(WATER_NEG_NEWANALYSIS_12022024) |  |
|  | 0.036975934 |
|  | 0.120325272 |
|  | 0.790301597 |
|  | 0.000434061 |
|  | 0.005817218 |
|  | 1 |
|  | 7.19567E-11 |
|  | 1.72761E-07 |
|  | 2.35531E-05 |
|  | 1.50931E-09 |
|  | 1.53964E-07 |
|  | 1 |
|  | 0.941935293 |
|  | 0.289338068 |
|  | 0.65973932 |
|  | 0.445214305 |
|  | 0.027355279 |
|  | 0.740386316 |
|  | 0.058556097 |
|  | 0.911311241 |
|  | 0.504177176 |
|  | 0.005013725 |
|  | 1 |
|  | 0.073450073 |
|  | 0.044649635 |
|  | 2.97289E-07 |
|  | 5.82393E-06 |
|  | 1.21416E-05 |
|  | 0.019716293 |
|  | 0.008623282 |
|  | 0.000201464 |
|  | 0.000205886 |
|  | 0.003268729 |

|  |
| --- |
| 0.364738826 |
| 1 |
| 1 |
| 0.837758194 |
| 0.021045548 |
| 0.94957543 |
| 0.737679956 |
| 0.433777541 |
| 0.06076251 |
| 1 |
| 0.099912281 |
| 0.041930926 |
| 0.08896773 |
| 0.037204345 |
| 0.026900213 |
| 0.145697714 |
| 4.89985E-06 |
| 0.438623801 |
| 0.056703777 |
| 0.360333725 |
| 0.049623004 |
| 0.40146101 |
| 1 |
| 0.003661942 |
| 0.050945714 |
| 0.008526608 |
| 1 |
| 0.076669002 |
| 0.000679524 |
| 0.021705296 |
| 0.017965068 |
| 1.70712E-07 |
| 0.038396827 |
| 0.546890157 |
| 1 |
| 0.393124628 |
| 1 |
| 0.001325503 |
| 1 |
| 0.010217501 |
| 1 |
| 1 |
| 0.243734008 |
| 0.281144232 |
| 0.811817759 |
| 0.020996478 |
| 0.002797461 |
| 3.61387E-05 |
| 0.002076806 |
| 0.978285154 |
| 0.046889199 |
| 0.525814557 |
| 0.016493276 |
| 0.009434245 |
| 0.376622267 |
| 1.14844E-08 |
| 3.27529E-05 |
| 0.010899169 |
| 0.005085252 |
| 0.093323213 |
| 0.072411192 |
| 0.004231742 |
| 0.01725548 |
| 0.175228857 |
| 0.804917303 |
| 9.49097E-05 |
| 0.078047128 |
| 0.85681806 |
| 0.552288435 |
| 0.003936184 |
| 0.806975243 |
| 1 |
| 0.070579342 |
| 0.071490582 |
| 0.358366232 |
| 3.45289E-09 |
| 0.01401318 |
| 4.11246E-06 |
| 1.68732E-09 |
| 8.69336E-07 |
| 0.33817806 |
| 0.274016228 |
| 0.999775061 |
| 0.042733609 |
| 0.146675194 |
| 0.776939751 |
| 0.00680639 |
| 1 |
| 0.031960646 |
| 1 |
| 0.683624591 |
| 0.036102451 |
| 0.317999663 |
| 0.004756853 |
| 0.665653787 |
| 0.348024968 |
| 0.272377409 |
| 1.10536E-07 |
| 1 |
| 5.4037E-10 |
| 0.382712447 |
| 9.79411E-05 |
| 9.84978E-05 |
| 0.045909868 |
| 0.786557157 |
| 1 |
| 0.023435525 |
| 0.516535989 |
| 0.322059408 |
| 0.919393056 |
| 0.049325195 |
| 1 |
| 0.353167018 |
| 0.002161772 |
| 1 |
| 2.05814E-10 |
| 0.858428127 |
| 0.053152008 |
| 1 |
| 5.41894E-07 |
| 7.24002E-10 |
| 1 |
| 0.001763891 |
| 0.044660836 |
| 0.02234617 |
| 0.374824984 |
| 1 |
| 0.909157194 |
| 0.00271177 |
| 0.000266729 |
| 0.542690395 |
| 5.70417E-05 |
| 0.397908812 |
| 0.010202897 |
| 4.26995E-05 |
| 2.87787E-09 |
| 1.14935E-07 |
| 7.19567E-11 |
| 0.084876268 |
| 1 |

|  |  |
| --- | --- |
|  | 3.53363E-05 |
|  | 9.50557E-06 |
|  | 0.002605939 |
|  | 0.298551592 |
|  | 1 |
|  | 0.012608771 |
|  | 0.002884013 |
|  | 0.005255148 |
