## Supplementary file 4 for "A lipid compendium of a metabolically compromised bacterium provides insights into lipid acquisition, biosynthesis, and metabolism"

### Supplementary Spreadsheet 4: MRM (mass reaction monitoring) parameters fo

or targeted lipid analysis including transitions, collision energies, and instrumenta

#### l settings in triple quadrupole

| Compound group | Compound name | Compound formula | Ion speci | CAS | z | Monoisot | ISTD? |
|---|---|---|---|---|---|---|---|
| AcylCarnitine(12:0) | AcylCarnitine(12:0) | C42H82NO10P | [M+H]+ | | 1 | | FALSE |
| AcylCarnitine(14:0) | AcylCarnitine(14:0) | C42H82NO10P | [M+H]+ | | 1 | | FALSE |
| AcylCarnitine(14:1) | AcylCarnitine(14:1) | C42H82NO10P | [M+H]+ | | 1 | | FALSE |
| AcylCarnitine(14:2) | AcylCarnitine(14:2) | C42H82NO10P | [M+H]+ | | 1 | | FALSE |
| AcylCarnitine(16:0) | AcylCarnitine(16:0) | C42H82NO10P | [M+H]+ | | 1 | | FALSE |
| AcylCarnitine(16:1) | AcylCarnitine(16:1) | C42H82NO10P | [M+H]+ | | 1 | | FALSE |
| AcylCarnitine(18:0) | AcylCarnitine(18:0) | C42H82NO10P | [M+H]+ | | 1 | | FALSE |
| AcylCarnitine(18:1) | AcylCarnitine(18:1) | C42H82NO10P | [M+H]+ | | 1 | | FALSE |
| AcylCarnitine(18:2) | AcylCarnitine(18:2) | C42H82NO10P | [M+H]+ | | 1 | | FALSE |
| AcylCarnitine(20:0) | AcylCarnitine(20:0) | C42H82NO10P | [M+H]+ | | 1 | | FALSE |
| AcHexCmE(18:1)_a | AcHexCmE(18:1)_a | C52H91O7 | [M+H]+ | | 1 | | FALSE |
| AcHexCmE(18:1)_b | AcHexCmE(18:1)_b | C52H91O7 | [M+NH4]+ | | 1 | | FALSE |
| C-Gal(a) | C-Gal(a) | $C_{33}H_{56}O_6$ | [M+Na]+ | | 1 | | FALSE |
| C-Gal(b) | C-Gal(b) | $C_{33}H_{56}O_6$ | [M+Na]+ | | 1 | | FALSE |
| BbGL-1(a) | BbGL-1(a) | $C_{49}H_{86}O_7$ | [M+NH4]+ | | 1 | | FALSE |
| BbGL-1(b) | BbGL-1(b) | $C_{49}H_{86}O_7$ | [M+NH4]+ | | 1 | | FALSE |
| CE(16:0) | CE(16:0) | C42H82NO10P | [M+NH4]+ | | 1 | | FALSE |
| CE(16:1) | CE(16:1) | C42H82NO10P | [M+NH4]+ | | 1 | | FALSE |
| CE(16:2) | CE(16:2) | C42H82NO10P | [M+NH4]+ | | 1 | | FALSE |
| CE(17:0) | CE(17:0) | C42H82NO10P | [M+NH4]+ | | 1 | | FALSE |
| CE(17:1) | CE(17:1) | C42H82NO10P | [M+NH4]+ | | 1 | | FALSE |
| CE(18:0) | CE(18:0) | C42H82NO10P | [M+NH4]+ | | 1 | | FALSE |
| CE(18:1) | CE(18:1) | C42H82NO10P | [M+NH4]+ | | 1 | | FALSE |
| CE(18:2) | CE(18:2) | C42H82NO10P | [M+NH4]+ | | 1 | | FALSE |
| CE(18:3) | CE(18:3) | C42H82NO10P | [M+NH4]+ | | 1 | | FALSE |
| CE(20:1) | CE(20:1) | C42H82NO10P | [M+NH4]+ | | 1 | | FALSE |
| CE(20:2) | CE(20:2) | C42H82NO10P | [M+NH4]+ | | 1 | | FALSE |
| CE(20:3) | CE(20:3) | C42H82NO10P | [M+NH4]+ | | 1 | | FALSE |
| CE(20:4) | CE(20:4) | C42H82NO10P | [M+NH4]+ | | 1 | | FALSE |
| CE(20:5) | CE(20:5) | C42H82NO10P | [M+NH4]+ | | 1 | | FALSE |
| CE(22:0) | CE(22:0) | C42H82NO10P | [M+NH4]+ | | 1 | | FALSE |
| CE(22:1) | CE(22:1) | C42H82NO10P | [M+NH4]+ | | 1 | | FALSE |
| CE(22:4) | CE(22:4) | C42H82NO10P | [M+NH4]+ | | 1 | | FALSE |
| CE(22:5) (n3) | CE(22:5) (n3) | C42H82NO10P | [M+NH4]+ | | 1 | | FALSE |
| CE(22:6) | CE(22:6) | C42H82NO10P | [M+NH4]+ | | 1 | | FALSE |
| CE(24:0) | CE(24:0) | C42H82NO10P | [M+NH4]+ | | 1 | | FALSE |
| CE(24:1) | CE(24:1) | C42H82NO10P | [M+NH4]+ | | 1 | | FALSE |

| | | | | | |
|---|---|---|---|---|---|
| CE(24:4) | CE(24:4) | C42H82NO10P | [M+NH4]+ | 1 | FALSE |
| CE(24:5) | CE(24:5) | C42H82NO10P | [M+NH4]+ | 1 | FALSE |
| CE(24:6) | CE(24:6) | C42H82NO10P | [M+NH4]+ | 1 | FALSE |
| CmE(18:3) | CmE(18:3) | $C_{46}H_{76}O_2$ | [M+NH4]+ | 1 | FALSE |
| Cer(d16:1/16:0) | Cer(d16:1/16:0) | | [M+H]+ | 1 | FALSE |
| Cer(d16:1/18:0) | Cer(d16:1/18:0) | | [M+H]+ | 1 | FALSE |
| Cer(d16:1/20:0) | Cer(d16:1/20:0) | | [M+H]+ | 1 | FALSE |
| Cer(d16:1/22:0) | Cer(d16:1/22:0) | | [M+H]+ | 1 | FALSE |
| Cer(d16:1/23:0) | Cer(d16:1/23:0) | | [M+H]+ | 1 | FALSE |
| Cer(d16:1/24:0) | Cer(d16:1/24:0) | | [M+H]+ | 1 | FALSE |
| Cer(d16:1/24:1) | Cer(d16:1/24:1) | | [M+H]+ | 1 | FALSE |
| Cer(d18:1/14:0) | Cer(d18:1/14:0) | | [M+H]+ | 1 | FALSE |
| Cer(d18:1/16:0) | Cer(d18:1/16:0) | | [M+H]+ | 1 | FALSE |
| Cer(d18:1/18:0) | Cer(d18:1/18:0) | | [M+H]+ | 1 | FALSE |
| Cer(d18:1/19:0) | Cer(d18:1/19:0) | | [M+H]+ | 1 | FALSE |
| Cer(d18:1/20:0) | Cer(d18:1/20:0) | | [M+H]+ | 1 | FALSE |
| Cer(d18:1/21:0) | Cer(d18:1/21:0) | | [M+H]+ | 1 | FALSE |
| Cer(d18:1/22:0) | Cer(d18:1/22:0) | | [M+H]+ | 1 | FALSE |
| Cer(d18:1/23:0) | Cer(d18:1/23:0) | | [M+H]+ | 1 | FALSE |
| Cer(d18:1/24:0) | Cer(d18:1/24:0) | | [M+H]+ | 1 | FALSE |
| Cer(d18:1/24:1) | Cer(d18:1/24:1) | | [M+H]+ | 1 | FALSE |
| Cer(d18:1/26:0) | Cer(d18:1/26:0) | | [M+H]+ | 1 | FALSE |
| Cer(d18:2/14:0) | Cer(d18:2/14:0) | | [M+H]+ | 1 | FALSE |
| Cer(d18:2/16:0) | Cer(d18:2/16:0) | | [M+H]+ | 1 | FALSE |
| Cer(d18:2/18:0) | Cer(d18:2/18:0) | | [M+H]+ | 1 | FALSE |
| Cer(d18:2/20:0) | Cer(d18:2/20:0) | | [M+H]+ | 1 | FALSE |
| Cer(d18:2/21:0) | Cer(d18:2/21:0) | | [M+H]+ | 1 | FALSE |
| Cer(d18:2/22:0) | Cer(d18:2/22:0) | | [M+H]+ | 1 | FALSE |
| Cer(d18:2/23:0) | Cer(d18:2/23:0) | | [M+H]+ | 1 | FALSE |
| Cer(d18:2/24:0) | Cer(d18:2/24:0) | | [M+H]+ | 1 | FALSE |
| Cer(d18:2/24:1) | Cer(d18:2/24:1) | | [M+H]+ | 1 | FALSE |
| Cer(d18:2/26:0) | Cer(d18:2/26:0) | | [M+H]+ | 1 | FALSE |
| Cer(d20:1/22:0) | Cer(d20:1/22:0) | | [M+H]+ | 1 | FALSE |
| Cer(d20:1/23:0) | Cer(d20:1/23:0) | | [M+H]+ | 1 | FALSE |
| Cer(d20:1/24:0) | Cer(d20:1/24:0) | | [M+H]+ | 1 | FALSE |
| Cer(d20:1/24:1) | Cer(d20:1/24:1) | | [M+H]+ | 1 | FALSE |
| Cer(d20:1/26:0) | Cer(d20:1/26:0) | | [M+H]+ | 1 | FALSE |
| Cer1P(d18:1/16:0) | Cer1P(d18:1/16:0) | | [M+H]+ | 1 | FALSE |

| | | | | | |
|---|---|---|---|---|---|
| Cer(d18:1_24:2) | Cer(d18:1_24:2) | | [M+H]+ | 1 | FALSE |
| Cer(d27:1) | Cer(d27:1) | | [M+H]+ | 1 | FALSE |
| Cer(d30:0) | Cer(d30:0) | | [M+H]+ | 1 | FALSE |
| Cer(d31:1) | Cer(d31:1) | | [M+H]+ | 1 | FALSE |
| Cer(d32:0) | Cer(d32:0) | | [M+H]+ | 1 | FALSE |
| Cer(d33:1) | Cer(d33:1) | | [M+H]+ | 1 | FALSE |
| Cer(d34:0) | Cer(d34:0) | | [M+H]+ | 1 | FALSE |
| Cer(d34:1) | Cer(d34:1) | | [M+H]+ | 1 | FALSE |
| Cer(d36:0) | Cer(d36:0) | | [M+H]+ | 1 | FALSE |
| Cer(m18:0_16:0) | Cer(m18:0_16:0) | | [M+H]+ | 1 | FALSE |
| Cer(m36:0) | Cer(m36:0) | | [M+H]+ | 1 | FALSE |
| Cer(t20:1_16:0) | Cer(t20:1_16:0) | | [M+H]+ | 1 | FALSE |
| Cer(t20:1_18:0) | Cer(t20:1_18:0) | | [M+H]+ | 1 | FALSE |
| COH | COH | C42H82NO10P | [M+NH4]+ | 1 | FALSE |
| DE(16:0) | DE(16:0) | C42H82NO10P | [M+NH4]+ | 1 | FALSE |
| DE(18:1) | DE(18:1) | C42H82NO10P | [M+NH4]+ | 1 | FALSE |
| DE(18:2) | DE(18:2) | C42H82NO10P | [M+NH4]+ | 1 | FALSE |
| DE(20:4) | DE(20:4) | C42H82NO10P | [M+NH4]+ | 1 | FALSE |
| DE(20:5) | DE(20:5) | C42H82NO10P | [M+NH4]+ | 1 | FALSE |
| DE(22:6) | DE(22:6) | C42H82NO10P | [M+NH4]+ | 1 | FALSE |
| DG(14:0_16:0) | DG(14:0_16:0) | C42H82NO10P | [M+NH4]+ | 1 | FALSE |
| DG(14:0_16:0) | DG(14:0_16:0) | C42H82NO10P | [M+NH4]+ | 1 | FALSE |
| DG(14:0_18:2) | DG(14:0_18:2) | C42H82NO10P | [M+NH4]+ | 1 | FALSE |
| DG(14:0_18:2) | DG(14:0_18:2) | C42H82NO10P | [M+NH4]+ | 1 | FALSE |
| DG(16:0_16:0) | DG(16:0_16:0) | C42H82NO10P | [M+NH4]+ | 1 | FALSE |
| DG(16:0_16:1) | DG(16:0_16:1) | C42H82NO10P | [M+NH4]+ | 1 | FALSE |
| DG(16:0_16:1) | DG(16:0_16:1) | C42H82NO10P | [M+NH4]+ | 1 | FALSE |
| DG(16:0_18:1) | DG(16:0_18:1) | C42H82NO10P | [M+NH4]+ | 1 | FALSE |
| DG(16:0_18:1) | DG(16:0_18:1) | C42H82NO10P | [M+NH4]+ | 2 | FALSE |
| DG(16:0_18:2) | DG(16:0_18:2) | C42H82NO10P | [M+NH4]+ | 1 | FALSE |
| DG(16:0_18:2) | DG(16:0_18:2) | C42H82NO10P | [M+NH4]+ | 1 | FALSE |
| DG(16:0_20:4) | DG(16:0_20:4) | C42H82NO10P | [M+NH4]+ | 1 | FALSE |
| DG(16:0_20:4) | DG(16:0_20:4) | C42H82NO10P | [M+NH4]+ | 1 | FALSE |
| DG(16:0_22:5) | DG(16:0_22:5) | C42H82NO10P | [M+NH4]+ | 1 | FALSE |
| DG(16:0_22:5) | DG(16:0_22:5) | C42H82NO10P | [M+NH4]+ | 1 | FALSE |
| DG(16:0_22:6) | DG(16:0_22:6) | C42H82NO10P | [M+NH4]+ | 1 | FALSE |
| DG(16:0_22:6) | DG(16:0_22:6) | C42H82NO10P | [M+NH4]+ | 1 | FALSE |
| DG(16:1_18:1) | DG(16:1_18:1) | C42H82NO10P | [M+NH4]+ | 1 | FALSE |

| | | | | | |
|---|---|---|---|---|---|
| DG(16:1_18:1) | DG(16:1_18:1) | C42H82NO10P | [M+NH4]+ | 1 | FALSE |
| DG(18:0_18:1) | DG(18:0_18:1) | C42H82NO10P | [M+NH4]+ | 1 | FALSE |
| DG(18:0_18:1) | DG(18:0_18:1) | C42H82NO10P | [M+NH4]+ | 1 | FALSE |
| DG(18:0_18:2) | DG(18:0_18:2) | C42H82NO10P | [M+NH4]+ | 1 | FALSE |
| DG(18:0_18:2) | DG(18:0_18:2) | C42H82NO10P | [M+NH4]+ | 1 | FALSE |
| DG(18:0_20:4) | DG(18:0_20:4) | C42H82NO10P | [M+NH4]+ | 1 | FALSE |
| DG(18:0_20:4) | DG(18:0_20:4) | C42H82NO10P | [M+NH4]+ | 1 | FALSE |
| DG(18:1_18:1) | DG(18:1_18:1) | C42H82NO10P | [M+NH4]+ | 1 | FALSE |
| DG(18:1_18:2) | DG(18:1_18:2) | C42H82NO10P | [M+NH4]+ | 1 | FALSE |
| DG(18:1_18:2) | DG(18:1_18:2) | C42H82NO10P | [M+NH4]+ | 1 | FALSE |
| DG(18:1_20:3) | DG(18:1_20:3) | C42H82NO10P | [M+NH4]+ | 1 | FALSE |
| DG(18:1_20:3) | DG(18:1_20:3) | C42H82NO10P | [M+NH4]+ | 1 | FALSE |

| Precurso | MS1 res | Product m/ | MS2 res | Dwell (m | Fragmen | CE (V) | Polarity |
|---|---|---|---|---|---|---|---|
| 344.3 | Unit | 85.1 | Unit | 20 | 166 | 30 | Positive |
| 372.3 | Unit | 85.1 | Unit | 20 | 166 | 30 | Positive |
| 370.3 | Unit | 85.1 | Unit | 20 | 166 | 30 | Positive |
| 368.3 | Unit | 85.1 | Unit | 20 | 166 | 30 | Positive |
| 400.4 | Unit | 85.1 | Unit | 20 | 166 | 30 | Positive |
| 398.3 | Unit | 85.1 | Unit | 20 | 166 | 30 | Positive |
| 428.4 | Unit | 85.1 | Unit | 20 | 166 | 30 | Positive |
| 426.4 | Unit | 85.1 | Unit | 20 | 166 | 30 | Positive |
| 424.3 | Unit | 85.1 | Unit | 20 | 166 | 30 | Positive |
| 456.4 | Unit | 85.1 | Unit | 20 | 166 | 30 | Positive |
| 827.7 | Unit | 383.3 | Unit | 20 | 166 | 23 | Positive |
| 844.7 | Unit | 383.3 | Unit | 20 | 166 | 23 | Positive |
| 571.4 | | 369.3 | Unit | 20 | 166 | 23 | Positive |
| 571.4 | | 203.1 | Unit | 20 | 166 | 23 | Positive |
| 804.7 | | 369.3 | Unit | 20 | 166 | 23 | Positive |
| 804.7 | | 401.3 | Unit | 20 | 166 | 23 | Positive |
| 642.6 | Unit | 369.3 | Unit | 20 | 166 | 10 | Positive |
| 640.6 | Unit | 369.3 | Unit | 20 | 166 | 10 | Positive |
| 638.6 | Unit | 369.3 | Unit | 20 | 166 | 10 | Positive |
| 656.6 | Unit | 369.3 | Unit | 20 | 166 | 10 | Positive |
| 654.6 | Unit | 369.3 | Unit | 20 | 166 | 10 | Positive |
| 670.7 | Unit | 369.3 | Unit | 20 | 166 | 10 | Positive |
| 668.6 | Unit | 369.3 | Unit | 20 | 166 | 10 | Positive |
| 666.6 | Unit | 369.3 | Unit | 20 | 166 | 10 | Positive |
| 664.6 | Unit | 369.3 | Unit | 20 | 166 | 10 | Positive |
| 696.7 | Unit | 369.3 | Unit | 20 | 166 | 10 | Positive |
| 694.7 | Unit | 369.3 | Unit | 20 | 166 | 10 | Positive |
| 692.6 | Unit | 369.3 | Unit | 20 | 166 | 10 | Positive |
| 690.6 | Unit | 369.3 | Unit | 20 | 166 | 10 | Positive |
| 688.6 | Unit | 369.3 | Unit | 20 | 166 | 10 | Positive |
| 726.7 | Unit | 369.3 | Unit | 20 | 166 | 10 | Positive |
| 724.7 | Unit | 369.3 | Unit | 20 | 166 | 10 | Positive |
| 718.7 | Unit | 369.3 | Unit | 20 | 166 | 10 | Positive |
| 716.6 | Unit | 369.3 | Unit | 20 | 166 | 10 | Positive |
| 714.6 | Unit | 369.3 | Unit | 20 | 166 | 10 | Positive |
| 754.7 | Unit | 369.3 | Unit | 20 | 166 | 10 | Positive |
| 752.7 | Unit | 369.3 | Unit | 20 | 166 | 10 | Positive |

| | | | | | | | |
|---|---|---|---|---|---|---|---|
| 746.7 | Unit | 369.3 | Unit | 20 | 166 | 10 | Positive |
| 744.7 | Unit | 369.3 | Unit | 20 | 166 | 10 | Positive |
| 742.7 | Unit | 369.3 | Unit | 20 | 166 | 10 | Positive |
| 678.6 | Unit | 383.3 | Unit | 20 | 166 | 20 | Positive |
| 510.6 | Unit | 236.3 | Unit | 20 | 166 | 10 | Positive |
| 538.6 | Unit | 236.3 | Unit | 20 | 166 | 10 | Positive |
| 566.6 | Unit | 236.3 | Unit | 20 | 166 | 29 | Positive |
| 594.6 | Unit | 236.3 | Unit | 20 | 166 | 29 | Positive |
| 608.6 | Unit | 236.3 | Unit | 20 | 166 | 29 | Positive |
| 622.6 | Unit | 236.3 | Unit | 20 | 166 | 29 | Positive |
| 620.6 | Unit | 236.3 | Unit | 20 | 166 | 29 | Positive |
| 510.5 | Unit | 264.3 | Unit | 20 | 166 | 29 | Positive |
| 538.5 | Unit | 264.3 | Unit | 20 | 166 | 29 | Positive |
| 566.6 | Unit | 264.3 | Unit | 20 | 166 | 29 | Positive |
| 580.6 | Unit | 264.3 | Unit | 20 | 166 | 29 | Positive |
| 594.6 | Unit | 264.3 | Unit | 20 | 166 | 29 | Positive |
| 608.6 | Unit | 264.3 | Unit | 20 | 166 | 29 | Positive |
| 622.6 | Unit | 264.3 | Unit | 20 | 166 | 29 | Positive |
| 636.6 | Unit | 264.3 | Unit | 20 | 166 | 29 | Positive |
| 650.6 | Unit | 264.3 | Unit | 20 | 166 | 29 | Positive |
| 648.6 | Unit | 264.3 | Unit | 20 | 166 | 29 | Positive |
| 678.6 | Unit | 264.3 | Unit | 20 | 166 | 29 | Positive |
| 508.5 | Unit | 262.3 | Unit | 20 | 166 | 23 | Positive |
| 536.5 | Unit | 262.3 | Unit | 20 | 166 | 23 | Positive |
| 564.6 | Unit | 262.3 | Unit | 20 | 166 | 23 | Positive |
| 592.6 | Unit | 262.3 | Unit | 20 | 166 | 23 | Positive |
| 606.6 | Unit | 262.3 | Unit | 20 | 166 | 23 | Positive |
| 620.6 | Unit | 262.3 | Unit | 20 | 166 | 23 | Positive |
| 634.6 | Unit | 262.3 | Unit | 20 | 166 | 23 | Positive |
| 648.6 | Unit | 262.3 | Unit | 20 | 166 | 23 | Positive |
| 646.6 | Unit | 262.3 | Unit | 20 | 166 | 23 | Positive |
| 676.6 | Unit | 262.3 | Unit | 20 | 166 | 23 | Positive |
| 650.6 | Unit | 292.3 | Unit | 20 | 166 | 29 | Positive |
| 664.6 | Unit | 292.3 | Unit | 20 | 166 | 29 | Positive |
| 678.6 | Unit | 292.3 | Unit | 20 | 166 | 29 | Positive |
| 676.6 | Unit | 292.3 | Unit | 20 | 166 | 29 | Positive |
| 706.6 | Unit | 292.3 | Unit | 20 | 166 | 29 | Positive |
| 618.4 | Unit | 264.3 | Unit | 20 | 166 | 29 | Positive |

| | | | | | | |
|---|---|---|---|---|---|---|
| 646.6 | Unit | 264.3 | Unit | 20 | 166 | 29 Positive |
| 440.4 | Unit | 264.3 | Unit | 20 | 166 | 29 Positive |
| 484.5 | Unit | 264.3 | Unit | 20 | 166 | 29 Positive |
| 496.5 | Unit | 264.3 | Unit | 20 | 166 | 29 Positive |
| 512.5 | Unit | 264.3 | Unit | 20 | 166 | 29 Positive |
| 524.5 | Unit | 264.3 | Unit | 20 | 166 | 29 Positive |
| 540.5 | Unit | 264.3 | Unit | 20 | 166 | 29 Positive |
| 538.5 | Unit | 264.3 | Unit | 20 | 166 | 29 Positive |
| 568.6 | Unit | 264.3 | Unit | 20 | 166 | 29 Positive |
| 524.5 | Unit | 264.3 | Unit | 20 | 166 | 29 Positive |
| 552.6 | Unit | 264.3 | Unit | 20 | 166 | 29 Positive |
| 582.5 | Unit | 264.3 | Unit | 20 | 166 | 29 Positive |
| 610.6 | Unit | 264.3 | Unit | 20 | 166 | 29 Positive |
| 369.4 | Unit | 161.2 | Unit | 20 | 166 | 23 Positive |
| 640.8 | Unit | 367.4 | Unit | 20 | 166 | 12 Positive |
| 666.8 | Unit | 367.4 | Unit | 20 | 166 | 12 Positive |
| 664.8 | Unit | 367.4 | Unit | 20 | 166 | 12 Positive |
| 688.8 | Unit | 367.4 | Unit | 20 | 166 | 12 Positive |
| 686.8 | Unit | 367.4 | Unit | 20 | 166 | 12 Positive |
| 712.8 | Unit | 367.4 | Unit | 20 | 166 | 12 Positive |
| 558.5 | Unit | 285.2 | Unit | 20 | 166 | 21 Positive |
| 558.5 | Unit | 313.3 | Unit | 20 | 166 | 21 Positive |
| 582.5 | Unit | 285.2 | Unit | 20 | 166 | 21 Positive |
| 582.5 | Unit | 337.3 | Unit | 20 | 166 | 21 Positive |
| 586.5 | Unit | 313.2 | Unit | 20 | 166 | 21 Positive |
| 584.5 | Unit | 313.2 | Unit | 20 | 166 | 21 Positive |
| 584.5 | Unit | 311.3 | Unit | 20 | 166 | 21 Positive |
| 612.6 | Unit | 313.3 | Unit | 20 | 166 | 21 Positive |
| 612.6 | Unit | 339.3 | Unit | 20 | 166 | 21 Positive |
| 610.5 | Unit | 313.2 | Unit | 20 | 166 | 21 Positive |
| 610.5 | Unit | 337.3 | Unit | 20 | 166 | 21 Positive |
| 634.5 | Unit | 313.2 | Unit | 20 | 166 | 21 Positive |
| 634.5 | Unit | 361.3 | Unit | 20 | 166 | 21 Positive |
| 660.6 | Unit | 313.3 | Unit | 20 | 166 | 21 Positive |
| 660.6 | Unit | 387.3 | Unit | 20 | 166 | 21 Positive |
| 658.5 | Unit | 313.2 | Unit | 20 | 166 | 21 Positive |
| 658.5 | Unit | 385.3 | Unit | 20 | 166 | 21 Positive |
| 610.5 | Unit | 339.2 | Unit | 20 | 166 | 21 Positive |

| | | | | | | | |
|---|---|---|---|---|---|---|---|
| 610.5 | Unit | 311.3 | Unit | 20 | 166 | 21 | Positive |
| 640.6 | Unit | 341.3 | Unit | 20 | 166 | 21 | Positive |
| 640.6 | Unit | 339.2 | Unit | 20 | 166 | 21 | Positive |
| 638.6 | Unit | 341.3 | Unit | 20 | 166 | 21 | Positive |
| 638.6 | Unit | 337.3 | Unit | 20 | 166 | 21 | Positive |
| 662.6 | Unit | 341.3 | Unit | 20 | 166 | 21 | Positive |
| 662.6 | Unit | 361.3 | Unit | 20 | 166 | 21 | Positive |
| 638.6 | Unit | 339.3 | Unit | 20 | 166 | 21 | Positive |
| 636.6 | Unit | 339.3 | Unit | 20 | 166 | 21 | Positive |
| 636.6 | Unit | 337.3 | Unit | 20 | 166 | 21 | Positive |
| 662.6 | Unit | 363.2894 | Unit | 20 | 166 | 21 | Positive |
| 662.6 | Unit | 339.3 | Unit | 20 | 166 | 21 | Positive |

| Compound group | Compour | Compour | Ion speci | CAS | z | Monoisot | ISTD | Precurso | MS1 res | Product r | MS2 res |
|---|---|---|---|---|---|---|---|---|---|---|---|
| SQDG_NEG | SQDG 16:0/16:0 | | [M-H]- | | 1 | 794.5 | FALSE | 793.5 | Unit | 81 | Unit |
| SQDG_NEG | SQDG 16:0/16:0 | | [M-H]- | | 1 | 794.5 | FALSE | 793.5 | Unit | 225 | Unit |
| SQDG_NEG | SQDG 16:0/16:1 | | [M-H]- | | 1 | 792.5 | FALSE | 791.5 | Unit | 81 | Unit |
| SQDG_NEG | SQDG 16:0/16:1 | | [M-H]- | | 1 | 792.5 | FALSE | 791.5 | Unit | 225 | Unit |
| SQDGNH4 | SQDG 16:0/16:1 | | [M+H]+ | | 1 | 809.5 | FALSE | 810.5 | Unit | 313.3 | Unit |
| SQDGNH4 | SQDG 16:0/16:1 | | [M+H]+ | | 1 | 809.5 | FALSE | 810.5 | Unit | 311.3 | Unit |
| SQDG_NEG | SQDG 16:0/18:0 | | [M-H]- | | 1 | 822.5 | FALSE | 821.5 | Unit | 81 | Unit |
| SQDG_NEG | SQDG 16:0/18:0 | | [M-H]- | | 1 | 822.5 | FALSE | 821.5 | Unit | 225 | Unit |
| SQDGNH4 | SQDG 16:0/18:0 | | [M+H]+ | | 1 | 839.6 | FALSE | 840.6 | Unit | 313.3 | Unit |
| SQDGNH4 | SQDG 16:0/18:0 | | [M+H]+ | | 1 | 839.6 | FALSE | 840.6 | Unit | 341.3 | Unit |
| SQDG_NEG | SQDG 16:0/18:1 | | [M-H]- | | 1 | 820.5 | FALSE | 819.5 | Unit | 81 | Unit |
| SQDG_NEG | SQDG 16:0/18:1 | | [M-H]- | | 1 | 820.5 | FALSE | 819.5 | Unit | 225 | Unit |
| SQDGNH4 | SQDG 16:0/18:1 | | [M+H]+ | | 1 | 837.6 | FALSE | 838.6 | Unit | 313.3 | Unit |
| SQDGNH4 | SQDG 16:0/18:1 | | [M+H]+ | | 1 | 837.6 | FALSE | 838.6 | Unit | 339.3 | Unit |
| SQDG_NEG | SQDG 16:0/18:2 | | [M-H]- | | 1 | 818.5 | FALSE | 817.5 | Unit | 81 | Unit |
| SQDG_NEG | SQDG 16:0/18:2 | | [M-H]- | | 1 | 818.5 | FALSE | 817.5 | Unit | 225 | Unit |
| SQDGNH4 | SQDG 16:0/18:2 | | [M+H]+ | | 1 | 836.6 | FALSE | 837.6 | Unit | 313.3 | Unit |
| SQDGNH4 | SQDG 16:0/18:2 | | [M+H]+ | | 1 | 836.6 | FALSE | 837.6 | Unit | 337.3 | Unit |
| SQDG_NEG | SQDG 16:0/18:3 | | [M-H]- | | 1 | 816.5 | FALSE | 815.5 | Unit | 81 | Unit |
| SQDG_NEG | SQDG 16:0/18:3 | | [M-H]- | | 1 | 816.5 | FALSE | 815.5 | Unit | 225 | Unit |
| SQDGNH4 | SQDG 16:0/18:3 | | [M+H]+ | | 1 | 833.5 | FALSE | 834.5 | Unit | 313.3 | Unit |
| SQDGNH4 | SQDG 16:0/18:3 | | [M+H]+ | | 1 | 833.5 | FALSE | 834.5 | Unit | 335.3 | Unit |
| SQDG_NEG | SQDG 16:0/22:6 | | [M-H]- | | 1 | 866.5 | FALSE | 865.5 | Unit | 81 | Unit |
| SQDG_NEG | SQDG 16:0/22:6 | | [M-H]- | | 1 | 866.5 | FALSE | 865.5 | Unit | 225 | Unit |
| SQDGNH4 | SQDG 16:0/22:6 | | [M+H]+ | | 1 | 883.6 | FALSE | 884.6 | Unit | 313.3 | Unit |
| SQDGNH4 | SQDG 16:0/22:6 | | [M+H]+ | | 1 | 883.6 | FALSE | 884.6 | Unit | 385.3 | Unit |
| SQDG_NEG | SQDG 16:1/16:1 | | [M-H]- | | 1 | 790.5 | FALSE | 789.5 | Unit | 81 | Unit |
| SQDG_NEG | SQDG 16:1/16:1 | | [M-H]- | | 1 | 790.5 | FALSE | 789.5 | Unit | 225 | Unit |
| SQDG_NEG | SQDG 16:1/18:1 | | [M-H]- | | 1 | 818.5 | FALSE | 817.5 | Unit | 81 | Unit |
| SQDG_NEG | SQDG 16:1/18:1 | | [M-H]- | | 1 | 818.5 | FALSE | 817.5 | Unit | 225 | Unit |
| SQDGNH4 | SQDG 16:1/18:1 | | [M+H]+ | | 1 | 835.6 | FALSE | 836.6 | Unit | 311.3 | Unit |
| SQDGNH4 | SQDG 16:1/18:1 | | [M+H]+ | | 1 | 835.6 | FALSE | 836.6 | Unit | 339.3 | Unit |
| SQDG_NEG | SQDG 16:1/18:2 | | [M-H]- | | 1 | 816.5 | FALSE | 815.5 | Unit | 81 | Unit |
| SQDG_NEG | SQDG 16:1/18:2 | | [M-H]- | | 1 | 816.5 | FALSE | 815.5 | Unit | 225 | Unit |
| SQDGNH4 | SQDG 16:1/18:2 | | [M+H]+ | | 1 | 833.5 | FALSE | 834.5 | Unit | 311.3 | Unit |
| SQDGNH4 | SQDG 16:1/18:2 | | [M+H]+ | | 1 | 833.5 | FALSE | 834.5 | Unit | 337.3 | Unit |
| SQDG_NEG | SQDG 16:1/20:3 | | [M-H]- | | 1 | 842.5 | FALSE | 841.5 | Unit | 81 | Unit |

| | | | | | | | | | |
|---|---|---|---|---|---|---|---|---|---|
| SQDG_NEG | SQDG 16:1/20:3 | [M-H]- | 1 | 842.5 | FALSE | 841.5 | Unit | 225 | Unit |
| SQDGNH4 | SQDG 16:1/20:3 | [M+H]+ | 1 | 859.6 | FALSE | 860.6 | Unit | 311.3 | Unit |
| SQDGNH4 | SQDG 16:1/20:3 | [M+H]+ | 1 | 859.6 | FALSE | 860.6 | Unit | 363.3 | Unit |
| SQDG_NEG | SQDG 16:1/22:6 | [M-H]- | 1 | 864.5 | FALSE | 863.5 | Unit | 81 | Unit |
| SQDG_NEG | SQDG 16:1/22:6 | [M-H]- | 1 | 864.5 | FALSE | 863.5 | Unit | 225 | Unit |
| SQDGNH4 | SQDG 16:1/22:6 | [M+H]+ | 1 | 881.5 | FALSE | 882.5 | Unit | 311.3 | Unit |
| SQDGNH4 | SQDG 16:1/22:6 | [M+H]+ | 1 | 881.5 | FALSE | 882.5 | Unit | 385.3 | Unit |
| SQDG_NEG | SQDG 18:0/18:0 | [M-H]- | 1 | 850.6 | FALSE | 849.6 | Unit | 81 | Unit |
| SQDG_NEG | SQDG 18:0/18:0 | [M-H]- | 1 | 850.6 | FALSE | 849.6 | Unit | 225 | Unit |
| SQDG_NEG | SQDG 18:0/18:1 | [M-H]- | 1 | 848.6 | FALSE | 847.6 | Unit | 81 | Unit |
| SQDG_NEG | SQDG 18:0/18:1 | [M-H]- | 1 | 848.6 | FALSE | 847.6 | Unit | 225 | Unit |
| SQDGNH4 | SQDG 18:1/18:1 | [M+H]+ | 1 | 863.6 | FALSE | 864.6 | Unit | 339.3 | Unit |
| SQDG_NEG | SQDG 18:1/18:2 | [M-H]- | 1 | 844.5 | FALSE | 843.5 | Unit | 81 | Unit |
| SQDG_NEG | SQDG 18:1/18:2 | [M-H]- | 1 | 844.5 | FALSE | 843.5 | Unit | 225 | Unit |
| SQDG_NEG | SQDG 18:1/20:0 | [M-H]- | 1 | 876.6 | FALSE | 875.6 | Unit | 81 | Unit |
| SQDG_NEG | SQDG 18:1/20:0 | [M-H]- | 1 | 876.6 | FALSE | 875.6 | Unit | 225 | Unit |
| SQDGNH4 | SQDG 18:1/20:0 | [M+H]+ | 1 | 893.6 | FALSE | 894.6 | Unit | 339.3 | Unit |
| SQDGNH4 | SQDG 18:1/20:0 | [M+H]+ | 1 | 893.6 | FALSE | 894.6 | Unit | 369.3 | Unit |
| SQDG_NEG | SQDG 18:1/20:1 | [M-H]- | 1 | 874.6 | FALSE | 873.6 | Unit | 81 | Unit |
| SQDG_NEG | SQDG 18:1/20:1 | [M-H]- | 1 | 874.6 | FALSE | 873.6 | Unit | 225 | Unit |
| SQDGNH4 | SQDG 18:1/20:1 | [M+H]+ | 1 | 891.6 | FALSE | 892.6 | Unit | 339.3 | Unit |
| SQDGNH4 | SQDG 18:1/20:1 | [M+H]+ | 1 | 891.6 | FALSE | 892.6 | Unit | 367.3 | Unit |
| SQDG_NEG | SQDG 18:1/20:2 | [M-H]- | 1 | 872.6 | FALSE | 871.6 | Unit | 81 | Unit |
| SQDG_NEG | SQDG 18:1/20:2 | [M-H]- | 1 | 872.6 | FALSE | 871.6 | Unit | 225 | Unit |
| SQDGNH4 | SQDG 18:1/20:2 | [M+H]+ | 1 | 889.6 | FALSE | 890.6 | Unit | 339.3 | Unit |
| SQDGNH4 | SQDG 18:1/20:2 | [M+H]+ | 1 | 889.6 | FALSE | 890.6 | Unit | 365.3 | Unit |
| SQDG_NEG | SQDG 18:1/20:3 | [M-H]- | 1 | 870.5 | FALSE | 869.5 | Unit | 81 | Unit |
| SQDG_NEG | SQDG 18:1/20:3 | [M-H]- | 1 | 870.5 | FALSE | 869.5 | Unit | 225 | Unit |
| SQDGNH4 | SQDG 18:1/20:3 | [M+H]+ | 1 | 887.6 | FALSE | 888.6 | Unit | 339.3 | Unit |
| SQDGNH4 | SQDG 18:1/20:3 | [M+H]+ | 1 | 887.6 | FALSE | 888.6 | Unit | 363.3 | Unit |
| SQDG_NEG | SQDG 18:1/20:4 | [M-H]- | 1 | 868.5 | FALSE | 867.5 | Unit | 81 | Unit |
| SQDG_NEG | SQDG 18:1/20:4 | [M-H]- | 1 | 868.5 | FALSE | 867.5 | Unit | 225 | Unit |
| SQDGNH4 | SQDG 18:1/20:4 | [M+H]+ | 1 | 885.6 | FALSE | 886.6 | Unit | 339.3 | Unit |
| SQDGNH4 | SQDG 18:1/20:4 | [M+H]+ | 1 | 885.6 | FALSE | 886.6 | Unit | 361.3 | Unit |
| SQDG_NEG | SQDG 18:1/22:1 | [M-H]- | 1 | 902.6 | FALSE | 901.6 | Unit | 81 | Unit |
| SQDG_NEG | SQDG 18:1/22:1 | [M-H]- | 1 | 902.6 | FALSE | 901.6 | Unit | 225 | Unit |
| SQDGNH4 | SQDG 18:1/22:1 | [M+H]+ | 1 | 919.6 | FALSE | 920.6 | Unit | 339.3 | Unit |
| SQDGNH4 | SQDG 18:1/22:1 | [M+H]+ | 1 | 919.6 | FALSE | 920.6 | Unit | 395.3 | Unit |

| | | | | | | | | | |
|---|---|---|---|---|---|---|---|---|---|
| SQDG_NEG | SQDG 18:1/22:2 | [M-H]- | 1 | 900.6 | FALSE | 899.6 | Unit | 81 | Unit |
| SQDG_NEG | SQDG 18:1/22:2 | [M-H]- | 1 | 900.6 | FALSE | 899.6 | Unit | 225 | Unit |
| SQDGNH4 | SQDG 18:1/22:2 | [M+H]+ | 1 | 917.6 | FALSE | 918.6 | Unit | 339.3 | Unit |
| SQDGNH4 | SQDG 18:1/22:2 | [M+H]+ | 1 | 917.6 | FALSE | 918.6 | Unit | 393.3 | Unit |
| SQDG_NEG | SQDG 18:1/22:3 | [M-H]- | 1 | 898.6 | FALSE | 897.6 | Unit | 81 | Unit |
| SQDG_NEG | SQDG 18:1/22:3 | [M-H]- | 1 | 898.6 | FALSE | 897.6 | Unit | 225 | Unit |
| SQDGNH4 | SQDG 18:1/22:3 | [M+H]+ | 1 | 915.6 | FALSE | 916.6 | Unit | 339.3 | Unit |
| SQDGNH4 | SQDG 18:1/22:3 | [M+H]+ | 1 | 915.6 | FALSE | 916.6 | Unit | 391.3 | Unit |
| SQDGNH4 | SQDG 18:1/22:4 | [M+H]+ | 1 | 913.6 | FALSE | 914.6 | Unit | 339.3 | Unit |
| SQDGNH4 | SQDG 18:1/22:4 | [M+H]+ | 1 | 913.6 | FALSE | 914.6 | Unit | 389.3 | Unit |
| SQDG_NEG | SQDG 18:1/22:5 | [M-H]- | 1 | 894.5 | FALSE | 893.5 | Unit | 81 | Unit |
| SQDG_NEG | SQDG 18:1/22:5 | [M-H]- | 1 | 894.5 | FALSE | 893.5 | Unit | 225 | Unit |
| SQDGNH4 | SQDG 18:1/22:5 | [M+H]+ | 1 | 911.6 | FALSE | 912.6 | Unit | 339.3 | Unit |
| SQDGNH4 | SQDG 18:1/22:5 | [M+H]+ | 1 | 911.6 | FALSE | 912.6 | Unit | 387.3 | Unit |
| SQDG_NEG | SQDG 18:1/22:6 | [M-H]- | 1 | 892.5 | FALSE | 891.5 | Unit | 81 | Unit |
| SQDG_NEG | SQDG 18:1/22:6 | [M-H]- | 1 | 892.5 | FALSE | 891.5 | Unit | 225 | Unit |
| SQDGNH4 | SQDG 18:1/22:6 | [M+H]+ | 1 | 909.6 | FALSE | 910.6 | Unit | 339.3 | Unit |
| SQDGNH4 | SQDG 18:1/22:6 | [M+H]+ | 1 | 909.6 | FALSE | 910.6 | Unit | 385.3 | Unit |
| SQDG_NEG | SQDG 18:2/18:2 | [M-H]- | 1 | 842.5 | FALSE | 841.5 | Unit | 81 | Unit |
| SQDG_NEG | SQDG 18:2/18:2 | [M-H]- | 1 | 842.5 | FALSE | 841.5 | Unit | 225 | Unit |
| SQDGNH4 | SQDG 18:2/18:2 | [M+H]+ | 1 | 859.6 | FALSE | 860.6 | Unit | 337.3 | Unit |
| SQDG_NEG | SQDG 18:2/20:3 | [M-H]- | 1 | 868.5 | FALSE | 867.5 | Unit | 81 | Unit |
| SQDG_NEG | SQDG 18:2/20:3 | [M-H]- | 1 | 868.5 | FALSE | 867.5 | Unit | 225 | Unit |
| SQDGNH4 | SQDG 18:2/20:3 | [M+H]+ | 1 | 885.6 | FALSE | 886.6 | Unit | 337.3 | Unit |
| SQDGNH4 | SQDG 18:2/20:3 | [M+H]+ | 1 | 885.6 | FALSE | 886.6 | Unit | 363.3 | Unit |
| SQDG_NEG | SQDG 18:2/22:6 | [M-H]- | 1 | 890.5 | FALSE | 889.5 | Unit | 81 | Unit |
| SQDG_NEG | SQDG 18:2/22:6 | [M-H]- | 1 | 890.5 | FALSE | 889.5 | Unit | 225 | Unit |
| SQDGNH4 | SQDG 18:2/22:6 | [M+H]+ | 1 | 907.6 | FALSE | 908.6 | Unit | 337.3 | Unit |
| SQDGNH4 | SQDG 18:2/22:6 | [M+H]+ | 1 | 907.6 | FALSE | 908.6 | Unit | 385.3 | Unit |

| Dwell (m | Fragmen | CE (V) | Polarity |
|---|---|---|---|
| 5 | 250 | 60 | Negative |
| 5 | 250 | 60 | Negative |
| 5 | 250 | 60 | Negative |
| 5 | 250 | 60 | Negative |
| 5 | 185 | 26 | Positive |
| 5 | 185 | 26 | Positive |
| 5 | 250 | 60 | Negative |
| 5 | 250 | 60 | Negative |
| 5 | 185 | 26 | Positive |
| 5 | 185 | 26 | Positive |
| 5 | 250 | 60 | Negative |
| 5 | 250 | 60 | Negative |
| 5 | 185 | 26 | Positive |
| 5 | 185 | 26 | Positive |
| 5 | 250 | 60 | Negative |
| 5 | 250 | 60 | Negative |
| 5 | 185 | 26 | Positive |
| 5 | 185 | 26 | Positive |
| 5 | 250 | 60 | Negative |
| 5 | 250 | 60 | Negative |
| 5 | 185 | 26 | Positive |
| 5 | 185 | 26 | Positive |
| 5 | 250 | 60 | Negative |
| 5 | 250 | 60 | Negative |
| 5 | 185 | 26 | Positive |
| 5 | 185 | 26 | Positive |
| 5 | 250 | 60 | Negative |
| 5 | 250 | 60 | Negative |
| 5 | 250 | 60 | Negative |
| 5 | 250 | 60 | Negative |
| 5 | 185 | 26 | Positive |
| 5 | 185 | 26 | Positive |
| 5 | 250 | 60 | Negative |
| 5 | 250 | 60 | Negative |
| 5 | 185 | 26 | Positive |
| 5 | 185 | 26 | Positive |
| 5 | 250 | 60 | Negative |

| | | | |
|---|---|---|---|
| 5 | 250 | 60 | Negative |
| 5 | 185 | 26 | Positive |
| 5 | 185 | 26 | Positive |
| 5 | 250 | 60 | Negative |
| 5 | 250 | 60 | Negative |
| 5 | 185 | 26 | Positive |
| 5 | 185 | 26 | Positive |
| 5 | 250 | 60 | Negative |
| 5 | 250 | 60 | Negative |
| 5 | 250 | 60 | Negative |
| 5 | 250 | 60 | Negative |
| 5 | 185 | 26 | Positive |
| 5 | 250 | 60 | Negative |
| 5 | 250 | 60 | Negative |
| 5 | 250 | 60 | Negative |
| 5 | 250 | 60 | Negative |
| 5 | 185 | 26 | Positive |
| 5 | 185 | 26 | Positive |
| 5 | 250 | 60 | Negative |
| 5 | 250 | 60 | Negative |
| 5 | 185 | 26 | Positive |
| 5 | 185 | 26 | Positive |
| 5 | 250 | 60 | Negative |
| 5 | 250 | 60 | Negative |
| 5 | 185 | 26 | Positive |
| 5 | 185 | 26 | Positive |
| 5 | 250 | 60 | Negative |
| 5 | 250 | 60 | Negative |
| 5 | 185 | 26 | Positive |
| 5 | 185 | 26 | Positive |
| 5 | 250 | 60 | Negative |
| 5 | 250 | 60 | Negative |
| 5 | 185 | 26 | Positive |
| 5 | 185 | 26 | Positive |
| 5 | 250 | 60 | Negative |
| 5 | 250 | 60 | Negative |
| 5 | 185 | 26 | Positive |
| 5 | 185 | 26 | Positive |

| | | | |
|---|---|---|---|
| 5 | 250 | 60 | Negative |
| 5 | 250 | 60 | Negative |
| 5 | 185 | 26 | Positive |
| 5 | 185 | 26 | Positive |
| 5 | 250 | 60 | Negative |
| 5 | 250 | 60 | Negative |
| 5 | 185 | 26 | Positive |
| 5 | 185 | 26 | Positive |
| 5 | 185 | 26 | Positive |
| 5 | 185 | 26 | Positive |
| 5 | 250 | 60 | Negative |
| 5 | 250 | 60 | Negative |
| 5 | 185 | 26 | Positive |
| 5 | 185 | 26 | Positive |
| 5 | 250 | 60 | Negative |
| 5 | 250 | 60 | Negative |
| 5 | 185 | 26 | Positive |
| 5 | 185 | 26 | Positive |
| 5 | 250 | 60 | Negative |
| 5 | 250 | 60 | Negative |
| 5 | 185 | 26 | Positive |
| 5 | 250 | 60 | Negative |
| 5 | 250 | 60 | Negative |
| 5 | 185 | 26 | Positive |
| 5 | 185 | 26 | Positive |
| 5 | 250 | 60 | Negative |
| 5 | 250 | 60 | Negative |
| 5 | 185 | 26 | Positive |
| 5 | 185 | 26 | Positive |

| Compound group | Compound name | Compound formula | Ion speci | CAS | z | Monoisot | ISTD | Precurso |
|---|---|---|---|---|---|---|---|---|
| DG(18:2_20:4) | DG(18:2_20:4) | C42H82NO10P | [M+NH4]+ | | 1 | | FALSE | 658.5 |
| MGDG(16:0_16:1)a | MGDG(16:0_16:1)a | C41H77O10 | [M+H]+ | | 1 | | FALSE | 729.5 |
| MGDG(16:0_16:1)b | MGDG(16:0_16:1)b | C41H77O10 | [M+NH4]+ | | 1 | | FALSE | 746.6 |
| MGDG(18:0/18:0)a | MGDG(18:0/18:0)a | | [M+NH4]+ | | 1 | | FALSE | 760.6 |
| MGDG(18:0/18:0)b | MGDG(18:0/18:0)b | | [M+H]+ | | 1 | | FALSE | 743.6 |
| BisMeLPA(12:0e)a | BisMeLPA(12:0e)a | | [M+H]+ | | 1 | | FALSE | 369.2 |
| BisMeLPA(12:0e)b | BisMeLPA(12:0e)b | | [M+H]+ | | 1 | | FALSE | 369.2 |
| CarE(14:0+O)a | CarE(14:0+O)a | C21 H41 N O5 | [M+H]+ | | 1 | | FALSE | 388.3 |
| CarE(16:1)b | CarE(16:1)b | C23 H43 N O4 | [M+H]+ | | 1 | | FALSE | 398.3 |
| dhCer(d18:0/16:0) | dhCer(d18:0/16:0) | | [M+H]+ | | 1 | | FALSE | 540.5 |
| dhCer(d18:0/18:0) | dhCer(d18:0/18:0) | | [M+H]+ | | 1 | | FALSE | 568.6 |
| dhCer(d18:0/20:0) | dhCer(d18:0/20:0) | | [M+H]+ | | 1 | | FALSE | 596.6 |
| dhCer(d18:0/22:0) | dhCer(d18:0/22:0) | | [M+H]+ | | 1 | | FALSE | 624.6 |
| dhCer(d18:0/24:0) | dhCer(d18:0/24:0) | | [M+H]+ | | 1 | | FALSE | 652.7 |
| dhCer(d18:0/24:1) | dhCer(d18:0/24:1) | | [M+H]+ | | 1 | | FALSE | 650.6 |
| dhCer(d18:1/18:0) | dhCer(d18:1/18:0) | | [M+H]+ | | 1 | | FALSE | 566.6 |
| dhCer(d18:1/20:0) | dhCer(d18:1/20:0) | | [M+H]+ | | 1 | | FALSE | 594.6 |
| dhCer(d18:1/22:0) | dhCer(d18:1/22:0) | | [M+H]+ | | 1 | | FALSE | 622.6 |
| dhCer(d18:1/24:0) | dhCer(d18:1/24:0) | | [M+H]+ | | 1 | | FALSE | 650.7 |
| dhCer(d18:1/24:1) | dhCer(d18:1/24:1) | | [M+H]+ | | 1 | | FALSE | 648.6 |
| dhCer(d18:1/24:2) | dhCer(d18:1/24:2) | | [M+H]+ | | 1 | | FALSE | 646.6 |
| Cer(d18:1_26:0) | Cer(d18:1_26:0) | | [M+H]+ | | 1 | | FALSE | 678.7 |
| Cer(d18:2_24:0) | Cer(d18:2_24:0) | | [M+H]+ | | 1 | | FALSE | 648.6 |
| Cer(d18:2_28:0) | Cer(d18:2_28:0) | | [M+H]+ | | 1 | | FALSE | 704.7 |
| Cer(d19:2_31:0+O) | Cer(d19:2_31:0+O) | | [M+H]+ | | 1 | | FALSE | 760.8 |
| Cer(d20:2_26:0+O) | Cer(d20:2_26:0+O) | | [M+H]+ | | 1 | | FALSE | 704.7 |
| Cer(t18:0_26:0) | Cer(t18:0_26:0) | | [M+H]+ | | 1 | | FALSE | 696.7 |
| Cer(t20:0_25:0+O) | Cer(t20:0_25:0+O) | | [M+H]+ | | 1 | | FALSE | 710.7 |
| GM1(d18:1/16:0) | GM1(d18:1/16:0) | | [M+2H]2+ | | 1 | | FALSE | 760.1 |
| GM3(d18:1/16:0) | GM3(d18:1/16:0) | | [M+H]+ | | 1 | | FALSE | 1153.7 |
| GM3(d18:1/18:0) | GM3(d18:1/18:0) | | [M+H]+ | | 1 | | FALSE | 1181.8 |
| GM3(d18:1/20:0) | GM3(d18:1/20:0) | | [M+H]+ | | 1 | | FALSE | 1209.8 |
| GM3(d18:1/22:0) | GM3(d18:1/22:0) | | [M+H]+ | | 1 | | FALSE | 1237.8 |
| GM3(d18:1/24:0) | GM3(d18:1/24:0) | | [M+H]+ | | 1 | | FALSE | 1265.8 |
| GM3(d18:1/24:1) | GM3(d18:1/24:1) | | [M+H]+ | | 1 | | FALSE | 1263.8 |
| GM3(d18:2/24:1) | GM3(d18:2/24:1) | | [M+H]+ | | 1 | | FALSE | 1261.8 |
| Hex2Cer(d18:1/16:0) | Hex2Cer(d18:1/16:0) | | [M+H]+ | | 1 | | FALSE | 862.6 |

| | | | | | | |
|---|---|---|---|---|---|---|
| Hex2Cer(d18:1/18:0) | Hex2Cer(d18:1/18:0) | | [M+H]+ | 1 | FALSE | 890.7 |
| Hex2Cer(d18:1/20:0) | Hex2Cer(d18:1/20:0) | | [M+H]+ | 1 | FALSE | 918.7 |
| Hex2Cer(d18:1/22:0) | Hex2Cer(d18:1/22:0) | | [M+H]+ | 1 | FALSE | 946.7 |
| Hex2Cer(d18:1/24:0) | Hex2Cer(d18:1/24:0) | | [M+H]+ | 1 | FALSE | 974.8 |
| Hex2Cer(d18:1/24:1) | Hex2Cer(d18:1/24:1) | | [M+H]+ | 1 | FALSE | 972.7 |
| Hex3Cer(d18:1/16:0) | Hex3Cer(d18:1/16:0) | | [M+H]+ | 1 | FALSE | 1024.7 |
| Hex3Cer(d18:1/18:0) | Hex3Cer(d18:1/18:0) | | [M+H]+ | 1 | FALSE | 1052.7 |
| Hex3Cer(d18:1/20:0) | Hex3Cer(d18:1/20:0) | | [M+H]+ | 1 | FALSE | 1080.7 |
| Hex3Cer(d18:1/22:0) | Hex3Cer(d18:1/22:0) | | [M+H]+ | 1 | FALSE | 1108.8 |
| Hex3Cer(d18:1/24:0) | Hex3Cer(d18:1/24:0) | | [M+H]+ | 1 | FALSE | 1136.8 |
| Hex3Cer(d18:1/24:1) | Hex3Cer(d18:1/24:1) | | [M+H]+ | 1 | FALSE | 1134.8 |
| HexCer(d16:1/18:0) | HexCer(d16:1/18:0) | | [M+H]+ | 1 | FALSE | 700.6 |
| HexCer(d16:1/20:0) | HexCer(d16:1/20:0) | | [M+H]+ | 1 | FALSE | 728.6 |
| HexCer(d16:1/22:0) | HexCer(d16:1/22:0) | | [M+H]+ | 1 | FALSE | 756.7 |
| HexCer(d16:1/24:0) | HexCer(d16:1/24:0) | | [M+H]+ | 1 | FALSE | 784.7 |
| HexCer(d18:1/16:0) | HexCer(d18:1/16:0) | | [M+H]+ | 1 | FALSE | 700.6 |
| HexCer(d18:1/18:0) | HexCer(d18:1/18:0) | | [M+H]+ | 1 | FALSE | 728.6 |
| HexCer(d18:1/20:0) | HexCer(d18:1/20:0) | | [M+H]+ | 1 | FALSE | 756.6 |
| HexCer(d18:1/22:0) | HexCer(d18:1/22:0) | | [M+H]+ | 1 | FALSE | 784.7 |
| HexCer(d18:1/24:0) | HexCer(d18:1/24:0) | | [M+H]+ | 1 | FALSE | 812.7 |
| HexCer(d18:1/24:1) | HexCer(d18:1/24:1) | | [M+H]+ | 1 | FALSE | 810.7 |
| HexCer(d18:2/18:0) | HexCer(d18:2/18:0) | | [M+H]+ | 1 | FALSE | 726.6 |
| HexCer(d18:2/20:0) | HexCer(d18:2/20:0) | | [M+H]+ | 1 | FALSE | 754.6 |
| HexCer(d18:2/22:0) | HexCer(d18:2/22:0) | | [M+H]+ | 1 | FALSE | 782.7 |
| HexCer(d18:2/24:0) | HexCer(d18:2/24:0) | | [M+H]+ | 1 | FALSE | 810.7 |
| LPC(12:0) | LPC(12:0) | C20H46N2O7P | [M+H]+ | 1 | FALSE | 457.3 |
| LPC(14:0) [sn1] | LPC(14:0) [sn1] | C42H82NO10P | [M+H]+ | 1 | FALSE | 468.3 |
| LPC(15:0) [sn1] | LPC(15:0) [sn1] | C42H82NO10P | [M+H]+ | 1 | FALSE | 482.3 |
| LPC(15-MHDA) [sn1] | LPC(15-MHDA) [sn1] | C42H82NO10P | [M+H]+ | 1 | FALSE | 510.4 |
| LPC(15-MHDA) [sn1] | LPC(15-MHDA) [sn1] | C42H82NO10P | [M+H]+ | 1 | FALSE | 510.4 |
| LPC(15-MHDA) [sn2] | LPC(15-MHDA) [sn2] | C42H82NO10P | [M+H]+ | 1 | FALSE | 510.4 |
| LPC(16:0) [sn1] | LPC(16:0) [sn1] | C42H82NO10P | [M+H]+ | 1 | FALSE | 496.3 |
| LPC(16:1) [sn1] | LPC(16:1) [sn1] | C42H82NO10P | [M+H]+ | 1 | FALSE | 494.3 |
| LPC(16:2) [sn1] | LPC(16:2) [sn1] | C42H82NO10P | [M+H]+ | 1 | FALSE | 492.3 |
| LPC(18:0) [sn1] | LPC(18:0) [sn1] | C42H82NO10P | [M+H]+ | 1 | FALSE | 524.4 |
| LPC(18:1) [sn1] | LPC(18:1) [sn1] | C42H82NO10P | [M+H]+ | 1 | FALSE | 522.4 |
| LPC(18:2) [sn1] | LPC(18:2) [sn1] | C42H82NO10P | [M+H]+ | 1 | FALSE | 520.3 |
| LPC(18:3) [sn1] (b) | LPC(18:3) [sn1] (b) | C42H82NO10P | [M+H]+ | 1 | FALSE | 518.3 |

| | | | | | | |
|---|---|---|---|---|---|---|
| LPC(18:3) [sn2] (a) | LPC(18:3) [sn2] (a) | C42H82NO10P | [M+H]+ | 1 | FALSE | 518.3 |
| LPC(18:4) [sn2] (a) | LPC(18:4) [sn2] (a) | C42H82NO10P | [M+H]+ | 1 | FALSE | 516.3 |
| LPC(20:0) [sn1] | LPC(20:0) [sn1] | C42H82NO10P | [M+H]+ | 1 | FALSE | 552.4 |
| LPC(20:1) [sn1] | LPC(20:1) [sn1] | C42H82NO10P | [M+H]+ | 1 | FALSE | 550.4 |
| LPC(20:2) [sn1] | LPC(20:2) [sn1] | C42H82NO10P | [M+H]+ | 1 | FALSE | 548.4 |
| LPC(20:3) [sn2] | LPC(20:3) [sn2] | C42H82NO10P | [M+H]+ | 1 | FALSE | 546.4 |
| LPC(20:4) [sn1] | LPC(20:4) [sn1] | C42H82NO10P | [M+H]+ | 1 | FALSE | 544.3 |
| LPC(20:4) [sn2] | LPC(20:4) [sn2] | C42H82NO10P | [M+H]+ | 1 | FALSE | 544.3 |
| LPC(20:5) [sn1] | LPC(20:5) [sn1] | C42H82NO10P | [M+H]+ | 1 | FALSE | 542.3 |
| LPC(22:0) [sn1] | LPC(22:0) [sn1] | C42H82NO10P | [M+H]+ | 1 | FALSE | 580.4 |
| LPC(22:1) [sn1] | LPC(22:1) [sn1] | C42H82NO10P | [M+H]+ | 1 | FALSE | 578.4 |
| LPC(22:4) [sn1] | LPC(22:4) [sn1] | C42H82NO10P | [M+H]+ | 1 | FALSE | 572.4 |
| LPC(22:5) (n3) [sn1] | LPC(22:5) (n3) [sn1] [ | C42H82NO10P | [M+H]+ | 1 | FALSE | 570.4 |
| LPC(22:5) [sn1] (n3)/ | LPC(22:5) [sn1] (n3)/L | C42H82NO10P | [M+H]+ | 1 | FALSE | 570.4 |
| LPC(22:5) [sn1] (n6) | LPC(22:5) [sn1] (n6) | C42H82NO10P | [M+H]+ | 1 | FALSE | 570.4 |
| LPC(22:5) [sn2] (n3) | LPC(22:5) [sn2] (n3) | C42H82NO10P | [M+H]+ | 1 | FALSE | 570.4 |
| LPC(22:6) [sn1] | LPC(22:6) [sn1] | C42H82NO10P | [M+H]+ | 1 | FALSE | 568.3 |
| LPC(24:0) [sn1] | LPC(24:0) [sn1] | C42H82NO10P | [M+H]+ | 1 | FALSE | 608.5 |
| LPC(26:0) [sn1] | LPC(26:0) [sn1] | C42H82NO10P | [M+H]+ | 1 | FALSE | 636.5 |
| LPC(32:1) [sn1] | LPC(32:1) [sn1] | C42H82NO10P | [M+H]+ | 1 | FALSE | 718.6 |
| LPC(O-16:0) | LPC(O-16:0) | C42H82NO10P | [M+H]+ | 1 | FALSE | 482.4 |
| LPC(O-18:0) | LPC(O-18:0) | C42H82NO10P | [M+H]+ | 1 | FALSE | 510.4 |
| LPC(O-18:1) | LPC(O-18:1) | C42H82NO10P | [M+H]+ | 1 | FALSE | 508.4 |
| LPC(O-20:0) | LPC(O-20:0) | C42H82NO10P | [M+H]+ | 1 | FALSE | 538.4 |
| LPC(O-20:1) | LPC(O-20:1) | C42H82NO10P | [M+H]+ | 1 | FALSE | 536.4 |
| LPC(O-22:0) | LPC(O-22:0) | C42H82NO10P | [M+H]+ | 1 | FALSE | 566.5 |
| LPC(O-22:1) | LPC(O-22:1) | C42H82NO10P | [M+H]+ | 1 | FALSE | 564.4 |
| LPC(O-24:0) | LPC(O-24:0) | C42H82NO10P | [M+H]+ | 1 | FALSE | 594.5 |
| LPC(O-24:1) | LPC(O-24:1) | C42H82NO10P | [M+H]+ | 1 | FALSE | 592.5 |
| LPC(O-24:2) | LPC(O-24:2) | C42H82NO10P | [M+H]+ | 1 | FALSE | 590.5 |
| LPC(P-16:0) | LPC(P-16:0) | C42H82NO10P | [M+H]+ | 1 | FALSE | 480.3 |
| LPC(P-17:0) (a) | LPC(P-17:0) (a) | C42H82NO10P | [M+H]+ | 1 | FALSE | 494.3 |
| LPC(P-18:0) | LPC(P-18:0) | C42H82NO10P | [M+H]+ | 1 | FALSE | 508.3 |
| LPC(P-18:1) | LPC(P-18:1) | C42H82NO10P | [M+H]+ | 1 | FALSE | 506.3 |
| LPC(P-20:0) | LPC(P-20:0) | C42H82NO10P | [M+H]+ | 1 | FALSE | 536.3 |
| MePC(29:0) | MePC(29:0) | $C_{38}H_{76}NO_8P$ | [M+H]+ | 1 | FALSE | 722.5 |
| MePC(32:1) | MePC(32:1) | $C_{41}H_{80}NO_8P$ | [M+H]+ | 1 | FALSE | 762.6 |
| MePC(35:0) | MePC(35:0) | $C_{44}H_{88}NO_8P$ | [M+H]+ | 1 | FALSE | 802.6 |

| | | | | | | |
|---|---|---|---|---|---|---|
| LPE(16:0) [sn1] | LPE(16:0) [sn1] | C42H82NO10P | [M+H]+ | 1 | FALSE | 454.3 |
| LPE(18:0) [sn1] | LPE(18:0) [sn1] | C42H82NO10P | [M+H]+ | 1 | FALSE | 482.3 |
| LPE(18:1) [sn1] | LPE(18:1) [sn1] | C42H82NO10P | [M+H]+ | 1 | FALSE | 480.3 |
| LPE(18:2) [sn1] | LPE(18:2) [sn1] | C42H82NO10P | [M+H]+ | 1 | FALSE | 478.3 |
| LPE(20:4) [sn2] | LPE(20:4) [sn2] | C42H82NO10P | [M+H]+ | 1 | FALSE | 502.3 |
| LPE(22:6) [sn2] | LPE(22:6) [sn2] | C42H82NO10P | [M+H]+ | 1 | FALSE | 526.3 |
| LPE(P-16:0) | LPE(P-16:0) | C42H82NO10P | [M+H]+ | 1 | FALSE | 438.3 |
| LPE(P-18:0) | LPE(P-18:0) | C42H82NO10P | [M+H]+ | 1 | FALSE | 466.3 |
| LPE(P-18:1) | LPE(P-18:1) | C42H82NO10P | [M+H]+ | 1 | FALSE | 464.3 |

| MS1 res | Product r | MS2 res | Dwell (m | Fragmen | CE (V) | Polarity |
|---|---|---|---|---|---|---|
| Unit | 337.2 | Unit | 20 | 166 | 21 | Positive |
| Unit | 257.2 | Unit | 20 | 166 | 21 | Positive |
| Unit | 313.2 | Unit | 20 | 166 | 21 | Positive |
| Unit | 341.3 | Unit | 20 | 166 | 21 | Positive |
| Unit | 284.3 | Unit | 20 | 166 | 21 | Positive |
| Unit | 153 | Unit | 20 | 166 | 21 | Positive |
| Unit | 255.2 | Unit | 20 | 166 | 21 | Positive |
| Unit | 85.02 | Unit | 20 | 166 | 21 | Positive |
| Unit | 85.02 | Unit | 20 | 166 | 21 | Positive |
| Unit | 284.3 | Unit | 20 | 166 | 31 | Positive |
| Unit | 284.3 | Unit | 20 | 166 | 31 | Positive |
| Unit | 284.3 | Unit | 20 | 166 | 31 | Positive |
| Unit | 284.3 | Unit | 20 | 166 | 31 | Positive |
| Unit | 284.3 | Unit | 20 | 166 | 31 | Positive |
| Unit | 284.3 | Unit | 20 | 166 | 31 | Positive |
| Unit | 282.3 | Unit | 20 | 166 | 31 | Positive |
| Unit | 282.3 | Unit | 20 | 166 | 31 | Positive |
| Unit | 282.3 | Unit | 20 | 166 | 31 | Positive |
| Unit | 282.3 | Unit | 20 | 166 | 31 | Positive |
| Unit | 282.3 | Unit | 20 | 166 | 31 | Positive |
| Unit | 282.3 | Unit | 20 | 166 | 31 | Positive |
| Unit | 282.3 | Unit | 20 | 166 | 31 | Positive |
| Unit | 280.3 | Unit | 20 | 166 | 31 | Positive |
| Unit | 280.3 | Unit | 20 | 166 | 31 | Positive |
| Unit | 264.3 | Unit | 20 | 166 | 31 | Positive |
| Unit | 264.3 | Unit | 20 | 166 | 31 | Positive |
| Unit | 264.3 | Unit | 20 | 166 | 31 | Positive |
| Unit | 264.3 | Unit | 20 | 166 | 31 | Positive |
| Unit | 366.2 | Unit | 20 | 166 | 31 | Positive |
| Unit | 264.3 | Unit | 20 | 166 | 57 | Positive |
| Unit | 264.3 | Unit | 20 | 166 | 57 | Positive |
| Unit | 264.3 | Unit | 20 | 166 | 57 | Positive |
| Unit | 264.3 | Unit | 20 | 166 | 57 | Positive |
| Unit | 264.3 | Unit | 20 | 166 | 57 | Positive |
| Unit | 264.3 | Unit | 20 | 166 | 57 | Positive |
| Unit | 262.3 | Unit | 20 | 166 | 57 | Positive |
| Unit | 264.3 | Unit | 20 | 166 | 53 | Positive |

| | | | | | | |
|---|---|---|---|---|---|---|
| Unit | 264.3 | Unit | 20 | 166 | 53 | Positive |
| Unit | 264.3 | Unit | 20 | 166 | 53 | Positive |
| Unit | 264.3 | Unit | 20 | 166 | 53 | Positive |
| Unit | 264.3 | Unit | 20 | 166 | 53 | Positive |
| Unit | 264.3 | Unit | 20 | 166 | 53 | Positive |
| Unit | 264.3 | Unit | 20 | 166 | 53 | Positive |
| Unit | 264.3 | Unit | 20 | 166 | 53 | Positive |
| Unit | 264.3 | Unit | 20 | 166 | 53 | Positive |
| Unit | 264.3 | Unit | 20 | 166 | 53 | Positive |
| Unit | 264.3 | Unit | 20 | 166 | 53 | Positive |
| Unit | 264.3 | Unit | 20 | 166 | 53 | Positive |
| Unit | 236.3 | Unit | 20 | 166 | 33 | Positive |
| Unit | 236.3 | Unit | 20 | 166 | 33 | Positive |
| Unit | 236.3 | Unit | 20 | 166 | 33 | Positive |
| Unit | 236.3 | Unit | 20 | 166 | 33 | Positive |
| Unit | 264.3 | Unit | 20 | 166 | 33 | Positive |
| Unit | 264.3 | Unit | 20 | 166 | 33 | Positive |
| Unit | 264.3 | Unit | 20 | 166 | 33 | Positive |
| Unit | 264.3 | Unit | 20 | 166 | 33 | Positive |
| Unit | 264.3 | Unit | 20 | 166 | 33 | Positive |
| Unit | 264.3 | Unit | 20 | 166 | 33 | Positive |
| Unit | 262.3 | Unit | 20 | 166 | 33 | Positive |
| Unit | 262.3 | Unit | 20 | 166 | 33 | Positive |
| Unit | 262.3 | Unit | 20 | 166 | 33 | Positive |
| Unit | 262.3 | Unit | 20 | 166 | 33 | Positive |
| Unit | 184.1 | Unit | 20 | 166 | 21 | Positive |
| Unit | 184.1 | Unit | 20 | 166 | 21 | Positive |
| Unit | 184.1 | Unit | 20 | 166 | 21 | Positive |
| Unit | 184.1 | Unit | 20 | 166 | 21 | Positive |
| Unit | 104.1 | Unit | 20 | 166 | 21 | Positive |
| Unit | 184.1 | Unit | 20 | 166 | 21 | Positive |
| Unit | 184.1 | Unit | 20 | 166 | 21 | Positive |
| Unit | 184.1 | Unit | 20 | 166 | 21 | Positive |
| Unit | 184.1 | Unit | 20 | 166 | 21 | Positive |
| Unit | 184.1 | Unit | 20 | 166 | 21 | Positive |
| Unit | 184.1 | Unit | 20 | 166 | 21 | Positive |
| Unit | 184.1 | Unit | 20 | 166 | 21 | Positive |
| Unit | 184.1 | Unit | 20 | 166 | 21 | Positive |

| | | | | | | |
|---|---|---|---|---|---|---|
| Unit | 184.1 | Unit | 20 | 166 | 21 | Positive |
| Unit | 184.1 | Unit | 20 | 166 | 21 | Positive |
| Unit | 184.1 | Unit | 20 | 166 | 21 | Positive |
| Unit | 184.1 | Unit | 20 | 166 | 21 | Positive |
| Unit | 184.1 | Unit | 20 | 166 | 21 | Positive |
| Unit | 184.1 | Unit | 20 | 166 | 21 | Positive |
| Unit | 184.1 | Unit | 20 | 166 | 21 | Positive |
| Unit | 184.1 | Unit | 20 | 166 | 21 | Positive |
| Unit | 184.1 | Unit | 20 | 166 | 21 | Positive |
| Unit | 184.1 | Unit | 20 | 166 | 21 | Positive |
| Unit | 184.1 | Unit | 20 | 166 | 21 | Positive |
| Unit | 184.1 | Unit | 20 | 166 | 21 | Positive |
| Unit | 184.1 | Unit | 20 | 166 | 21 | Positive |
| Unit | 184.1 | Unit | 20 | 166 | 21 | Positive |
| Unit | 184.1 | Unit | 20 | 166 | 21 | Positive |
| Unit | 184.1 | Unit | 20 | 166 | 21 | Positive |
| Unit | 184.1 | Unit | 20 | 166 | 21 | Positive |
| Unit | 184.1 | Unit | 20 | 166 | 21 | Positive |
| Unit | 184.1 | Unit | 20 | 166 | 21 | Positive |
| Unit | 184.1 | Unit | 20 | 166 | 21 | Positive |
| Unit | 184.1 | Unit | 20 | 166 | 21 | Positive |
| Unit | 184.1 | Unit | 20 | 166 | 21 | Positive |
| Unit | 184.1 | Unit | 20 | 166 | 21 | Positive |
| Unit | 184.1 | Unit | 20 | 166 | 21 | Positive |
| Unit | 184.1 | Unit | 20 | 166 | 21 | Positive |
| Unit | 184.1 | Unit | 20 | 166 | 21 | Positive |
| Unit | 184.1 | Unit | 20 | 166 | 21 | Positive |
| Unit | 184.1 | Unit | 20 | 166 | 21 | Positive |
| Unit | 184.1 | Unit | 20 | 166 | 21 | Positive |
| Unit | 184.1 | Unit | 20 | 166 | 21 | Positive |
| Unit | 184.1 | Unit | 20 | 166 | 21 | Positive |
| Unit | 184.1 | Unit | 20 | 166 | 21 | Positive |
| Unit | 184.1 | Unit | 20 | 166 | 21 | Positive |
| Unit | 184.1 | Unit | 20 | 166 | 21 | Positive |
| Unit | 184.1 | Unit | 20 | 166 | 21 | Positive |
| Unit | 184.1 | Unit | 20 | 166 | 21 | Positive |
| Unit | 184.1 | Unit | 20 | 166 | 21 | Positive |
| Unit | 184.1 | Unit | 20 | 166 | 21 | Positive |

| | | | | | | |
|---|---|---|---|---|---|---|
| Unit | 313.3 | Unit | 20 | 166 | 17 | Positive |
| Unit | 341.3 | Unit | 20 | 166 | 17 | Positive |
| Unit | 339.3 | Unit | 20 | 166 | 17 | Positive |
| Unit | 337.3 | Unit | 20 | 166 | 17 | Positive |
| Unit | 361.3 | Unit | 20 | 166 | 17 | Positive |
| Unit | 385.3 | Unit | 20 | 166 | 17 | Positive |
| Unit | 266.4 | Unit | 20 | 166 | 19 | Positive |
| Unit | 294.4 | Unit | 20 | 166 | 19 | Positive |
| Unit | 292.4 | Unit | 20 | 166 | 19 | Positive |

| label | Compound group | Compoui | Ion speci | CAS | z | Monoisot | ISTD | Precurso |
|---|---|---|---|---|---|---|---|---|
| BMPNH4 | BMP_14:0/18:1 | | [M+NH4]+ | | | | FALSE | 738.6 |
| BMPNH4 | BMP_14:0/18:1 | | [M+NH4]+ | | | | FALSE | 738.6 |
| BMPNH4 | BMP_16:0/16:0 | | [M+NH4]+ | | | | FALSE | 740.5 |
| BMPNH4 | BMP_16:0/16:1 | | [M+NH4]+ | | | | FALSE | 738.5 |
| BMPNH4 | BMP_16:0/16:1 dup | | [M+NH4]+ | | | | FALSE | 738.5 |
| BMPNH4 | BMP_16:0/18:0 | | [M+NH4]+ | | | | FALSE | 768.5 |
| BMPNH4 | BMP_16:0/18:0 | | [M+NH4]+ | | | | FALSE | 768.5 |
| BMPNH4 | BMP_16:0/18:1 | | [M+NH4]+ | | | | FALSE | 766.6 |
| BMPNH4 | BMP_16:0/18:1 | | [M+NH4]+ | | | | FALSE | 766.6 |
| BMPNH4 | BMP(16:0/18:2) | | [M+NH4]+ | | | | FALSE | 764.5 |
| BMPNH4 | BMP(16:0/18:2) | | [M+NH4]+ | | | | FALSE | 764.5 |
| BMPNH4 | BMP_16:0/18:1 | | [M+NH4]+ | | | | FALSE | 766.6 |
| BMPNH4 | BMP_16:0/22:6 | | [M+NH4]+ | | | | FALSE | 812.5 |
| BMPNH4 | BMP_16:0/22:6 | | [M+NH4]+ | | | | FALSE | 812.5 |
| BMPNH4 | BMP_16:1/16:1 | | [M+NH4]+ | | | | FALSE | 736.5 |
| BMPNH4 | BMP_16:1/18:1 | | [M+NH4]+ | | | | FALSE | 764.5 |
| BMPNH4 | BMP_16:1/18:1 | | [M+NH4]+ | | | | FALSE | 764.5 |
| BMPNH4 | BMP_16:1/18:2 | | [M+NH4]+ | | | | FALSE | 762.5 |
| BMPNH4 | BMP_16:1/20:3 | | [M+NH4]+ | | | | FALSE | 788.6 |
| BMPNH4 | BMP_16:1/22:6 | | [M+NH4]+ | | | | FALSE | 810.5 |
| BMPNH4 | BMP_16:1/22:6 | | [M+NH4]+ | | | | FALSE | 810.5 |
| BMPNH4 | BMP_18:0/18:0 | | [M+NH4]+ | | | | FALSE | 796.6 |
| BMPNH4 | BMP_18:0/18:1 | | [M+NH4]+ | | | | FALSE | 794.6 |
| BMPNH4 | BMP_18:0/18:1 | | [M+NH4]+ | | | | FALSE | 794.6 |
| BMPNH4 | BMP_18:1/18:1 | | [M+NH4]+ | | | | FALSE | 792.6 |
| BMPNH4 | BMP_18:1/18:2 | | [M+NH4]+ | | | | FALSE | 790.6 |
| BMPNH4 | BMP_18:1/18:2 | | [M+NH4]+ | | | | FALSE | 790.6 |
| BMPNH4 | BMP_18:1/20:0 | | [M+NH4]+ | | | | FALSE | 822.6 |
| BMPNH4 | BMP_18:1/20:1 | | [M+NH4]+ | | | | FALSE | 820.6 |
| BMPNH4 | BMP_18:1/20:1 | | [M+NH4]+ | | | | FALSE | 820.6 |
| BMPNH4 | BMP_18:1/20:2 | | [M+NH4]+ | | | | FALSE | 818.6 |
| BMPNH4 | BMP_18:1/20:3 | | [M+NH4]+ | | | | FALSE | 816.6 |
| BMPNH4 | BMP_18:1/20:4 | | [M+NH4]+ | | | | FALSE | 814.6 |
| BMPNH4 | BMP_18:1/22:1 | | [M+NH4]+ | | | | FALSE | 848.6 |
| BMPNH4 | BMP_18:1/22:2 | | [M+NH4]+ | | | | FALSE | 846.6 |
| BMPNH4 | BMP_18:1/22:3 | | [M+NH4]+ | | | | FALSE | 844.6 |
| BMPNH4 | BMP_18:1/22:4 | | [M+NH4]+ | | | | FALSE | 842.6 |

| | | | | | | |
|---|---|---|---|---|---|---|
| BMPNH4 | BMP_18:1/22:5 | | [M+NH4]+ | | FALSE | 840.6 |
| BMPNH4 | BMP_18:1/22:6 | | [M+NH4]+ | | FALSE | 838.6 |
| BMPNH4 | BMP_18:1/22:6 | | [M+NH4]+ | | FALSE | 838.6 |
| BMPNH4 | BMP_18:2/18:2 | | [M+NH4]+ | | FALSE | 788.5 |
| BMPNH4 | BMP_18:2/20:3 | | [M+NH4]+ | | FALSE | 814.6 |
| BMPNH4 | BMP_18:2/22:6 | | [M+NH4]+ | | FALSE | 836.5 |
| BMPNH4 | BMP_18:4/22:6 | | [M+NH4]+ | | FALSE | 832.5 |
| BMPNH4 | BMP_20:4/20:4 | | [M+NH4]+ | | FALSE | 836.5 |
| BMPNH4 | BMP_20:4/22:6 | | [M+NH4]+ | | FALSE | 860.5 |
| BMPNH4 | BMP_20:5/22:6 | | [M+NH4]+ | | FALSE | 858.5 |
| BMPNH4 | BMP_22:5/22:5 | | [M+NH4]+ | | FALSE | 888.6 |
| BMPNH4 | BMP_22:5/22:6 | | [M+NH4]+ | | FALSE | 886.6 |
| BMPNH4 | BMP_22:6/22:6 | | [M+NH4]+ | | FALSE | 884.5 |
| HemiBMI | HB_16:0/18:1/18:1 | C60H11 | [M+NH4]+ | | FALSE | 1030.8 |
| HemiBMI | HB_18:0/18:1/18:1 | C60H11 | [M+NH4]+ | | FALSE | 1058.8 |
| HemiBMI | HB_18:1/18:1/18:1 | C60H11 | [M+NH4]+ | 1 | FALSE | 1056.8 |
| LPG | LPG_16:0 | | [M-H]- | | FALSE | 483.3 |
| LPG | LPG_18:0 | | [M-H]- | | FALSE | 511.3 |
| LPG | LPG_18:1 | C24H47 | [M-H]- | 1 | FALSE | 509.3 |
| LPG | LPG_18:2 | | [M-H]- | | FALSE | 507.3 |
| LPG | LPG_20:4 | | [M-H]- | | FALSE | 531.3 |
| LPG | LPG_20:5 | | [M-H]- | | FALSE | 529.3 |
| LPG | LPG_22:4 | | [M-H]- | | FALSE | 559.3 |
| LPG | LPG_22:5 | | [M-H]- | | FALSE | 557.3 |
| LPG | LPG_22:6 | | [M-H]- | | FALSE | 555.3 |
| PC | PC_18:1 | | [M+H]+ | | FALSE | 786.6 |
| PG NH4 | PG_14:0/16:0 | | [M+NH4]+ | | FALSE | 712.4 |
| PG NH4 | PG_14:0/18:1 | | [M+NH4]+ | | FALSE | 738.6 |
| PG NH4 | PG_16:0/16:0 | | [M+NH4]+ | | FALSE | 740.5 |
| PG NH4 | PG_16:0/16:1 | | [M+NH4]+ | | FALSE | 738.5 |
| PG NH4 | PG_16:0/18:0 | | [M+NH4]+ | | FALSE | 768.5 |
| PG NH4 | PG_16:0/18:1 | | [M+NH4]+ | | FALSE | 766.6 |
| PG NH4 | PG_16:0/20:4 S18:2 and S16:1/20:3 | | [M+NH4]+ | | FALSE | 788.6 |
| PG NH4 | PG_16:0/22:6 | | [M+NH4]+ | | FALSE | 812.5 |
| PG NH4 | PG_16:1/16:1 | | [M+NH4]+ | | FALSE | 736.5 |
| PG NH4 | PG_16:1/18:1 | | [M+NH4]+ | | FALSE | 764.5 |
| PG NH4 | PG_16:1/18:2 | | [M+NH4]+ | | FALSE | 762.5 |
| PG NH4 | PG_16:1/22:6 | | [M+NH4]+ | | FALSE | 810.5 |

| | | | | |
|---|---|---|---|---|
| PG NH4 | PG_18:0/18:0 | [M+NH4]+ | FALSE | 796.6 |
| PG NH4 | PG_18:0/18:1 | [M+NH4]+ | FALSE | 794.6 |
| PG NH4 | PG_18:1/18:1 | [M+NH4]+ | FALSE | 792.6 |
| PG NH4 | PG_18:1/18:2 | [M+NH4]+ | FALSE | 790.6 |
| PG NH4 | PG_18:1/20:0 S18:0/20:1 | [M+NH4]+ | FALSE | 822.6 |
| PG NH4 | PG_18:1/20:1 | [M+NH4]+ | FALSE | 820.6 |
| PG NH4 | PG_18:1/20:2 | [M+NH4]+ | FALSE | 818.6 |
| PG NH4 | PG_18:1/20:3 | [M+NH4]+ | FALSE | 816.6 |
| PG NH4 | PG_18:1/20:4 S18:2/20:3 | [M+NH4]+ | FALSE | 814.6 |
| PG NH4 | PG_18:1/22:1 | [M+NH4]+ | FALSE | 848.6 |
| PG NH4 | PG_18:1/22:2 | [M+NH4]+ | FALSE | 846.6 |
| PG NH4 | PG_18:1/22:3 | [M+NH4]+ | FALSE | 844.6 |
| PG NH4 | PG_18:1/22:4 | [M+NH4]+ | FALSE | 842.6 |
| PG NH4 | PG_18:1/22:5 | [M+NH4]+ | FALSE | 840.6 |
| PG NH4 | PG_18:1/22:6 | [M+NH4]+ | FALSE | 838.6 |
| PG NH4 | PG_18:2/22:6 | [M+NH4]+ | FALSE | 836.5 |
| PG NH4 | PG_18:4/22:6 | [M+NH4]+ | FALSE | 832.5 |
| PG NH4 | PG_20:4/22:6 | [M+NH4]+ | FALSE | 860.5 |
| PG NH4 | PG_20:5/22:6 | [M+NH4]+ | FALSE | 858.5 |
| PG NH4 | PG_22:4/22:6/ S22:5 | [M+NH4]+ | FALSE | 888.6 |
| PG NH4 | PG_22:5/22:6 | [M+NH4]+ | FALSE | 886.6 |
| PG NH4 | PG_22:6/22:6 | [M+NH4]+ | FALSE | 884.5 |
| PC | POPC | [M+H]+ | FALSE | 760.6 |
| PC | POPC | [M+H]+ | FALSE | 760.6 |

| MS1 res | Product r | MS2 res | Dwell (m | Fragmen | CE (V) | Polarity |
|---|---|---|---|---|---|---|
| Unit | 285.3 | Unit | 5 | 150 | 27 | Positive |
| Unit | 339.3 | Unit | 5 | 150 | 27 | Positive |
| Unit | 313.5 | Unit | 5 | 150 | 27 | Positive |
| Unit | 311.5 | Unit | 5 | 150 | 27 | Positive |
| Unit | 313.5 | Unit | 5 | 150 | 27 | Positive |
| Unit | 313.5 | Unit | 5 | 150 | 27 | Positive |
| Unit | 341.3 | Unit | 5 | 150 | 27 | Positive |
| Unit | 313.5 | Unit | 5 | 150 | 27 | Positive |
| Unit | 339.3 | Unit | 5 | 150 | 27 | Positive |
| Unit | 313.5 | Unit | 5 | 150 | 27 | Positive |
| Unit | 337.3 | Unit | 5 | 150 | 27 | Positive |
| Unit | 339.3 | Unit | 5 | 150 | 27 | Positive |
| Unit | 313.5 | Unit | 5 | 150 | 27 | Positive |
| Unit | 385.3 | Unit | 5 | 150 | 27 | Positive |
| Unit | 311.5 | Unit | 5 | 150 | 27 | Positive |
| Unit | 311.5 | Unit | 5 | 150 | 27 | Positive |
| Unit | 339.3 | Unit | 5 | 150 | 27 | Positive |
| Unit | 337.3 | Unit | 5 | 150 | 27 | Positive |
| Unit | 311.3 | Unit | 5 | 150 | 27 | Positive |
| Unit | 311.3 | Unit | 5 | 150 | 27 | Positive |
| Unit | 385.3 | Unit | 5 | 150 | 27 | Positive |
| Unit | 341.3 | Unit | 5 | 150 | 27 | Positive |
| Unit | 339.3 | Unit | 5 | 150 | 27 | Positive |
| Unit | 341.3 | Unit | 5 | 150 | 27 | Positive |
| Unit | 339.3 | Unit | 5 | 150 | 27 | Positive |
| Unit | 337.3 | Unit | 5 | 150 | 27 | Positive |
| Unit | 339.3 | Unit | 5 | 150 | 27 | Positive |
| Unit | 339.3 | Unit | 5 | 150 | 27 | Positive |
| Unit | 339.3 | Unit | 5 | 150 | 27 | Positive |
| Unit | 367.3 | Unit | 5 | 150 | 27 | Positive |
| Unit | 339.3 | Unit | 5 | 150 | 27 | Positive |
| Unit | 363.3 | Unit | 5 | 150 | 27 | Positive |
| Unit | 361.3 | Unit | 5 | 150 | 27 | Positive |
| Unit | 339.3 | Unit | 5 | 150 | 27 | Positive |
| Unit | 339.3 | Unit | 5 | 150 | 27 | Positive |
| Unit | 339.3 | Unit | 5 | 150 | 27 | Positive |
| Unit | 339.3 | Unit | 5 | 150 | 27 | Positive |

| | | | | | | |
|---|---|---|---|---|---|---|
| Unit | 339.3 | Unit | 5 | 150 | 27 | Positive |
| Unit | 339.3 | Unit | 5 | 150 | 27 | Positive |
| Unit | 385.3 | Unit | 5 | 150 | 27 | Positive |
| Unit | 337.3 | Unit | 5 | 150 | 27 | Positive |
| Unit | 363.3 | Unit | 5 | 150 | 27 | Positive |
| Unit | 385.3 | Unit | 5 | 150 | 27 | Positive |
| Unit | 385.6 | Unit | 5 | 150 | 27 | Positive |
| Unit | 357.3 | Unit | 5 | 150 | 27 | Positive |
| Unit | 385.3 | Unit | 5 | 150 | 27 | Positive |
| Unit | 385.3 | Unit | 5 | 150 | 27 | Positive |
| Unit | 387.3 | Unit | 5 | 150 | 27 | Positive |
| Unit | 387.3 | Unit | 5 | 150 | 27 | Positive |
| Unit | 385.3 | Unit | 5 | 150 | 27 | Positive |
| Unit | 577.5 | Unit | 5 | 143 | 27 | Positive |
| Unit | 605.5 | Unit | 5 | 143 | 27 | Positive |
| Unit | 603.5 | Unit | 5 | 143 | 27 | Positive |
| Unit | 255.2 | Unit | 5 | 163 | 32 | Negative |
| Unit | 283.2 | Unit | 5 | 163 | 32 | Negative |
| Unit | 281.2 | Unit | 5 | 163 | 32 | Negative |
| Unit | 279.2 | Unit | 5 | 163 | 32 | Negative |
| Unit | 303.2 | Unit | 5 | 163 | 32 | Negative |
| Unit | 301.2 | Unit | 5 | 163 | 32 | Negative |
| Unit | 331.2 | Unit | 5 | 163 | 32 | Negative |
| Unit | 329.2 | Unit | 5 | 163 | 32 | Negative |
| Unit | 327.2 | Unit | 5 | 163 | 32 | Negative |
| Unit | 104.1 | Unit | 5 | 180 | 24 | Positive |
| Unit | 523.4 | Unit | 5 | 110 | 9 | Positive |
| Unit | 549.5 | Unit | 5 | 110 | 9 | Positive |
| Unit | 551.5 | Unit | 5 | 110 | 9 | Positive |
| Unit | 549.5 | Unit | 5 | 110 | 9 | Positive |
| Unit | 579.5 | Unit | 5 | 110 | 9 | Positive |
| Unit | 577.5 | Unit | 5 | 110 | 9 | Positive |
| Unit | 599.5 | Unit | 5 | 110 | 9 | Positive |
| Unit | 623.5 | Unit | 5 | 110 | 9 | Positive |
| Unit | 547.5 | Unit | 5 | 110 | 9 | Positive |
| Unit | 575.5 | Unit | 5 | 110 | 9 | Positive |
| Unit | 573.5 | Unit | 5 | 110 | 9 | Positive |
| Unit | 621.5 | Unit | 5 | 110 | 9 | Positive |

| | | | | | | |
|---|---|---|---|---|---|---|
| Unit | 607.5 | Unit | 5 | 110 | 9 | Positive |
| Unit | 605.5 | Unit | 5 | 110 | 9 | Positive |
| Unit | 603.5 | Unit | 5 | 110 | 9 | Positive |
| Unit | 601.5 | Unit | 5 | 110 | 9 | Positive |
| Unit | 633.5 | Unit | 5 | 110 | 9 | Positive |
| Unit | 631.5 | Unit | 5 | 110 | 9 | Positive |
| Unit | 629.6 | Unit | 5 | 110 | 9 | Positive |
| Unit | 627.5 | Unit | 5 | 110 | 9 | Positive |
| Unit | 625.5 | Unit | 5 | 110 | 9 | Positive |
| Unit | 659.5 | Unit | 5 | 110 | 9 | Positive |
| Unit | 657.5 | Unit | 5 | 110 | 9 | Positive |
| Unit | 655.5 | Unit | 5 | 110 | 9 | Positive |
| Unit | 653.5 | Unit | 5 | 110 | 9 | Positive |
| Unit | 651.5 | Unit | 5 | 110 | 9 | Positive |
| Unit | 649.5 | Unit | 5 | 110 | 9 | Positive |
| Unit | 647.5 | Unit | 5 | 110 | 9 | Positive |
| Unit | 643.5 | Unit | 5 | 110 | 9 | Positive |
| Unit | 671.5 | Unit | 5 | 110 | 9 | Positive |
| Unit | 669.5 | Unit | 5 | 110 | 9 | Positive |
| Unit | 699.5 | Unit | 5 | 110 | 9 | Positive |
| Unit | 697.5 | Unit | 5 | 110 | 9 | Positive |
| Unit | 695.5 | Unit | 5 | 110 | 9 | Positive |
| Unit | 86.1 | Unit | 5 | 215 | 50 | Positive |
| Unit | 124.9 | Unit | 5 | 215 | 50 | Positive |

| Compound group | Compour | Compour | Ion speci | CAS | z | Monoisot | ISTD | Precurso |
|---|---|---|---|---|---|---|---|---|
| PE(16:0_16:0) | PE(16:0_ | C42H82N | [M+H]+ | | 1 | | FALSE | 692.5 |
| PE(16:0_16:1) | PE(16:0_ | C42H82N | [M+H]+ | | 1 | | FALSE | 690.5 |
| PE(16:0_18:1) | PE(16:0_ | C42H82N | [M+H]+ | | 1 | | FALSE | 718.5 |
| PE(16:0_18:2) | PE(16:0_ | C42H82N | [M+H]+ | | 1 | | FALSE | 716.5 |
| PE(16:0_18:3) (a) | PE(16:0_ | C42H82N | [M+H]+ | | 1 | | FALSE | 714.5 |
| PE(16:0_20:3) | PE(16:0_ | C42H82N | [M+H]+ | | 1 | | FALSE | 742.5 |
| PE(16:0_20:4) | PE(16:0_ | C42H82N | [M+H]+ | | 1 | | FALSE | 740.5 |
| PE(16:0_20:5) | PE(16:0_ | C42H82N | [M+H]+ | | 1 | | FALSE | 738.5 |
| PE(16:0_22:6) | PE(16:0_ | C42H82N | [M+H]+ | | 1 | | FALSE | 764.5 |
| PE(16:1_18:2) | PE(16:1_ | C42H82N | [M+H]+ | | 1 | | FALSE | 714.5 |
| PE(16:1_20:4) | PE(16:1_ | C42H82N | [M+H]+ | | 1 | | FALSE | 738.5 |
| PE(17:0_18:1) | PE(17:0_ | C42H82N | [M+H]+ | | 1 | | FALSE | 732.6 |
| PE(17:0_18:2) | PE(17:0_ | C42H82N | [M+H]+ | | 1 | | FALSE | 730.5 |
| PE(17:0_20:4) | PE(17:0_ | C42H82N | [M+H]+ | | 1 | | FALSE | 754.6 |
| PE(18:0_18:1) | PE(18:0_ | C42H82N | [M+H]+ | | 1 | | FALSE | 746.6 |
| PE(18:0_18:2) | PE(18:0_ | C42H82N | [M+H]+ | | 1 | | FALSE | 744.6 |
| PE(18:0_20:3) (a) | PE(18:0_ | C42H82N | [M+H]+ | | 1 | | FALSE | 770.6 |
| PE(18:0_20:4) | PE(18:0_ | C42H82N | [M+H]+ | | 1 | | FALSE | 768.6 |
| PE(18:0_22:4) | PE(18:0_ | C42H82N | [M+H]+ | | 1 | | FALSE | 796.6 |
| PE(18:0_22:5) (n3) | PE(18:0_ | C42H82N | [M+H]+ | | 1 | | FALSE | 794.6 |
| PE(18:0_22:5) (n6) | PE(18:0_ | C42H82N | [M+H]+ | | 1 | | FALSE | 794.6 |
| PE(18:0_22:6) | PE(18:0_ | C42H82N | [M+H]+ | | 1 | | FALSE | 792.6 |
| PE(18:1_18:1) | PE(18:1_ | C42H82N | [M+H]+ | | 1 | | FALSE | 744.6 |
| PE(18:1_18:2) | PE(18:1_ | C42H82N | [M+H]+ | | 1 | | FALSE | 742.5 |
| PE(18:1_22:6) (a) | PE(18:1_ | C42H82N | [M+H]+ | | 1 | | FALSE | 790.5 |
| PE(20:0_20:4) | PE(20:0_ | C42H82N | [M+H]+ | | 1 | | FALSE | 796.6 |
| PE(36:0) | PE(36:0) | C42H82N | [M+H]+ | | 1 | | FALSE | 748.6 |
| PE(38:5) (a) | PE(38:5) | C42H82N | [M+H]+ | | 1 | | FALSE | 766.5 |
| PE(O-16:0/18:2) | PE(O-16: | C42H82N | [M+H]+ | | 1 | | FALSE | 702.5 |
| PE(O-16:0/20:3) | PE(O-16: | C42H82N | [M+H]+ | | 1 | | FALSE | 728.6 |
| PE(O-16:0/20:4) | PE(O-16: | C42H82N | [M+H]+ | | 1 | | FALSE | 726.5 |
| PE(O-16:0/22:4) | PE(O-16: | C42H82N | [M+H]+ | | 1 | | FALSE | 754.6 |
| PE(O-16:0/22:6) | PE(O-16: | C42H82N | [M+H]+ | | 1 | | FALSE | 750.6 |
| PE(O-18:0/20:4) | PE(O-18: | C42H82N | [M+H]+ | | 1 | | FALSE | 754.6 |
| PE(O-18:0/22:5) (a) | PE(O-18: | C42H82N | [M+H]+ | | 1 | | FALSE | 780.6 |
| PE(O-18:0/22:6) | PE(O-18: | C42H82N | [M+H]+ | | 1 | | FALSE | 778.5 |
| PE(O-18:1/18:2) | PE(O-18: | C42H82N | [M+H]+ | | 1 | | FALSE | 728.6 |

| | | | | | |
|---|---|---|---|---|---|
| PE(O-18:1/22:6) | PE(O-18: | C42H82I | [M+H]+ | 1 | FALSE | 776.6 |
| PE(O-34:1) | PE(O-34: | C42H82I | [M+H]+ | 1 | FALSE | 704.6 |
| PE(O-36:5) | PE(O-36: | C42H82I | [M+H]+ | 1 | FALSE | 724.5 |
| PE(O-38:5) (a) | PE(O-38: | C42H82I | [M+H]+ | 1 | FALSE | 752.6 |
| PE(O-38:5) (b) | PE(O-38: | C42H82I | [M+H]+ | 1 | FALSE | 752.6 |
| PE(P-15:0/20:4) (a) | PE(P-15: | C42H82I | [M+H]+ | 1 | FALSE | 710.5 |
| PE(P-15:0/22:6) (a) | PE(P-15: | C42H82I | [M+H]+ | 1 | FALSE | 734.5 |
| PE(P-16:0/18:1) | PE(P-16: | C42H82I | [M+H]+ | 1 | FALSE | 702.5 |
| PE(P-16:0/18:2) | PE(P-16: | C42H82I | [M+H]+ | 1 | FALSE | 700.5 |
| PE(P-16:0/18:3) | PE(P-16: | C42H82I | [M+H]+ | 1 | FALSE | 698.5 |
| PE(P-16:0/20:3) (a) | PE(P-16: | C42H82I | [M+H]+ | 1 | FALSE | 726.5 |
| PE(P-16:0/20:4) | PE(P-16: | C42H82I | [M+H]+ | 1 | FALSE | 724.5 |
| PE(P-16:0/20:5) | PE(P-16: | C42H82I | [M+H]+ | 1 | FALSE | 722.5 |
| PE(P-16:0/22:4) | PE(P-16: | C42H82I | [M+H]+ | 1 | FALSE | 752.6 |
| PE(P-16:0/22:5) (n3) | PE(P-16: | C42H82I | [M+H]+ | 1 | FALSE | 750.5 |
| PE(P-16:0/22:5) (n6) | PE(P-16: | C42H82I | [M+H]+ | 1 | FALSE | 750.5 |
| PE(P-16:0/22:6) | PE(P-16: | C42H82I | [M+H]+ | 1 | FALSE | 748.5 |
| PE(P-17:0/20:4) (a) | PE(P-17: | C42H82I | [M+H]+ | 1 | FALSE | 738.6 |
| PE(P-17:0/22:6) (a) | PE(P-17: | C42H82I | [M+H]+ | 1 | FALSE | 762.6 |
| PE(P-18:0/18:1) | PE(P-18: | C42H82I | [M+H]+ | 1 | FALSE | 730.6 |
| PE(P-18:0/18:2) | PE(P-18: | C42H82I | [M+H]+ | 1 | FALSE | 728.6 |
| PE(P-18:0/18:3) | PE(P-18: | C42H82I | [M+H]+ | 1 | FALSE | 726.5 |
| PE(P-18:0/20:3) (a) | PE(P-18: | C42H82I | [M+H]+ | 1 | FALSE | 754.5 |
| PE(P-18:0/20:4) | PE(P-18: | C42H82I | [M+H]+ | 1 | FALSE | 752.6 |
| PE(P-18:0/20:5) | PE(P-18: | C42H82I | [M+H]+ | 1 | FALSE | 750.5 |
| PE(P-18:0/22:4) | PE(P-18: | C42H82I | [M+H]+ | 1 | FALSE | 780.6 |
| PE(P-18:0/22:5) (n3) | PE(P-18: | C42H82I | [M+H]+ | 1 | FALSE | 778.5 |
| PE(P-18:0/22:5) (n6) | PE(P-18: | C42H82I | [M+H]+ | 1 | FALSE | 778.5 |
| PE(P-18:0/22:6) | PE(P-18: | C42H82I | [M+H]+ | 1 | FALSE | 776.6 |
| PE(P-18:1/18:1) (a) | PE(P-18: | C42H82I | [M+H]+ | 1 | FALSE | 728.6 |
| PE(P-18:1/18:2) (a) | PE(P-18: | C42H82I | [M+H]+ | 1 | FALSE | 726.5 |
| PE(P-18:1/18:3) | PE(P-18: | C42H82I | [M+H]+ | 1 | FALSE | 724.5 |
| PE(P-18:1/20:3) (a) | PE(P-18: | C42H82I | [M+H]+ | 1 | FALSE | 752.5 |
| PE(P-18:1/20:4) (a) | PE(P-18: | C42H82I | [M+H]+ | 1 | FALSE | 750.5 |
| PE(P-18:1/20:5) (a) | PE(P-18: | C42H82I | [M+H]+ | 1 | FALSE | 748.5 |
| PE(P-18:1/22:4) | PE(P-18: | C42H82I | [M+H]+ | 1 | FALSE | 778.5 |
| PE(P-18:1/22:5) (a) | PE(P-18: | C42H82I | [M+H]+ | 1 | FALSE | 776.6 |
| PE(P-18:1/22:6) (a) | PE(P-18: | C42H82I | [M+H]+ | 1 | FALSE | 774.5 |

| | | | | | | |
|---|---|---|---|---|---|---|
| PE(P-19:0/20:4) (a) | PE(P-19: | C42H82N | [M+H]+ | 1 | FALSE | 766.6 |
| PE(P-20:0/18:1) | PE(P-20: | C42H82N | [M+H]+ | 1 | FALSE | 758.6 |
| PE(P-20:0/18:2) | PE(P-20: | C42H82N | [M+H]+ | 1 | FALSE | 756.6 |
| PE(P-20:0/20:4) | PE(P-20: | C42H82N | [M+H]+ | 1 | FALSE | 780.6 |
| PE(P-20:0/22:6) | PE(P-20: | C42H82N | [M+H]+ | 1 | FALSE | 804.6 |
| PE(P-20:1/20:4) | PE(P-20: | C42H82N | [M+H]+ | 1 | FALSE | 778.5 |
| PE(P-20:1/22:6) (a) | PE(P-20: | C42H82N | [M+H]+ | 1 | FALSE | 802.6 |
| PE(18:1_20:4) | PE(18:1_ | C42H82N | [M+H]+ | 1 | FALSE | 766.5 |
| PE(19:1_19:1) | PE(19:1_ | C42H82N | [M+H]+ | 1 | FALSE | 772.6 |
| PG(34:1) | PG(34:1) | C42H82N | [M+NH4]+ | 1 | FALSE | 766.6 |
| PG(34:2) | PG(34:2) | C42H82N | [M+NH4]+ | 1 | FALSE | 792.6 |
| PG(36:1) | PG(36:1) | C42H82N | [M+NH4]+ | 1 | FALSE | 764.6 |
| PG(36:2) | PG(36:2) | C42H82N | [M+NH4]+ | 1 | FALSE | 794.6 |
| PG(16:0_17:0) | PG(16:0_ | C42H82N | [M+NH4]+ | 1 | FALSE | 737.5 |
| PI(15-MHDA_18:1)/PI(17:0_18:1) | PI(15-MH | C42H82N | [M+NH4]+ | 1 | FALSE | 868.6 |
| PI(15-MHDA_18:2)/PI(17:0_18:2) | PI(15-MH | C42H82N | [M+NH4]+ | 1 | FALSE | 866.6 |
| PI(15-MHDA_20:4)/PI(17:0_20:4) | PI(15-MH | C42H82N | [M+NH4]+ | 1 | FALSE | 890.6 |
| PI(16:0/16:0) | PI(16:0/1 | C42H82N | [M+NH4]+ | 1 | FALSE | 828.6 |
| PI(16:0_16:1) | PI(16:0_1 | C42H82N | [M+NH4]+ | 1 | FALSE | 826.5 |
| PI(16:0_20:3) (a) | PI(16:0_2 | C42H82N | [M+NH4]+ | 1 | FALSE | 878.6 |
| PI(16:0_20:4) | PI(16:0_2 | C42H82N | [M+NH4]+ | 1 | FALSE | 876.6 |
| PI(18:0_18:1) | PI(18:0_1 | C42H82N | [M+NH4]+ | 1 | FALSE | 882.6 |
| PI(18:0_20:2) | PI(18:0_2 | C42H82N | [M+NH4]+ | 1 | FALSE | 908.6 |
| PI(18:0_20:3) (a) | PI(18:0_2 | C42H82N | [M+NH4]+ | 1 | FALSE | 906.6 |
| PI(18:0_20:4) | PI(18:0_2 | C42H82N | [M+NH4]+ | 1 | FALSE | 904.6 |
| PI(18:0_22:4) | PI(18:0_2 | C42H82N | [M+NH4]+ | 1 | FALSE | 932.6 |
| PI(18:0_22:5) (n3) | PI(18:0_2 | C42H82N | [M+NH4]+ | 1 | FALSE | 930.6 |
| PI(18:0_22:5) (n6) | PI(18:0_2 | C42H82N | [M+NH4]+ | 1 | FALSE | 930.6 |
| PI(18:0_22:6) | PI(18:0_2 | C42H82N | [M+NH4]+ | 1 | FALSE | 928.6 |
| Sitosterolester(18:3) | Sitosterol | C29H49 | [M+H]+ | 1 | FALSE | 675.61 |

| MS1 res | Product r | MS2 res | Dwell (m | Fragmen | CE (V) | Polarity |
|---|---|---|---|---|---|---|
| Unit | 551.5 | Unit | 20 | 166 | 17 | Positive |
| Unit | 549.5 | Unit | 20 | 166 | 17 | Positive |
| Unit | 577.5 | Unit | 20 | 166 | 17 | Positive |
| Unit | 575.5 | Unit | 20 | 166 | 17 | Positive |
| Unit | 573.5 | Unit | 20 | 166 | 17 | Positive |
| Unit | 601.5 | Unit | 20 | 166 | 17 | Positive |
| Unit | 599.5 | Unit | 20 | 166 | 17 | Positive |
| Unit | 597.5 | Unit | 20 | 166 | 17 | Positive |
| Unit | 623.5 | Unit | 20 | 166 | 17 | Positive |
| Unit | 573.5 | Unit | 20 | 166 | 17 | Positive |
| Unit | 597.5 | Unit | 20 | 166 | 17 | Positive |
| Unit | 591.5 | Unit | 20 | 166 | 17 | Positive |
| Unit | 589.5 | Unit | 20 | 166 | 17 | Positive |
| Unit | 613.5 | Unit | 20 | 166 | 17 | Positive |
| Unit | 605.6 | Unit | 20 | 166 | 17 | Positive |
| Unit | 603.5 | Unit | 20 | 166 | 17 | Positive |
| Unit | 629.6 | Unit | 20 | 166 | 17 | Positive |
| Unit | 627.5 | Unit | 20 | 166 | 17 | Positive |
| Unit | 655.6 | Unit | 20 | 166 | 17 | Positive |
| Unit | 653.6 | Unit | 20 | 166 | 17 | Positive |
| Unit | 653.6 | Unit | 20 | 166 | 17 | Positive |
| Unit | 651.5 | Unit | 20 | 166 | 17 | Positive |
| Unit | 603.5 | Unit | 20 | 166 | 17 | Positive |
| Unit | 601.5 | Unit | 20 | 166 | 17 | Positive |
| Unit | 649.5 | Unit | 20 | 166 | 17 | Positive |
| Unit | 655.6 | Unit | 20 | 166 | 17 | Positive |
| Unit | 607.6 | Unit | 20 | 166 | 17 | Positive |
| Unit | 625.5 | Unit | 20 | 166 | 17 | Positive |
| Unit | 561.5 | Unit | 20 | 166 | 17 | Positive |
| Unit | 587.5 | Unit | 20 | 166 | 17 | Positive |
| Unit | 585.5 | Unit | 20 | 166 | 17 | Positive |
| Unit | 613.6 | Unit | 20 | 166 | 17 | Positive |
| Unit | 609.5 | Unit | 20 | 166 | 17 | Positive |
| Unit | 613.6 | Unit | 20 | 166 | 17 | Positive |
| Unit | 639.6 | Unit | 20 | 166 | 17 | Positive |
| Unit | 637.5 | Unit | 20 | 166 | 17 | Positive |
| Unit | 587.5 | Unit | 20 | 166 | 17 | Positive |

| | | | | | | |
|---|---|---|---|---|---|---|
| Unit | 635.5 | Unit | 20 | 166 | 17 | Positive |
| Unit | 563.5 | Unit | 20 | 166 | 17 | Positive |
| Unit | 583.5 | Unit | 20 | 166 | 17 | Positive |
| Unit | 611.5 | Unit | 20 | 166 | 17 | Positive |
| Unit | 611.5 | Unit | 20 | 166 | 17 | Positive |
| Unit | 361.3 | Unit | 20 | 166 | 17 | Positive |
| Unit | 385.3 | Unit | 20 | 166 | 17 | Positive |
| Unit | 339.3 | Unit | 20 | 166 | 17 | Positive |
| Unit | 337.3 | Unit | 20 | 166 | 17 | Positive |
| Unit | 335.3 | Unit | 20 | 166 | 17 | Positive |
| Unit | 363.3 | Unit | 20 | 166 | 17 | Positive |
| Unit | 361.3 | Unit | 20 | 166 | 17 | Positive |
| Unit | 359.3 | Unit | 20 | 166 | 17 | Positive |
| Unit | 389.3 | Unit | 20 | 166 | 17 | Positive |
| Unit | 387.3 | Unit | 20 | 166 | 17 | Positive |
| Unit | 387.3 | Unit | 20 | 166 | 17 | Positive |
| Unit | 385.3 | Unit | 20 | 166 | 17 | Positive |
| Unit | 361.3 | Unit | 20 | 166 | 17 | Positive |
| Unit | 385.3 | Unit | 20 | 166 | 17 | Positive |
| Unit | 339.3 | Unit | 20 | 166 | 17 | Positive |
| Unit | 337.3 | Unit | 20 | 166 | 17 | Positive |
| Unit | 335.3 | Unit | 20 | 166 | 17 | Positive |
| Unit | 363.3 | Unit | 20 | 166 | 17 | Positive |
| Unit | 361.3 | Unit | 20 | 166 | 17 | Positive |
| Unit | 359.3 | Unit | 20 | 166 | 17 | Positive |
| Unit | 389.3 | Unit | 20 | 166 | 17 | Positive |
| Unit | 387.3 | Unit | 20 | 166 | 17 | Positive |
| Unit | 387.3 | Unit | 20 | 166 | 17 | Positive |
| Unit | 385.3 | Unit | 20 | 166 | 17 | Positive |
| Unit | 339.3 | Unit | 20 | 166 | 17 | Positive |
| Unit | 337.3 | Unit | 20 | 166 | 17 | Positive |
| Unit | 335.3 | Unit | 20 | 166 | 17 | Positive |
| Unit | 363.3 | Unit | 20 | 166 | 17 | Positive |
| Unit | 361.3 | Unit | 20 | 166 | 17 | Positive |
| Unit | 359.3 | Unit | 20 | 166 | 17 | Positive |
| Unit | 389.3 | Unit | 20 | 166 | 17 | Positive |
| Unit | 387.3 | Unit | 20 | 166 | 17 | Positive |
| Unit | 385.3 | Unit | 20 | 166 | 17 | Positive |

| | | | | | | |
|---|---|---|---|---|---|---|
| Unit | 361.3 | Unit | 20 | 166 | 17 | Positive |
| Unit | 339.3 | Unit | 20 | 166 | 17 | Positive |
| Unit | 337.3 | Unit | 20 | 166 | 17 | Positive |
| Unit | 361.3 | Unit | 20 | 166 | 17 | Positive |
| Unit | 385.3 | Unit | 20 | 166 | 17 | Positive |
| Unit | 361.3 | Unit | 20 | 166 | 17 | Positive |
| Unit | 385.3 | Unit | 20 | 166 | 17 | Positive |
| Unit | 577.5 | Unit | 20 | 166 | 17 | Positive |
| Unit | 591.5 | Unit | 20 | 166 | 17 | Positive |
| Unit | 577.5 | Unit | 20 | 166 | 17 | Positive |
| Unit | 603.5 | Unit | 20 | 166 | 21 | Positive |
| Unit | 575.5 | Unit | 20 | 166 | 21 | Positive |
| Unit | 605.6 | Unit | 20 | 166 | 21 | Positive |
| Unit | 565.5 | Unit | 20 | 166 | 21 | Positive |
| Unit | 591.6 | Unit | 20 | 166 | 17 | Positive |
| Unit | 589.6 | Unit | 20 | 166 | 17 | Positive |
| Unit | 613.6 | Unit | 20 | 166 | 17 | Positive |
| Unit | 551.6 | Unit | 20 | 166 | 17 | Positive |
| Unit | 549.5 | Unit | 20 | 166 | 17 | Positive |
| Unit | 601.6 | Unit | 20 | 166 | 17 | Positive |
| Unit | 599.6 | Unit | 20 | 166 | 17 | Positive |
| Unit | 605.6 | Unit | 20 | 166 | 17 | Positive |
| Unit | 631.6 | Unit | 20 | 166 | 17 | Positive |
| Unit | 629.6 | Unit | 20 | 166 | 17 | Positive |
| Unit | 627.6 | Unit | 20 | 166 | 17 | Positive |
| Unit | 655.6 | Unit | 20 | 166 | 17 | Positive |
| Unit | 653.6 | Unit | 20 | 166 | 17 | Positive |
| Unit | 653.6 | Unit | 20 | 166 | 17 | Positive |
| Unit | 651.6 | Unit | 20 | 166 | 17 | Positive |
| Unit | 397.4 | Unit | 20 | 166 | 23 | Positive |

| Compound group | Compound name | Compour | Ion speci | CAS | z | Monoisot | ISTD | Precurso | MS1 res |
|---|---|---|---|---|---|---|---|---|---|
| Ubiquinone | Ubiquinone | C42H82I | [M+NH4]+ | | 1 | | FALSE | 880.7 | Unit |
| TG(6:0_12:0_18:3) | TG(6:0_11:2_18:3) | C42H82I | [M+NH4]+ | | 1 | | FALSE | 650.5 | Unit |
| TG(6:0_12:0_18:3) | TG(6:0_11:2_18:3) | C42H82I | [M+NH4]+ | | 1 | | FALSE | 650.5 | Unit |
| TG(6:0_12:0_18:3) | TG(6:0_11:2_18:3) | C42H82I | [M+NH4]+ | | 1 | | FALSE | 650.5 | Unit |
| TG(22:1_10:0_10:0) | TG(22:1_10:0_10:( | C42H82I | [M+NH4]+ | | 1 | | FALSE | 738.66 | Unit |
| TG(22:1_10:0_10:0) | TG(22:1_10:0_10:( | C42H82I | [M+NH4]+ | | 1 | | FALSE | 738.66 | Unit |
| TG(22:1_10:0_10:0) | TG(22:1_10:0_10:( | C42H82I | [M+NH4]+ | | 1 | | FALSE | 738.66 | Unit |
| TG(18:4_18:2_18:2) | TG(18:4_18:2_18:2 | C42H82I | [M+NH4]+ | | 1 | | FALSE | 892.7 | Unit |
| TG(18:4_18:2_18:2) | TG(18:4_18:2_18:2 | C42H82I | [M+NH4]+ | | 1 | | FALSE | 892.7 | Unit |
| TG(18:4_18:2_18:2) | TG(18:4_18:2_18:2 | C42H82I | [M+NH4]+ | | 1 | | FALSE | 892.7 | Unit |
| TG(18:4_16:0_18:2) | TG(18:4_16:0_18:2 | C42H82I | [M+NH4]+ | | 1 | | FALSE | 868.7 | Unit |
| TG(18:4_16:0_18:2) | TG(18:4_16:0_18:2 | C42H82I | [M+NH4]+ | | 1 | | FALSE | 868.7 | Unit |
| TG(18:4_16:0_18:2) | TG(18:4_16:0_18:2 | C42H82I | [M+NH4]+ | | 1 | | FALSE | 868.7 | Unit |
| TG(18:3_18:2_18:2) | TG(18:3_18:2_18:2 | C42H82I | [M+NH4]+ | | 1 | | FALSE | 894.8 | Unit |
| TG(18:3_18:2_18:2) | TG(18:3_18:2_18:2 | C42H82I | [M+NH4]+ | | 1 | | FALSE | 894.8 | Unit |
| TG(18:3_18:2_18:2) | TG(18:3_18:2_18:2 | C42H82I | [M+NH4]+ | | 1 | | FALSE | 894.8 | Unit |
| TG(18:3_18:2_16:0) | TG(18:3_18:2_16:( | C42H82I | [M+NH4]+ | | 1 | | FALSE | 870.8 | Unit |
| TG(18:3_18:2_16:0) | TG(18:3_18:2_16:( | C42H82I | [M+NH4]+ | | 2 | | FALSE | 870.8 | Unit |
| TG(18:3_18:2_16:0) | TG(18:3_18:2_16:( | C42H82I | [M+NH4]+ | | 3 | | FALSE | 870.8 | Unit |
| TG(18:2_18:2_20:4) | TG(18:2_18:2_20:4 | C42H82I | [M+NH4]+ | | 1 | | FALSE | 920.8 | Unit |
| TG(18:2_18:2_20:4) | TG(18:2_18:2_20:4 | C42H82I | [M+NH4]+ | | 2 | | FALSE | 920.8 | Unit |
| TG(18:2_18:2_20:4) | TG(18:2_18:2_20:4 | C42H82I | [M+NH4]+ | | 3 | | FALSE | 920.8 | Unit |
| TG(18:2_18:2_18:2) | TG(18:2_18:2_18:2 | C42H82I | [M+NH4]+ | | 1 | | FALSE | 896.8 | Unit |
| TG(18:2_18:2_16:0) | TG(18:2_18:2_16:( | C42H82I | [M+NH4]+ | | 1 | | FALSE | 872.8 | Unit |
| TG(18:2_18:2_16:0) | TG(18:2_18:2_16:( | C42H82I | [M+NH4]+ | | 1 | | FALSE | 872.8 | Unit |
| TG(18:2_18:2_16:0) | TG(18:2_18:2_16:( | C42H82I | [M+NH4]+ | | 1 | | FALSE | 872.8 | Unit |
| TG(18:2_18:2_15:0) | TG(18:2_18:2_15:( | C42H82I | [M+NH4]+ | | 1 | | FALSE | 858.8 | Unit |
| TG(18:2_18:2_15:0) | TG(18:2_18:2_15:( | C42H82I | [M+NH4]+ | | 1 | | FALSE | 858.8 | Unit |
| TG(18:1e_16:0_16:0) | TG(18:1e_16:0_16 | C42H82I | [M+NH4]+ | | 1 | | FALSE | 850.8 | Unit |
| TG(18:1e_16:0_16:0) | TG(18:1e_16:0_16 | C42H82I | [M+NH4]+ | | 1 | | FALSE | 850.8 | Unit |
| TG(18:1_18:2_20:3) | TG(18:1_18:2_20:3 | C42H82I | [M+NH4]+ | | 1 | | FALSE | 924.8 | Unit |
| TG(18:1_18:2_20:3) | TG(18:1_18:2_20:3 | C42H82I | [M+NH4]+ | | 1 | | FALSE | 924.8 | Unit |
| TG(18:1_18:2_20:3) | TG(18:1_18:2_20:3 | C42H82I | [M+NH4]+ | | 1 | | FALSE | 924.8 | Unit |
| TG(18:1_18:2_18:2) | TG(18:1_18:2_18:2 | C42H82I | [M+NH4]+ | | 1 | | FALSE | 898.8 | Unit |
| TG(18:1_18:2_18:2) | TG(18:1_18:2_18:2 | C42H82I | [M+NH4]+ | | 1 | | FALSE | 898.8 | Unit |
| TG(18:1_18:2_18:2) | TG(18:1_18:2_18:2 | C42H82I | [M+NH4]+ | | 1 | | FALSE | 898.8 | Unit |
| TG(18:1_18:1_22:6) | TG(18:1_18:1_22:6 | C42H82I | [M+NH4]+ | | 1 | | FALSE | 948.8 | Unit |

| | | | | | | | |
|---|---|---|---|---|---|---|---|
| TG(18:1_18:1_22:6) | TG(18:1_18:1_22: | C42H82N | [M+NH4]+ | 1 | FALSE | 948.8 | Unit |
| TG(18:1_18:1_22:6) | TG(18:1_18:1_22: | C42H82N | [M+NH4]+ | 1 | FALSE | 948.8 | Unit |
| TG(18:1_18:1_18:3) | TG(18:1_18:1_18: | C42H82N | [M+NH4]+ | 1 | FALSE | 898.8 | Unit |
| TG(18:1_18:1_18:3) | TG(18:1_18:1_18: | C42H82N | [M+NH4]+ | 1 | FALSE | 898.8 | Unit |
| TG(18:1_18:1_18:3) | TG(18:1_18:1_18: | C42H82N | [M+NH4]+ | 1 | FALSE | 898.8 | Unit |
| TG(18:1_18:1_18:1) | TG(18:1_18:1_18: | C42H82N | [M+NH4]+ | 1 | FALSE | 902.8 | Unit |
| TG(18:1_18:1_18:1) | TG(18:1_18:1_18: | C42H82N | [M+NH4]+ | 1 | FALSE | 902.8 | Unit |
| TG(18:1_18:1_18:1) | TG(18:1_18:1_18: | C42H82N | [M+NH4]+ | 1 | FALSE | 902.8 | Unit |
| TG(18:0_18:1_22:6) | TG(18:0_18:1_22: | C42H82N | [M+NH4]+ | 1 | FALSE | 950.8 | Unit |
| TG(18:0_18:1_22:6) | TG(18:0_18:1_22: | C42H82N | [M+NH4]+ | 1 | FALSE | 950.8 | Unit |
| TG(18:0_18:1_22:6) | TG(18:0_18:1_22: | C42H82N | [M+NH4]+ | 1 | FALSE | 950.8 | Unit |
| TG(18:0_18:1_18:3) | TG(18:0_18:1_18: | C42H82N | [M+NH4]+ | 1 | FALSE | 900.8 | Unit |
| TG(18:0_18:1_18:3) | TG(18:0_18:1_18: | C42H82N | [M+NH4]+ | 1 | FALSE | 900.8 | Unit |
| TG(18:0_18:1_18:3) | TG(18:0_18:1_18: | C42H82N | [M+NH4]+ | 1 | FALSE | 900.8 | Unit |
| TG(18:0_16:0_18:3) | TG(18:0_16:0_18: | C42H82N | [M+NH4]+ | 1 | FALSE | 874.8 | Unit |
| TG(18:0_16:0_18:3) | TG(18:0_16:0_18: | C42H82N | [M+NH4]+ | 1 | FALSE | 874.8 | Unit |
| TG(18:0_16:0_18:3) | TG(18:0_16:0_18: | C42H82N | [M+NH4]+ | 1 | FALSE | 874.8 | Unit |
| TG(18:0_16:0_16:0) | TG(18:0_16:0_16: | C42H82N | [M+NH4]+ | 1 | FALSE | 852.8 | Unit |
| TG(18:0_16:0_16:0) | TG(18:0_16:0_16: | C42H82N | [M+NH4]+ | 1 | FALSE | 852.8 | Unit |
| TG(18:0_16:0_16:0) | TG(18:0_16:0_16: | C42H82N | [M+NH4]+ | 1 | FALSE | 852.8 | Unit |
| TG(16:1_18:2_18:3) | TG(16:1_18:2_18: | C42H82N | [M+NH4]+ | 1 | FALSE | 868.7 | Unit |
| TG(16:1_18:2_18:3) | TG(16:1_18:2_18: | C42H82N | [M+NH4]+ | 1 | FALSE | 868.7 | Unit |
| TG(16:1_18:2_18:3) | TG(16:1_18:2_18: | C42H82N | [M+NH4]+ | 1 | FALSE | 868.7 | Unit |
| TG(16:1_18:2_15:0) | TG(16:1_18:2_15: | C42H82N | [M+NH4]+ | 1 | FALSE | 832.7 | Unit |
| TG(16:1_18:2_15:0) | TG(16:1_18:2_15: | C42H82N | [M+NH4]+ | 1 | FALSE | 832.7 | Unit |
| TG(16:1_18:2_15:0) | TG(16:1_18:2_15: | C42H82N | [M+NH4]+ | 1 | FALSE | 832.7 | Unit |
| TG(16:1_14:1_15:0) | TG(16:1_14:1_15: | C42H82N | [M+NH4]+ | 1 | FALSE | 778.7 | Unit |
| TG(16:1_14:1_15:0) | TG(16:1_14:1_15: | C42H82N | [M+NH4]+ | 1 | FALSE | 778.7 | Unit |
| TG(16:1_14:1_15:0) | TG(16:1_14:1_15: | C42H82N | [M+NH4]+ | 1 | FALSE | 778.7 | Unit |
| TG(16:0_18:2_22:4) | TG(16:0_18:2_22: | C42H82N | [M+NH4]+ | 1 | FALSE | 924.8 | Unit |
| TG(16:0_18:2_22:4) | TG(16:0_18:2_22: | C42H82N | [M+NH4]+ | 1 | FALSE | 924.8 | Unit |
| TG(16:0_18:2_22:4) | TG(16:0_18:2_22: | C42H82N | [M+NH4]+ | 1 | FALSE | 924.8 | Unit |
| TG(16:0_18:1_22:6) | TG(16:0_18:1_22: | C42H82N | [M+NH4]+ | 1 | FALSE | 922.8 | Unit |
| TG(16:0_18:1_22:6) | TG(16:0_18:1_22: | C42H82N | [M+NH4]+ | 1 | FALSE | 922.8 | Unit |
| TG(16:0_18:1_22:6) | TG(16:0_18:1_22: | C42H82N | [M+NH4]+ | 1 | FALSE | 922.8 | Unit |
| TG(16:0_18:1_18:3) | TG(16:0_18:1_18: | C42H82N | [M+NH4]+ | 1 | FALSE | 872.8 | Unit |
| TG(16:0_18:1_18:3) | TG(16:0_18:1_18: | C42H82N | [M+NH4]+ | 2 | FALSE | 872.8 | Unit |
| TG(16:0_18:1_18:3) | TG(16:0_18:1_18: | C42H82N | [M+NH4]+ | 3 | FALSE | 872.8 | Unit |

| | | | | | | | |
|---|---|---|---|---|---|---|---|
| TG(15:0_18:2_18:3) | TG(15:0_18:2_18:3 | C42H82N | [M+NH4]+ | 1 | FALSE | 856.7 | Unit |
| TG(15:0_18:2_18:3) | TG(15:0_18:2_18:3 | C42H82N | [M+NH4]+ | 1 | FALSE | 856.7 | Unit |
| TG(15:0_18:2_18:3) | TG(15:0_18:2_18:3 | C42H82N | [M+NH4]+ | 1 | FALSE | 856.7 | Unit |
| TG(14:0_14:1_13:0) | TG(14:0_14:1_13:( | C42H82N | [M+NH4]+ | 1 | FALSE | 724.6 | Unit |
| TG(14:0_14:1_13:0) | TG(14:0_14:1_13:( | C42H82N | [M+NH4]+ | 1 | FALSE | 724.6 | Unit |
| TG(14:0_14:1_13:0) | TG(14:0_14:1_13:( | C42H82N | [M+NH4]+ | 1 | FALSE | 724.6 | Unit |
| TG(12:1e_6:0_18:3) | TG(12:1e_6:0_18:3 | C42H82N | [M+NH4]+ | 1 | FALSE | 634.5 | Unit |
| TG(12:0_14:0_16:0) | TG(12:0_14:0_16:( | C42H82N | [M+NH4]+ | 1 | FALSE | 740.7 | Unit |
| TG(12:0_14:0_16:0) | TG(12:0_14:0_16:( | C42H82N | [M+NH4]+ | 1 | FALSE | 740.7 | Unit |
| TG(12:0_14:0_16:0) | TG(12:0_14:0_16:( | C42H82N | [M+NH4]+ | 1 | FALSE | 740.7 | Unit |
| TG(12:0_12:0_14:0) | TG(12:0_12:0_14:( | C42H82N | [M+NH4]+ | 1 | FALSE | 684.6 | Unit |
| TG(12:0_12:0_14:0) | TG(12:0_12:0_14:( | C42H82N | [M+NH4]+ | 1 | FALSE | 684.6 | Unit |

| Product r | MS2 res | Dwell (m | Fragmen | CE (V) | Polarity |
|---|---|---|---|---|---|
| 197.0 | Unit | 20 | 166 | 17 | Positive |
| 517.4 | Unit | 20 | 166 | 21 | Positive |
| 433.3 | Unit | 20 | 166 | 21 | Positive |
| 355.3 | Unit | 20 | 166 | 21 | Positive |
| 549.5 | Unit | 20 | 166 | 21 | Positive |
| 549.5 | Unit | 20 | 166 | 21 | Positive |
| 383.3 | Unit | 20 | 166 | 21 | Positive |
| 599.5 | Unit | 20 | 166 | 21 | Positive |
| 595.5 | Unit | 20 | 166 | 21 | Positive |
| 595.5 | Unit | 20 | 166 | 21 | Positive |
| 595.5 | Unit | 20 | 166 | 21 | Positive |
| 575.5 | Unit | 20 | 166 | 21 | Positive |
| 571.5 | Unit | 20 | 166 | 21 | Positive |
| 599.5 | Unit | 20 | 166 | 21 | Positive |
| 597.5 | Unit | 20 | 166 | 21 | Positive |
| 597.5 | Unit | 20 | 166 | 21 | Positive |
| 597.5 | Unit | 20 | 166 | 21 | Positive |
| 575.5 | Unit | 20 | 166 | 21 | Positive |
| 573.5 | Unit | 20 | 166 | 21 | Positive |
| 623.5 | Unit | 20 | 166 | 21 | Positive |
| 623.5 | Unit | 20 | 166 | 21 | Positive |
| 599.5 | Unit | 20 | 166 | 21 | Positive |
| 599.5 | Unit | 20 | 166 | 21 | Positive |
| 599.5 | Unit | 20 | 166 | 21 | Positive |
| 575.5 | Unit | 20 | 166 | 21 | Positive |
| 575.5 | Unit | 20 | 166 | 21 | Positive |
| 599.5 | Unit | 20 | 166 | 21 | Positive |
| 561.5 | Unit | 20 | 166 | 21 | Positive |
| 551.5 | Unit | 20 | 166 | 21 | Positive |
| 577.5 | Unit | 20 | 166 | 21 | Positive |
| 625.5 | Unit | 20 | 166 | 21 | Positive |
| 627.5 | Unit | 20 | 166 | 21 | Positive |
| 601.5 | Unit | 20 | 166 | 21 | Positive |
| 599.5 | Unit | 20 | 166 | 21 | Positive |
| 601.5 | Unit | 20 | 166 | 21 | Positive |
| 601.5 | Unit | 20 | 166 | 21 | Positive |
| 649.5 | Unit | 20 | 166 | 21 | Positive |

| | | | | | |
|---|---|---|---|---|---|
| 649.5 | Unit | 20 | 166 | 21 | Positive |
| 603.5 | Unit | 20 | 166 | 21 | Positive |
| 603.5 | Unit | 20 | 166 | 21 | Positive |
| 599.5 | Unit | 20 | 166 | 21 | Positive |
| 599.5 | Unit | 20 | 166 | 21 | Positive |
| 603.5 | Unit | 20 | 166 | 21 | Positive |
| 603.5 | Unit | 20 | 166 | 21 | Positive |
| 603.5 | Unit | 20 | 166 | 21 | Positive |
| 651.5 | Unit | 20 | 166 | 21 | Positive |
| 649.5 | Unit | 20 | 166 | 21 | Positive |
| 605.6 | Unit | 20 | 166 | 21 | Positive |
| 605.6 | Unit | 20 | 166 | 21 | Positive |
| 601.5 | Unit | 20 | 166 | 21 | Positive |
| 599.5 | Unit | 20 | 166 | 21 | Positive |
| 601.5 | Unit | 20 | 166 | 21 | Positive |
| 579.5 | Unit | 20 | 166 | 21 | Positive |
| 573.5 | Unit | 20 | 166 | 21 | Positive |
| 551.5 | Unit | 20 | 166 | 21 | Positive |
| 579.5 | Unit | 20 | 166 | 21 | Positive |
| 579.5 | Unit | 20 | 166 | 21 | Positive |
| 597.5 | Unit | 20 | 166 | 21 | Positive |
| 573.5 | Unit | 20 | 166 | 21 | Positive |
| 571.5 | Unit | 20 | 166 | 21 | Positive |
| 573.5 | Unit | 20 | 166 | 21 | Positive |
| 561.5 | Unit | 20 | 166 | 21 | Positive |
| 535.5 | Unit | 20 | 166 | 21 | Positive |
| 507.4 | Unit | 20 | 166 | 21 | Positive |
| 519.4 | Unit | 20 | 166 | 21 | Positive |
| 535.5 | Unit | 20 | 166 | 21 | Positive |
| 651.5 | Unit | 20 | 166 | 21 | Positive |
| 627.5 | Unit | 20 | 166 | 21 | Positive |
| 575.5 | Unit | 20 | 166 | 21 | Positive |
| 649.5 | Unit | 20 | 166 | 21 | Positive |
| 623.5 | Unit | 20 | 166 | 21 | Positive |
| 577.5 | Unit | 20 | 166 | 21 | Positive |
| 599.5 | Unit | 20 | 166 | 21 | Positive |
| 577.5 | Unit | 20 | 166 | 21 | Positive |
| 573.5 | Unit | 20 | 166 | 21 | Positive |

| | | | | | |
|---|---|---|---|---|---|
| 597.5 | Unit | 20 | 166 | 21 | Positive |
| 559.5 | Unit | 20 | 166 | 21 | Positive |
| 561.5 | Unit | 20 | 166 | 21 | Positive |
| 479.4 | Unit | 20 | 166 | 21 | Positive |
| 481.4 | Unit | 20 | 166 | 21 | Positive |
| 493.4 | Unit | 20 | 166 | 21 | Positive |
| 335.3 | Unit | 20 | 166 | 21 | Positive |
| 523.5 | Unit | 20 | 166 | 21 | Positive |
| 495.4 | Unit | 20 | 166 | 21 | Positive |
| 467.4 | Unit | 20 | 166 | 21 | Positive |
| 467.4 | Unit | 20 | 166 | 21 | Positive |
| 439.4 | Unit | 20 | 166 | 21 | Positive |

| Compound group | Compound name | Compound fc | Ion speci | CAS | z | Monoisot | ISTD |
|---|---|---|---|---|---|---|---|
| PI(18:1_18:2) | PI(18:1_18:2) | C42H82NO1 | [M+NH4]+ | | 1 | | FALSE |
| PI(20:0_20:4) | PI(20:0_20:4) | C42H82NO1 | [M+NH4]+ | | 1 | | FALSE |
| PI(34:0) | PI(34:0) | C42H82NO1 | [M+NH4]+ | | 1 | | FALSE |
| PI(34:1) | PI(34:1) | C42H82NO1 | [M+NH4]+ | | 1 | | FALSE |
| PI(36:2) | PI(36:2) | C42H82NO1 | [M+NH4]+ | | 1 | | FALSE |
| PI(37:6) | PI(37:6) | C42H82NO1 | [M+NH4]+ | | 1 | | FALSE |
| PI(38:5) (a) | PI(38:5) (a) | C42H82NO1 | [M+NH4]+ | | 1 | | FALSE |
| PI(38:6) | PI(38:6) | C42H82NO1 | [M+NH4]+ | | 1 | | FALSE |
| PI(39:6) | PI(39:6) | C42H82NO1 | [M+NH4]+ | | 1 | | FALSE |
| PS(36:1) | PS(36:1) | C42H82NO1 | [M+H]+ | | 1 | | FALSE |
| PS(36:2) | PS(36:2) | C42H82NO1 | [M+H]+ | | 1 | | FALSE |
| PS(38:3) | PS(38:3) | C42H82NO1 | [M+H]+ | | 1 | | FALSE |
| PS(38:4) | PS(38:4) | C42H82NO1 | [M+H]+ | | 1 | | FALSE |
| PS(38:5) | PS(38:5) | C42H82NO1 | [M+H]+ | | 1 | | FALSE |
| PS(40:5) | PS(40:5) | C42H82NO1 | [M+H]+ | | 1 | | FALSE |
| PS(40:6) | PS(40:6) | C42H82NO1 | [M+H]+ | | 1 | | FALSE |
| S1P(d16:1) | S1P(d16:1) | | [M+H]+ | | 1 | | FALSE |
| S1P(d18:0) | S1P(d18:0) | | [M+H]+ | | 1 | | FALSE |
| S1P(d18:1) | S1P(d18:1) | | [M+H]+ | | 1 | | FALSE |
| S1P(d18:2) | S1P(d18:2) | | [M+H]+ | | 1 | | FALSE |
| SM(34:3) | SM(34:3) | | [M+H]+ | | 1 | | FALSE |
| SM(38:3) (a) | SM(38:3) (a) | | [M+H]+ | | 1 | | FALSE |
| SM(40:3) (a) | SM(40:3) (a) | | [M+H]+ | | 1 | | FALSE |
| SM(41:0) | SM(41:0) | | [M+H]+ | | 1 | | FALSE |
| SM(41:1) (a) | SM(41:1) (a) | | [M+H]+ | | 1 | | FALSE |
| SM(43:1) | SM(43:1) | | [M+H]+ | | 1 | | FALSE |
| SM(43:2) (c) | SM(43:2) (c) | | [M+H]+ | | 1 | | FALSE |
| SM(44:1) | SM(44:1) | | [M+H]+ | | 1 | | FALSE |
| SM(44:2) | SM(44:2) | | [M+H]+ | | 1 | | FALSE |
| SM(44:3) | SM(44:3) | | [M+H]+ | | 1 | | FALSE |
| SM(d16:1/23:0)/SM(d17:1/22 | SM(d16:1/23:0)/SM(d17:1/22:0) | | [M+H]+ | | 1 | | FALSE |
| SM(d16:1/24:1) | SM(d16:1/24:1) | | [M+H]+ | | 1 | | FALSE |
| SM(d18:0/14:0) | SM(d18:0/14:0) | | [M+H]+ | | 1 | | FALSE |
| SM(d18:0/16:0) | SM(d18:0/16:0) | | [M+H]+ | | 1 | | FALSE |
| SM(d18:0/22:0) | SM(d18:0/22:0) | | [M+H]+ | | 1 | | FALSE |
| SM(d18:1/14:0)/SM(d16:1/16 | SM(d18:1/14:0)/SM(d16:1/16:0) | | [M+H]+ | | 1 | | FALSE |
| SM(d18:1/16:0) | SM(d18:1/16:0) | | [M+H]+ | | 1 | | FALSE |

| | | | | | |
|---|---|---|---|---|---|
| SM(d18:1/18:0)/SM(d16:1/20 | SM(d18:1/18:0)/SM(d16:1/20:0) | | [M+H]+ | 1 | FALSE |
| SM(d18:1/20:0)/SM(d16:1/22 | SM(d18:1/20:0)/SM(d16:1/22:0) | | [M+H]+ | 1 | FALSE |
| SM(d18:1/22:0)/SM(d16:1/24 | SM(d18:1/22:0)/SM(d16:1/24:0) | | [M+H]+ | 1 | FALSE |
| SM(d18:1/23:0)/SM(d17:1/24 | SM(d18:1/23:0)/SM(d17:1/24:0) | | [M+H]+ | 1 | FALSE |
| SM(d18:1/24:0) | SM(d18:1/24:0) | | [M+H]+ | 1 | FALSE |
| SM(d18:1/24:1) | SM(d18:1/24:1) | | [M+H]+ | 1 | FALSE |
| SM(d18:2/14:0) | SM(d18:2/14:0) | | [M+H]+ | 1 | FALSE |
| SM(d18:2/16:0) | SM(d18:2/16:0) | | [M+H]+ | 1 | FALSE |
| SM(d18:2/17:0) | SM(d18:2/17:0) | | [M+H]+ | 1 | FALSE |
| SM(d18:2/18:0) | SM(d18:2/18:0) | | [M+H]+ | 1 | FALSE |
| SM(d18:2/18:1) | SM(d18:2/18:1) | | [M+H]+ | 1 | FALSE |
| SM(d18:2/20:0) | SM(d18:2/20:0) | | [M+H]+ | 1 | FALSE |
| SM(d18:2/22:0) | SM(d18:2/22:0) | | [M+H]+ | 1 | FALSE |
| SM(d18:2/23:0) | SM(d18:2/23:0) | C42H79O10I | [M+H]+ | 1 | FALSE |
| SM(d18:2/24:0) | SM(d18:2/24:0) | | [M+H]+ | 1 | FALSE |
| SM(d19:0_23:1) | SM(d19:0_23:1) | | [M+H]+ | 1 | FALSE |
| SM(d20:1_20:1) | SM(d20:1_20:1) | | [M+H]+ | 1 | FALSE |
| SM(d31:1) | SM(d31:1) | | [M+H]+ | 1 | FALSE |
| SM(d34:3) | SM(d34:3) | | [M+H]+ | 1 | FALSE |
| SM(d38:4) | SM(d38:4) | | [M+H]+ | 1 | FALSE |
| SM(d39:2) | SM(d39:2) | | [M+H]+ | 1 | FALSE |
| SM(d40:1) | SM(d40:1) | | [M+H]+ | 1 | FALSE |
| SM(d40:3) | SM(d40:3) | | [M+H]+ | 1 | FALSE |
| SM(d41:3) | SM(d41:3) | | [M+H]+ | 1 | FALSE |
| SM(d42:2) | SM(d42:2) | | [M+H]+ | 1 | FALSE |
| SM(d43:2) | SM(d43:2) | | [M+H]+ | 1 | FALSE |
| SM(d44:5) | SM(d44:5) | | [M+H]+ | 1 | FALSE |
| Sph(d16:1) | Sph(d16:1) | C42H82NO1 | [M+H]+ | 1 | FALSE |
| Sph(d18:1) | Sph(d18:1) | C42H79O10I | [M+H]+ | 1 | FALSE |
| Sph(d18:2) | Sph(d18:2) | | [M+H]+ | 1 | FALSE |
| Sulfatide (d18:1:/16:0(OH)) | Sulfatide (d18:1:/16:0(OH)) | | [M+H]+ | 1 | FALSE |
| Sulfatide (d18:1:/16:0) | Sulfatide (d18:1:/16:0) | | [M+H]+ | 1 | FALSE |
| Sulfatide (d18:1:/24:0(OH)) | Sulfatide (d18:1:/24:0(OH)) | | [M+H]+ | 1 | FALSE |
| Sulfatide (d18:1:/24:0) | Sulfatide (d18:1:/24:0) | | [M+H]+ | 1 | FALSE |
| Sulfatide (d18:1:/24:1(OH)) | Sulfatide (d18:1:/24:1(OH)) | | [M+H]+ | 1 | FALSE |
| Sulfatide (d18:1:/24:1) | Sulfatide (d18:1:/24:1) | | [M+H]+ | 1 | FALSE |
| TG(48:0) [NL-16:0] | TG(48:0) [NL-16:0] | C42H82NO1 | [M+NH4]+ | 1 | FALSE |
| TG(48:1) [NL-18:1] | TG(48:1) [NL-18:1] | C42H82NO1 | [M+NH4]+ | 1 | FALSE |

| | | | | |
|---|---|---|---|---|
| TG(48:2) [NL-14:1] | TG(48:2) [NL-14:1] | C42H82NO1 [M+NH4]+ | 1 | FALSE |
| TG(48:2) [NL-16:0] | TG(48:2) [NL-16:0] | C42H82NO1 [M+NH4]+ | 1 | FALSE |
| TG(48:2) [NL-16:1] | TG(48:2) [NL-16:1] | C42H82NO1 [M+NH4]+ | 1 | FALSE |
| TG(48:2) [NL-18:1] | TG(48:2) [NL-18:1] | C42H82NO1 [M+NH4]+ | 1 | FALSE |
| TG(48:3) [NL-16:1] | TG(48:3) [NL-16:1] | C42H82NO1 [M+NH4]+ | 1 | FALSE |
| TG(48:3) [NL-18:2] | TG(48:3) [NL-18:2] | C42H82NO1 [M+NH4]+ | 1 | FALSE |
| TG(49:1) [NL-15:0] | TG(49:1) [NL-15:0] | C42H82NO1 [M+NH4]+ | 1 | FALSE |
| TG(50:0) [NL-18:0] | TG(50:0) [NL-18:0] | C42H82NO1 [M+NH4]+ | 1 | FALSE |
| TG(50:1) [NL-14:0] | TG(50:1) [NL-14:0] | C42H82NO1 [M+NH4]+ | 1 | FALSE |
| TG(50:1) [NL-18:1] | TG(50:1) [NL-18:1] | C42H82NO1 [M+NH4]+ | 1 | FALSE |
| TG(50:2) [NL-18:0] | TG(50:2) [NL-18:0] | C42H82NO1 [M+NH4]+ | 1 | FALSE |
| TG(50:2) [NL-18:1] | TG(50:2) [NL-18:1] | C42H82NO1 [M+NH4]+ | 1 | FALSE |
| TG(50:2) [NL-18:2] | TG(50:2) [NL-18:2] | C42H82NO1 [M+NH4]+ | 1 | FALSE |
| TG(50:3) [NL-14:1] | TG(50:3) [NL-14:1] | C42H82NO1 [M+NH4]+ | 1 | FALSE |
| TG(50:3) [NL-16:1] | TG(50:3) [NL-16:1] | C42H82NO1 [M+NH4]+ | 1 | FALSE |
| TG(50:3) [NL-18:1] | TG(50:3) [NL-18:1] | C42H82NO1 [M+NH4]+ | 1 | FALSE |
| TG(50:4) [NL-14:0] | TG(50:4) [NL-14:0] | C42H82NO1 [M+NH4]+ | 1 | FALSE |
| TG(52:1) [NL-18:0] | TG(52:1) [NL-18:0] | C42H82NO1 [M+NH4]+ | 1 | FALSE |
| TG(52:2) [NL-16:0] | TG(52:2) [NL-16:0] | C42H82NO1 [M+NH4]+ | 1 | FALSE |
| TG(52:3) [NL-16:1] | TG(52:3) [NL-16:1] | C42H82NO1 [M+NH4]+ | 1 | FALSE |
| TG(52:3) [NL-18:2] | TG(52:3) [NL-18:2] | C42H82NO1 [M+NH4]+ | 1 | FALSE |
| TG(52:4) [NL-16:0] | TG(52:4) [NL-16:0] | C42H82NO1 [M+NH4]+ | 1 | FALSE |
| TG(52:4) [NL-18:1] | TG(52:4) [NL-18:1] | C42H82NO1 [M+NH4]+ | 1 | FALSE |
| TG(54:0) [NL-18:0] | TG(54:0) [NL-18:0] | C42H82NO1 [M+NH4]+ | 1 | FALSE |
| TG(54:1) [NL-18:1] | TG(54:1) [NL-18:1] | C42H82NO1 [M+NH4]+ | 1 | FALSE |
| TG(54:2) [NL-18:0] | TG(54:2) [NL-18:0] | C42H82NO1 [M+NH4]+ | 1 | FALSE |
| TG(54:3) [NL-18:1] | TG(54:3) [NL-18:1] | C42H82NO1 [M+NH4]+ | 1 | FALSE |
| TG(54:4) [NL-18:0] | TG(54:4) [NL-18:0] | C42H82NO1 [M+NH4]+ | 1 | FALSE |
| TG(54:4) [NL-18:2] | TG(54:4) [NL-18:2] | C42H82NO1 [M+NH4]+ | 1 | FALSE |
| TG(54:5) [NL-18:1] | TG(54:5) [NL-18:1] | C42H82NO1 [M+NH4]+ | 1 | FALSE |
| TG(54:6) [NL-18:2] | TG(54:6) [NL-18:2] | C42H82NO1 [M+NH4]+ | 1 | FALSE |
| TG(56:6) [NL-20:4] | TG(56:6) [NL-20:4] | C42H82NO1 [M+NH4]+ | 1 | FALSE |
| TG(56:8) [NL-20:4] | TG(56:8) [NL-20:4] | C42H82NO1 [M+NH4]+ | 1 | FALSE |
| TG(58:8) [NL-22:6] | TG(58:8) [NL-22:6] | C42H82NO1 [M+NH4]+ | 1 | FALSE |
| TG(O-50:1) [NL-16:0] | TG(O-50:1) [NL-16:0] | C42H82NO1 [M+NH4]+ | 1 | FALSE |
| TG(O-52:0) [NL-16:0] | TG(O-52:0) [NL-16:0] | C42H82NO1 [M+NH4]+ | 1 | FALSE |
| TG(O-52:2) [NL-16:0] | TG(O-52:2) [NL-16:0] | C42H82NO1 [M+NH4]+ | 1 | FALSE |

| Precurso | MS1 res | Product r | MS2 res | Dwell (m: | Fragmen | CE (V) | Polarity |
|---|---|---|---|---|---|---|---|
| 878.6 | Unit | 601.6 | Unit | 20 | 166 | 17 | Positive |
| 932.6 | Unit | 655.6 | Unit | 20 | 166 | 17 | Positive |
| 856.6 | Unit | 579.6 | Unit | 20 | 166 | 17 | Positive |
| 854.6 | Unit | 577.6 | Unit | 20 | 166 | 17 | Positive |
| 880.6 | Unit | 603.6 | Unit | 20 | 166 | 17 | Positive |
| 886.6 | Unit | 609.6 | Unit | 20 | 166 | 17 | Positive |
| 902.6 | Unit | 625.6 | Unit | 20 | 166 | 17 | Positive |
| 900.6 | Unit | 623.6 | Unit | 20 | 166 | 17 | Positive |
| 914.6 | Unit | 637.6 | Unit | 20 | 166 | 17 | Positive |
| 790.6 | Unit | 605.6 | Unit | 20 | 166 | 25 | Positive |
| 788.5 | Unit | 603.5 | Unit | 20 | 166 | 25 | Positive |
| 814.6 | Unit | 629.6 | Unit | 20 | 166 | 25 | Positive |
| 812.5 | Unit | 627.5 | Unit | 20 | 166 | 25 | Positive |
| 810.5 | Unit | 625.5 | Unit | 20 | 166 | 25 | Positive |
| 838.6 | Unit | 653.6 | Unit | 20 | 166 | 25 | Positive |
| 836.5 | Unit | 651.5 | Unit | 20 | 166 | 25 | Positive |
| 352.2 | Unit | 236.3 | Unit | 20 | 166 | 16 | Positive |
| 382.2 | Unit | 284.3 | Unit | 20 | 166 | 11 | Positive |
| 380.2 | Unit | 264.3 | Unit | 20 | 166 | 16 | Positive |
| 378.2 | Unit | 262.3 | Unit | 20 | 166 | 16 | Positive |
| 699.5 | Unit | 184.1 | Unit | 20 | 166 | 25 | Positive |
| 755.6 | Unit | 184.1 | Unit | 20 | 166 | 25 | Positive |
| 783.6 | Unit | 184.1 | Unit | 20 | 166 | 25 | Positive |
| 803.7 | Unit | 184.1 | Unit | 20 | 166 | 25 | Positive |
| 801.7 | Unit | 184.1 | Unit | 20 | 166 | 25 | Positive |
| 829.7 | Unit | 184.1 | Unit | 20 | 166 | 25 | Positive |
| 827.7 | Unit | 184.1 | Unit | 20 | 166 | 25 | Positive |
| 843.6 | Unit | 184.1 | Unit | 20 | 166 | 25 | Positive |
| 841.6 | Unit | 184.1 | Unit | 20 | 166 | 25 | Positive |
| 839.6 | Unit | 184.1 | Unit | 20 | 166 | 25 | Positive |
| 773.7 | Unit | 184.1 | Unit | 20 | 166 | 25 | Positive |
| 785.7 | Unit | 184.1 | Unit | 20 | 166 | 25 | Positive |
| 677.6 | Unit | 184.1 | Unit | 20 | 166 | 25 | Positive |
| 705.6 | Unit | 184.1 | Unit | 20 | 166 | 25 | Positive |
| 789.7 | Unit | 184.1 | Unit | 20 | 166 | 25 | Positive |
| 675.5 | Unit | 184.1 | Unit | 20 | 166 | 25 | Positive |
| 703.6 | Unit | 184.1 | Unit | 20 | 166 | 25 | Positive |

| | | | | | | | |
|---|---|---|---|---|---|---|---|
| 731.6 | Unit | 184.1 | Unit | 20 | 166 | 25 | Positive |
| 759.6 | Unit | 184.1 | Unit | 20 | 166 | 25 | Positive |
| 787.7 | Unit | 184.1 | Unit | 20 | 166 | 25 | Positive |
| 801.7 | Unit | 184.1 | Unit | 20 | 166 | 25 | Positive |
| 815.7 | Unit | 184.1 | Unit | 20 | 166 | 25 | Positive |
| 813.7 | Unit | 184.1 | Unit | 20 | 166 | 25 | Positive |
| 673.5 | Unit | 184.1 | Unit | 20 | 166 | 25 | Positive |
| 701.6 | Unit | 184.1 | Unit | 20 | 166 | 25 | Positive |
| 715.6 | Unit | 184.1 | Unit | 20 | 166 | 25 | Positive |
| 729.6 | Unit | 184.1 | Unit | 20 | 166 | 25 | Positive |
| 727.6 | Unit | 184.1 | Unit | 20 | 166 | 25 | Positive |
| 757.6 | Unit | 184.1 | Unit | 20 | 166 | 25 | Positive |
| 785.7 | Unit | 184.1 | Unit | 20 | 166 | 25 | Positive |
| 799.7 | Unit | 184.1 | Unit | 20 | 166 | 25 | Positive |
| 813.7 | Unit | 184.1 | Unit | 20 | 166 | 25 | Positive |
| 815.7 | Unit | 184.1 | Unit | 20 | 166 | 25 | Positive |
| 785.7 | Unit | 184.1 | Unit | 20 | 166 | 25 | Positive |
| 661.5 | Unit | 184.1 | Unit | 20 | 166 | 25 | Positive |
| 699.5 | Unit | 184.1 | Unit | 20 | 166 | 25 | Positive |
| 753.6 | Unit | 184.1 | Unit | 20 | 166 | 25 | Positive |
| 771.6 | Unit | 184.1 | Unit | 20 | 166 | 25 | Positive |
| 787.7 | Unit | 184.1 | Unit | 20 | 166 | 25 | Positive |
| 783.6 | Unit | 184.1 | Unit | 20 | 166 | 25 | Positive |
| 797.7 | Unit | 184.1 | Unit | 20 | 166 | 25 | Positive |
| 813.7 | Unit | 184.1 | Unit | 20 | 166 | 25 | Positive |
| 827.7 | Unit | 184.1 | Unit | 20 | 166 | 25 | Positive |
| 835.7 | Unit | 184.1 | Unit | 20 | 166 | 25 | Positive |
| 272.3 | Unit | 254.3 | Unit | 20 | 166 | 8 | Positive |
| 300.3 | Unit | 282.3 | Unit | 20 | 166 | 8 | Positive |
| 298.3 | Unit | 280.3 | Unit | 20 | 166 | 8 | Positive |
| 796.8 | Unit | 264.3 | Unit | 20 | 166 | 56 | Positive |
| 780.8 | Unit | 264.3 | Unit | 20 | 166 | 56 | Positive |
| 908.8 | Unit | 264.3 | Unit | 20 | 166 | 56 | Positive |
| 892.8 | Unit | 264.3 | Unit | 20 | 166 | 56 | Positive |
| 906.8 | Unit | 264.3 | Unit | 20 | 166 | 56 | Positive |
| 890.8 | Unit | 264.3 | Unit | 20 | 166 | 56 | Positive |
| 824.8 | Unit | 551.5 | Unit | 20 | 166 | 21 | Positive |
| 822.8 | Unit | 523.5 | Unit | 20 | 166 | 21 | Positive |

| | | | | | | | |
|---|---|---|---|---|---|---|---|
| 820.8 | Unit | 577.6 | Unit | 20 | 166 | 21 | Positive |
| 820.8 | Unit | 547.5 | Unit | 20 | 166 | 21 | Positive |
| 820.8 | Unit | 549.5 | Unit | 20 | 166 | 21 | Positive |
| 820.8 | Unit | 521.5 | Unit | 20 | 166 | 21 | Positive |
| 818.8 | Unit | 547.5 | Unit | 20 | 166 | 21 | Positive |
| 818.8 | Unit | 521.5 | Unit | 20 | 166 | 21 | Positive |
| 836.8 | Unit | 577.5 | Unit | 20 | 166 | 21 | Positive |
| 852.8 | Unit | 551.5 | Unit | 20 | 166 | 21 | Positive |
| 850.8 | Unit | 605.6 | Unit | 20 | 166 | 21 | Positive |
| 850.8 | Unit | 551.5 | Unit | 20 | 166 | 21 | Positive |
| 848.8 | Unit | 547.5 | Unit | 20 | 166 | 21 | Positive |
| 848.8 | Unit | 549.5 | Unit | 20 | 166 | 21 | Positive |
| 848.8 | Unit | 551.5 | Unit | 20 | 166 | 21 | Positive |
| 846.8 | Unit | 603.6 | Unit | 20 | 166 | 21 | Positive |
| 846.8 | Unit | 575.6 | Unit | 20 | 166 | 21 | Positive |
| 846.8 | Unit | 547.5 | Unit | 20 | 166 | 21 | Positive |
| 844.8 | Unit | 599.5 | Unit | 20 | 166 | 21 | Positive |
| 878.8 | Unit | 577.5 | Unit | 20 | 166 | 21 | Positive |
| 876.8 | Unit | 603.6 | Unit | 20 | 166 | 21 | Positive |
| 874.8 | Unit | 603.6 | Unit | 20 | 166 | 21 | Positive |
| 874.8 | Unit | 577.6 | Unit | 20 | 166 | 21 | Positive |
| 872.8 | Unit | 599.6 | Unit | 20 | 166 | 21 | Positive |
| 872.8 | Unit | 573.6 | Unit | 20 | 166 | 21 | Positive |
| 908.9 | Unit | 607.6 | Unit | 20 | 166 | 21 | Positive |
| 906.9 | Unit | 607.6 | Unit | 20 | 166 | 21 | Positive |
| 904.9 | Unit | 603.6 | Unit | 20 | 166 | 21 | Positive |
| 902.9 | Unit | 603.6 | Unit | 20 | 166 | 21 | Positive |
| 900.8 | Unit | 599.5 | Unit | 20 | 166 | 21 | Positive |
| 900.9 | Unit | 603.9 | Unit | 20 | 166 | 21 | Positive |
| 898.9 | Unit | 599.6 | Unit | 20 | 166 | 21 | Positive |
| 896.9 | Unit | 599.6 | Unit | 20 | 166 | 21 | Positive |
| 924.9 | Unit | 603.6 | Unit | 20 | 166 | 21 | Positive |
| 920.9 | Unit | 599.6 | Unit | 20 | 166 | 21 | Positive |
| 948.9 | Unit | 603.7 | Unit | 20 | 166 | 21 | Positive |
| 836.8 | Unit | 563.5 | Unit | 20 | 166 | 21 | Positive |
| 866.8 | Unit | 593.6 | Unit | 20 | 166 | 21 | Positive |
| 862.8 | Unit | 589.6 | Unit | 20 | 166 | 21 | Positive |

| Compour | Compour | Compour | Ion speci | CAS | z | Monoisot | ISTD | Precursor n | MS1 res | Product r | MS2 res | Dwell (m |
|---|---|---|---|---|---|---|---|---|---|---|---|---|
| PC(12:0_ | PC(12:0_18:2) | | [M+H]+ | | 1 | | FALSE | 702.5069 | Unit | 184.1 | Unit | 20 |
| PC(14:0_ | PC(14:0_14:0) | | [M+H]+ | | 1 | | FALSE | 678.5069 | Unit | 184.1 | Unit | 20 |
| PC(14:0_ | PC(14:0_18:2) | | [M+H]+ | | 1 | | FALSE | 730.5382 | Unit | 184.1 | Unit | 20 |
| PC(14:0_ | PC(14:0_18:3) | | [M+H]+ | | 1 | | FALSE | 728.5225 | Unit | 184.1 | Unit | 20 |
| PC(14:0_ | PC(14:0_18:3) | | [M+H]+ | | 1 | | FALSE | 728.5225 | Unit | 184.1 | Unit | 20 |
| PC(14:0_ | PC(14:0_20:4) | | [M+H]+ | | 1 | | FALSE | 754.5382 | Unit | 184.1 | Unit | 20 |
| PC(15:0_ | PC(15:0_16:0) | | [M+H]+ | | 1 | | FALSE | 720.5538 | Unit | 184.1 | Unit | 20 |
| PC(15:0_ | PC(15:0_16:0) | | [M+H]+ | | 1 | | FALSE | 720.5538 | Unit | 184.1 | Unit | 20 |
| PC(15:0_ | PC(15:0_16:0) | | [M+H]+ | | 1 | | FALSE | 720.5538 | Unit | 184.1 | Unit | 20 |
| PC(15:0_ | PC(15:0_16:1) | | [M+H]+ | | 1 | | FALSE | 718.5382 | Unit | 184.1 | Unit | 20 |
| PC(15:0_ | PC(15:0_18:1) | | [M+H]+ | | 1 | | FALSE | 746.5695 | Unit | 184.1 | Unit | 20 |
| PC(15:0_ | PC(15:0_18:2) | | [M+H]+ | | 1 | | FALSE | 744.5538 | Unit | 184.1 | Unit | 20 |
| PC(15:0_ | PC(15:0_18:2) | | [M+H]+ | | 1 | | FALSE | 744.5538 | Unit | 184.1 | Unit | 20 |
| PC(15:0_ | PC(15:0_18:3) | | [M+H]+ | | 1 | | FALSE | 742.5382 | Unit | 184.1 | Unit | 20 |
| PC(15:0_ | PC(15:0_20:4) | | [M+H]+ | | 1 | | FALSE | 768.5538 | Unit | 184.1 | Unit | 20 |
| PC(15:0_ | PC(15:0_22:6) | | [M+H]+ | | 1 | | FALSE | 792.5538 | Unit | 184.1 | Unit | 20 |
| PC(16:0_ | PC(16:0_13:0) | | [M+H]+ | | 1 | | FALSE | 692.5225 | Unit | 184.1 | Unit | 20 |
| PC(16:0_ | PC(16:0_14:0) | | [M+H]+ | | 1 | | FALSE | 706.5382 | Unit | 184.1 | Unit | 20 |
| PC(16:0_ | PC(16:0_14:1) | | [M+H]+ | | 1 | | FALSE | 704.5225 | Unit | 184.1 | Unit | 20 |
| PC(16:0_ | PC(16:0_16:0) | | [M+H]+ | | 1 | | FALSE | 734.5695 | Unit | 184.1 | Unit | 20 |
| PC(16:0_ | PC(16:0_16:0) | | [M+H]+ | | 1 | | FALSE | 734.5695 | Unit | 184.1 | Unit | 20 |
| PC(16:0_ | PC(16:0_16:1) | | [M+H]+ | | 1 | | FALSE | 732.5538 | Unit | 184.1 | Unit | 20 |
| PC(16:0_ | PC(16:0_16:1) | | [M+H]+ | | 1 | | FALSE | 732.5538 | Unit | 184.1 | Unit | 20 |
| PC(16:0_ | PC(16:0_17:0) | | [M+H]+ | | 1 | | FALSE | 748.5851 | Unit | 184.1 | Unit | 20 |
| PC(16:0_ | PC(16:0_17:0) | | [M+H]+ | | 1 | | FALSE | 748.5851 | Unit | 184.1 | Unit | 20 |
| PC(16:0_ | PC(16:0_18:1) | | [M+H]+ | | 1 | | FALSE | 760.5851 | Unit | 184.1 | Unit | 20 |
| PC(16:0_ | PC(16:0_18:1) | | [M+H]+ | | 1 | | FALSE | 760.5851 | Unit | 184.1 | Unit | 20 |
| PC(16:0_ | PC(16:0_18:1) | | [M+H]+ | | 1 | | FALSE | 760.5851 | Unit | 184.1 | Unit | 20 |
| PC(16:0_ | PC(16:0_18:1) | | [M+H]+ | | 1 | | FALSE | 760.5851 | Unit | 184.1 | Unit | 20 |
| PC(16:0_ | PC(16:0_18:2) | | [M+H]+ | | 1 | | FALSE | 758.5695 | Unit | 184.1 | Unit | 20 |
| PC(16:0_ | PC(16:0_18:3) | | [M+H]+ | | 1 | | FALSE | 756.5538 | Unit | 184.1 | Unit | 20 |
| PC(16:0_ | PC(16:0_18:3) | | [M+H]+ | | 1 | | FALSE | 756.5538 | Unit | 184.1 | Unit | 20 |
| PC(16:0_ | PC(16:0_18:3) | | [M+H]+ | | 1 | | FALSE | 756.5538 | Unit | 184.1 | Unit | 20 |
| PC(16:0_ | PC(16:0_18:3) | | [M+H]+ | | 1 | | FALSE | 756.5538 | Unit | 184.1 | Unit | 20 |
| PC(16:0_ | PC(16:0_22:6) | | [M+H]+ | | 1 | | FALSE | 806.5695 | Unit | 184.1 | Unit | 20 |
| PC(16:0_ | PC(16:0_8:0) | | [M+H]+ | | 1 | | FALSE | 622.4443 | Unit | 184.1 | Unit | 20 |
| PC(16:1_ | PC(16:1_18:1) | | [M+H]+ | | 1 | | FALSE | 758.5695 | Unit | 184.1 | Unit | 20 |

| | | | | | | | | | |
|---|---|---|---|---|---|---|---|---|---|
| PC(16:1_ | PC(16:1_18:1) | [M+H]+ | 1 | FALSE | 758.5695 | Unit | 184.1 | Unit | 20 |
| PC(16:1_ | PC(16:1_20:3) | [M+H]+ | 1 | FALSE | 782.5695 | Unit | 184.1 | Unit | 20 |
| PC(16:1e | PC(16:1e_18:1) | [M+H]+ | 1 | FALSE | 744.5902 | Unit | 184.1 | Unit | 20 |
| PC(16:1e | PC(16:1e_20:3) | [M+H]+ | 1 | FALSE | 768.5902 | Unit | 184.1 | Unit | 20 |
| PC(16:1e | PC(16:1e_22:5) | [M+H]+ | 1 | FALSE | 792.5902 | Unit | 184.1 | Unit | 20 |
| PC(16:1e | PC(16:1e_22:5) | [M+H]+ | 1 | FALSE | 792.5902 | Unit | 184.1 | Unit | 20 |
| PC(17:0_ | PC(17:0_18:1) | [M+H]+ | 1 | FALSE | 774.6008 | Unit | 184.1 | Unit | 20 |
| PC(17:0_ | PC(17:0_18:1) | [M+H]+ | 1 | FALSE | 774.6008 | Unit | 184.1 | Unit | 20 |
| PC(17:0_ | PC(17:0_18:2) | [M+H]+ | 1 | FALSE | 772.5851 | Unit | 184.1 | Unit | 20 |
| PC(17:0_ | PC(17:0_18:2) | [M+H]+ | 1 | FALSE | 772.5851 | Unit | 184.1 | Unit | 20 |
| PC(17:0_ | PC(17:0_18:2) | [M+H]+ | 1 | FALSE | 772.5851 | Unit | 184.1 | Unit | 20 |
| PC(17:0_ | PC(17:0_18:2) | [M+H]+ | 1 | FALSE | 772.5851 | Unit | 184.1 | Unit | 20 |
| PC(17:0_ | PC(17:0_18:3) | [M+H]+ | 1 | FALSE | 770.5695 | Unit | 184.1 | Unit | 20 |
| PC(17:0_ | PC(17:0_20:3) | [M+H]+ | 1 | FALSE | 798.6008 | Unit | 184.1 | Unit | 20 |
| PC(17:0_ | PC(17:0_20:3) | [M+H]+ | 1 | FALSE | 798.6008 | Unit | 184.1 | Unit | 20 |
| PC(17:1_ | PC(17:1_18:2) | [M+H]+ | 1 | FALSE | 770.5695 | Unit | 184.1 | Unit | 20 |
| PC(17:1_ | PC(17:1_18:2) | [M+H]+ | 1 | FALSE | 770.5695 | Unit | 184.1 | Unit | 20 |
| PC(18:0_ | PC(18:0_16:0) | [M+H]+ | 1 | FALSE | 762.6008 | Unit | 184.1 | Unit | 20 |
| PC(18:0_ | PC(18:0_18:1) | [M+H]+ | 1 | FALSE | 788.6164 | Unit | 184.1 | Unit | 20 |
| PC(18:0_ | PC(18:0_18:1) | [M+H]+ | 1 | FALSE | 788.6164 | Unit | 184.1 | Unit | 20 |
| PC(18:0_ | PC(18:0_18:3) | [M+H]+ | 1 | FALSE | 784.5851 | Unit | 184.1 | Unit | 20 |
| PC(18:0_ | PC(18:0_18:3) | [M+H]+ | 1 | FALSE | 784.5851 | Unit | 184.1 | Unit | 20 |
| PC(18:0_ | PC(18:0_20:4) | [M+H]+ | 1 | FALSE | 810.6008 | Unit | 184.1 | Unit | 20 |
| PC(18:0_ | PC(18:0_22:5) | [M+H]+ | 1 | FALSE | 836.6164 | Unit | 184.1 | Unit | 20 |
| PC(18:0_ | PC(18:0_8:0) | [M+H]+ | 1 | FALSE | 650.4756 | Unit | 184.1 | Unit | 20 |
| PC(18:1_ | PC(18:1_18:2) | [M+H]+ | 1 | FALSE | 784.5851 | Unit | 184.1 | Unit | 20 |
| PC(18:1_ | PC(18:1_20:4) | [M+H]+ | 1 | FALSE | 808.5851 | Unit | 184.1 | Unit | 20 |
| PC(18:2_ | PC(18:2_18:2) | [M+H]+ | 1 | FALSE | 782.5695 | Unit | 184.1 | Unit | 20 |
| PC(18:2e | PC(18:2e_20:4) | [M+H]+ | 1 | FALSE | 792.5902 | Unit | 184.1 | Unit | 20 |
| PC(18:3_ | PC(18:3_18:2) | [M+H]+ | 1 | FALSE | 780.5538 | Unit | 184.1 | Unit | 20 |
| PC(19:0_ | PC(19:0_20:4) | [M+H]+ | 1 | FALSE | 824.6164 | Unit | 184.1 | Unit | 20 |
| PC(19:1_ | PC(19:1_18:1) | [M+H]+ | 1 | FALSE | 800.6164 | Unit | 184.1 | Unit | 20 |
| PC(20:0e | PC(20:0e_18:2) | [M+H]+ | 1 | FALSE | 800.6528 | Unit | 184.1 | Unit | 20 |
| PC(20:5_ | PC(20:5_18:2) | [M+H]+ | 1 | FALSE | 804.5538 | Unit | 184.1 | Unit | 20 |
| PC(22:3) | PC(22:3) | [M+H]+ | 1 | FALSE | 588.366 | Unit | 184.1 | Unit | 20 |
| PC(22:5_ | PC(22:5_14:1) | [M+H]+ | 1 | FALSE | 778.5382 | Unit | 184.1 | Unit | 20 |
| PC(22:5_ | PC(22:5_14:1) | [M+H]+ | 1 | FALSE | 778.5382 | Unit | 184.1 | Unit | 20 |
| PC(26:1) | PC(26:1) | [M+H]+ | 1 | FALSE | 648.4599 | Unit | 184.1 | Unit | 20 |

| | | | | | | | | | |
|---|---|---|---|---|---|---|---|---|---|
| PC(26:2) | PC(26:2) | [M+H]+ | 1 | FALSE | 646.4443 | Unit | 184.1 | Unit | 20 |
| PC(31:0) | PC(31:0) | [M+H]+ | 1 | FALSE | 720.5538 | Unit | 184.1 | Unit | 20 |
| PC(31:1) | PC(31:1) | [M+H]+ | 1 | FALSE | 718.5382 | Unit | 184.1 | Unit | 20 |
| PC(31:1e | PC(31:1e) | [M+H]+ | 1 | FALSE | 718.5382 | Unit | 184.1 | Unit | 20 |
| PC(32:1) | PC(32:1) | [M+H]+ | 1 | FALSE | 732.5538 | Unit | 184.1 | Unit | 20 |
| PC(32:1) | PC(32:1) | [M+H]+ | 1 | FALSE | 732.5538 | Unit | 184.1 | Unit | 20 |
| PC(32:1) | PC(32:1) | [M+H]+ | 1 | FALSE | 732.5538 | Unit | 184.1 | Unit | 20 |
| PC(33:2e | PC(33:2e) | [M+H]+ | 1 | FALSE | 744.5538 | Unit | 184.1 | Unit | 20 |
| PC(33:3) | PC(33:3) | [M+H]+ | 1 | FALSE | 742.5382 | Unit | 184.1 | Unit | 20 |
| PC(33:3e | PC(33:3e) | [M+H]+ | 1 | FALSE | 742.5382 | Unit | 184.1 | Unit | 20 |
| PC(34:0) | PC(34:0) | [M+H]+ | 1 | FALSE | 762.6008 | Unit | 184.1 | Unit | 20 |
| PC(34:0) | PC(34:0) | [M+H]+ | 1 | FALSE | 762.6008 | Unit | 184.1 | Unit | 20 |
| PC(34:1) | PC(34:1) | [M+H]+ | 1 | FALSE | 760.5851 | Unit | 184.1 | Unit | 20 |
| PC(34:2) | PC(34:2) | [M+H]+ | 1 | FALSE | 758.5695 | Unit | 184.1 | Unit | 20 |
| PC(35:2e | PC(35:2e) | [M+H]+ | 1 | FALSE | 772.5851 | Unit | 184.1 | Unit | 20 |
| PC(36:2) | PC(36:2) | [M+H]+ | 1 | FALSE | 786.6008 | Unit | 184.1 | Unit | 20 |
| PC(36:2) | PC(36:2) | [M+H]+ | 1 | FALSE | 786.6008 | Unit | 184.1 | Unit | 20 |
| PC(36:2) | PC(36:2) | [M+H]+ | 1 | FALSE | 786.6008 | Unit | 184.1 | Unit | 20 |
| PC(36:5) | PC(36:5) | [M+H]+ | 1 | FALSE | 780.5538 | Unit | 184.1 | Unit | 20 |
| PC(36:5) | PC(36:5) | [M+H]+ | 1 | FALSE | 780.5538 | Unit | 184.1 | Unit | 20 |
| PC(37:4e | PC(37:4e) | [M+H]+ | 1 | FALSE | 796.5851 | Unit | 184.1 | Unit | 20 |
| PC(37:5) | PC(37:5) | [M+H]+ | 1 | FALSE | 794.5695 | Unit | 184.1 | Unit | 20 |
| PC(37:5e | PC(37:5e) | [M+H]+ | 1 | FALSE | 794.5695 | Unit | 184.1 | Unit | 20 |
| PC(37:5e | PC(37:5e) | [M+H]+ | 1 | FALSE | 794.5695 | Unit | 184.1 | Unit | 20 |
| PC(37:5e | PC(37:5e) | [M+H]+ | 1 | FALSE | 794.5695 | Unit | 184.1 | Unit | 20 |
| PC(37:5e | PC(37:5e) | [M+H]+ | 1 | FALSE | 794.5695 | Unit | 184.1 | Unit | 20 |
| PC(38:4) | PC(38:4) | [M+H]+ | 1 | FALSE | 810.6008 | Unit | 184.1 | Unit | 20 |
| PC(38:5) | PC(38:5) | [M+H]+ | 1 | FALSE | 808.5851 | Unit | 184.1 | Unit | 20 |
| PC(38:5) | PC(38:5) | [M+H]+ | 1 | FALSE | 808.5851 | Unit | 184.1 | Unit | 20 |
| PC(38:7) | PC(38:7) | [M+H]+ | 1 | FALSE | 804.5538 | Unit | 184.1 | Unit | 20 |
| PC(38:7) | PC(38:7) | [M+H]+ | 1 | FALSE | 804.5538 | Unit | 184.1 | Unit | 20 |
| PC(40:3e | PC(40:3e) | [M+H]+ | 1 | FALSE | 840.6477 | Unit | 184.1 | Unit | 20 |
| PC(40:4) | PC(40:4) | [M+H]+ | 1 | FALSE | 838.6321 | Unit | 184.1 | Unit | 20 |
| PC(44:6e | PC(44:6e) | [M+H]+ | 1 | FALSE | 890.6634 | Unit | 184.1 | Unit | 20 |

| Fragmen | CE (V) | Polarity |
|---|---|---|
| 166 | 17 | Positive |
| 166 | 17 | Positive |
| 166 | 17 | Positive |
| 166 | 17 | Positive |
| 166 | 17 | Positive |
| 166 | 17 | Positive |
| 166 | 17 | Positive |
| 166 | 17 | Positive |
| 166 | 17 | Positive |
| 166 | 17 | Positive |
| 166 | 17 | Positive |
| 166 | 17 | Positive |
| 166 | 17 | Positive |
| 166 | 17 | Positive |
| 166 | 17 | Positive |
| 166 | 17 | Positive |
| 166 | 17 | Positive |
| 166 | 17 | Positive |
| 166 | 17 | Positive |
| 166 | 17 | Positive |
| 166 | 17 | Positive |
| 166 | 17 | Positive |
| 166 | 17 | Positive |
| 166 | 17 | Positive |
| 166 | 17 | Positive |
| 166 | 17 | Positive |
| 166 | 17 | Positive |
| 166 | 17 | Positive |
| 166 | 17 | Positive |
| 166 | 17 | Positive |
| 166 | 17 | Positive |
| 166 | 17 | Positive |
| 166 | 17 | Positive |
| 166 | 17 | Positive |
| 166 | 17 | Positive |
| 166 | 17 | Positive |
| 166 | 17 | Positive |

| | | |
|---|---|---|
| 166 | 17 | Positive |
| 166 | 17 | Positive |
| 166 | 17 | Positive |
| 166 | 17 | Positive |
| 166 | 17 | Positive |
| 166 | 17 | Positive |
| 166 | 17 | Positive |
| 166 | 17 | Positive |
| 166 | 17 | Positive |
| 166 | 17 | Positive |
| 166 | 17 | Positive |
| 166 | 17 | Positive |
| 166 | 17 | Positive |
| 166 | 17 | Positive |
| 166 | 17 | Positive |
| 166 | 17 | Positive |
| 166 | 17 | Positive |
| 166 | 17 | Positive |
| 166 | 17 | Positive |
| 166 | 17 | Positive |
| 166 | 17 | Positive |
| 166 | 17 | Positive |
| 166 | 17 | Positive |
| 166 | 17 | Positive |
| 166 | 17 | Positive |
| 166 | 17 | Positive |
| 166 | 17 | Positive |
| 166 | 17 | Positive |
| 166 | 17 | Positive |
| 166 | 17 | Positive |
| 166 | 17 | Positive |
| 166 | 17 | Positive |
| 166 | 17 | Positive |
| 166 | 17 | Positive |
| 166 | 17 | Positive |
| 166 | 17 | Positive |
| 166 | 17 | Positive |
| 166 | 17 | Positive |

| | | |
|---|---|---|
| 166 | 17 | Positive |
| 166 | 17 | Positive |
| 166 | 17 | Positive |
| 166 | 17 | Positive |
| 166 | 17 | Positive |
| 166 | 17 | Positive |
| 166 | 17 | Positive |
| 166 | 17 | Positive |
| 166 | 17 | Positive |
| 166 | 17 | Positive |
| 166 | 17 | Positive |
| 166 | 17 | Positive |
| 166 | 17 | Positive |
| 166 | 17 | Positive |
| 166 | 17 | Positive |
| 166 | 17 | Positive |
| 166 | 17 | Positive |
| 166 | 17 | Positive |
| 166 | 17 | Positive |
| 166 | 17 | Positive |
| 166 | 17 | Positive |
| 166 | 17 | Positive |
| 166 | 17 | Positive |
| 166 | 17 | Positive |
| 166 | 17 | Positive |
| 166 | 17 | Positive |
| 166 | 17 | Positive |
| 166 | 17 | Positive |
| 166 | 17 | Positive |
| 166 | 17 | Positive |
| 166 | 17 | Positive |
| 166 | 17 | Positive |
| 166 | 17 | Positive |
| 166 | 17 | Positive |

| Compound group | Compounc | Compounc | Ion species | CAS | z | Monoisoto\| | ISTD? | Precursor n | MS1 res | Product m/ |
|---|---|---|---|---|---|---|---|---|---|---|
| LPE(P-20:0) | LPE(P-20:0 | C42H82NC | [M+H]+ | | 1 | | FALSE | 494.3 | Unit | 322.4 |
| LPI(18:0) [sn1] | LPI(18:0) [s | C42H82NC | [M+NH4]+ | | 1 | | FALSE | 618.3 | Unit | 341.3 |
| LPI(18:1) [sn1] | LPI(18:1) [s | C42H82NC | [M+NH4]+ | | 1 | | FALSE | 616.3 | Unit | 339.3 |
| LPI(18:1) [sn2] | LPI(18:1) [s | C42H82NC | [M+NH4]+ | | 1 | | FALSE | 616.3 | Unit | 339.3 |
| LPI(18:2) [sn1] | LPI(18:2) [s | C42H82NC | [M+NH4]+ | | 1 | | FALSE | 614.3 | Unit | 337.3 |
| LPI(20:4) [sn1] | LPI(20:4) [s | C42H82NC | [M+NH4]+ | | 1 | | FALSE | 638.3 | Unit | 361.3 |
| PC(14:0_16:0) | PC(14:0_16:0) | | [M+H]+ | | 1 | | FALSE | 706.5 | Unit | 184.1 |
| PC(14:0_20:4) | PC(14:0_20:4) | | [M+H]+ | | 1 | | FALSE | 754.5 | Unit | 184.1 |
| PC(14:0_22:6) | PC(14:0_22:6) | | [M+H]+ | | 1 | | FALSE | 778.5 | Unit | 184.1 |
| PC(15-MHDA_18:1) | PC(15-MHDA_18:1) | | [M+H]+ | | 1 | | FALSE | 774.6 | Unit | 184.1 |
| PC(15-MHDA_22:6) | PC(15-MHDA_22:6) | | [M+H]+ | | 1 | | FALSE | 820.6 | Unit | 184.1 |
| PC(16:0/16:0) | PC(16:0/16:0) | | [M+H]+ | | 1 | | FALSE | 734.6 | Unit | 184.1 |
| PC(16:0_18:0) | PC(16:0_18:0) | | [M+H]+ | | 1 | | FALSE | 762.6 | Unit | 184.1 |
| PC(16:0_18:1) | PC(16:0_18:1) | | [M+H]+ | | 1 | | FALSE | 760.6 | Unit | 184.1 |
| PC(16:0_18:2) | PC(16:0_18:2) | | [M+H]+ | | 1 | | FALSE | 758.6 | Unit | 184.1 |
| PC(16:0_18:3) (a) | PC(16:0_18:3) (a) | | [M+H]+ | | 1 | | FALSE | 756.6 | Unit | 184.1 |
| PC(16:0_18:3) (b) | PC(16:0_18:3) (b) | | [M+H]+ | | 1 | | FALSE | 756.6 | Unit | 184.1 |
| PC(16:0_20:3) (a) | PC(16:0_20:3) (a) | | [M+H]+ | | 1 | | FALSE | 784.6 | Unit | 184.1 |
| PC(16:0_20:4) | PC(16:0_20:4) | | [M+H]+ | | 1 | | FALSE | 782.6 | Unit | 184.1 |
| PC(16:0_20:5) | PC(16:0_20:5) | | [M+H]+ | | 1 | | FALSE | 780.6 | Unit | 184.1 |
| PC(16:0_22:6) | PC(16:0_22:6) | | [M+H]+ | | 1 | | FALSE | 806.6 | Unit | 184.1 |
| PC(16:1_18:2) | PC(16:1_18:2) | | [M+H]+ | | 1 | | FALSE | 756.6 | Unit | 184.1 |
| PC(16:1_20:4) | PC(16:1_20:4) | | [M+H]+ | | 1 | | FALSE | 780.6 | Unit | 184.1 |
| PC(16:1_22:6) | PC(16:1_22:6) | | [M+H]+ | | 1 | | FALSE | 804.6 | Unit | 184.1 |
| PC(18:0_18:1) | PC(18:0_18:1) | | [M+H]+ | | 1 | | FALSE | 788.6 | Unit | 184.1 |
| PC(18:0_18:2) | PC(18:0_18:2) | | [M+H]+ | | 1 | | FALSE | 786.6 | Unit | 184.1 |
| PC(18:0_20:3) | PC(18:0_20:3) | | [M+H]+ | | 1 | | FALSE | 812.6 | Unit | 184.1 |
| PC(18:0_20:4) | PC(18:0_20:4) | | [M+H]+ | | 1 | | FALSE | 810.6 | Unit | 184.1 |
| PC(18:0_22:4) | PC(18:0_22:4) | | [M+H]+ | | 1 | | FALSE | 838.6 | Unit | 184.1 |
| PC(18:0_22:5) (n3)/PC(20:1_20:4) | PC(18:0_22:5) (n3)/PC | | [M+H]+ | | 1 | | FALSE | 836.6 | Unit | 184.1 |
| PC(18:0_22:5) (n6) | PC(18:0_22:5) (n6) | | [M+H]+ | | 1 | | FALSE | 836.6 | Unit | 184.1 |
| PC(18:0_22:6) | PC(18:0_22:6) | | [M+H]+ | | 1 | | FALSE | 834.6 | Unit | 184.1 |
| PC(18:1_18:1) | PC(18:1_18:1) | | [M+H]+ | | 1 | | FALSE | 786.6 | Unit | 184.1 |
| PC(18:1_18:2) | PC(18:1_18:2) | | [M+H]+ | | 1 | | FALSE | 784.6 | Unit | 184.1 |
| PC(18:1_20:3) | PC(18:1_20:3) | | [M+H]+ | | 1 | | FALSE | 810.6 | Unit | 184.1 |
| PC(18:1_22:6) (a) | PC(18:1_22:6) (a) | | [M+H]+ | | 1 | | FALSE | 832.6 | Unit | 184.1 |
| PC(18:1_22:6) (b) | PC(18:1_22:6) (b) | | [M+H]+ | | 1 | | FALSE | 832.6 | Unit | 184.1 |

| | | | | | | | |
|---|---|---|---|---|---|---|---|
| PC(18:2_18:2) | PC(18:2_18:2) | [M+H]+ | 1 | FALSE | 782.6 | Unit | 184.1 |
| PC(18:2_20:5) | PC(18:2_20:5) | [M+H]+ | 1 | FALSE | 804.6 | Unit | 184.1 |
| PC(20:0_20:4) | PC(20:0_20:4) | [M+H]+ | 1 | FALSE | 838.6 | Unit | 184.1 |
| PC(28:0) | PC(28:0) | [M+H]+ | 1 | FALSE | 678.5 | Unit | 184.1 |
| PC(32:1) | PC(32:1) | [M+H]+ | 1 | FALSE | 732.6 | Unit | 184.1 |
| PC(32:2) | PC(32:2) | [M+H]+ | 1 | FALSE | 730.5 | Unit | 184.1 |
| PC(33:0) (a) | PC(33:0) (a) | [M+H]+ | 1 | FALSE | 748.6 | Unit | 184.1 |
| PC(34:5) | PC(34:5) | [M+H]+ | 1 | FALSE | 752.5 | Unit | 184.1 |
| PC(36:0) | PC(36:0) | [M+H]+ | 1 | FALSE | 790.6 | Unit | 184.1 |
| PC(36:6) (a) | PC(36:6) (a) | [M+H]+ | 1 | FALSE | 778.5 | Unit | 184.1 |
| PC(38:2) | PC(38:2) | [M+H]+ | 1 | FALSE | 814.6 | Unit | 184.1 |
| PC(38:4) (b) | PC(38:4) (b) | [M+H]+ | 1 | FALSE | 810.6 | Unit | 184.1 |
| PC(38:5) (a) | PC(38:5) (a) | [M+H]+ | 1 | FALSE | 808.6 | Unit | 184.1 |
| PC(38:5) (b) | PC(38:5) (b) | [M+H]+ | 1 | FALSE | 808.6 | Unit | 184.1 |
| PC(38:6) (a) | PC(38:6) (a) | [M+H]+ | 1 | FALSE | 806.6 | Unit | 184.1 |
| PC(38:7) (c) | PC(38:7) (c) | [M+H]+ | 1 | FALSE | 804.6 | Unit | 184.1 |
| PC(40:7) (a) | PC(40:7) (a) | [M+H]+ | 1 | FALSE | 832.6 | Unit | 184.1 |
| PC(40:8) | PC(40:8) | [M+H]+ | 1 | FALSE | 830.6 | Unit | 184.1 |
| PC(O-16:0/16:0) | PC(O-16:0/16:0) | [M+H]+ | 1 | FALSE | 720.6 | Unit | 184.1 |
| PC(O-16:0/20:3) | PC(O-16:0/20:3) | [M+H]+ | 1 | FALSE | 770.6 | Unit | 184.1 |
| PC(O-16:0/20:4) | PC(O-16:0/20:4) | [M+H]+ | 1 | FALSE | 768.6 | Unit | 184.1 |
| PC(O-16:0/22:6) | PC(O-16:0/22:6) | [M+H]+ | 1 | FALSE | 792.6 | Unit | 184.1 |
| PC(O-18:0/18:1) | PC(O-18:0/18:1) | [M+H]+ | 1 | FALSE | 774.6 | Unit | 184.1 |
| PC(O-18:0/18:2) | PC(O-18:0/18:2) | [M+H]+ | 1 | FALSE | 772.6 | Unit | 184.1 |
| PC(O-18:0/20:4) | PC(O-18:0/20:4) | [M+H]+ | 1 | FALSE | 796.6 | Unit | 184.1 |
| PC(O-18:0/22:6) | PC(O-18:0/22:6) | [M+H]+ | 1 | FALSE | 820.6 | Unit | 184.1 |
| PC(O-18:1/18:1) | PC(O-18:1/18:1) | [M+H]+ | 1 | FALSE | 772.6 | Unit | 184.1 |
| PC(O-18:1/18:2) | PC(O-18:1/18:2) | [M+H]+ | 1 | FALSE | 770.6 | Unit | 184.1 |
| PC(O-32:1) | PC(O-32:1) | [M+H]+ | 1 | FALSE | 718.5 | Unit | 184.1 |
| PC(O-32:2) | PC(O-32:2) | [M+H]+ | 1 | FALSE | 716.6 | Unit | 184.1 |
| PC(O-34:1) | PC(O-34:1) | [M+H]+ | 1 | FALSE | 746.6 | Unit | 184.1 |
| PC(O-34:2) | PC(O-34:2) | [M+H]+ | 1 | FALSE | 744.6 | Unit | 184.1 |
| PC(O-34:4) | PC(O-34:4) | [M+H]+ | 1 | FALSE | 740.6 | Unit | 184.1 |
| PC(O-35:4) | PC(O-35:4) | [M+H]+ | 1 | FALSE | 754.5 | Unit | 184.1 |
| PC(O-36:0) | PC(O-36:0) | [M+H]+ | 1 | FALSE | 776.6 | Unit | 184.1 |
| PC(O-36:5) | PC(O-36:5) | [M+H]+ | 1 | FALSE | 766.5 | Unit | 184.1 |
| PC(O-38:5) | PC(O-38:5) | [M+H]+ | 1 | FALSE | 794.6 | Unit | 184.1 |
| PC(O-40:5) | PC(O-40:5) | [M+H]+ | 1 | FALSE | 822.6 | Unit | 184.1 |

| | | | | | | | |
|---|---|---|---|---|---|---|---|
| PC(O-40:7) (a) | PC(O-40:7) (a) | [M+H]+ | 1 | FALSE | 818.6 | Unit | 184.1 |
| PC(O-40:7) (b) | PC(O-40:7) (b) | [M+H]+ | 1 | FALSE | 818.6 | Unit | 184.1 |
| PC(P-16:0/14:0) | PC(P-16:0/14:0) | [M+H]+ | 1 | FALSE | 690.4 | Unit | 184.1 |
| PC(P-16:0/16:0) | PC(P-16:0/16:0) | [M+H]+ | 1 | FALSE | 718.5 | Unit | 184.1 |
| PC(P-16:0/16:1) | PC(P-16:0/16:1) | [M+H]+ | 1 | FALSE | 716.6 | Unit | 184.1 |
| PC(P-16:0/18:0) | PC(P-16:0/18:0) | [M+H]+ | 1 | FALSE | 746.6 | Unit | 184.1 |
| PC(P-16:0/18:1) | PC(P-16:0/18:1) | [M+H]+ | 1 | FALSE | 744.6 | Unit | 184.1 |
| PC(P-16:0/18:2) | PC(P-16:0/18:2) | [M+H]+ | 1 | FALSE | 742.5 | Unit | 184.1 |
| PC(P-16:0/18:3) | PC(P-16:0/18:3) | [M+H]+ | 1 | FALSE | 740.6 | Unit | 184.1 |
| PC(P-16:0/20:4) | PC(P-16:0/20:4) | [M+H]+ | 1 | FALSE | 766.5 | Unit | 184.1 |
| PC(P-16:0/20:5) | PC(P-16:0/ C42H82NC | [M+H]+ | 1 | FALSE | 764.6 | Unit | 184.1 |
| PC(P-16:0/22:6) | PC(P-16:0/ C42H82NC | [M+H]+ | 1 | FALSE | 790.6 | Unit | 184.1 |
| PC(P-18:0/18:2) | PC(P-18:0/18:2) | [M+H]+ | 1 | FALSE | 770.6 | Unit | 184.1 |
| PC(P-18:0/20:4) | PC(P-18:0/ C42H82NC | [M+H]+ | 1 | FALSE | 794.6 | Unit | 184.1 |
| PC(P-18:0/22:5) | PC(P-18:0/ C42H82NC | [M+H]+ | 1 | FALSE | 820.6 | Unit | 184.1 |
| PC(P-18:0/22:6) | PC(P-18:0/ C42H82NC | [M+H]+ | 1 | FALSE | 818.6 | Unit | 184.1 |
| PC(P-18:1/18:1) | PC(P-18:1/18:1) | [M+H]+ | 1 | FALSE | 770.6 | Unit | 184.1 |
| PC(P-18:1/22:6) | PC(P-18:1/ C42H82NC | [M+H]+ | 1 | FALSE | 816.6 | Unit | 184.1 |
| PC(P-20:0/20:4) | PC(P-20:0/ C42H82NC | [M+H]+ | 1 | FALSE | 822.6 | Unit | 184.1 |
| PC(P-36:3) | PC(P-36:3) | [M+H]+ | 1 | FALSE | 768.5 | Unit | 184.1 |
| PC(P-38:5) (a) | PC(P-38:5) C42H82NC | [M+H]+ | 1 | FALSE | 792.6 | Unit | 184.1 |
| PC(P-38:5) (b) | PC(P-38:5) C42H82NC | [M+H]+ | 1 | FALSE | 792.6 | Unit | 184.1 |
| PE(15-MHDA_18:1) | PE(15-MHD C42H82NC | [M+H]+ | 1 | FALSE | 732.6 | Unit | 591.5 |
| PE(15-MHDA_18:2) | PE(15-MHD C42H82NC | [M+H]+ | 1 | FALSE | 730.5 | Unit | 589.5 |
| PE(15-MHDA_20:4) | PE(15-MHD C42H82NC | [M+H]+ | 1 | FALSE | 754.6 | Unit | 613.5 |
| PE(15-MHDA_22:6) | PE(15-MHD C42H82NC | [M+H]+ | 1 | FALSE | 778.5 | Unit | 637.5 |

| MS2 res | Dwell (ms) | Fragmentor | CE (V) | Polarity |
|---|---|---|---|---|
| Unit | 20 | 166 | 19 | Positive |
| Unit | 20 | 166 | 17 | Positive |
| Unit | 20 | 166 | 17 | Positive |
| Unit | 20 | 166 | 17 | Positive |
| Unit | 20 | 166 | 17 | Positive |
| Unit | 20 | 166 | 17 | Positive |
| Unit | 20 | 166 | 21 | Positive |
| Unit | 20 | 166 | 21 | Positive |
| Unit | 20 | 166 | 21 | Positive |
| Unit | 20 | 166 | 21 | Positive |
| Unit | 20 | 166 | 21 | Positive |
| Unit | 20 | 166 | 21 | Positive |
| Unit | 20 | 166 | 21 | Positive |
| Unit | 20 | 166 | 21 | Positive |
| Unit | 20 | 166 | 21 | Positive |
| Unit | 20 | 166 | 21 | Positive |
| Unit | 20 | 166 | 21 | Positive |
| Unit | 20 | 166 | 21 | Positive |
| Unit | 20 | 166 | 21 | Positive |
| Unit | 20 | 166 | 21 | Positive |
| Unit | 20 | 166 | 21 | Positive |
| Unit | 20 | 166 | 21 | Positive |
| Unit | 20 | 166 | 21 | Positive |
| Unit | 20 | 166 | 21 | Positive |
| Unit | 20 | 166 | 21 | Positive |
| Unit | 20 | 166 | 21 | Positive |
| Unit | 20 | 166 | 21 | Positive |
| Unit | 20 | 166 | 21 | Positive |
| Unit | 20 | 166 | 21 | Positive |
| Unit | 20 | 166 | 21 | Positive |
| Unit | 20 | 166 | 21 | Positive |
| Unit | 20 | 166 | 21 | Positive |
| Unit | 20 | 166 | 21 | Positive |
| Unit | 20 | 166 | 21 | Positive |
| Unit | 20 | 166 | 21 | Positive |
| Unit | 20 | 166 | 21 | Positive |
| Unit | 20 | 166 | 21 | Positive |

| | | | |
|---|---|---|---|
| Unit | 20 | 166 | 21 Positive |
| Unit | 20 | 166 | 21 Positive |
| Unit | 20 | 166 | 21 Positive |
| Unit | 20 | 166 | 21 Positive |
| Unit | 20 | 166 | 21 Positive |
| Unit | 20 | 166 | 21 Positive |
| Unit | 20 | 166 | 21 Positive |
| Unit | 20 | 166 | 21 Positive |
| Unit | 20 | 166 | 21 Positive |
| Unit | 20 | 166 | 21 Positive |
| Unit | 20 | 166 | 21 Positive |
| Unit | 20 | 166 | 21 Positive |
| Unit | 20 | 166 | 21 Positive |
| Unit | 20 | 166 | 21 Positive |
| Unit | 20 | 166 | 21 Positive |
| Unit | 20 | 166 | 21 Positive |
| Unit | 20 | 166 | 21 Positive |
| Unit | 20 | 166 | 21 Positive |
| Unit | 20 | 166 | 21 Positive |
| Unit | 20 | 166 | 21 Positive |
| Unit | 20 | 166 | 21 Positive |
| Unit | 20 | 166 | 21 Positive |
| Unit | 20 | 166 | 21 Positive |
| Unit | 20 | 166 | 21 Positive |
| Unit | 20 | 166 | 21 Positive |
| Unit | 20 | 166 | 21 Positive |
| Unit | 20 | 166 | 21 Positive |
| Unit | 20 | 166 | 21 Positive |
| Unit | 20 | 166 | 21 Positive |
| Unit | 20 | 166 | 21 Positive |
| Unit | 20 | 166 | 21 Positive |
| Unit | 20 | 166 | 21 Positive |
| Unit | 20 | 166 | 21 Positive |
| Unit | 20 | 166 | 21 Positive |
| Unit | 20 | 166 | 21 Positive |
| Unit | 20 | 166 | 21 Positive |
| Unit | 20 | 166 | 21 Positive |
| Unit | 20 | 166 | 21 Positive |

| | | | | |
|---|---|---|---|---|
| Unit | 20 | 166 | 21 | Positive |
| Unit | 20 | 166 | 21 | Positive |
| Unit | 20 | 166 | 21 | Positive |
| Unit | 20 | 166 | 21 | Positive |
| Unit | 20 | 166 | 21 | Positive |
| Unit | 20 | 166 | 21 | Positive |
| Unit | 20 | 166 | 21 | Positive |
| Unit | 20 | 166 | 21 | Positive |
| Unit | 20 | 166 | 21 | Positive |
| Unit | 20 | 166 | 21 | Positive |
| Unit | 20 | 166 | 21 | Positive |
| Unit | 20 | 166 | 21 | Positive |
| Unit | 20 | 166 | 21 | Positive |
| Unit | 20 | 166 | 21 | Positive |
| Unit | 20 | 166 | 21 | Positive |
| Unit | 20 | 166 | 21 | Positive |
| Unit | 20 | 166 | 21 | Positive |
| Unit | 20 | 166 | 21 | Positive |
| Unit | 20 | 166 | 21 | Positive |
| Unit | 20 | 166 | 21 | Positive |
| Unit | 20 | 166 | 21 | Positive |
| Unit | 20 | 166 | 21 | Positive |
| Unit | 20 | 166 | 17 | Positive |
| Unit | 20 | 166 | 17 | Positive |
| Unit | 20 | 166 | 17 | Positive |
| Unit | 20 | 166 | 17 | Positive |

| Compound group | Compound formula | Ion speci | CAS | z | Monoisot | ISTD | Precurso | MS1 res | Product r |
|---|---|---|---|---|---|---|---|---|---|
| DAG 16:0/18:0 | | [M+H]+ | | 1 | | FALSE | 614.5 | Unit | 313 |
| DAG 16:0/18:1 | | [M+H]+ | | 1 | | FALSE | 612.5 | Unit | 313 |
| DAG 16:0/20:4 | | [M+H]+ | | 1 | | FALSE | 634.5 | Unit | 313 |
| DAG 18:0/18:0 | | [M+H]+ | | 1 | | FALSE | 642.5 | Unit | 341 |
| DAG 18:0/18:1 | | [M+H]+ | | 1 | | FALSE | 640.5 | Unit | 341 |
| DAG 18:0/20:4-d8 | C42H82NO10P | [M+NH4]+ | | 1 | | FALSE | 670.5 | Unit | 341 |
| DAG 18:0/22:6 | C42H82NO10P | [M+NH4]+ | | 1 | | FALSE | 686.5 | Unit | 341 |
| DAG 18:0/24:0 | | [M+H]+ | | 1 | | FALSE | 662.5 | Unit | 341 |
| DAG 18:1/18:1 | | [M+H]+ | | 1 | | FALSE | 638.5 | Unit | 339 |
| DAG 18:1/20:4 | | [M+H]+ | | 1 | | FALSE | 660.5 | Unit | 339 |
| DAG 18:1/22:6 | | [M+H]+ | | 1 | | FALSE | 684.5 | Unit | 339 |
| PIP2 26:0 | C42H82NO10P | [M+H]+ | | 1 | | FALSE | 885 | Unit | 401 |
| PIP2 28:0 | C42H82NO10P | [M+H]+ | | 1 | | FALSE | 913 | Unit | 401 |
| PIP2 28:1 | C42H82NO10P | [M+H]+ | | 1 | | FALSE | 911 | Unit | 401 |
| PIP2 30:0 | C42H82NO10P | [M+H]+ | | 1 | | FALSE | 941 | Unit | 401 |
| PIP2 30:1 | C42H82NO10P | [M+H]+ | | 1 | | FALSE | 939 | Unit | 401 |
| PIP2 32:0 | C42H82NO10P | [M+H]+ | | 1 | | FALSE | 969 | Unit | 401 |
| PIP2 32:1 | C42H82NO10P | [M+H]+ | | 1 | | FALSE | 967 | Unit | 401 |
| PIP2 32:2 | C42H82NO10P | [M+H]+ | | 1 | | FALSE | 965 | Unit | 401 |
| PIP2 34:0 | C42H82NO10P | [M+NH4]+ | | 1 | | FALSE | 997 | Unit | 401 |
| PIP2 34:1 | C42H82NO10P | [M+NH4]+ | | 1 | | FALSE | 995 | Unit | 401 |
| PIP2 34:2 | C42H82NO10P | [M+H]+ | | 1 | | FALSE | 993 | Unit | 401 |
| PIP2 36:0 | C42H82NO10P | [M+NH4]+ | | 1 | | FALSE | 1025 | Unit | 401 |
| PIP2 36:1 | C42H82NO10P | [M+NH4]+ | | 1 | | FALSE | 1023 | Unit | 401 |
| PIP2 36:2 | C42H82NO10P | [M+NH4]+ | | 1 | | FALSE | 1021 | Unit | 401 |
| PIP2 36:4 | C42H82NO10P | [M+NH4]+ | | 1 | | FALSE | 1017 | Unit | 401 |
| PIP2 38:4 | C42H82NO10P | [M+NH4]+ | | 1 | | FALSE | 1045 | Unit | 401 |
| PIP2 40:6 | C42H82NO10P | [M+NH4]+ | | 1 | | FALSE | 1069 | Unit | 401 |
| MGDG(16:0_18:1) | C42H82NO10P | [M-H]- | | 1 | | FALSE | 755.57 | Unit | 331.3 |
| MGDG(16:0_18:1) | C42H82NO10P | [M-H]- | | 1 | | FALSE | 755.57 | Unit | 357.3 |
| MGDG(16:0_18:2) | C42H82NO10P | [M-H]- | | 1 | | FALSE | 753.55 | Unit | 331.3 |
| MGDG(16:0_18:2) | C42H82NO10P | [M-H]- | | 1 | | FALSE | 753.55 | Unit | 355.3 |
| MGDG(18:1_18:1) | C42H82NO10P | [M-H]- | | 1 | | FALSE | 781.58 | Unit | 357.3 |
| MGDG(18:1_18:1) | C42H82NO10P | [M-H]- | | 1 | | FALSE | 781.58 | Unit | 357.3 |
| MGDG(18:1_18:2) | C42H82NO10P | [M-H]- | | 1 | | FALSE | 779.57 | Unit | 357.3 |
| MGDG(18:1_18:2) | C42H82NO10P | [M-H]- | | 1 | | FALSE | 779.57 | Unit | 355.3 |
| MGDG(42:4) | C42H82NO10P | [M-H]- | | 1 | | FALSE | 861.65 | Unit | 323.2 |
